# *MECP2*-mutant marmosets exhibit primate-specific phenotypes of Rett syndrome

**DOI:** 10.64898/2026.06.26.734672

**Authors:** Noriyuki Kishi, Junko Okahara, Kenya Sato, Daisuke Yoshimaru, Yoko Kurotaki, Kohei Onishi, Tsukasa Sanosaka, Rachel Henry, Taeko Ito, Misako Okuno, Edward J. Weinstein, Jill R. Crittenden, Hirotaka James Okano, Jun-ichi Hata, Jun Kohyama, Erika Sasaki, Tomomi Shimogori, Hideyuki Okano

## Abstract

We used genome editing to generate *MECP2*-knockout marmosets and establish a new primate model for Rett syndrome (RTT), a neurodevelopmental disorder. MRI analysis of this marmoset model revealed diminished cortico-cortical connections, particularly those originating in the prefrontal cortex. Detailed histological analysis revealed reduced dendritic arborization and synaptic density, particularly in the upper layers of the cerebral cortex. Cortical layer II/III excitatory neurons appeared to be immature based on single-nucleus RNA sequencing and spatial transcriptomics, while *PVALB*-positive interneurons exhibited upregulation of neuronal maturation-related transcripts. Moreover, the neurodevelopmental disease gene *GABAB receptor 2* (*GABBR2*) emerged as a conserved downstream target of MECP2, illuminating a cellular cascade that could be targeted in preclinical and therapeutic drug development. Many of the gene expression changes observed in the marmoset model are common in RTT patients, thereby highlighting molecular links from cell type-specific phenotypes in the marmoset model to the RTT clinical presentation.

## Introduction

Rett syndrome (RTT) is a severe neurodevelopmental disorder predominantly affecting females, with a prevalence rate of approximately one in 10,000–15,000 girls^1–3^. Following a period of apparently normal development for 6–18 months, girls with RTT exhibit regression of acquired skills, loss of purposeful hand movements, microcephaly, autistic behaviors, breathing abnormalities, repetitive movements, and severe cognitive and social impairments^1–3^.

Mutations in the *methyl-CpG binding protein 2* (*MECP2*) gene on the X chromosome are found in over 95% of RTT patients^4^ and typically arise de novo in the paternal germline^5^. Depending on the specific mutation and whether it is transmitted to a female or, more rarely, a male, *MECP2* dysfunction can also cause a variety of neurodevelopmental disorders, including autism spectrum disorder, childhood schizophrenia and X-linked cognitive disability^1–3,6^. Since the discovery of this gene, several lines of *Mecp2*-mutant mice have been established^7,8^, and studies of *MECP2*-mutant phenotypes have provided direct insight into the relationship between the regulation of gene expression and neurodevelopment. MECP2 was initially characterized as a transcriptional repressor that selectively binds to methyl-CpG dinucleotides in the mammalian genome, alongside co-factors such as HDAC1, SIN3A and NCoR1/2 co-repressor complex^9–11^. However, it is recognized that under certain conditions, MECP2 can also function as a transcriptional activator^9,12,13^. Recent research has also demonstrated that MECP2 is involved in chromatin structure^14,15^, miRNA processing^16^, transcriptional regulation through its interaction with methyl-CpA^17,18^ and MECP2 binding to specific gene enhancers in a DNA methylation-independent manner^19^. MECP2 regulates distinct molecular pathways in different cell types, and the variety of phenotypes reflects the involved cell type^20–23^. In the central nervous system (CNS), MECP2 expression increases with neural maturation and is particularly high in mature neurons, supporting its function in differentiated neurons^24,25^. Indeed, in response to neuronal activity, MECP2 dissociates from the promotor region of target genes, derepressing their expression^26–28^.

Understanding the function of MECP2 in the CNS and the pathological mechanisms of RTT led to clinical trials that were based on findings obtained in a mouse model. The administration of insulin-like growth factor 1 (IGF1) and related compounds significantly improved RTT-like phenotypes in the mouse model^29,30^. However, clinical trials using recombinant IGF1 as a potential treatment for RTT have failed to demonstrate therapeutic efficacy^31^. Although trofinetide (a tripeptide derived from enzymatic cleavage of IGF1) was recently approved by the FDA as the first drug for RTT^32^, its therapeutic effect in patients with Rett syndrome was limited compared to the that in the mouse model. This suggests that the mouse model does not always faithfully recapitulate the pathophysiology of RTT.

Although the availability of well-established gene manipulation techniques for mice have made mice the predominant mammalian model for human disease, their relatively underdeveloped brains are not optimal for studying human brain disorders. Notably, the non-human primate (NHP) brain, unlike the rodent brain, has a relatively large cerebral neocortex that can be used to model high-level human brain functions^33^. NHP models are thus key to the experimental study of psychiatric and neurodevelopmental brain disorders. Recently, the common marmoset (*Callithrix jacchus*) has attracted much attention as a model because of its overall small size, rapid development, sociability and genetic tractability^33,34^. Thus, we have generated a genetic model of RTT in marmosets and evaluated the significance of this model.

In the present study, to investigate whether *MECP2*-deficient marmosets are the most suitable model of RTT, we employed state-of-the-art genome editing techniques to generate the first marmoset models of RTT. Using genome editing, we disrupted the expression of two key MECP2 domains that are commonly impacted by de novo, disease-causing hypomorphic mutations in RTT^3,35^. The resulting *MECP2*-knockout marmosets were born without overt abnormalities. However, by the weaning period, RTT model marmosets displayed several behavioral and neurological phenotypes. MRI analyses of *MECP2-*deficient marmosets showed significant decreases in cortico-cortical connections, especially those stemming from the prefrontal cortex. Pyramidal neuron dendritic trees play an essential role in these connections, and morphological analyses showed that *MECP2*-deficient pyramidal neurons of the cerebral cortex had severely decreased dendritic arborization and spine density. Single-nucleus RNA sequencing (RNA-seq) results were consistent with impaired neuronal maturation in the prefrontal and occipital cortices of RTT marmosets, including diminished dendritic markers for excitatory pyramidal neurons and a potential excitation/inhibition imbalance. Notably, the identity and severity of gene dysregulation varied not only among cell types but also across cortical layers, as evidenced by spatial transcriptomics. Comparison of RNA-seq data in mouse and marmoset models of RTT indicated that the marmoset model more faithfully recapitulates the gene dysregulation found in the human syndrome. The genes commonly dysregulated in both humans and this primate model have a strong tendency to be associated with other human neurodevelopmental disorders. For example, we found that *GABBR2*, a downstream target of MECP2 that is known to be a causative gene for an RTT-like neurodevelopmental disorder^36,37^, was significantly downregulated in the prefrontal cortex of RTT model marmosets. Therefore, *MECP2*-mutant marmosets are a valid model of RTT, mimicking the molecular, cellular and behavioral indications of this disease in humans, and establish a new foundation for discovering and testing new therapies.

## Results

### *MECP2-*mutant marmosets are successfully established

To establish a nonhuman primate model of RTT, *MECP2*-mutant marmosets were generated using engineered zinc finger nucleases^38^ (ZFNs) and the CRISPR/Cas9 system^39,40^. The *MECP2* gene contains two critical domains: the methyl-CpG binding domain (MBD), which is responsible for binding to methylated CpG dinucleotides in DNA, and the transcriptional repression domain (TRD), which interacts with corepressors such as SIN3A and HDAC1. We simultaneously disrupted both the MBD and TRD by utilizing ZFNs and CRISPR/Cas9 to create frameshift mutations in exon 3, which encodes the essential region of the MBD. We first tested and validated exon 3 target sequences (Figures 1A and 1B) by using in vitro assays and experiments with fertilized marmoset eggs. ZFN mRNA or gRNA and Cas9 protein were injected into in vitro fertilized eggs, which were then transferred into the uteri of surrogate mothers to achieve either ZFN or CRISPR/Cas9 mutagenesis (Table S1).

**Figure 1.**
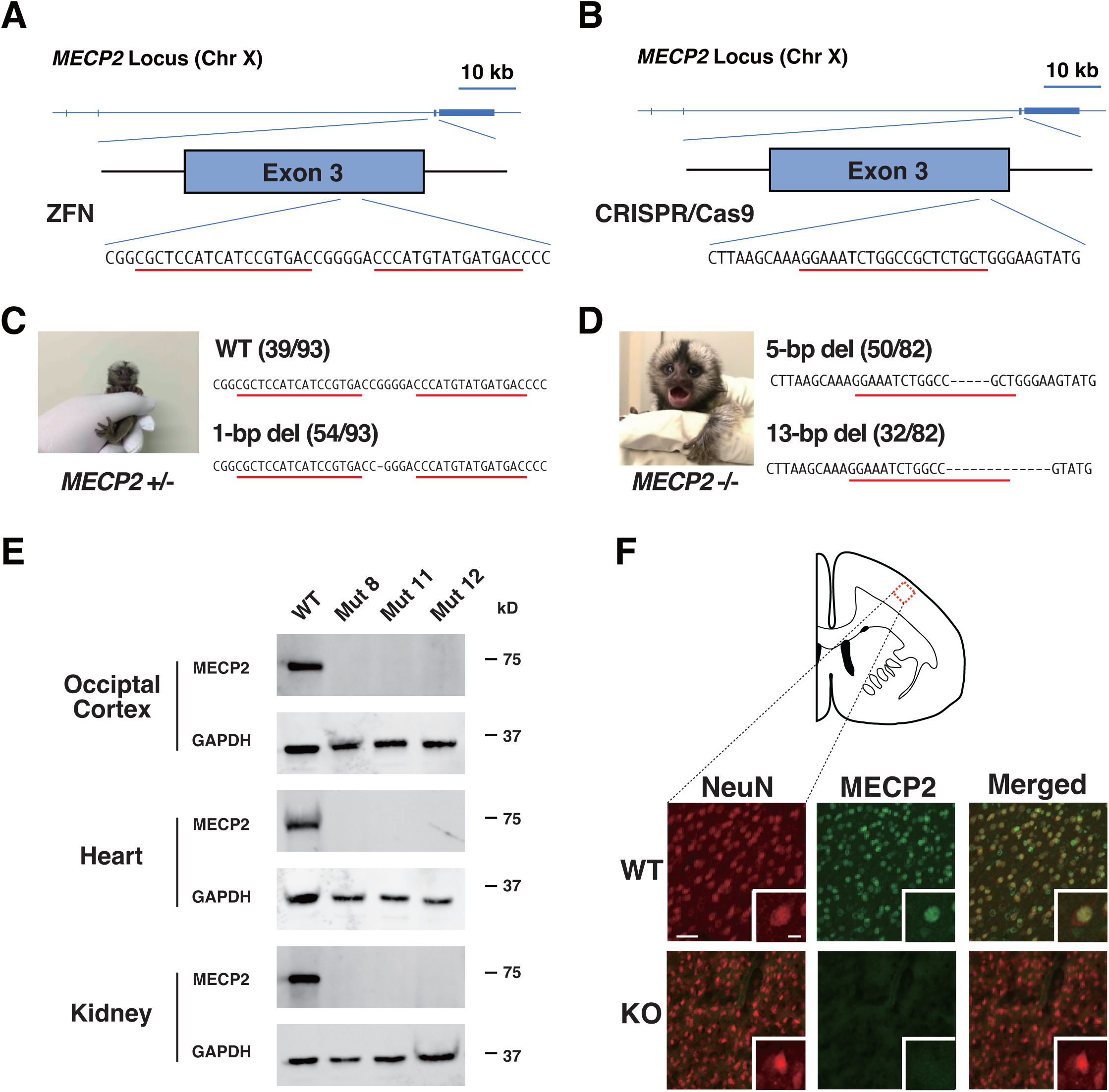
Generation of *MECP2*-mutant marmosets using genome editing technologies. (A) Schematic diagram of the pair of ZFNs targeting exon 3 of the marmoset *MECP2* locus. Red lines under the nucleotide sequence indicate the binding sequences of the ZFN pair. (B) Schematic diagram of the CRISPR/Cas9 gRNA targeting exon 3 of the marmoset *MECP2* locus. The red line under the nucleotide sequence indicates the gRNA targeting site. (C) Sequences of the modified *MECP2* locus in a *MECP2* heterozygous marmoset (Mut 1 on Postnatal Day 1 (P1)). A one-bp deletion was detected in 58 of 93 PCR products analyzed for genotyping. Red lines indicate the binding sequences of the ZFN pair. (D) Sequences of the modified *MECP2* locus in a *MECP2*-null marmoset (Mut 2 on P1). Out of 82 PCR products analyzed for genotyping, a 5-bp deletion was detected in 50 samples, and a 13-bp deletion was detected in 32 samples. (E) Western blot analysis of MECP2 in occipital cortex, heart, and kidney protein extracts from wild-type (WT) and *MECP2*-null (KO) marmosets (Mut 8, Mut 11 and Mut 12) showed that no MECP2 was detected in KO marmoset tissues. GAPDH was used as a loading control. (F) Immunostaining of the frontal cortex of WT and *MECP2*-null marmosets at 4 months of age using an anti-NeuN antibody (red) and anti-MECP2 antibody (green). Scale bar, 50 μm in the main frames and 10 μm in the insets.

Following ZFN mutagenesis, 11 marmosets were born, and one female marmoset with a heterozygous 1-bp deletion in exon 3 of *MECP2* (Mut 1, Figure 1C) survived. Mut 1 gave rise to two male *MECP2* hemizygous mutant marmosets (Mut 11 and Mut 12) and two female *MECP2* heterozygous mutant marmosets (Mut 4 and Mut 16), confirming germline transmission (Table S2). Following CRISPR/Cas9 mutagenesis, 12 of 15 marmosets carried *MECP2* mutations, demonstrating the high efficiency of this genome editing system in establishing the model. Of the 12 *MECP2* mutants established by CRISPR/Cas9, 10 had insertion or deletion mutations resulting in a frame shift and premature stop codon (Figure 1D and Table S3). Western blot and immunohistochemical analyses confirmed the absence of MECP2 protein in various tissues of these marmosets (Figures 1E and 1F). The other two mutant marmosets (Mut 5 and Mut 13) had a deletion that resulted in the expression of a MECP2 protein lacking four amino acids (Table S3); these marmosets had a lifespan approximately twice that of the other nine marmosets, indicative of a hypomorphic mutation.

To further evaluate the model, we examined whether the engineered nucleases induced off-target mutations. Based on sequence similarity to the target sequences, 4 off-target sequences for ZFN (Table S4) and 11 off-target sequences for CRISPR/Cas9 (Table S5) were identified. To examine the candidate off-targets, we performed whole-genome sequencing (WGS) analysis on the 13 founder mutant marmosets, and no mutations in these potential off-target sequences were detected in the *MECP2*-mutant marmosets (Tables S6 and S7). Next, we performed trio analysis for the four marmosets (Mut 1, Mut 2, Mut 13 and Mut 14) for which parental genomes were available (Figures S1A-D) by WGS, to distinguish de novo variants from inherited variants in those mutant marmosets. From this analysis, we identified 167, 18, 18 and 12 de novo variants in Mut1, Mut 2, Mut 13 and Mut 14, respectively (Figures S1E and F, Tables S8-11). When compared to the predicted off-target candidates with less stringent criteria (Tables S12 and S13), none of them overlapped with the identified de novo variants in the four *MECP2* mutant marmosets (Figures S1E and 1F), indicating that these de novo variants are unlikely to have been caused by ZFN or CRISPR/Cas9. Taken together, these results suggest that the engineered nucleases did not cause off-target effects on the genomes of the mutant marmosets.

A total of 17 *MECP2*-mutant marmosets have been born to date (3 heterozygotes, 2 hypomorphic mutants and 12 null mutants) (Tables 1 and S3).

### Loss of MECP2 leads to premature death and abnormal behavior related to RTT

Among the *MECP2*-mutant marmosets generated, we focused our analyses on the 12 *MECP2*-null marmosets generated by ZFNs and CRISPR/Cas9 (n = 3 females and n = 9 males) because of their pronounced phenotypes. Initially, *MECP2*-null marmosets required more frequent administration of artificial supplemental milk than wild-type marmosets; however, baby *MECP2*-null marmosets matched the weight gain of wild-type controls up to 3 months of age, except at 2 months of age (Figure 2A). To initially analyze the phenotype of the model marmosets, we examined their breathing patterns by whole-body flow plethysmograph, given that irregular breathing is a characteristic of RTT patients^1,41^; however, we did not detect any abnormal breathing, including apnea, at 3 months of age (Figure S2). These findings indicate that loss of *MECP2* does not cause gross abnormalities in initial postnatal development, which mimics the initial non-symptomatic phase of RTT.

**Figure 2.**
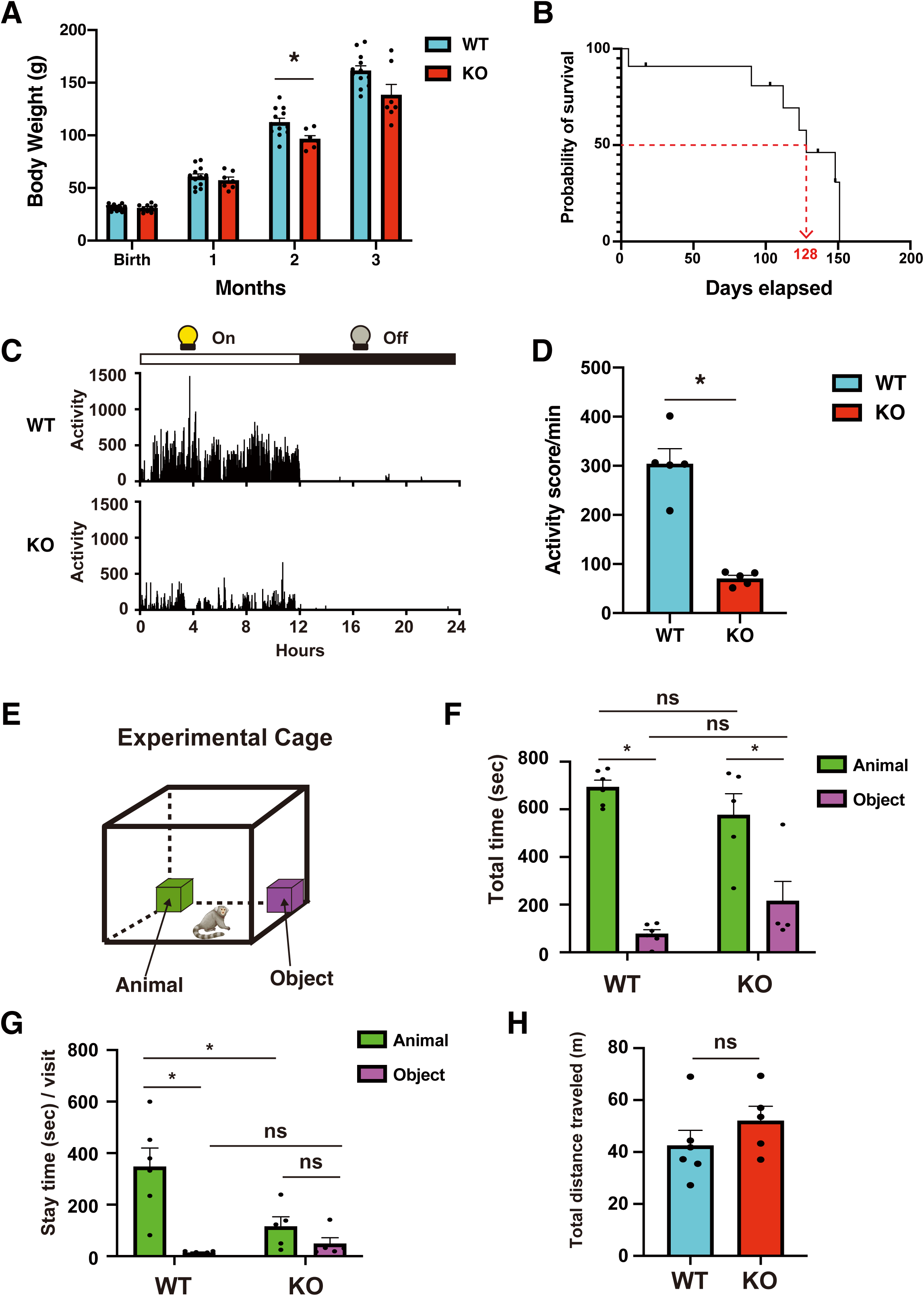
Loss of *MECP2* results in premature death and reduced voluntary movement. (A) Average body weight of wild-type (WT) and *MECP2*-null (KO) marmosets during the neonatal stages. Data are presented as the mean ± SEM. WT: n = 12, KO: n = 9, multiple t test corrected by the Holm‒Sidak method, * *p* < 0.05. (B) Survival probability curve of KO marmosets (n = 11). The median survival time was 128 days. (C) Activity scores of WT and KO marmosets at 4 months of age were monitored for 7 days while in their home cages using an Actiwatch-Mini® device. Activity was plotted for every minute of each day (light phase for 12 hours and dark phase for 12 hours). (D) KO marmosets display a robust reduction in voluntary activity in the light phase compared to that of WT marmosets. Data are presented as the mean ± SEM. Student’s t test, WT: n = 5, KO: n = 5, * *p* < 0.01. (E) Sociability test using a cage-like apparatus and a motion-sensing input device. Two transparent small boxes containing either an animal or an object were placed in the corners of the experimental cage. The test animal was placed in the apparatus and allowed to move freely for 15 min. (F) The total time spent in a compartment, i.e., near one of the small transparent boxes, during the sociability test was analyzed. (G and H) Average stay time per visit (G) and total distance traveled during a 15-min session (H) during the sociability test. Data are presented as the mean ± SEM. F and G: Two-way ANOVA, Sidak’s multiple comparisons test; H: Student’s t test, WT: n = 6, KO: n = 5, * *p* < 0.01.

However, after the 3-month weaning period, *MECP2*-null marmosets frequently died, with a median survival time of 128 days (Figure 2B). Pathologic examination identified aspiration pneumonia as the cause of death in three *MECP2*-null marmosets (Mut 3, Mut 6, and Mut 10) (Table 1). In addition, two marmosets, *MECP2*^+/-^ (Mut 4) and a MECP2 mosaic with an in-frame shift mutation (Mut 5), died from foreign body aspiration during anesthesia. We surmise that dysphagia led to fatal aspiration complications in *MECP2*-null marmosets as dysphagia commonly causes aspiration pneumonia and contributes to mortality in individuals with RTT^1,42^ and other cognitive and psychomotor disorders, such as Alzheimer’s, Parkinson’s and Huntington’s disease^43–45^.

**Table 1:** Summary of *MECP2* mutant marmosets.

| Animal ID | Gender | Genotype | Engineered Nuclease | Generation | Lifespan | Cause of death |
| --- | --- | --- | --- | --- | --- | --- |
| Mut 1 | Female | *MECP2* +/- | ZFN | F0 | alive (> 9 years) | n/a |
| Mut 2 | Female | *MECP2* -/- | CRISPR/Cas9 | F0 | 151 days | Unknown |
| Mut 3 | Male | *MECP2* -/Y | CRISPR/Cas9 | F0 | 112 days | Aspiration pneumonia |
| Mut 4 | Female | *MECP2* +/- | ZFN | F1 | 18 months | Accidental death from anesthesia |
| Mut 5 | Female | mosaic | CRISPR/Cas9 | F0 | 248 days | Accidental death from anesthesia |
| Mut 6 | Male | *MECP2* -/Y | CRISPR/Cas9 | F0 | 5 days | Aspiration pneumonia |
| Mut 7 | Female | *MECP2* -/- | CRISPR/Cas9 | F0 | 148 days | Pulmonary hypertension |
| Mut 8 | Male | *MECP2* -/Y | CRISPR/Cas9 | F0 | 148 days | Sacrificed for experiments |
| Mut 9 | Female | *MECP2* -/- | CRISPR/Cas9 | F0 | 90 days | Bacterial pneumonia |
| Mut 10 | Male | *MECP2* -/Y | CRISPR/Cas9 | F0 | 128 days | Aspiration pneumonia |
| Mut 11 | Male | *MECP2* -/Y | ZFN | F1 | 136 days | Sacrificed for experiments |
| Mut 12 | Male | *MECP2* -/Y | ZFN | F1 | 123 days | Interstitial pneumonia |
| Mut 13 | Female | *MECP2* -/12-bp del | CRISPR/Cas9 | F0 | 281 days | Unknown |
| Mut 14 | Male | *MECP2* -/Y | CRISPR/Cas9 | F0 | 17 days | Sacrificed for experiments |
| Mut 15 | Male | *MECP2* -/Y | CRISPR/Cas9 | F0 | 103 days | Sacrificed for experiments |
| Mut 16 | Female | *MECP2* +/- | ZFN | F1 | alive (> 1 year) | n/a |
| Mut 17 | Male | *MECP2* -/Y | CRISPR/Cas9 | F0 | 123 days | Sacrificed for experiments |

To determine whether *MECP2*-null marmosets mimic the behavioral deficits observed in humans with RTT, we evaluated their voluntary activity, sleep and sociability. Considering the short lifespans of the *MECP2*-null marmosets, our subsequent analyses focused primarily on the period prior to 4 months of age. First, we examined the voluntary activity of *MECP2*-null marmosets using an Actiwatch-Mini®, an activity monitor based on an accelerometer. The Actiwatch-Mini® was placed in a pocket attached to a jacket worn by the marmosets to record their activity during free movement both day and night for seven days. Both wild-type and *MECP2*-null marmosets exhibited high levels of activity during the day, while their activity decreased significantly during the night (Figure 2C). A comparison of daytime activity between wild-type and *MECP2*-null marmosets revealed a significant reduction in voluntary activity among *MECP2*-null marmosets compared to their wild-type controls (Figure 2D).

Next, as sleep disturbances are known to occur in RTT patients, we analyzed the activity patterns of *MECP2*-null marmosets during the night to identify potential sleep disorders. Several sleep parameters were examined, such as actual sleep time, movement time, wake bouts, and fragmentation index. However, we did not observe any sleep disturbances in the *MECP2*-null marmosets, at least at 4 months of age (Figure S3).

Then, since impaired sociability is a prominent feature of RTT^1^, we developed an apparatus to study sociability in marmosets (Figure 2E) consisting of an experimental enclosure equipped with a motion-sensing video camera and two small transparent boxes placed in adjacent corners. One of the boxes contained an object, and the other contained a wild-type cage-mate marmoset. After an acclimation period, the subject marmoset was placed in the center of the enclosure and allowed to freely explore for 15 min. Video images were analyzed to determine the amount of time spent in the compartment with either the small box containing a social stimulus or the box containing an object. Analysis of sociability revealed that wild-type marmosets spent more total time, as well as more time per visit, in the compartment with the social stimulus than in the one with the object (Figures 2F and 2G). Although the total time spent with each of the two stimuli did not differ between genotypes (Figure 2F), the length of the social visits varied. Whereas wild-type marmosets stayed in the compartment containing the social stimulus for an average of 347 seconds per visit, *MECP2*-null marmosets stayed for an average of only 115 seconds (Figure 2G). Although *MECP2-*null marmosets showed significantly less voluntary activity than wild-type animals, they traveled as much as wild-type marmosets during the sociability assay (Figure 2H), suggesting that their movement did not affect those assays. Together, these results indicate that *MECP2*-null marmosets spent more time moving in and out of the compartment containing the social stimulus than wild-type marmosets. In summary, while *MECP2*-null marmosets exhibit an overall preference for social stimuli over objects, their social interactions are more fragmented than those of their wild-type counterparts.

### *MECP2*-null marmosets exhibit stalled brain growth and reduced neuronal connectivity

Microcephaly is a major hallmark of RTT^1^; thus, we next examined postnatal brain growth in *MECP2*-null marmosets. First, to assess structural differences between wild-type and *MECP2*-mutant marmosets, we evaluated marmoset brains by MRI, obtained T2-weighted images, and analyzed changes in brain volume from 1 to 3 months of age (Figure 3A). At 1 month of age, there was no difference between groups in total brain volume. However, as the marmosets matured, brain growth stagnated in the *MECP2*-null group, resulting in 15% smaller total brain volume than the wild-type group by 3 months of age. Moreover, gray matter showed earlier vulnerability to a reduction in brain volume than white matter in *MECP2*-null marmosets, as previously reported in MRI studies of RTT patients^46^. Further volumetric comparisons focused on seven individual brain regions that we chose for their involvement in cognition and for subsequent histological and transcriptomic comparisons. Volume reductions were evident in all seven of these regions in *MECP2*-null marmosets relative to wild-type controls at 1 month of age (Figures 3B, S4-6 and Table S14). These volume reductions were not significant at 2 months of age (Figures 3B and S4) but reappeared at 3 months of age (Figures 3B and S6), when brain growth stagnation became more pronounced (Figure 3A) compared to that in wild-type marmosets.

**Figure 3.**
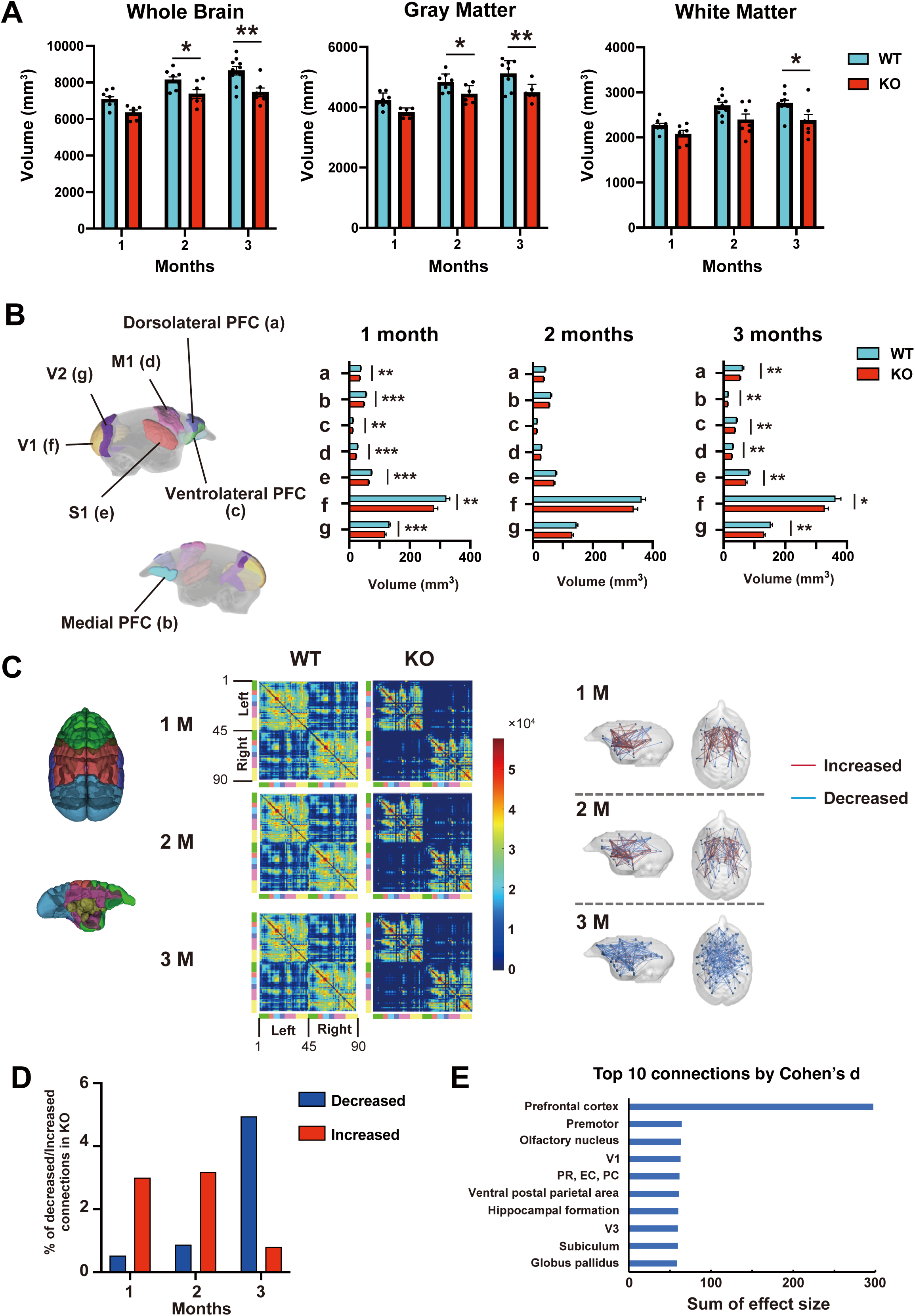
*MECP2*-null marmosets recapitulate microcephaly and show reduced connectivity between brain regions. (A) Comparison of whole brain, gray matter, and white matter volumes between wild-type (WT) and *MECP2*-null (KO) marmosets at neonatal stages. Data are presented as the mean ± SEM. Two-way ANOVA, Sidak’s multiple comparisons tests, * *p* < 0.05, ** *p* < 0.01. (B) Comparison of the volume of selected brain regions (a: dorsolateral prefrontal cortex (PFC), b: medial PFC, c: ventrolateral PFC, d: M1, e: S1, f: V1, g: V2) between WT and KO marmosets at neonatal stages. Data are presented as the mean ± SEM. Multiple t tests, FDR < 0.1. * *q* < 0.05, ** *q* < 0.01. (C) Changes over time in the structural neural connection matrix in WT and KO marmoset brains. The matrix is arranged as follows: top panel: WT (left) and KO brain (right) at 1 month of age; middle panel: WT (left) and KO brain (right) at 2 months of age; and bottom panel: WT (left) and KO brain (right) at 3 months of age. The matrix is divided into a total of 90 regions. For ease of visualization, each arrangement is shown with brain structures divided into six modules (frontal, parietal, temporal, occipital, cingulate gyrus, and subcortical). The colors of the brain diagrams correspond to the colors along the horizontal and vertical edges of the matrices. Connectivity maps showed that increases and decreases in neural connectivity between brain regions that showed significant changes and sufficient effect sizes (>0.8). This figure shows changes in KO brains compared to WT brains. Red indicates increased neural connectivity, and blue indicates decreased neural connectivity. (D) Change over time in the number of connections showing a statistically significant difference and sufficient effect size (> 0.8) between KO regions and WT regions, with increased connections and decreased connections shown separately. (E) Top 10 regions (nodes) used for neural connectivity that are significantly different in KO brains compared to WT brains at 3 months of age and have a sufficient difference in effect size (> 0.8), weighted by the sum of their respective effect sizes. PR, perirhinal cortex; EC, entorhinal cortex; and PC, piriform cortex.

To investigate the impact of reduced brain volume on neural network development in *MECP2*-null brains, we performed diffusion tensor imaging (DTI) to analyze connectivity between brain regions. DTI is a noninvasive imaging technique that can be used to visualize neuronal fiber tracts by detecting water diffusion^47^. The brain was divided into 90 regions (Figure 3C, left panel), and the connectivity between these brain regions was assessed (Figure 3C, middle panel). Increased connections were more prevalent in *MECP2*-null brains up to 2 months of age, but decreased connections dominated at 3 months of age (Figure 3C, right panel, and 3D), indicating that the pronounced stagnation of brain growth in *MECP2*-null marmosets at 3 months of age also leads to reduced neuronal connections.

Next, to assess the effect size of altered connectivity in *MECP2*-null brains at 3 months of age, we calculated Cohen’s d for each brain region. The standardized effect size was strikingly higher in the prefrontal cortex (PFC) than in other regions (Figure 3E), suggesting that reduced connectivity in the PFC due to *MECP2* loss may affect phenotypes associated with higher brain functions.

### Single-nucleus RNA-seq reveals that loss of *MECP2* dysregulates neuronal maturation

MECP2 deficiency is thought to result in various phenotypes through its dual role as a transcriptional repressor and activator^9,12,13^, providing an opportunity to directly investigate how gene expression changes drive phenotypes in *MECP2*-null marmosets. To this end, we chose to perform single-nucleus RNA-seq of the PFC, for which reduced size is known to correlate with RTT symptoms^46^, and of the occipital cortex (OC), a brain region in which there are comparative transcriptome data from RTT patients^48^. We collected tissue samples from both wild-type and *MECP2*-null marmosets at 4 months of age. Nuclei were isolated from frozen tissue samples, resulting in the successful sequencing of 14,965 nuclei from wild-type PFC samples, 12,632 nuclei from *MECP2*-null PFC samples, 10,855 nuclei from wild-type OC samples, and 14,968 nuclei from *MECP2*-null OC samples. The minimum requirement for inclusion was 500 uniquely expressed genes (Figures S7A and S8A). On average, the analyzed nuclei from the wild-type and *MECP2*-null PFC samples contained 11,919 and 11,823 transcripts per nucleus from 3,755 and 3,880 unique genes, respectively (Figures S7B and S7C), and the nuclei from the wild-type and *MECP2*-null OC samples contained 10,904 and 9,868 transcripts per nucleus from 3,741 and 3,686 unique genes, respectively (Figures S8B, and S8C).

UMAP plot analysis was performed using the R package Seurat^49^ to arrange the nuclei into 28 clusters for the PFC and the OC (Figures S7D and S8D). Each cluster was then assigned a specific cell type using known cell-type markers and additional single-nucleus RNA-seq data^50^ (Figures 4A, 4B, S7E, S7F, S8E and S8F). Within these cell-type clusters, our subsequent analyses focused on excitatory and inhibitory neuron clusters. Excitatory neurons were further categorized into upper-layer (UN: Ex_1 for the PFC in Figure 4A and Ex_1 for the OC in Figure 4B) and deep-layer (DN: Ex_3–8 for the PFC in Figure 4A and Ex_3–8 for the OC in Figure 4B) neurons, and differential gene expression analysis was performed to compare wild-type and *MECP2*-null neurons (Tables S15-S27).

**Figure 4.**
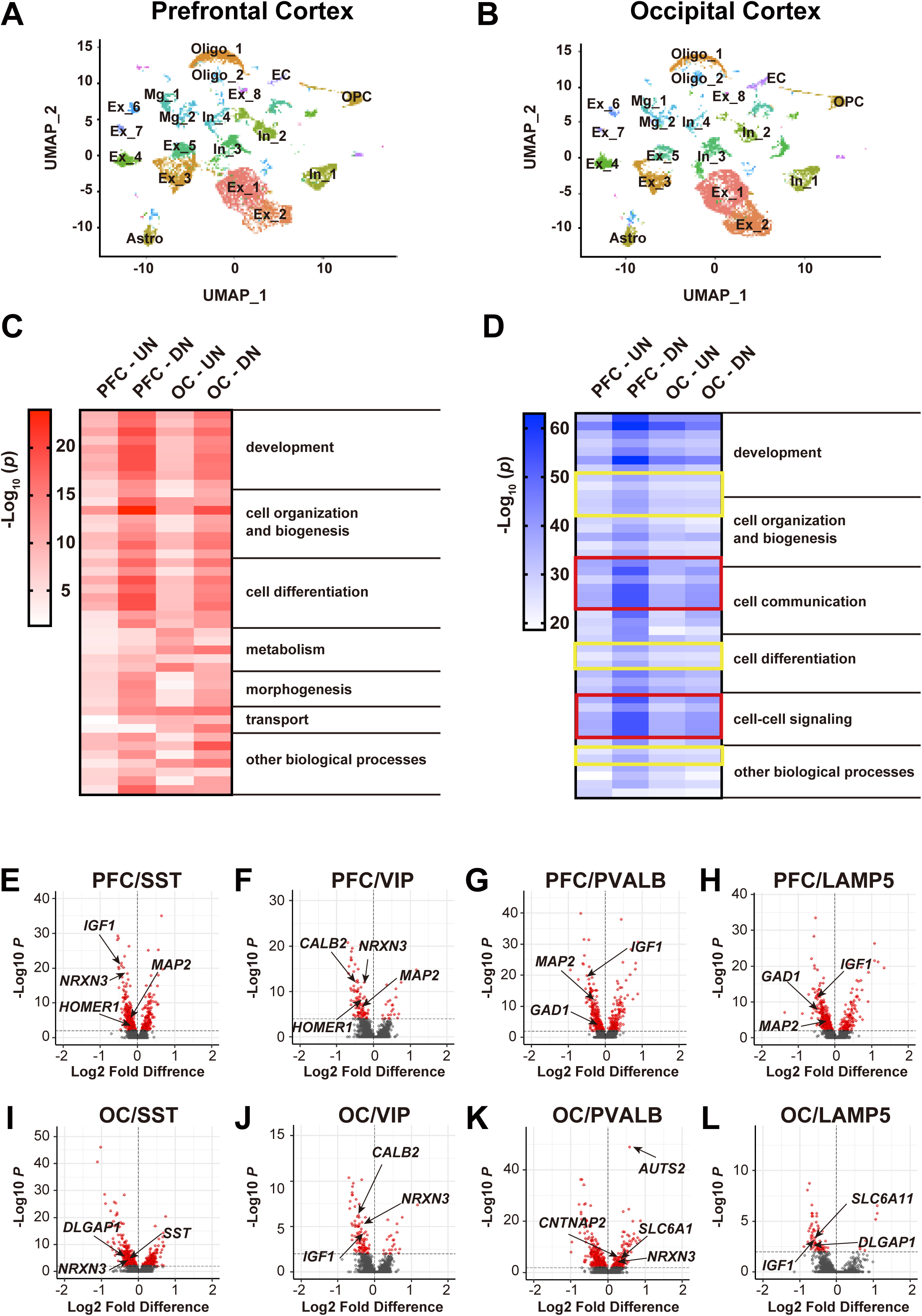
Single-nucleus RNA sequencing reveals that loss of *MECP2* affects neuronal maturation. (A and B) UMAP dimensional reduction of single-nucleus RNA sequencing of the PFC (A) and OC (B) from 3 wild-type and 3 *MECP2*-null brains. Cell types were identified by cell type-specific gene expression. Ex_1-8: Excitatory neurons, In_1: *SST*-positive inhibitory neurons, In_2: *VIP*-positive inhibitory neurons, In_3: *PVALB*-positive inhibitory neurons, In_4: *LAMP5*-positive inhibitory neurons, Oligo_1–2: oligodendrocyte, OPC: oligodendrocyte precursor cells, Astro: astrocyte, Mg_1-2: microglia, EC: endothelial cells. (C and D) Heatmap of GO enrichment analysis of upregulated (C) and downregulated (D) genes in *MECP2*-null upper layer excitatory neurons (UN) and deep layer excitatory neurons (DN) of the PFC and OC. The 30 most highly significant GO terms in differentially expressed genes in each excitatory neuron were grouped according to the GO slim list. GO terms related to dendritic morphology are highlighted in yellow boxes, and GO terms related to synapse organization are highlighted in red boxes. (E-L) Volcano plot showing the differentially expressed genes in *SST*-positive inhibitory neurons (E), *VIP*-positive inhibitory neurons (F), *PVALB*-positive inhibitory neurons (G) and *LAMP5*-positive inhibitory neurons (H) of the PFC and in *SST*-positive inhibitory neurons (I), *VIP*-positive inhibitory neurons (J), *PVALB*-positive inhibitory neurons (K) and *LAMP5*-positive inhibitory neurons (L) of the OC. Names of genes that are related to neuronal development and maturation are marked on the volcano plots.

To gain insights into the functional annotations of up- or downregulated differentially expressed genes (DEGs) in *MECP2*-null excitatory neurons, we performed gene ontology (GO) analysis. The top 30 GO categories were subclassified for each excitatory neuron category (Figures 4C and 4D, Tables S28 and S29). For upregulated genes, GO terms related to development or cell organization were significantly enriched, with differences across cortical areas and layers. In contrast, downregulated genes were significantly enriched in GO terms associated with neuronal development, cell communication and dendritic morphology and synapses (GO terms related to dendritic morphology and synapse organization are highlighted in yellow and red boxes, respectively, in Figure 4D). These results suggest that dysregulated gene expression due to the loss of *MECP2* disrupts the neuronal maturation process to varying degrees in different cell types.

Next, to investigate MECP2 function in inhibitory interneurons, we further analyzed our single-nucleus RNA-seq data. We identified several distinct interneurons and classified them into four known classes based on the expression of specific marker genes: *SST*, *PVALB*, *VIP*, and *LAMP5*. Gene expression analysis between wild-type and *MECP2*-null interneuron subtypes revealed differential effects of *MECP2* mutations on each subtype (Tables S30-37). Similar to the outcome for excitatory neurons, the loss of *MECP2* suppressed the expression of genes related to neuronal maturation in all but one interneuron subtype (Figures 4E-L). Specifically, *PVALB*-positive neurons showed an increase in neuronal maturity transcripts in the OC of *MECP2*-null marmosets relative to wild-type controls (Figure 4K).

### Altered gene expression due to loss of *MECP2* affects neuronal morphology and synaptogenesis

The decreased expression of genes associated with neuronal maturation in various neuronal subtypes of *MECP2*-null marmosets prompted us to look for morphological sequelae. We performed Golgi staining to visualize the complete structure of individual pyramidal excitatory neurons of the PFC (Figure 5A) and to examine their histological characteristics. Using a camera lucida device, we traced the morphology of pyramidal neurons in both the upper and deep cortical layers (Figure 5B) and subsequently analyzed dendritic arborization using Sholl analysis^51^ (Figures 5C-E and 5K-M).

**Figure 5.**
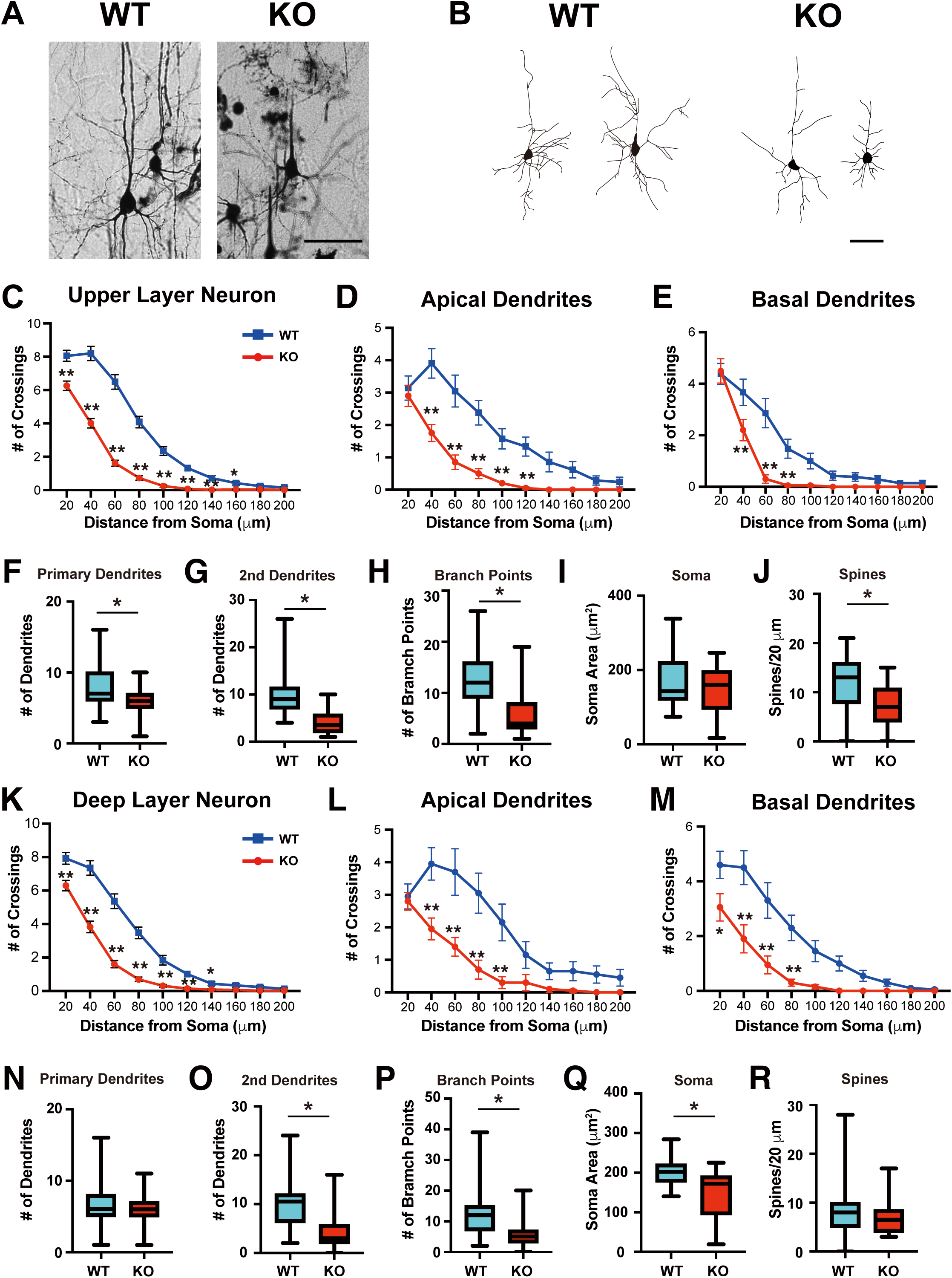
*MECP2*-null pyramidal neurons in both upper and deep cortical layers show reduced dendritic arborization. (A) Representative images of Golgi staining of pyramidal neurons in the upper cortical layer of the PFC of wild-type (WT) and *MECP2*-null marmosets at 4 months of age. Scale bar, 50 µm. (B) Representative morphological drawings of WT and *MECP2*-null upper layer pyramidal neurons in the PFC. Scale bar, 50 µm. (C-E) Arborization of total (C, WT: n = 69, *MECP2*-null: n = 81), apical (D, WT: n = 21, *MECP2-*null: n = 21), and basal (E, WT: n = 21, *MECP2*-null: n = 21) dendrites was measured by Sholl analysis of WT and *MECP2*-null upper-layer pyramidal neurons. Two-way ANOVA, Sidak’s multiple comparisons tests, * *p* < 0.05, ** *p* < 0.01. Data are presented as the mean ± SEM. (F-J) The number of primary dendrites (F, WT: n = 64, *MECP2*-null: n = 79), secondary dendrites (G, WT: n = 21, *MECP2*-null: n = 20) and branching points (H, WT: n = 64, *MECP2*-null: n = 79) as well as the soma size (I, WT: n = 25, *MECP2*-null: n = 43) and spine density (J, WT: n = 62, *MECP2*-null: n = 32) of WT and *MECP2*-null upper-layer pyramidal neurons. Student’s t test, * *p* < 0.01. Data are presented as the mean ± SEM. Box plots extend from 25th to 75th percentiles; central lines represent medial values; whiskers extend from minimum to maximum values. (K-M) Arborization of total (K, WT: n = 84, *MECP2*-null: n = 83), apical (L, WT: n = 20, *MECP2*-null: n = 20) and basal (M, WT: n = 20, *MECP2*-null: n = 20) dendrites was measured by Sholl analysis of WT and *MECP2*-null deep-layer pyramidal neurons. Two-way ANOVA, Sidak’s multiple comparisons test, * *p* < 0.05, ** *p* < 0.01. Data are presented as the mean ± SEM. (N-R) The number of primary dendrites (N, WT: n = 75, *MECP2*-null: n = 83), secondary dendrites (O, WT: n = 20, *MECP2*-null: n = 20) and branching points (P, WT: n = 75, *MECP2*-null: n = 83) as well as the soma size (Q, WT: n = 18, *MECP2*-null: n = 25) and spine density (R, WT: n = 68, *MECP2*-null: n = 18) of WT and *MECP2*-null deep-layer pyramidal neurons. Student’s t test, * *p* < 0.01. Data are presented as the mean ± SEM. Box plots extend from the 25th to the 75th percentile; central lines represent median values; whiskers extend from the minimum to the maximum value.

*MECP2*-null pyramidal neurons in both the upper and deep layers showed a significant reduction in overall dendritic arborization (Figures 5C and 5K), suggesting a decrease in inputs from other neurons. The dendritic tree of a pyramidal neuron can be divided into two distinct classes of dendrites: basal and apical dendrites^52^. Basal and proximal apical dendrites receive input from the local neuronal circuit, while the apical tufts of pyramidal neurons receive input from other cortical areas. To assess whether *MECP2* mutations affect apical or basal dendrites, we performed Sholl analysis specifically on apical and basal dendrites. Both apical and basal dendritic complexity was reduced in *MECP2*-null pyramidal neurons in both the upper and deep layers (Figures 5D, 5E, 5L and 5M), suggesting that loss of *MECP2* affects both local and distal neural connectivity.

We further characterized the morphological components of the neurons in *MECP2-*null and wild-type marmosets by measuring the number of primary (Figures 5F and 5N) and secondary dendrites (Figures 5G and 5O), branch points (Figures 5H and 5P), soma size (Figures 5I and 5Q), and spine density (Figures 5J and 5R). Consistent with the reduced dendritic arborization of *MECP2*-null pyramidal neurons, many of these morphological features were diminished by the loss of MECP2. Although both upper and deep layer neurons exhibited similar trends, primary dendrite and spine numbers were significantly reduced in upper layer neurons but not deep layer neurons. However, even though the synaptic density of neurons in deep layers of *MECP2*-null animals was not significantly different from that of wild-type neurons, the total number of synapses per neuron was reduced due to the significant decrease in dendritic number. Therefore, both upper and deep layer neurons in *MECP2*-null animals are likely to receive reduced input from other neurons, and these changes may underlie the reduced connectivity between brain regions at the whole-brain level captured by MRI.

### Spatial transcriptome analysis reveals layer-specific gene expression changes in RTT model marmosets

Our single-nucleus RNA-seq analysis revealed that the loss of *MECP2* leads to abnormal neuronal development by altering the expression of genes involved in neuronal maturation at the single-cell level. Although this analysis can classify cell types based on gene expression and analyze nuclear gene expression changes accordingly, we sought to directly measure where transcripts are localized in intact tissue to analyze changes in the spatial transcriptome caused by loss of MECP2. For this, we used 10x Visium spatial transcriptomics technology to analyze gene expression within tissue slices in which the spatial context of the cells is preserved. Frozen tissue sections from the PFC and OC of one wild-type marmoset and one *MECP2*-null marmoset at 3 months of age were evaluated (Figures 6A, S9 and S10). Sections were placed individually on a library preparation slide that holds 5000 spots with barcoded capture probes, and cDNA was subsequently synthesized from the captured mRNA, allowing for spatial mapping of the entire transcriptome. In the first round of analysis, the PFC was divided into three regions: dorsolateral PFC (dPFC), medial PFC (mPFC), and ventrolateral PFC (vPFC). Among these regions, the greatest number of DEGs between wild-type and *MECP2*-null tissue spots were in the dPFC. We then analyzed the dPFC region in finer detail. Tissue spots were grouped into seven clusters in the dPFC, and each cluster was assigned to cortical layers based on known layer-specific markers and histological locations (Figures 6B and 6C).

**Figure 6.**
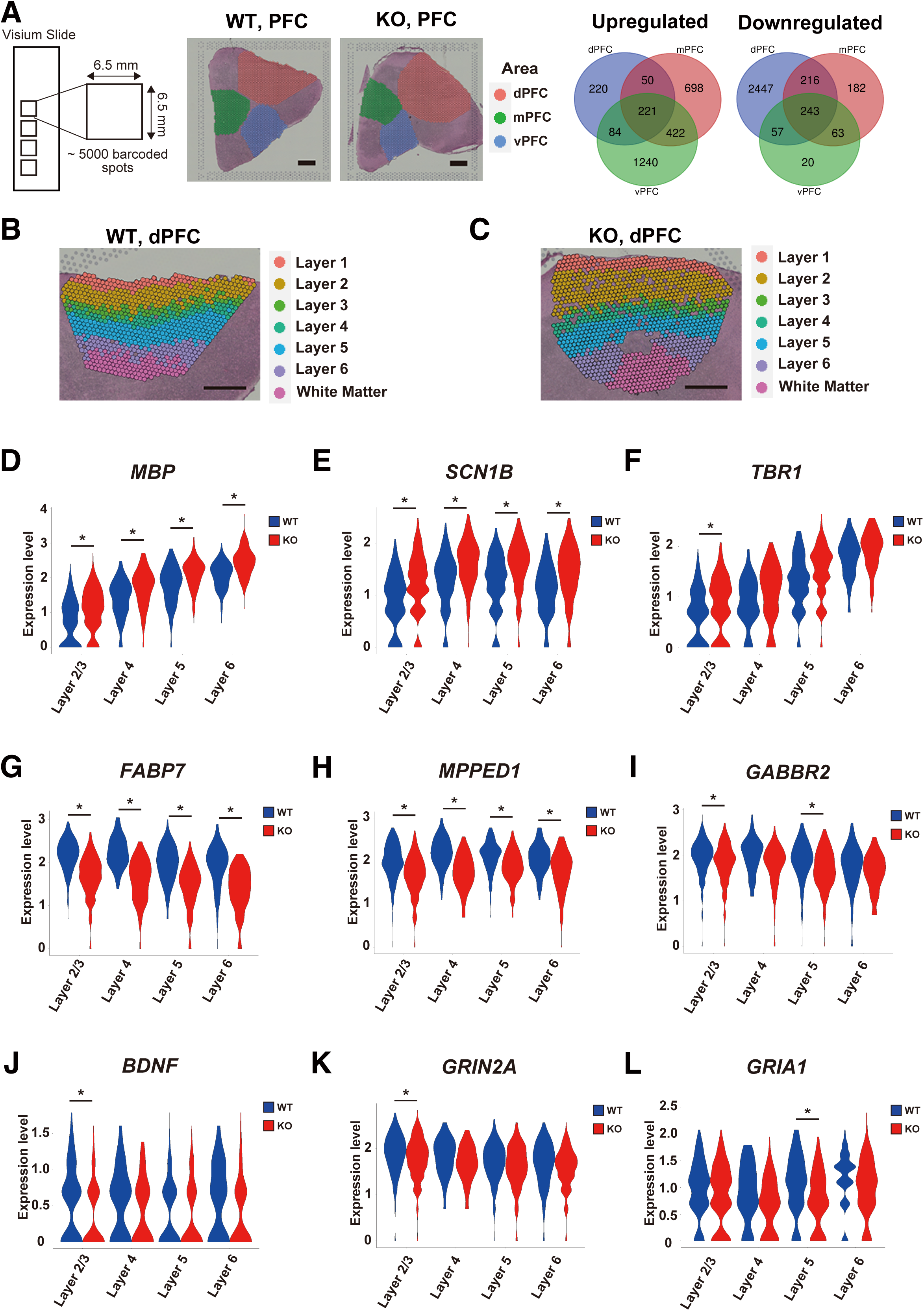
Spatial transcriptome analysis using Visium reveals that loss of *MECP2* causes distinct gene expression changes in different cortical layers. (A) PFC tissue slices from a male wild-type (WT) and a male *MECP2*-null marmoset at 3 months of age evaluated on a Visium platform; the spatial transcriptome was analyzed in three brain regions, the dorsolateral PFC (dPFC), medial PFC (mPFC) and ventrolateral PFC (vPFC). Comparisons of DEGs among the three brain regions. Scale bar, 1 mm. (B and C) Supervised annotation of dPFC layers of WT (B) and *MECP2*-null marmosets (C) based on cytoarchitecture and known gene markers. Scale bars, 1 mm. (D-L) Violin plots showing DEGs in the dPFC between WT and *MECP2*-null marmosets: *MBP* (D), *SCN1B* (E), *TBR1* (F), *FABP7* (G), *MPPED1* (H), *GABBR2* (I), *BDNF* (J), *GRIN2A* (K) and *GRIA1* (L). Wilcoxon rank sum test with the Benjamini‒Hochberg method, \**q* < 0.01.

We next compared gene expression in each layer as defined by the identities of transcripts detected in each spot between genotypes. We found distinct layer-specific patterns of gene dysregulation in RTT model marmosets (Figures 6D–6L and Tables S38-S41). In spots from *MECP2*-null marmosets, *MBP* and *SCN1B* were upregulated in both the upper and deep layers. *MBP* is predominantly expressed in oligodendrocytes and is a major component of the myelin sheath. Golli MBP, a product of the *MBP* gene, is expressed in both oligodendrocytes and neurons^53^. Interestingly, the promoter region of *MBP* that produces Golli MBP contains CpG islands^54^, suggesting the possibility that it is repressed by MECP2. Conversely, *TBR1* was upregulated in layer 2/3 of *MECP2*-null marmosets. *TBR1* encodes a transcription factor that normally defines the identity of layer 6 neurons, in part through the autism-linked gene *Auts2*^55^. The abnormal upregulation of *TBR1* in layer 2/3 could be related to a failure in migration of layer 6 neurons, a failure in cell identity specification, or mislocalization of the neuronal processes that harbor the transcript.

Among the downregulated genes in *MECP2*-null marmosets, *FABP7*, *MPPED1*, and *GABBR2* were downregulated in both the upper and deep layers, whereas others, such as *BDNF*, *GRIN2A* and *GRIA1,* showed layer-specific downregulation. *GABBR2* encodes a subunit of the GABA-B receptor, which is a G-protein coupled receptor involved in the regulation of inhibitory neurotransmission in the CNS. GABA signaling has been implicated in the pathogenesis of RTT through studies in *Mecp2*-mutant mice^20^. In particular, recent studies have shown that patients with *GABBR2* mutations exhibit Rett-like phenotypes^36,37^. *BDNF*, a gene encoding a neurotrophin critical for neuronal growth, differentiation, and survival, was specifically downregulated in only layer 2/3 of the *MECP2*-null marmoset brain. *BDNF* was the first MECP2 target gene to be identified in the CNS^26,28^, and BDNF overexpression was shown to evoke partial amelioration of the phenotypes of *Mecp2*-null mice^56^. This suggests that the downregulation of *BDNF* in layer 2/3 may affect the maturation of upper layer neurons. In summary, loss of *MECP2* alters gene expression in all layers of the PFC, with certain genes showing layer-specific dysregulation. These results identify precise regions to target for linking gene expression to *MECP2-*mutant phenotypes.

### *MECP2*-deficient marmosets share altered gene expression with patients with RTT

Our analysis showed that *MECP2*-null marmosets exhibit molecular, anatomical, and behavioral phenotypes that overlap with those of RTT patients. To further assess the relative value of our NHP model, we compared gene expression changes in excitatory neurons in the OC among *MECP2*-null marmosets, *Mecp2*-mutant mice, and RTT patients using previously analyzed single-nucleus RNA-seq data^48^ (Figures 7A, 7B, Tables S42 and S43). We first identified genes that were dysregulated in both the PFC and OC of RTT model marmosets, which yielded 1483 upregulated genes and 1604 downregulated genes. Notably, the marmoset model shared 223 upregulated genes (24%) and 355 downregulated genes (24%) with RTT patients, whereas the mouse model shared only 48 upregulated genes (5%) and 161 downregulated genes (11%) with RTT patients (Fig. 7C). This suggests that, in terms of molecular mechanisms, the marmoset model more closely resembles the pathophysiology of RTT patients than does the mouse model.

**Figure 7.**
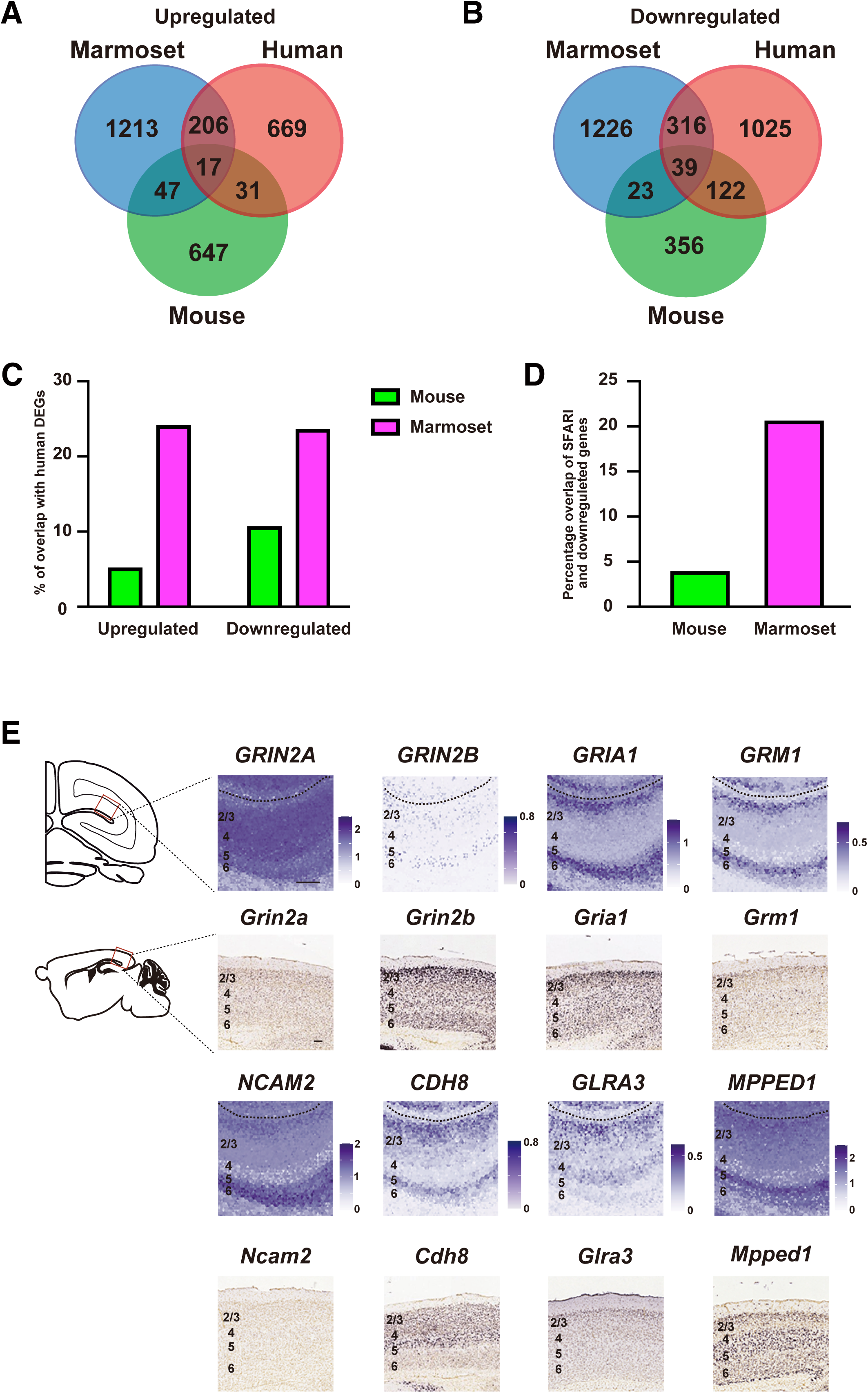
*MECP2-*mutant marmosets share genetic alterations with patients with Rett syndrome. (A and B) Venn diagram of the number of overlapping significantly upregulated (left) or downregulated (right) genes (Wilcoxon rank sum test with Benjamini-Yekutieli correction, *p* < 0.01) between wild-type (WT) and *MECP2*-null excitatory neurons in area V1 of the occipital cortex among humans, marmosets, and mice. (C) Percentage of up- or downregulated genes in mice and marmosets overlapping with human DEGs. (D) Percentage of downregulated genes overlapping with SFARI genes. (E) Expression of some genes that are downregulated in only marmosets and humans in WT OCs at 3 months of age, based on spatial expression levels from Visium analysis in the marmoset V1 area and expression of the corresponding mouse orthologs at P28 in the mouse V1 area cited from the Allen Brain Atlas^46^. The dotted line represents the calcarine sulcus. Scale bars: marmoset, 500 μm; mouse, 100 μm.

Importantly, the genes commonly downregulated in humans and marmosets included several genes associated with other neurodevelopmental disorders, such as *CUX2*, *BDNF,* and *GABBR2*. Among the 355 downregulated genes common only to humans and marmosets, 226 were SFARI genes associated with neurodevelopmental disorders^57^, whereas among the 161 downregulated genes common only to humans and mice, only 43 were SFARI genes (Figure 7D). This suggests that the *MECP2*-mutant marmoset, as a primate model of RTT, shares molecular mechanisms with other neurodevelopmental disorders.

Regarding the significant disparity in gene expression changes between rodent and primate models of RTT, we hypothesized that changes in the temporal and spatial pattern of gene expression contribute to these differences. To investigate this, we compared our spatial transcriptome data using Visium with the existing mouse gene atlas database^58^, focusing specifically on genes that were downregulated only in humans and marmosets (Figure 7E). Since 3 months is the weaning point for marmosets, we compared our spatial transcriptome data with the mouse expression data at postnatal day 28 (P28), which corresponds to the weaning time for mice. Considering that mutations in glutamate receptors are associated with a variety of neurological disorders, including neurodevelopmental disorders, we chose to evaluate this family of genes. Glutamate serves as the primary excitatory neurotransmitter in the brain and plays a critical role in various brain functions, including learning, memory, and synaptic plasticity. There are two main types of glutamate receptors: ionotropic glutamate receptors (iGluRs), such as AMPA and NMDA receptors, and metabotropic glutamate receptors (mGluRs). Importantly, recent studies suggest that NMDAR antagonists have therapeutic potential for RTT^59^. *GRIN2A*, *GRIN2B,* and *GRIA1* encode subunits of iGluRs, while *GRM1* encodes an mGluR. Mutations in these genes contribute to neurological disorders characterized by impaired intellectual development, epilepsy, and ataxia^60–62^. When we compared the expression of these genes between the marmoset OC at 3 months of age and the mouse OC at P28, distinct expression patterns emerged; for example, while *GRIN2A* was highly expressed and *GRIN2B* was weakly expressed in all layers in marmosets, *Grin2a* was weakly expressed and *Grin2b* was highly expressed in all layers in mice. We also compared the expression of *MPPED1*, *NCAM2*, *CDH8,* and *GLRA3* between mice and marmosets and observed distinct expression patterns during the weaning period in each species. *MPPED1* encodes a metallophosphodiesterase and is located in the 22q13 region that is deleted in human autosomal recessive diseases^63^. *NCAM2* and *CDH8* encode cell adhesion molecules and have been identified as autism candidate genes by genome-wide association studies^64^. *GLRA3* encodes a member of the glycine receptor superfamily, and a recent study identified GLRA3 as a novel candidate gene of neurodevelopmental disorders^65^. While *MPPED1* and *NCAM2* showed strong expression in deep layers and *CDH8* and *GLRA3* were expressed in upper layers 2/3 and deep layer 5 of marmosets, mouse homologs of these genes showed different expression patterns. These differences in the temporal and spatial patterns of downstream gene expression regulated by MECP2 suggest that *MECP2* mutations have different effects on neuronal maturation in different animal models, and these different patterns expression in different species would be expected to result in different outcomes regarding neuronal maturation. Considering that RTT is characterized by a failure of neuronal maturation and regression in behavioral development, an animal model with advanced NHP ontogeny is key for evaluating therapies for symptom reversal.

## Discussion

To better understand the link between *MECP2* loss and the molecular, cellular, and behavioral changes observed in human RTT patients, we generated and analyzed *MECP2*-mutant marmosets. We successfully created these *MECP2*-mutant marmosets through the use of genome editing technology, thereby introducing a significant nonhuman primate model for RTT research. In the predominantly female population of individuals with RTT, correlating the phenotypes with loss of *MECP2* function is challenging because of the mosaicism of X inactivation. This is not a confound in hemizygous males, but this population is very small because most mutations arise de novo in the paternal line and thus are typically passed only to daughters^5,66,67^. The rarity of inherited RTT-causing *MECP2* mutations in males, combined with the mosaicism of X inactivation in females, has also made it very difficult to assess sex differences in disease presentation. The use of CRISPR/Cas9 to directly generate homozygous female and hemizygous male marmosets in similar proportions provides an opportunity to properly evaluate genotype–phenotype correlations and the impact of sex on *MECP2* mutations.

The lethality associated with MECP2 deficiency varies among animal species. Although hemizygous male and homozygous female marmosets initially developed and grew normally until 3 months of age, the complete absence of *MECP2* led to premature death shortly after weaning. In mice, hemizygous males that lack *Mecp2* expression are born and survive to early adult stage^7,8^. In contrast, in humans, *MECP2* mutations that cause RTT in females cause severe symptoms in males, ranging from embryonic lethality to severe phenotypes and death in the first year of life^66,68^. Whereas major phenotypes in the mouse model typically manifest around sexual maturity and result in adult lethality^7,8^, the marmoset model displayed those phenotypes much earlier, around the weaning period, similar to RTT patients. Thus, compared to the mouse model, the marmoset model more closely reflects the clinical time course of RTT in humans.

Interestingly, we observed an intriguing neurodevelopmental phenomenon in *MECP2*-null marmosets during the seemingly non-symptomatic early neonatal period. At one month of age, there was no difference in total brain volume between the wild-type and *MECP2*-null marmosets, and no overt phenotypes were observed this early in development. Nevertheless, the volumes of many individual brain regions were already smaller in the model marmosets than in the controls by one month of age, suggesting that loss of MECP2 affects brain development during embryogenesis. Since MECP2 is highly expressed during postnatal neurodevelopment, its functions in the differentiated neuronal state have been extensively analyzed, revealing that MECP2 is involved in neuronal maturation and maintenance. However, MECP2 is also expressed at low levels in neural stem cells and immature neurons and plays an important role in embryonic neurogenesis. For example, a previous study revealed that MECP2 represses the transcription of long interspersed nuclear element-1 (L1) and that L1 neuronal transcription and retrotransposition are increased in *Mecp2*-null mice and neuronal progenitors derived from human iPS cells established from RTT patients^69^. The marmoset model provides an avenue to analyze the function of *MECP2* not only in postnatal primate neurodevelopment but also in the prenatal period.

To date, macaques with mutations within *MECP2* have been generated as primate models of RTT using TALEN-mediated gene targeting^70,71^, and macaques with lentiviral overexpression of *MECP2* have been developed to represent *MECP2* duplication disorder, which shares features with autism^72,73^. The TALEN-targeting technique generated 5 female mutant macaques with multiple de novo *MECP2* mutations. These monkeys exhibit several abnormalities that recapitulate some symptoms of RTT^74^. MRI analysis revealed a reduction in the volume of certain cortical and subcortical regions in the brains of the mutant macaques, and eye-tracking tests revealed social abnormalities that resemble clinical symptoms of RTT. While the marmoset model shares phenotypes with the macaque model, there are discrepancies between the two. For instance, although our MRI analyses clearly showed reductions in whole brain, gray matter, and white matter volume at three months of age and in the volume of many individual brain regions in the marmoset model, the macaque model did not show significant reductions in whole brain, gray matter, or white matter volume. In addition, while the marmoset model showed reduced voluntary activity during the daytime and no abnormalities during sleep, the macaque model showed normal daytime activity but increased wake bouts during sleep. These differences between the two primate models may be due to the degree of MECP2 dysfunction, in addition to species differences. *MECP2*-null marmosets completely lack MECP2 function, leading to robust and consistent phenotypes in mutant marmosets. In contrast, *MECP2* mutations are mosaic at the genomic level in the current macaque models^70,71,74^, resulting in variable residual MECP2 function among individuals and contributing to phenotypic variation. However, both primate models offer distinct advantages. Both *MECP2*-null and *MECP2*-heterozygous marmosets are available, so they can be used differently depending on the purpose of the research. Although *MECP2*-null marmosets display robust phenotypes, it is difficult to perform a long-term experiment due to their short lifespan. *MECP2* heterozygous marmosets display milder phenotypes than *MECP2*-null marmosets, and we could follow changes in the brain volume of *MECP2* heterozygotes over 5 years due to their longer lifespan than *MECP2*-null marmosets (Figure S11). Although statistical analysis was not possible due to the small number of individuals, the brain volume of the three female *MECP2* heterozygotes was comparable to that of wild-type marmosets up to 12 months of age but then appeared to stall, such that the brains of *MECP2* heterozygotes were smaller than those of wild-type marmosets from 18 months of age onward. In addition, like *MECP2*-null marmosets, *MECP2* heterozygotes also displayed reduced voluntary activity compared to wild-type controls at 1 year of age (Figure S12). The longer lifespan of *MECP2* heterozygous marmosets allows for long-term phenotypic analysis. Furthermore, because marmosets reach sexual maturity at 18 months, their reproductive capacity greatly facilitates the expansion of *MECP2* mutant marmosets. On the other hand, the macaque model allows for more complex analyses due to its evolutionary proximity to humans and its larger and more gyrated brains compared to marmosets. Since RTT patients show symptoms related to higher brain functions, such as the loss of purposeful hand movements, the macaque model is suitable for various complex behavioral analyses. By taking advantage of both of these two nonhuman primate models, it enables a better understanding of the pathogenesis of RTT patients.

### Translational relevance of *MECP2*-null marmosets

The development of therapies for RTT requires a bridge between preclinical findings and human clinical trials. As non-human primates, marmosets offer a unique advantage over mouse models because of their closer evolutionary relationship to humans^33^. The genetic, physiological, and behavioral similarities between marmosets and humans increase the translational relevance of findings obtained from studying RTT in marmoset models. Therefore, observations and therapeutic strategies derived from marmosets may be more readily applicable to human patients, facilitating the translation of research findings into clinical practice.

The larger size of marmosets compared to mice allows for more practical and precise pharmacological testing. This is particularly important when evaluating potential treatments for RTT. Marmosets can be used to assess the effectiveness and safety of various drug interventions, including novel compounds and therapeutic approaches, in a preclinical setting that more closely approximates human physiology than mice can offer. Such testing can provide valuable data on the potential efficacy, deliverability and side effects of candidate drugs, helping to guide clinical trials and accelerate the development of effective treatments for RTT.

To identify therapeutic targets for RTT, we investigated the downstream gene expression changes resulting from the loss of MECP2 using two techniques: single-nucleus RNA-seq and spatial transcriptome analysis of total mRNA. The combination of these two methods increases the reliability of the results and allows visualization of the spatial downregulation of genes associated with neuronal maturation in the *MECP2*-null cortex. Notably, when comparing the transcriptomes of RTT patients, mouse models, and marmoset models, we found a greater number of DEGs shared between humans and marmosets than between mice and humans. Among the DEGs commonly downregulated only in humans and marmosets, many have been implicated in other neurodevelopmental disorders, suggesting that RTT shares common pathological mechanisms with other neurodevelopmental disorders. For example, both single-nucleus RNA-seq and spatial transcriptome analysis identified *GABBR2*, *GRIN2A* and *GRIN2B* as commonly downregulated genes in humans and marmosets but not in mice. Recent studies have shown that mutations in these genes are present in *MECP2* mutation-negative patients with RTT-like phenotypes^36,37^, suggesting a direct involvement of these mutations in the pathological mechanisms of RTT. GABBR1 and GABBR2 form heterodimeric GABA receptors that inhibit neuronal activity through Gi-coupled second messenger systems^75^. GABBR2 is recognized for its capacity to maintain a delicate balance of excitatory and inhibitory neuronal signaling, exerting slow and long-lasting effects on neural development^76^. Thus, impairment of the MECP2–GABBR2 axis is likely to lead to an excitatory/inhibitory imbalance in patients with RTT and in the marmoset model, suggesting that it is a promising candidate therapeutic target for RTT patients. Moreover, previous studies have revealed that maturation of *Pvalb*-positive interneurons, which provide activity-dependent inhibitory input to pyramidal neurons to maintain excitatory/inhibitory balance^77^, is accelerated in the visual cortex of *Mecp2*-deficient mice^78,79^. *MECP2*-null marmosets also showed preservation of *PVALB*-positive interneuron identity relative to other cell types, possibly exacerbating the excitatory/inhibitory imbalance, which is recognized as the leading cellular and synaptic feature underlying various RTT neurological symptoms.

*GRIN2A* and *GRIN2B* encode subunits of the N-methyl-D-aspartate (NMDA) receptor (NMDAR), a glutamate-gated ion channel that is involved in the activity-dependent increase in synaptic transmission underlying memory and learning. Mutations in NMDARs are associated with a broad range of neurodevelopmental and psychiatric disorders^80–83^. Interestingly, the genes encoding NMDARs display very different expression patterns in the neonatal OC of marmosets than in mice, suggesting species-distinct roles in brain development. Moreover, we found *GRIN2A* and *GRIN2B* to be downregulated in the OC of the marmoset model, but they were not identified as DEGs in the comparable mouse data set. Since the relationship between MECP2 and these genes could not be detected in *Mecp2*-deficient mice, the *MECP2*-deficient marmoset we have created provides an important preclinical model for testing RTT treatments related to GABBR2, GRIN2A and GRIN2B function.

Finally, we would like to emphasize the potential clinical utility of *MECP2*-mutant marmosets. With accumulating knowledge about MECP2 function and the pathophysiology of RTT, efforts are underway to develop strategies for the treatment of this disease. Although the clinical trial of recombinant IGF1 did not show improvement in RTT patients^31^, trofinetide, a tripeptide derived from enzymatic cleavage of IGF1 that has additional anti-inflammatory properties, was recently approved by the FDA as the first drug for RTT^32^. Given that trofinetide has been recognized as an orphan drug, the imperative for continued advancement in the realm of RTT therapeutics is apparent. A phase 2 clinical trial, based in part on preclinical studies in mice^84–86^, for use of the NMDAR antagonist ketamine to treat RTT is completed, and data are under consideration for a phase 3 trial (ClinicalTrials.gov Identifier: <u>NCT03633058</u>) ^59^. Identification of the NMDAR-encoding genes *GRIN2A* and *GRIN2B* as DEGs exclusive to primates within the marmoset model provides robust support for related clinical investigations. Furthermore, our investigation has unveiled region-specific *MECP2* gene regulation in distinct layers of the brains of marmosets. This finding underscores the potential efficacy of region-specific gene targeting as an effective therapeutic approach. For instance, gene therapy employing viral vectors to deliver the *MECP2* gene may ameliorate *MECP2* deficiency in a more precisely targeted manner, contingent upon the specific behavioral deficits exhibited by individual patients. In our study, we found that marmosets, especially when considering their unique symptoms and behaviors, are a good fit for developing treatments tailored to each individual. The similarities between marmosets and humans make them a promising model for personalized therapies.

## Acknowledgments

This work was funded by the Funding Program for World-leading Innovative R&D on Science and Technology (FIRST Program), Brain Mapping by Integrated Neurotechnologies for Disease Studies (Brain/MINDS) and “Construction of System for Spread of Primate Model Animals” (to H.O. and E.S.) carried out under the Strategic Research Program for Brain Science from the Ministry of Education, Culture, Sports, Science, and Technology of Japan (MEXT) and the Japan Agency for Medical Research and Development (AMED) under grant numbers JP15dm0207001 and JP23wm0625001 (to H.O.). This work was also supported by the Rett Syndrome Support Organization and Brain/MINDS grant 24wm0625306h0001 (to N.K.). We thank Drs. Tomomi Aida, Yi Liu and Rudolf Jaenisch at MIT for their helpful comments on the manuscript and thank Ms. Yumi Ogawa, Kei Hagiya-Sakuma and Ayaka Oguchi for their excellent technical assistance.

## Author information

### Author contributions

N.K. and H.O. conceived and designed the study. J.O., K.S., Y.K., and E.S. established to the establishment of the *MECP2*-mutant marmoset lines. R.H. and E.J.W. designed and prepared the ZFN constructs. D.Y. and J.H. performed brain imaging and interpreted the data. T.I., M.O., and N.K. performed the behavioral and sleep analysis and interpreted the data. N.K. performed the pathological analysis and interpreted the data. N.K., J.O., T.S., and H.O. participated in critical discussions. N.K., K.O., T.S., J.K., and T.S. contributed to single nucleus RNA-seq and spatial transcriptome analyses. H.J.O., E.S., and H.O. contributed to the development of this project. E.S. and H.O. supervised the project and acquired financial support. N.K. and H.O. were involved in the administration of the project. The manuscript was prepared by N.K., T.S., J.C., and H.O. All authors contributed to editing the final version of the manuscript and approved submission of the manuscript.

### Data and materials availability

All single nucleus RNA-seq and Visium data reported in this study have been deposited in the NCBI Gene Expression Omnibus (GEO) under accession GSE239439. All WGS data reported in this study have been deposited in the NCBI Sequence Read Archive (SRA) under accession PRJNA1283618 (https://dataview.ncbi.nlm.nih.gov/object/PRJNA1283618?reviewer=h7614o4pgro6e2uea0i80gim6b).

## Supplemental Materials

### Materials and Methods

#### Animal Experiments

All animal experimental protocols were approved by the Animal Experimental Committee of RIKEN (approval number: W2022-2-030, W2022-2-036, W2021-2-037 and W2022-2-027) and the Institutional Animal Care and Use Committee of CIEM (approval number: 11028). Animal handling and experimental procedures were conducted in accordance with institutional guidelines for the use of laboratory animals at RIKEN and CIEM, which are consistent with the Guidelines for Proper Conduct of Animal Experiments by the Science Council of Japan (2006). Animal care was carried out in accordance with the recommendations of the Guide for the Care and Use of Laboratory Animals (National Research Council, 2011). Humane endpoints were determined based on general health conditions and behaviors (e.g., weight loss, inability to access food or water), with the approval of the Animal Experimental Committee of the RIKEN and the Institutional Animal Care and Use Committee of the CIEM. To determine the cause of death, the marmoset’s body was examined for pathology at CIEM’s Pathology Center.

#### Engineered nucleases

To generate Rett syndrome model marmosets, we designed two types of engineered nucleases, ZFN and CRISPR/Cas9, which target exon 3 of the marmoset *MECP2* gene. The ZFN constructs were obtained from Horizon Discovery (St. Louis, MO, USA). The ZFN pair binds to the target sequences shown in Figure 1A and was validated by microinjecting the mRNAs into fertilized marmoset eggs. A gRNA targeting marmoset *MECP2* was designed using the CRISPR Design website at http://crispr.mit.edu (currently not active), and the genome editing efficiency of the crRNA shown in Figure 1B was validated by comicroinjecting tracrRNA and Cas9 nuclease V3 (Integrated DNA Technologies, Japan) into fertilized marmoset eggs.

#### Off-target assay and WGS

Four potential off-target sites with homology to the ZFN binding sequences were retrieved from the whole marmoset genome database (GenBank: GCA_000004665.1) using the e-PCR tool^1^, allowing for up to 2 bp gapped alignments with up to 9 mismatches in the ZFN target sequences. Eleven potential off-target sites with homology to the 23 bp target sequence (gRNA + PAM) were retrieved using Cas-OFFinder (http://www.rgenome.net), allowing for ungapped alignments with up to three mismatches in the gRNA target sequence. WGS was performed with an Illumina NovaSeq 6000 at Rhelixa Inc, (Tokyo, Japan), using the NGS library of genomic DNA from the hair roots or umbilical cords of genetically modified marmosets. The reads were aligned to the marmoset genome reference (mCalJac4) using bwa-mem (v0.7.17-r1188, https://github.com/lh3/bwa) and duplicates were marked using Picard (v2.18.11, http://broadinstiture.github.io/picard/). After removing the duplicates, single nucleotide variants and short indels were called using freebayes (Version 1.3.4) and the raw variants were filtered with a quality filter, QUAL > 20. Trio analysis was performed for MECP2 mutant marmosets for which both parents’ genomes were available. The filtered variant files from the mutant marmosets, the mothers and the fathers were compared using ggVennDiagram (v1.2.0)^2^. To identify de novo variants, we used stringent filters of GQ > 30 and DP > 30. We also used another set of filtering to obtain the final set of de novo variants. Only variant sites in which all reads were reference or alternative alleles were judged as homozygous reference or alternative alleles, and the allele balance of the heterozygous allele had to be between 0.3 and 0.7.

#### Microinjection and embryo transfer

Unfertilized (GV) or fertilized (2PN) eggs were obtained, and embryo transfers were performed as previously described^3^. The *MECP2* eHiFi-ZFN expression plasmids were linearized using *Xba*I (Takara Bio Inc., Japan) and purified using a QIAquick PCR Purification kit (QIAGEN, Germany). The mRNAs were synthesized from the linearized template DNA using the mMESSAGE mMACHINE T7 transcription kit and Poly(A) tailing kit and purified using a MEGAclear transcription clean-up kit (Thermo Fisher Scientific, USA). Approximately 5–10 pl of 8 ng/µl ZFN-mRNA solution or a mixture of *MECP2* crRNA (17 ng/µl), tracrRNA (33 ng/µl) and Cas9 nuclease V3 (100 ng/µl) was microinjected into the cytoplasm of GV or 2PN eggs and cultured in vitro for 3–5 days using ISM1 (Discontinued, 11500010A, CooperSurgical, USA) or Sequential Cleav (83040010A, CooperSurgical). Embryos that developed to the 4-cell stage or beyond were transferred into the surrogate mother’s uterus, and pregnancy was confirmed by monitoring blood progesterone levels and using ultrasound imaging equipment. Animals with continued pregnancies periodically underwent ultrasound imaging, i.e., at a frequency of once a week, and founder animals were obtained by spontaneous delivery within the period of 139 to 148 days of gestation. After *MECP2* heterozygous female marmosets (Mut 1) reached sexual maturity, F1 neonates from the *MECP2* mutant marmosets were obtained using eggs from Mut1. After mating with wild-type male marmosets, embryos were collected from the founder marmosets by nonsurgical uterine flushing of naturally fertilized (NAT) embryos^4^. The ovulation cycles of recipient animals were synchronized, and the collected NAT embryos were transferred to them^5^. After embryo transfer, the recipients were tested for pregnancy by measuring plasma progesterone levels and conducting ultrasonography on day 15. For pregnant recipients, fetal development was monitored by ultrasonography once a week until the expected date of delivery.

#### Genomic PCR

To investigate mutations at the target sequences and off-target sequences, we performed genomic PCR. Genomic DNA was extracted from the hair roots or umbilical cords of the marmosets using the DNeasy Blood & Tissue Kit (QIAGEN). The primers used for detection of on-site sequences of *MECP2* ZFN and CRISPR/Cas9 were a forward primer 5′- AAGGACAAGCCCCTCAAGTT-3′ and a reverse primer 5′-TCCTGCTCCATGAGAGATCC-3′.

#### Western blot analysis

After euthanasia with an overdose of ketamine, tissue samples from the occipital cortex, heart, and kidney were collected from both 1 wild-type and 3 *MECP2*-null marmosets (Mut 8, Mut 11 and Mut 12), and 100 mg of each of the tissue samples was homogenized in 1 ml of 2x Laemmli sample buffer (Bio-Rad Laboratories, USA). Samples were loaded into 10% Mini-PROTEAN TGX gels (Bio-Rad Laboratories), and SDS‒PAGE was performed. Protein was transferred to Immun-Blot PVDF membranes (Bio-Rad Laboratories) using the Trans-Blot Turbo System (Bio-Rad Laboratories). After blocking with 4% skim milk in Tris-buffered saline with 0.1% Triton X-100 (TBST) for 30 min, the membrane was incubated in the following antibody solutions overnight at 4 °C: MECP2, 1:500 (Merck, USA); GAPDH, 1:500 (Medical & Biological Laboratories, Japan). Blots were then incubated in the following horseradish peroxidase-conjugated secondary antibodies for 30 min: anti-rabbit IgG, HRP-linked (Bio-Rad Laboratories), anti-mouse IgG, HRP-linked (Bio-Rad Laboratories). HRP was activated using Clarity Western ECL substrate (Bio-Rad Laboratories), and the signal was visualized by a LAS-3000 imaging system (Fujifilm, Japan). The raw data for the cropped western blot images shown in Figure 1E is shown in Figure S13.

#### Monitoring dark-/light-phase activity

An Actiwatch mini device (CamNtech, Ltd, U.K.) is an acceleration sensor-based activity monitor that can record the activity of animals under free movement. An Actiwatch mini device was placed in a pocket attached to a jacket that the marmosets wore or was attached to the neck using a ball chain. Activity data were collected for each marmoset over more than 7 days at a sampling interval of 1 min in units of activity counts. In the RIKEN animal facility, the marmosets were housed under a 12-h dark/12-h light cycle. Daily activity and sleep indexes were analyzed using Actiwatch Activity & Sleep Analysis 7 Software (CamNtech Ltd).

#### MRI scanning

We performed an MRI study to investigate the structural differences between the brains of wild-type and *MECP2* mutant marmosets. First, we evaluated changes in brain volume over time from 1 to 3 months of age. In addition, because achieving long-term survival is unlikely in *MECP2*-null marmosets, we used *MECP2* heterozygous marmosets to investigate changes over time in the whole brain, cerebellum, white matter, and cortex from 1 to 60 months of age. Then, we investigated changes in structural neuronal connections in *MECP2*-null marmoset brains. Age data and other information about the marmosets enrolled in each study are shown in Tables S44 and S45. MRI was performed using a 9.4T BioSpec 94/30 (Biospin GmbH, Germany) unit and a transmitting and receiving volume coil with an 86-mm inner diameter for marmoset brains. T2-weighted imaging and diffusion tensor imaging data were acquired to examine brain volume changes and changes in the neural structural network of these subjects. We obtained T2-weighted images (TR, 4000 ms; TE, 22 ms; rare factor, 4; field of view, 48 × 48 mm^2^; matrix, 178 × 178; no intersection gap; and acquisition time, 7.4 minutes) and diffusion-weighted images (DWI; repetition time, 3,000 ms; echo time, 25.6 ms; b value, 1000 s/mm^2^; gradient directions 32 axis; field of view, 44.8 × 44.8 mm^2^; matrix, 128 × 128; thickness, 0.7 mm; total scan time, 28.8 minutes) from each animal. The animals were scanned in a “sphinx-like” position on an imaging stretcher and administered a mixture of oxygen and 1.5%–1.8% concentrated isoflurane (Abbott Laboratories, USA). During the scan, heart rate, peripheral oxygen saturation (SpO_2_), respiration, and rectal temperature were monitored regularly to manage the animal’s physical condition.

#### Brain volume measurements

All marmoset data were subjected to a skull stripping process to extract brain images from the head. Volume measurements of the brain were then calculated in the following two ways:

1. A registration map derived from a white matter atlas, gray matter atlas, and a 90 brain structure atlas (Table S14) based on a predefined brain atlas^6^ was used to compare brain images and store the deformations.
2. The determinants from the deformation fields of each subject were used to calculate the volume of each structure.

Brain volumes of the whole brain, white matter, gray matter, and 90 regions across the whole brain were compared between wild-type and *MECP2*-null images. Brain volumes were measured at 1, 2, and 3 months of age. We used 2-way ANOVA with age as a covariate and further multivariate correction to evaluate statistically significant differences between groups. We used MATLAB and ITK-SNAP (ITK-SNAP 3.8.0, http://itksnap.org accessed on March 2021) software to segment brain regions and measure the volume of each region.

#### DTI analysis

First, we performed noise reduction on the obtained DWI data^7^. To estimate the inhomogeneous magnetic field, an additional image was acquired with b = 0 as the reversed-phase encoding of the DWI data^8^. Subsequently, the FSL software-based topup and eddy tools were used to correct for B0 field inhomogeneities, eddy currents, and intervolume motion^9^. In addition, the MRtrix software package^10^ was used to estimate high-quality fiber orientation distribution functions and constrained spherical deconvolution^11^ to obtain fiber tractograms. Whole-brain streamline tractography incorporated the anatomically constrained tractography (ACT) framework^12^ and generated 100 million streamlines. Streamline seeding was performed using dynamic seeding, a mechanism that constantly updates the probability of seeding from a particular location in the white matter based on the relative difference between the fiber density estimated from the diffusion model and the currently reconstructed streamline density. We used the spherical-deconvolution informed filtering of tractograms (SIFT2) algorithm as an optimization method to provide a quantitative and biologically meaningful measure of structural connectivity throughout the brain^13^. This algorithm is included in the MRtrix3 software package. We used these whole-brain neural streamlines and 90 brain structure atlases to estimate structural neural connectivity between all brain regions. Moreover, we investigated differences in neural connectivity between wild-type marmosets and *MECP2*-null marmosets at each month of age. In addition, we extracted the neural connections between brain regions that showed statistically significant changes and sufficient effect sizes (> 0.8) monthly to evaluate the relationship between the monthly increases and decreases in neural connectivity. The regions (nodes) used for connections between the same extracted brain regions were weighted by the sum of their respective effect sizes, and the brain regions that seemed to influence differences in network organization were calculated. In this case, we performed the t test with Benjamini‒Hochberg (BH) multiple-testing correction for the significance difference test and calculated the effect size by Cohen’s d.

#### Single-nucleus RNA sequencing

After euthanasia with an overdose of ketamine (50 mg/kg, i.m.), brain tissues were dissected from three wild-type and three *MECP2*-null marmosets, flash frozen in liquid nitrogen for 2 minutes and subsequently kept at -80 °C before nuclear isolation. Brain tissue samples from the PFC and OC were cut into small pieces of < 5 mm and homogenized using a glass Dounce tissue grinder in 0.5 ml of ice-cold EZ PREP buffer (Sigma, USA). Single nuclei were isolated according to the Frankenstein protocol (dx.doi.org/10.17504/protocols.io.bqxymxpw). Isolated single nuclei were sorted using an SH800S Cell Sorter (Sony, Japan) or BD FACSAria II (BD Biosciences, U.S.A.) based on PI fluorescence. Libraries were constructed using the microdroplet-based platform Chromium Single Cell 3’ v3.1 (10× Genomics, USA) according to the manufacturer’s instructions. A DNBSEQ-G400 (MGI, China) or a NovaSeq 6000 (Illumina, USA) was used to sequence 28-and 91-base paired-end reads. Sequenced reads were aligned to the reference genome (Callithrix_jacchus_1700 (mCalJac4)) using Cell Ranger v6.0.1 software (10× Genomics). The data were processed and analyzed using Seurat package v4.1.3^14^. The raw counts were log-normalized and scaled to 10,000 transcripts per cell. Principal component analysis was then performed, and the top 30 principal components were used in implementing the FindClusters function and RunUMAP function. Identifications of cell types were determined by visualizing known marker gene expression within each identified cluster. Excitatory neurons were marked by the expression of *Vesicular Glutamate Transporter 1* (*SLC17A7*) and *glutamate ionotropic receptor NMDA type subunit 1* (*GRIN1*) and were further separated into subtypes by additional single-nucleus RNA-seq data^15^. Interneurons were marked by *Glutamate Decarboxylase 1* (*GAD1*) and *Glutamate Decarboxylase 2* (*GAD2*) and were further separated into subtypes by the expression of *Parvalbumin* (*PVALB*), *Vasoactive Intestinal Peptide* (*VIP*), *Lysosomal Associated Membrane Protein Family Member 5* (*LAMP5*) and *Somatostatin* (*SST*). Astrocytes were marked by the expression of *Aquaporin 4* (*AQP4*) and *Glial Fibrillary Acidic protein* (*GFAP*). Oligodendrocytes were marked by the expression of *Myelin Basic Protein* (*MBP*) and *Proteolipid Protein 1* (*PLP1*). Oligodendrocyte precursor cells were marked by the expression of *Platelet Derived Growth Factor Receptor Alpha* (*PDGFRA*) and *Chondroitin Sulfate Proteoglycan 4* (*CSPG4*). Microglia was marked by the expression of *C-X3-C Motif Chemokine Receptor 1* (*CX3CR1*) and *Integrin Subunit Alpha M* (*ITGAM*). Endothelial cells were identified based on the expression of *Von Willebrand Factor* (VWF) and *TEK Receptor Tyrosine Kinase* (*TEK*). DEGs between wild-type and *MECP2*-null cell types were identified using the FindMarkers function. Single-nucleus RNA-seq transcriptome data on *Mecp2* mutant mice and RTT patients were obtained from a previous paper^16^. Gene ontology analysis was performed using g:Profiler^17^ (Fisher’s one-tailed test, adjusted *p* value < 0.05), and the top 30 GO terms were classified using CateGOrizer^18^.

#### Visium analysis

The marmosets were deeply anesthetized with a lethal dose of mixed anesthesia (0.4 mg/kg medetomidine, 2 mg/kg midazolam, and 5 mg/kg butorphanol, i.p.), followed by secobarbital (100 mg/kg, i.p.). After three failed attempts to elicit a foot withdrawal reflex, the brain was dissected out and placed in ice-cold phosphate-buffered saline (PBS). The brain was immediately coronally cut into 3–4 mm thick slices, and the coronal brain slices were frozen in OCT compound (Sakura Finetech, Japan). Coronal cryosections of 10 mm thickness (CM1950, Leica Biosystems, Germany) were mounted on Visium Tissue Optimization Slides (PN-1000192, 10x) and Visium Spatial Gene Expression Slides (PN-1000185, 10x) following the company’s instructions. The subsequent experimental work was conducted by KOTAI Biotechnologies (Osaka, Japan). The detailed experimental conditions and results are summarized in Table S46. In brief, the Visium Spatial Tissue Optimization Reagent Kit (PN-1000192, 10x) was used for tissue optimization, and the Visium Spatial Gene Expression Reagent Kit (PN-1000186, 10x), Library Construction Kit (PN-1000215, 10x), and Dual Index Kit TT Set A (PN-1000215) were used for gene expression analysis (GEX slide). Total RNA was extracted using an RNeasy Mini Kit (QIAGEN), and the RNA Integrity Number (RIN) was calculated using Tapestation (Agilent). For sequencing, a NovaSeq S4 system on a NovaSeq 6000 (Illumina, USA) was used. Raw FASTQ files and staining images were processed by Space Ranger software version 1.3.1, with Callithrix_jacchus_1700 (mCalJac4) as the reference genome. The data were processed and analyzed using Seurat package v4.1.3 and Loupe Browser 6.4.1. The raw counts were normalized using the SCTransform function. Principal component analysis was then performed, and the top 30 principal components were used in the implementation of the FindNeighbors function and RunUMAP function. Identifications of cortical layers were determined by known marker gene expression and histological locations. DEGs between wild-type and *MECP2*-null cortical layers were identified using the FindMarkers function. Gene expression in Figure 7E was visualized using the BayesSpace function^19^.

#### Immunohistochemistry

For immunohistochemistry, wild-type and *MECP2*-null male marmosets at 4 months of age were euthanized with an overdose of ketamine (50 mg/kg, i.m.), and brain tissues were fixed in 4% paraformaldehyde (PFA) in PBS buffer for two days, cryoprotected for 3 days in 30% sucrose in PBS, and embedded in OCT compound (Sakura Finetech). Frozen brain tissues were sectioned coronally at 50 μm on a freezing microtome (Leica Biosystems). Sections were incubated in blocking solution (PBS, 0.5% BSA, 1% goat serum, and 0.3% Triton X-100) for 30 min at room temperature and then incubated in primary antibody overnight at 4 °C. Primary antibodies were used at the following concentrations: rabbit anti-MECP2 antibody (1:200, Merck) and mouse anti-NeuN antibody (1:200, Merck). Sections were washed three times in PBS for 10 min and then incubated with secondary antibody for 2 hours at room temperature. Secondary antibodies were used at the following concentrations: goat anti-rabbit IgG antibody Alexa Fluor 488 (1:500, Thermo Fisher Scientific) and goat anti-mouse IgG antibody Alexa Fluor 555 (1:500, Thermo Fisher Scientific). Sections were washed three times in PBS, rinsed in distilled deionized water (DDW), and mounted on glass slides.

#### Sociability test

The social testing apparatus consisted of an experimental cage (width 72 cm × height 72 cm × depth 60 cm) with a motion-sensing video camera (Ohara & Co., Japan). Two transparent acrylic small boxes (width 16 cm × height 16 cm × depth 20 cm) with either a cage-mate marmoset or an object (a roll of packing tape) were placed in the corners of the experimental cage. The location of the marmoset/object in the left or right corner was systematically alternated in three trials. After the subject marmoset was habituated to the experimental cage without a cage-mate marmoset or an object for 1 hour, the subject marmoset was first placed in the middle of the experimental cage and allowed to explore the entire cage for a 15 min session. The movement of the subject marmosets was recorded by a motion sensing video camera, and the position of the subject marmosets was analyzed using Marmo Detector version 2.0 (Ohara & Co., Japan). The cage space was divided into 12 compartments (width 24 cm × height 36 cm × depth 30 cm), and the amount of time spent in the two compartments with the two transparent acrylic small boxes was recorded.

#### Golgi staining

For dendritic morphology, soma size and spine density analyses, 3 wild-type and 2 *MECP2*-null male marmosets were analyzed at 4 months of age. Marmosets were euthanized with ketamine overdose, and their brains were fixed in 4% paraformaldehyde (PFA) in PBS buffer for two days and sectioned at 100 μm with a vibrating microtome VT1200S (Leica Biosystems). Brain sections were processed according to the protocol of the sliceGolgi kit (Bioenno Tech, USA). Pyramidal neurons were randomly selected for analysis and drawn by an investigator blinded to genotype using a camera lucida device. Concentric circles in 20-μm radius increments were drawn around the center of the reconstructed soma, and the number of dendrites crossing each circle was counted for Sholl analysis. Images of the soma and spine were captured using a Keyence BZ-X710 microscope (Keyence, Japan), and soma size was measured using BZ-X Viewer software (Keyence).

#### Breathing analysis

The breathing pattern of unrestrained marmosets was analyzed using a whole-body flow plethysmograph (EMKA Technologies, France). In a whole-body flow plethysmograph, a constant flow pump connected to the animal chamber provided a continuous inflow of fresh air avoid interrupting the recording, and the experimental room was maintained at 25 °C. The spirogram was obtained by recording the pressure difference between the subject and the reference chambers for 15 min. The signal was amplified, filtered, input into an analog-to-digital converter (sampling frequency 1 kHz) and stored on a PC disk. The data were analyzed by IOX2 software (EMKA Technologies). Because calls and movements of marmosets disrupt the spirogram, we videotaped the chamber and analyzed breathing only when the marmosets were quiet.

#### Statistical analysis

Statistical analyses were performed using Prism 8 (Graph Pad, USA), Seurat^14^, MATLAB (MathWorks, Inc., USA), or Microsoft Excel (Microsoft, USA). Data were analyzed using the statistical methods noted in the figure legends. Values are expressed as the mean ± SEM unless noted otherwise.

## Figure Legends

**Figure S1.**
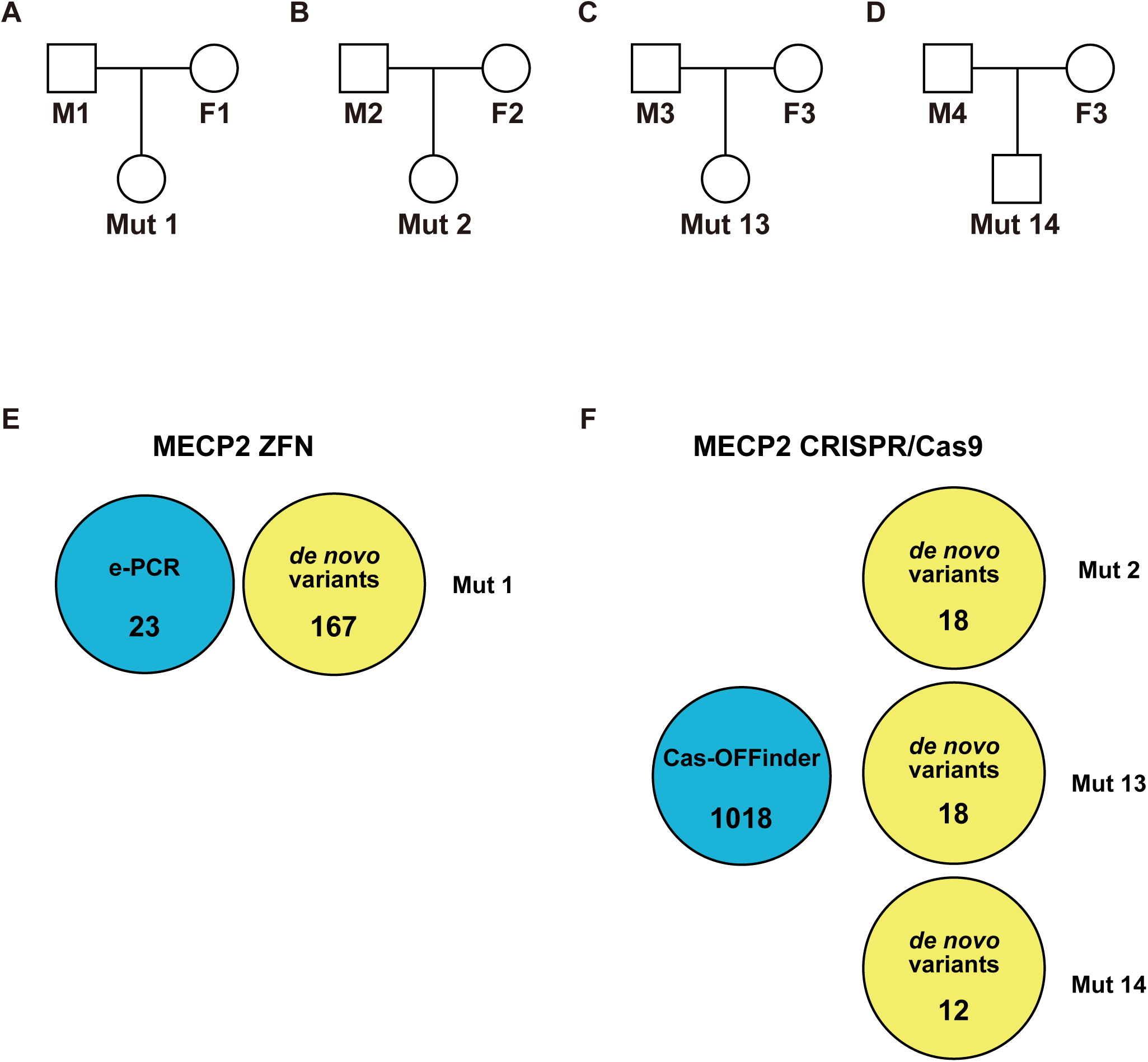
Off-target analysis. (A-D) Family tree of four *MECP2*-null marmoset families. Square, males; oval, females. (E) Venn diagram of in silico predicted potential MECP2 ZFN off-target sites (e-PCR, length < 100, mismatch < 11, gap < 3) and de novo mutations of Mut 1. (F) Venn diagram of in silico predicted potential MECP2 CRISPR/Cas9 off-target sites (Cas-OFFinder, mismatch < 6) and de novo mutations of Mut 2, Mut 13 and Mut 14.

**Figure S2.**
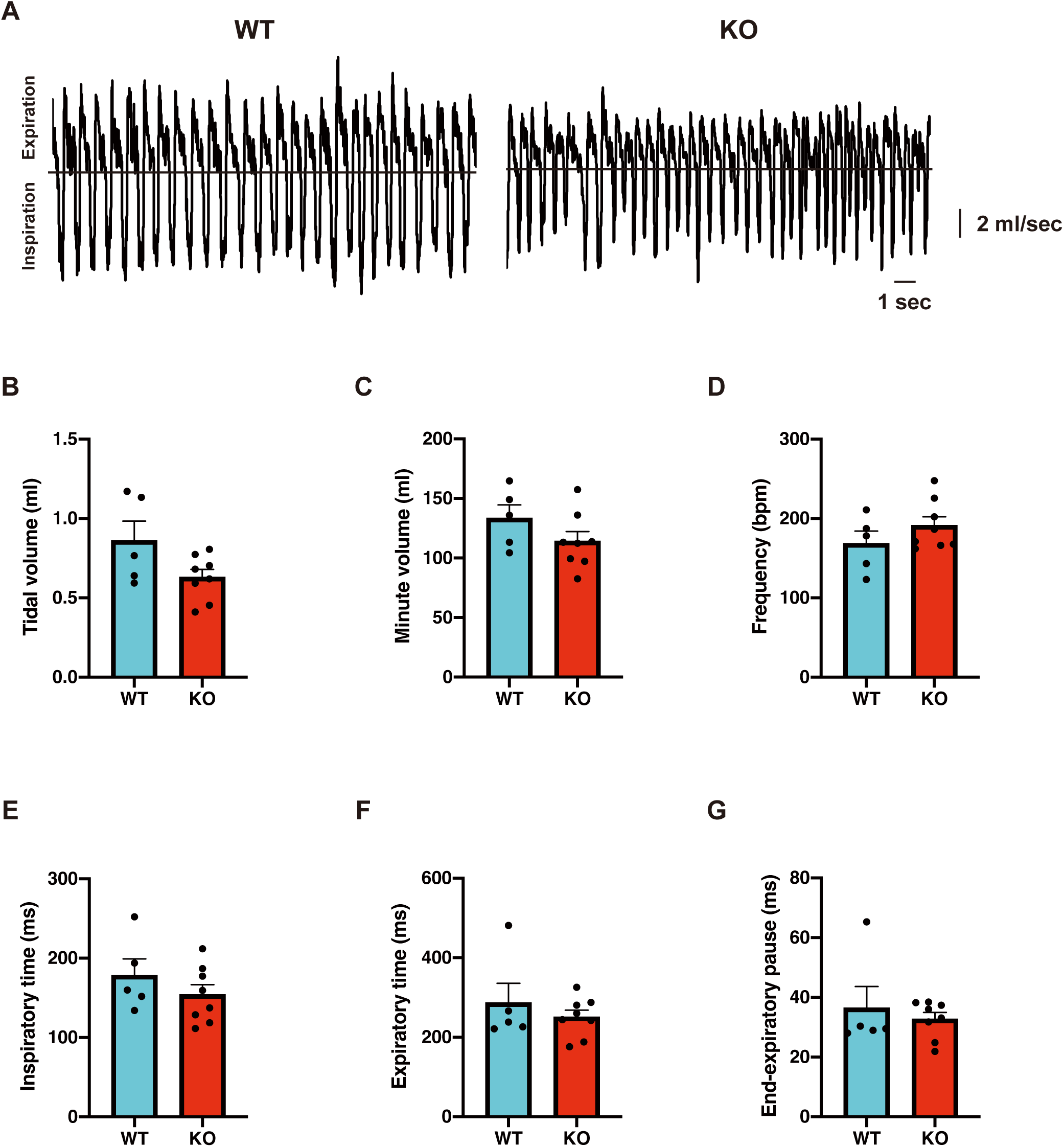
Breathing patterns of *MECP2*-null marmosets using a whole-body flow plethysmograph. (A) Plethysmographic recording of breathing at 3 months of age in wild-type (WT) and *MECP2*-null marmosets (KO). (B-G) No differences were detected in tidal volume (B), minute volume (C), respirator rate (D), inspiratory time (E), expiratory time (F) or end-expiratory pause (G) in *MECP2*-null marmosets compared to wild-type marmosets. Values represent the mean ± SEM, Student’s t test, WT: n = 5, KO: n = 8.

**Figure S3.**
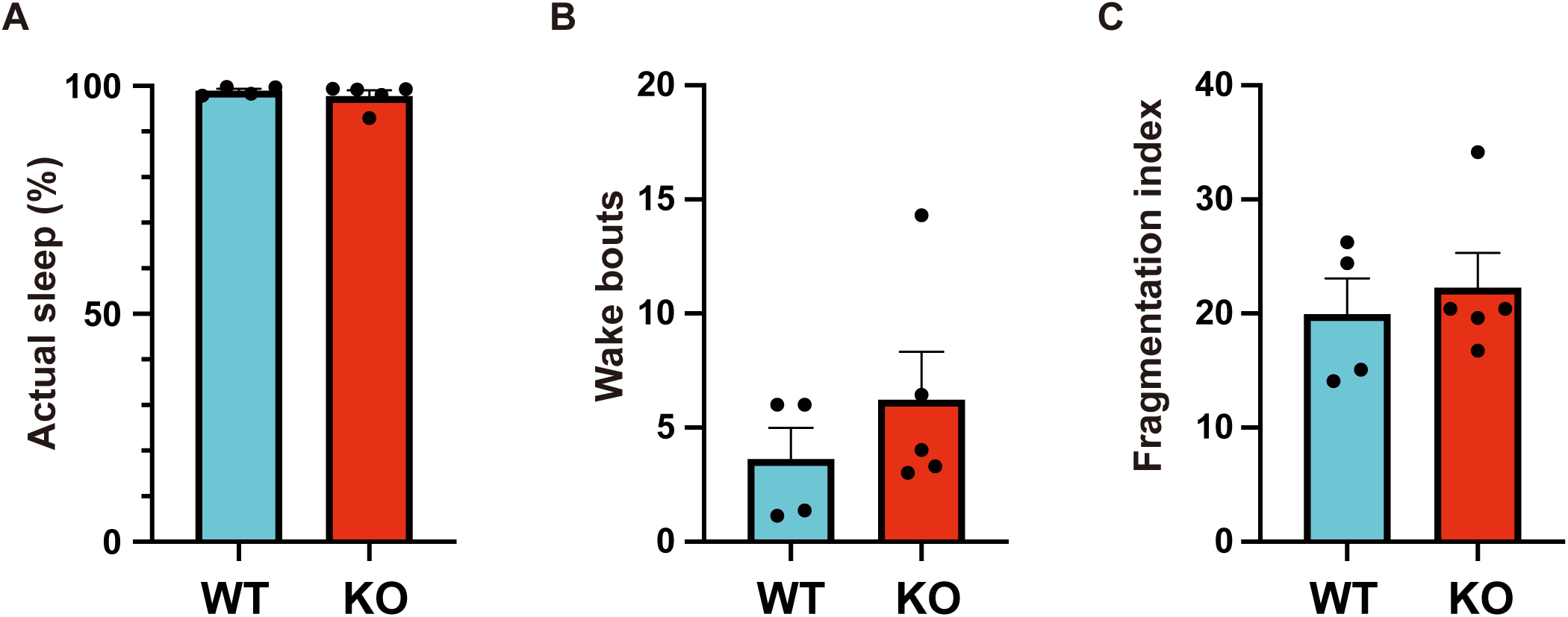
No sleep disturbances were detected in the *MECP2*-null marmosets. (A-C) Using an Actiwatch mini device, the percentage of actual sleep time during the 12 h dark phase (A), number of wake bouts (B) and fragmentation index, an indicator of restlessness (C), during the dark phase were analyzed; *MECP2*-null marmosets (KO) displayed no sleep disturbances compared to wild-type marmosets (WT). Values represent the mean ± SEM, Student’s t test, WT: n = 5, KO: n = 5.

**Figure S4.**
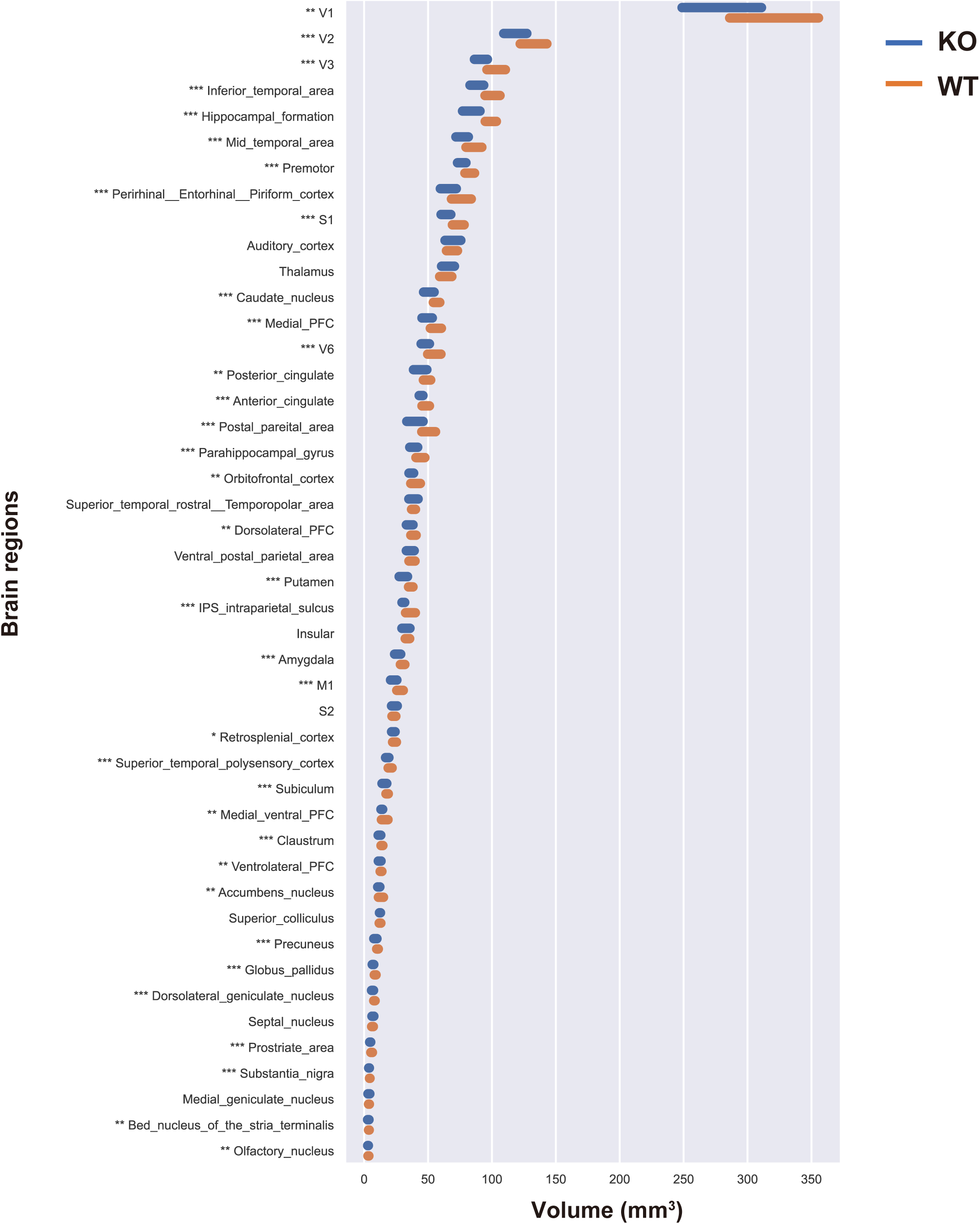
Comparison of the volumes of 45 brain regions between wild-type and *MECP2*-null marmosets at 1 month of age. The volumes of 45 brain regions (see Table 14) of wild-type (WT) and *MECP2*-null (KO) marmosets were analyzed by MRI at 1 month of age. The length of the bars in each region indicates the width of the standard deviation. Multiple t tests, FDR < 0.1, * *q* < 0.1, ** *q* < 0.05, *** *q* < 0.01.

**Figure S5.**
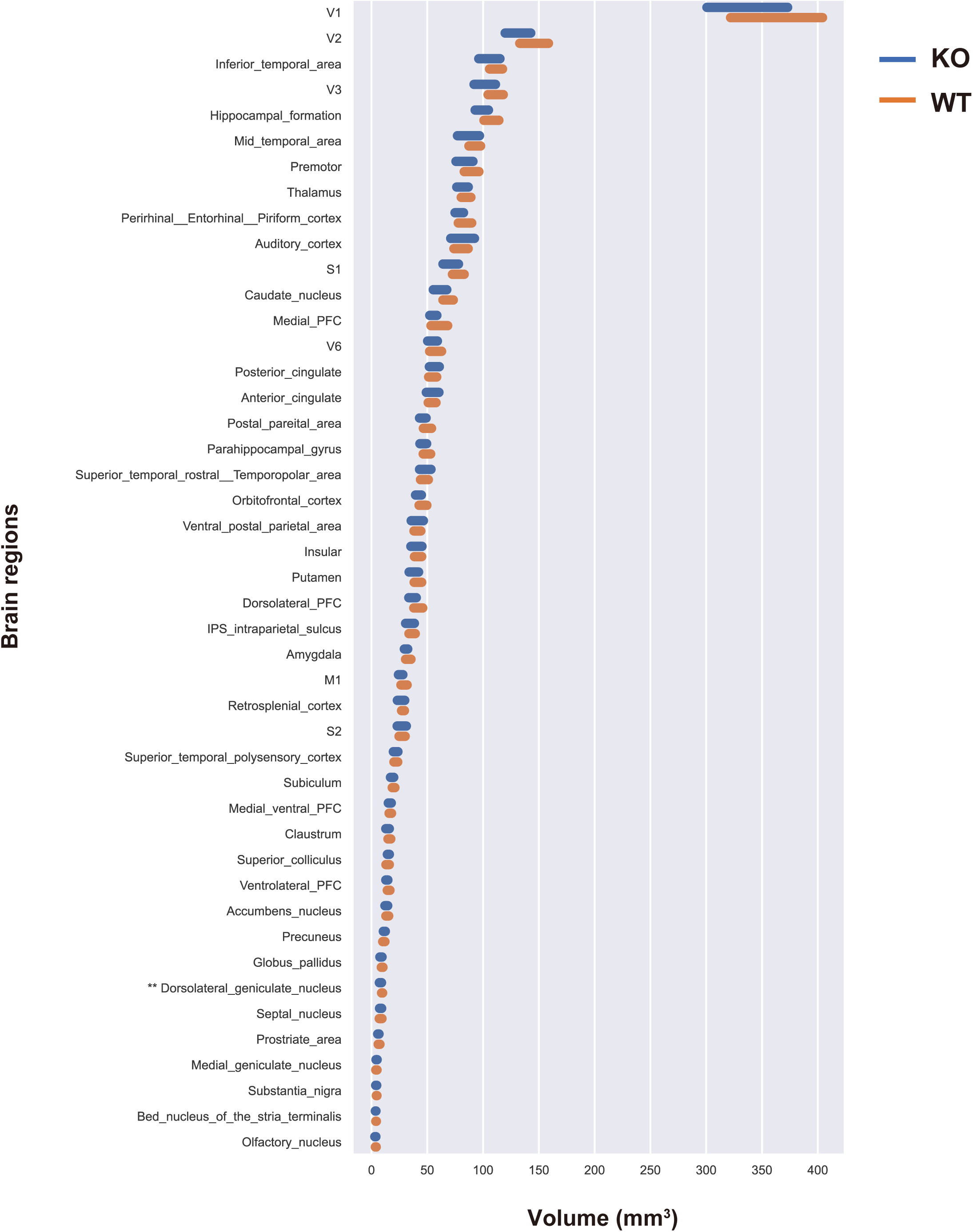
Comparison of the volumes of 45 brain regions between wild-type and *MECP2*-null marmosets at 2 months of age. The volumes of 45 brain regions (see Table 14) of wild-type (WT) and *MECP2*-null (KO) marmosets were analyzed by MRI at 2 months of age. The length of the bars in each region indicates the width of the standard deviation. Multiple t tests, FDR < 0.1, * *q* < 0.1, ** *q* < 0.05, *** *q* < 0.01.

**Figure S6.**
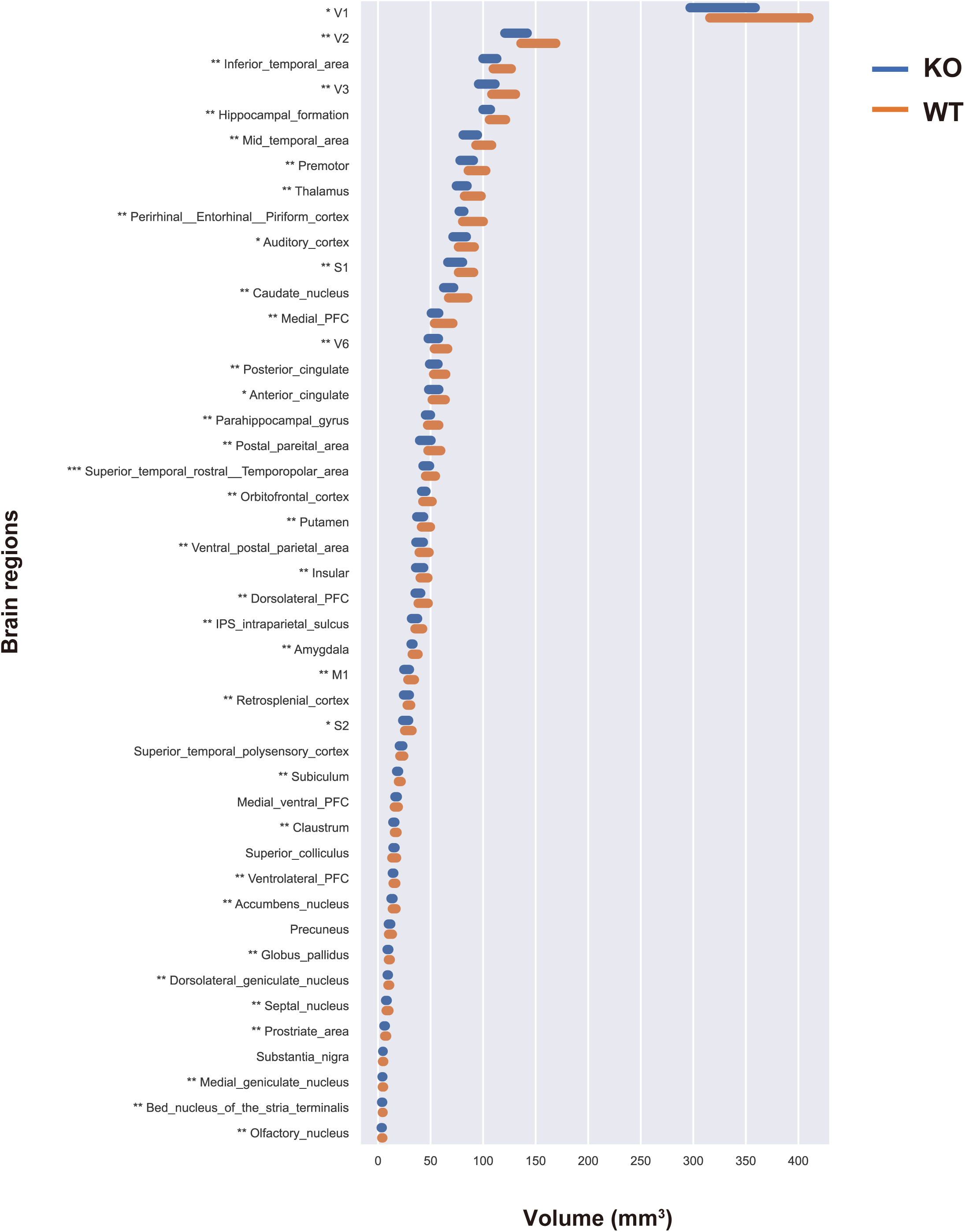
Comparison of the volumes of 45 brain regions between wild-type and *MECP2*-null marmosets at 3 months of age. The volumes of 45 brain regions (see Table 14) of wild-type (WT) and *MECP2*-null (KO) marmosets were analyzed by MRI at 3 months of age. The length of the bars in each region indicates the width of the standard deviation. Multiple t tests, FDR < 0.1, * *q* < 0.1, ** *q* < 0.05, *** *q* < 0.01.

**Figure S7.**
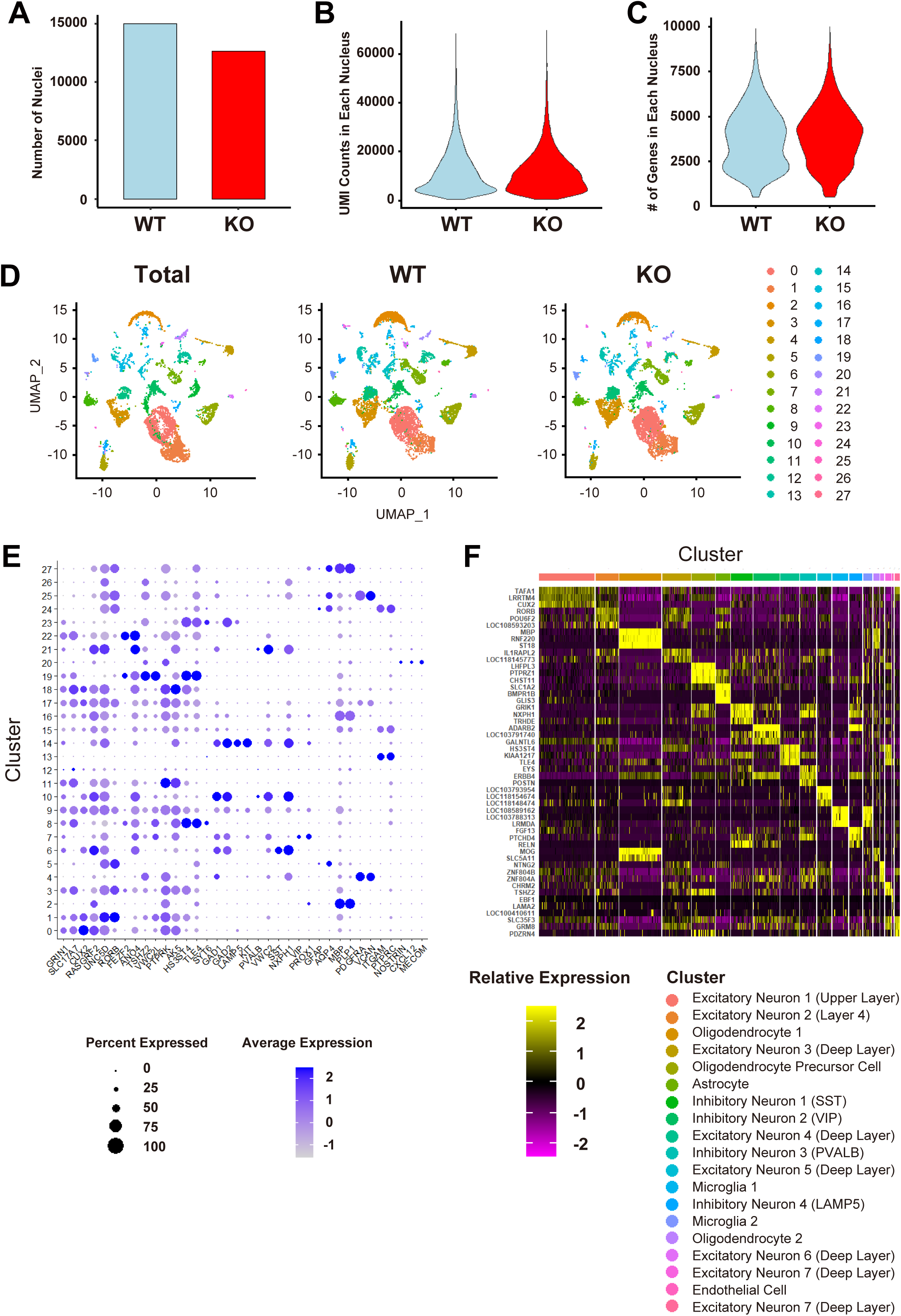
Summary and quality control analyses for single-nucleus RNA-Seq using PFC tissues. (A-C) Bar graph displaying the number of nuclei (A) and violin plots displaying the distribution of unique molecular identifiers (UMIs) (B) and the number of genes (C) detected in wild-type (WT) and *MECP2*-null (KO) PFC tissues obtained at 4 months of age. (D) UMAP dimensional reduction of single-nucleus RNA sequencing of PFC. (E) Dot plot of scaled expression of selected marker genes in the identified clusters. The size of the dot indicates the percentage of nuclei that expressed a given gene within a cluster, while the color encodes the average expression level across all nuclei within a cluster. (F) Heatmap revealing the scaled expression of the top 3 differentially expressed genes for each cell type.

**Figure S8.**
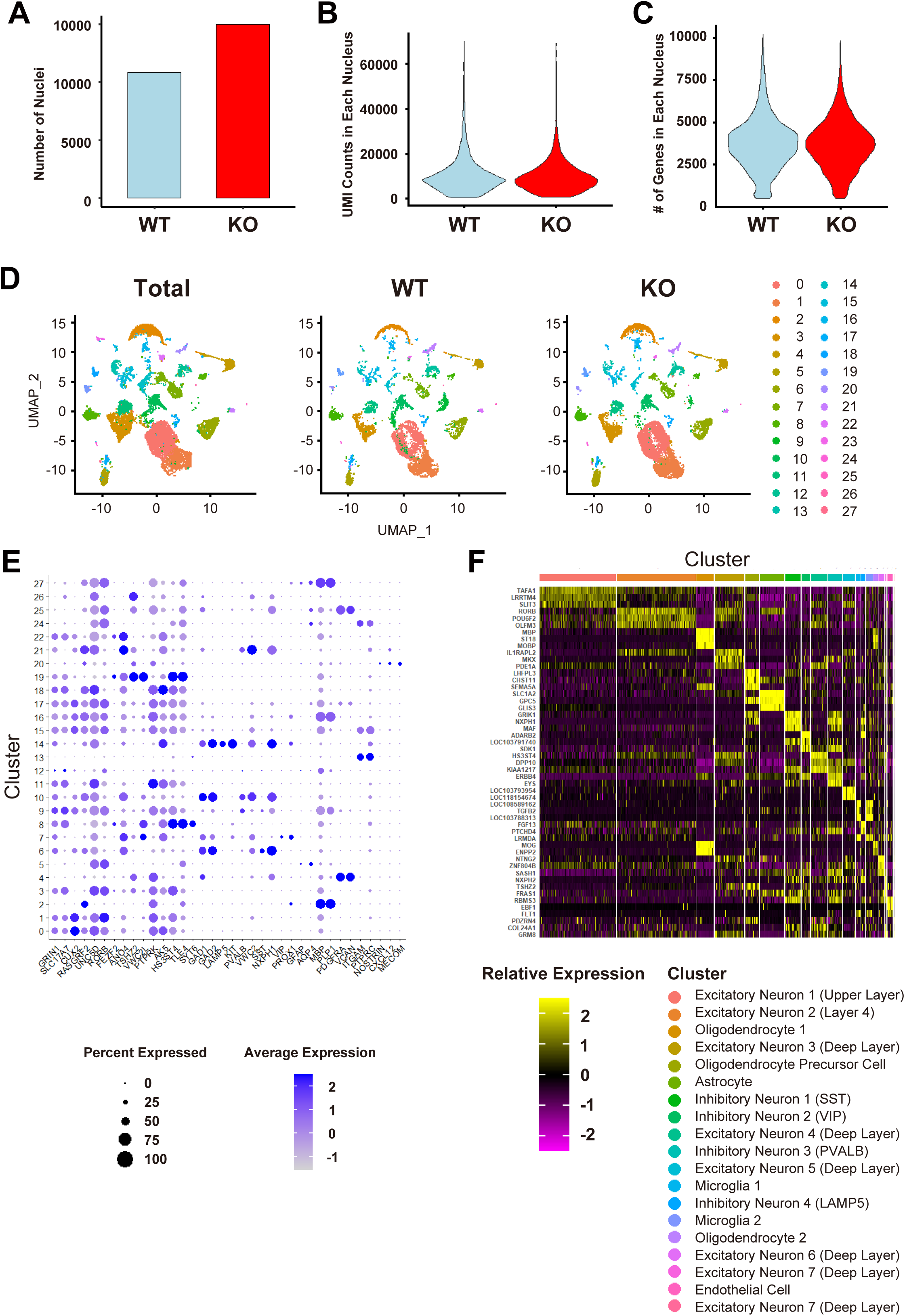
Summary and quality control analyses for single-nucleus RNA-Seq using OC tissues. (A-C) Bar graph displaying the number of nuclei (A) and violin plots displaying the distribution of UMIs (B) and the number of genes (C) detected in wild-type (WT) and *MECP2*-null (KO) OC tissues obtained at 4 months of age. (D) UMAP dimensional reduction of single-nucleus RNA sequencing of OC. (E) Dot plot of scaled expression of selected marker genes in the identified clusters. The size of the dot encodes the percentage of nuclei that expressed a given gene within a cluster, while the color encodes the average expression level across all nuclei within a cluster. (F) Heatmap revealing the scaled expression of the top 3 differentially expressed genes for each cell type.

**Figure S9.**
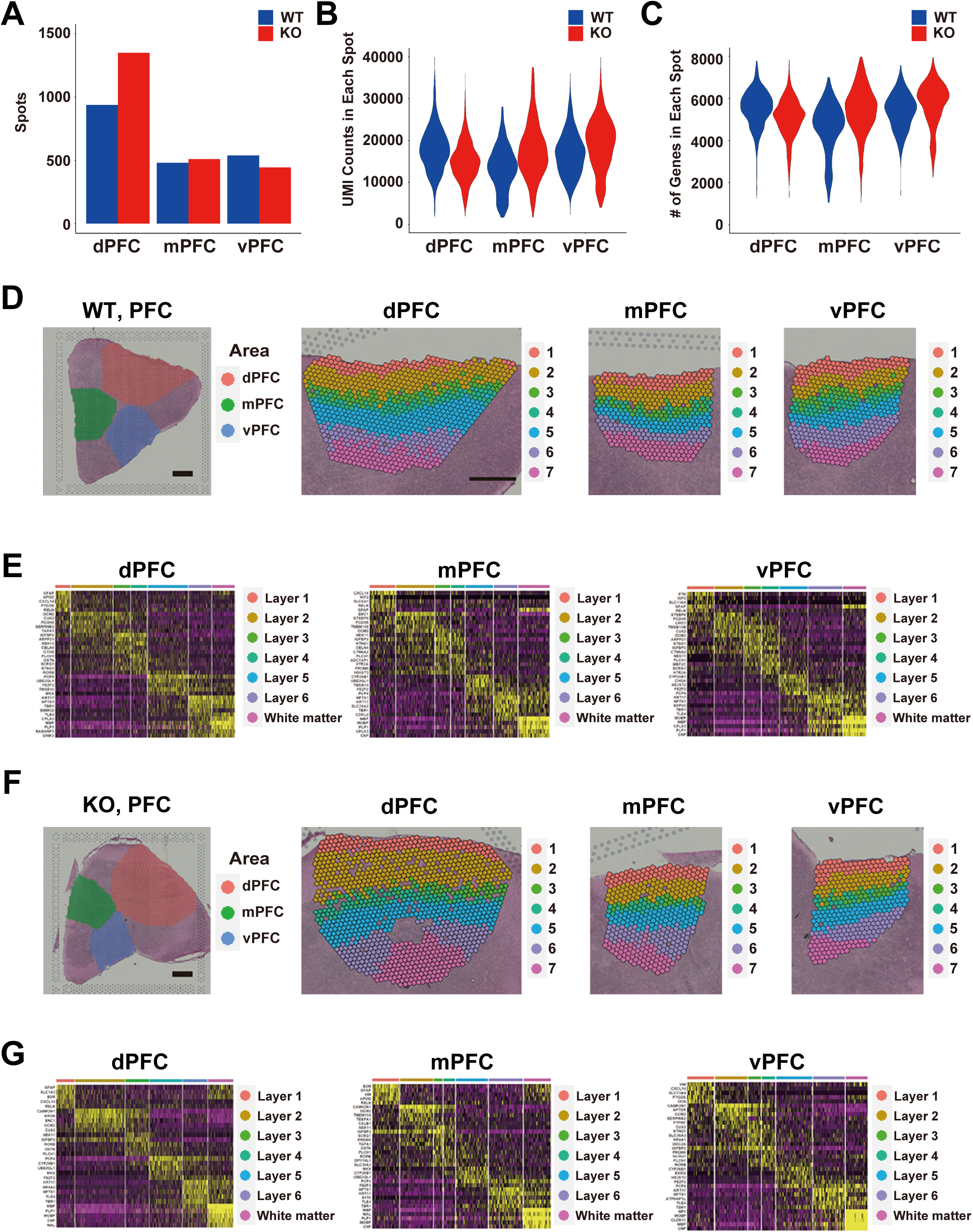
Summary and quality control analyses for Visium spatial transcriptome using PFC tissues. (A-C) Bar graph displaying the number of nuclei (A) and violin plots displaying the distribution of UMIs (B) and the number of genes (C) detected in the wild-type (WT) and *MECP2*-null (KO) dPFC, mPFC and vPFC at 3 months of age. (D and E) Supervised annotation of wild-type PFC layers (D) based on selected layer-specific gene markers and cytoarchitecture and heatmaps revealing the scaled expression of 5 uniquely expressed genes for each layer (E). (F and G) Supervised annotation of *MECP2*-null PFC layers (F) based on selected layer-specific gene markers and cytoarchitecture and heatmaps revealing the scaled expression of 5 uniquely expressed genes for each layer (G).

**Figure S10.**
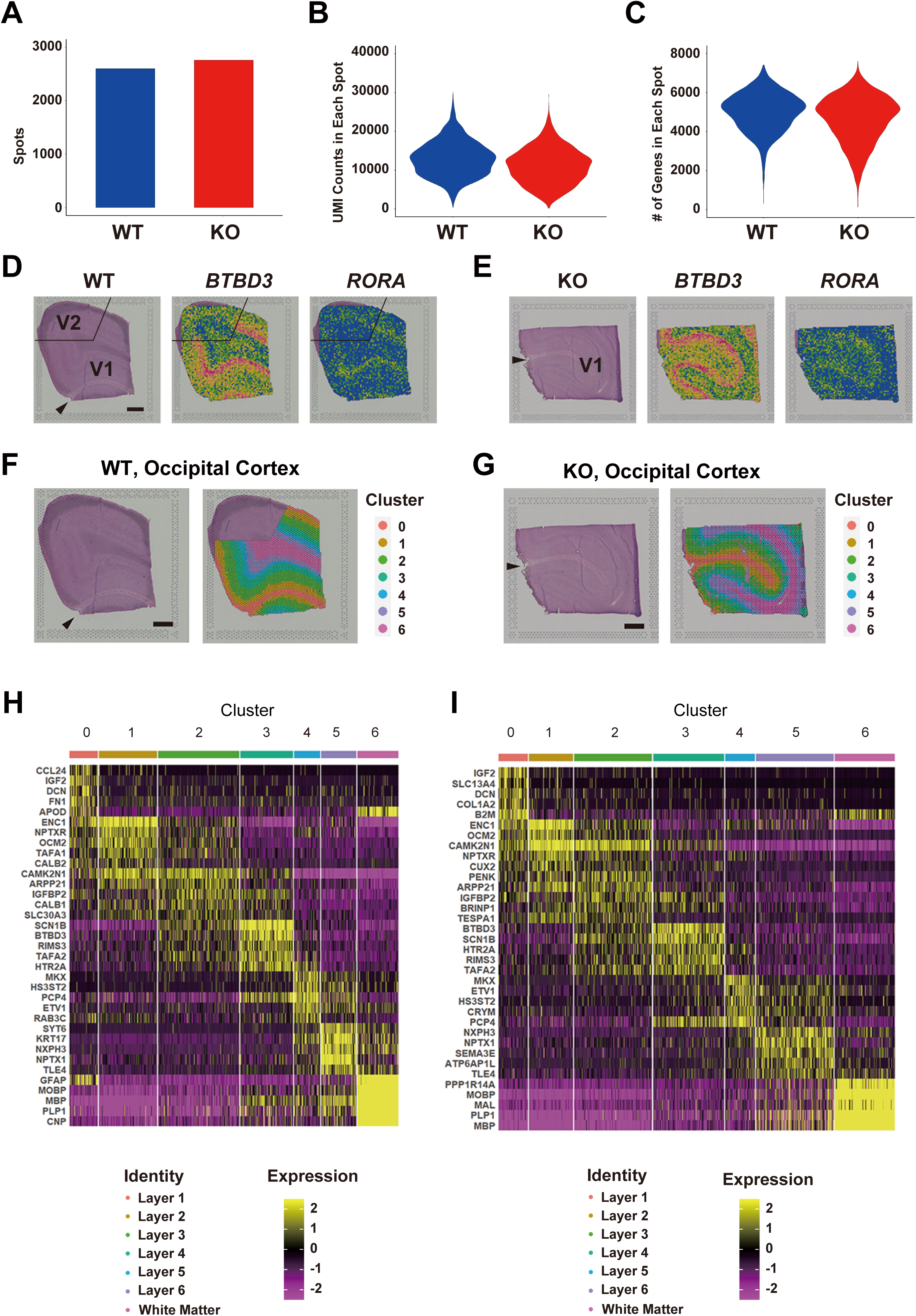
Summary and quality control analyses for Visium spatial transcriptome using OC tissues. (A-C) Bar graph displaying the number of nuclei (A) and violin plots displaying the distribution of UMIs (B) and the number of genes (C) detected in wild-type (WT) and *MECP2*-null (KO) OC samples obtained at 3 months of age. (D and E) To determine the cortical area of V1 and V2, the expression of *BTBD3* and *RORA* was analyzed in wild-type (D) and *MECP2*-null marmosets (E). (F-I) Supervised annotation of wild-type V1 layers (F) based on selected layer-specific gene markers and cytoarchitecture and heatmaps revealing the scaled expression of 5 uniquely expressed genes for each layer (H). Supervised annotation of *MECP2*-null V1 layers (F) based on selected layer-specific gene markers and cytoarchitecture and heatmaps revealing the scaled expression of 5 uniquely expressed genes for each layer (H).

**Figure S11.**
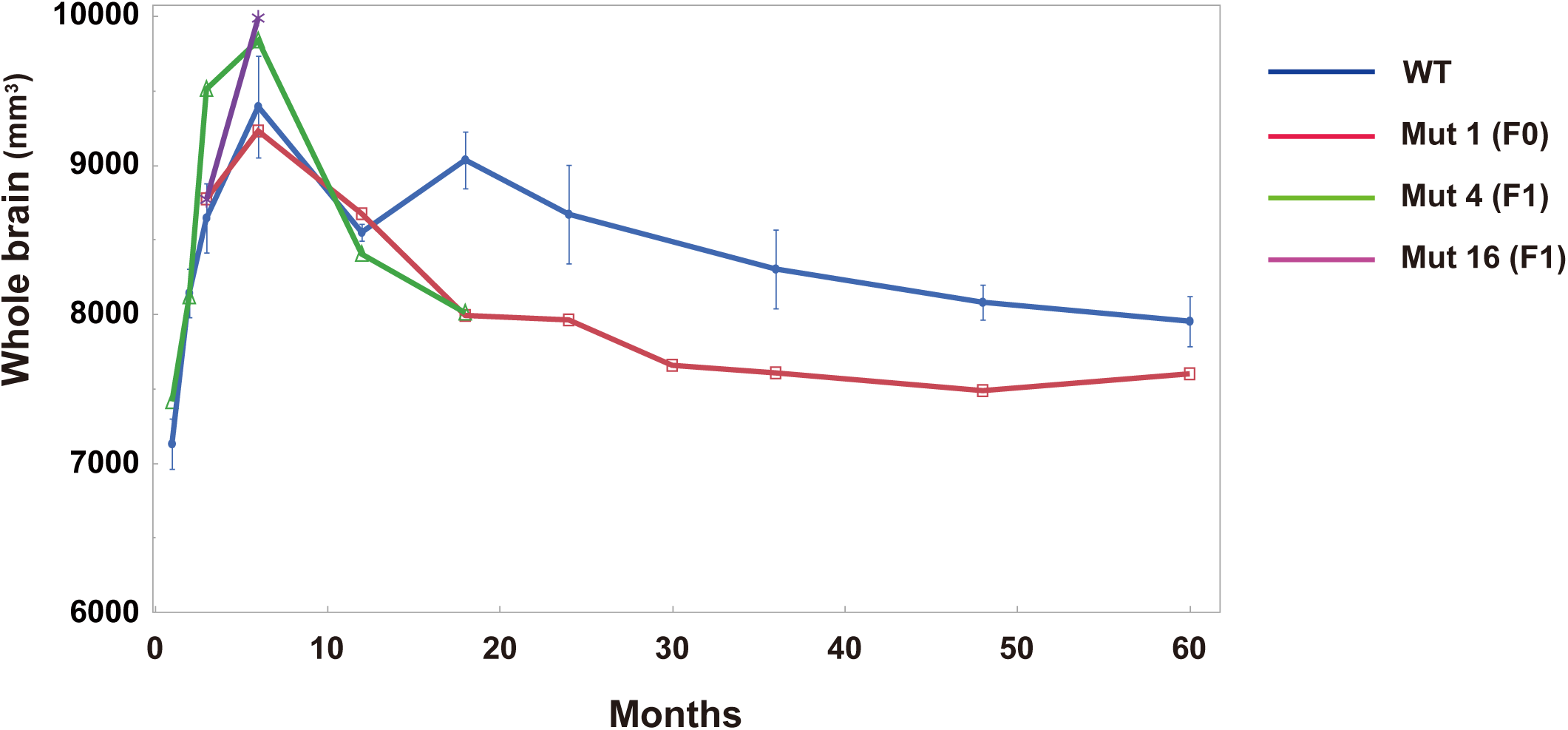
Brain growth of *MECP2* heterozygous marmosets. Whole-brain volumes of wild-type (WT) and three female *MECP2* heterozygous marmosets, Mut 1 (founder), Mut 4 (F1 generation) and Mut 16 (F1 generation), were analyzed based on MRI data obtained monthly from 1 to 60 months of age. Values represent the mean ± SD of the brain volume of WT marmosets.

**Figure S12.**
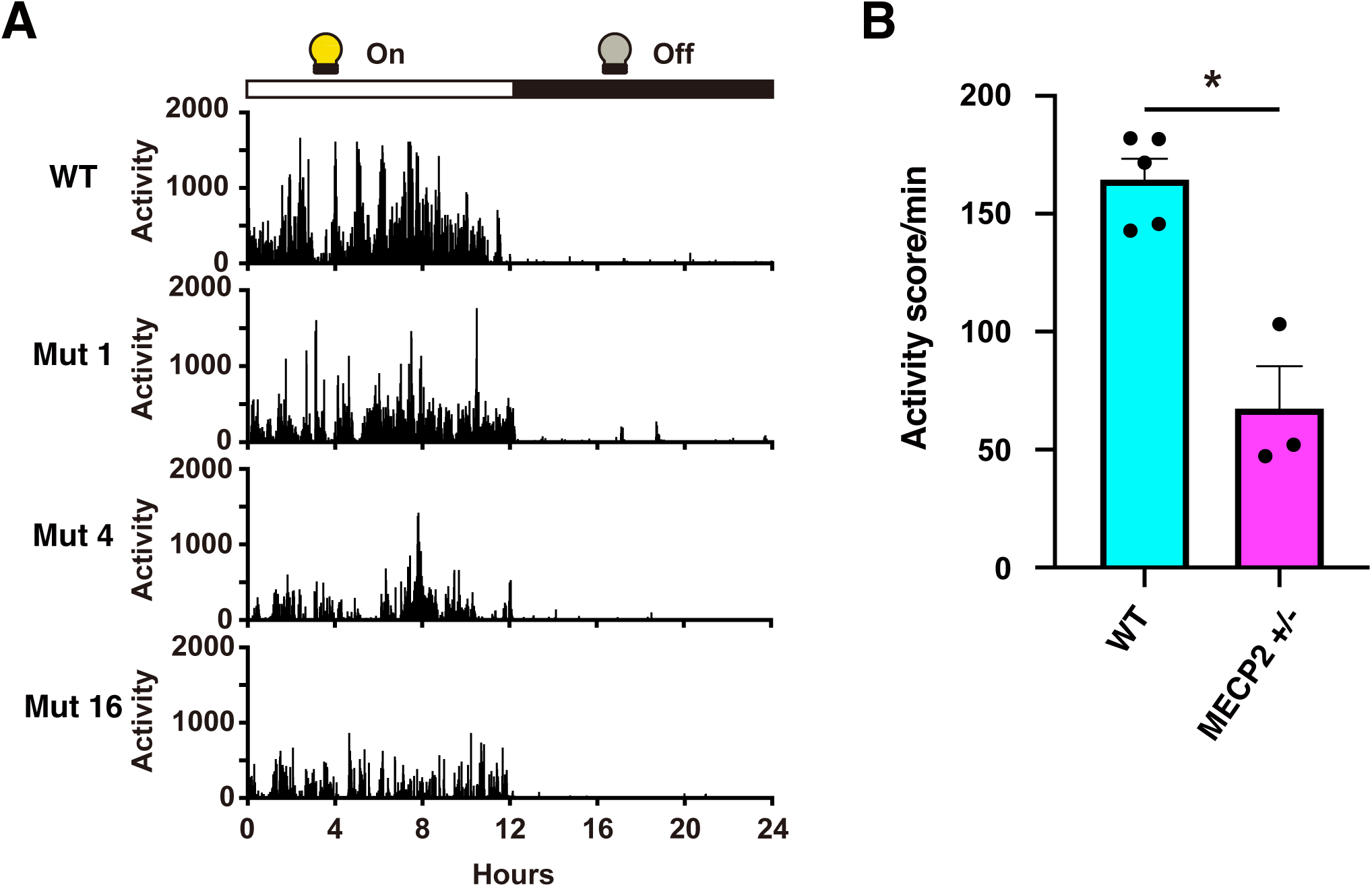
Activity of *MECP2* heterozygous marmosets. (A) Activity scores of wild-type (WT) and 3 *MECP2* heterozygous marmosets (Mut 1, Mut 4 and Mut 16) at 1 year of age were monitored for 7 days while in their home cages using an Actiwatch-Mini® device. Activity was plotted for every minute of each day (light phase for 12 hours and dark phase for 12 hours). (B) The three *MECP2* heterozygous marmosets showed a reduction in voluntary activity in the light phase compared to WT marmosets. Data are presented as the mean ± SEM. Student’s T test, WT: n = 5, *MECP2* +/-: n = 3, * *p* < 0.01.

**Figure S13.**
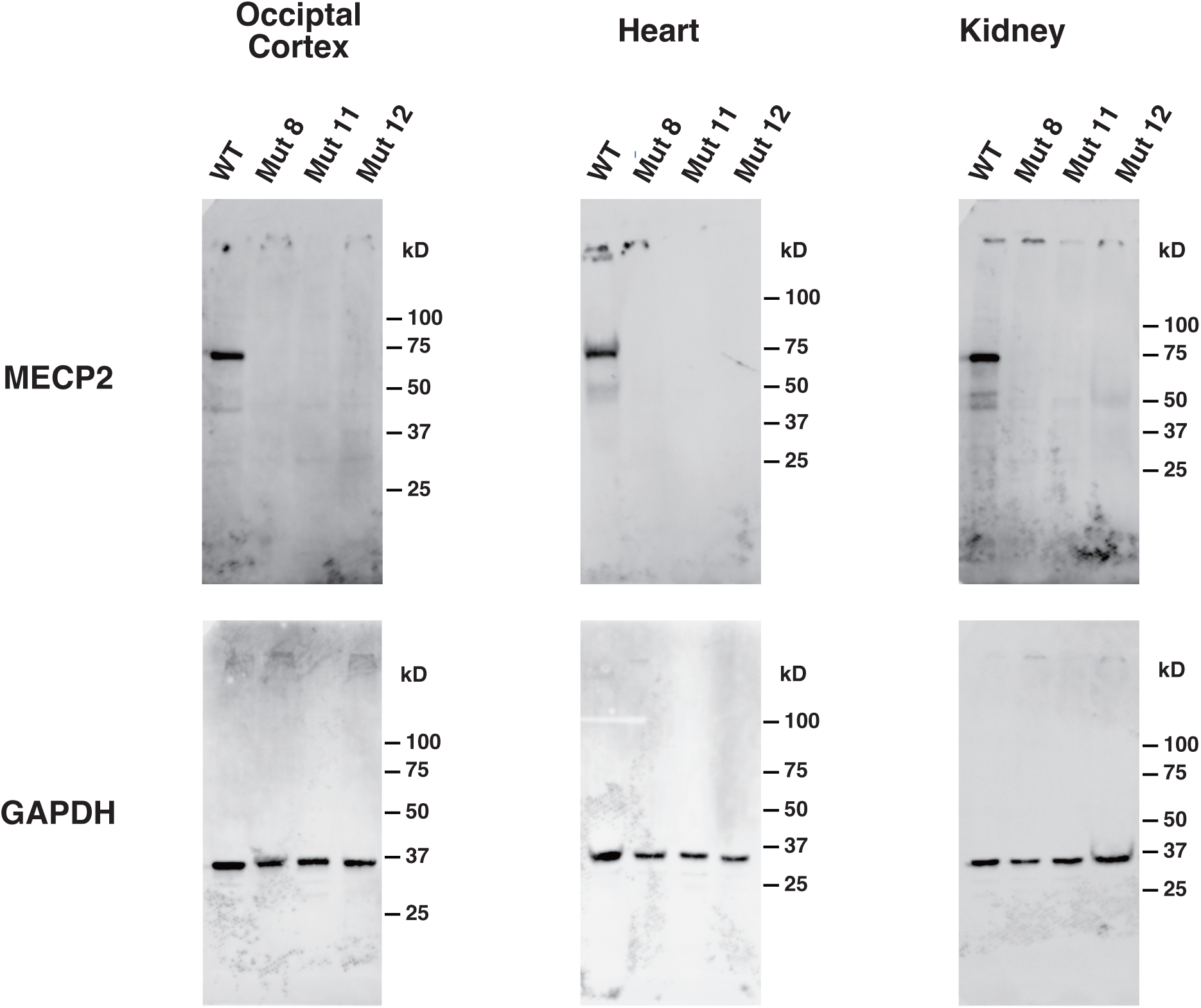
Raw data for cropped western blot images shown in Figure 1E. The raw data for cropped western blot images of MECP2 and GAPDH in occipital cortex, heart, and kidney protein extracts from WT and KO marmosets (Mut 8, Mut 11 and Mut 12) shown in Figure 1E.

---

**Table S1:** Summary of microinjection and obtained neonates.

| Construction | Transferred embryos | Recipients | Pregnancies (%) | Neonates | Mutant animals |
| --- | --- | --- | --- | --- | --- |
| eHiFi-ZFN | 88 | 86 | 13 (15%) | M: 8, F: 3 | F: 1 |
| CRISPR/Cas9 | 136 | 57 | 21 (36%) | M: 10, F: 5 | M: 7, F: 5 |
| Total | 224 | 143 | 34 (24%) | M: 18, F: 8 | M: 7, F: 6 |

M, male; F, female

**Table S2:**
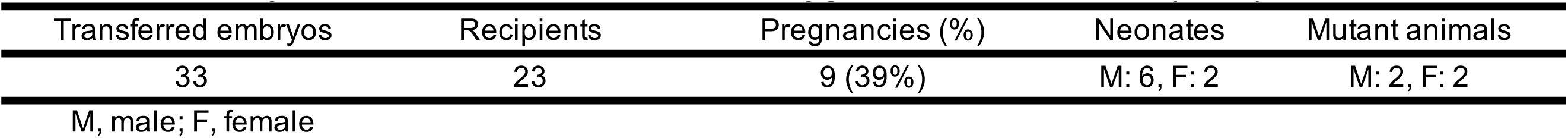
Summary of F1 neonates from *MECP2* heterozygous founder marmoset (Mut 1)

| Transferred embryos | Recipients | Pregnancies (%) | Neonates | Mutant animals |
| --- | --- | --- | --- | --- |
| 33 | 23 | 9 (39%) | M: 6, F: 2 | M: 2, F: 2 |

M, male; F, female

**Table S3.**
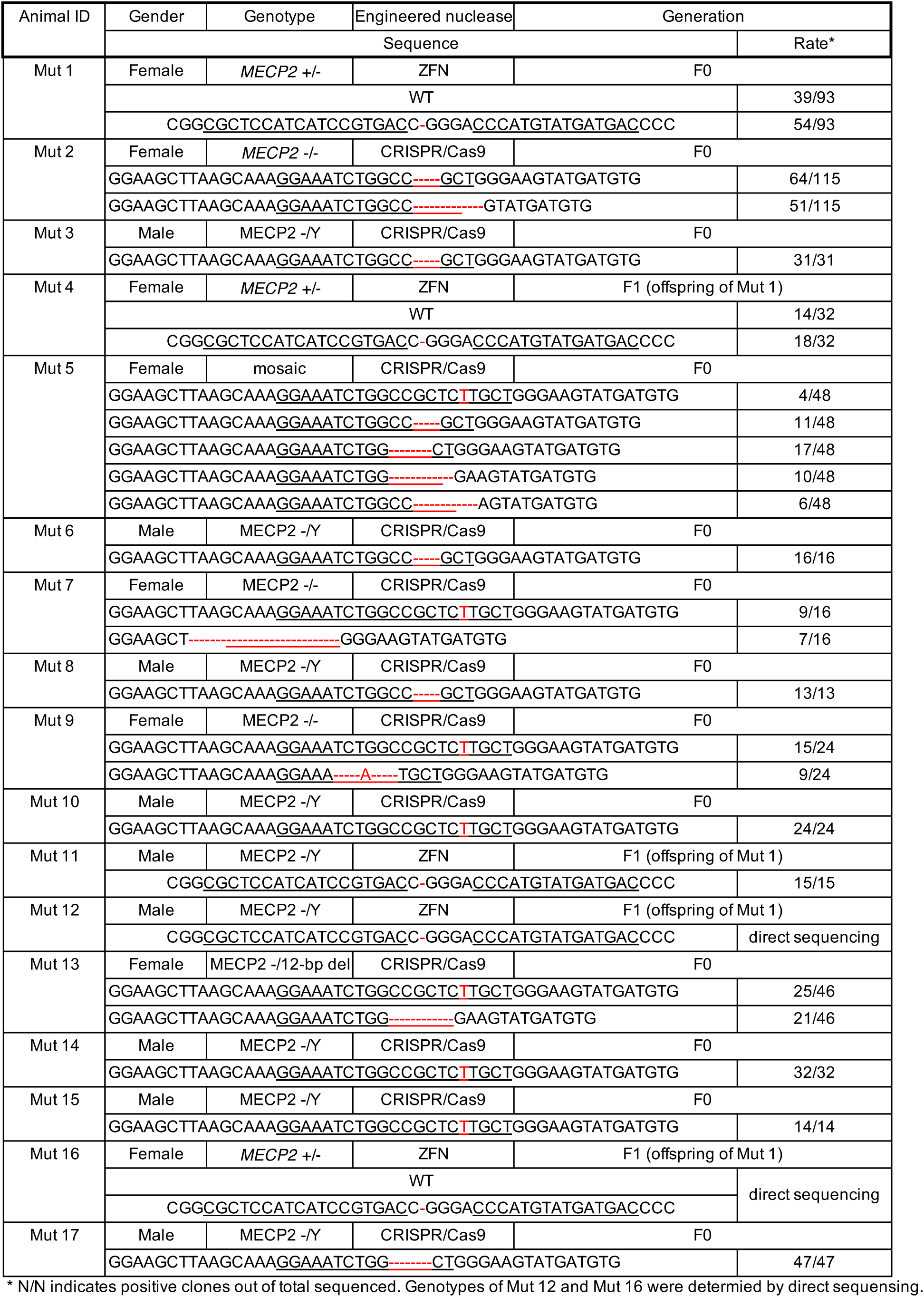
Summary of mutations in the target sequences in the *MECP2* locus.

**Table S4.**
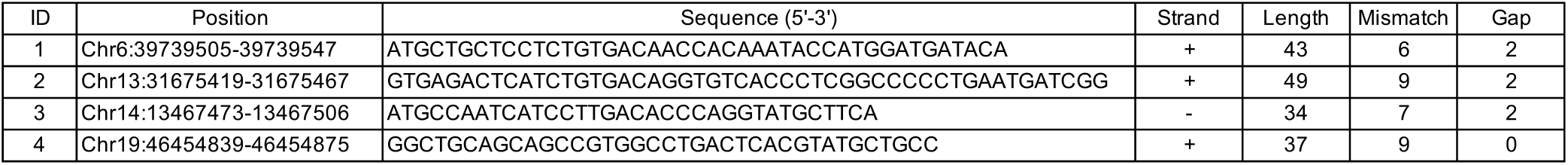
Putative Off-Target Sequences of MECP2 ZFN (32 < Length < 50, Mismatch < 11, Gap < 3)

**Table S5.**
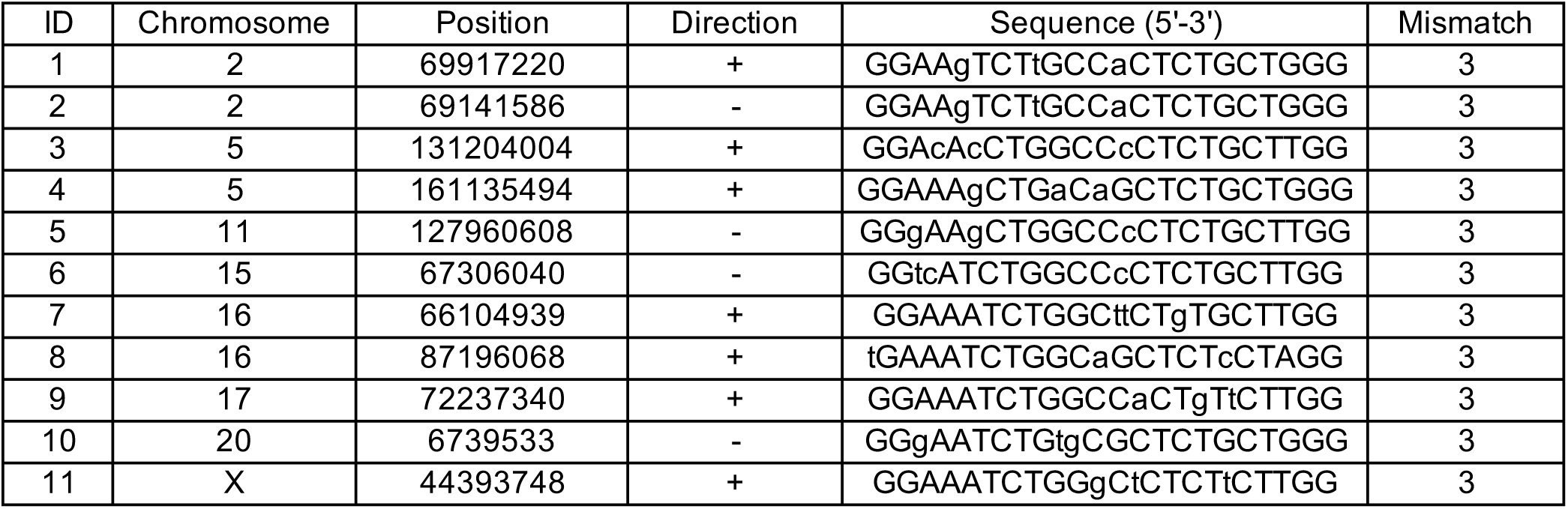
Putative Off-Target Sites of MECP2 CRISPR/Cas9 (Mismatch < 4)

**Table S6.**
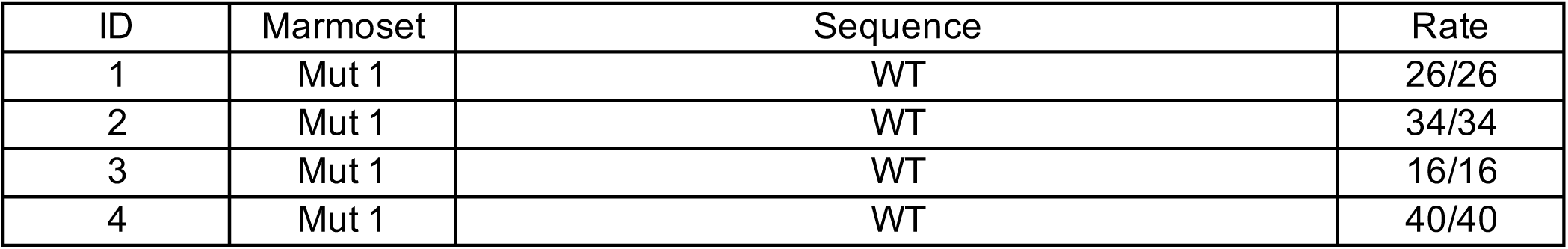
Summary of the Alleles for Putative Off-Target Sites of MECP2 ZFN.

| ID | Marmoset | Sequence | Rate |
| --- | --- | --- | --- |
| 1 | Mut 1 | WT | 26/26 |
| 2 | Mut 1 | WT | 34/34 |
| 3 | Mut 1 | WT | 16/16 |
| 4 | Mut 1 | WT | 40/40 |

**Table S7.**
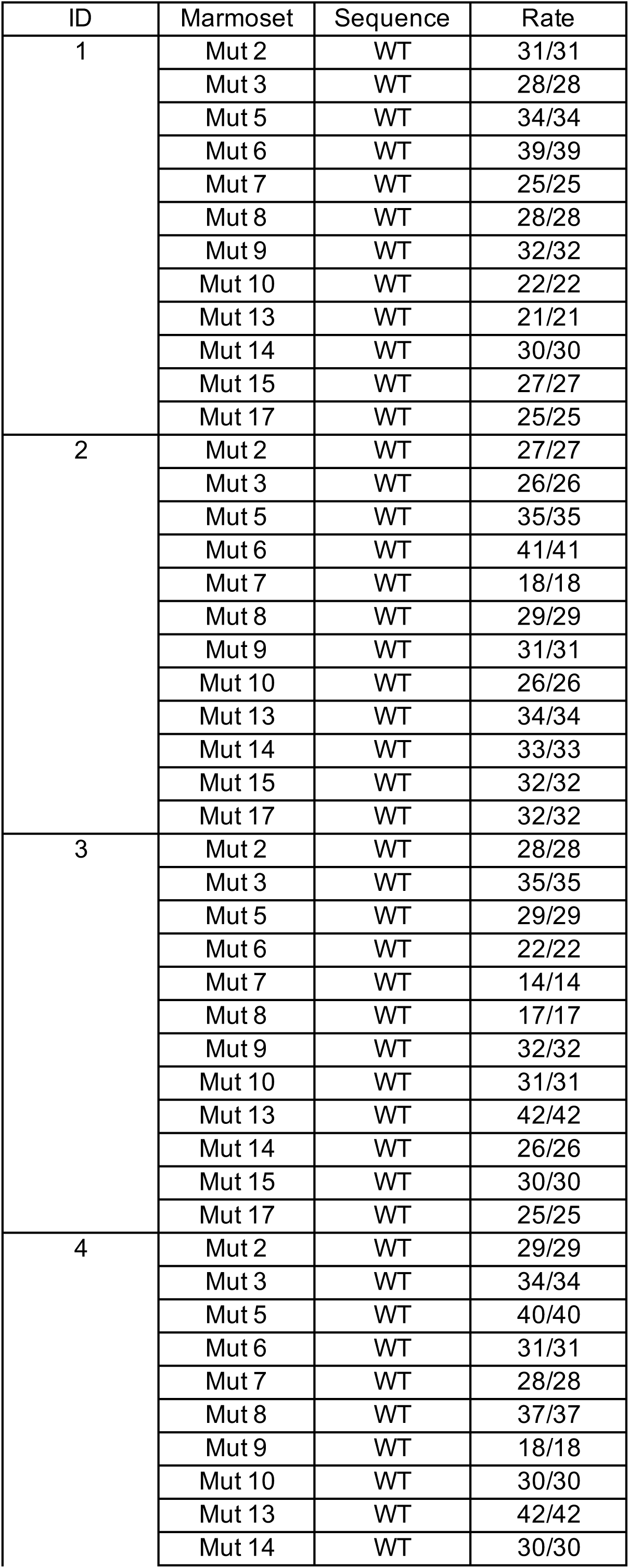
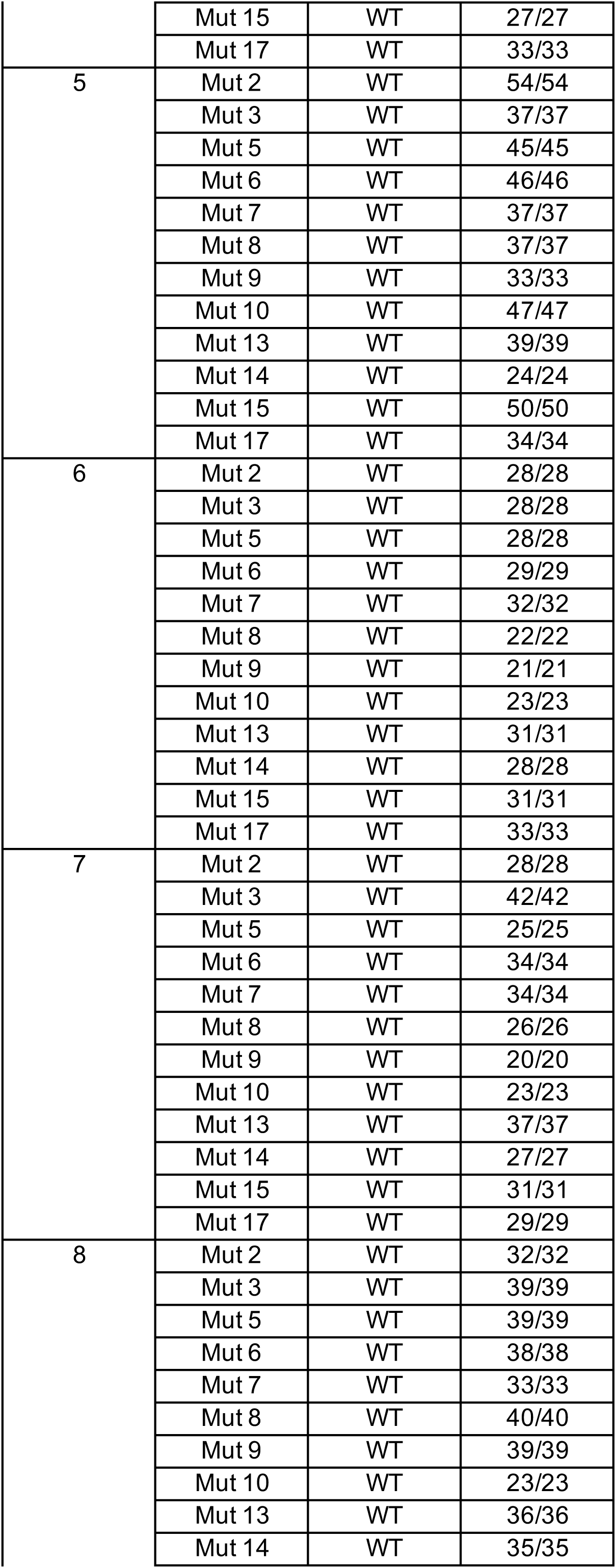
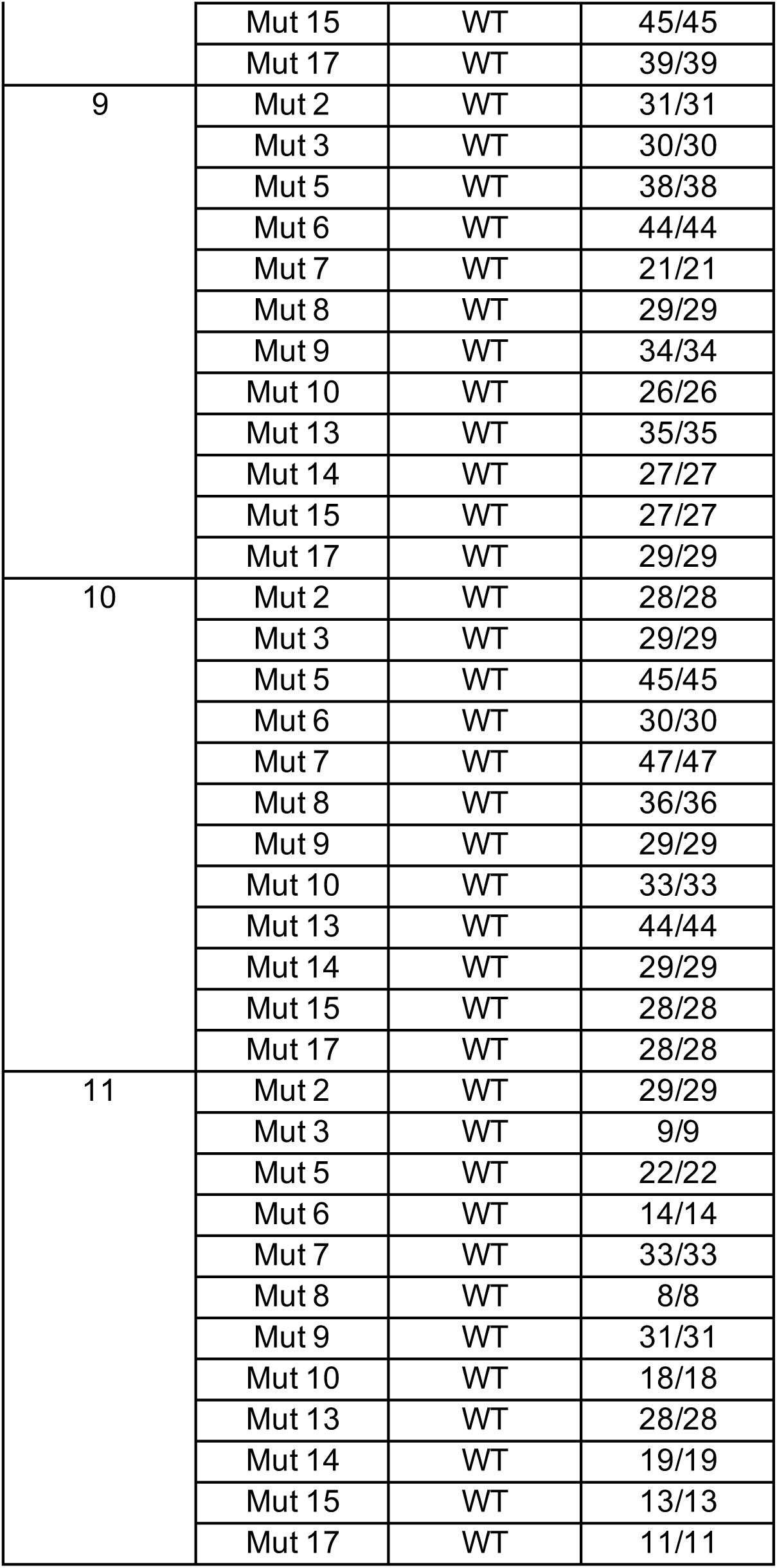
Summary of the Alleles for Putative Off-Target Sites of MECP2 CRISPR/Cas9.

**Table S8.**
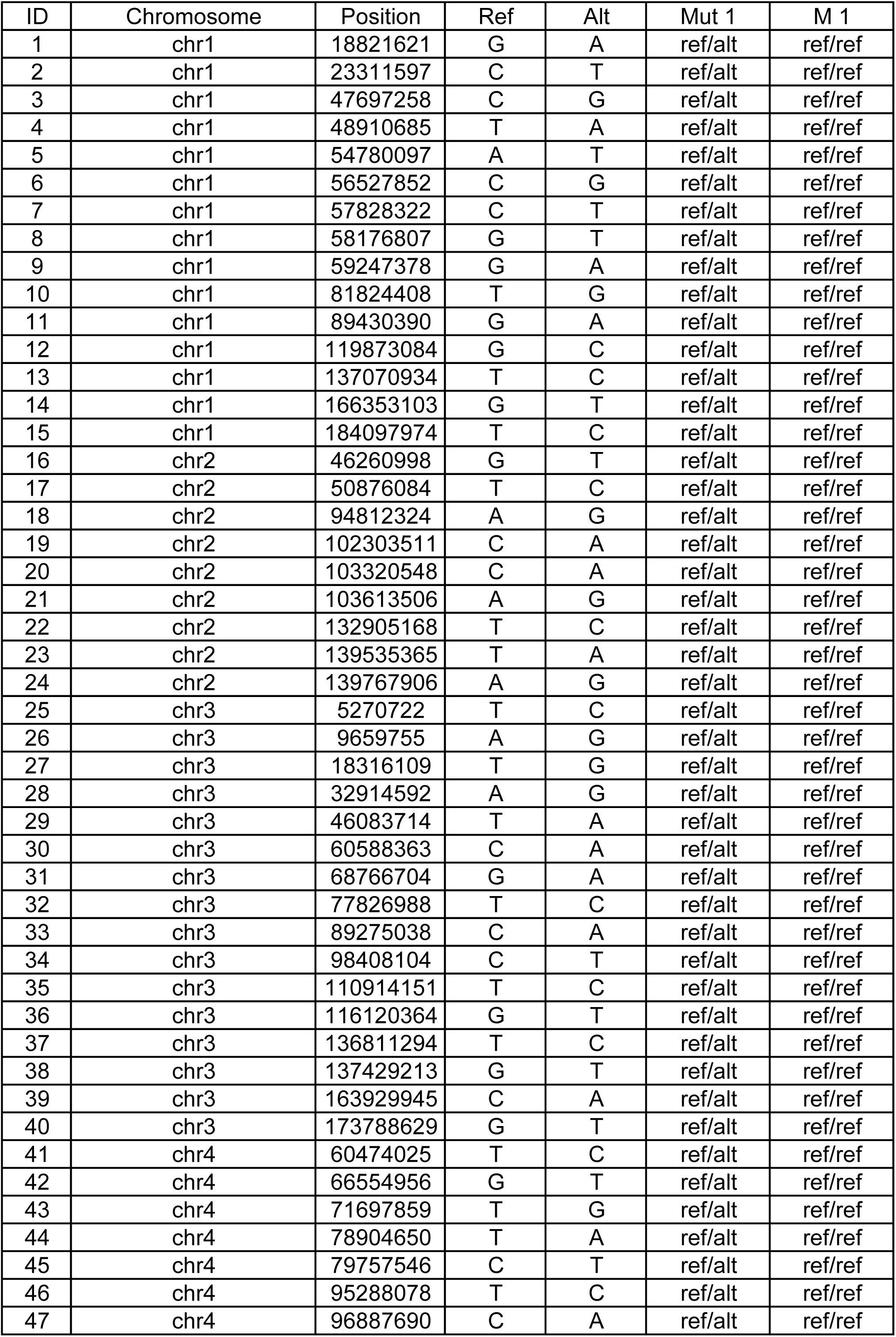
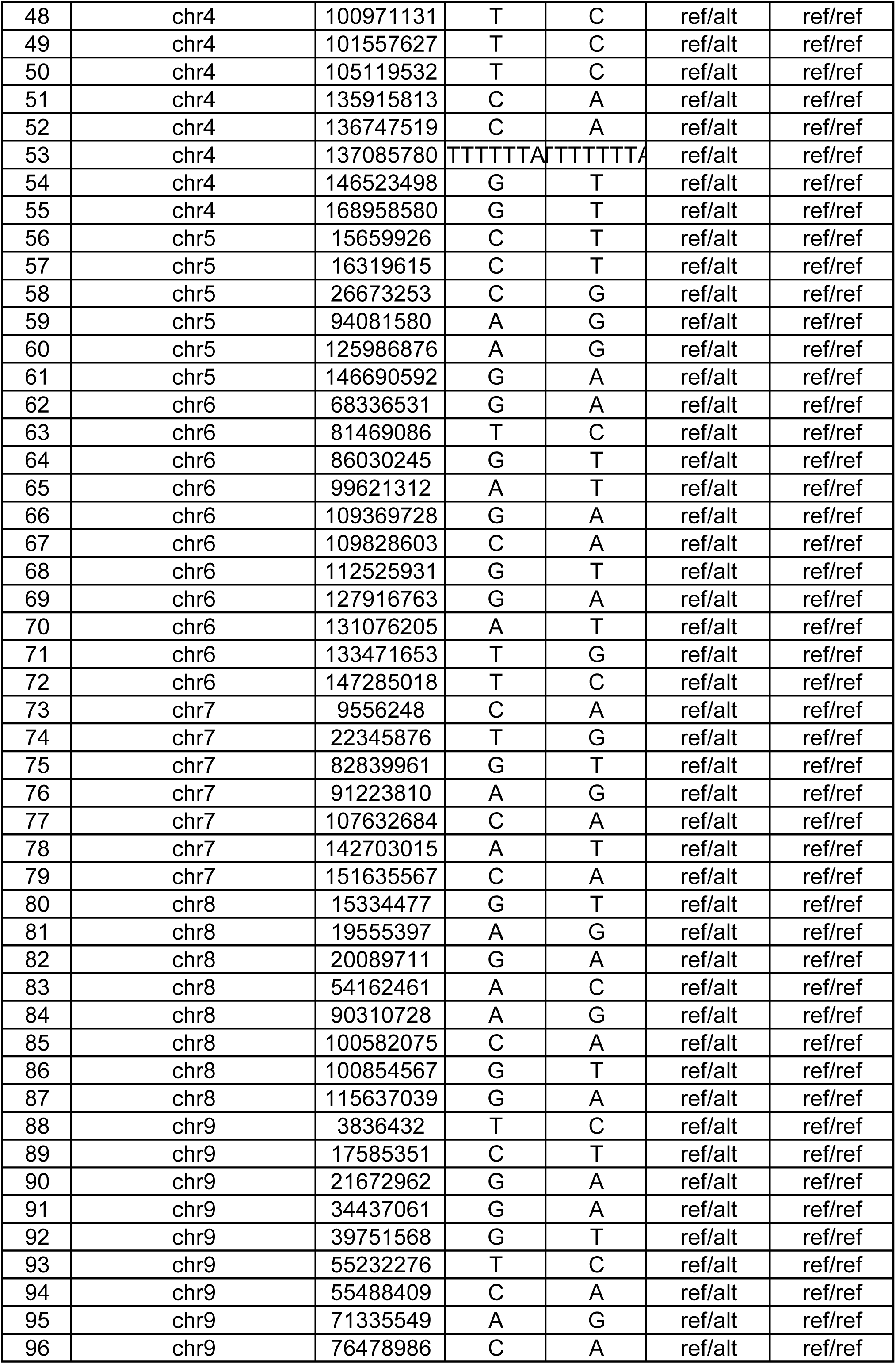
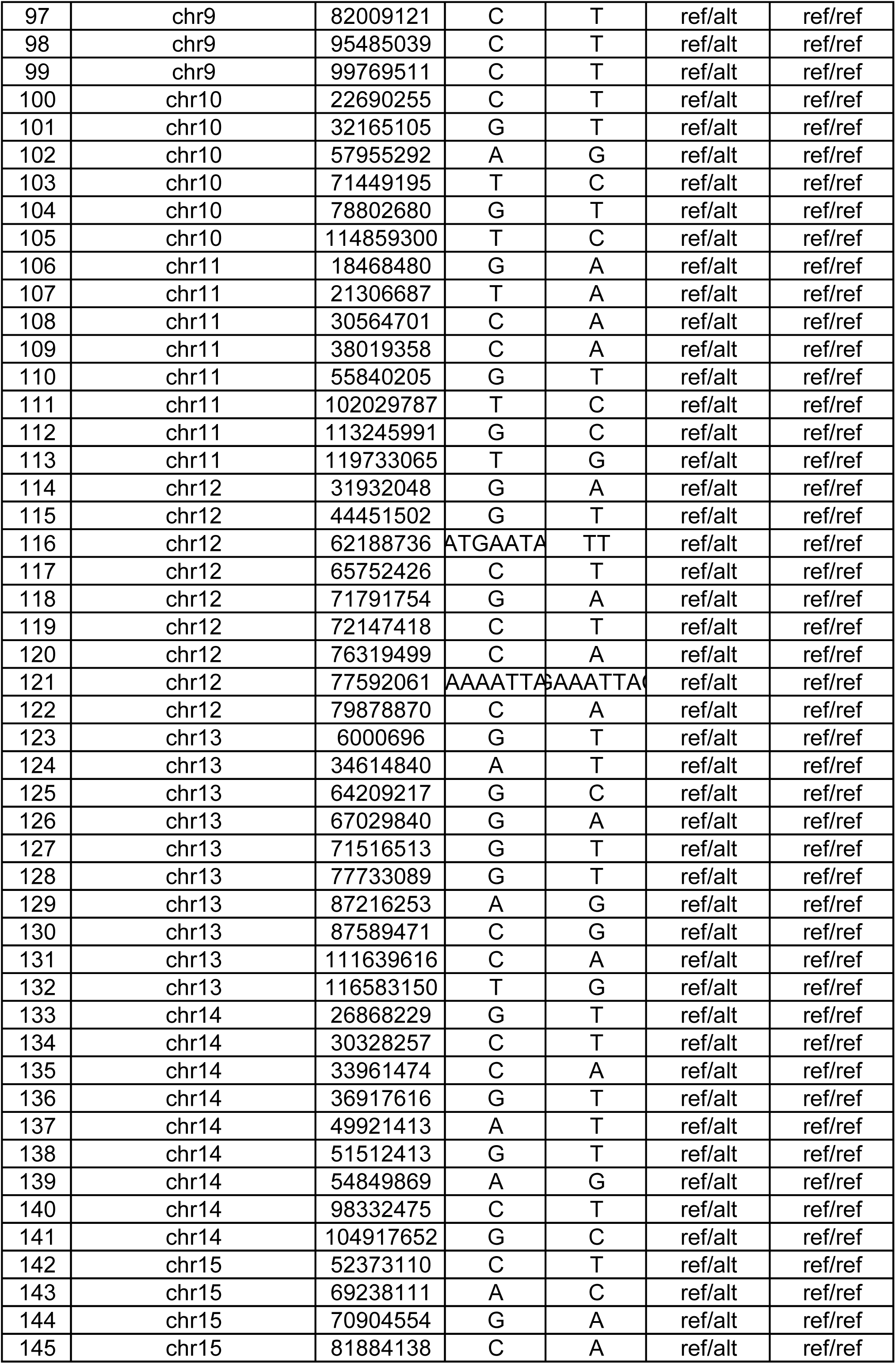
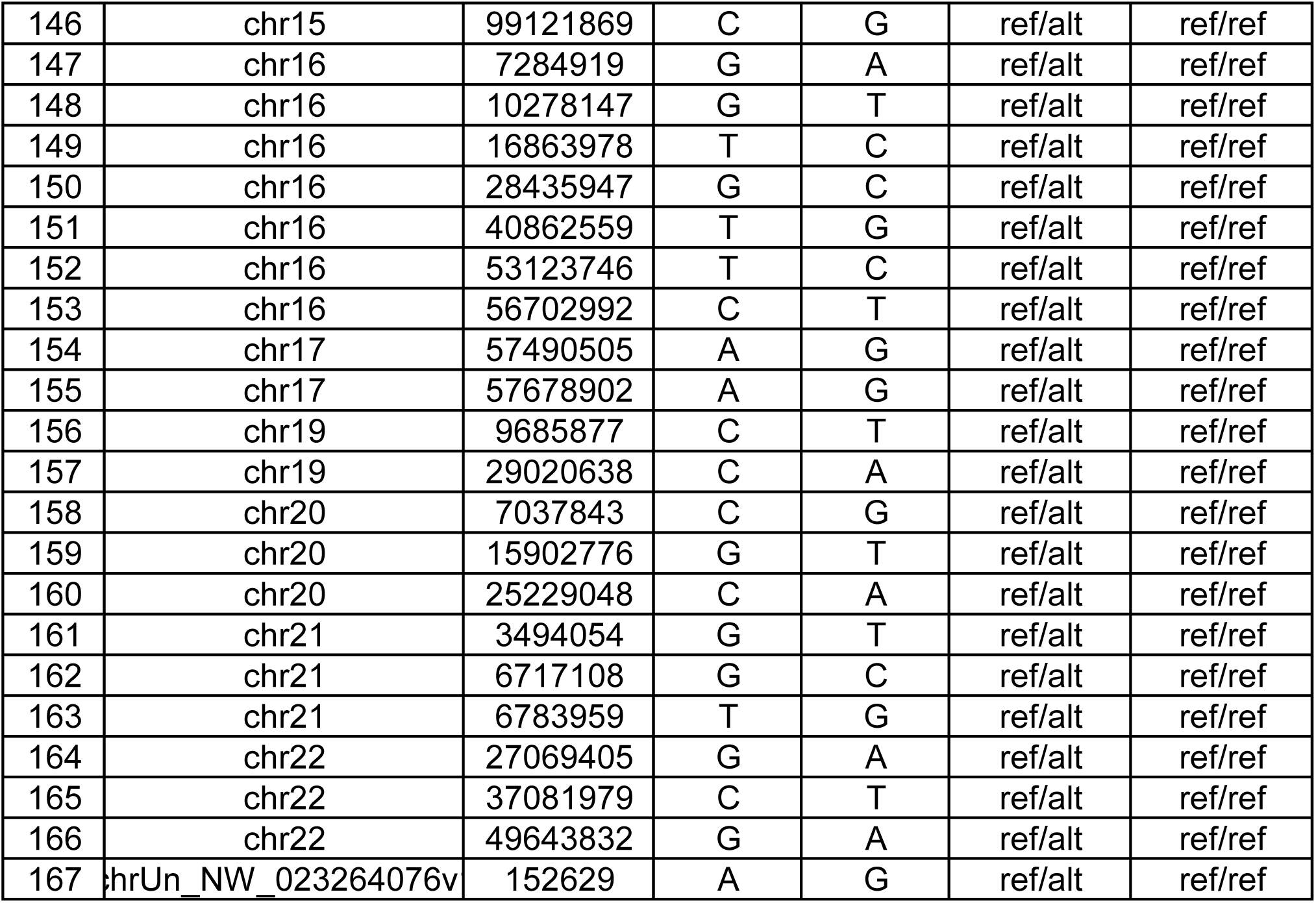
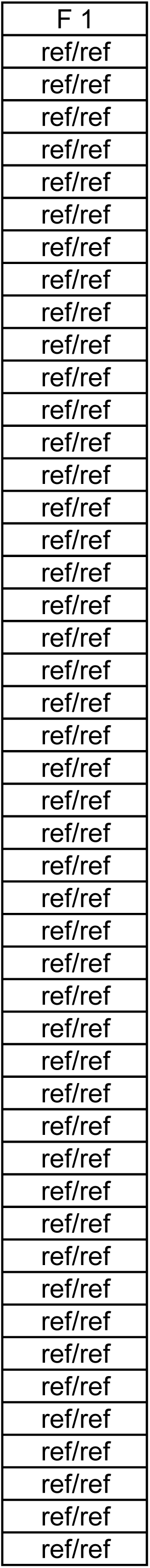
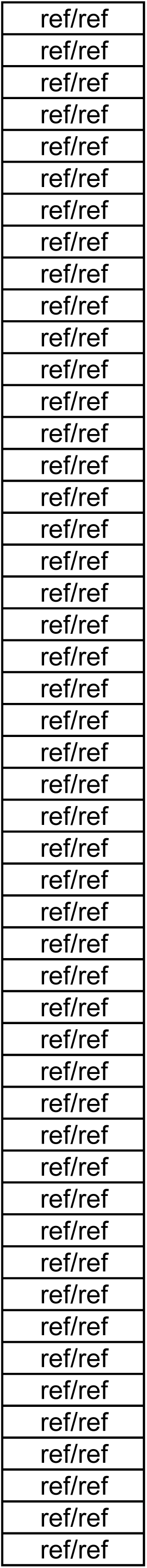
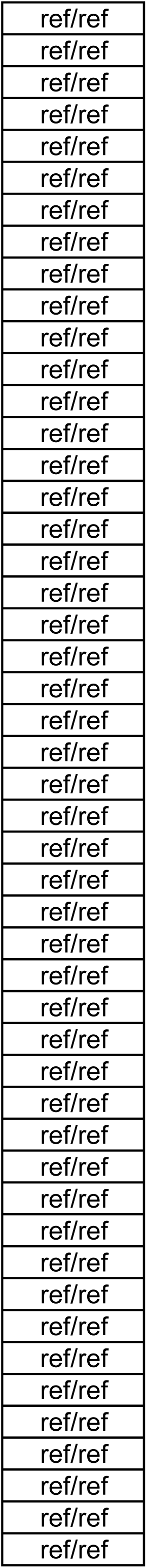
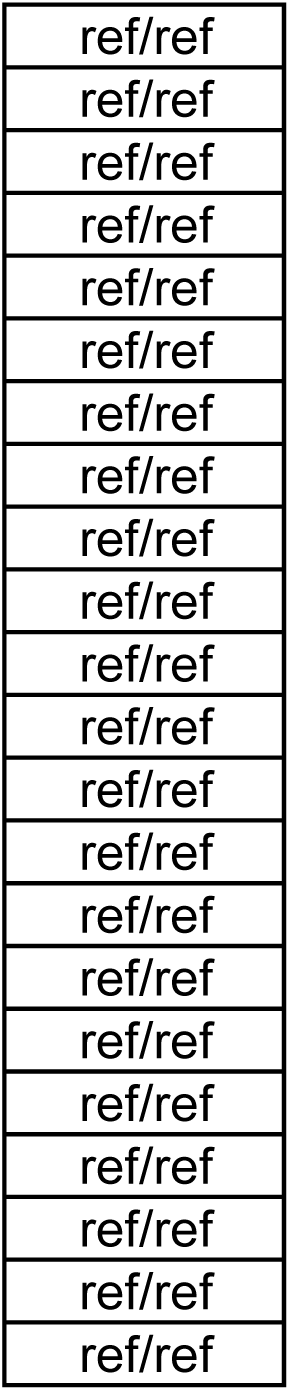
*de novo* variants of Mut 1.

**Table S9.** *de novo* variants of Mut 2.

| ID | Chromosome | Position | Ref | Alt | Mut 2 | M2 | F2 |
| --- | --- | --- | --- | --- | --- | --- | --- |
| 1 | chr2 | 164989104 | T | C | ref/alt | ref/ref | ref/ref |
| 2 | chr3 | 30615359 | C | T | alt/alt | ref/ref | ref/alt |
| 3 | chr4 | 105062083 | G | C | ref/alt | ref/ref | ref/ref |
| 4 | chr6 | 60236382 | G | A | ref/alt | ref/ref | ref/ref |
| 5 | chr6 | 66766930 | C | G | ref/alt | ref/ref | ref/ref |
| 6 | chr6 | 69936908 | A | G | ref/alt | ref/ref | ref/ref |
| 7 | chr7 | 123898746 | A | G | ref/alt | ref/ref | ref/ref |
| 8 | chr12 | 88914088 | C | A | ref/alt | ref/ref | ref/ref |
| 9 | chr13 | 14264596 | A | G | ref/alt | ref/ref | ref/ref |
| 10 | chr13 | 54347239 | C | A | ref/alt | ref/ref | ref/ref |
| 11 | chr15 | 15142919 | C | T | ref/alt | ref/ref | ref/ref |
| 12 | chr15 | 49628139 | C | A | ref/alt | ref/ref | ref/ref |
| 13 | chr16 | 24549225 | T | A | ref/alt | ref/ref | ref/ref |
| 14 | chr16 | 73745154 | A | C | ref/alt | ref/ref | ref/ref |
| 15 | chr16 | 91652968 | A | T | ref/alt | ref/ref | ref/ref |
| 16 | chr17 | 55938431 | GCG | GCA | ref/alt | ref/ref | ref/ref |
| 17 | chr20 | 16453582 | G | C | ref/alt | ref/ref | ref/ref |
| 18 | chrUn_NW_023264148v1 | 124987 | T | A | ref/alt | ref/ref | ref/ref |

**Table S10.** *de novo* variants of Mut 13.

| ID | Chromosome | Position | Ref | Alt | Mut 13 | M3 | F3 |
| --- | --- | --- | --- | --- | --- | --- | --- |
| 1 | chr1 | 145358199 | G | T | ref/alt | ref/ref | ref/ref |
| 2 | chr3 | 21400216 | C | G | ref/alt | ref/ref | ref/ref |
| 3 | chr3 | 62803807 | G | A | ref/alt | ref/ref | ref/ref |
| 4 | chr4 | 58235173 | A | G | ref/alt | ref/ref | ref/ref |
| 5 | chr5 | 56978499 | G | A | ref/alt | ref/ref | ref/ref |
| 6 | chr5 | 84469641 | C | G | ref/alt | ref/ref | ref/ref |
| 7 | chr7 | 37852778 | C | T | ref/alt | ref/ref | ref/ref |
| 8 | chr9 | 18077744 | A | G | ref/alt | ref/ref | ref/ref |
| 9 | chr10 | 64830070 | C | G | ref/alt | ref/ref | ref/ref |
| 10 | chr11 | 8892488 | G | A | ref/alt | ref/ref | ref/ref |
| 11 | chr12 | 34106114 | T | A | ref/alt | ref/ref | ref/ref |
| 12 | chr13 | 53454694 | TC | CT | ref/alt | ref/ref | ref/ref |
| 13 | chr14 | 67379168 | C | A | ref/alt | ref/ref | ref/ref |
| 14 | chr16 | 53929970 | C | T | ref/alt | ref/ref | ref/ref |
| 15 | chr16 | 73647350 | A | T | ref/alt | ref/ref | ref/ref |
| 16 | chr17 | 21040857 | C | T | ref/alt | ref/ref | ref/ref |
| 17 | chr21 | 6453150 | G | T | ref/alt | ref/ref | ref/ref |
| 18 | chrUn_NW_023264968v1 | 541431 | C | T | alt/alt | ref/ref | alt/alt |

**Table S11.** *de novo* variants of Mut 14.

| ID | Chromosome | Position | Ref | Alt | Mut 14 | M4 | F3 |
| --- | --- | --- | --- | --- | --- | --- | --- |
| 1 | chr2 | 66132982 | C | T | ref/alt | ref/ref | ref/ref |
| 2 | chr2 | 77515937 | C | A | ref/alt | ref/ref | ref/ref |
| 3 | chr3 | 131063184 | C | G | ref/alt | ref/ref | ref/ref |
| 4 | chr4 | 118818150 | G | A | ref/alt | ref/ref | ref/ref |
| 5 | chr7 | 7899050 | A | G | ref/alt | ref/ref | ref/ref |
| 6 | chr8 | 58961434 | T | C | ref/alt | ref/ref | ref/ref |
| 7 | chr11 | 3240912 | C | G | ref/alt | ref/ref | ref/ref |
| 8 | chr11 | 13620954 | C | T | ref/alt | ref/ref | ref/ref |
| 9 | chr15 | 11969624 | T | C | ref/alt | ref/ref | ref/ref |
| 10 | chr16 | 39170225 | C | T | ref/alt | ref/ref | ref/ref |
| 11 | chr17 | 41578117 | C | T | ref/alt | ref/ref | ref/ref |
| 12 | chr20 | 6427629 | C | T | ref/alt | ref/ref | ref/ref |

**Table S12.** Putative Off-Target Sequences of MECP2 ZFN (Length < 100, Mismatch < 11, Gap < 3)

| ID | Position | Strand | Length | Mismatch | Gap |
| --- | --- | --- | --- | --- | --- |
| 1 | chr1:173,501,007-173,501,059 | + | 53 | 8 | 1 |
| 2 | chr1:122,023,555-122,023,637 | + | 83 | 10 | 1 |
| 3 | chr2:80,779,024-80,779,118 | + | 95 | 8 | 2 |
| 4 | chr2:80,988,196-80,988,290 | - | 95 | 10 | 1 |
| 5 | chr6:139,155,333-139,155,387 | + | 55 | 9 | 2 |
| 6 | chr6:35,181,617-35,181,701 | + | 85 | 8 | 0 |
| 7 | chr6:39,739,505-39,739,547 | - | 43 | 6 | 2 |
| 8 | chr7:126,451,583-126,451,666 | - | 84 | 8 | 1 |
| 9 | chr7:71,051,442-71,051,527 | - | 86 | 8 | 1 |
| 10 | chr8:88,472,735-88,472,823 | + | 89 | 8 | 1 |
| 11 | chr9:49,902,830-49,902,901 | + | 72 | 10 | 1 |
| 12 | chr9:4,976,472-4,976,533 | - | 62 | 10 | 1 |
| 13 | chr10:107,810,817-107,810,876 | - | 60 | 9 | 1 |
| 14 | chr10:85,645,791-85,645,877 | - | 87 | 7 | 1 |
| 15 | chr10:14,812,145-14,812,214 | - | 70 | 9 | 1 |
| 16 | chr13:31,675,419-31,675,467 | + | 49 | 9 | 2 |
| 17 | chr14:13,467,473-13,467,506 | + | 34 | 7 | 2 |
| 18 | chr15:76,329,234-76,329,298 | + | 65 | 8 | 1 |
| 19 | chr15:71,475,834-71,475,889 | - | 56 | 9 | 1 |
| 20 | chr16:90,862,063-90,862,161 | - | 99 | 9 | 2 |
| 21 | chr19:46,454,839-46,454,875 | + | 37 | 9 | 0 |
| 22 | chr19:33,859,473-33,859,527 | - | 55 | 9 | 2 |
| 23 | chrX:474,425-474,488 | - | 64 | 9 | 2 |

**Table S13.**
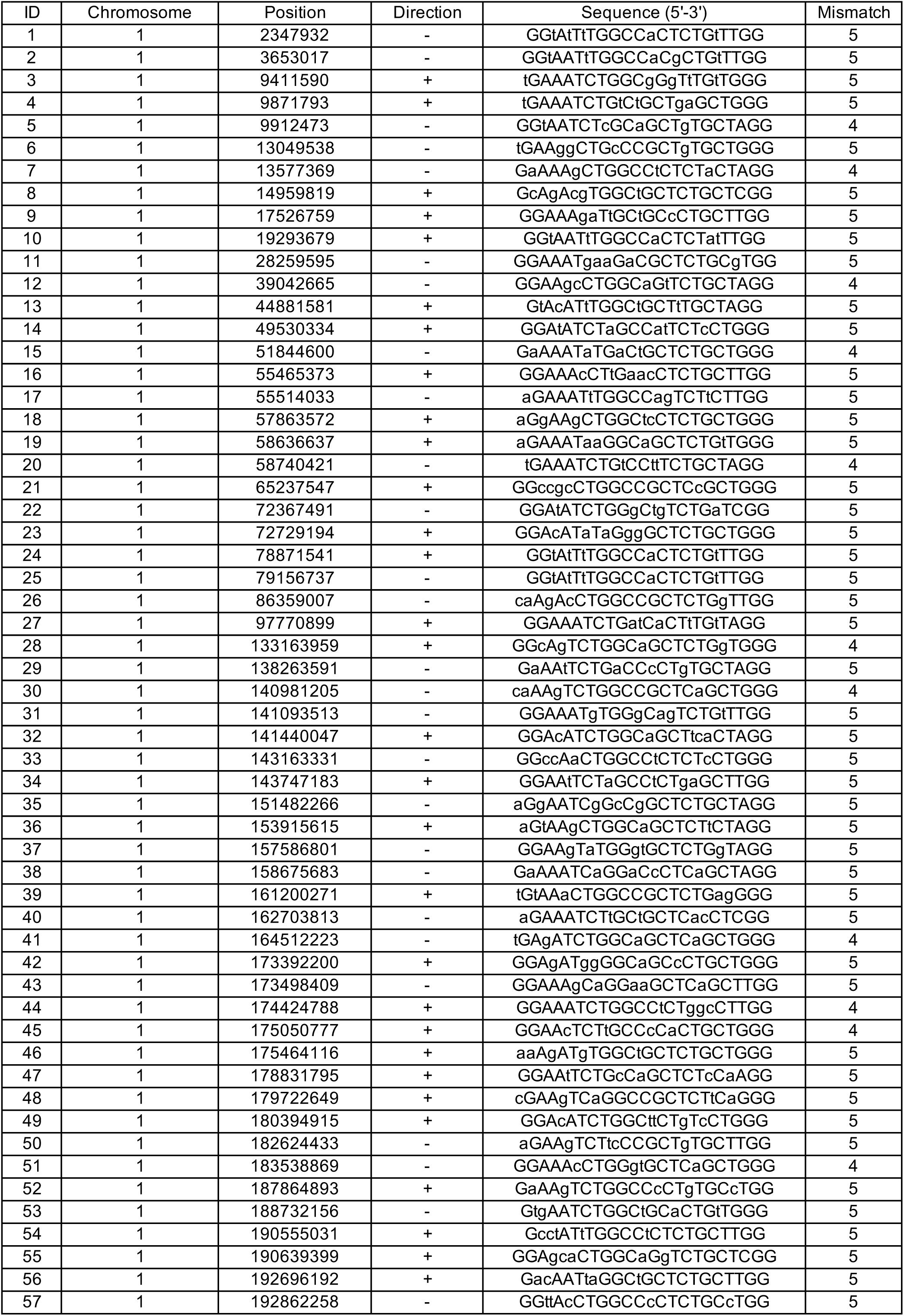
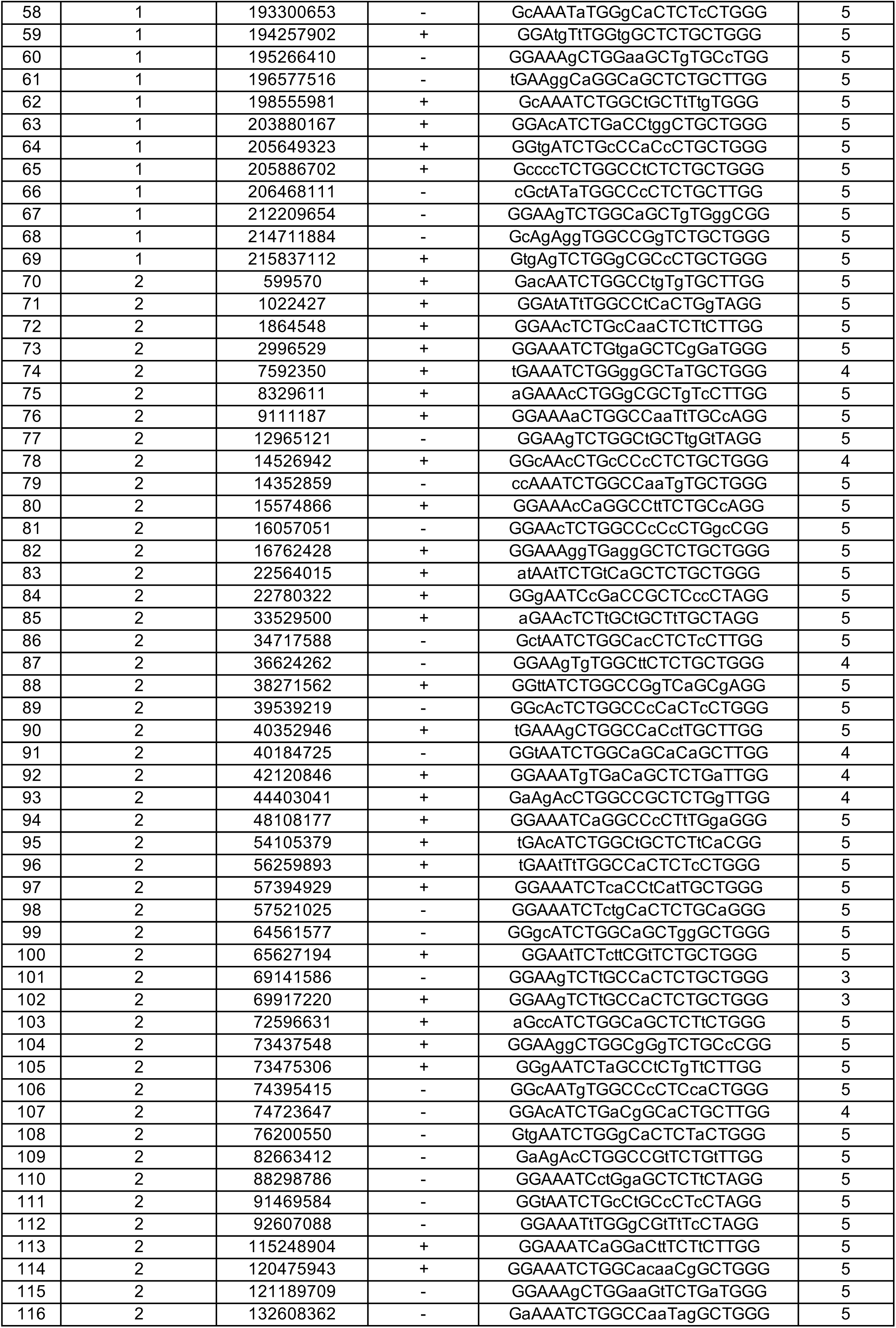
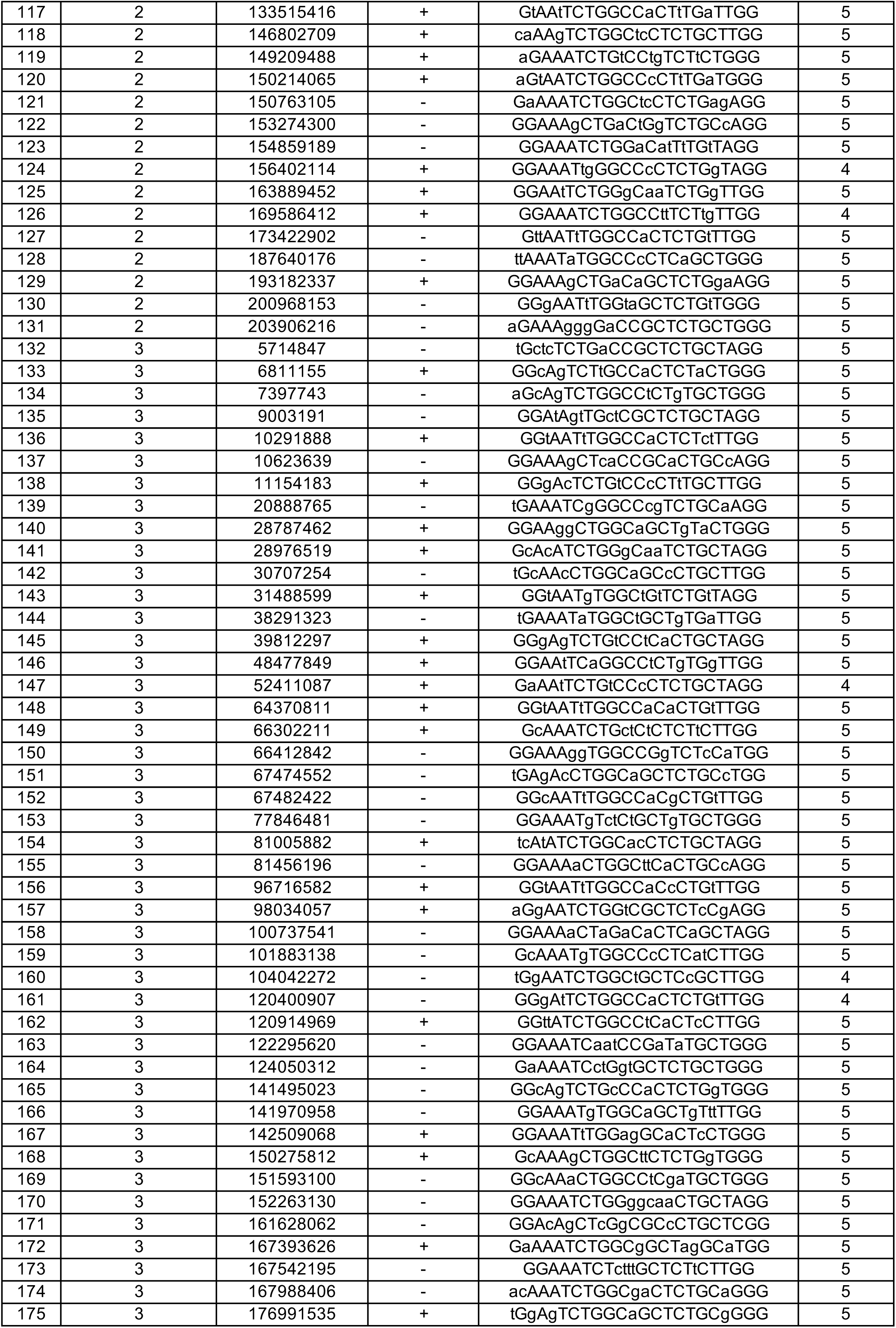
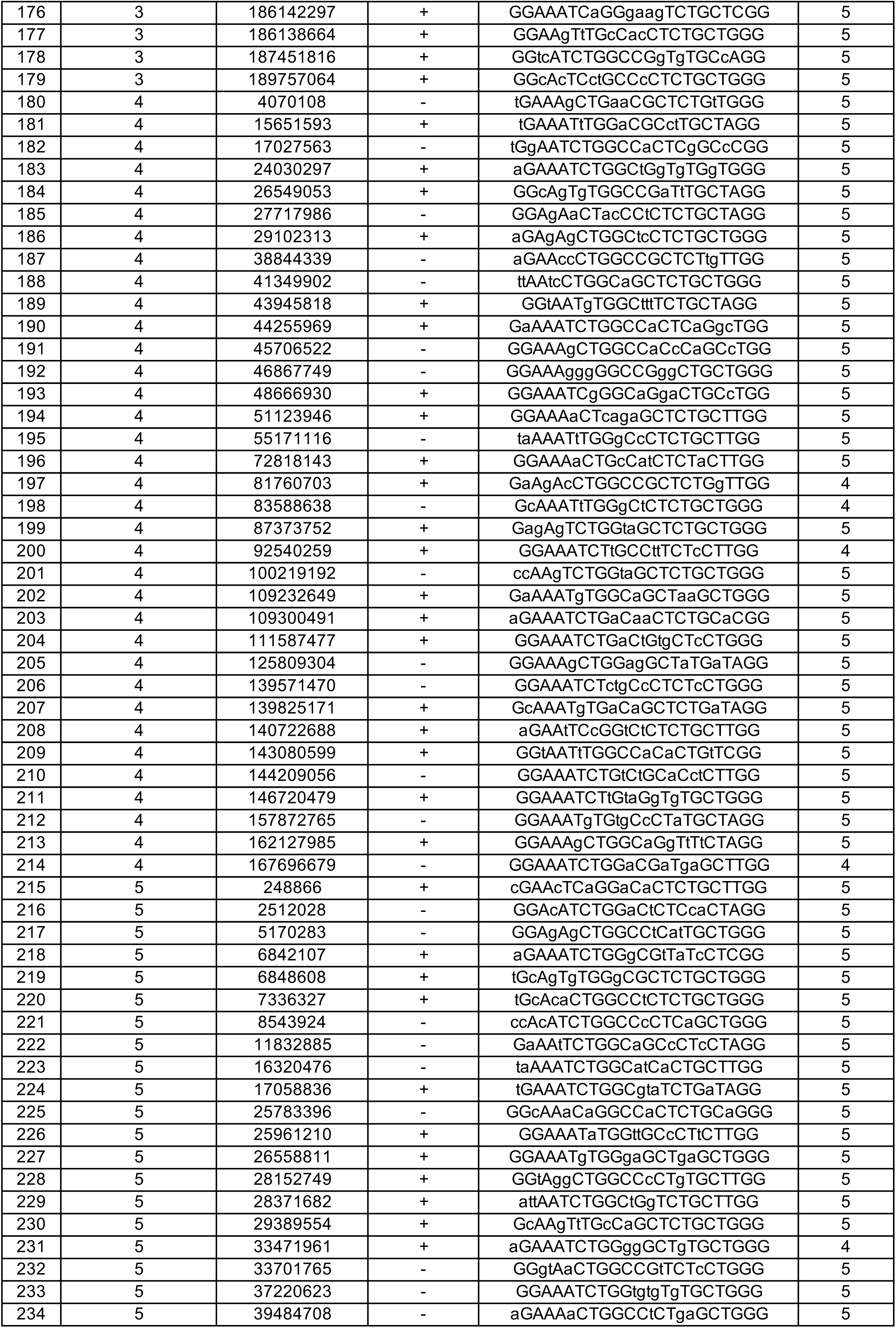
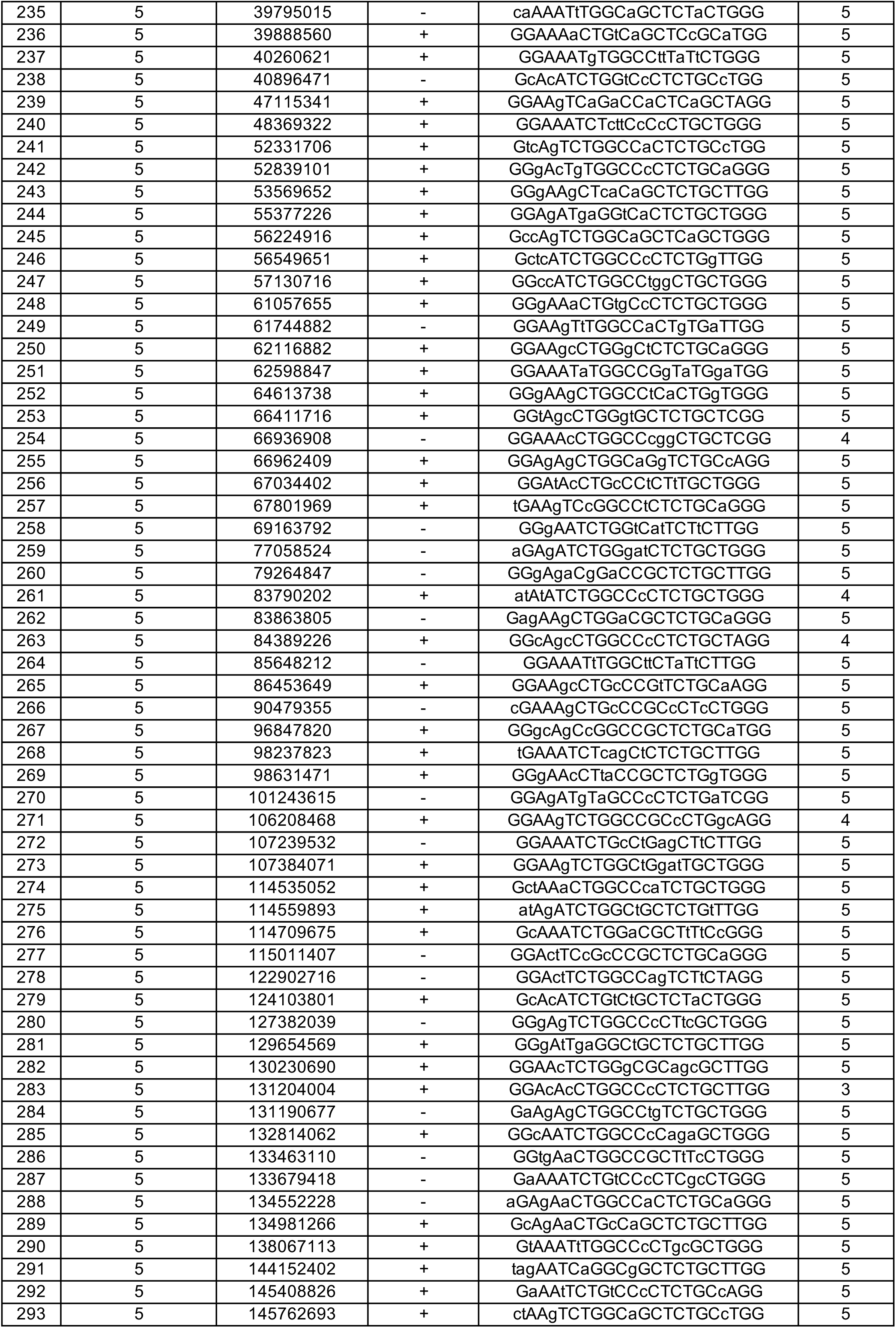
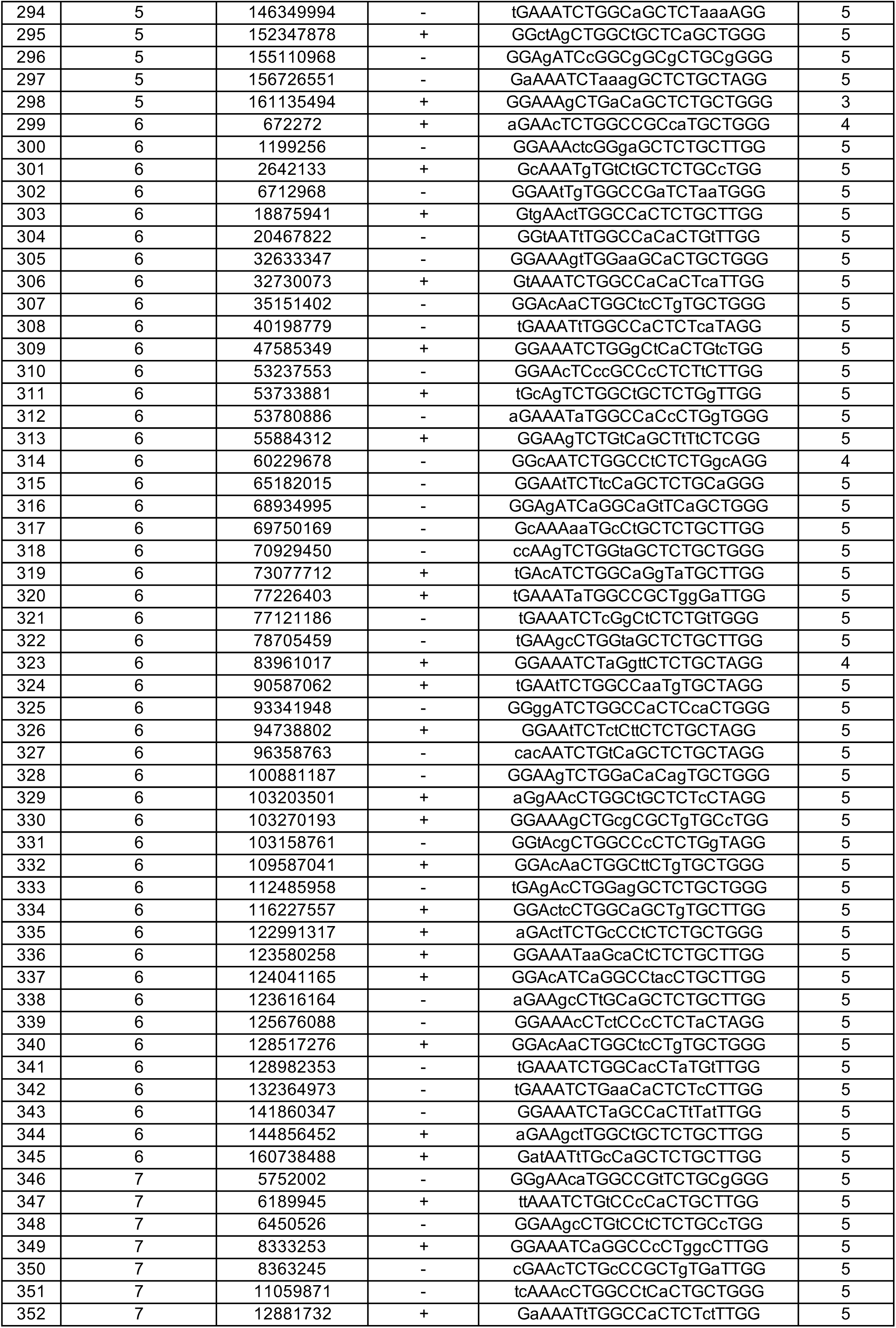
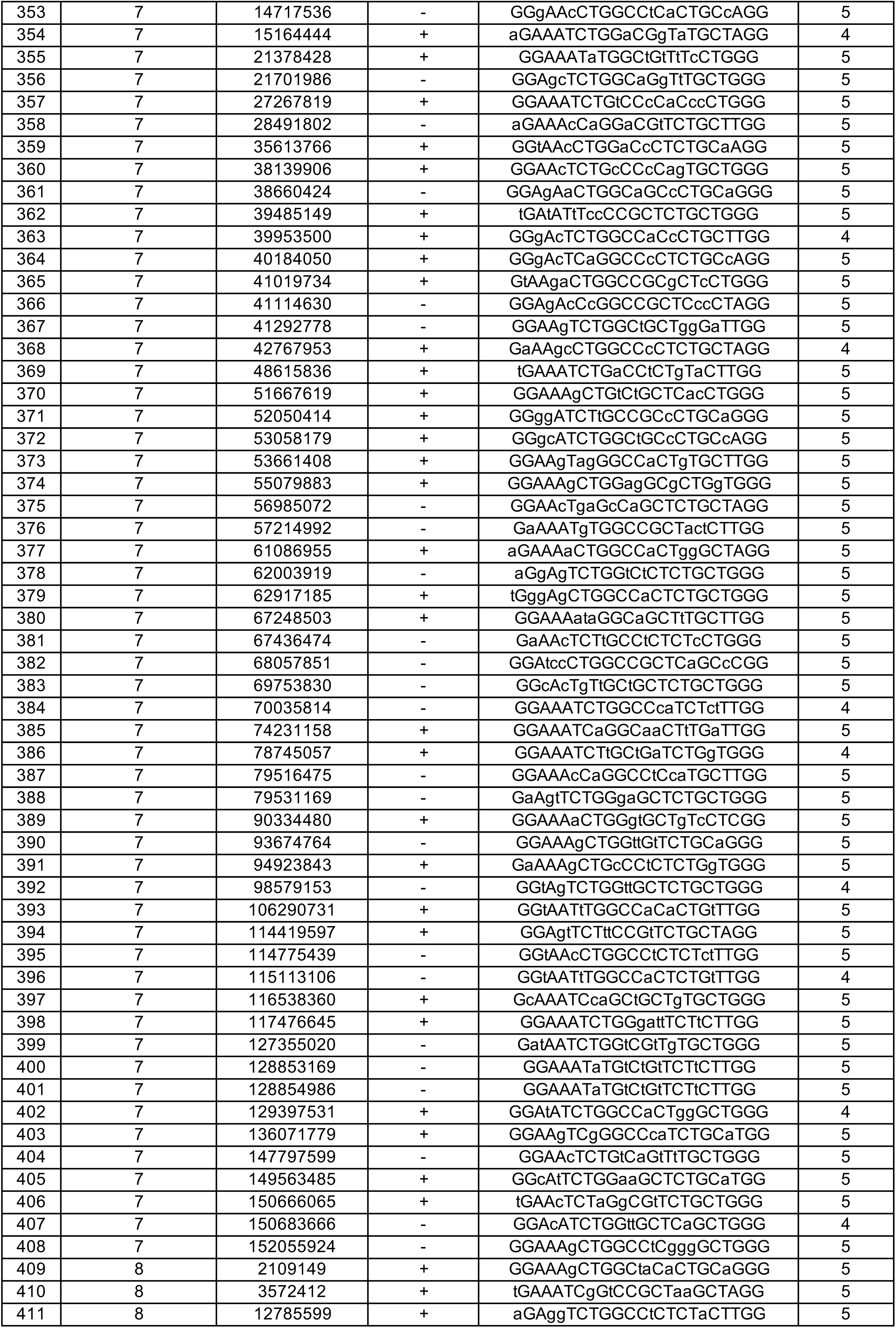
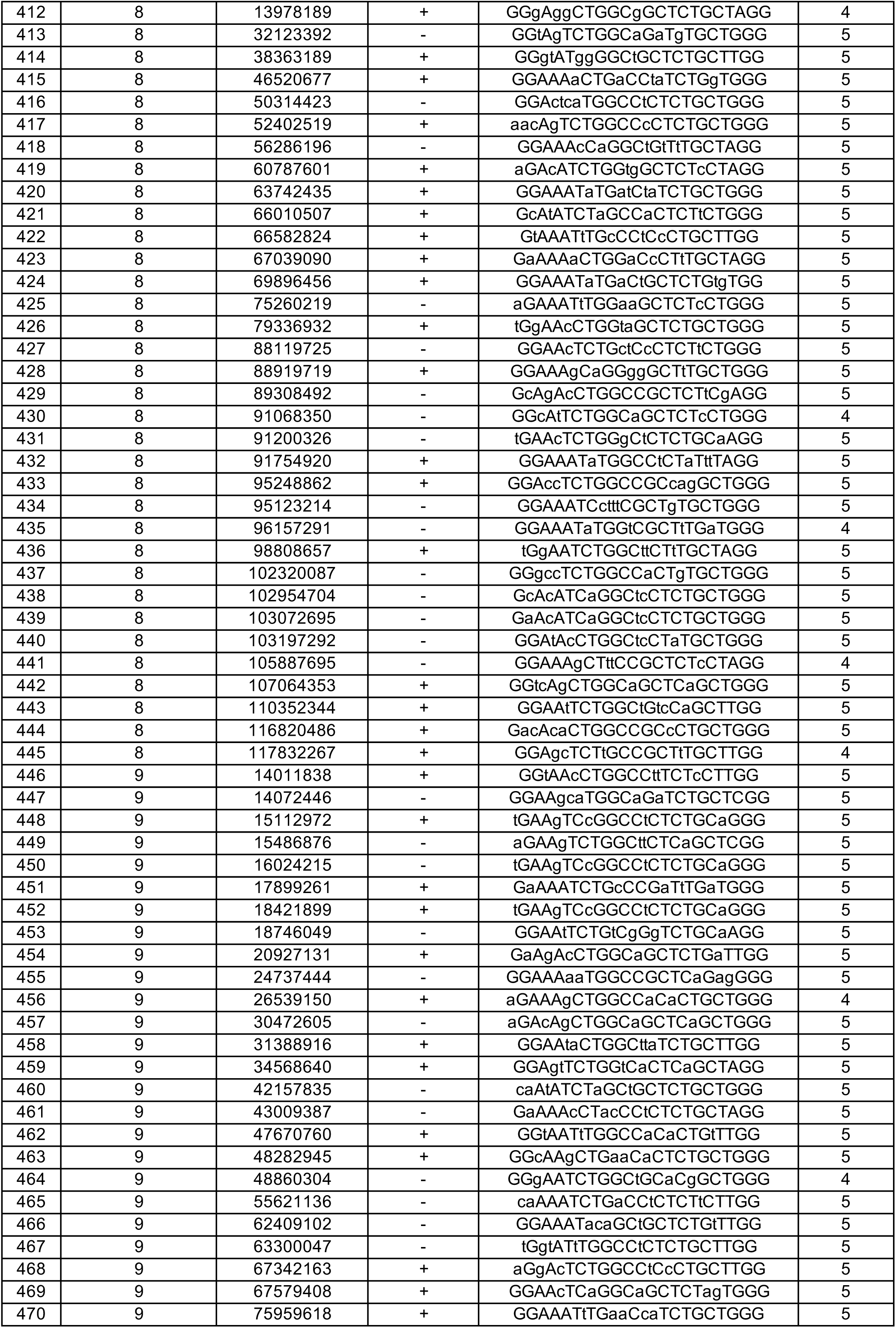
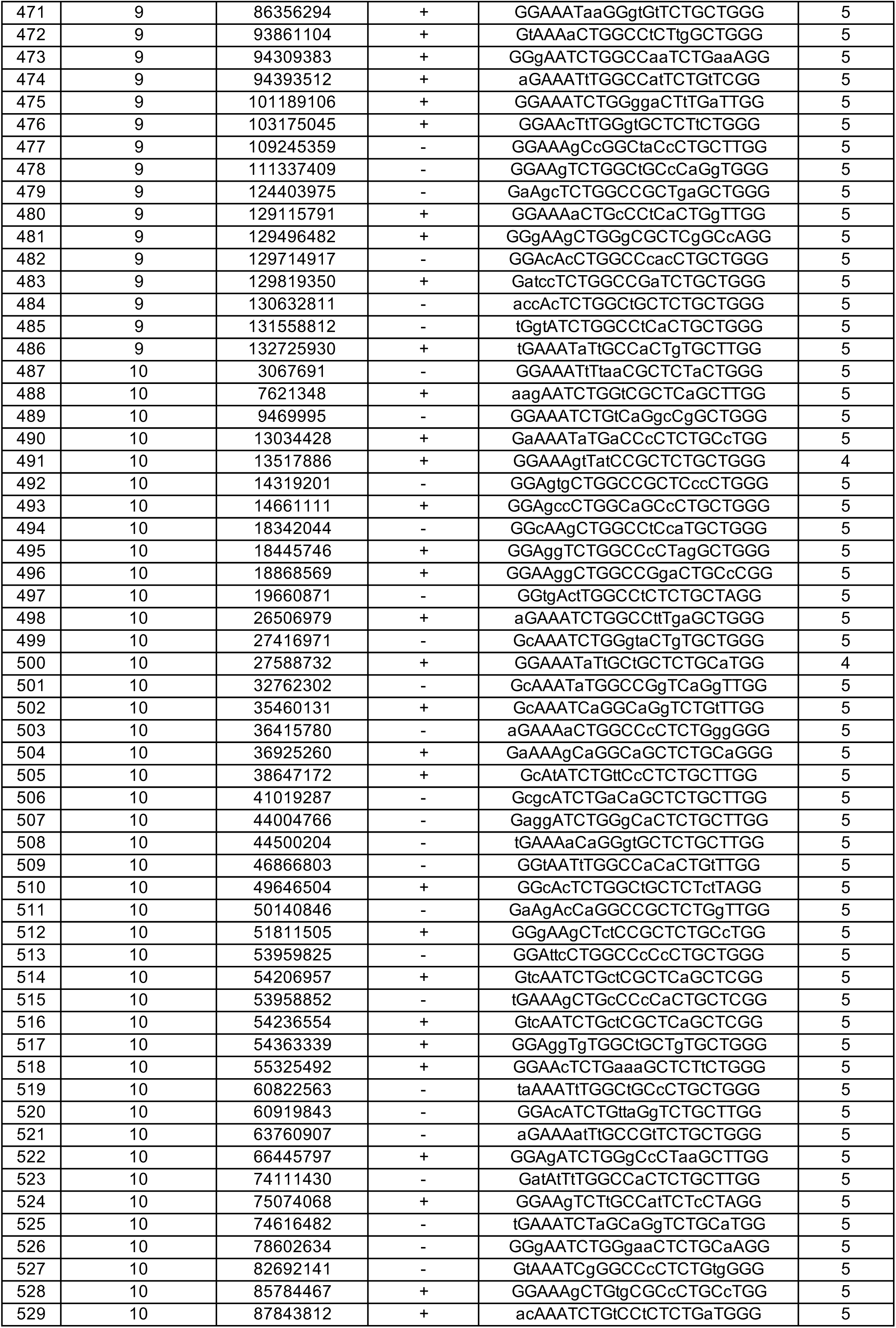
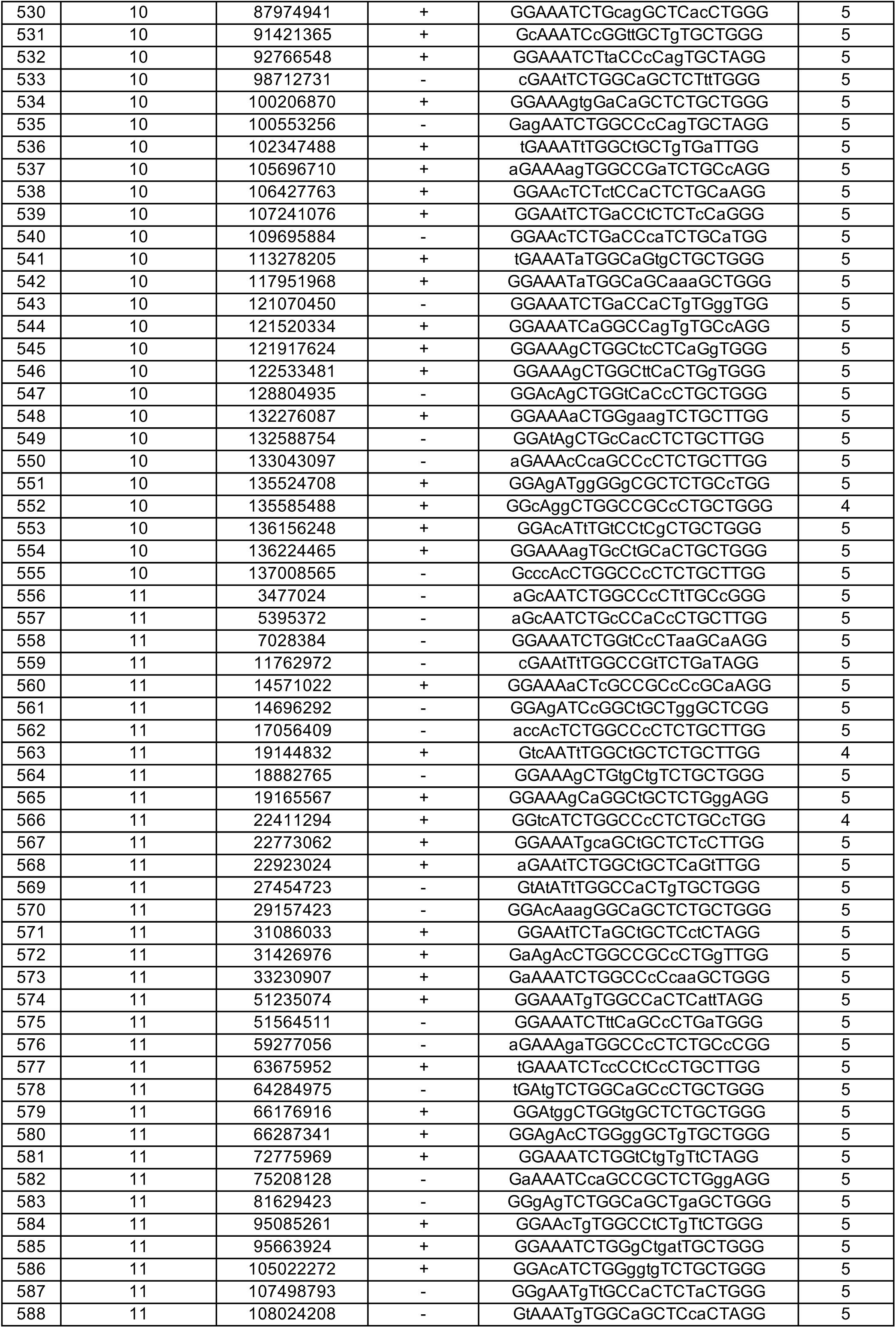
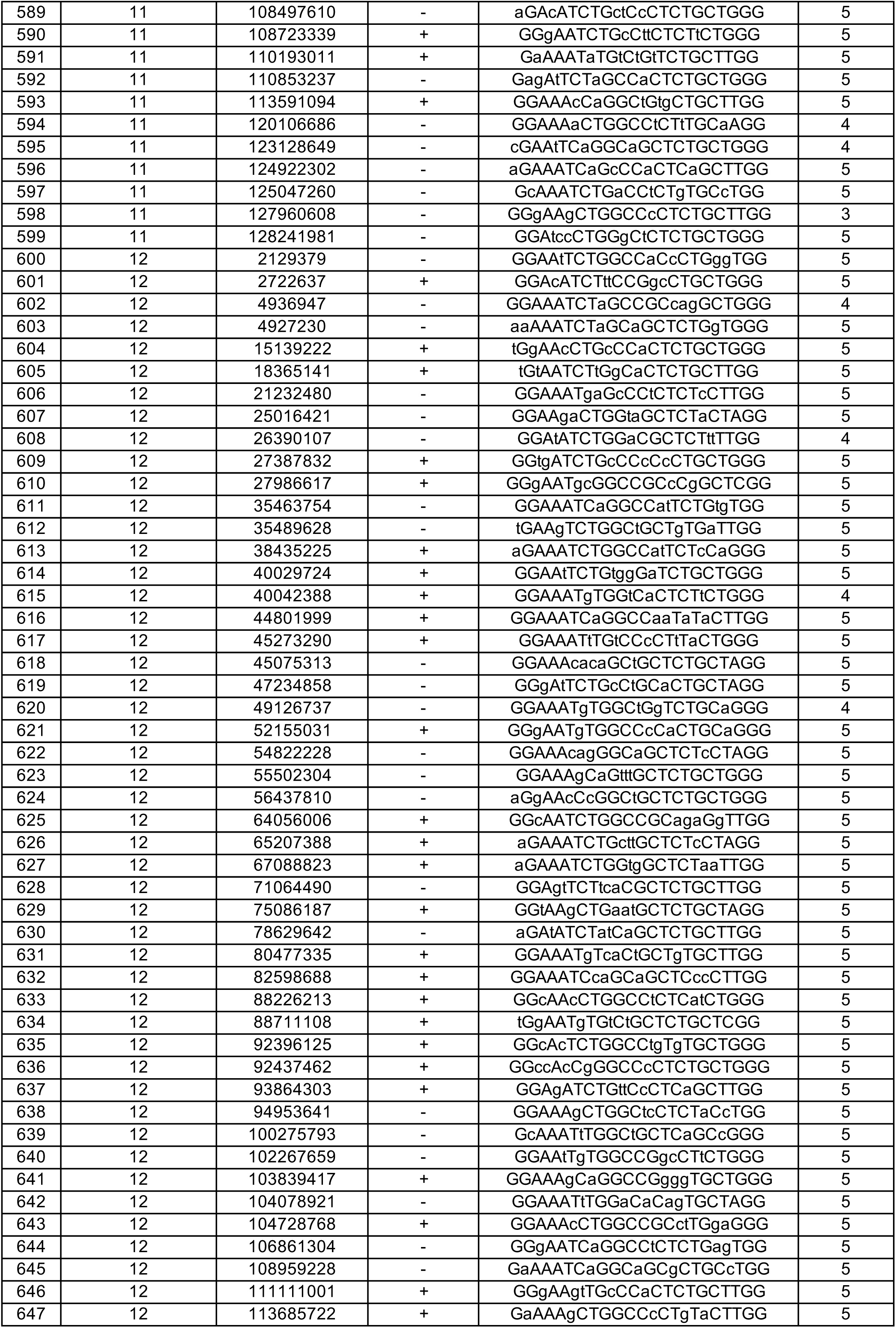
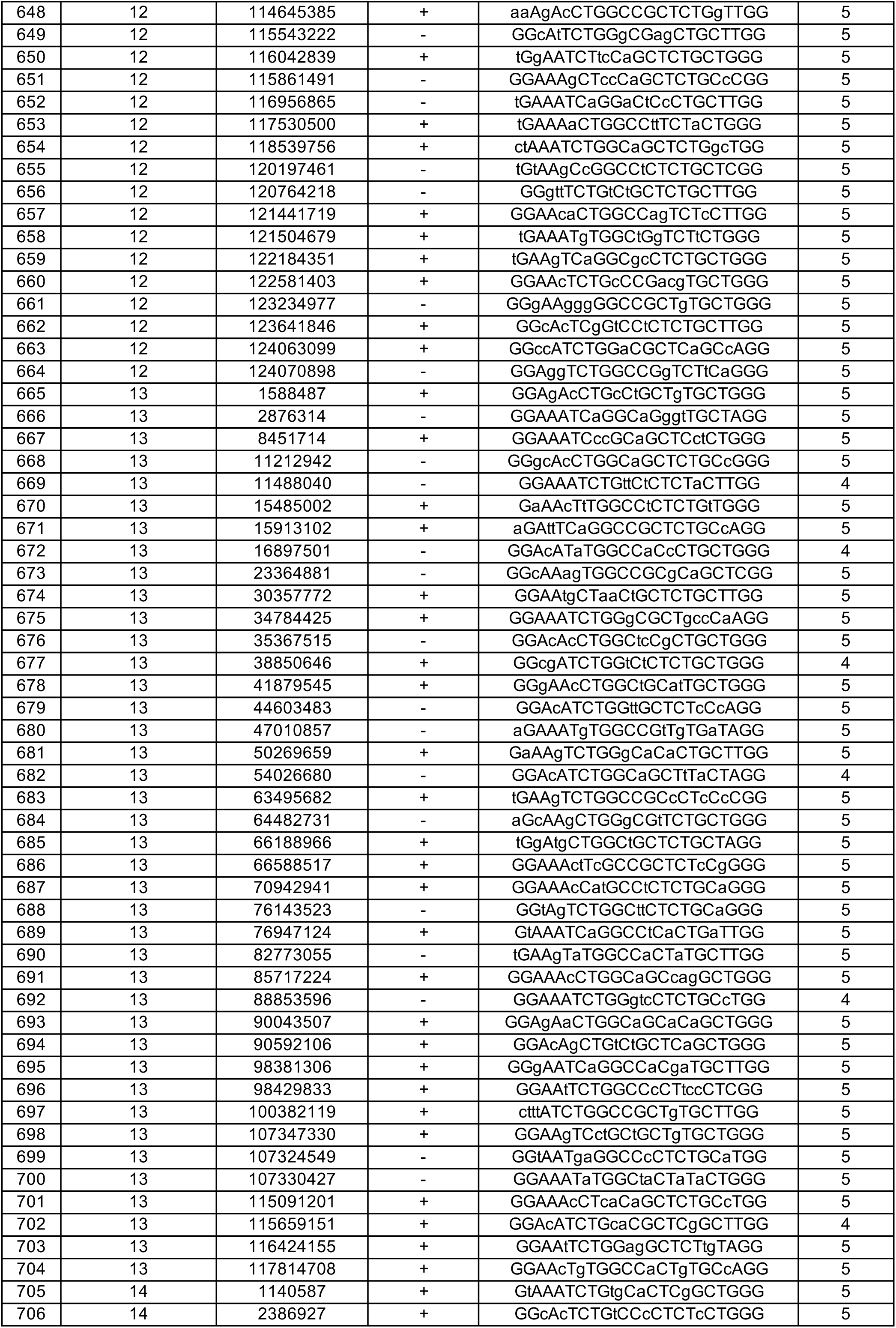
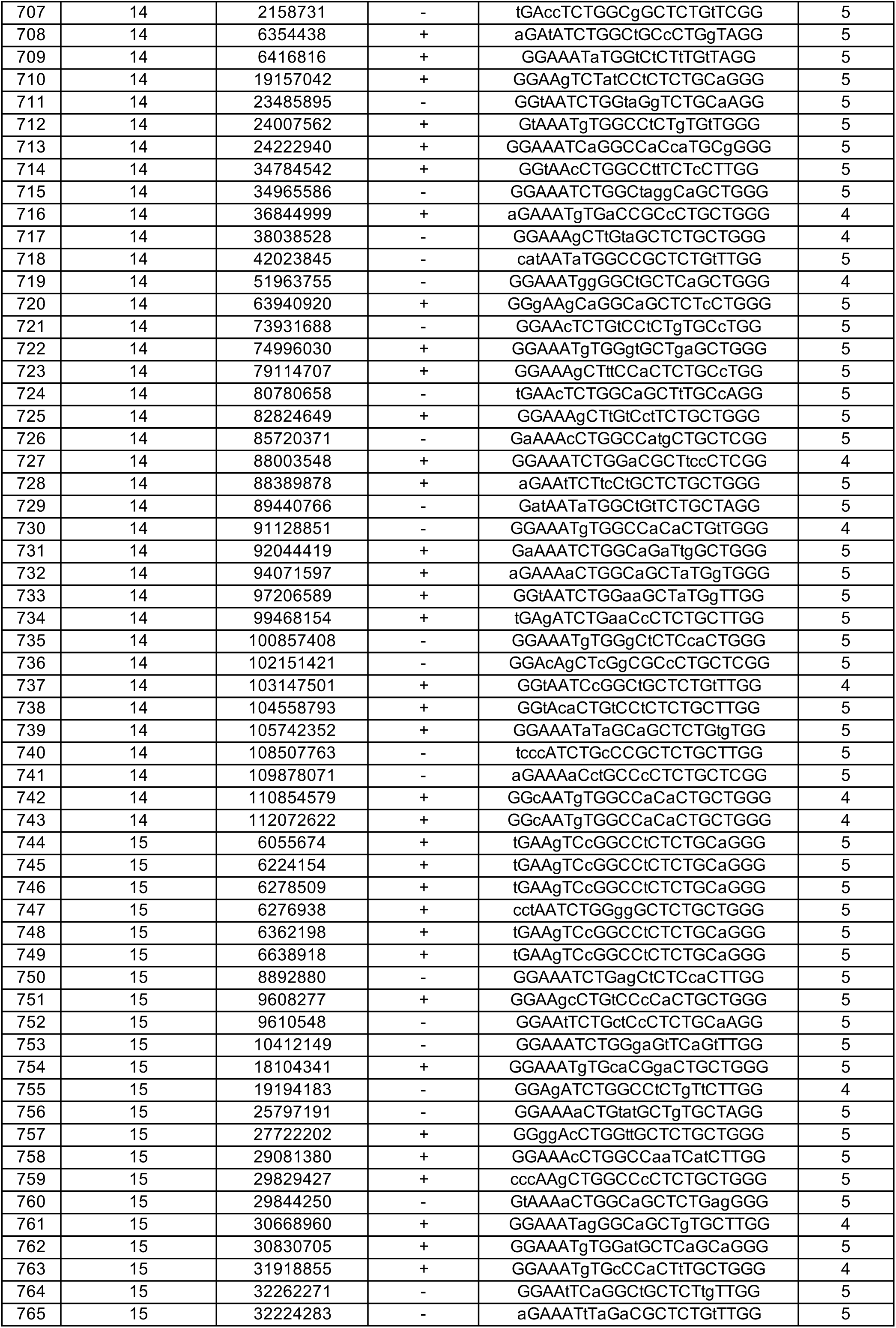
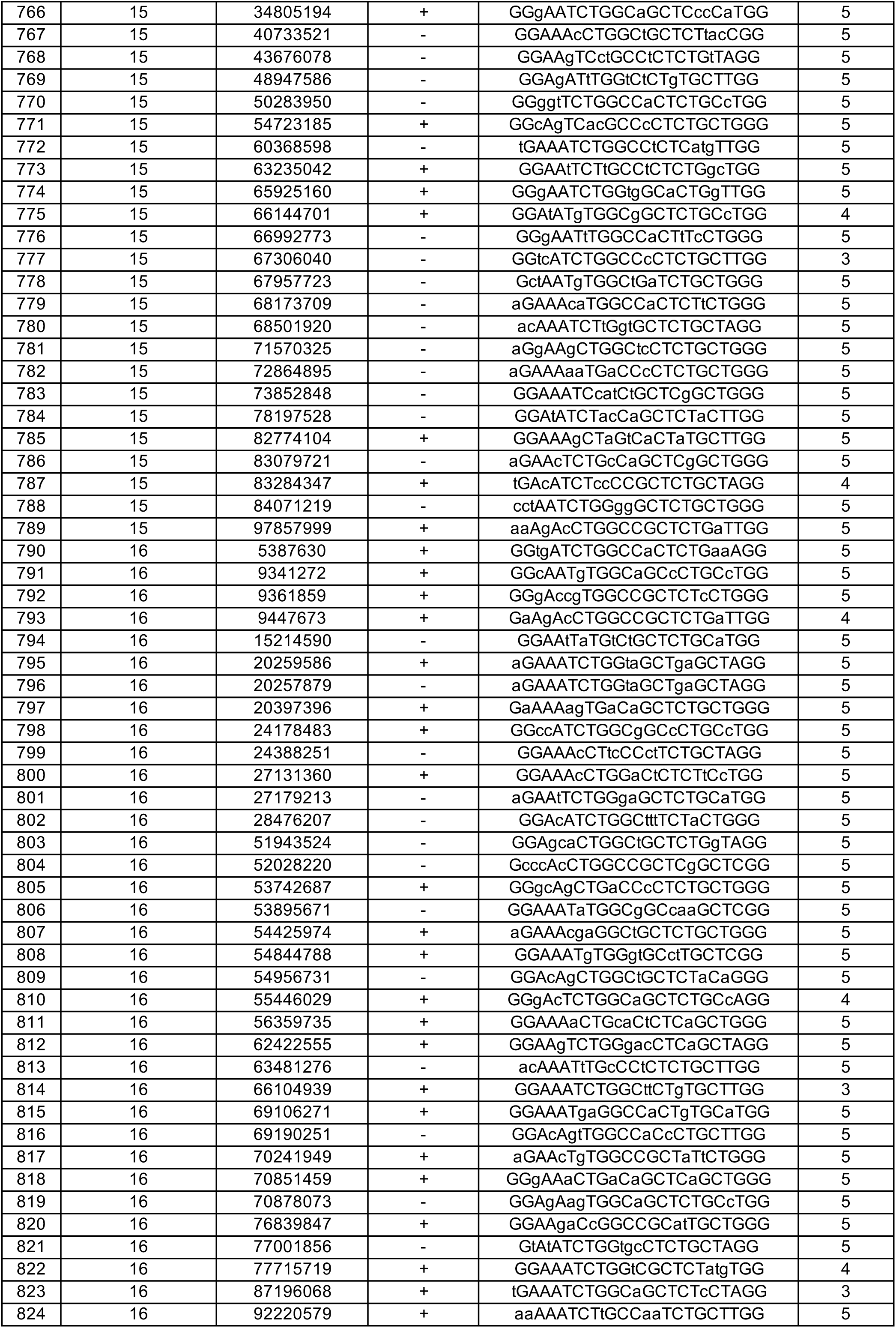
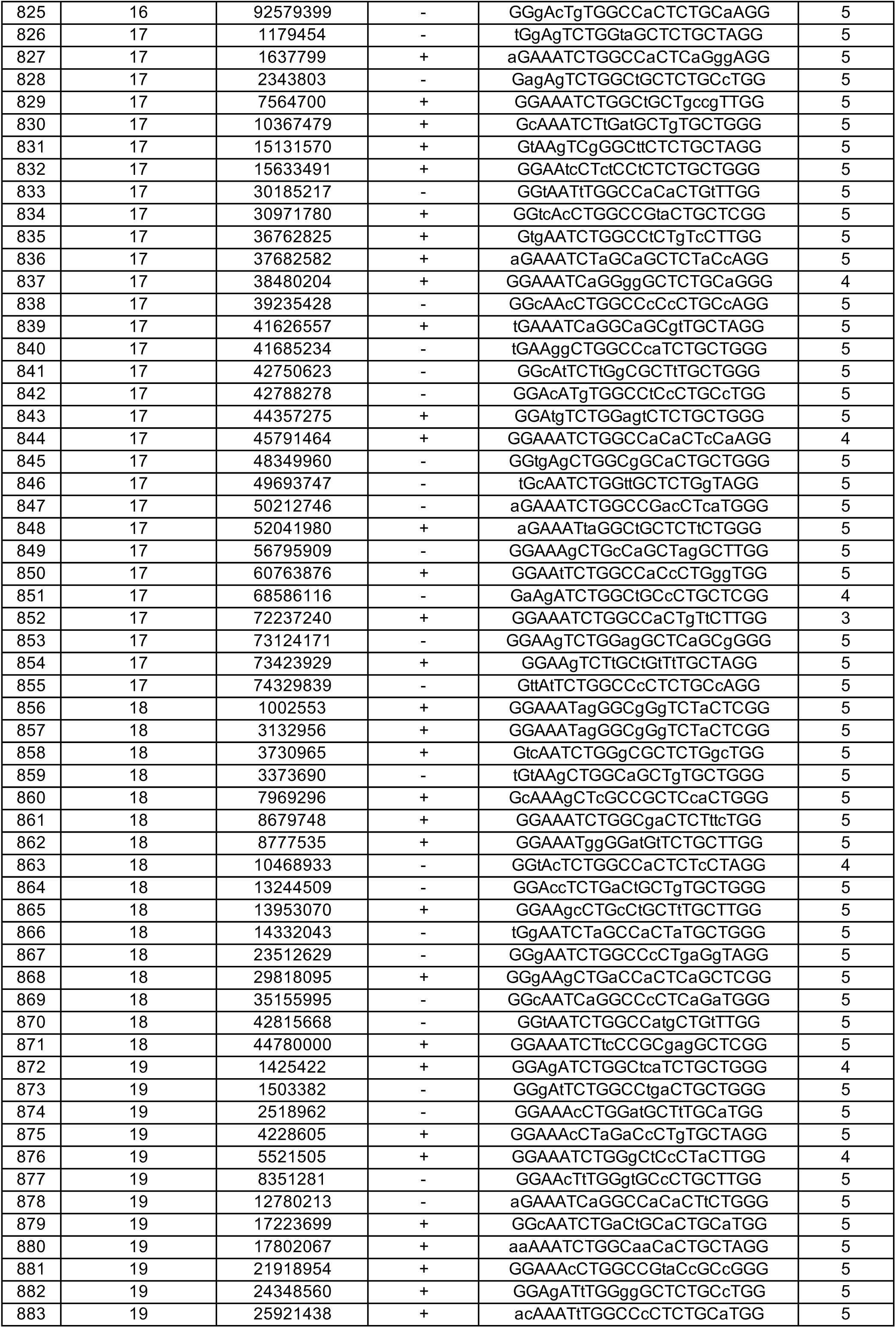
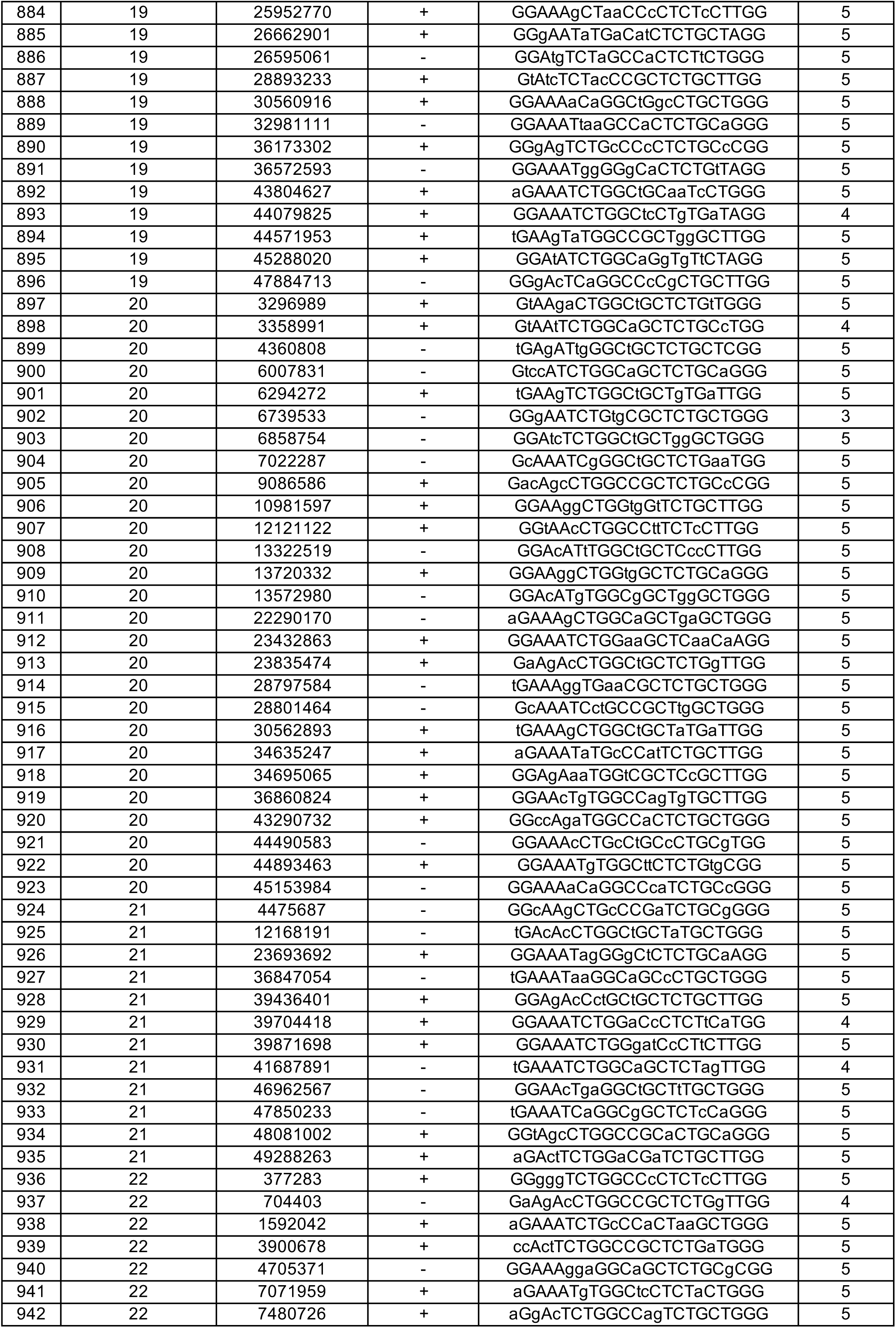
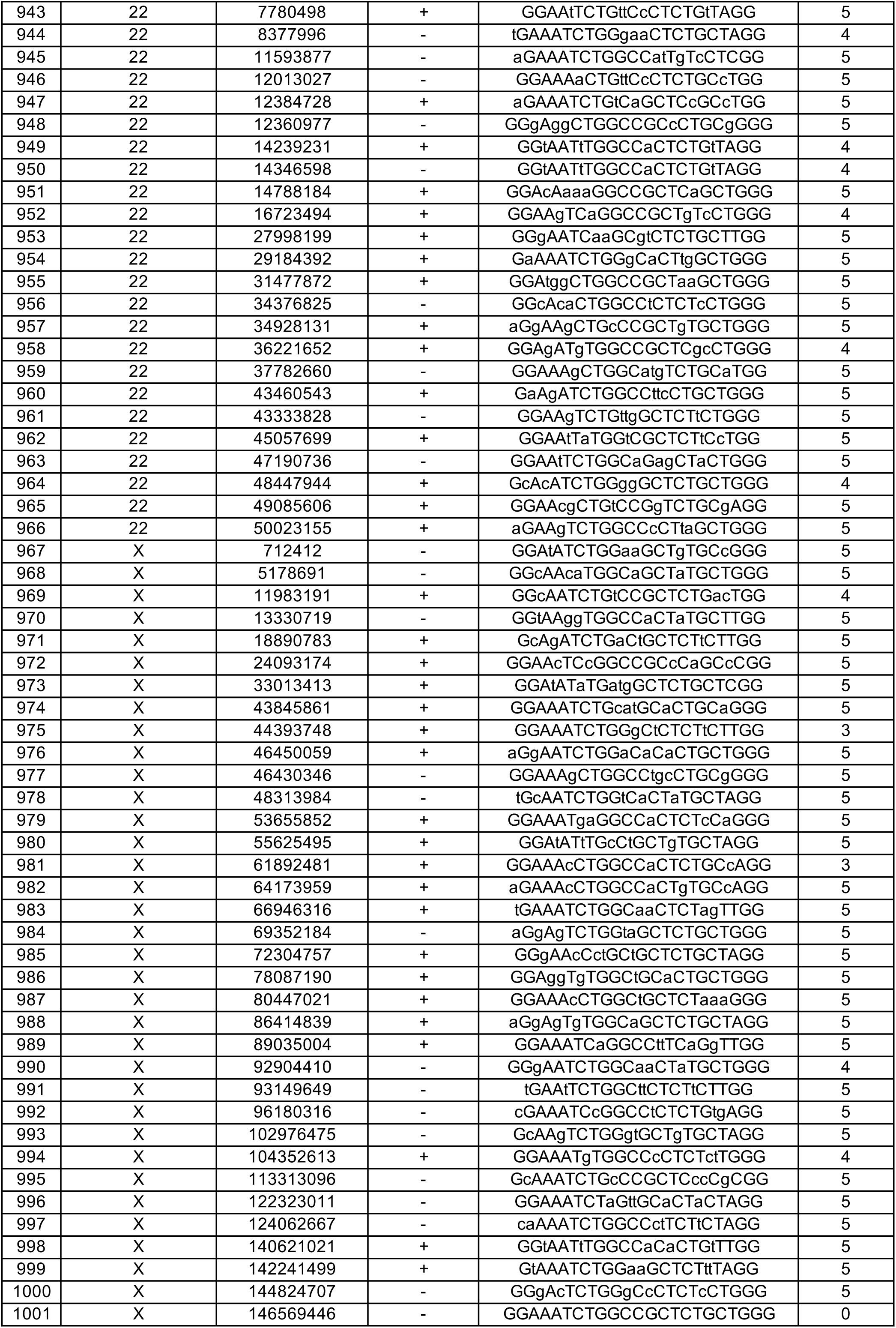
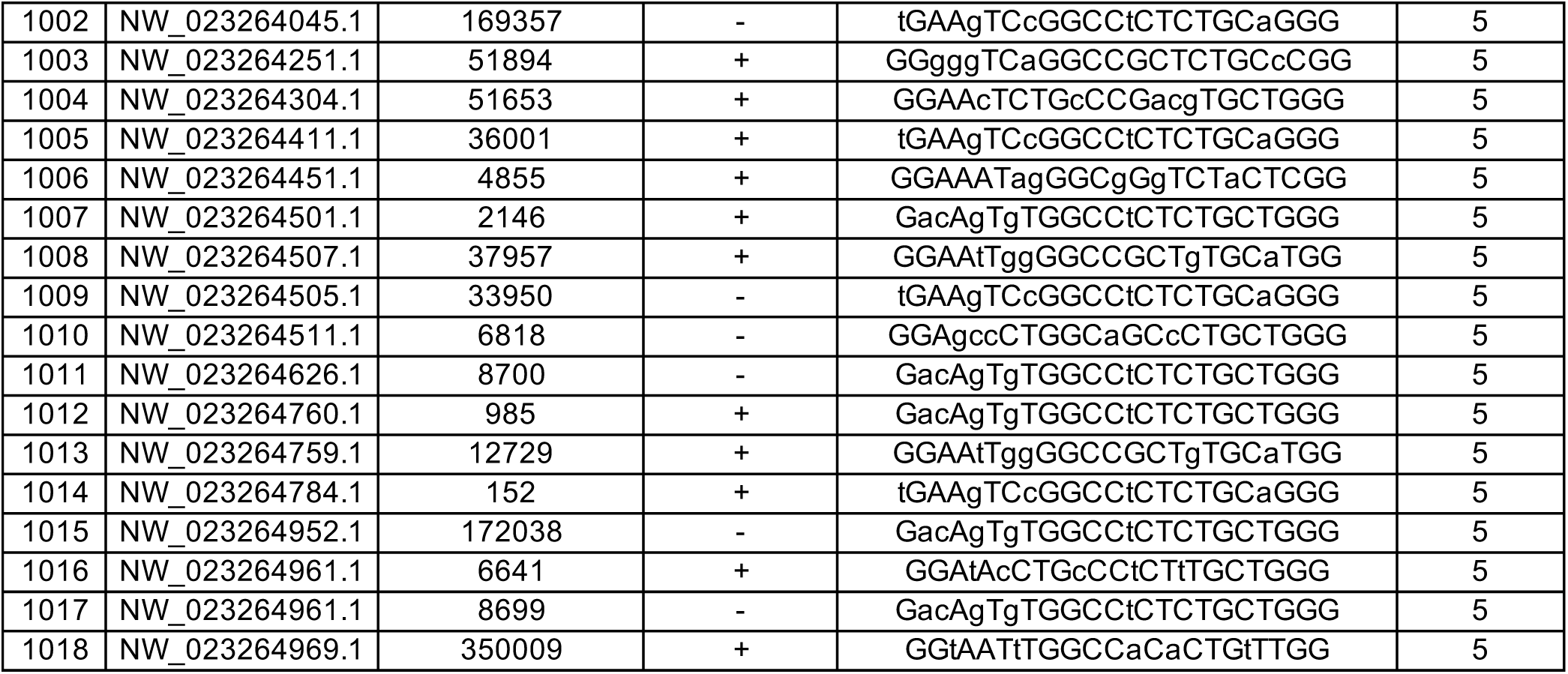
Putative Off-Target Sequences of MECP2 CRISPR/Cas9 (Mismatch < 6)

**Table S14.**
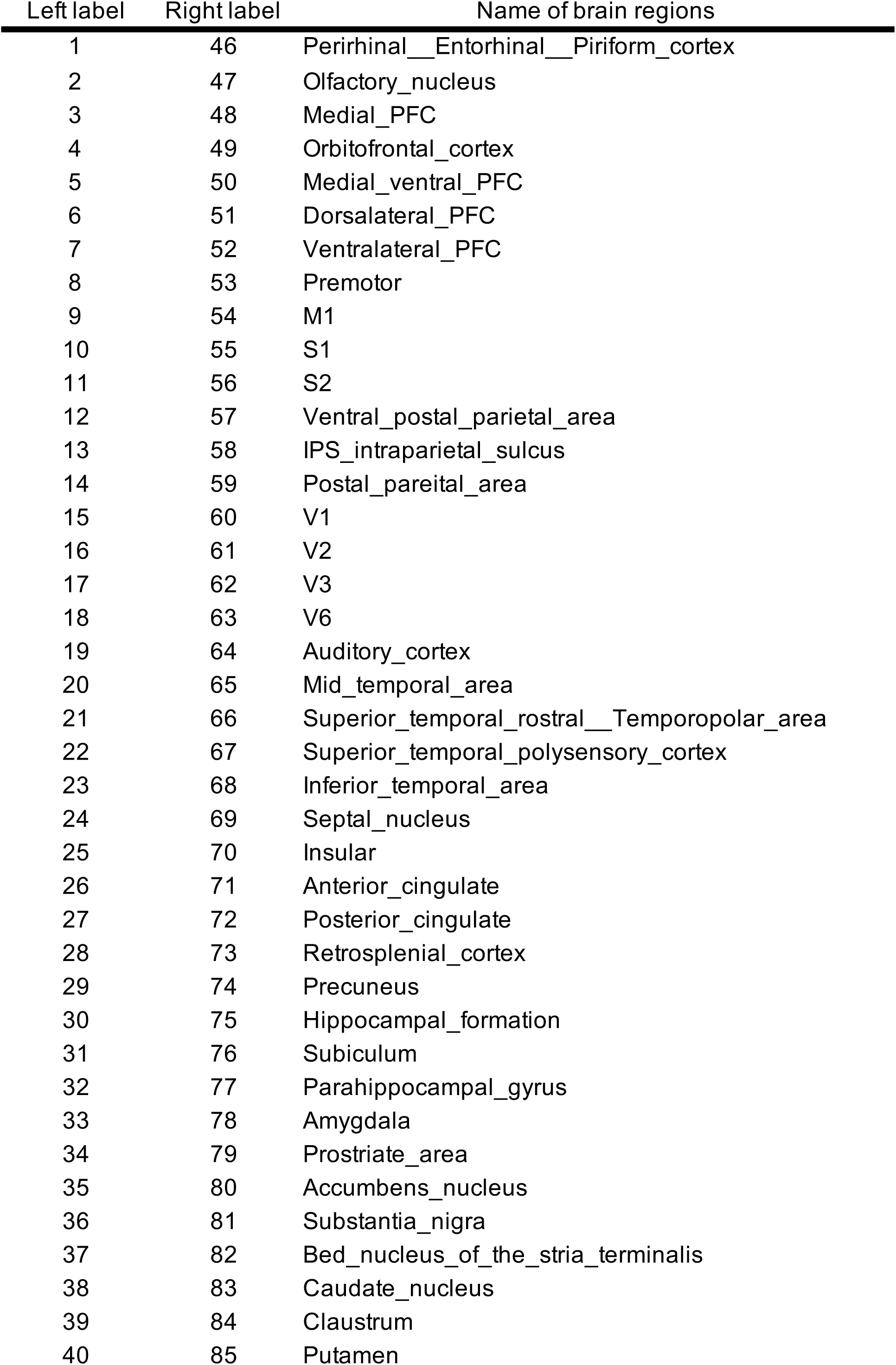
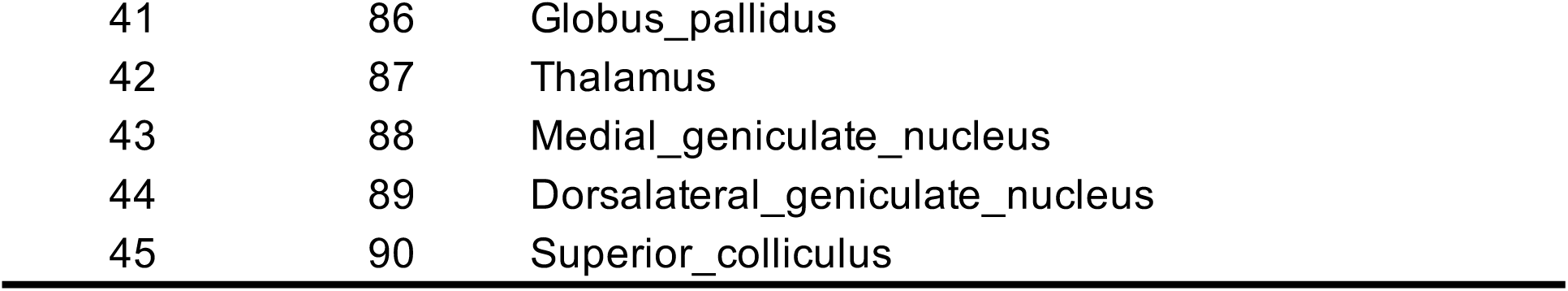
Names of the 45 brain regions.

**Table S15.**
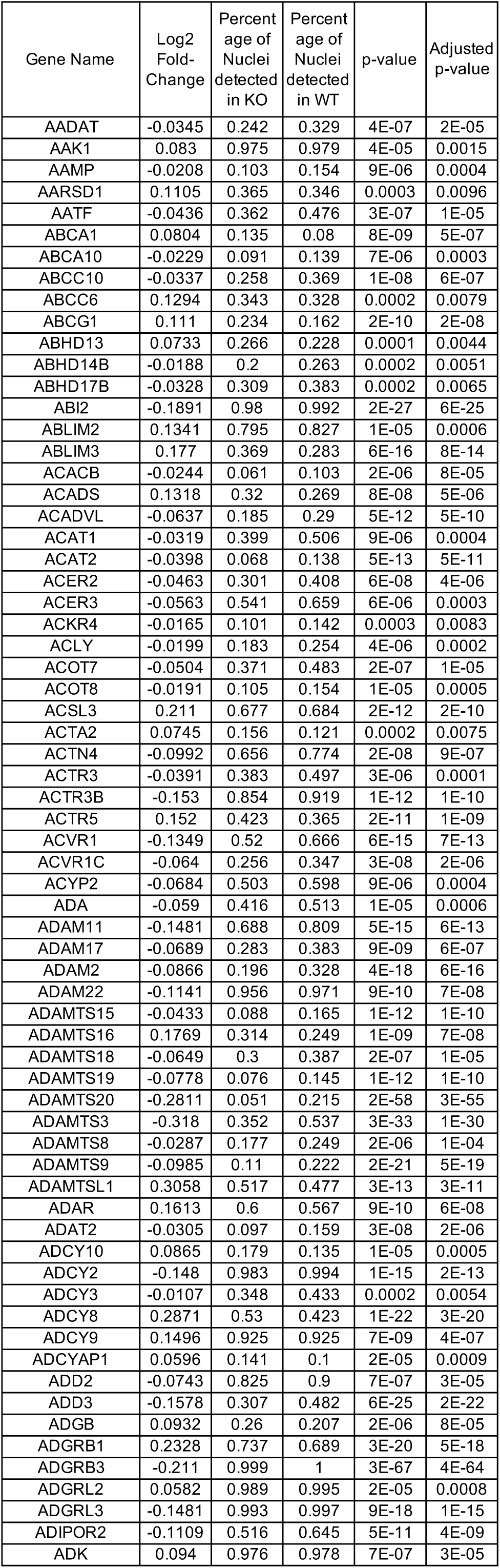
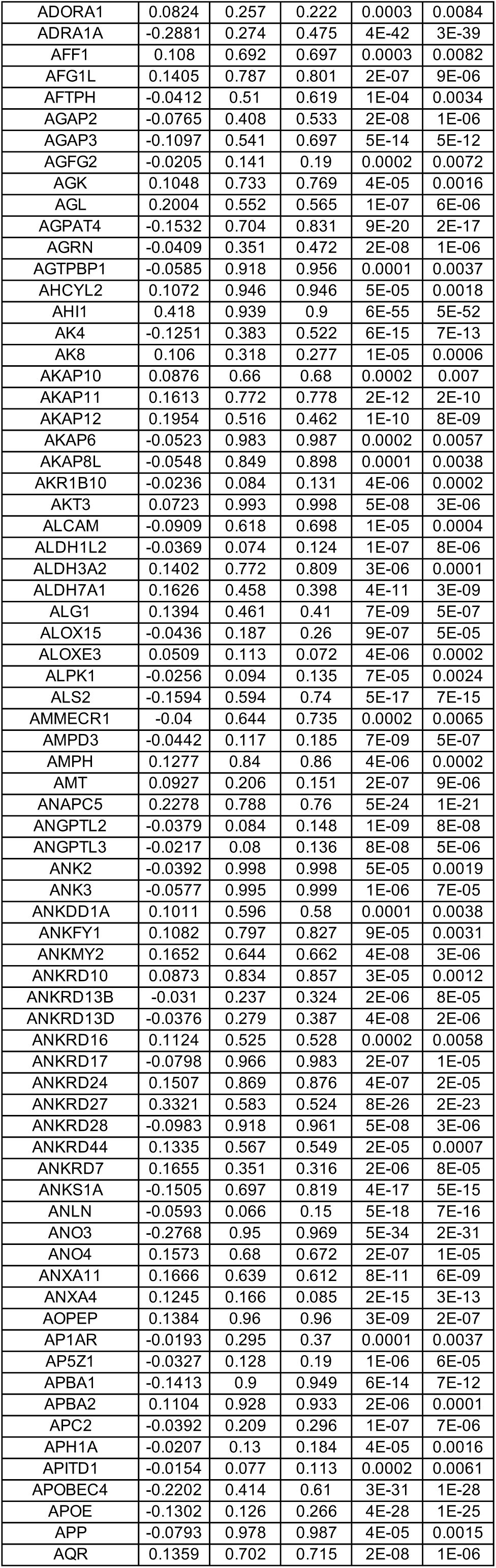
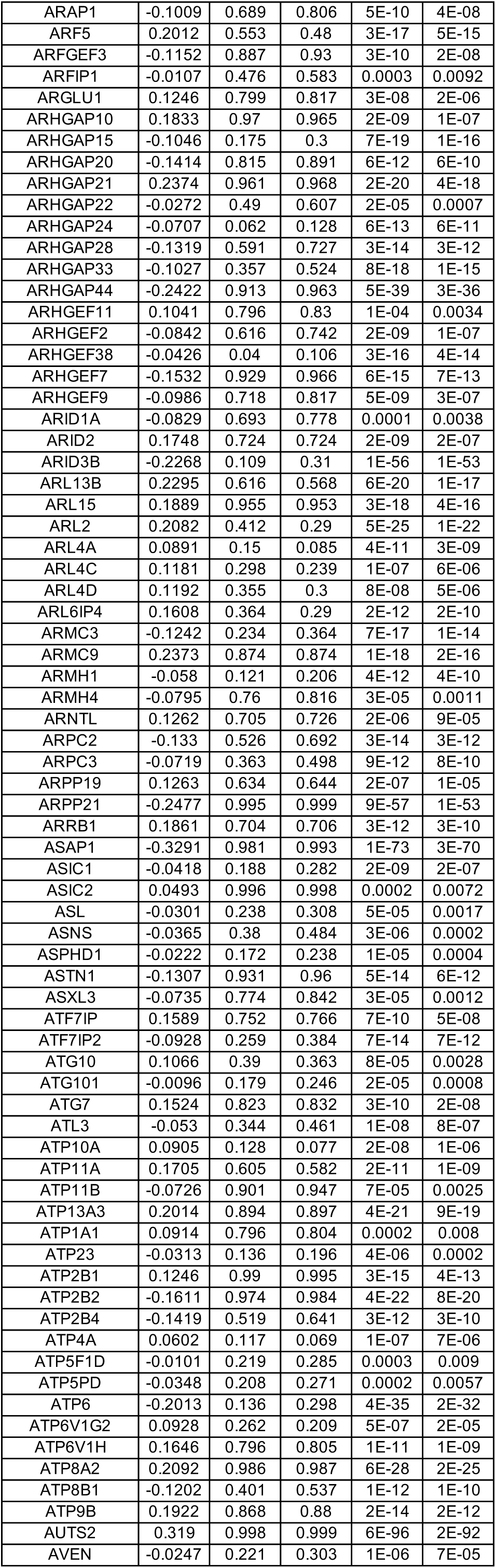
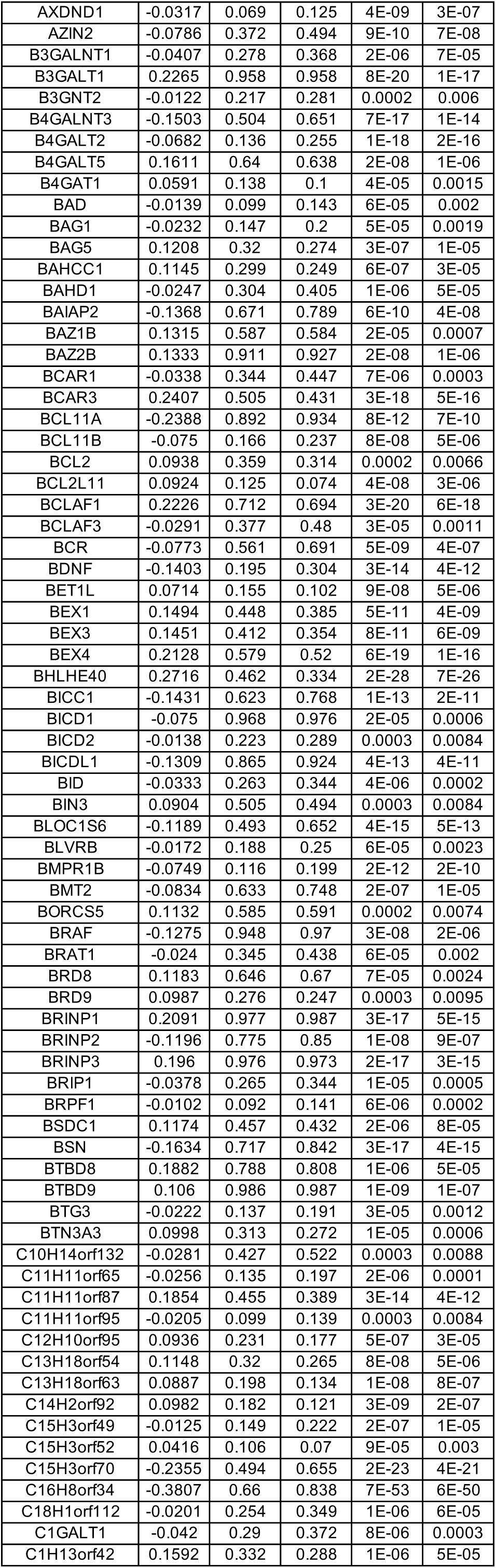
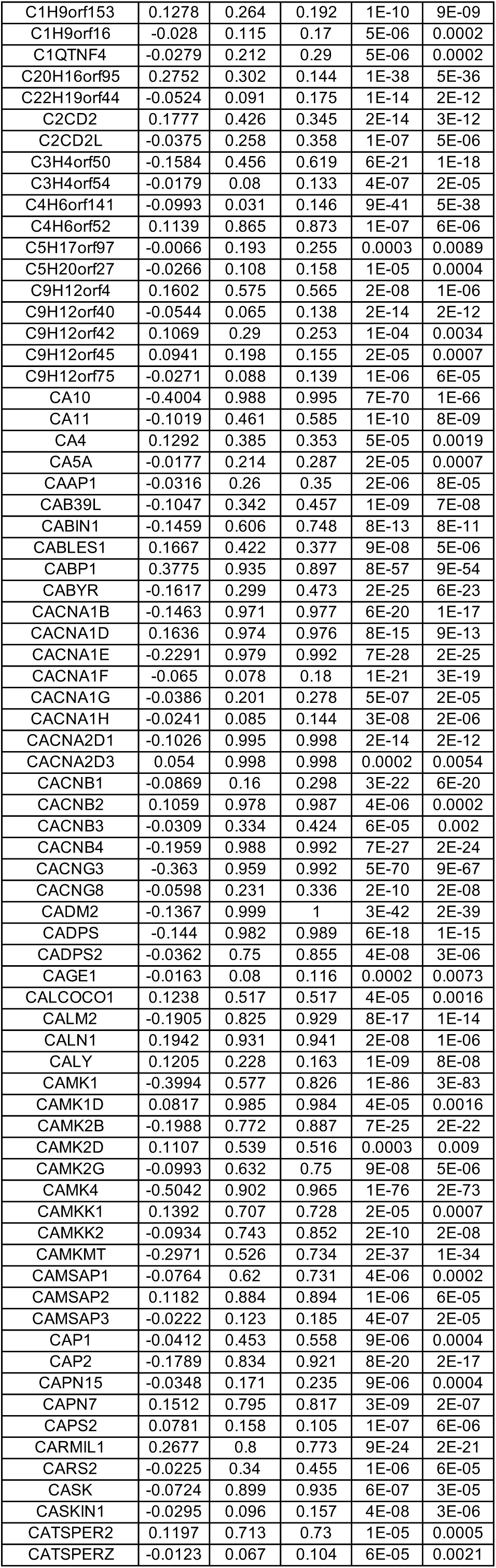
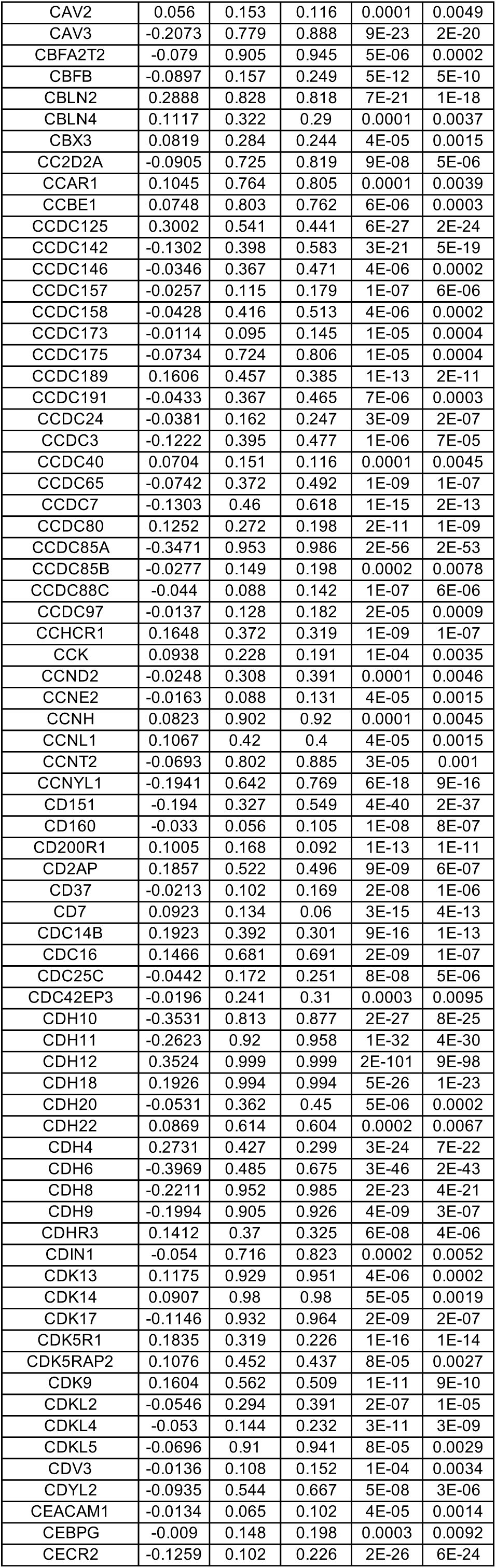
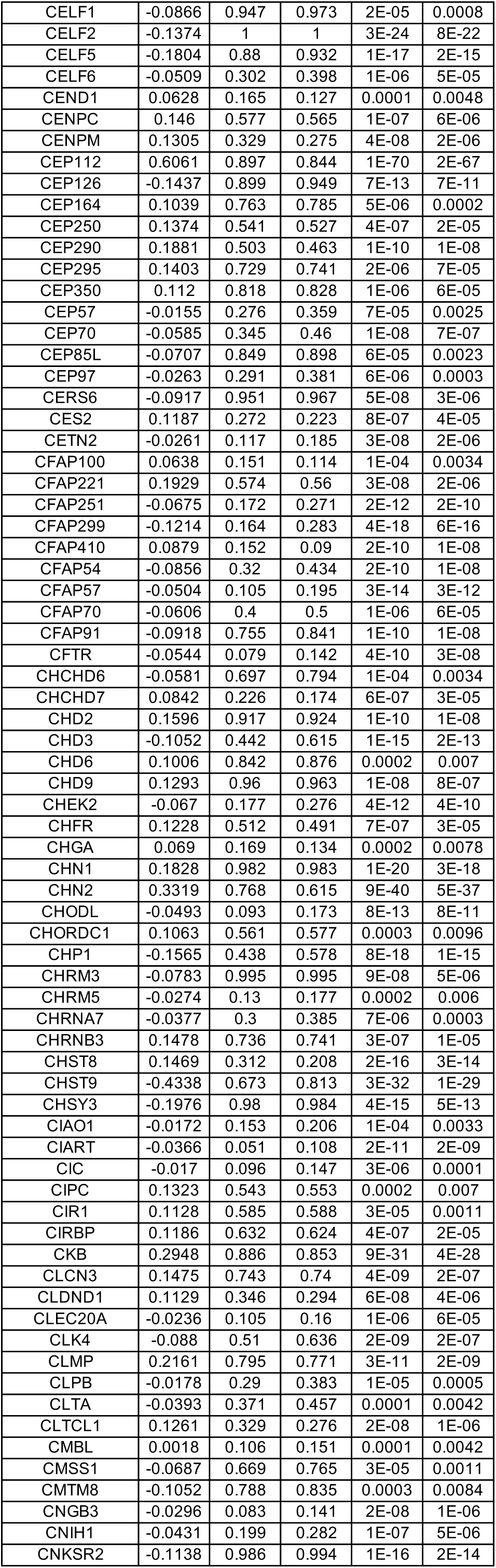
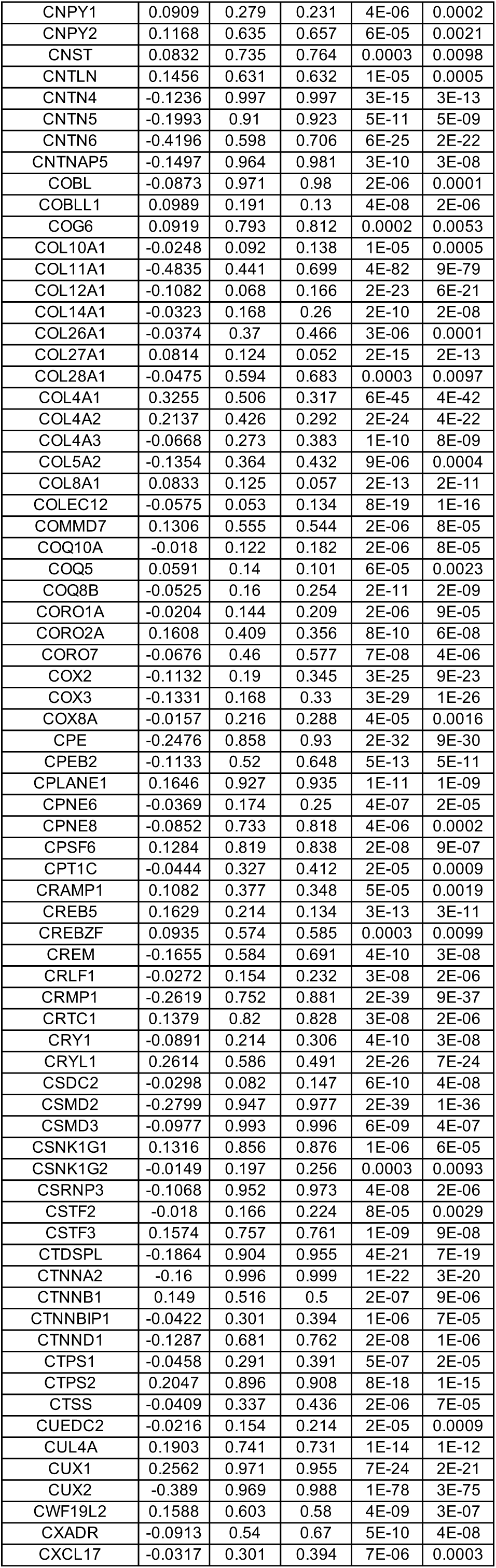
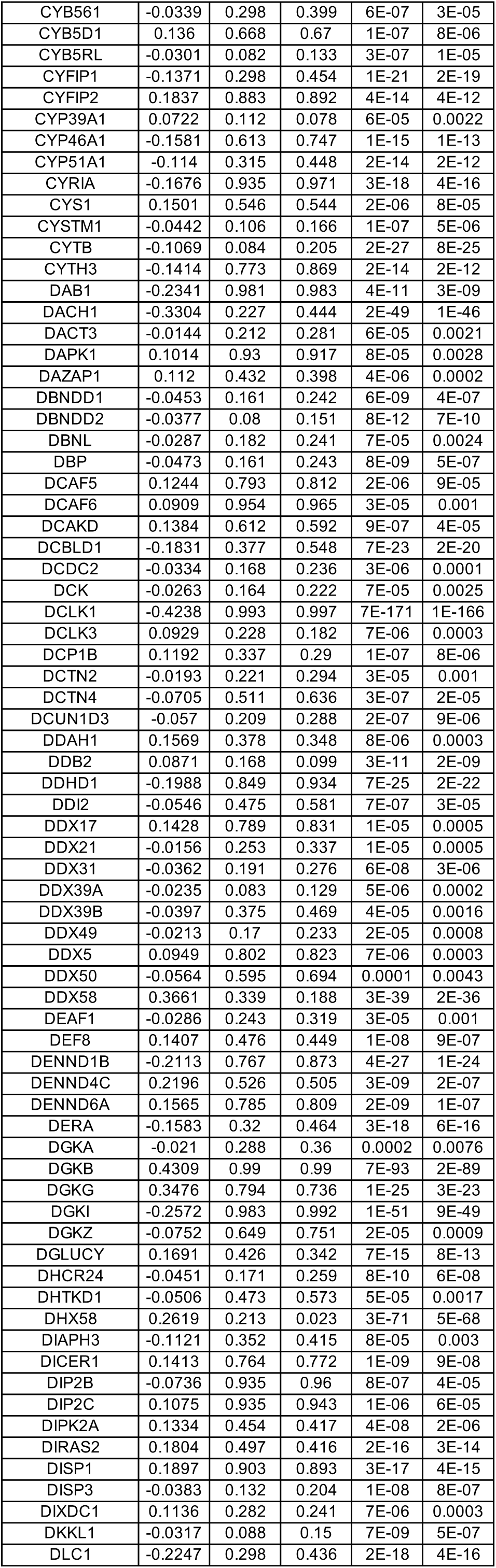
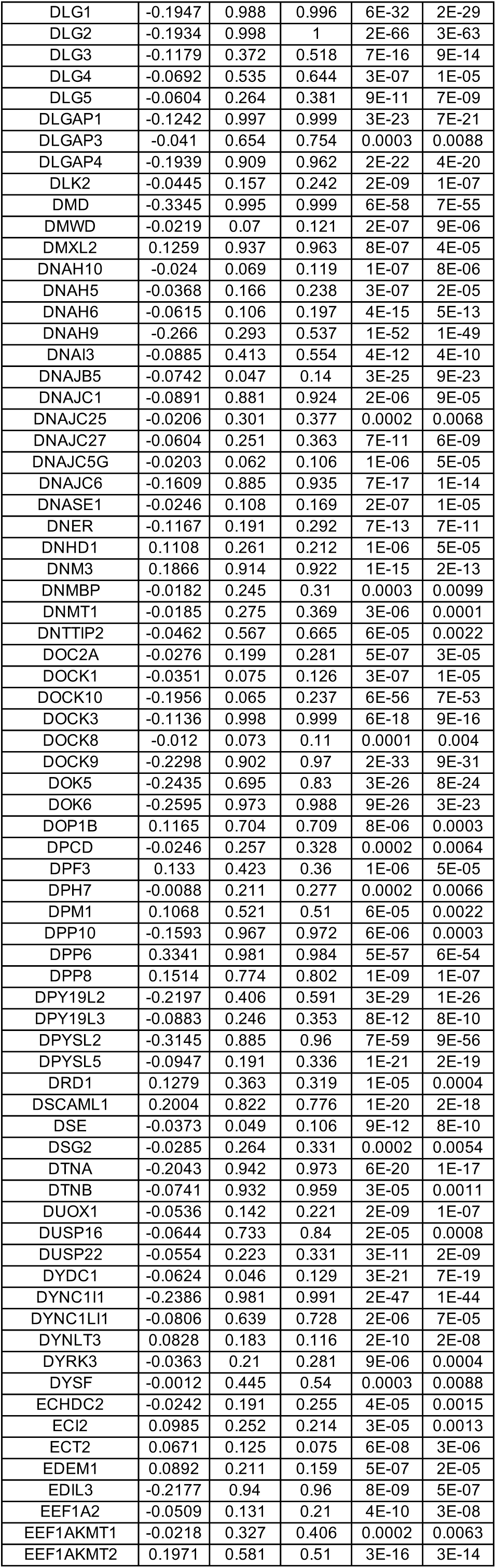
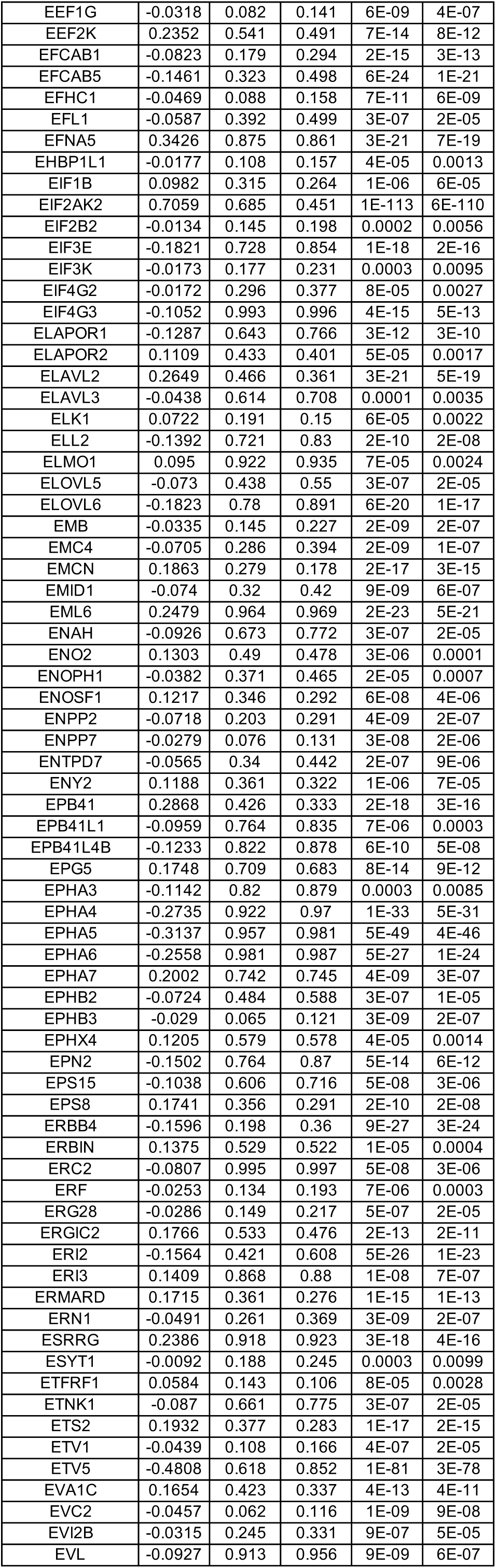
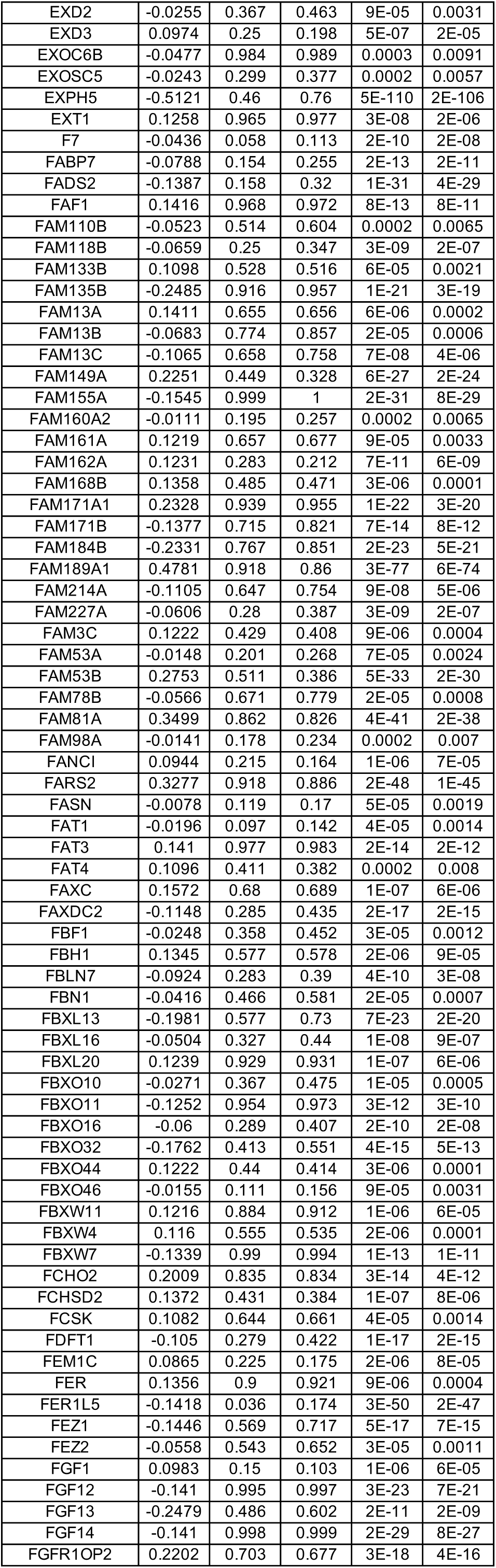
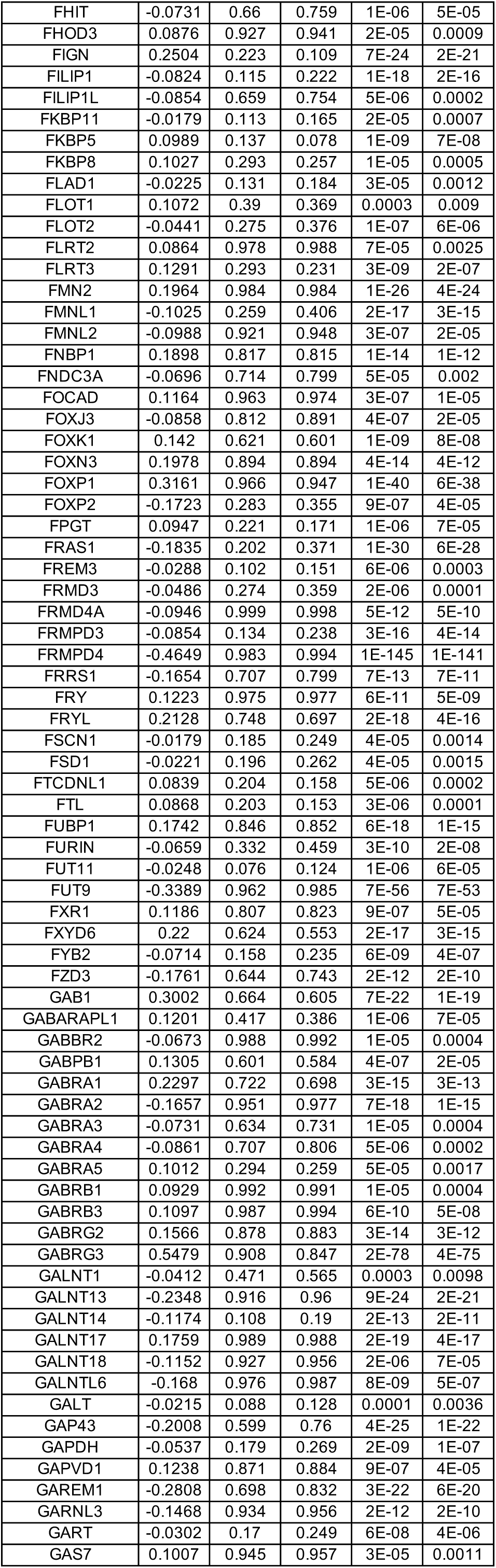
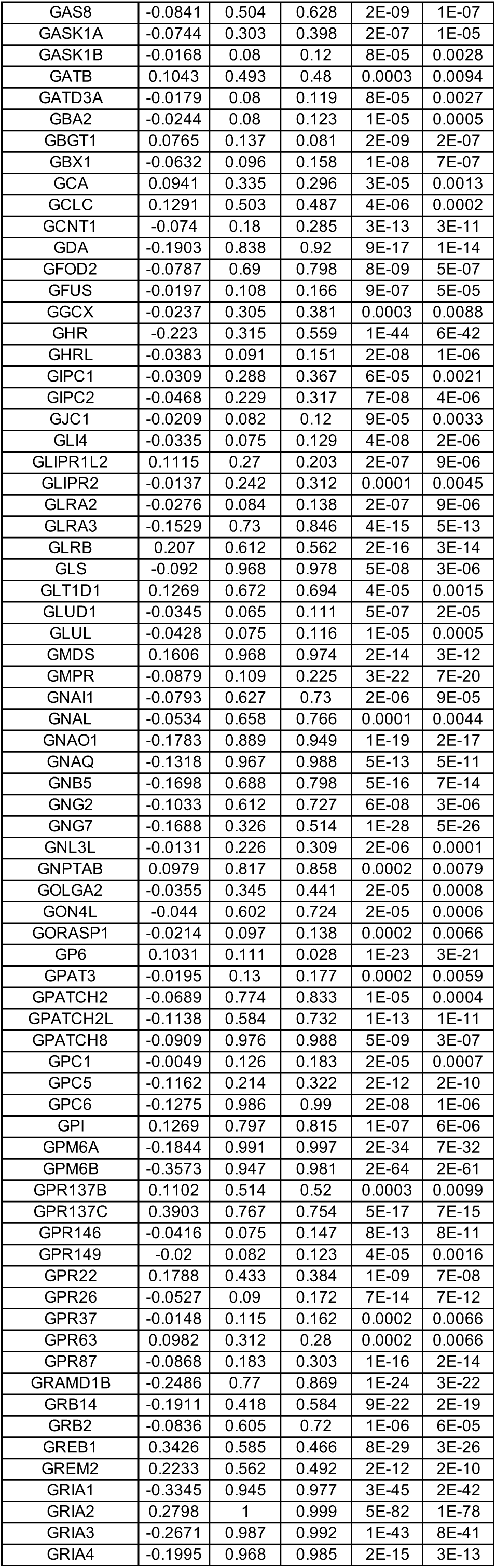
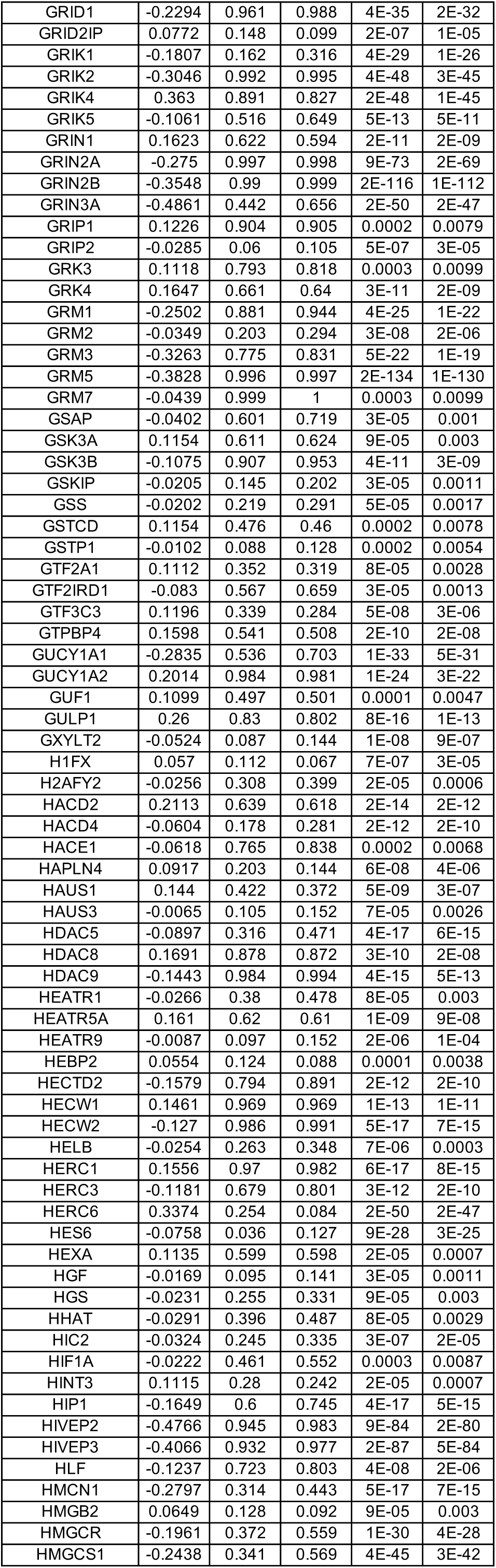
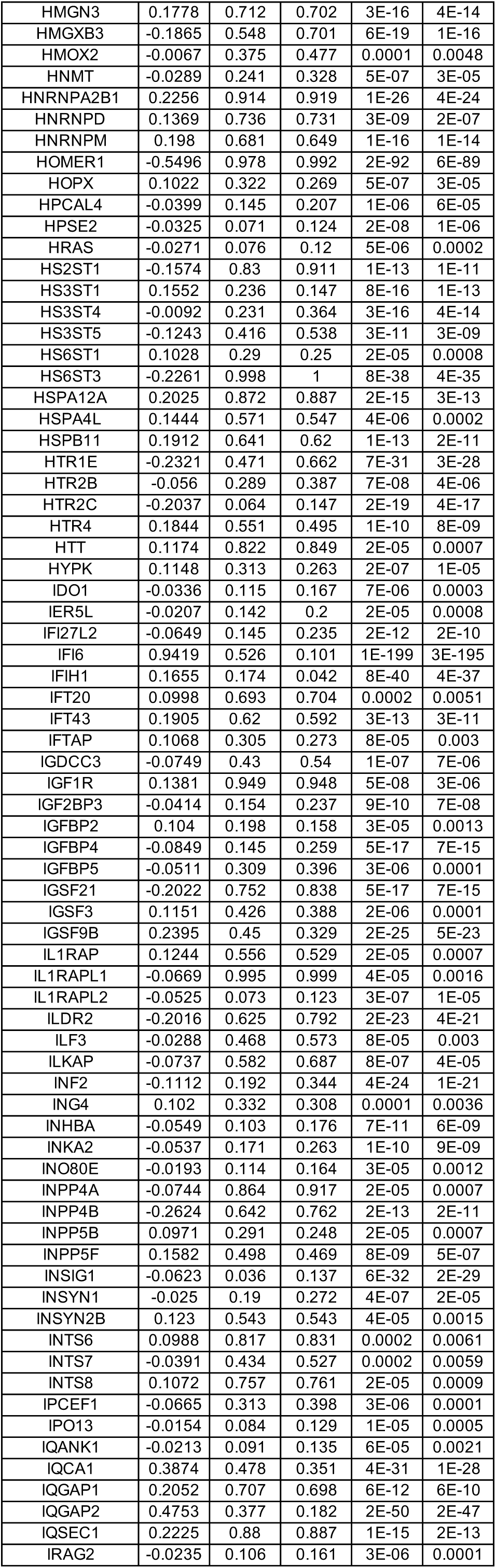
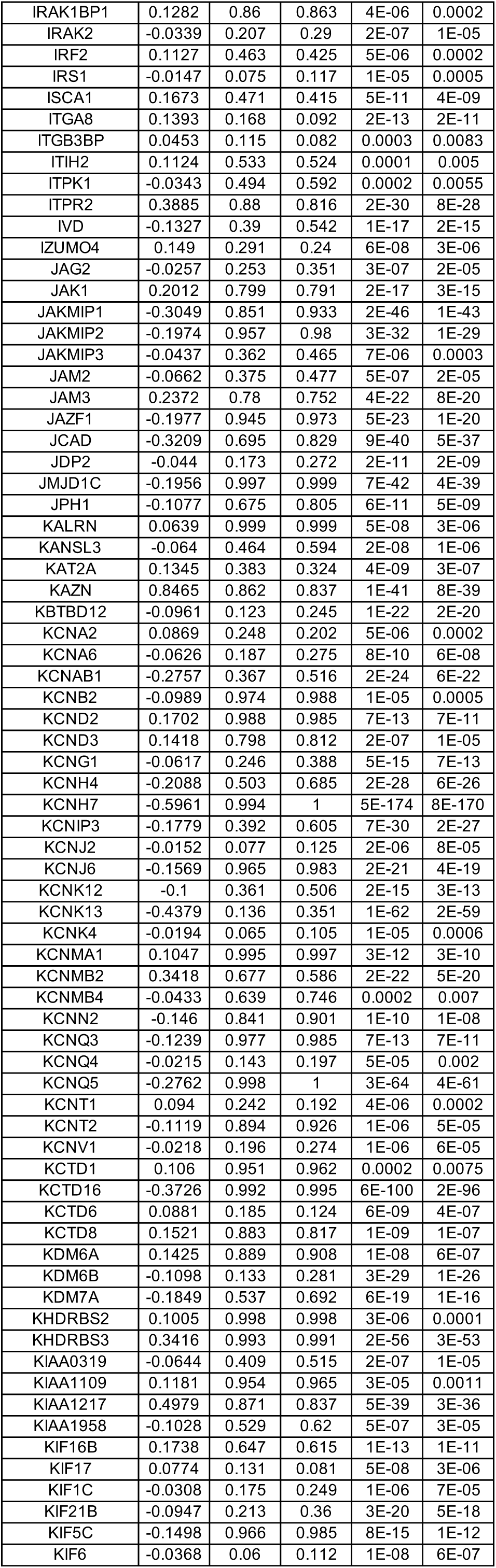
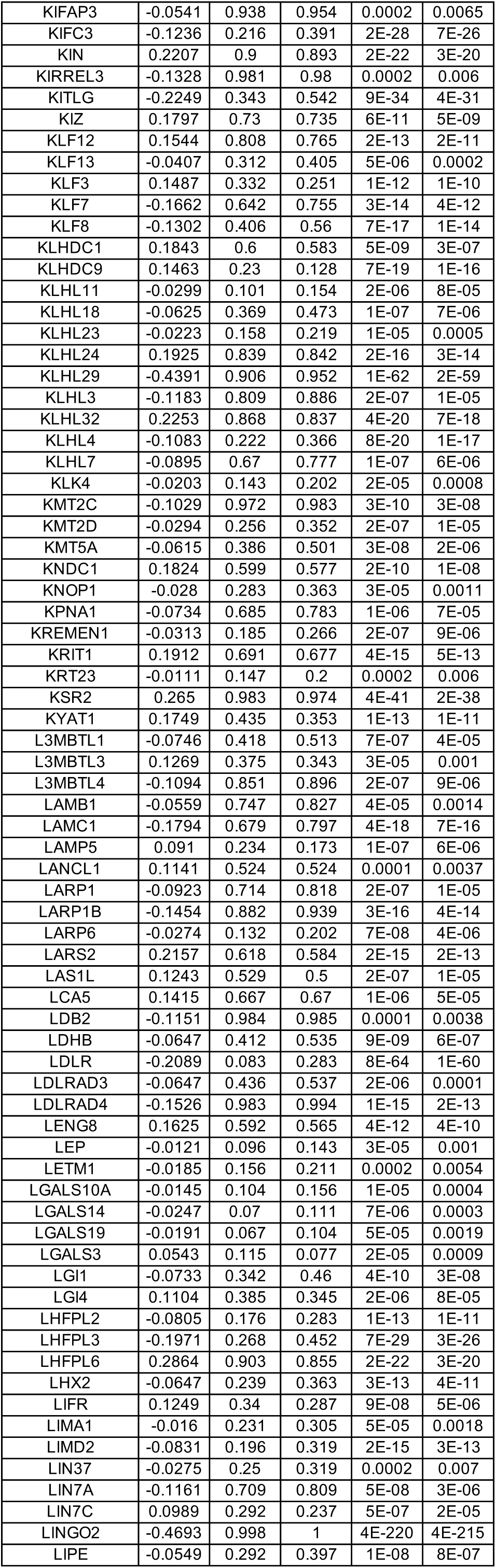
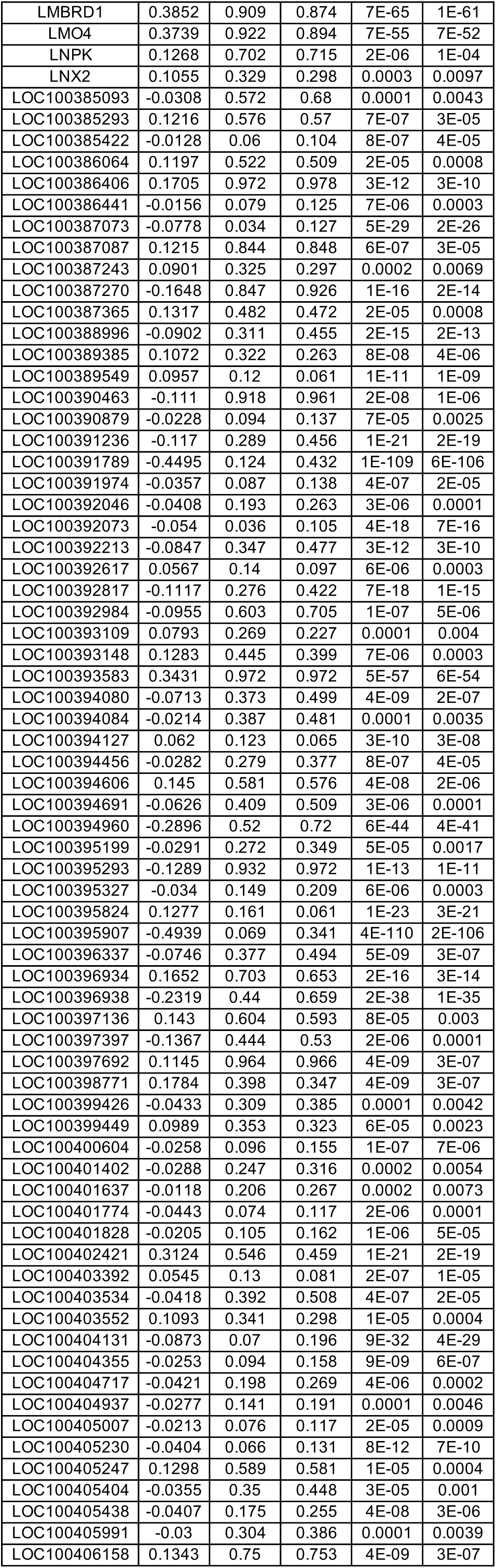
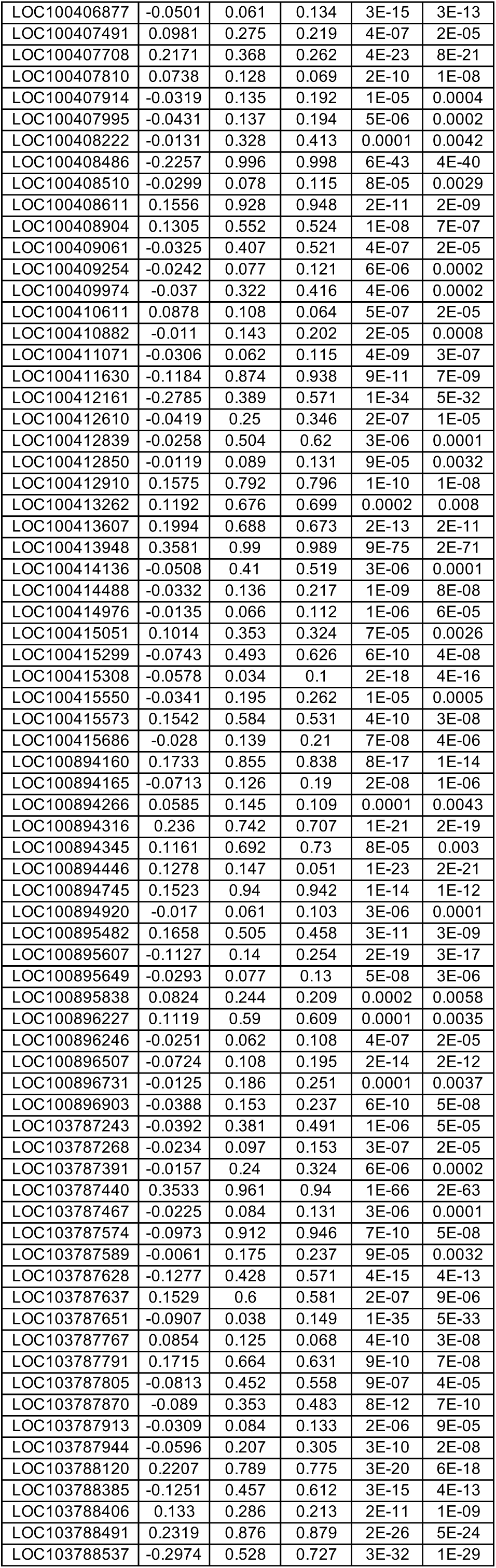
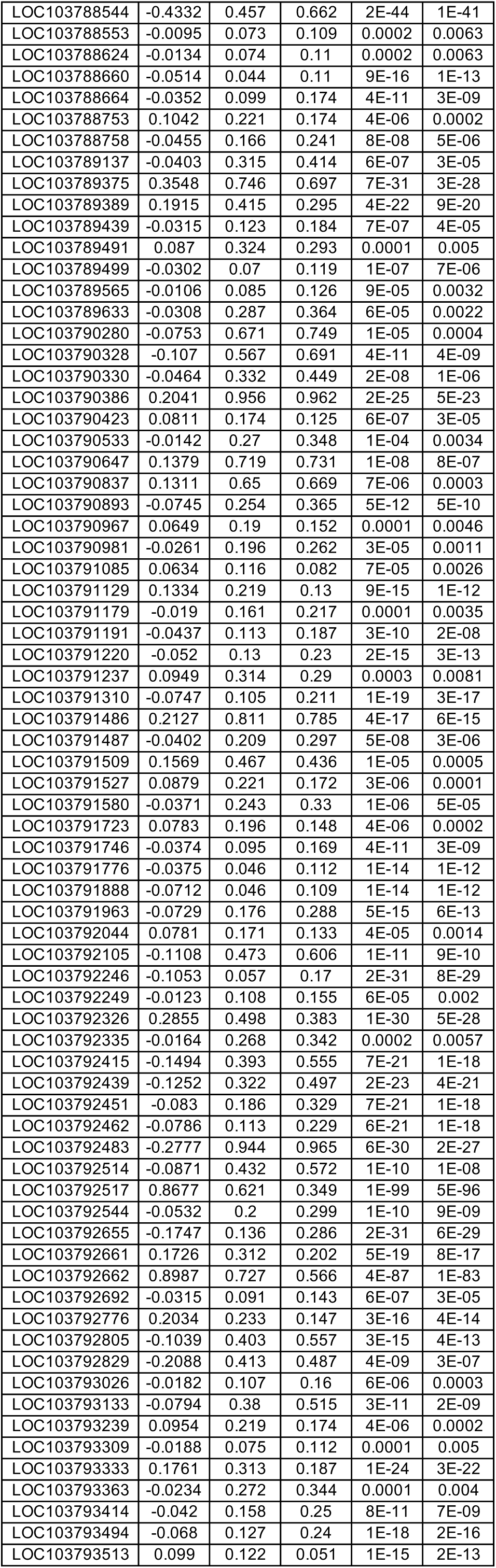
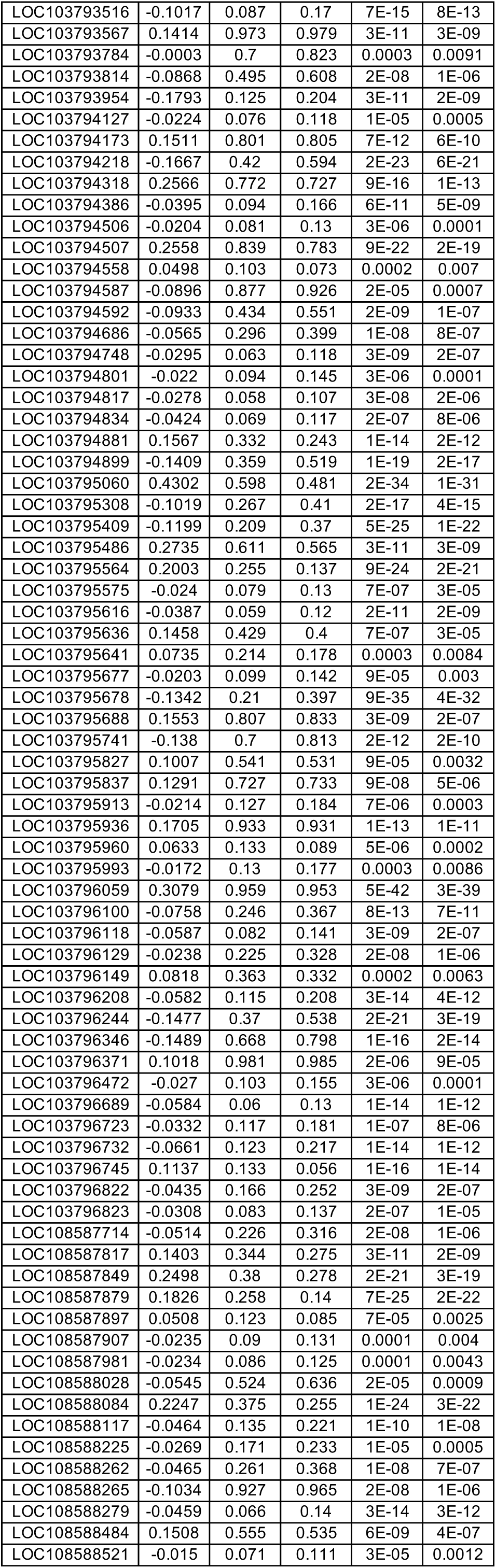
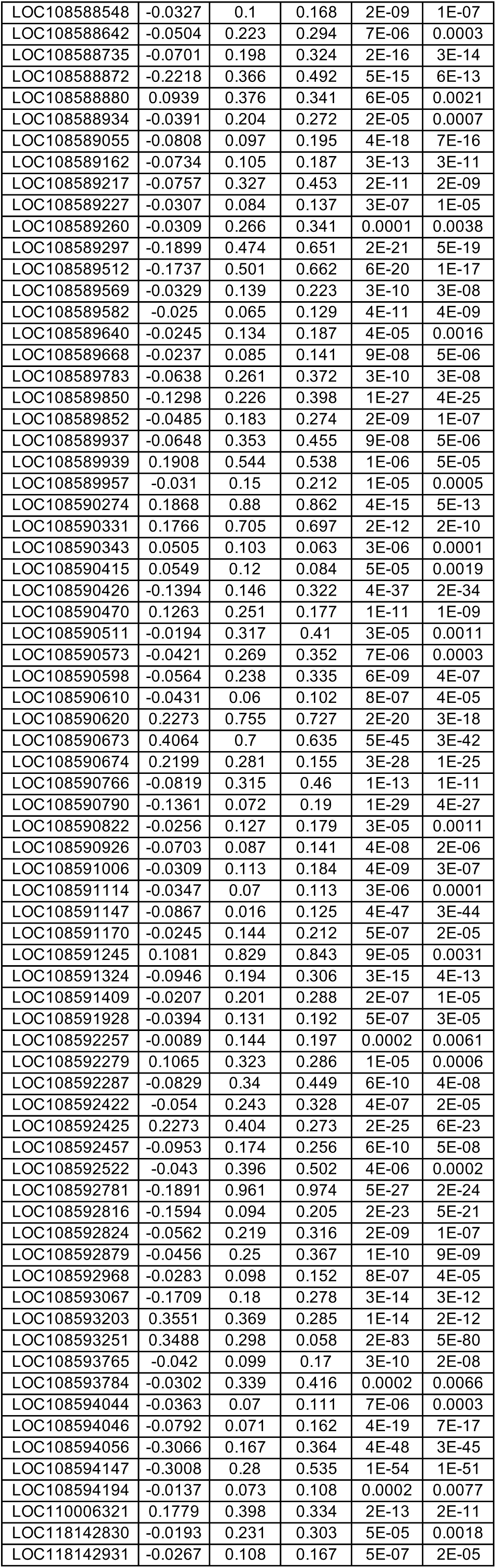
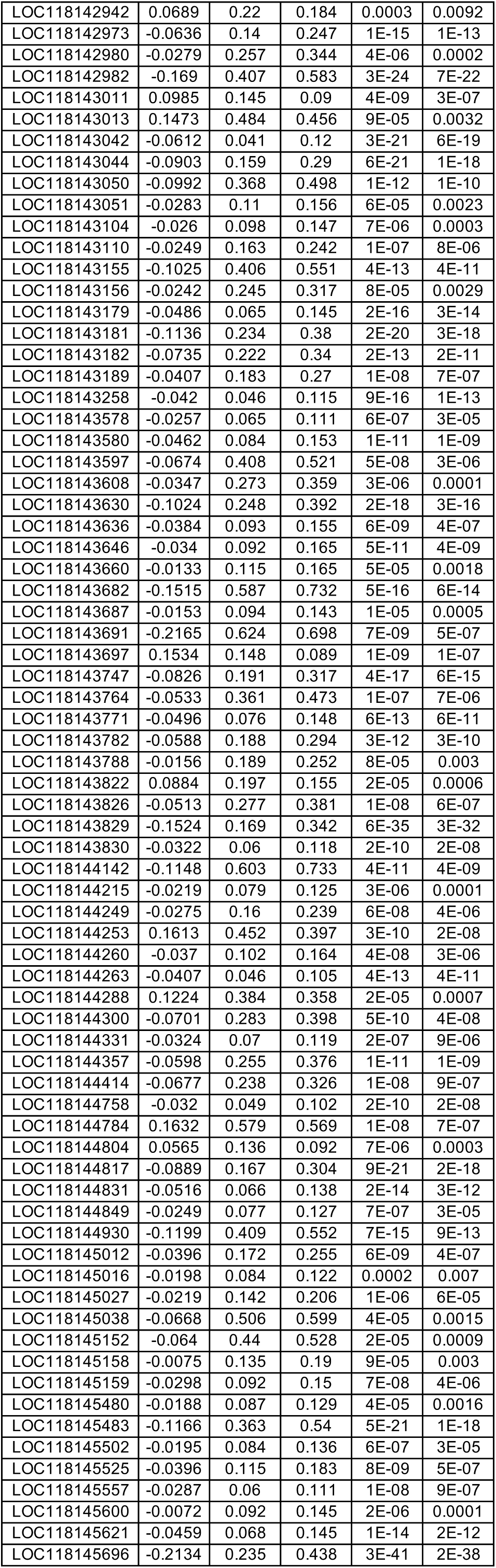
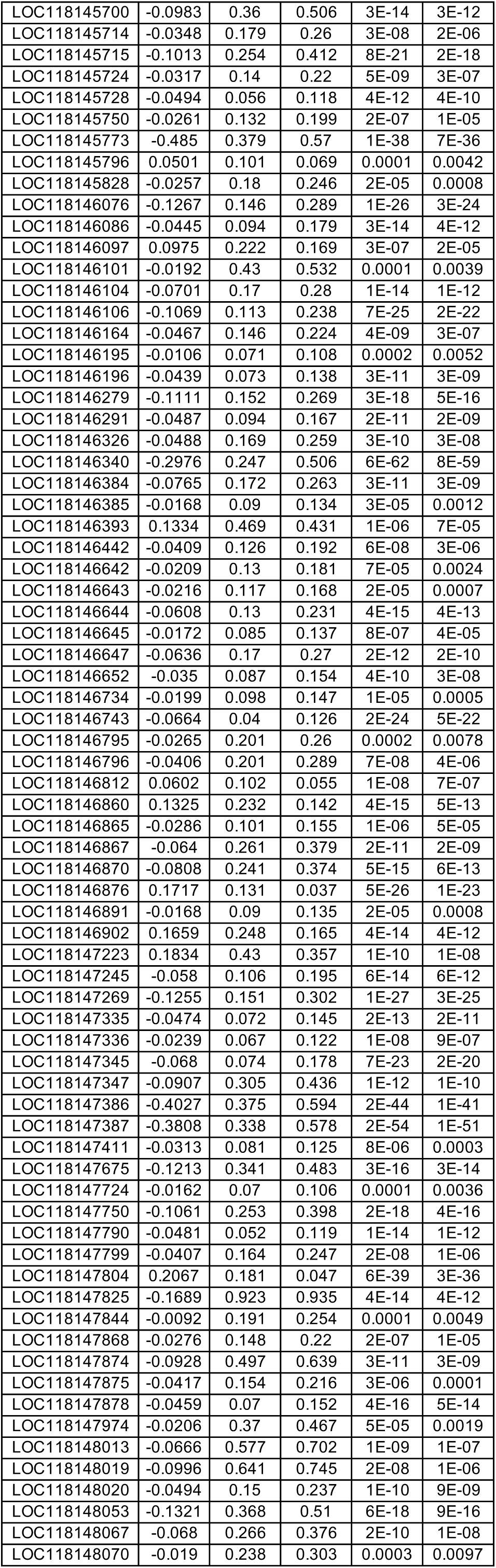
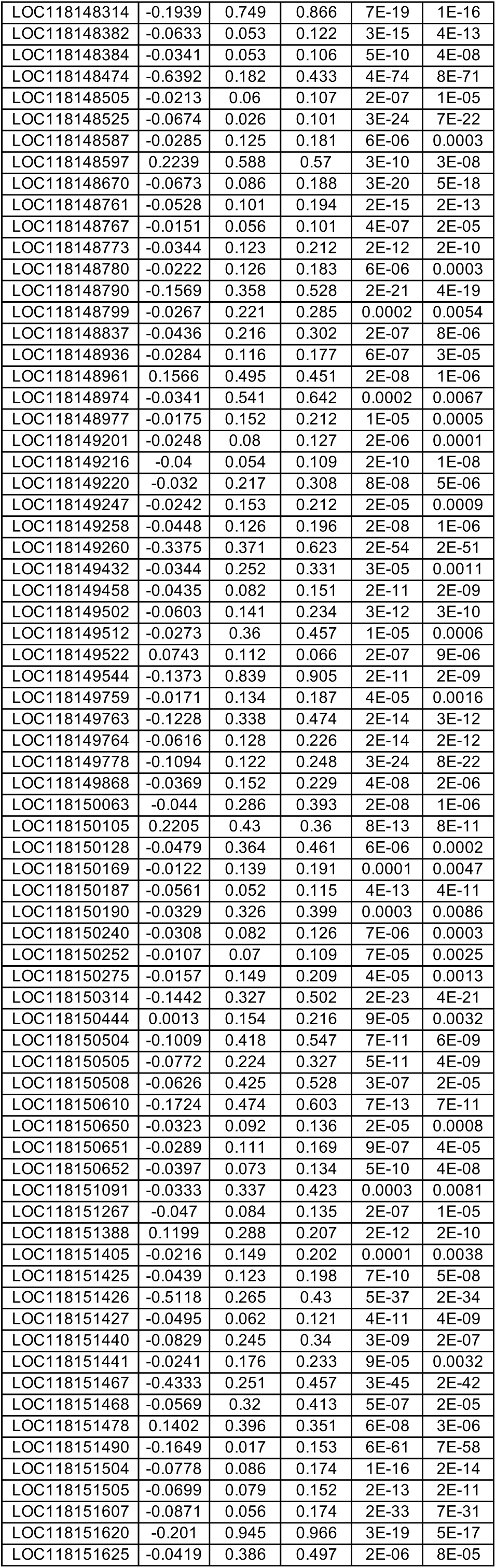
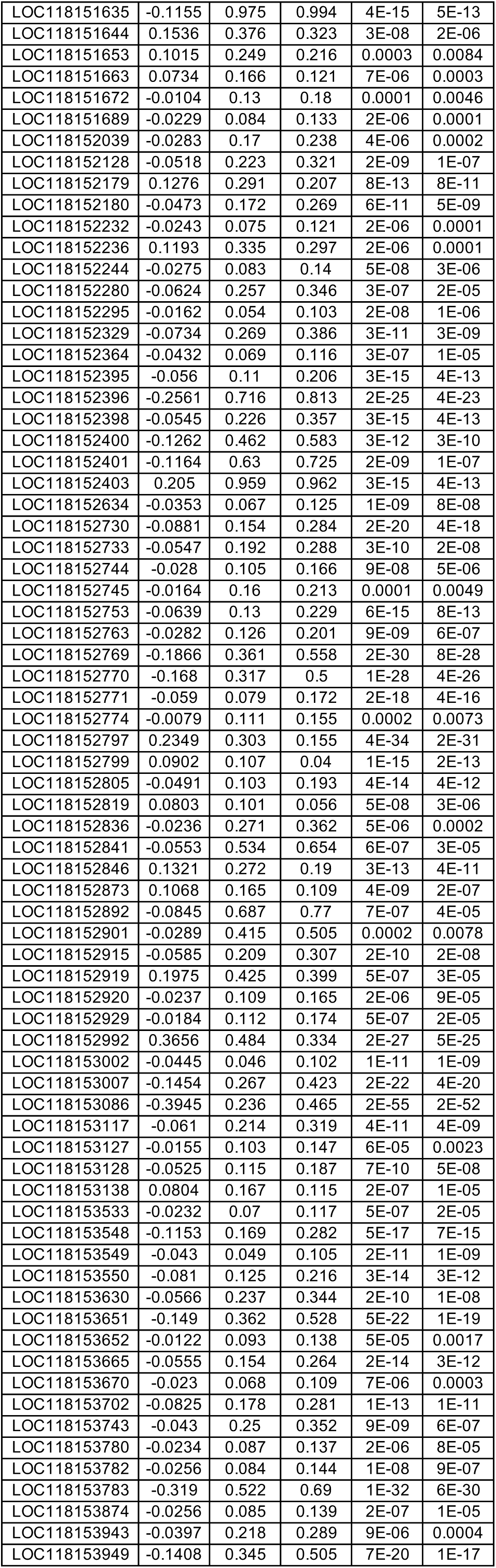
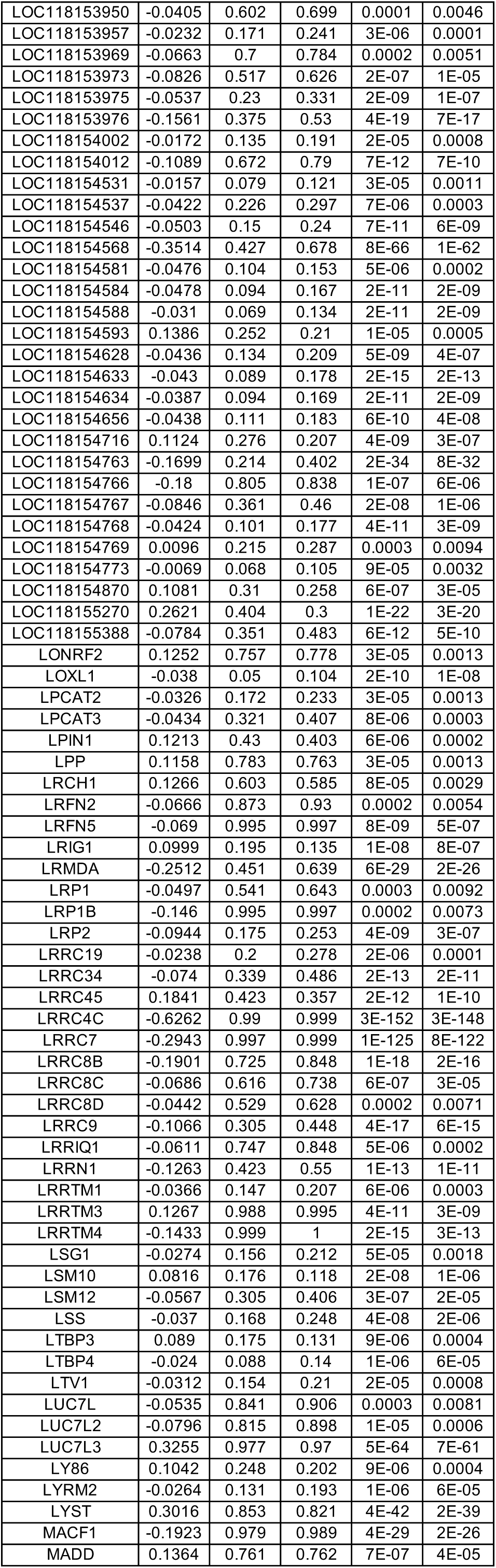
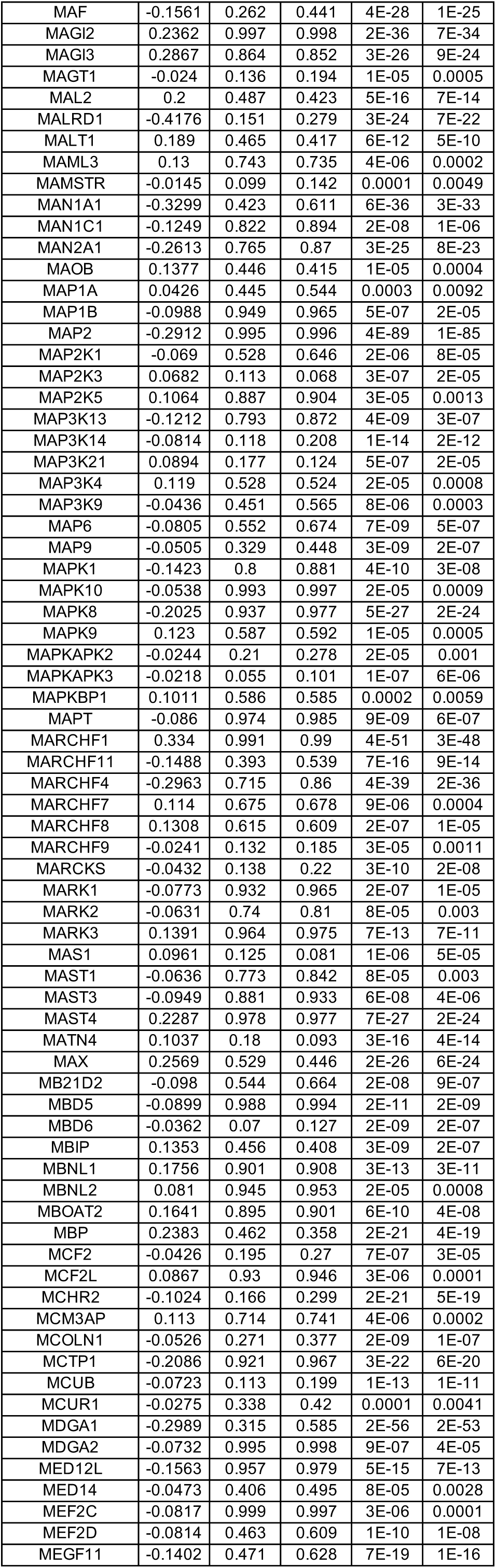
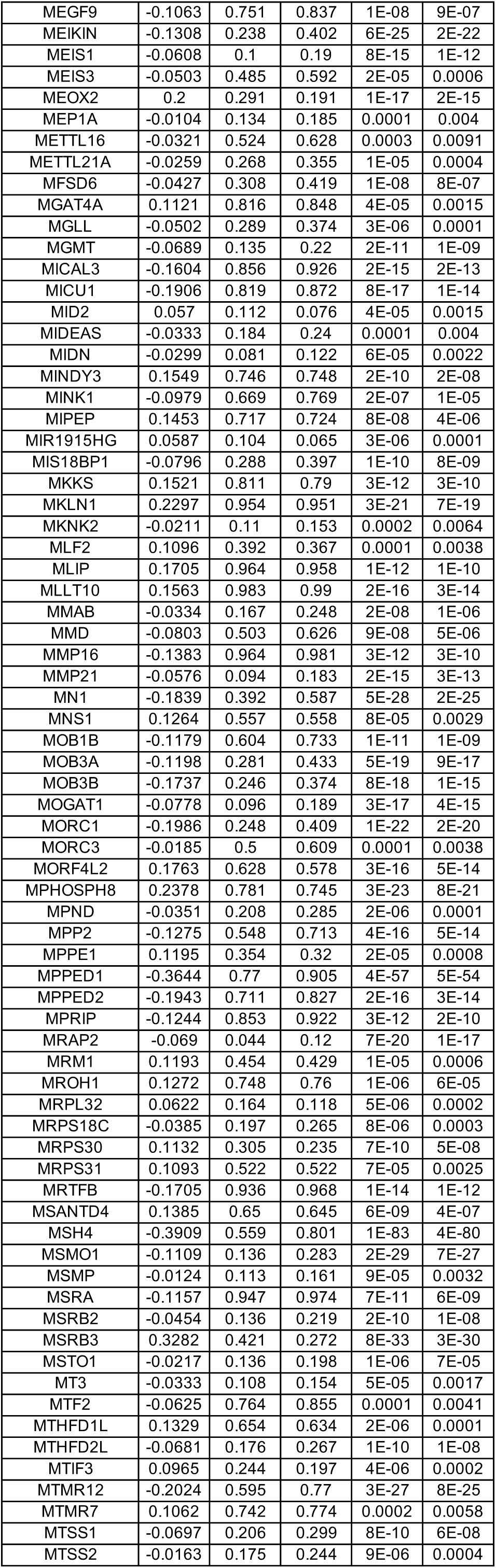
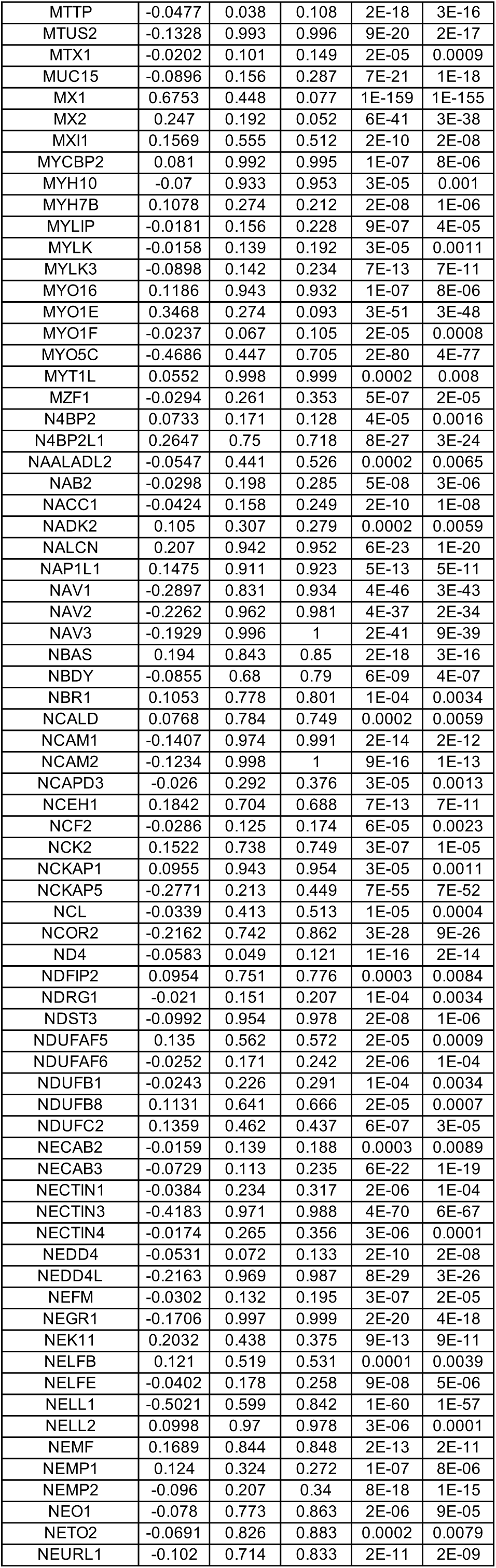
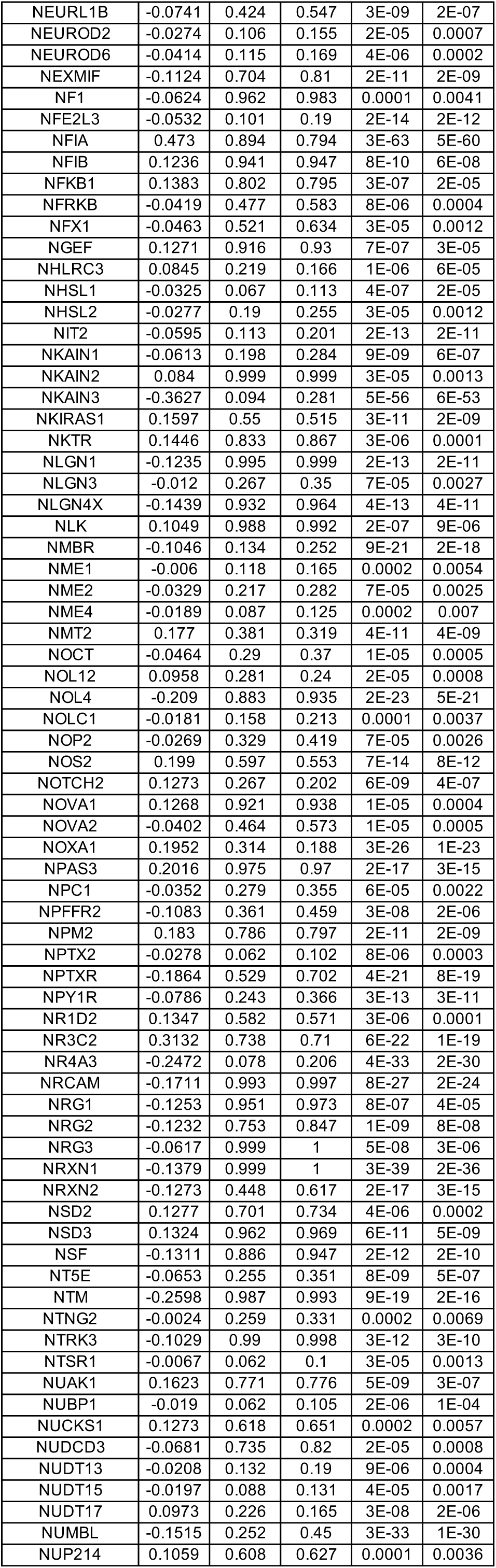
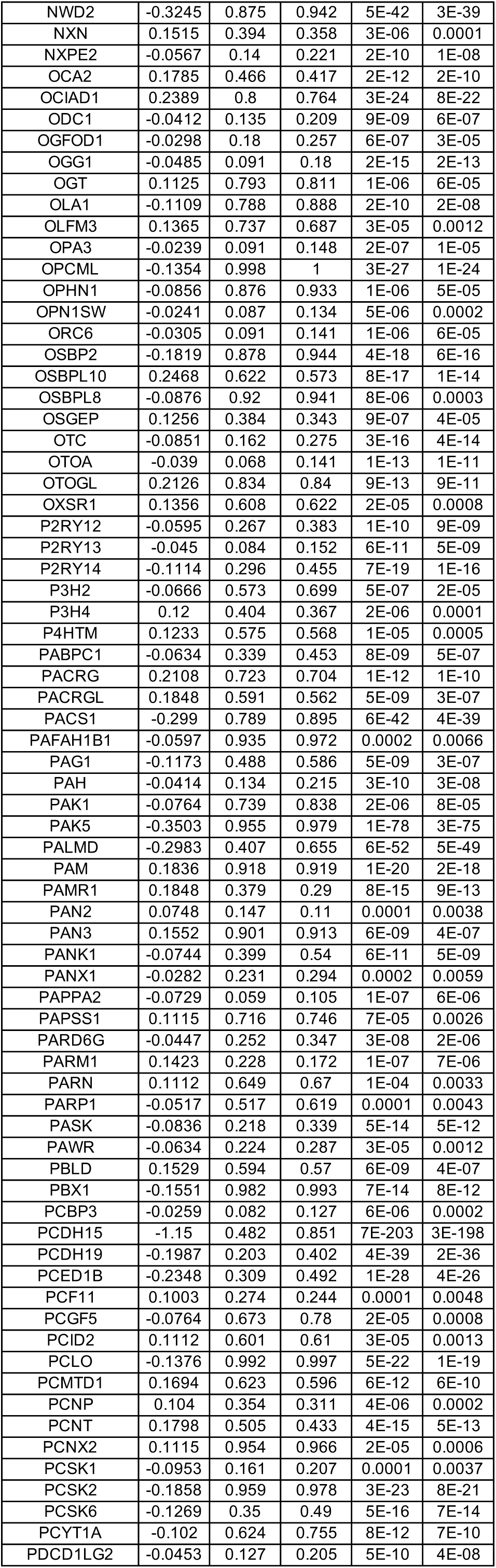
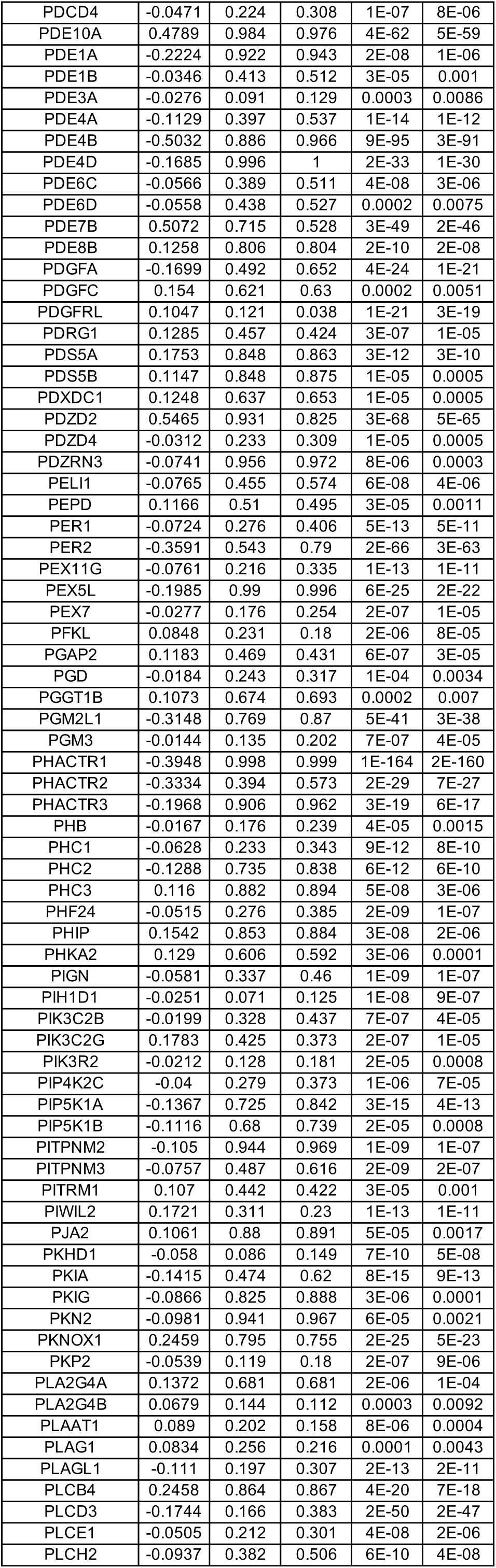
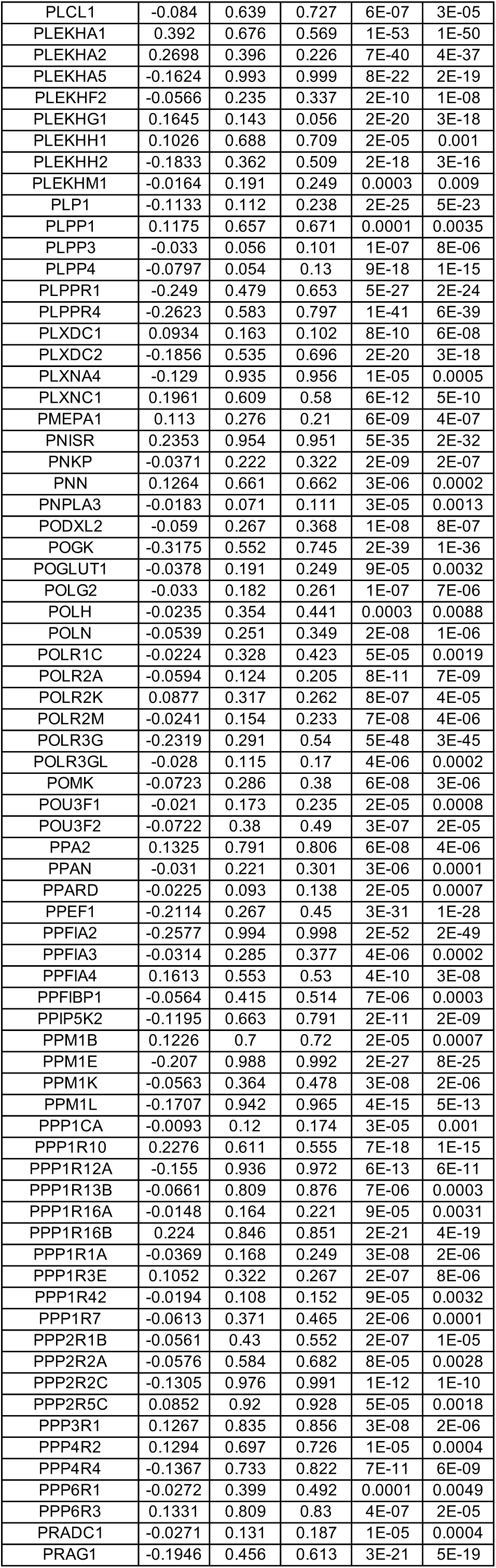
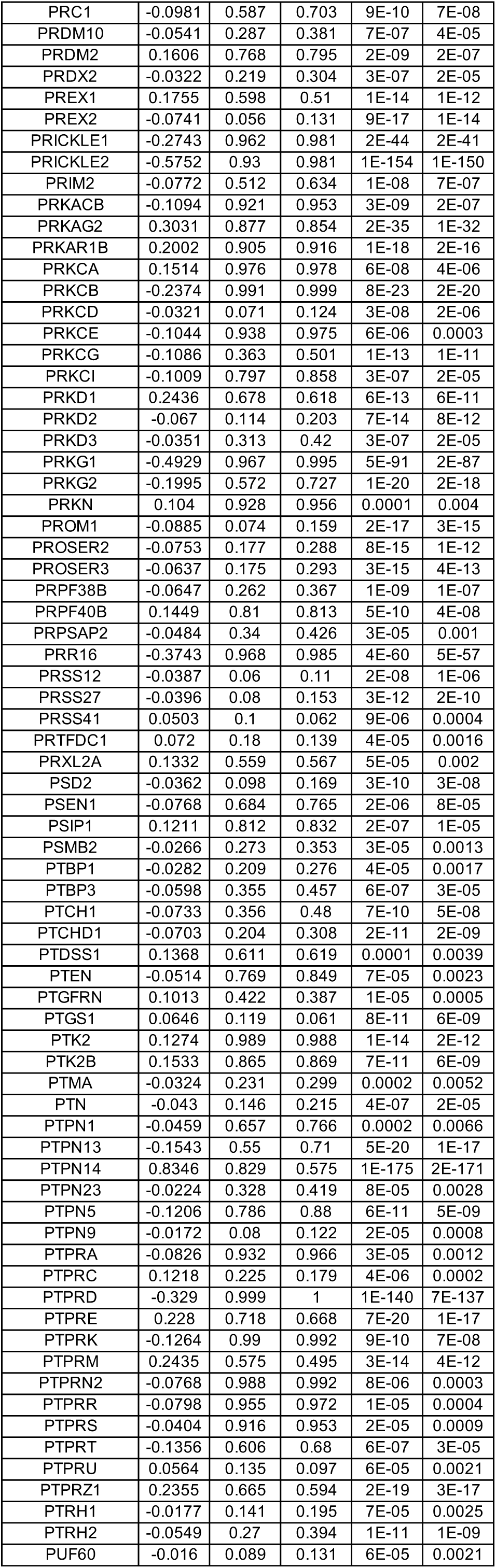
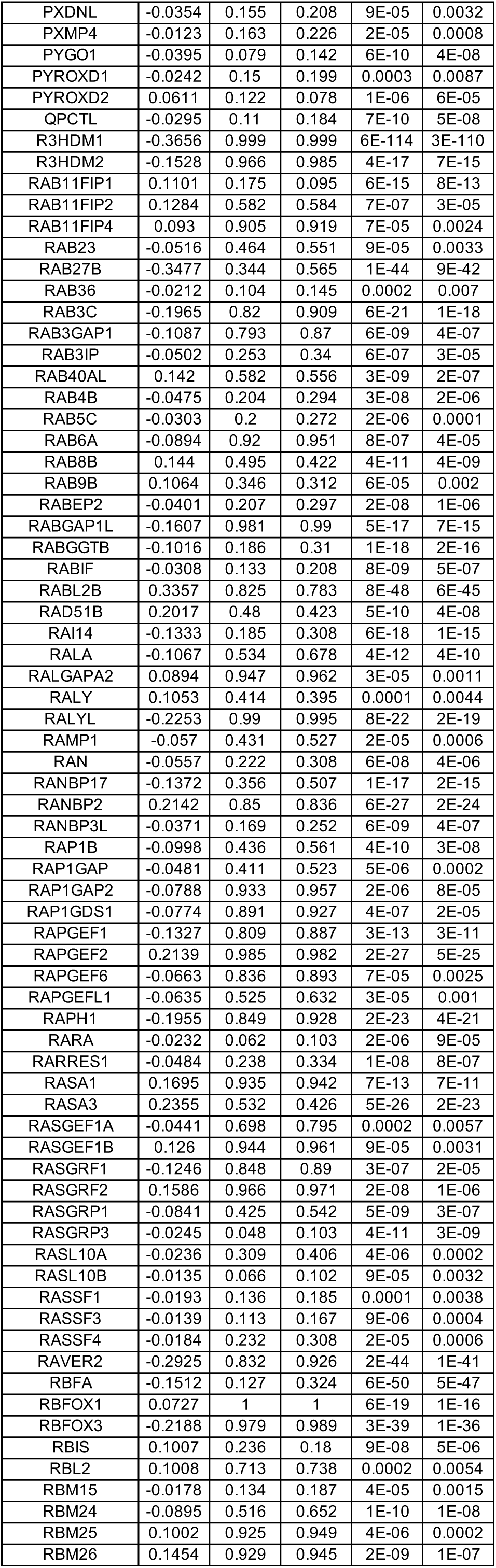
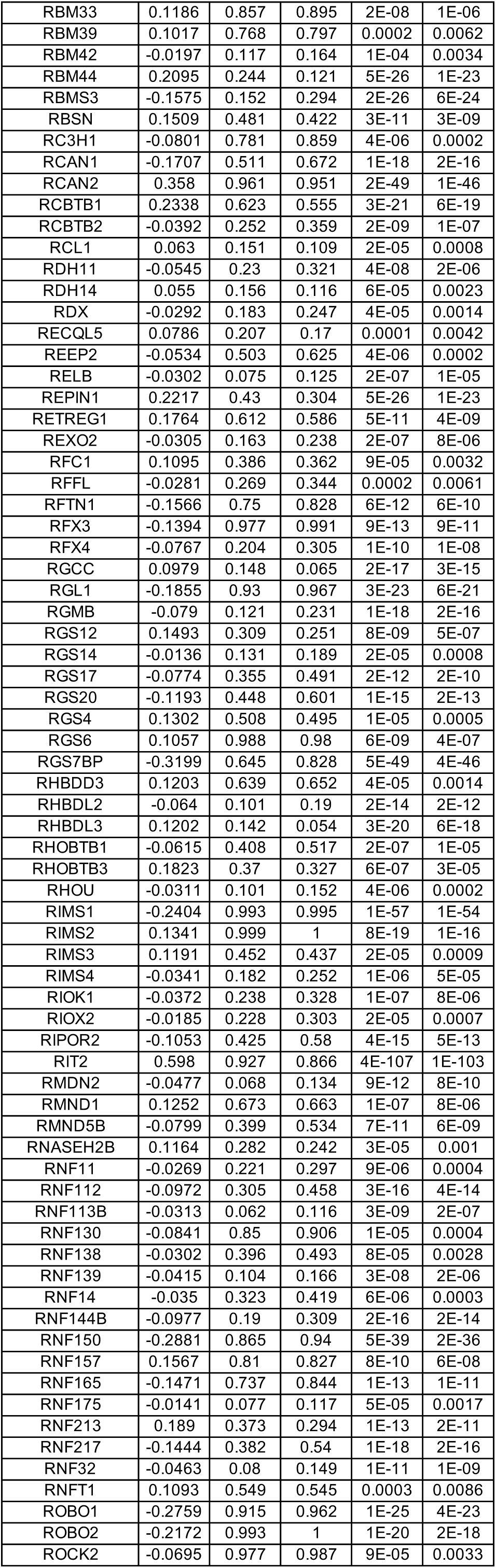
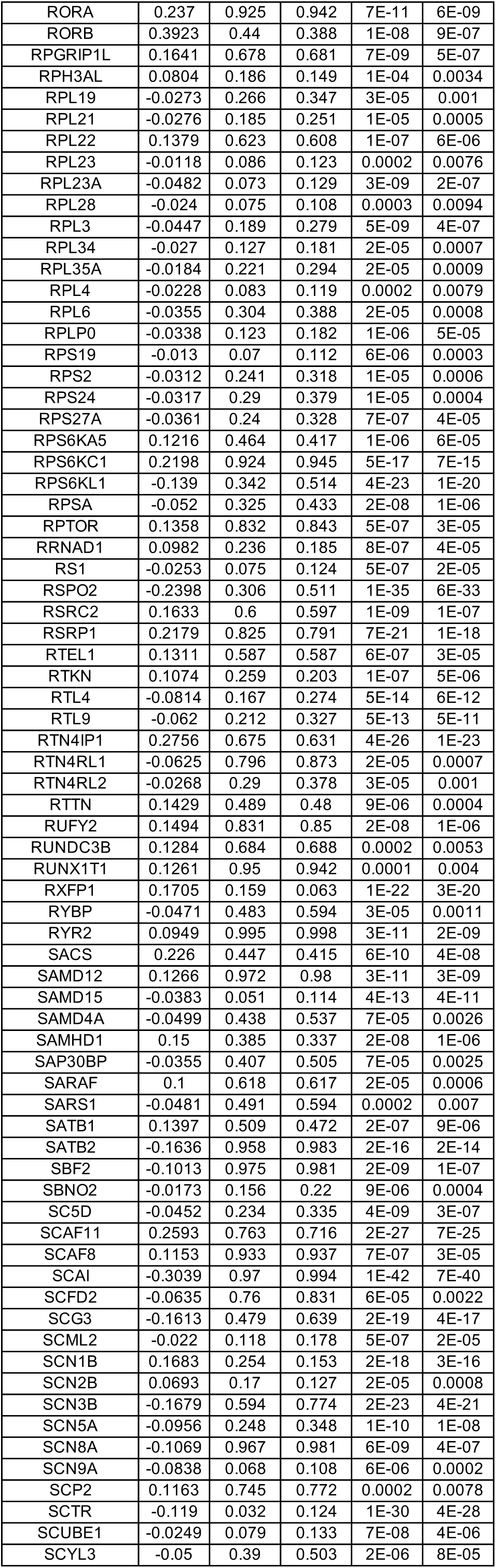
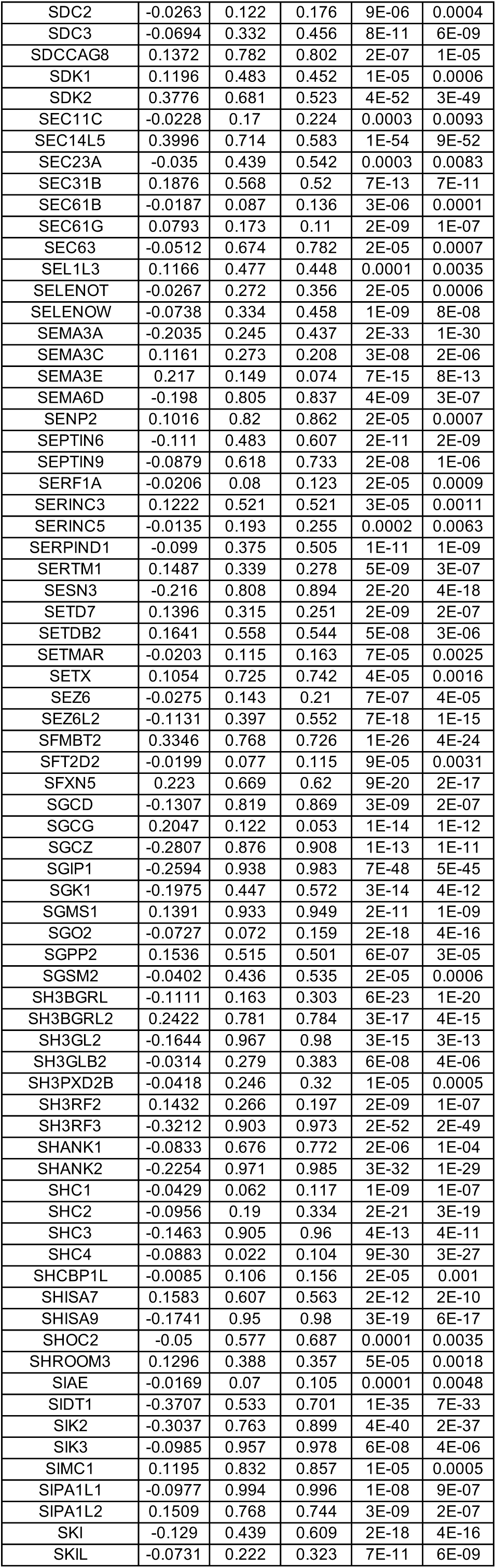
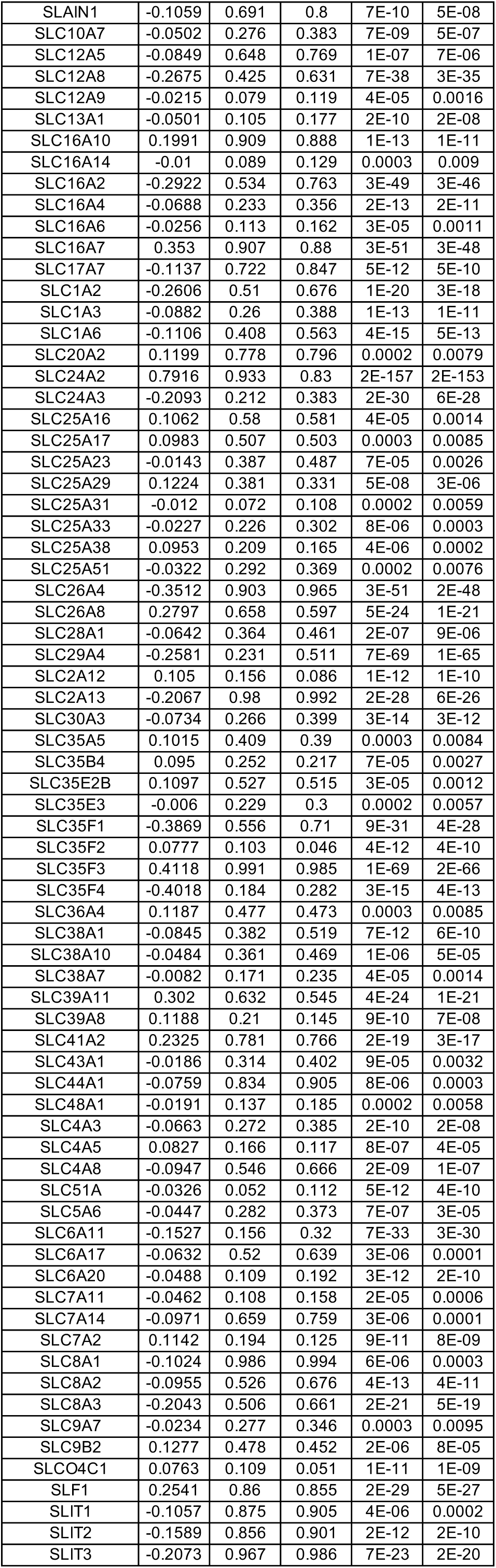
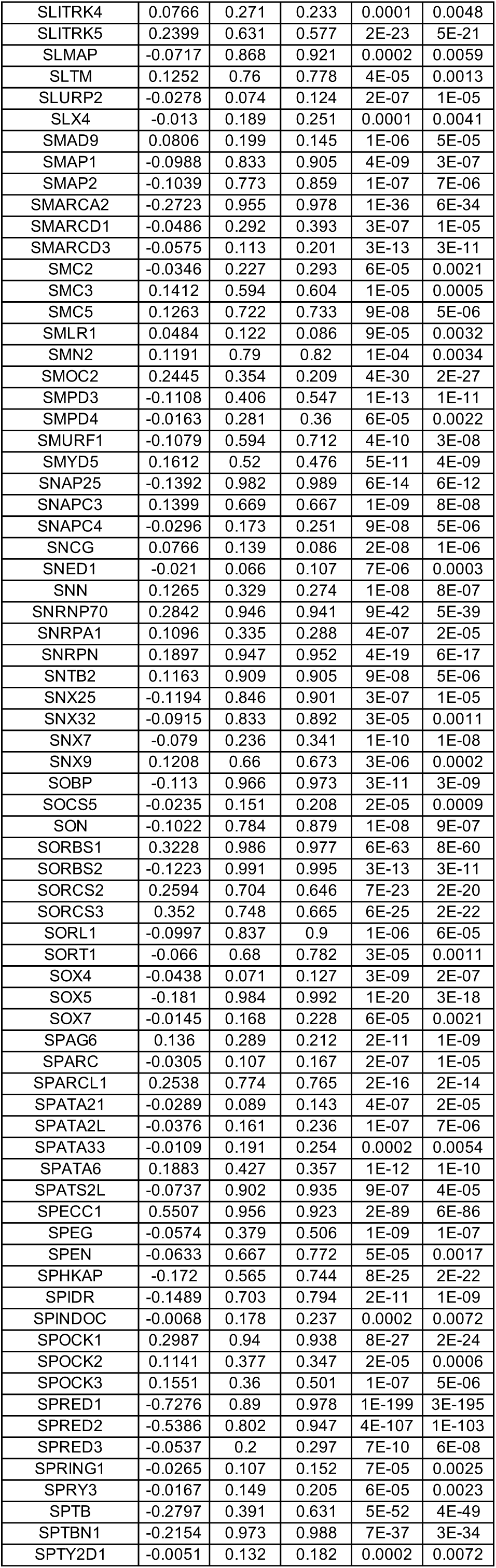
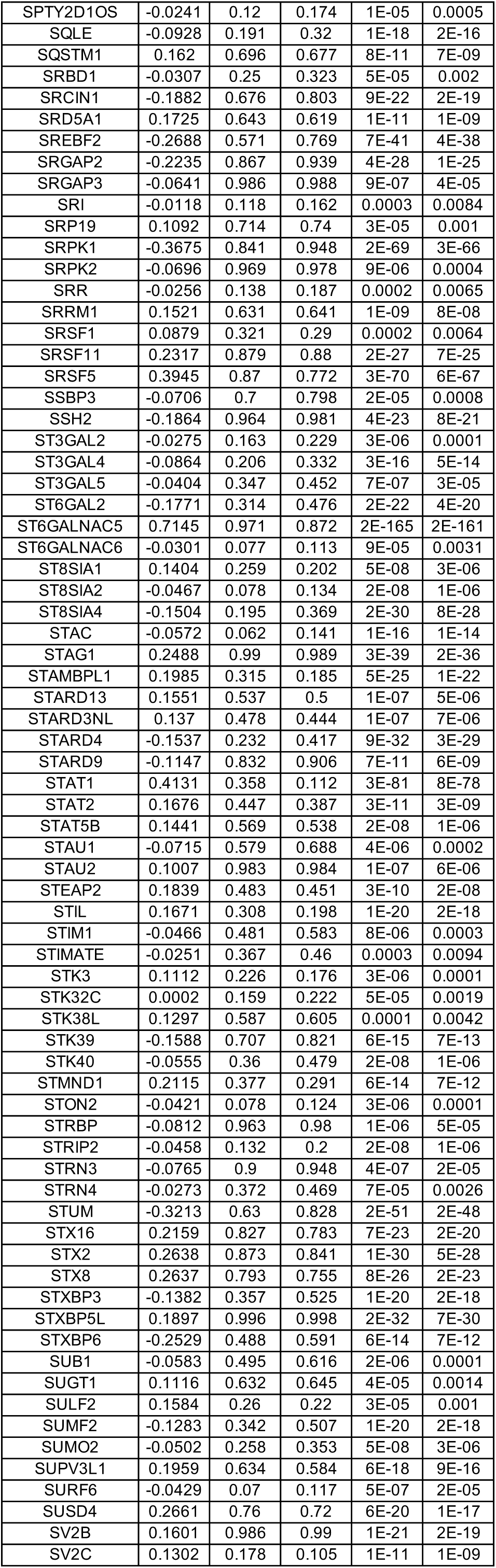
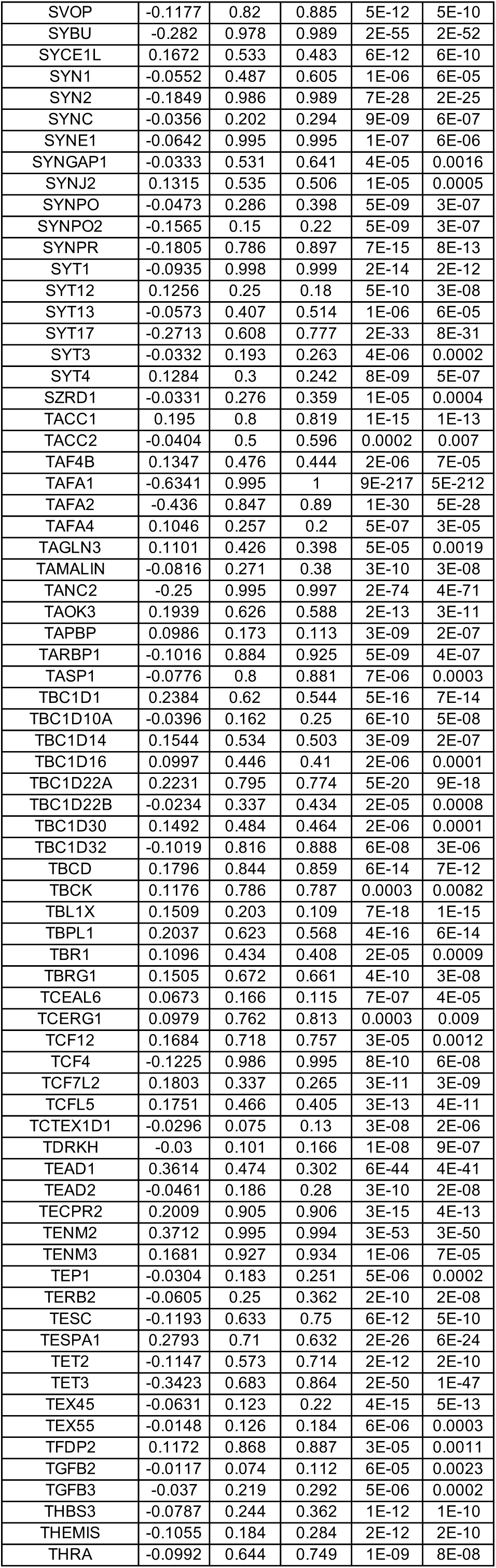
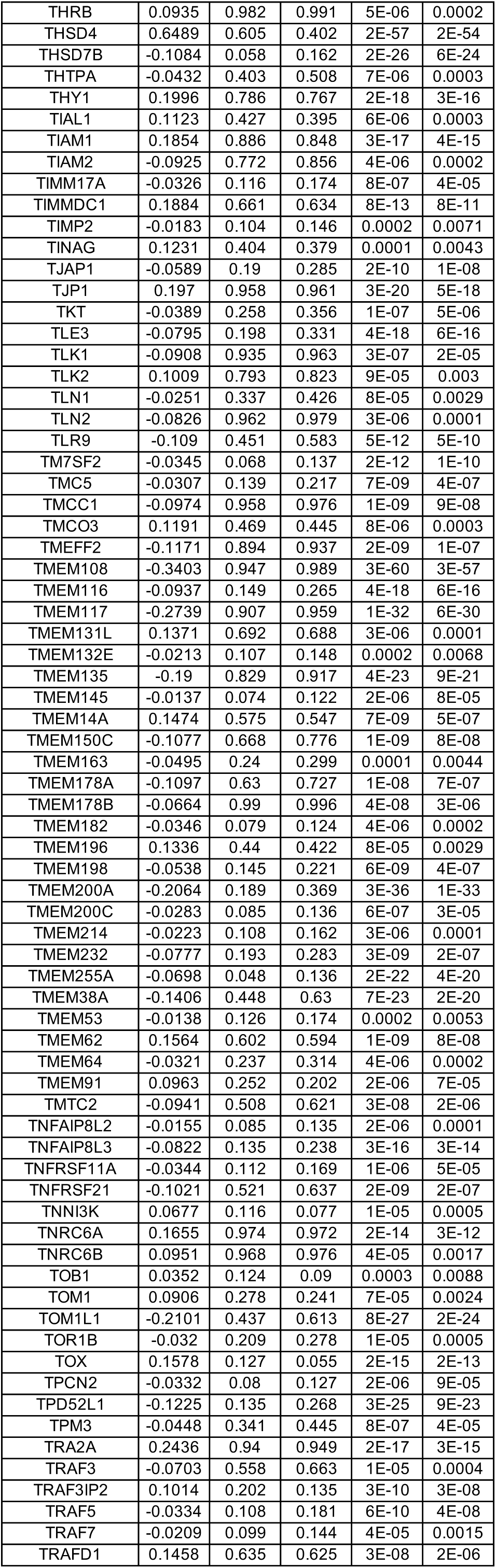
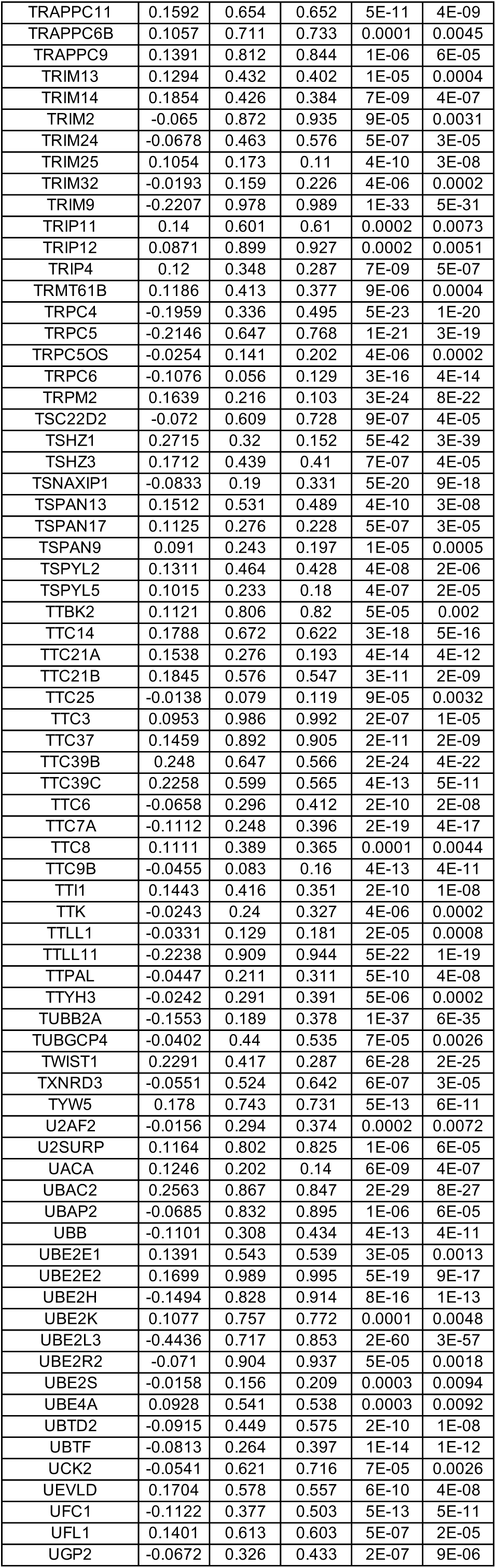
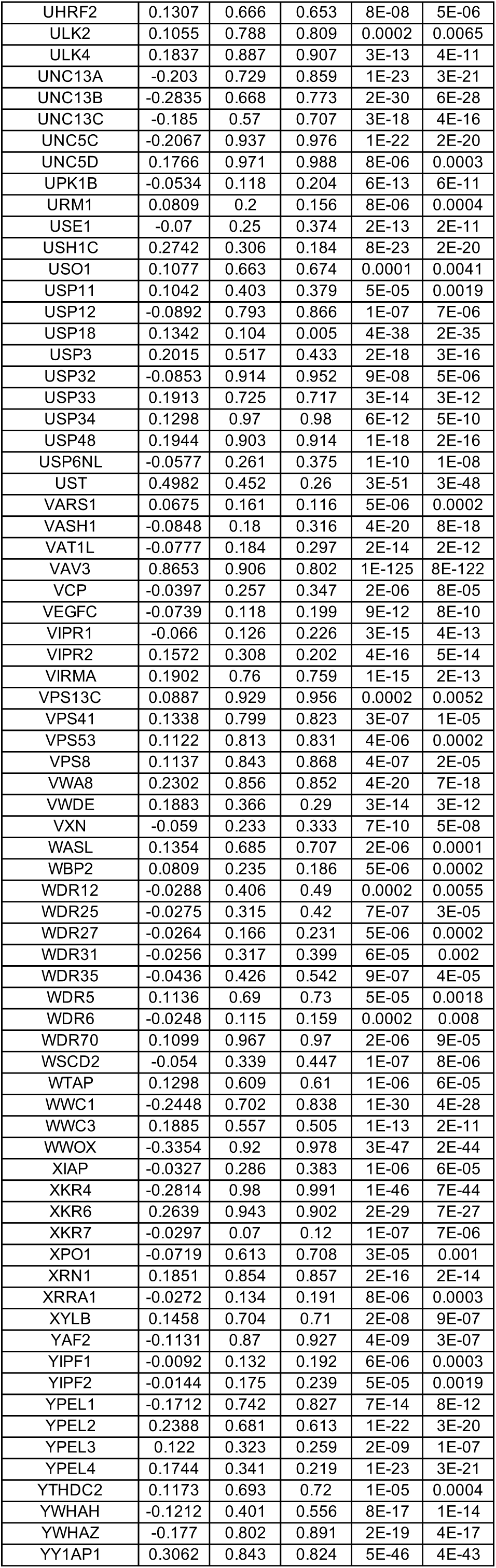
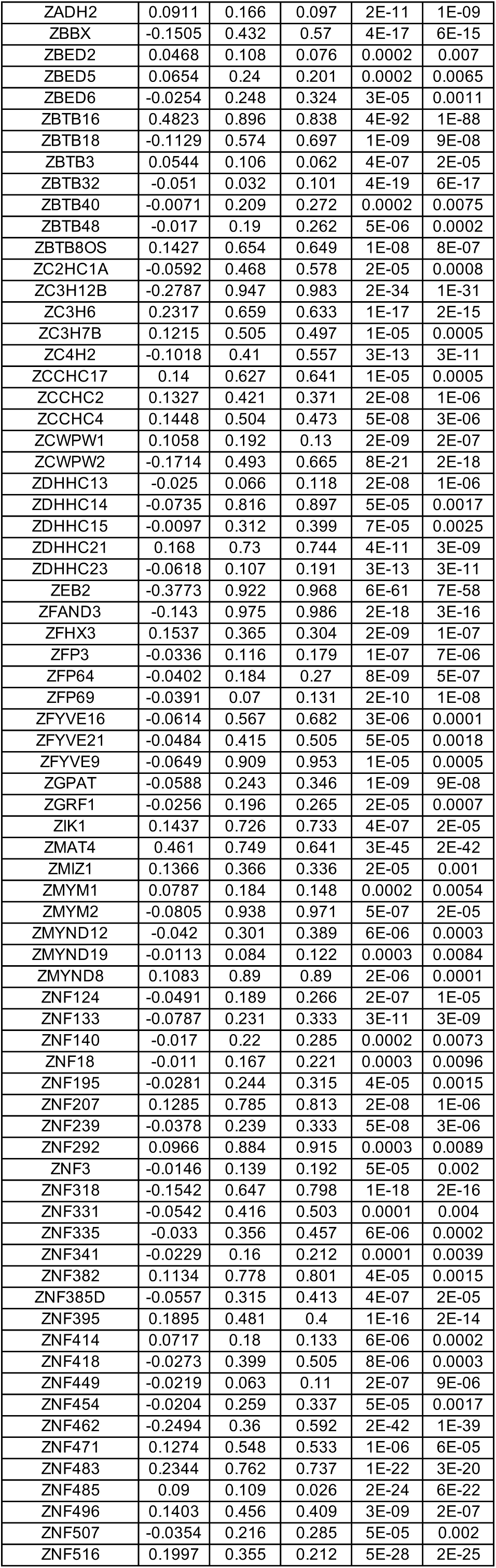
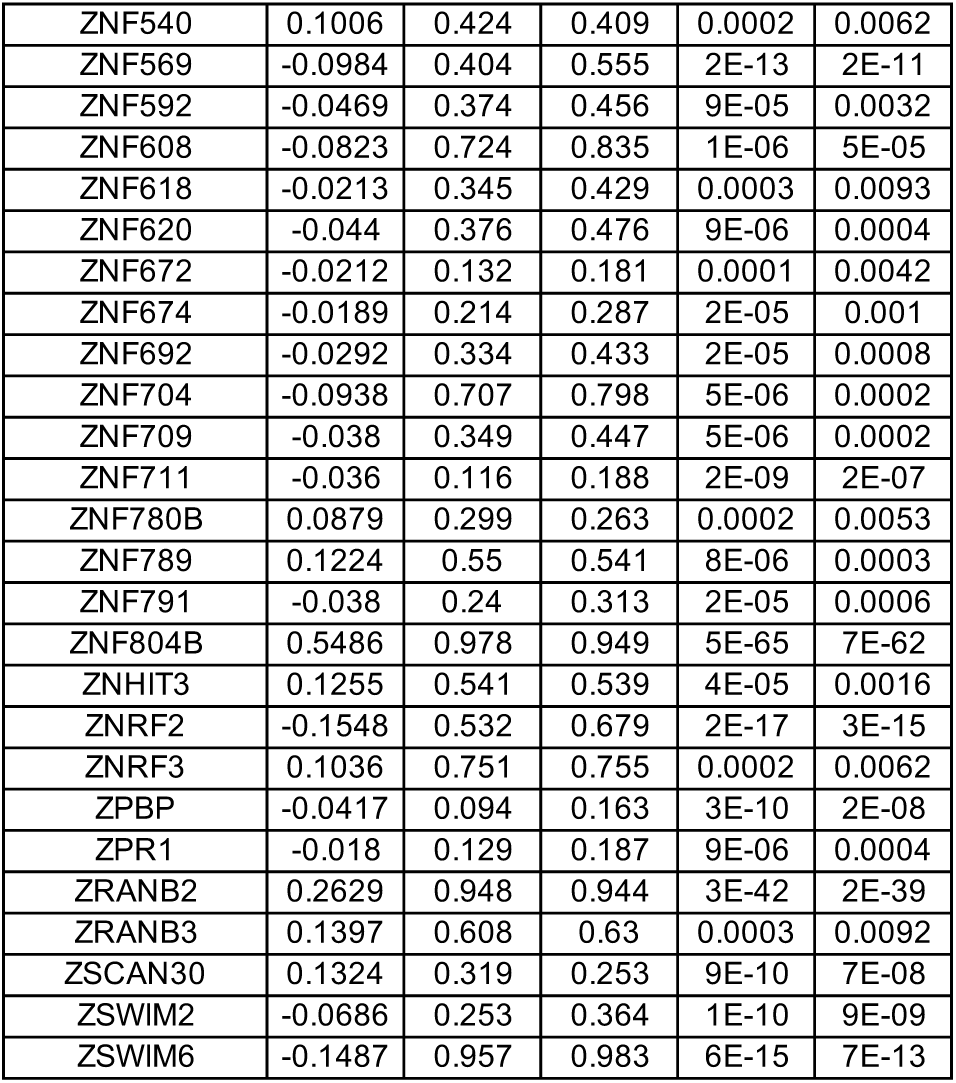
DEGs between wild-type (WT) and MECP2-null (KO) upper layer excitatory neurons (Ex_1) of PFC (adjusted p-value < 0.01)

**Table S16.**
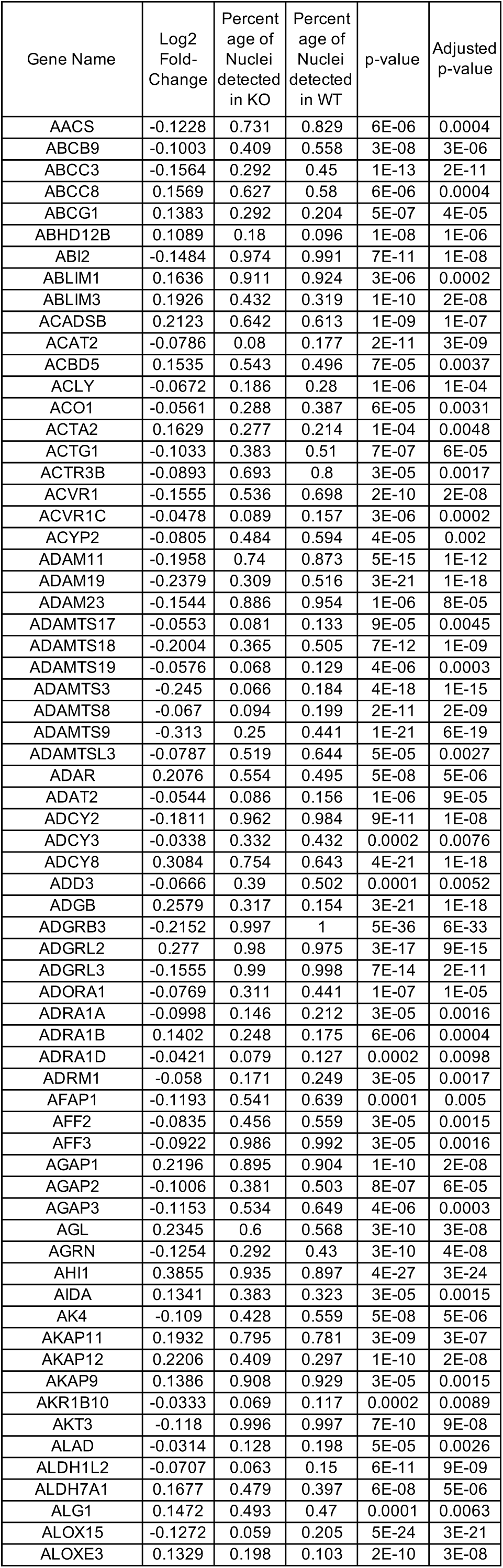
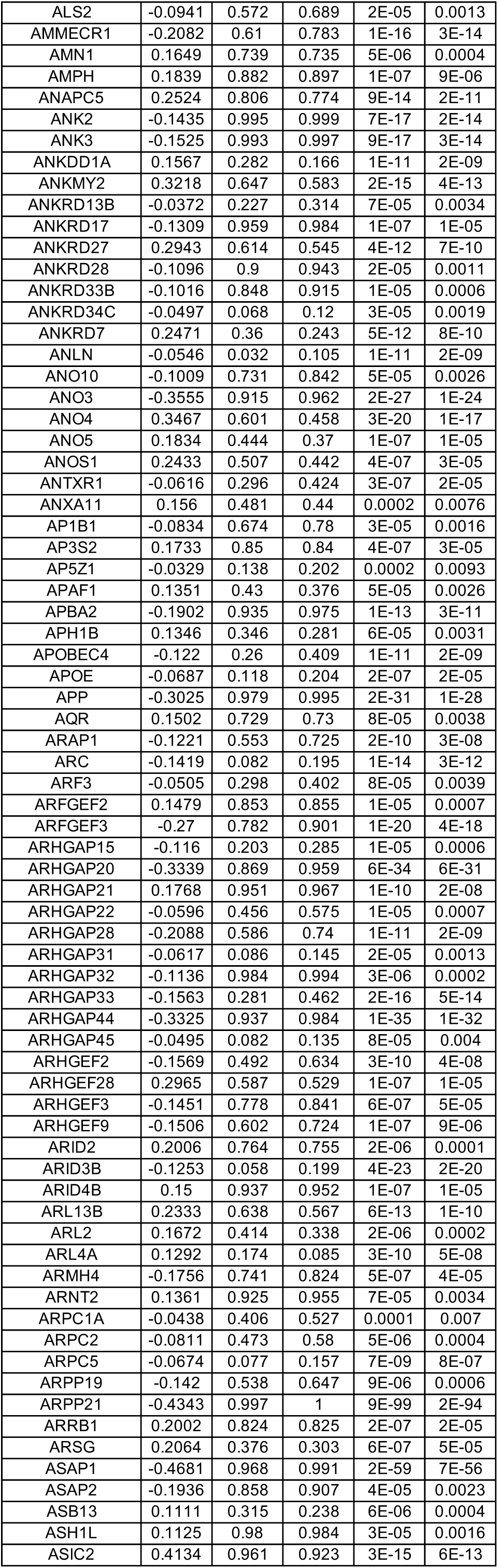
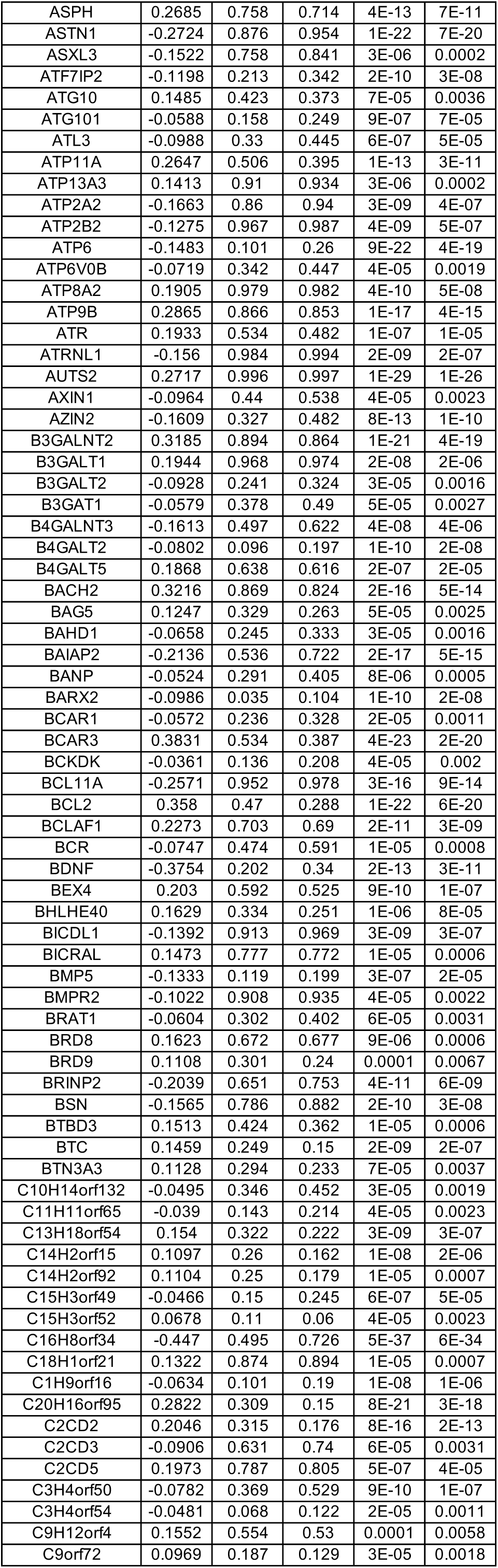
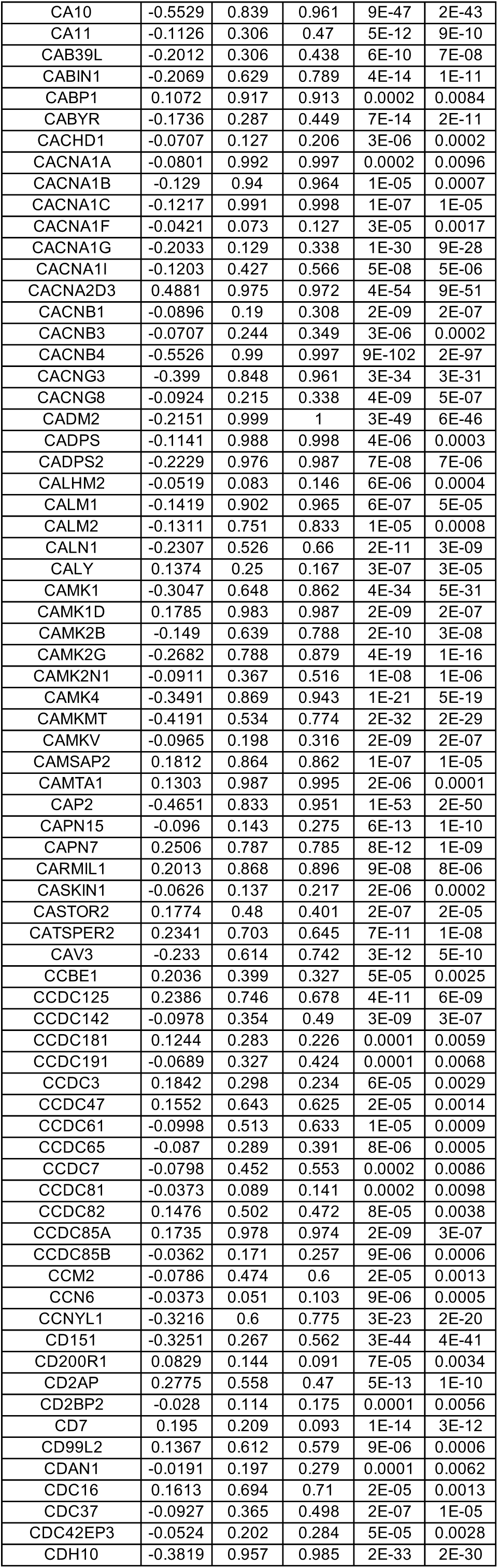
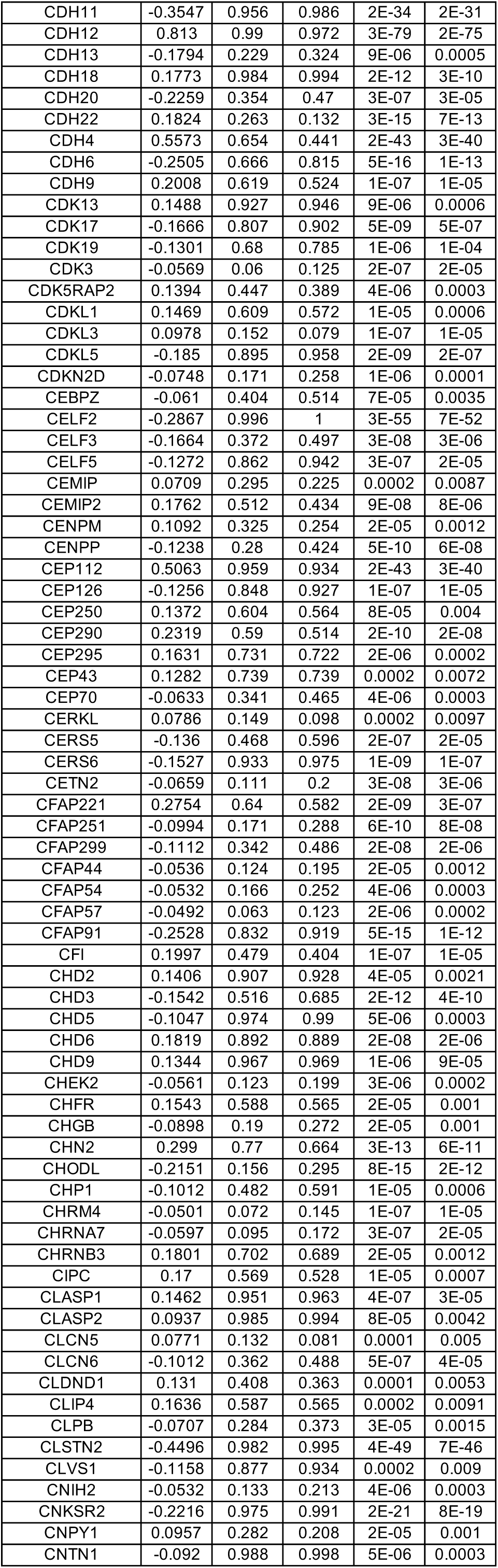
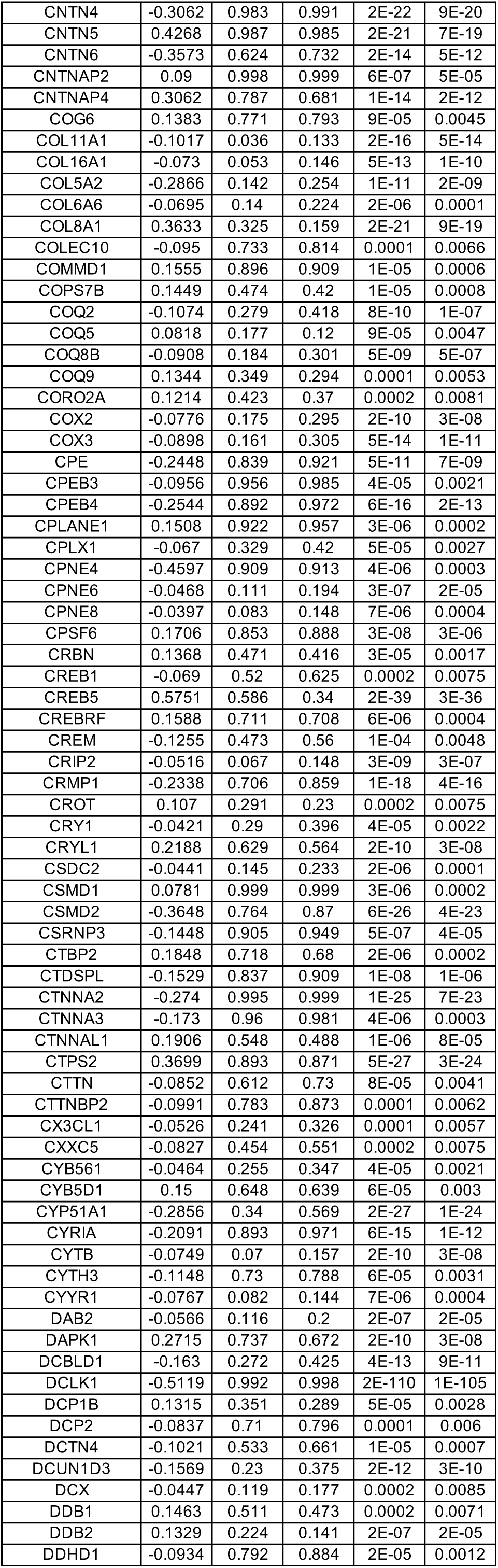
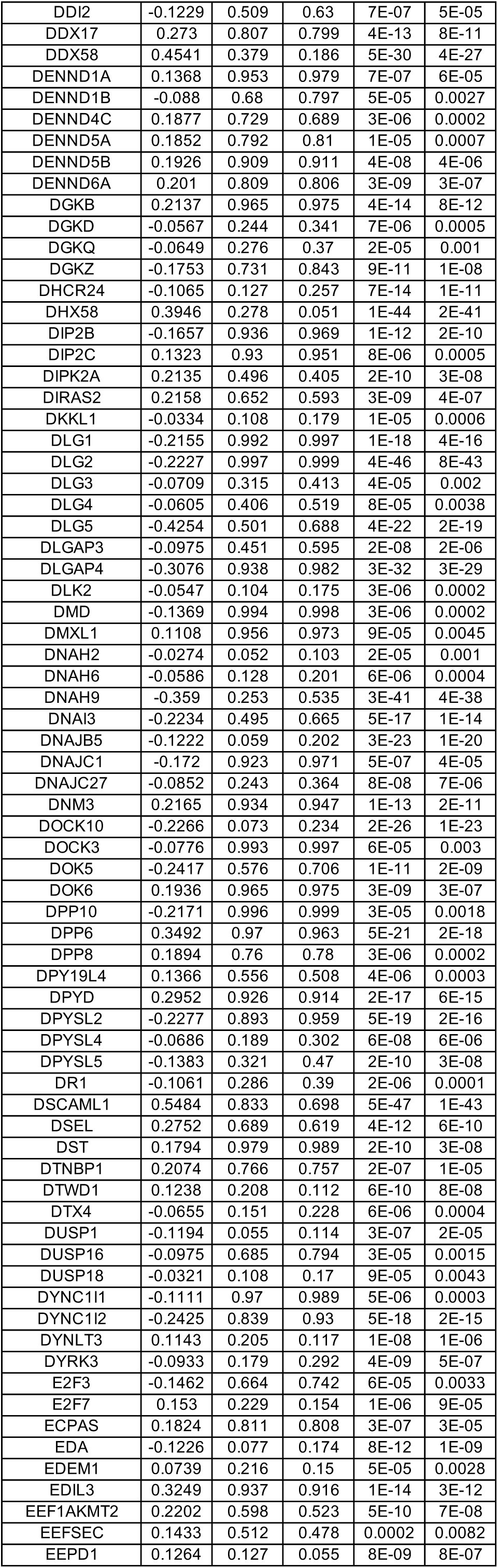
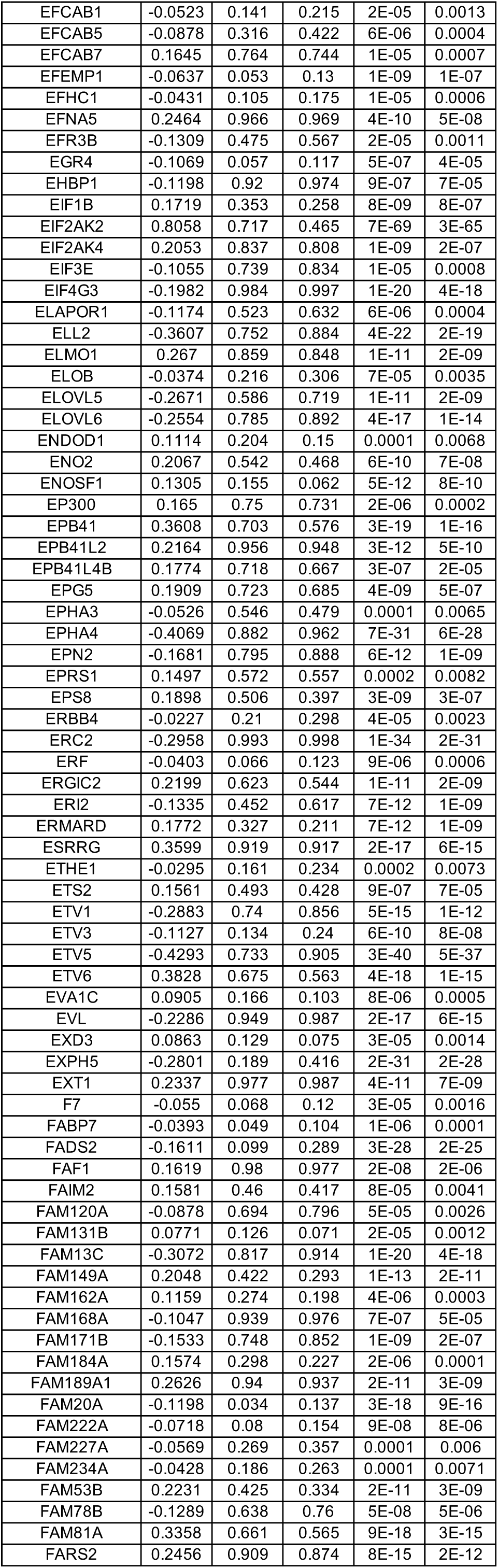
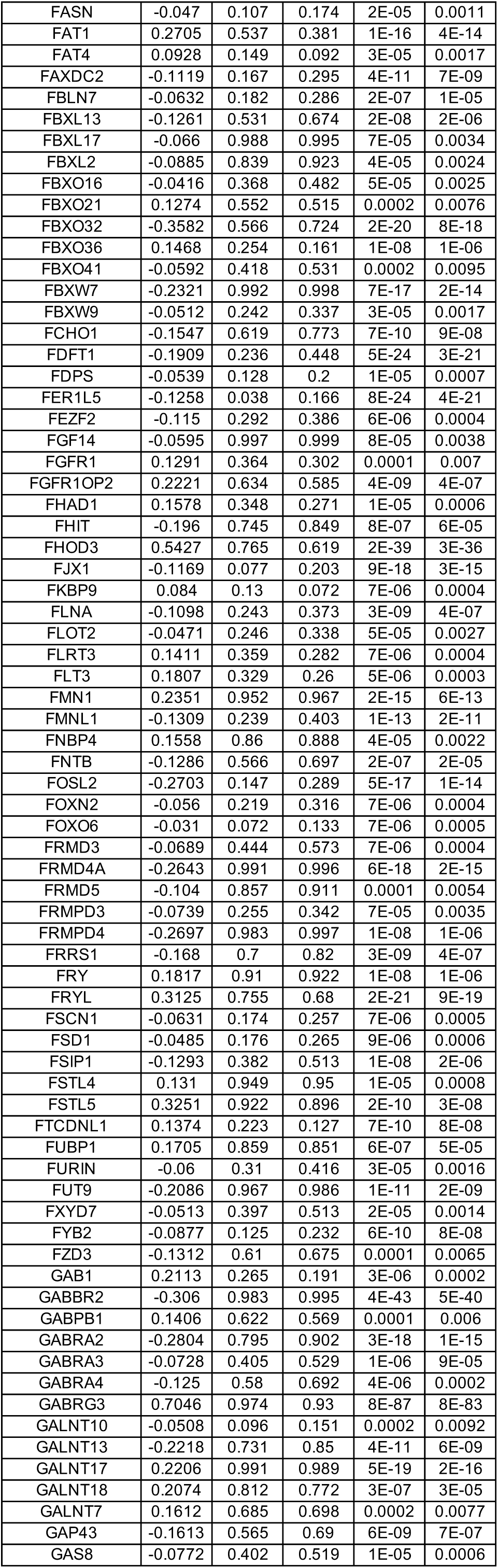
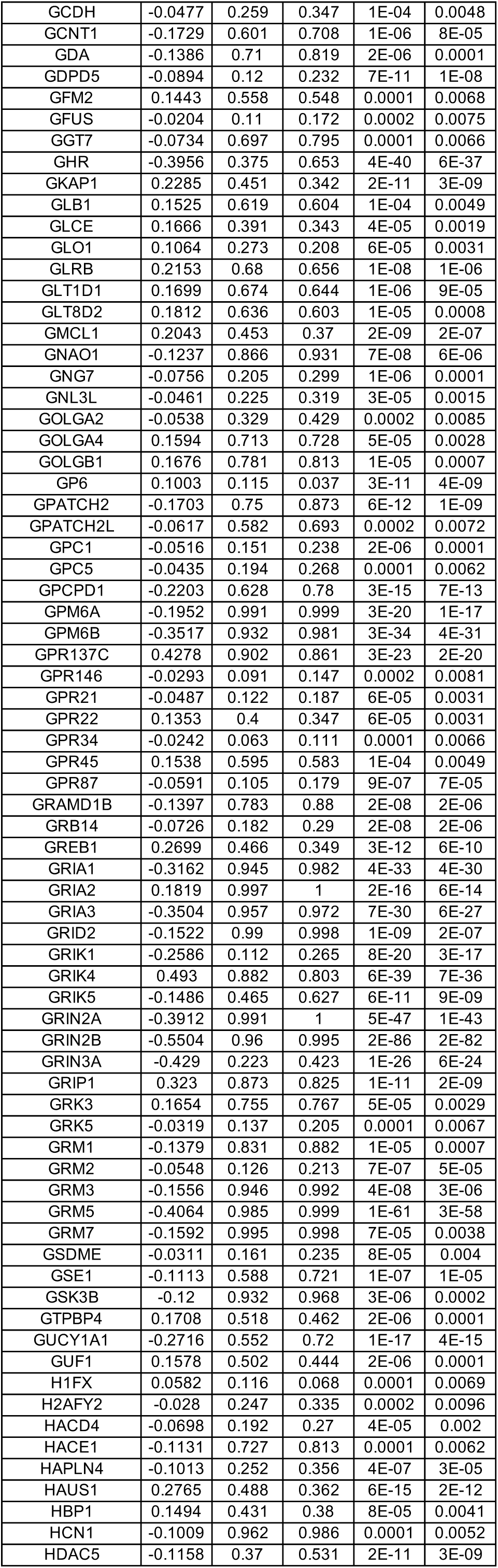
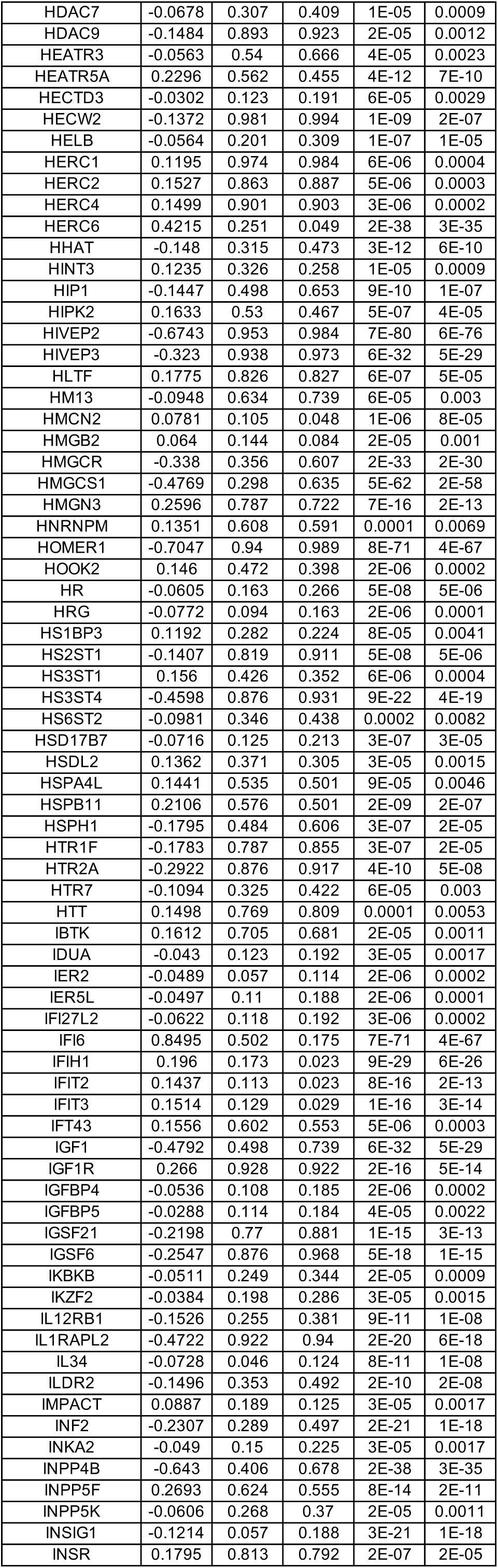
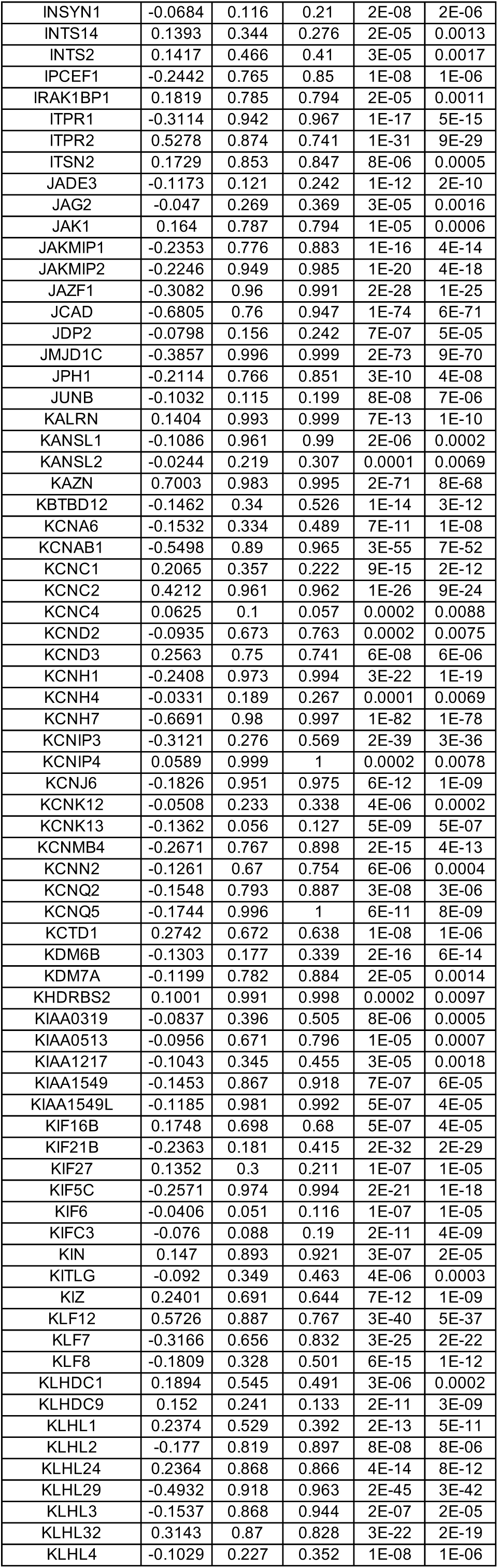
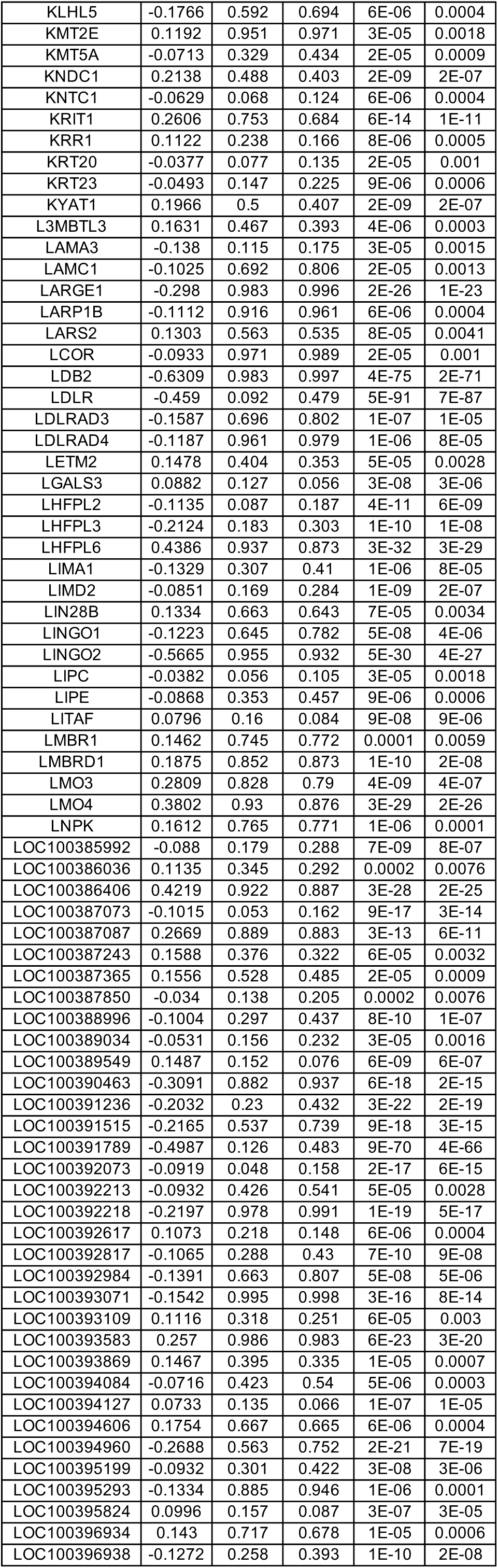
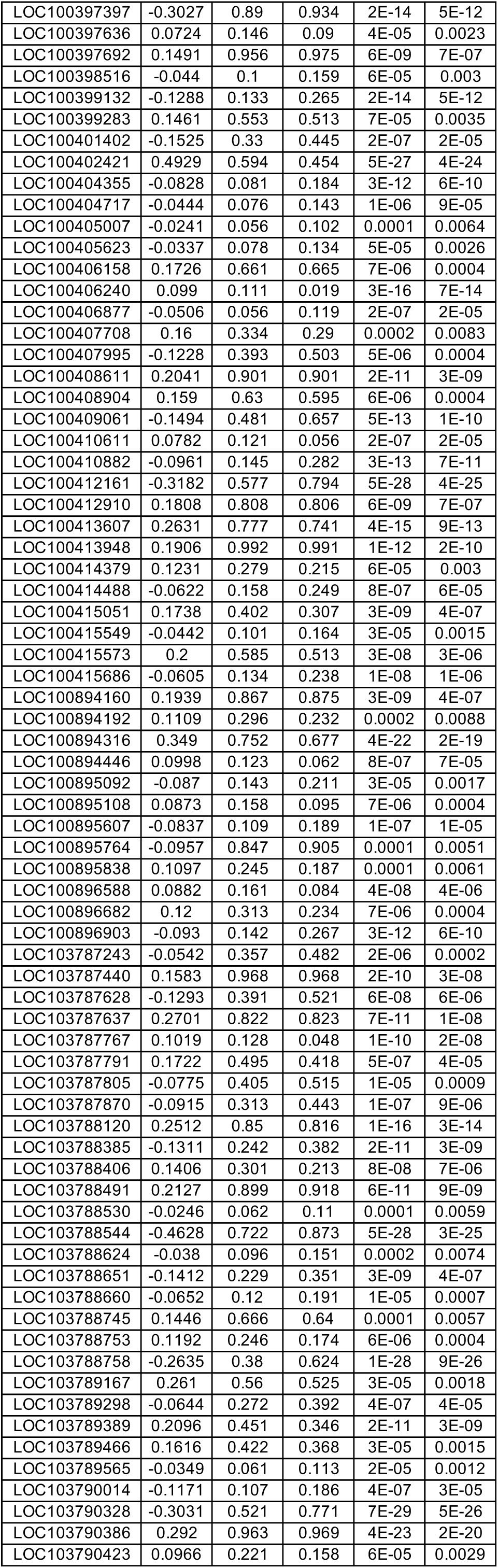
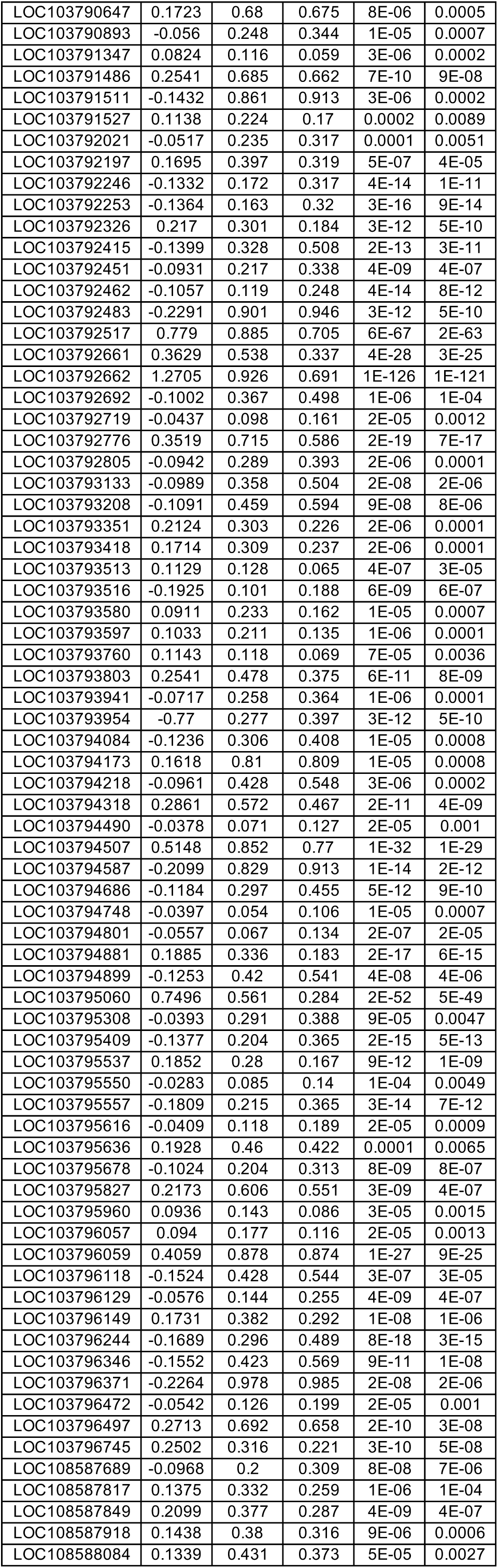
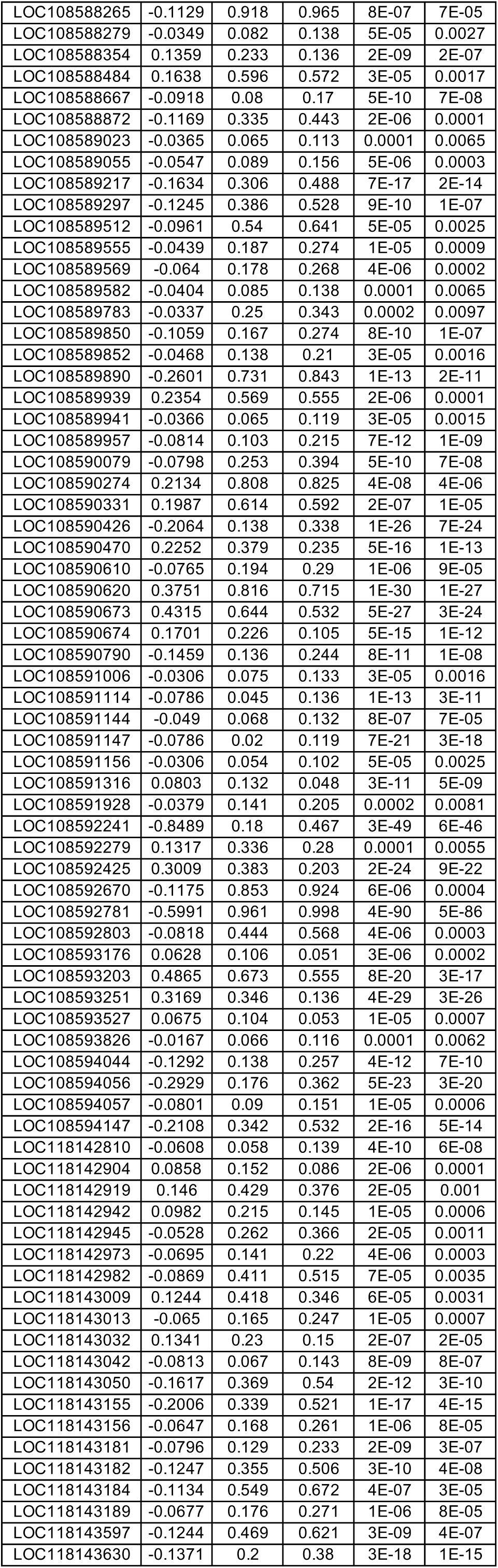
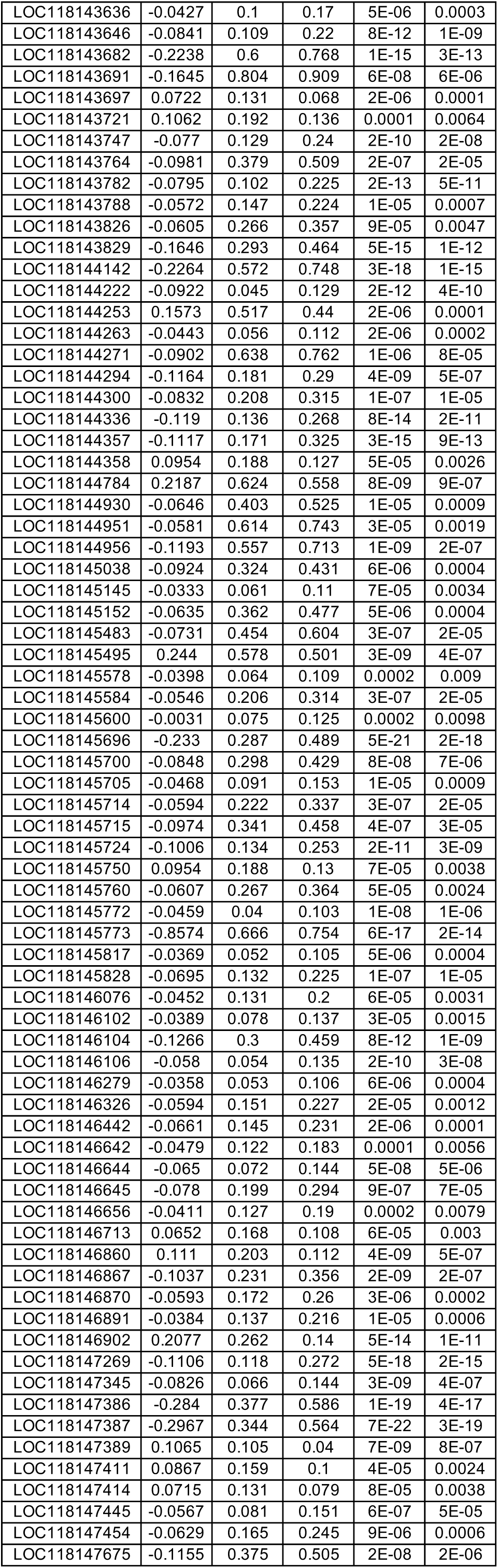
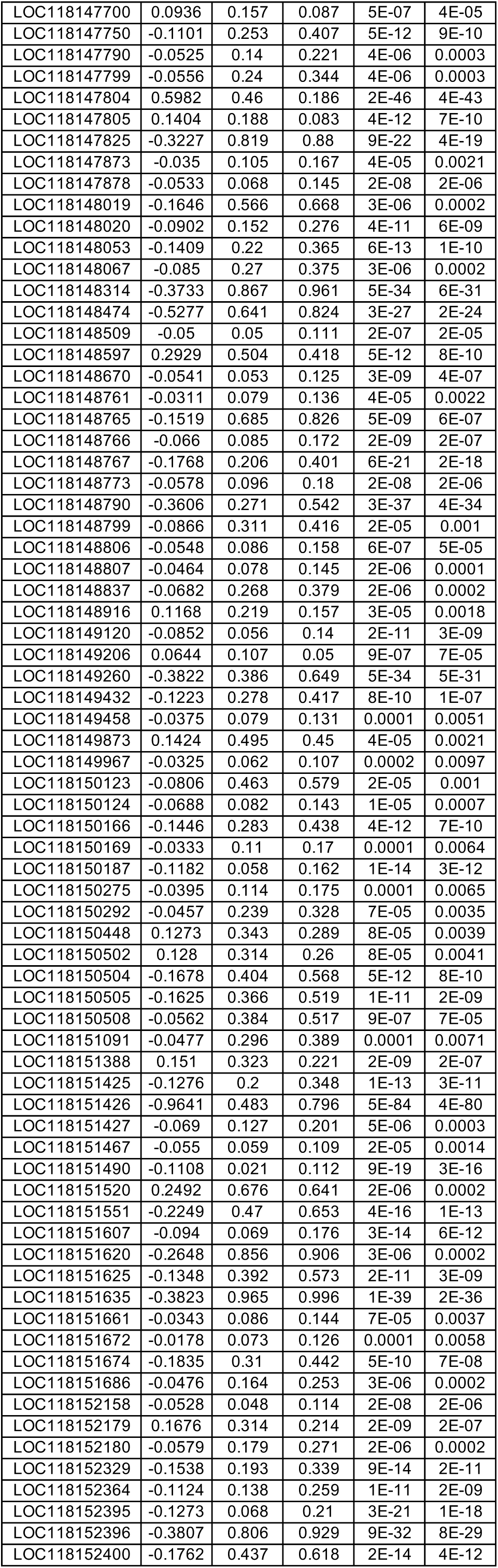
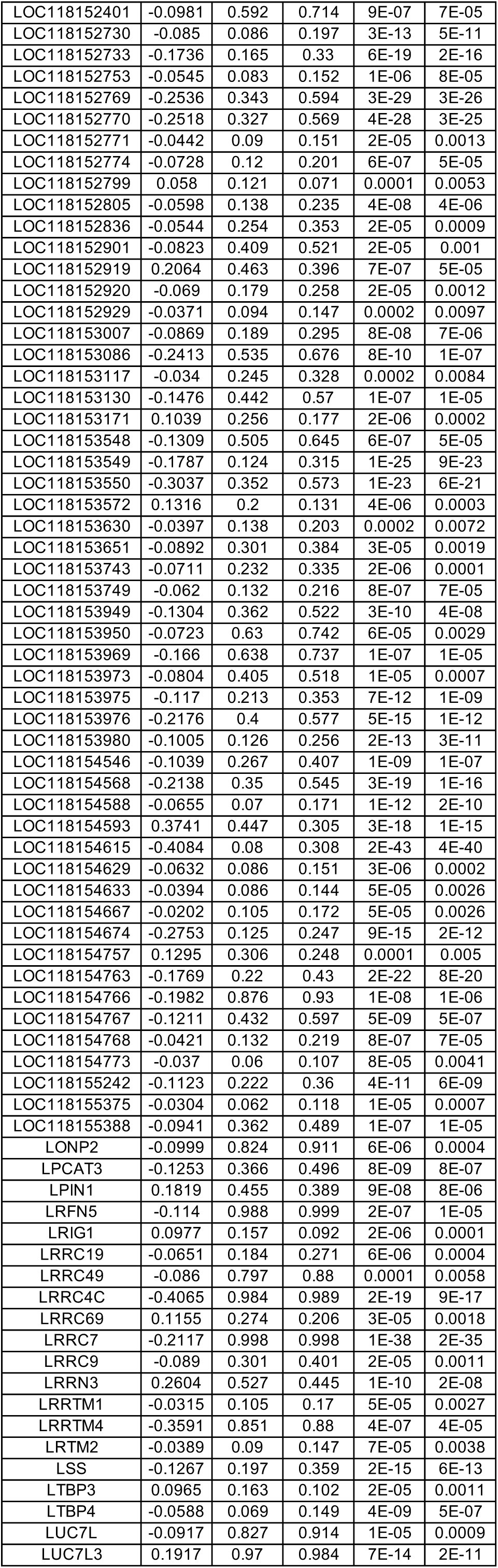
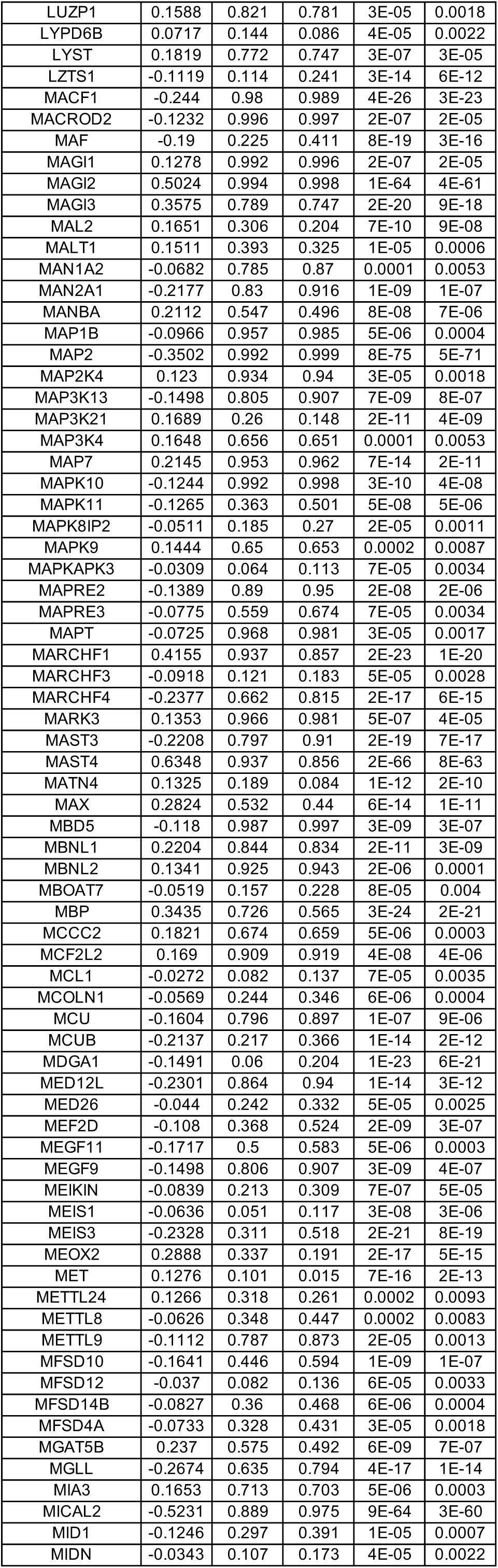
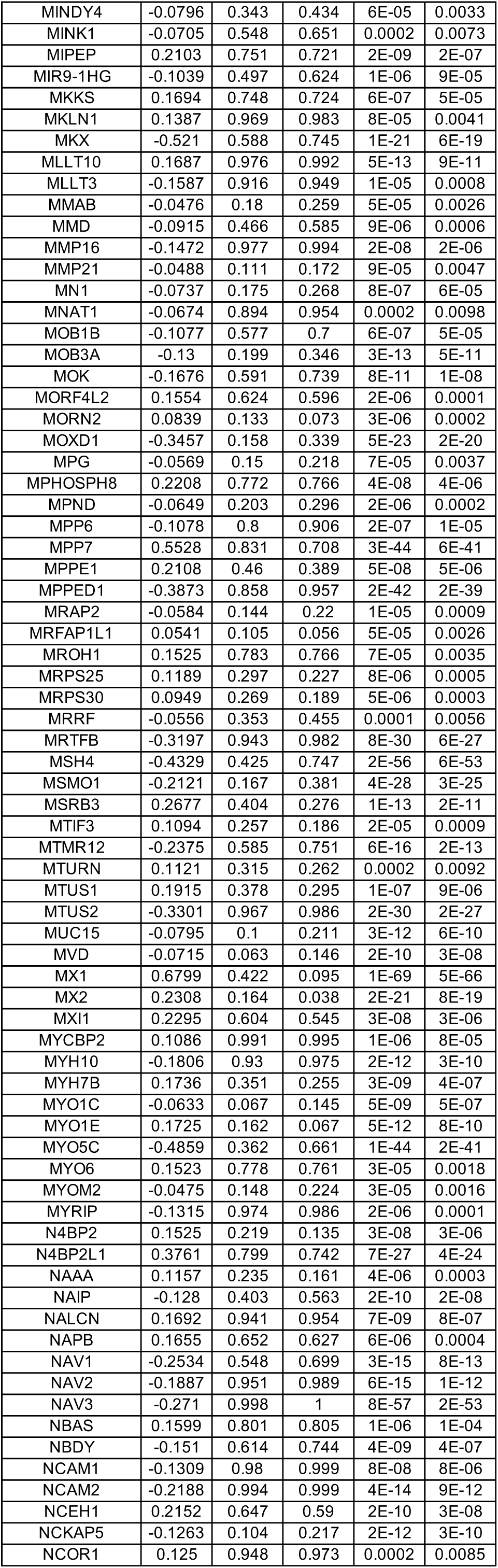
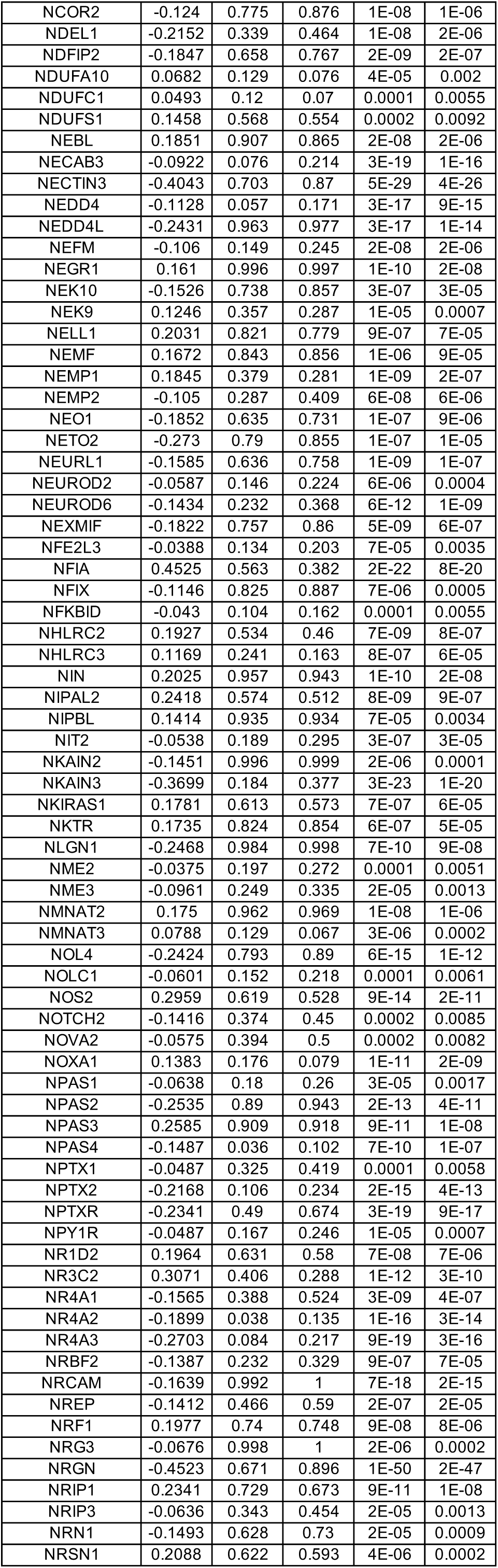
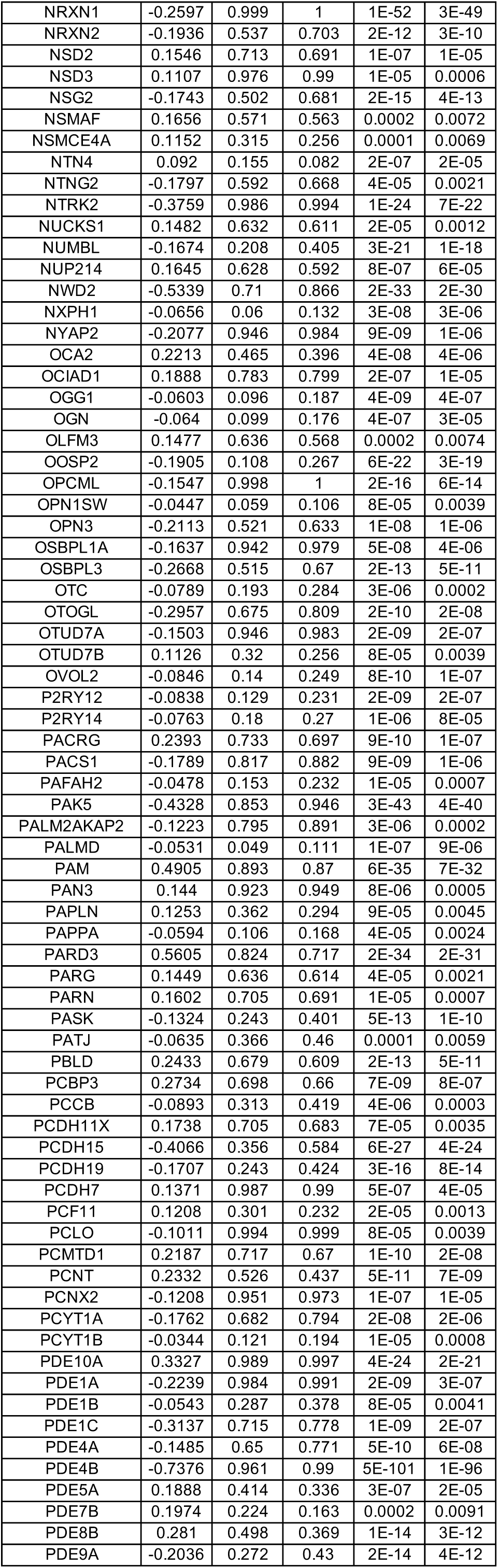
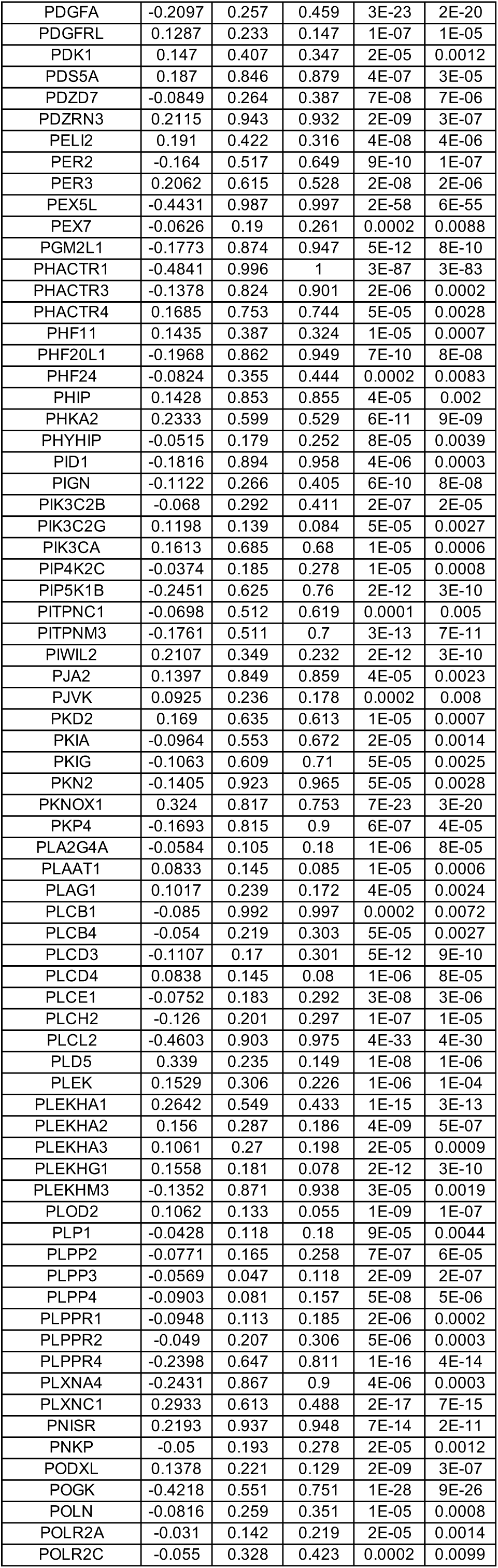
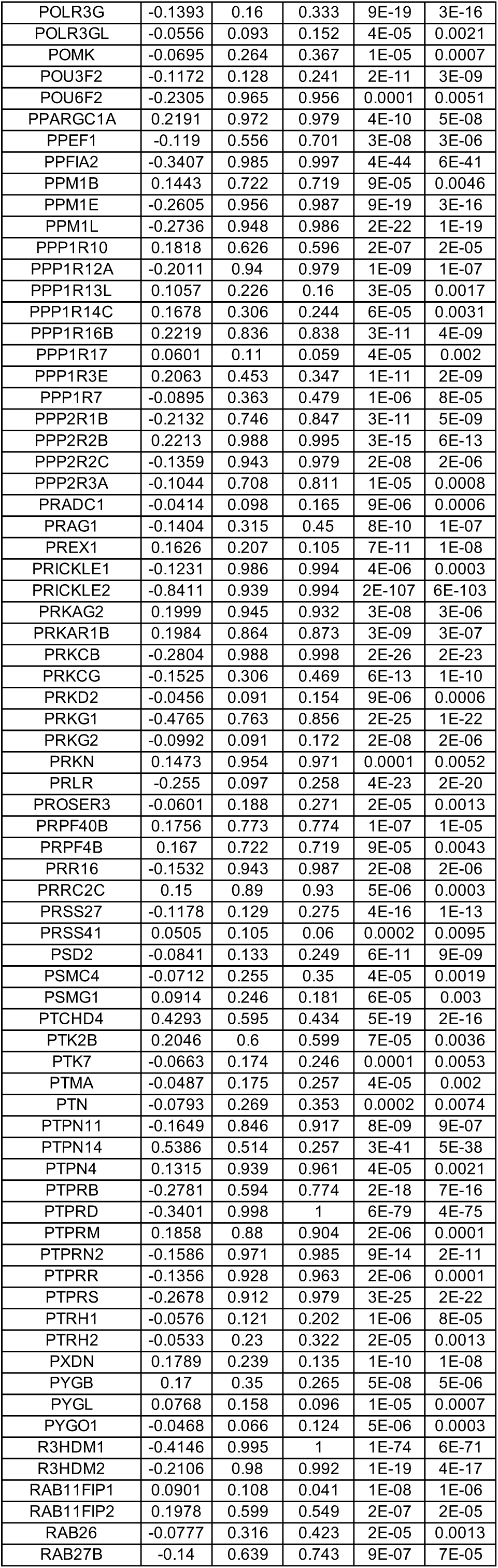
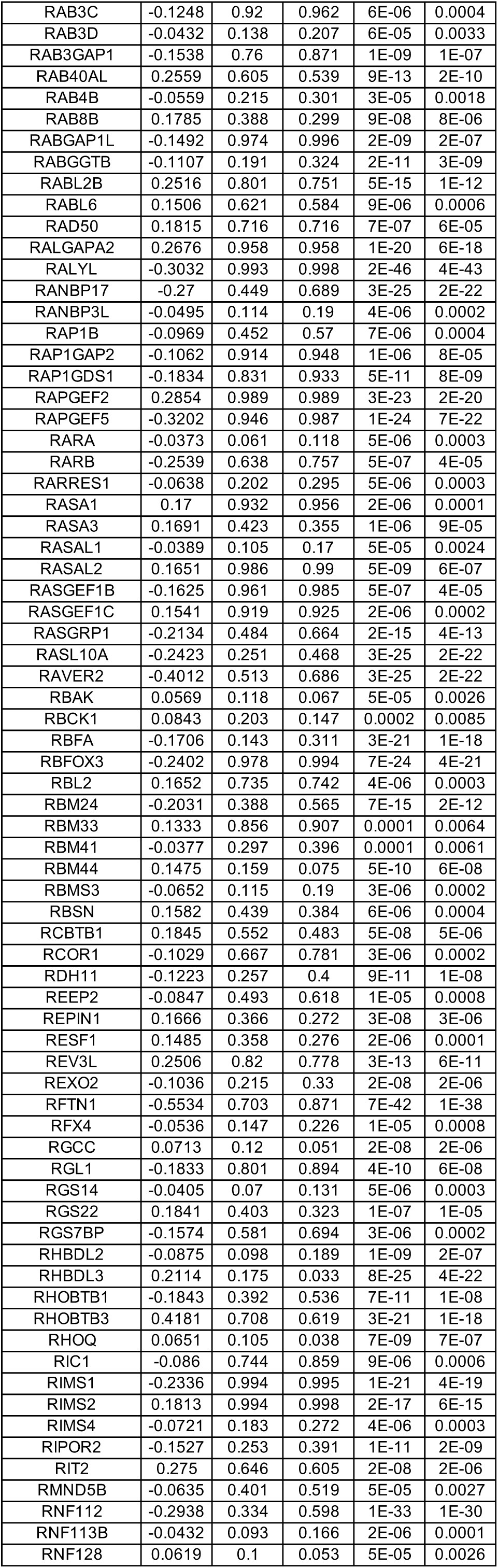
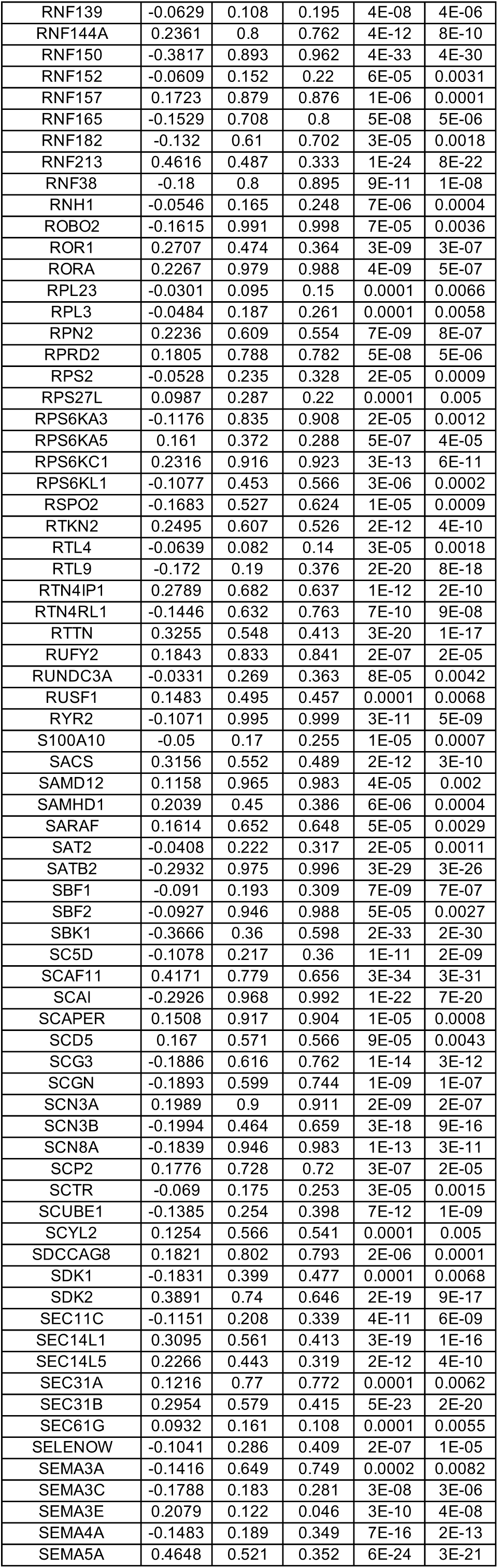
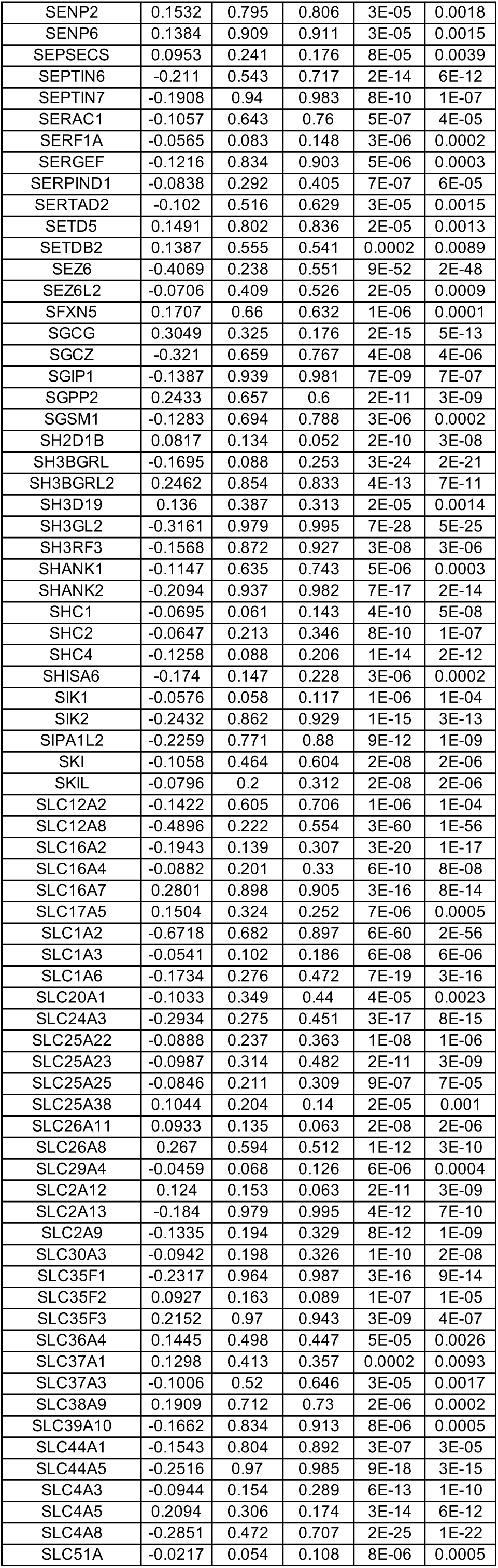
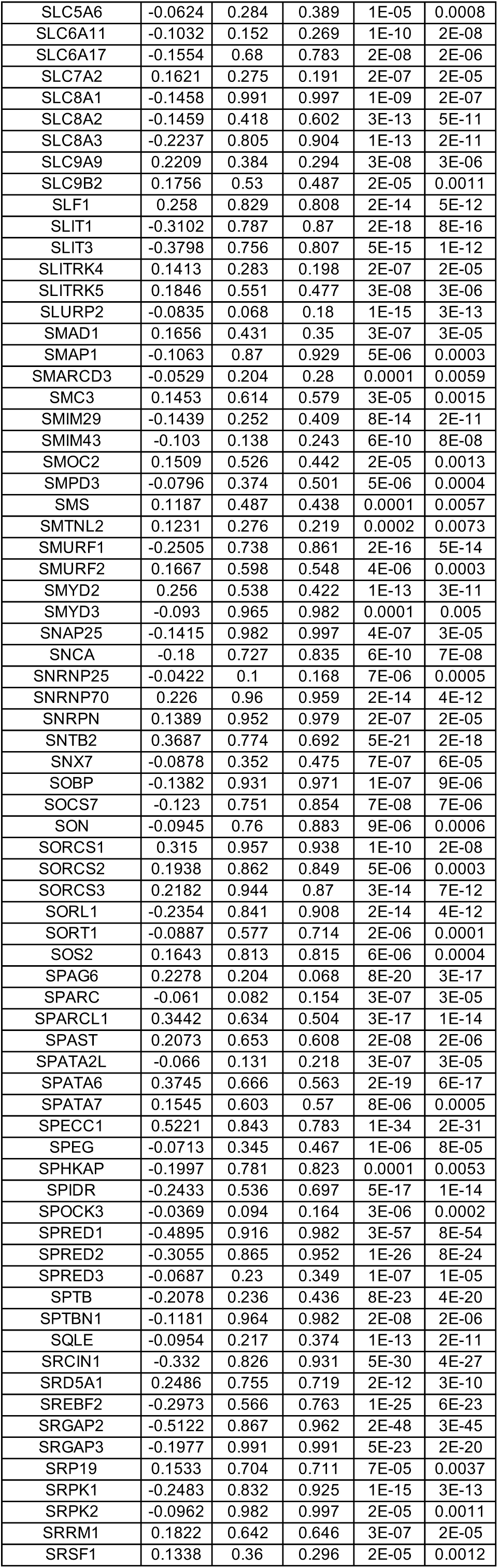
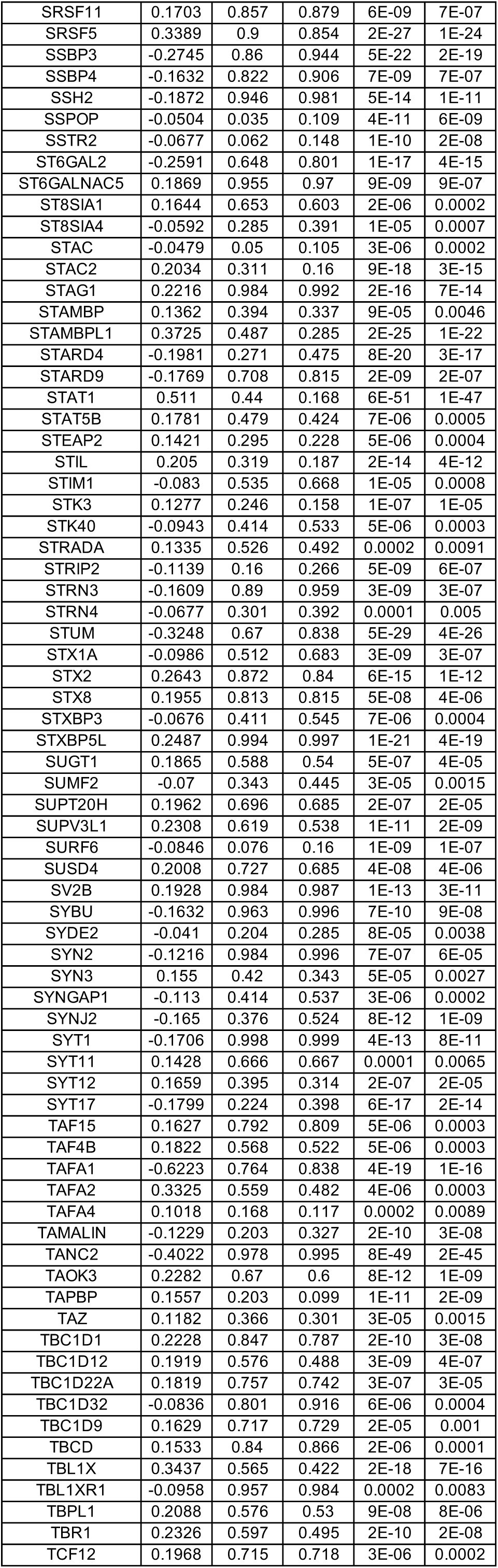
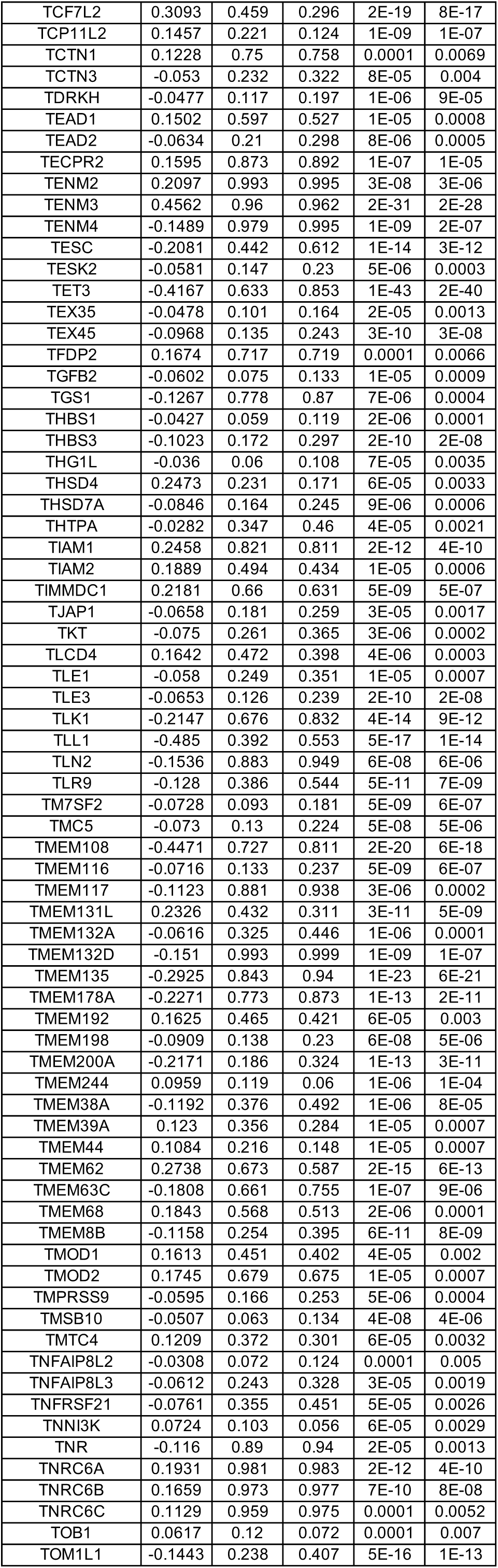
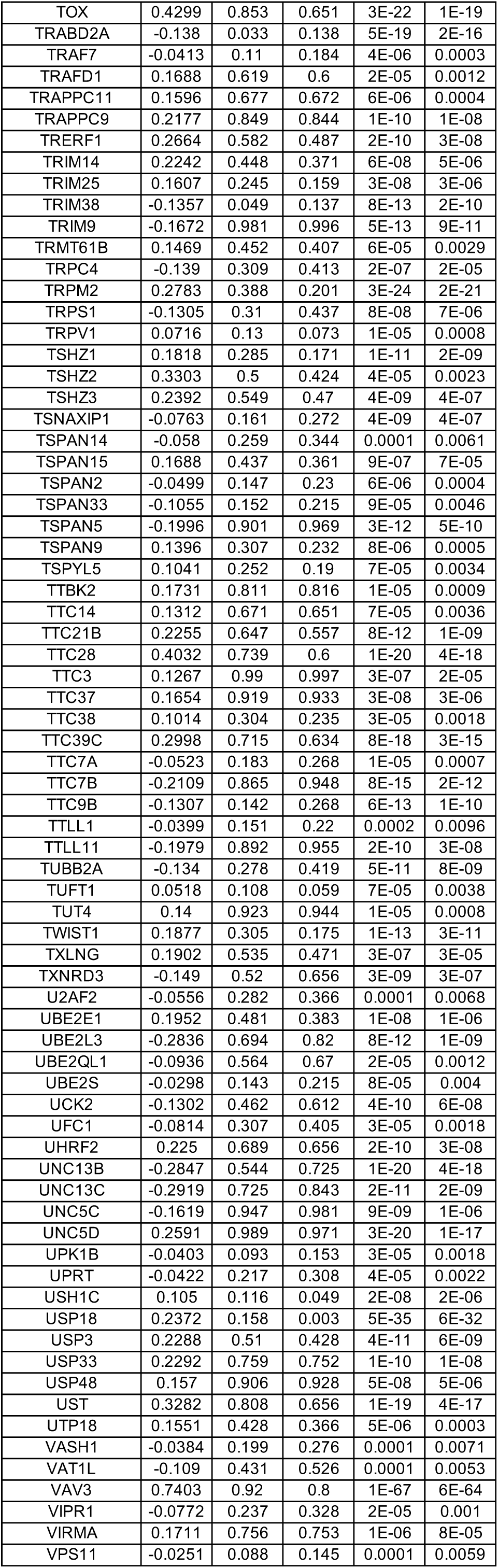
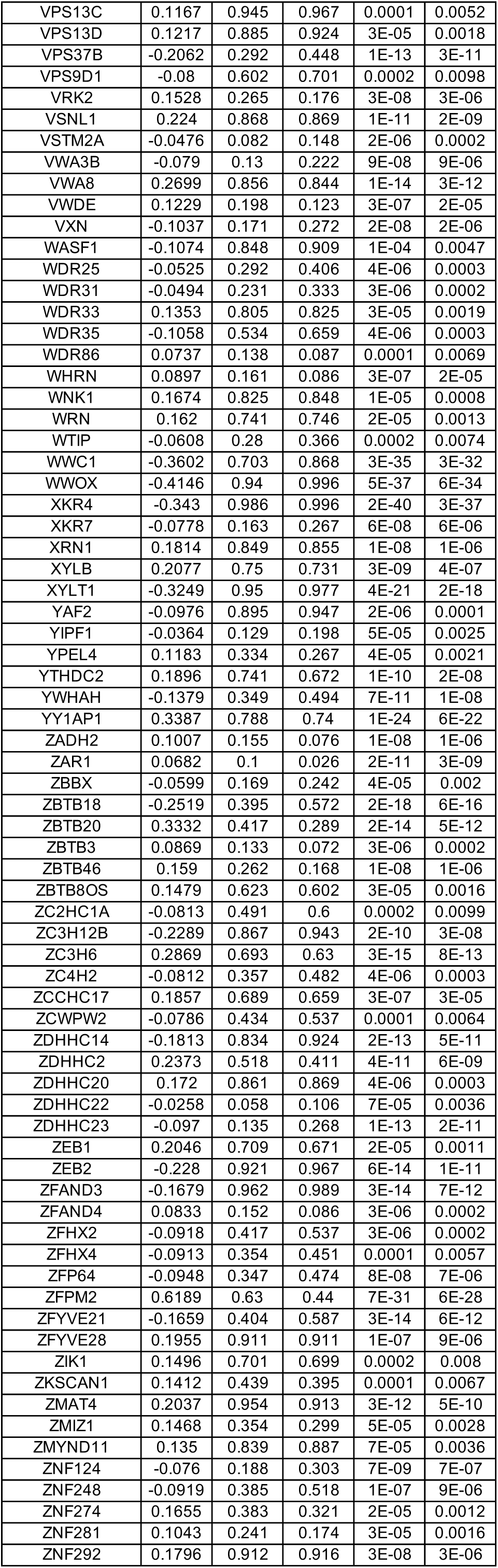
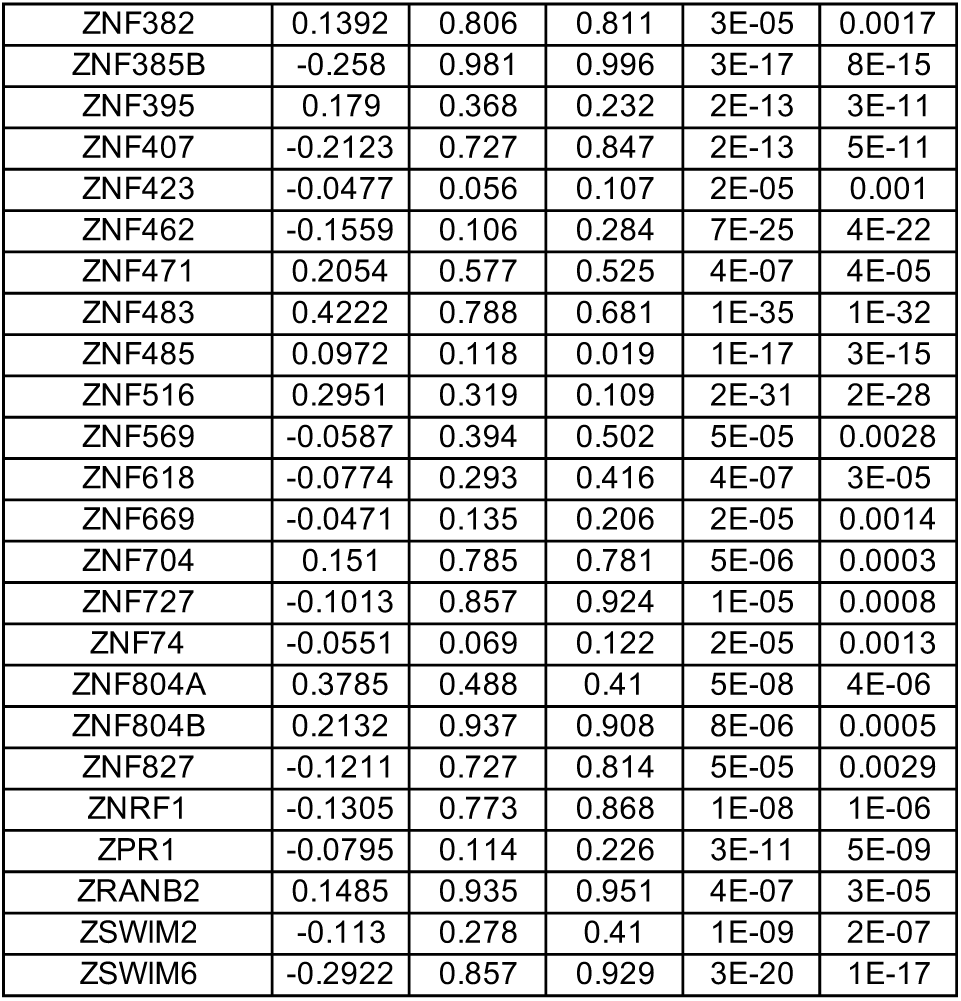
DEGs between wild-type (WT) and MECP2-null (KO) upper layer excitatory neurons (Ex_3) of PFC (adjusted p-value < 0.01)

**Table S17.**
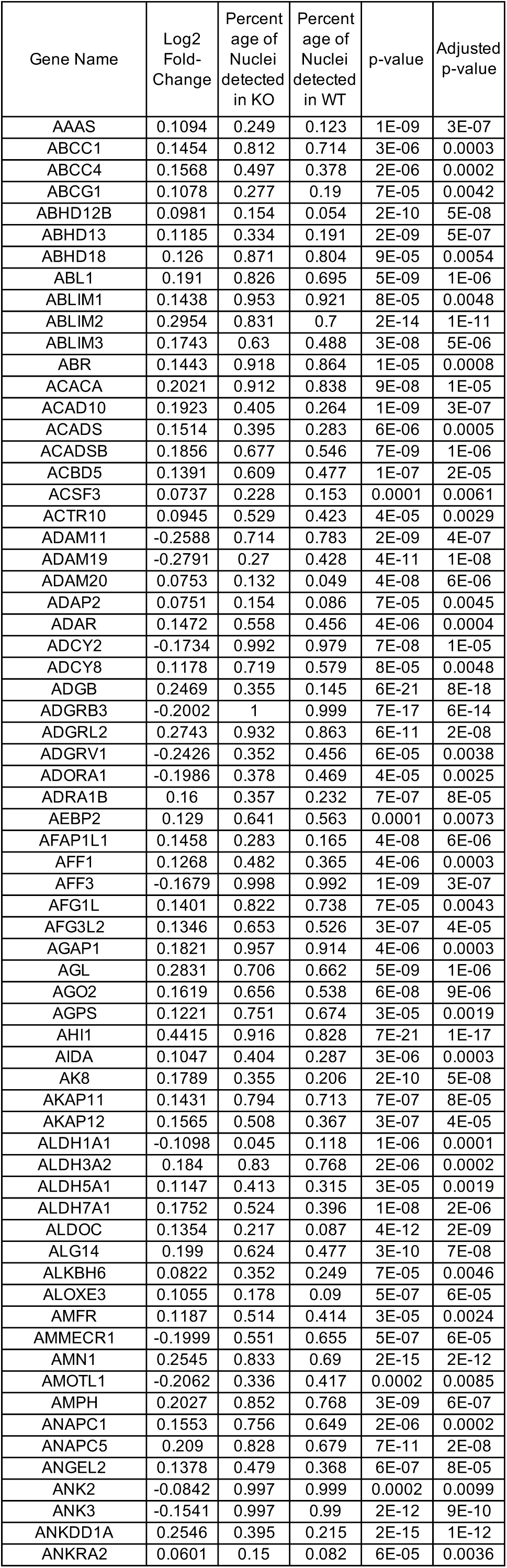
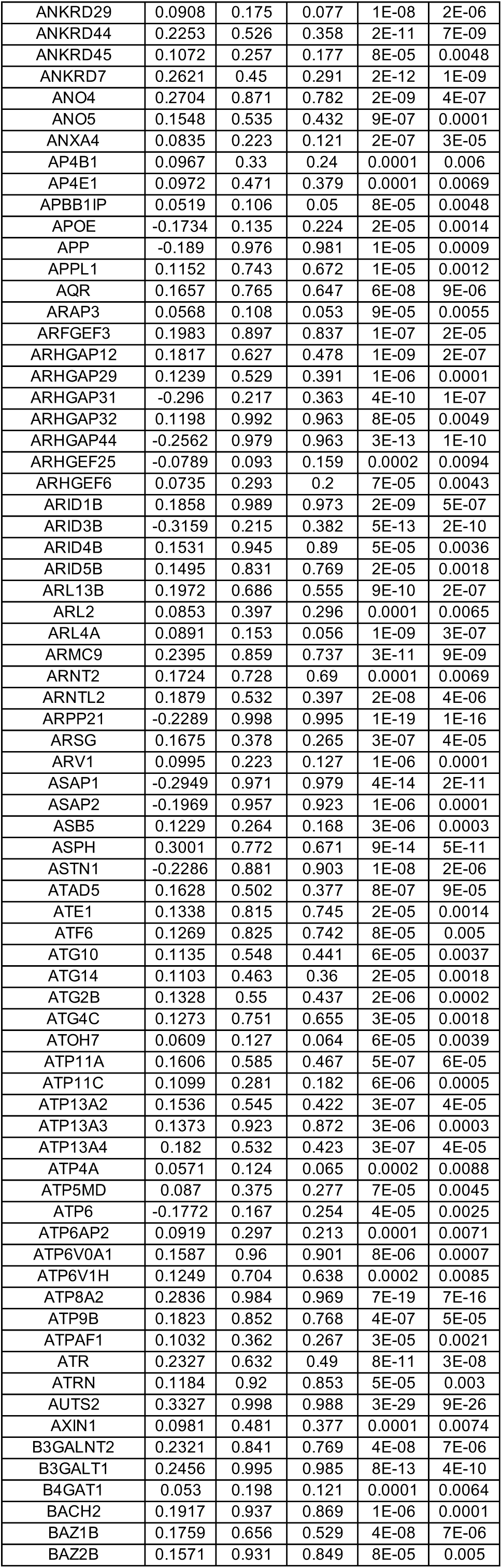
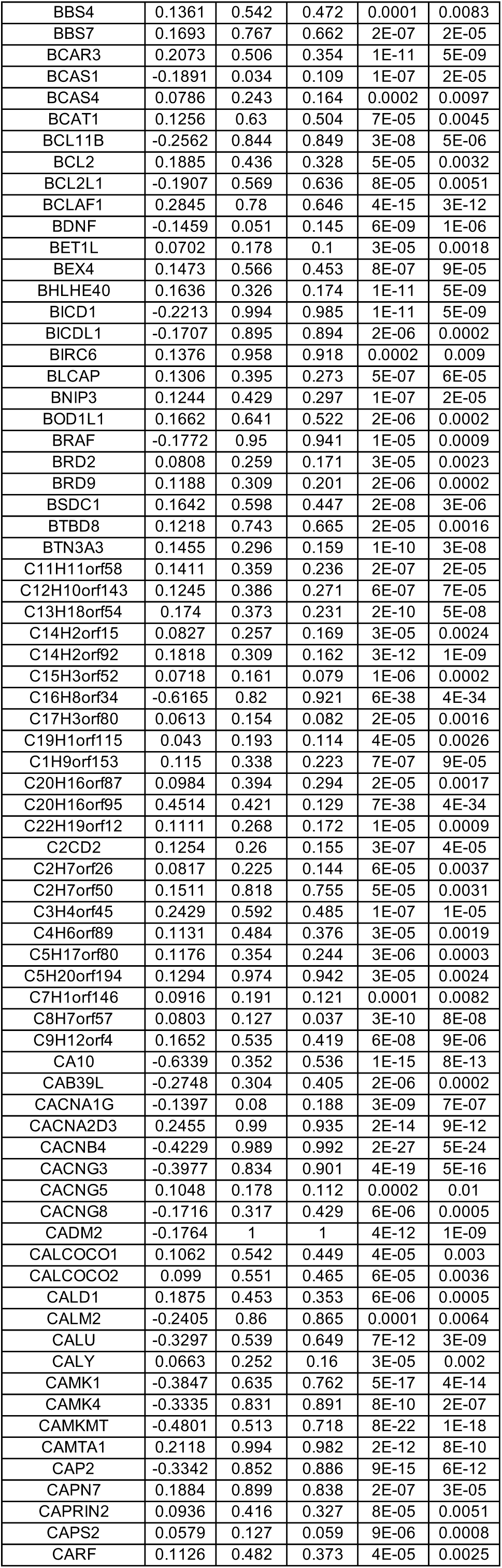
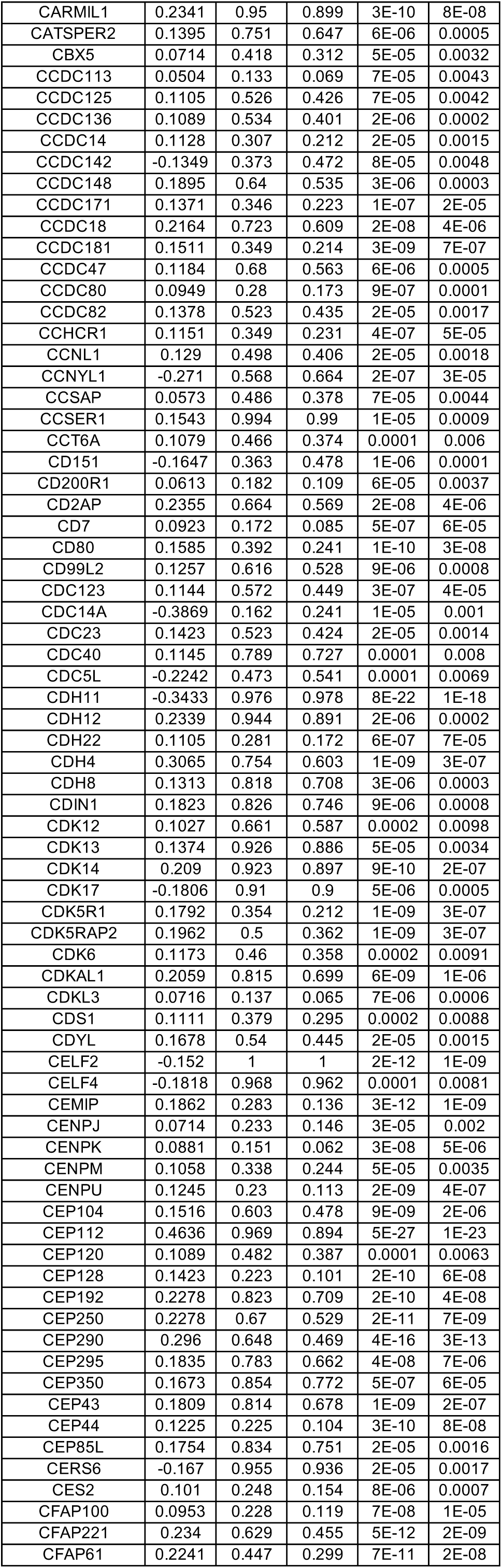
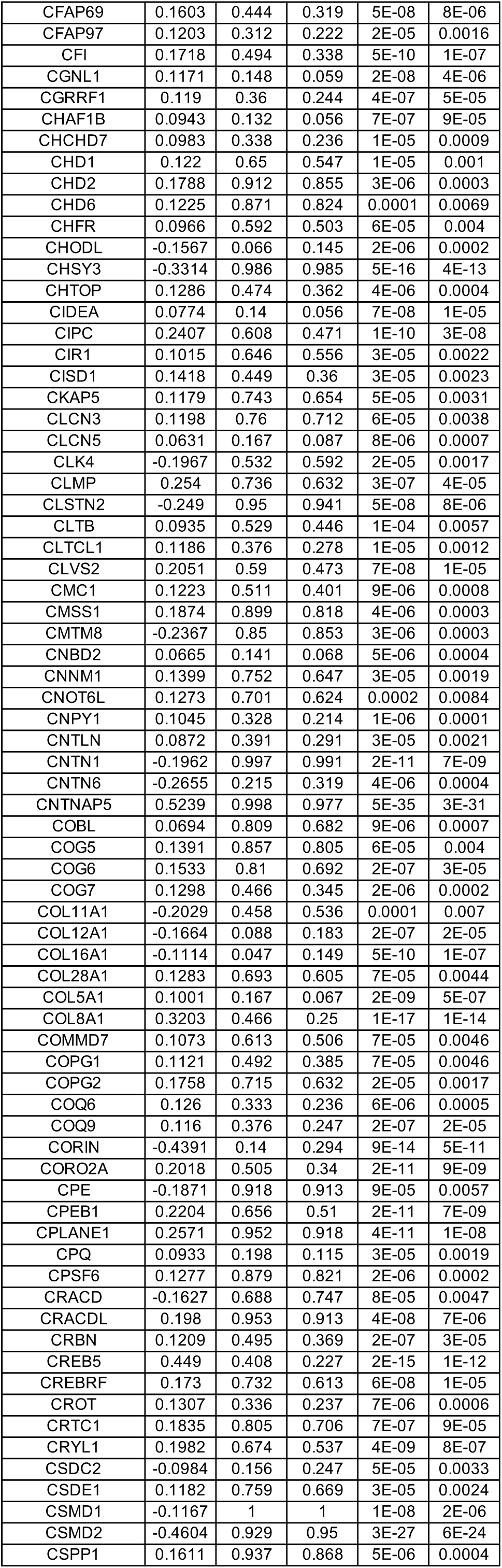
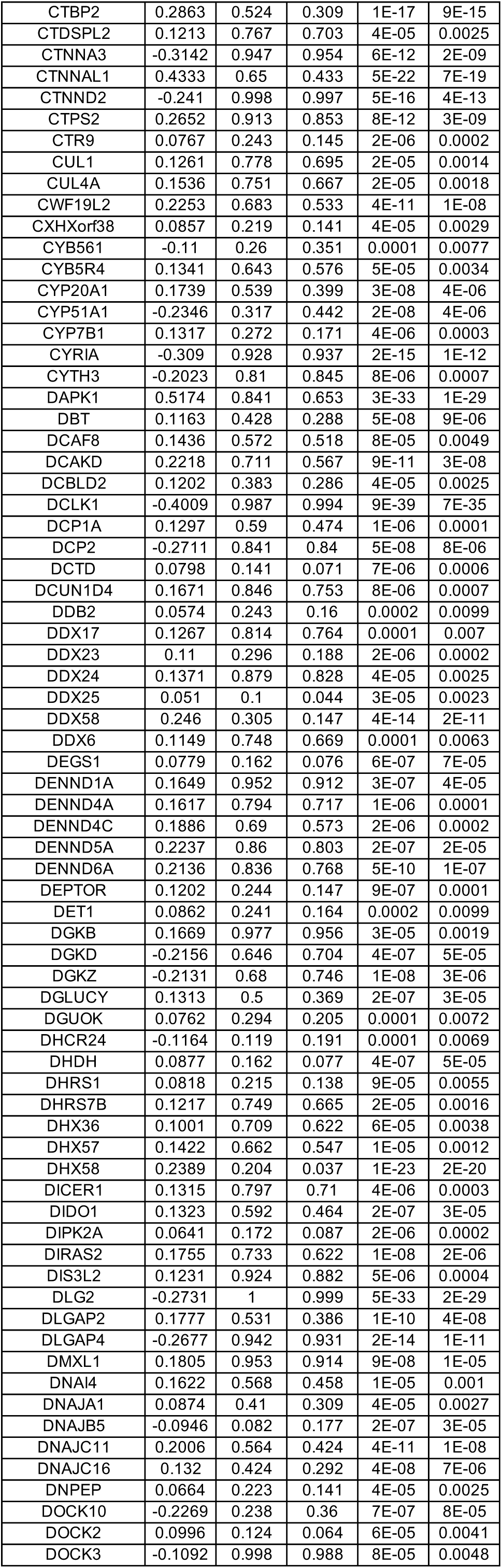
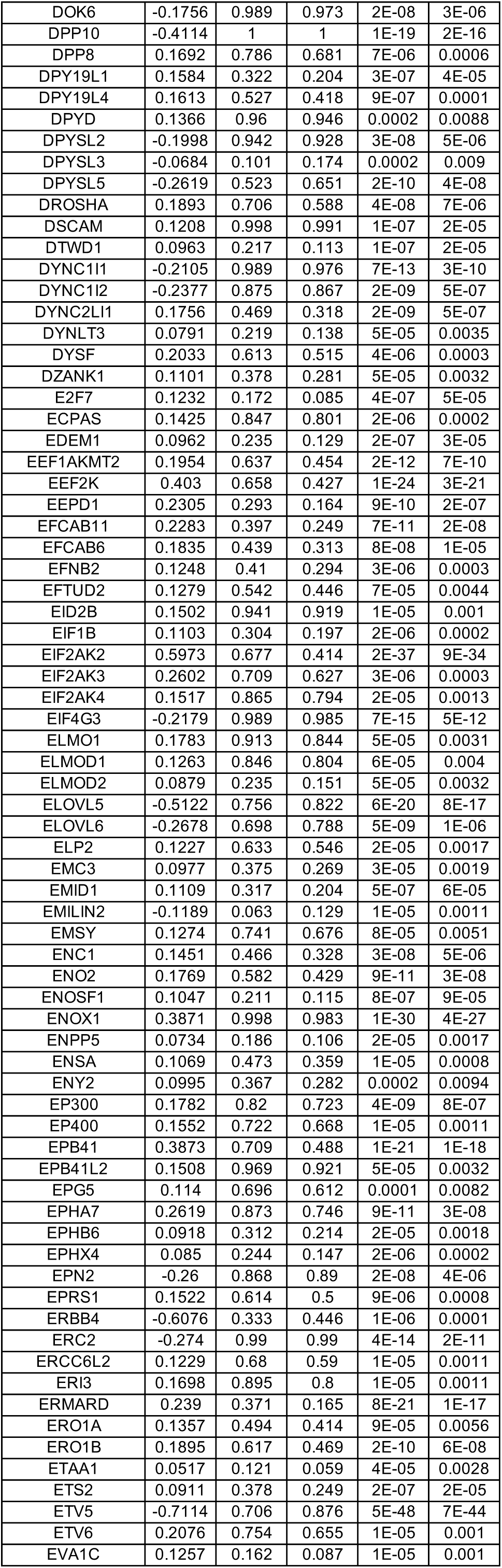
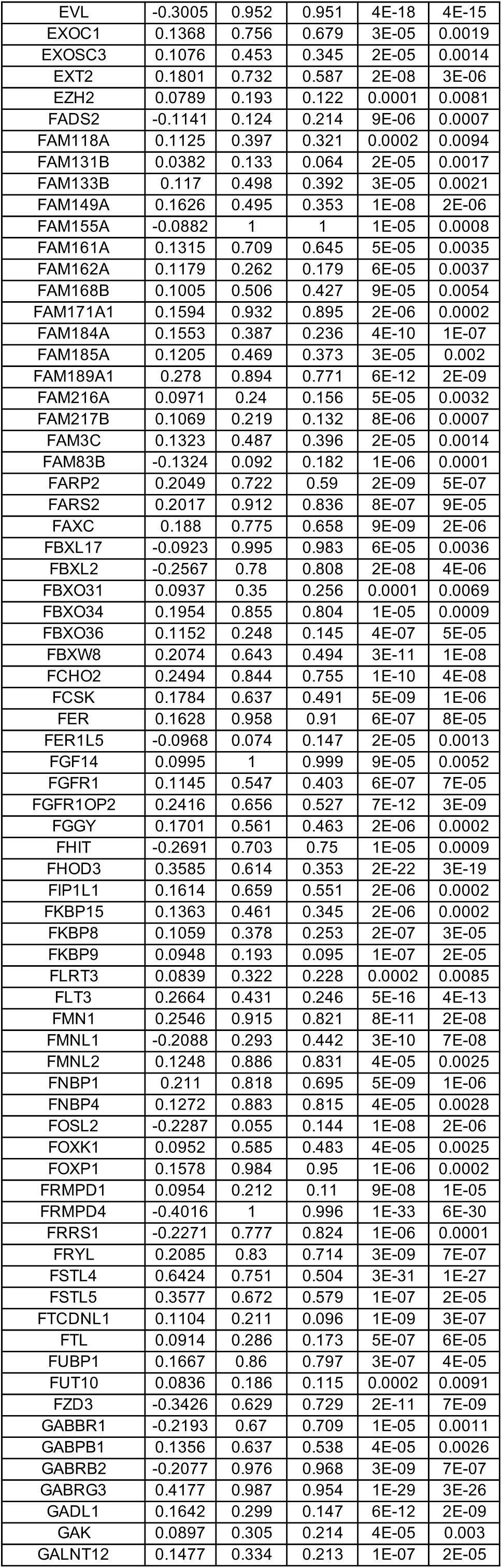
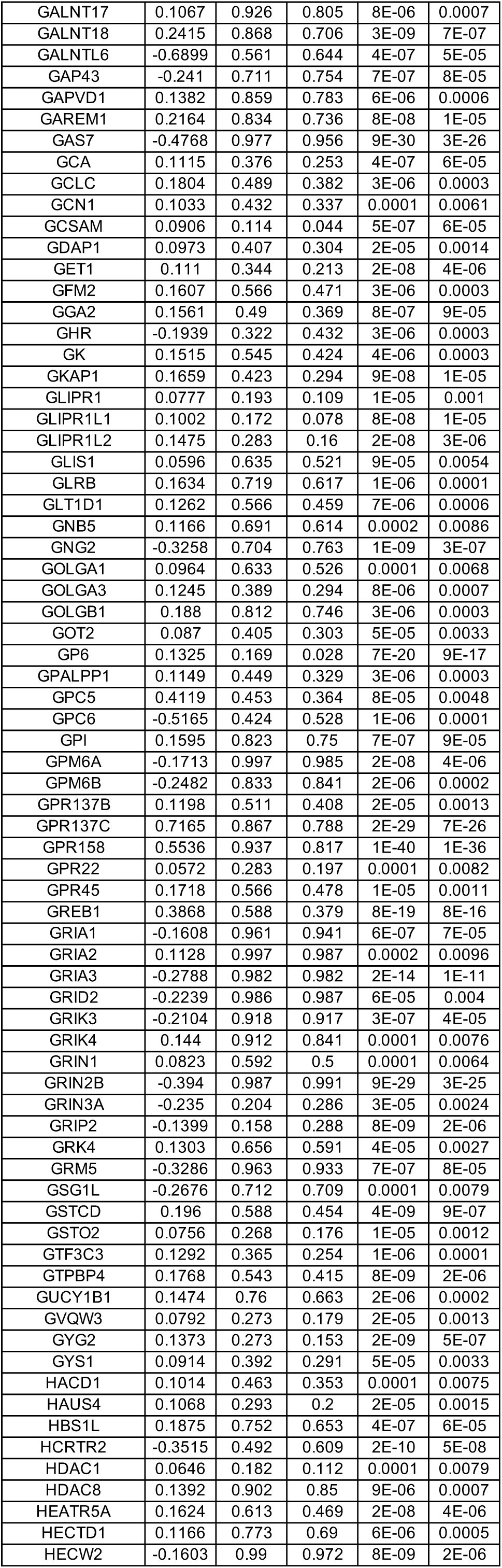
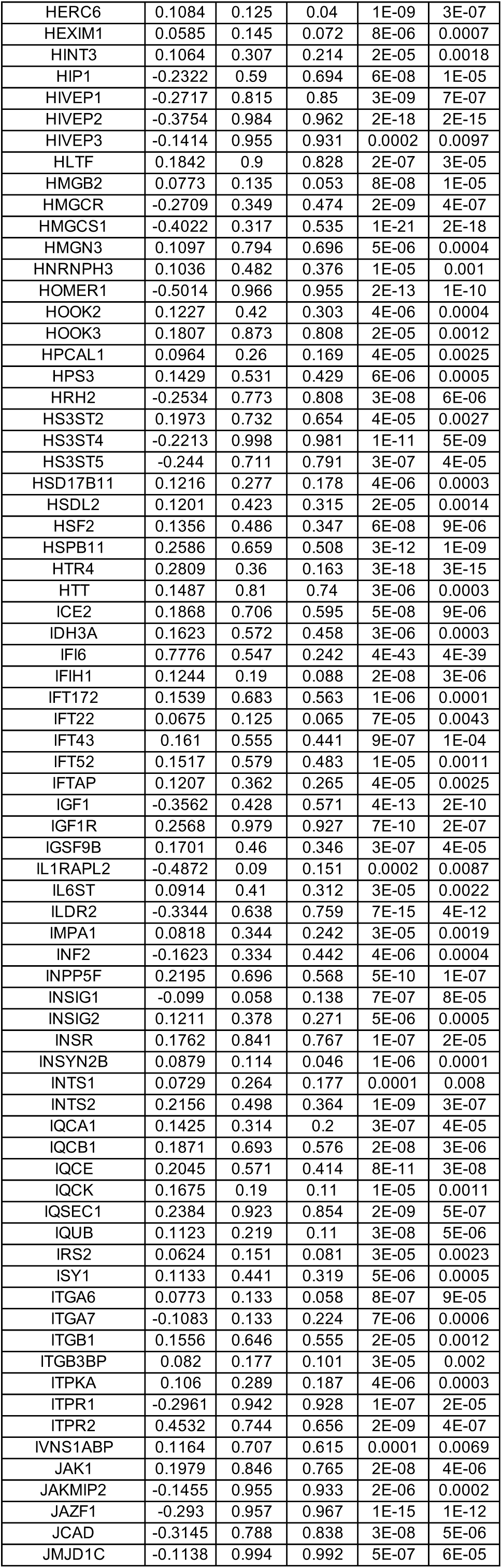
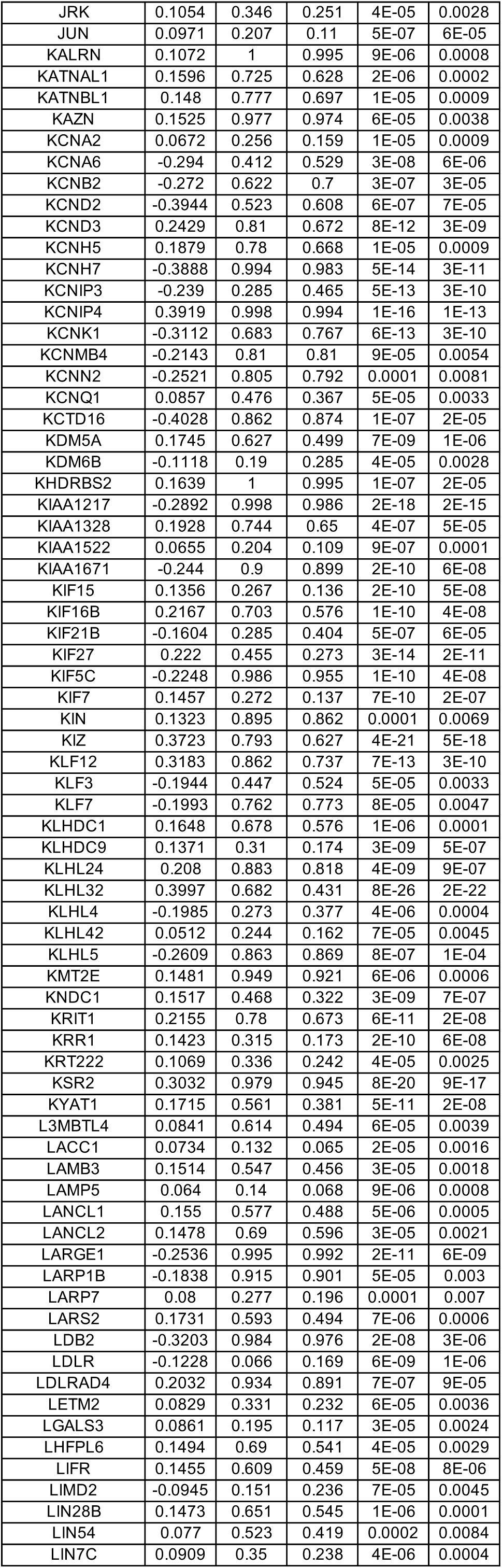
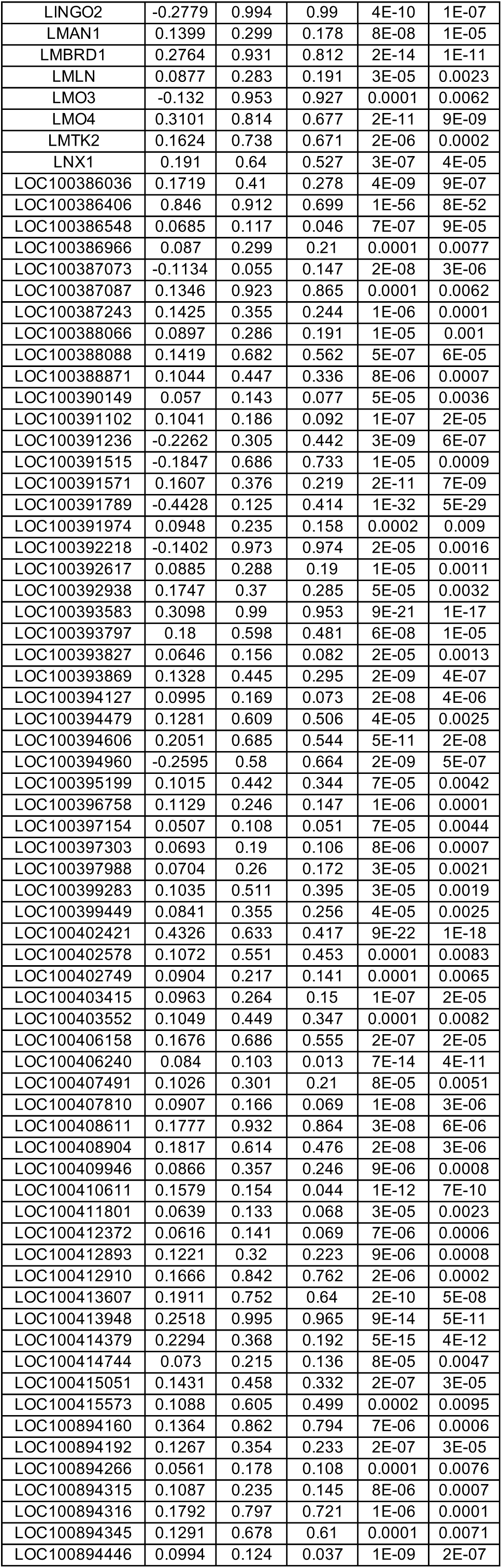
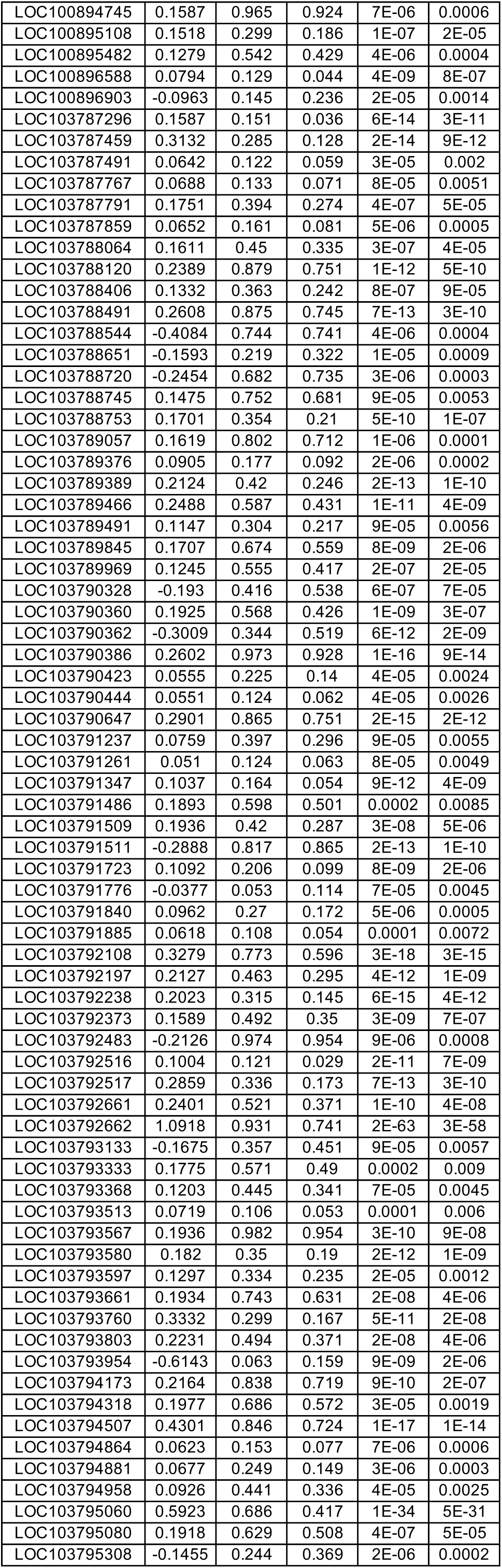
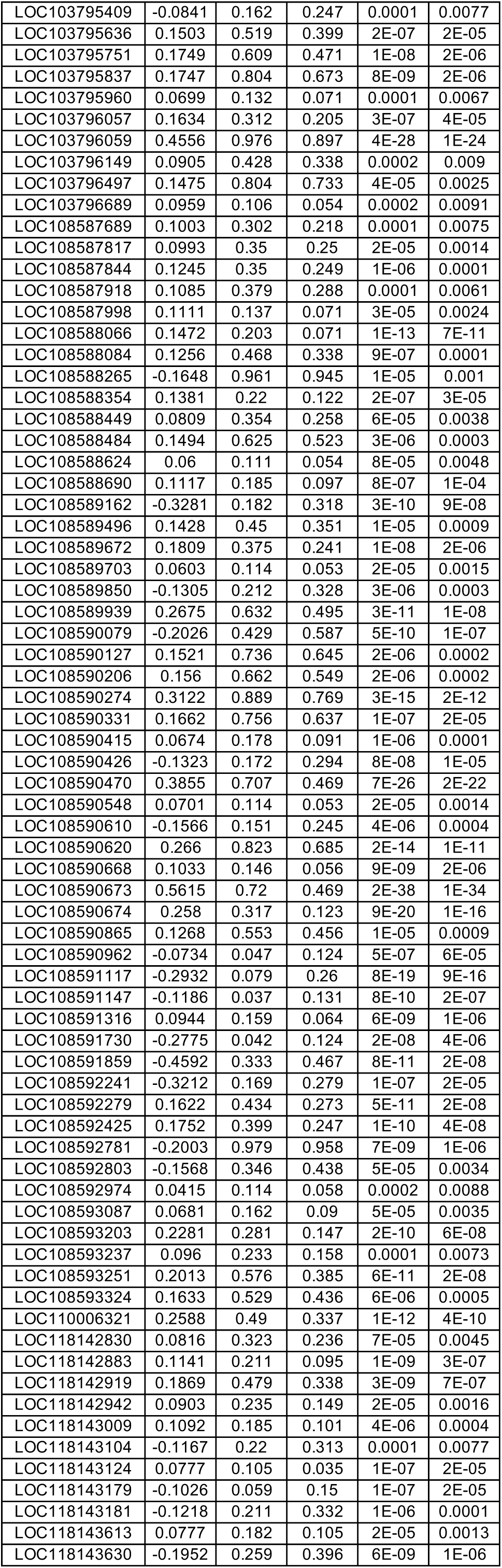
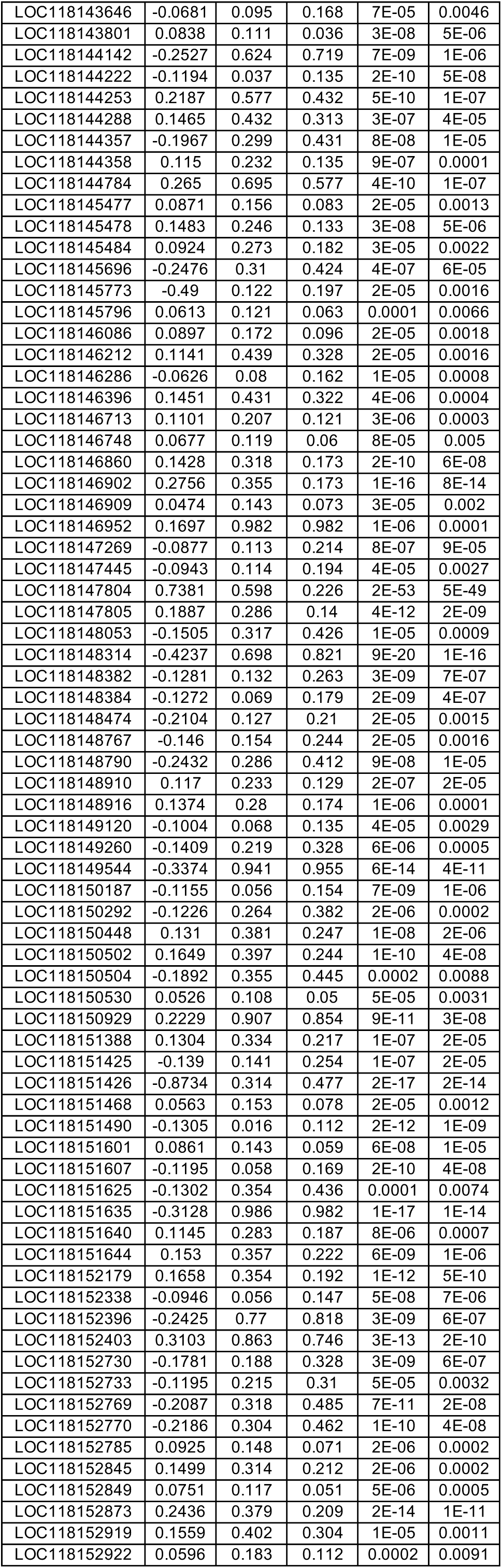
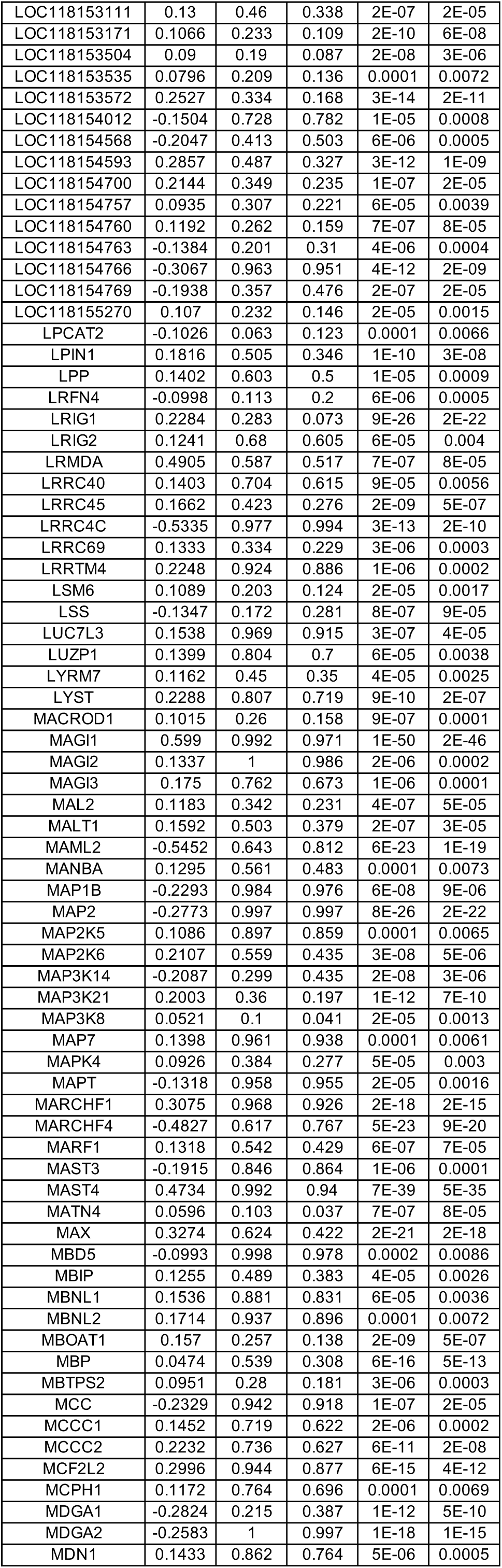
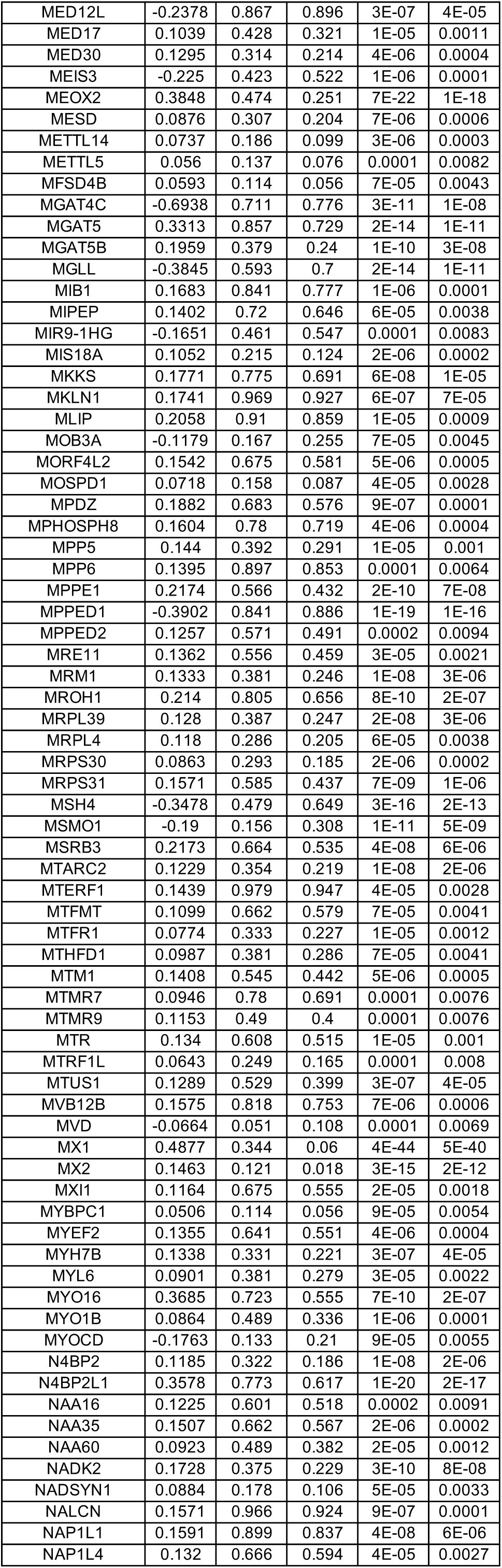
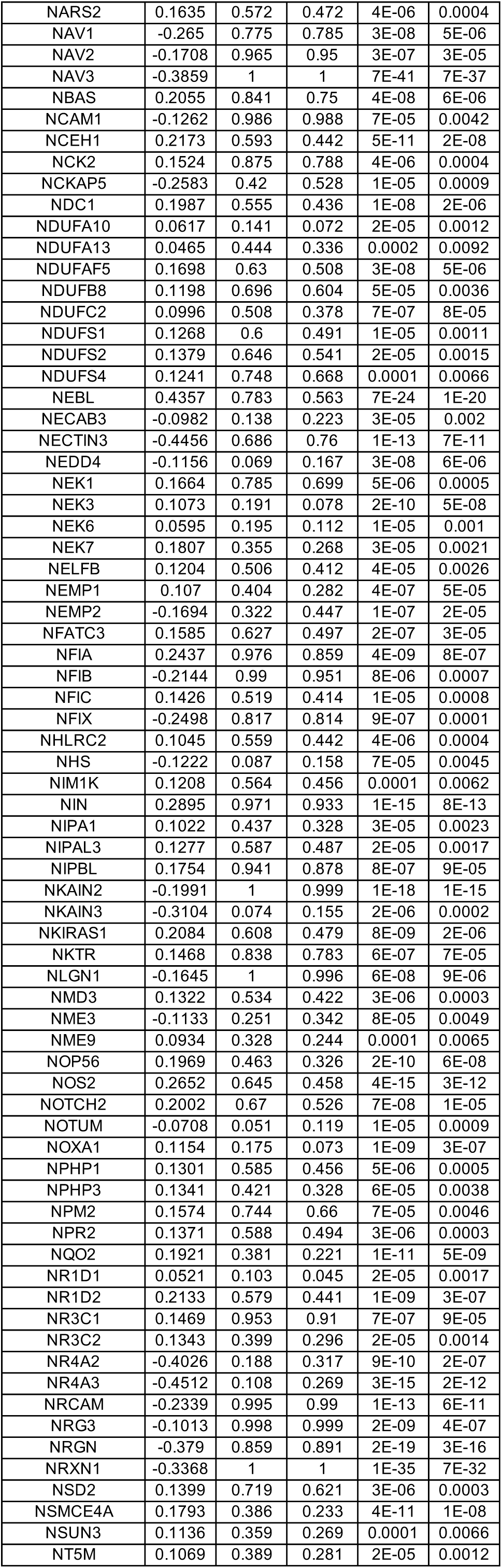
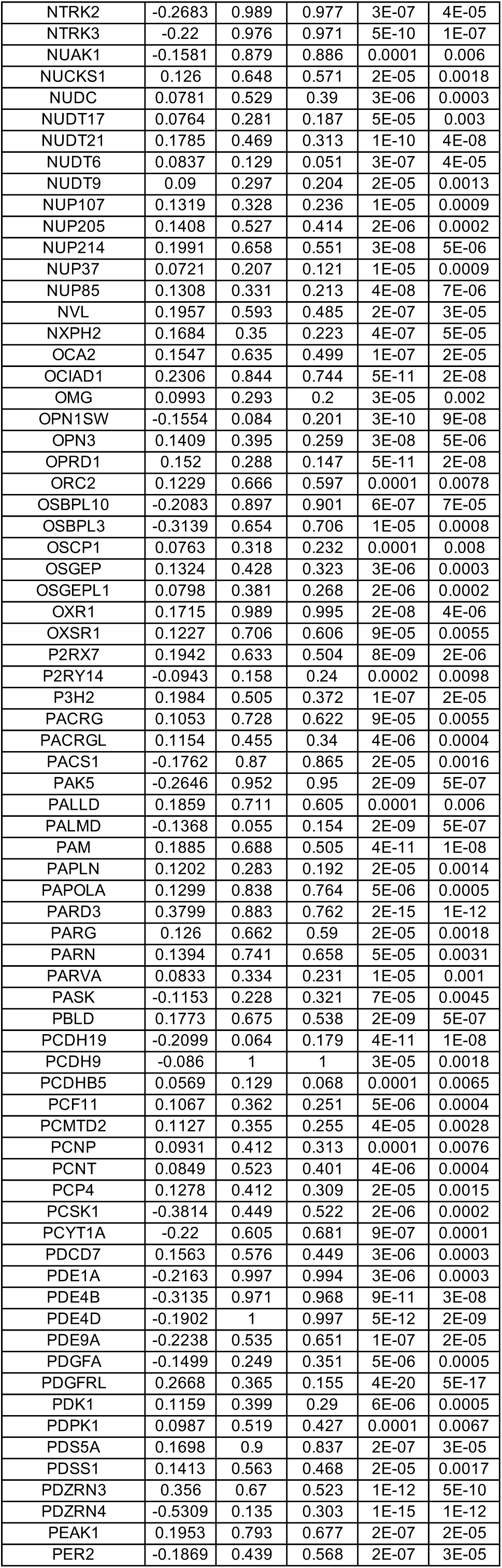
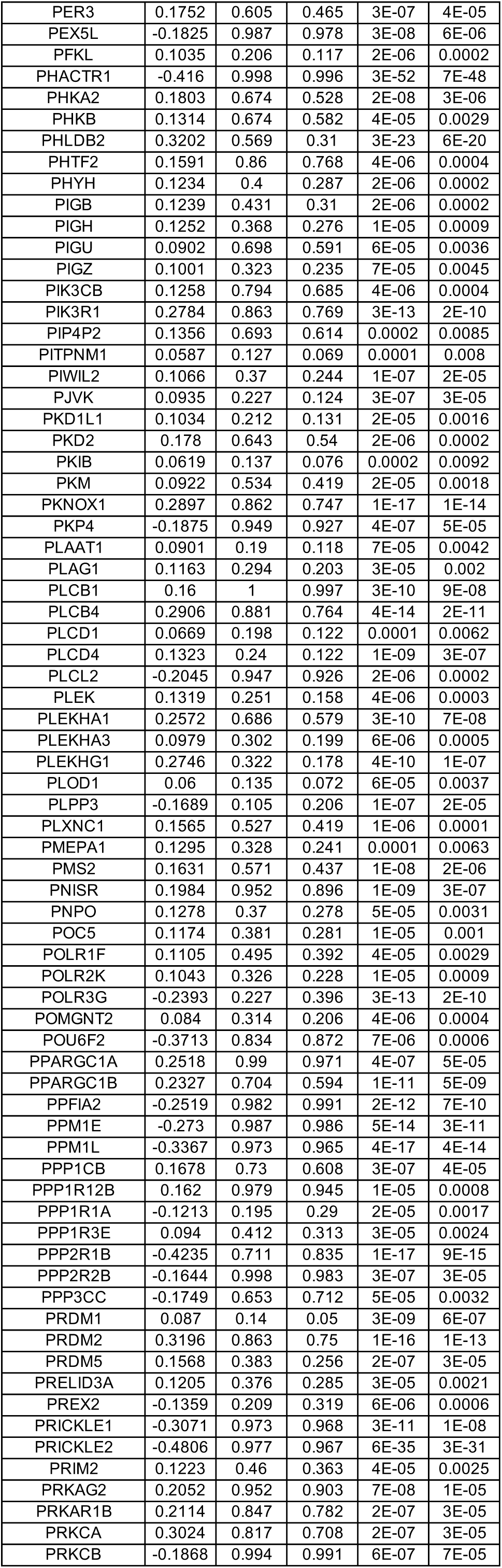
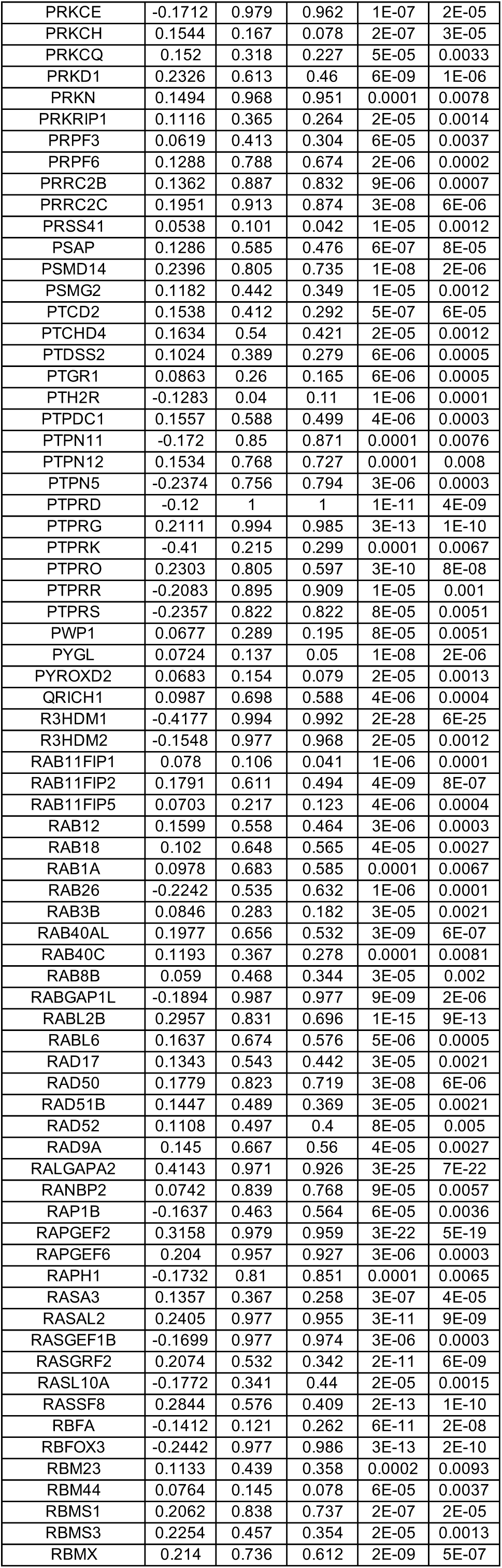
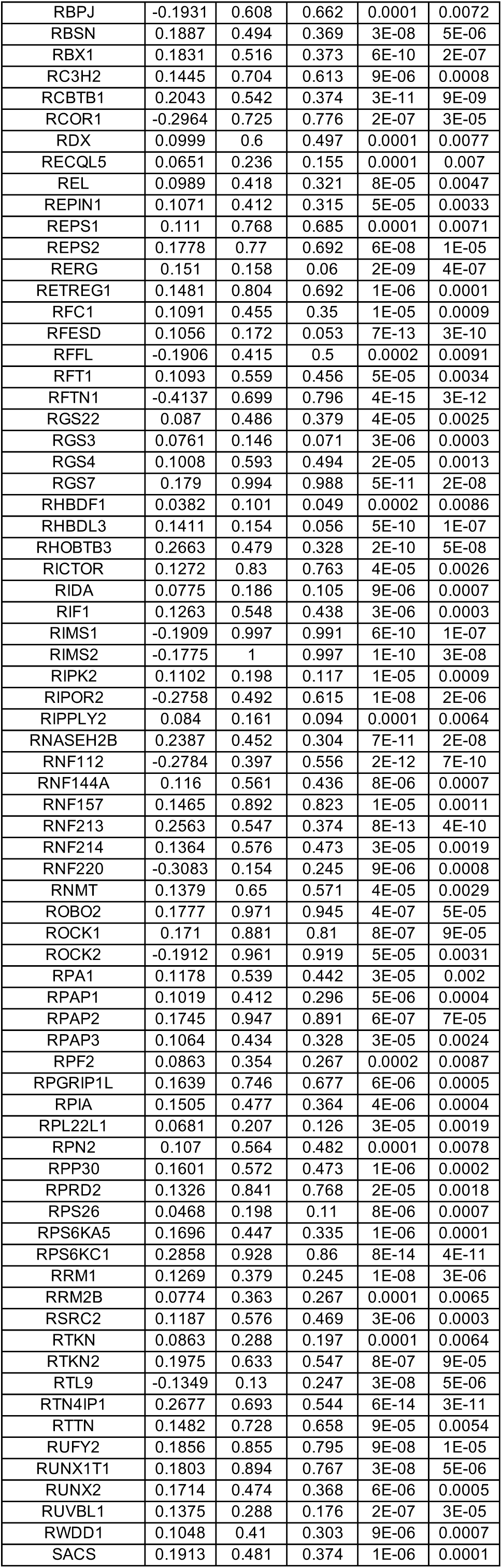
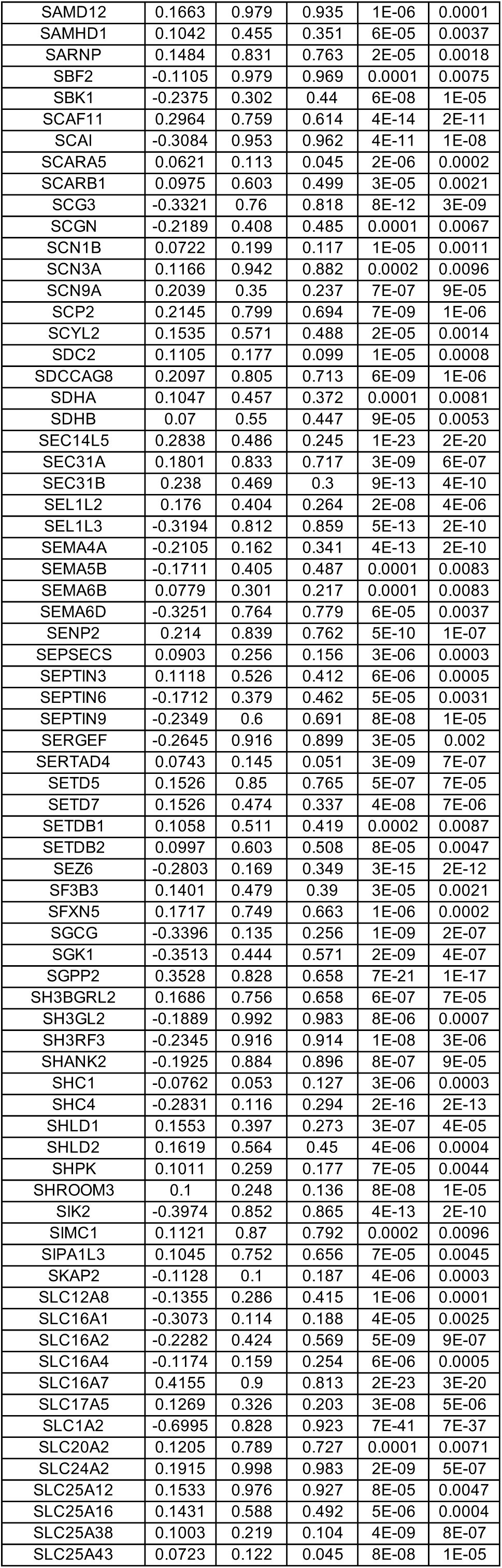
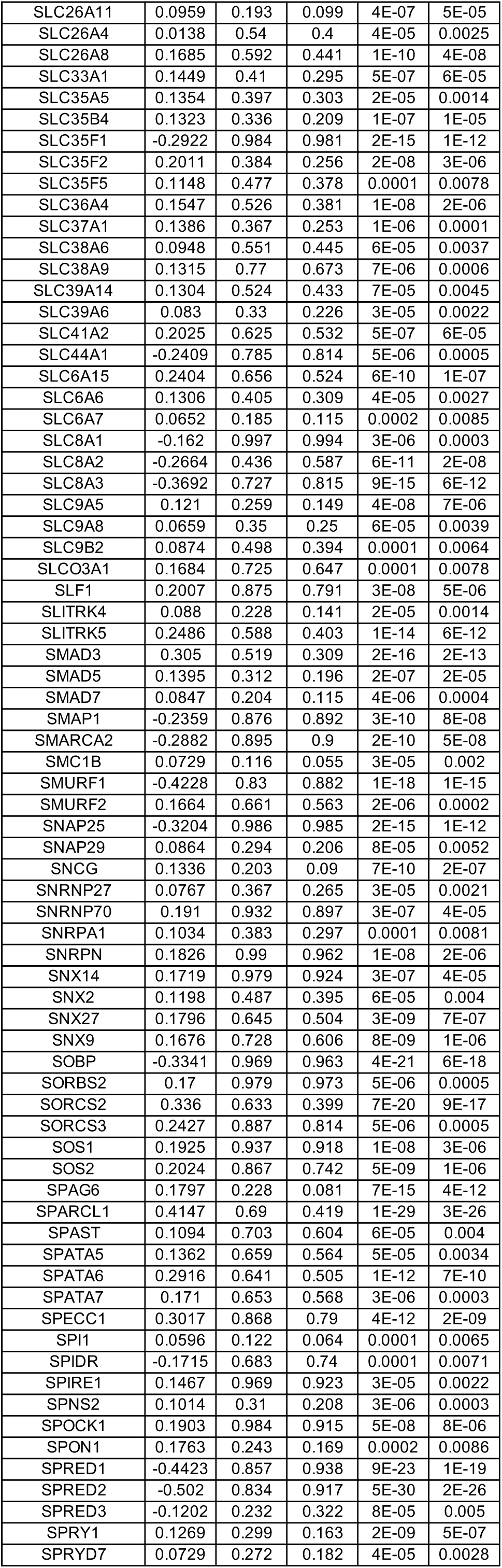
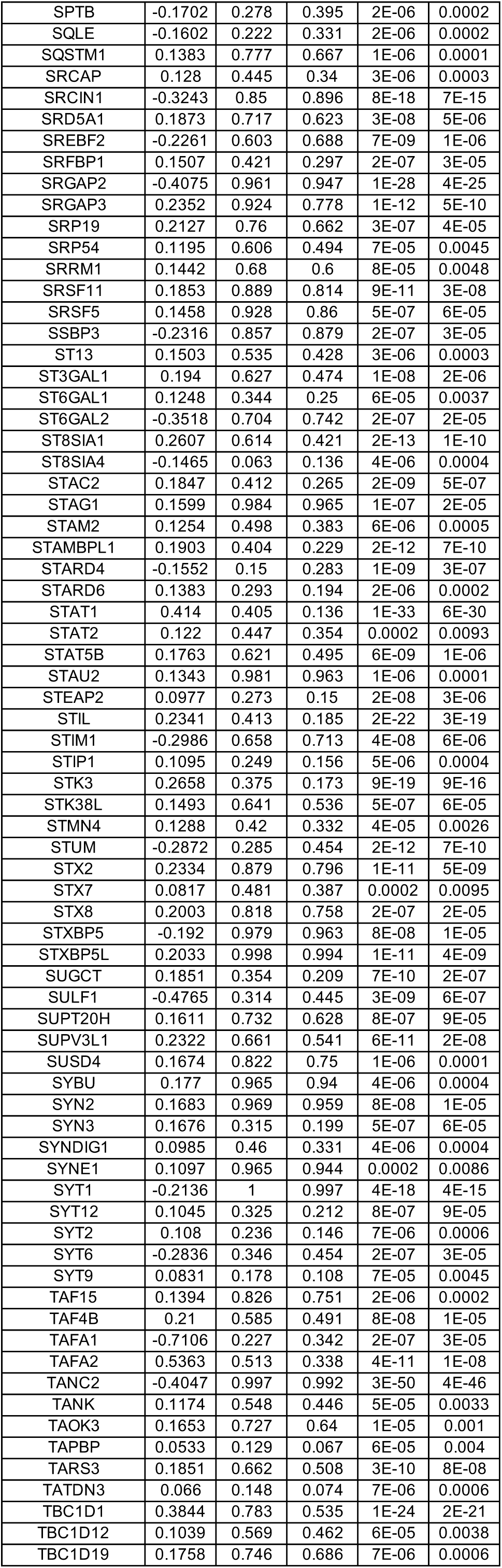
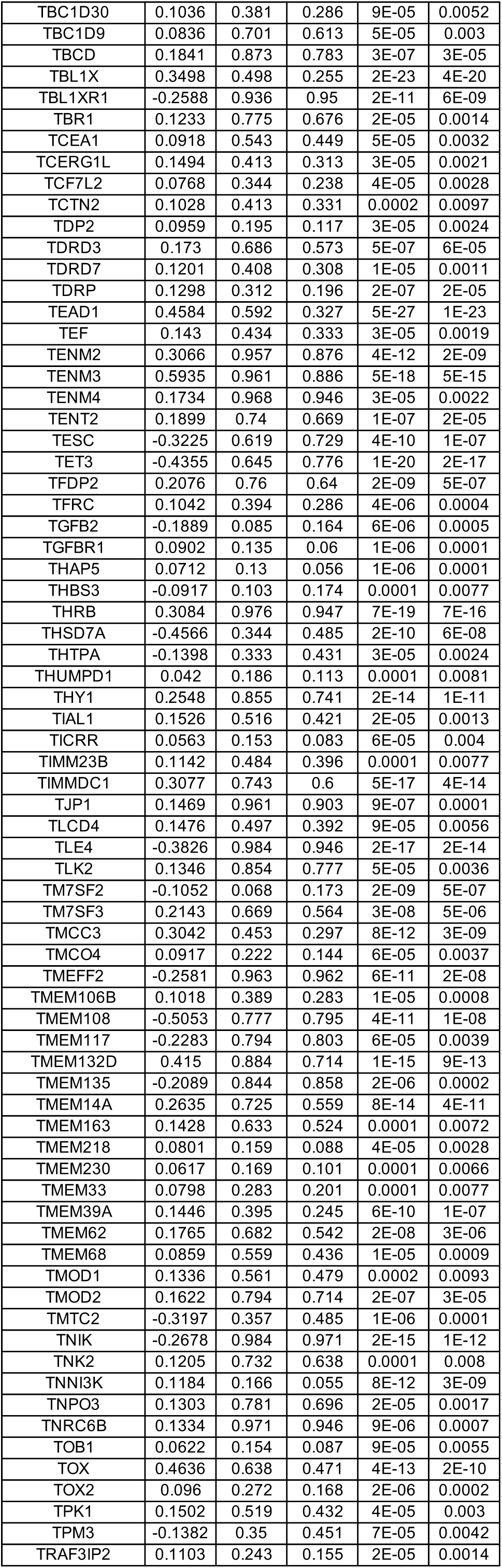
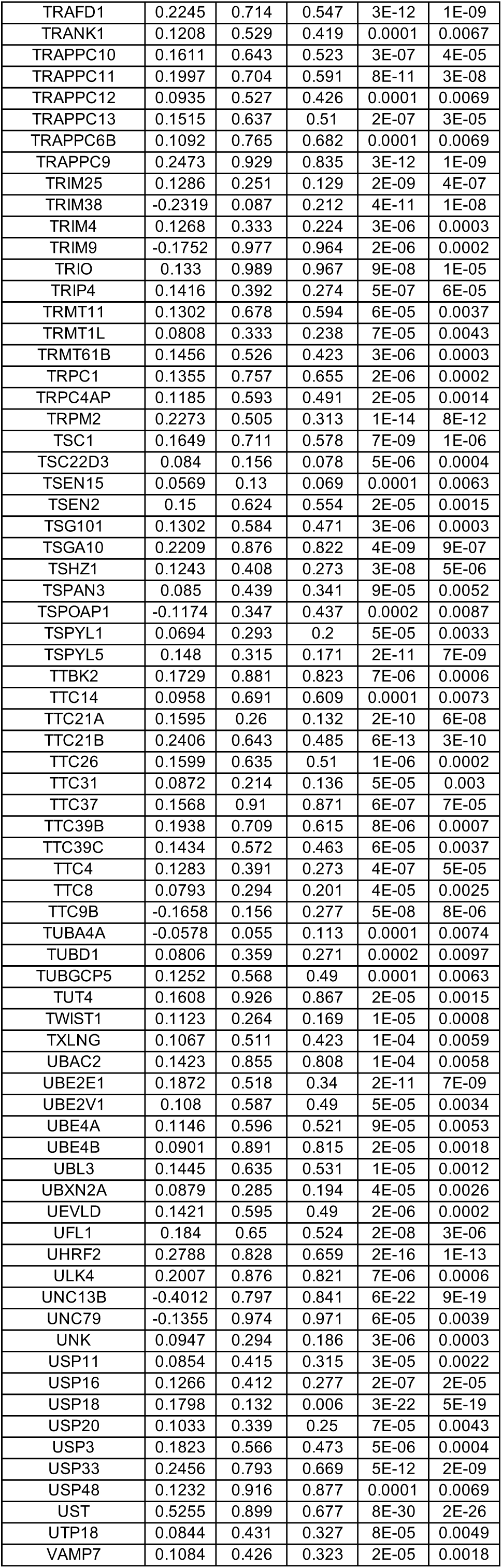
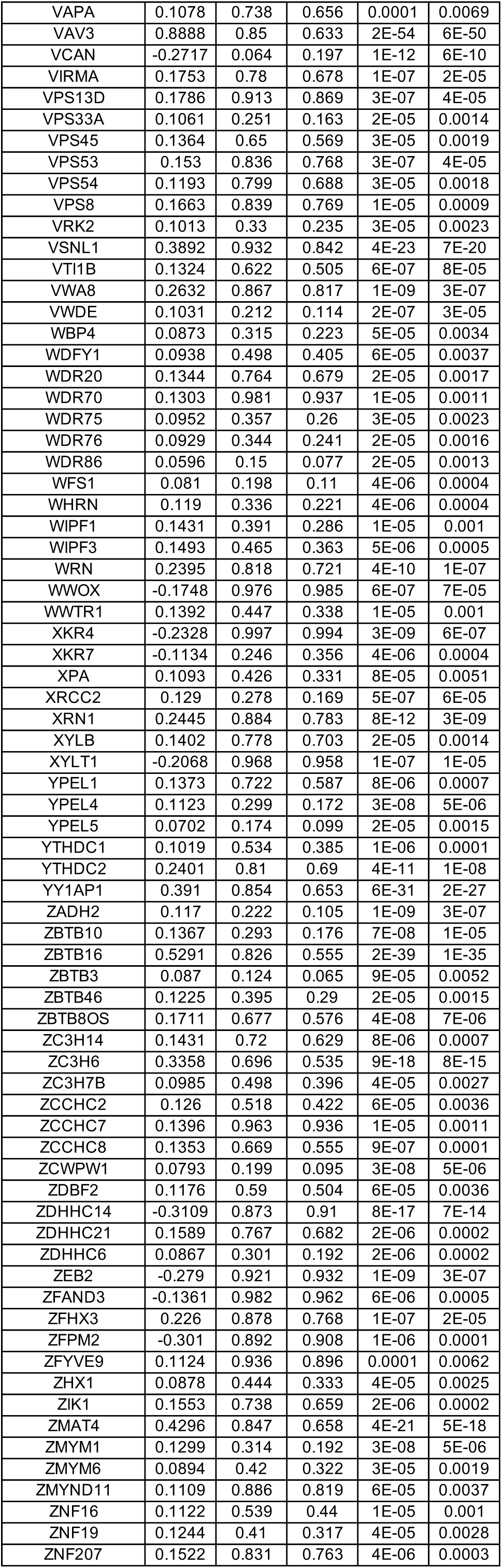
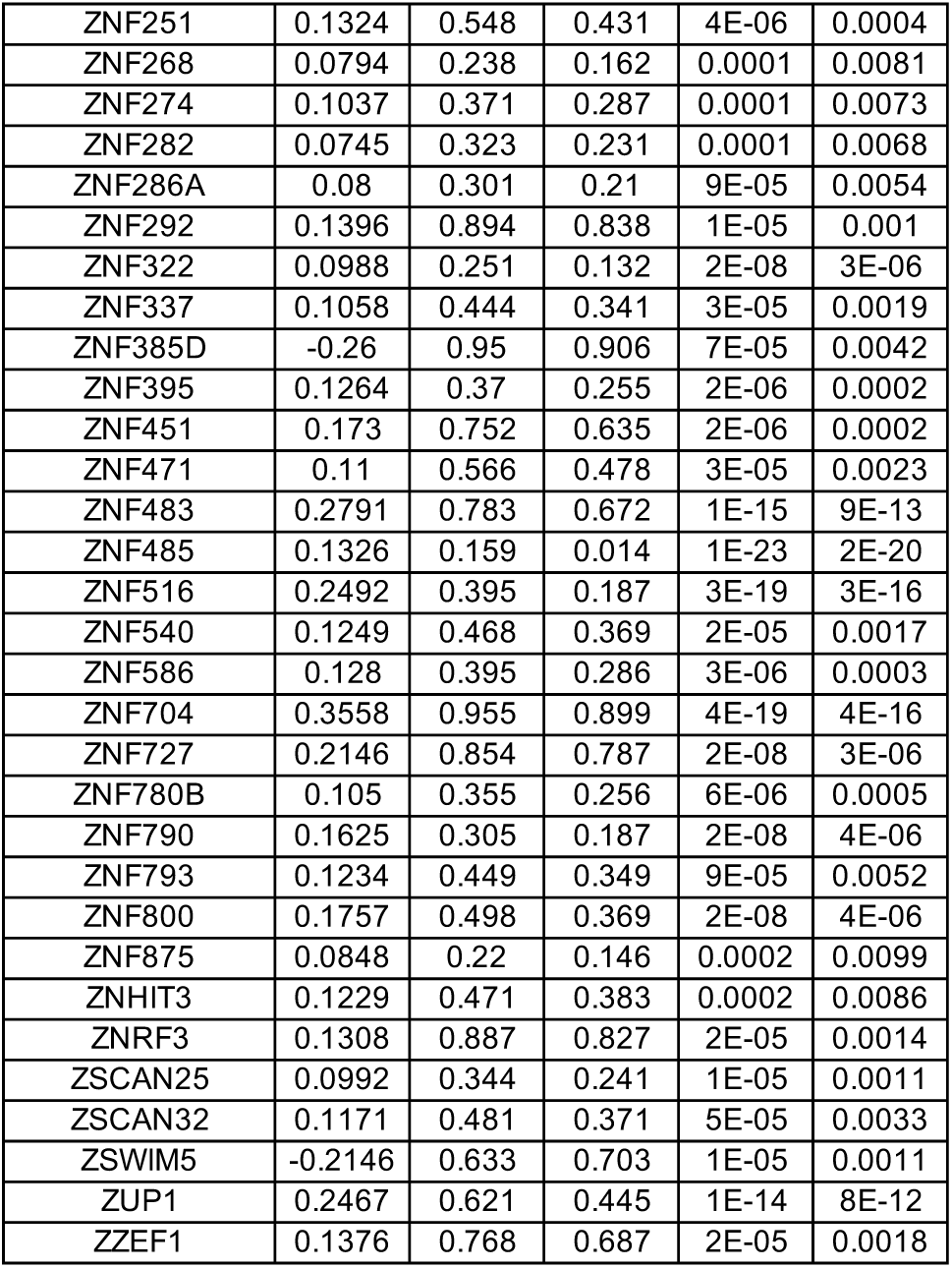
DEGs between wild-type (WT) and MECP2-null (KO) upper layer excitatory neurons (Ex_4) of PFC (adjusted p-value < 0.01)

**Table S18.**
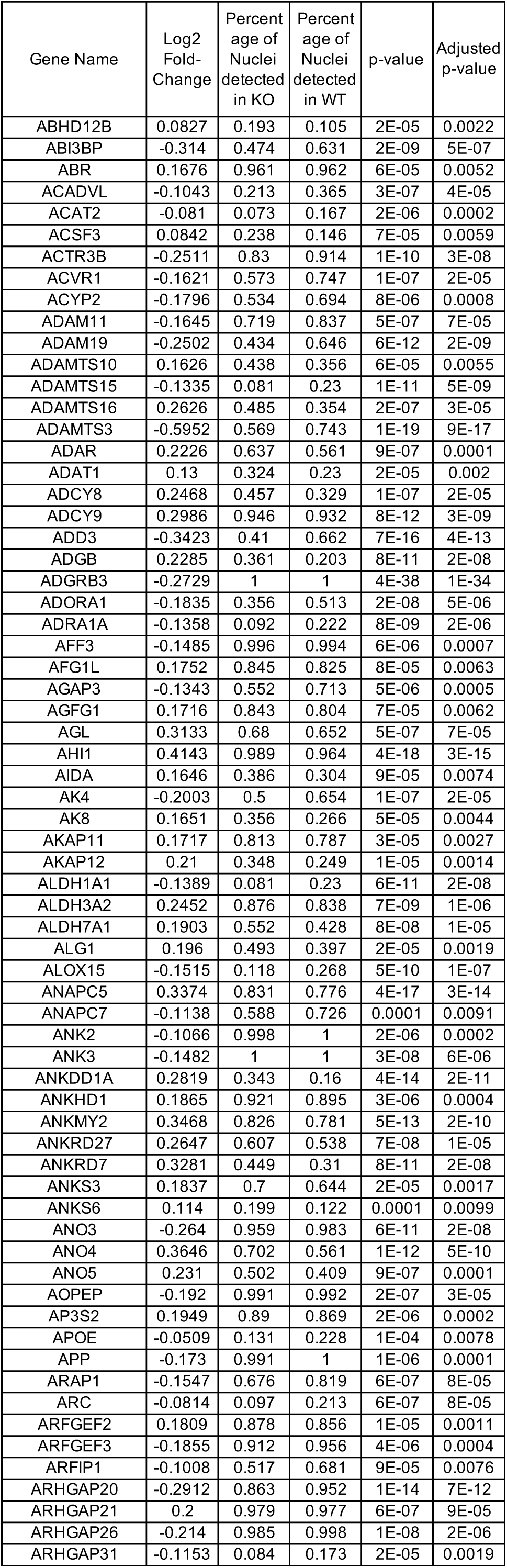
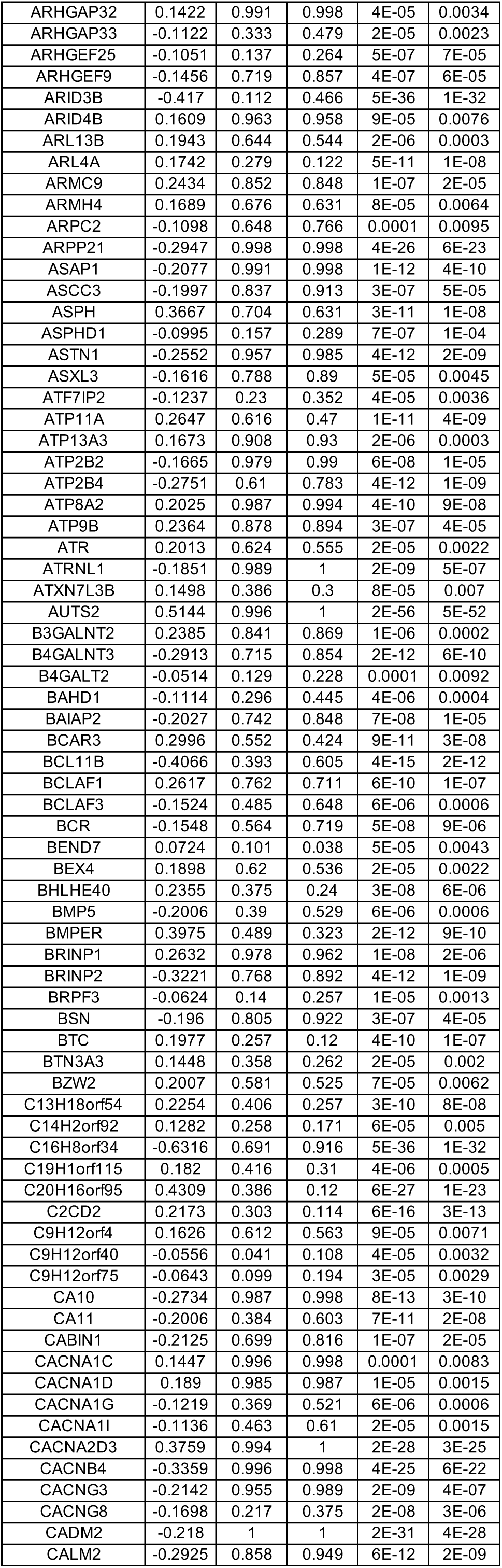
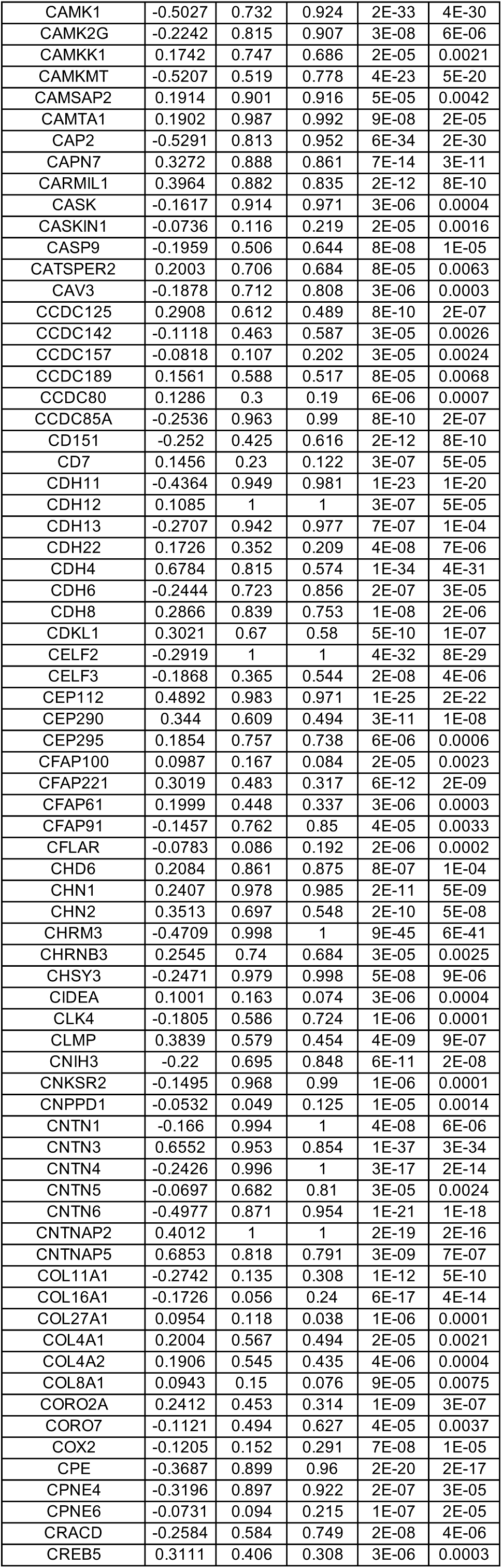
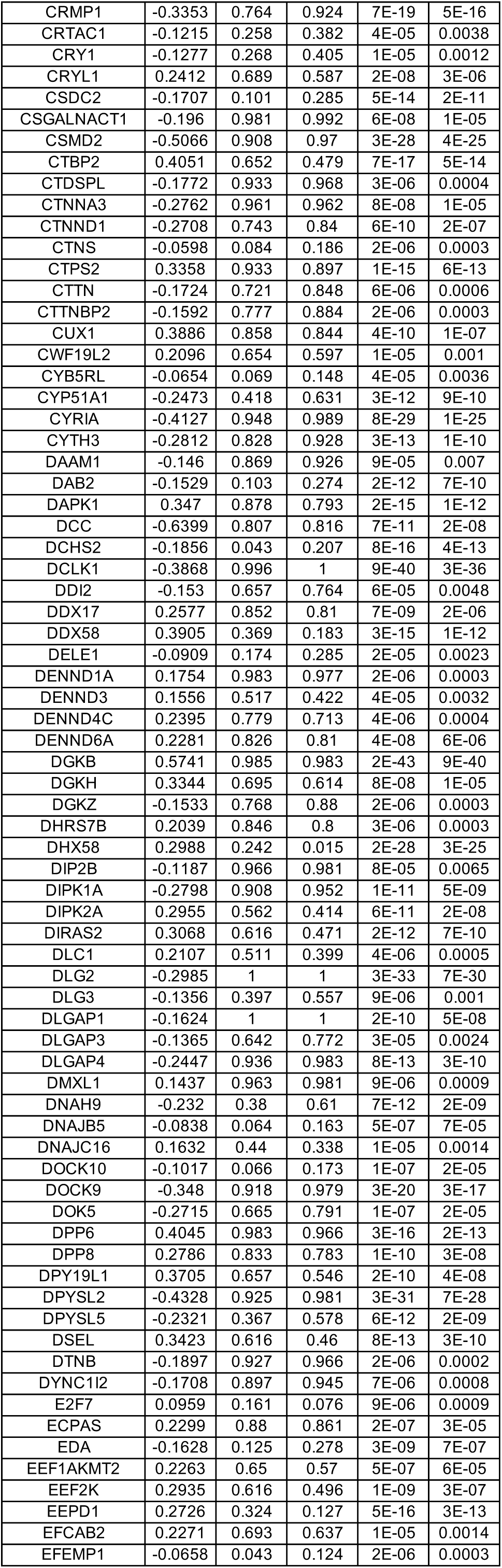
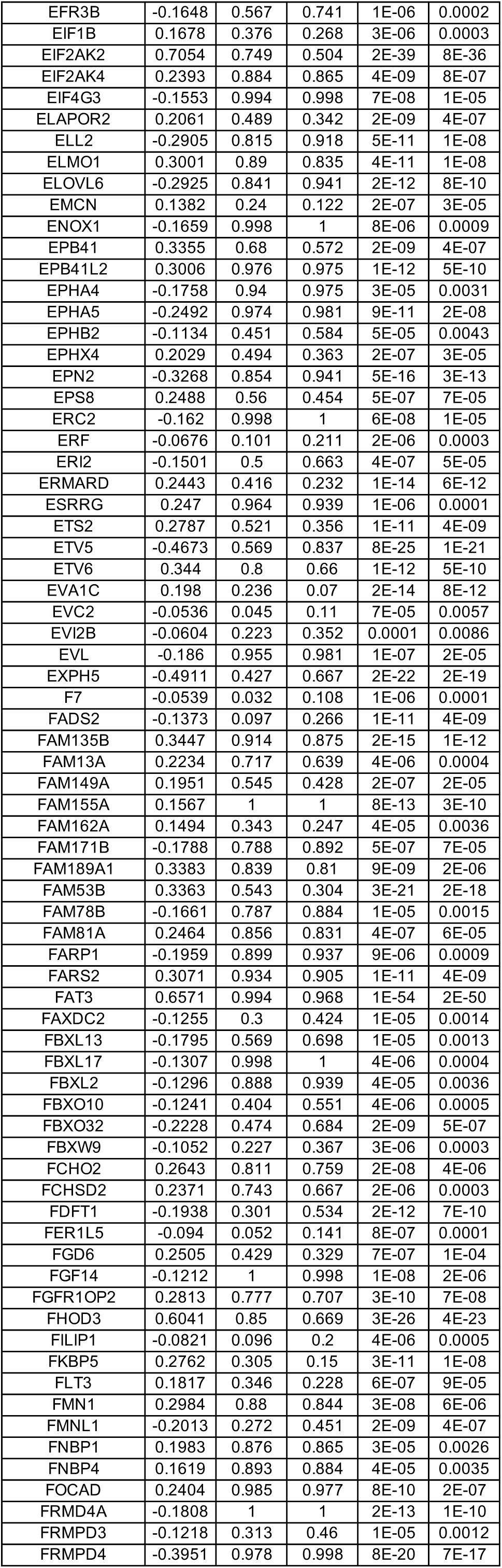
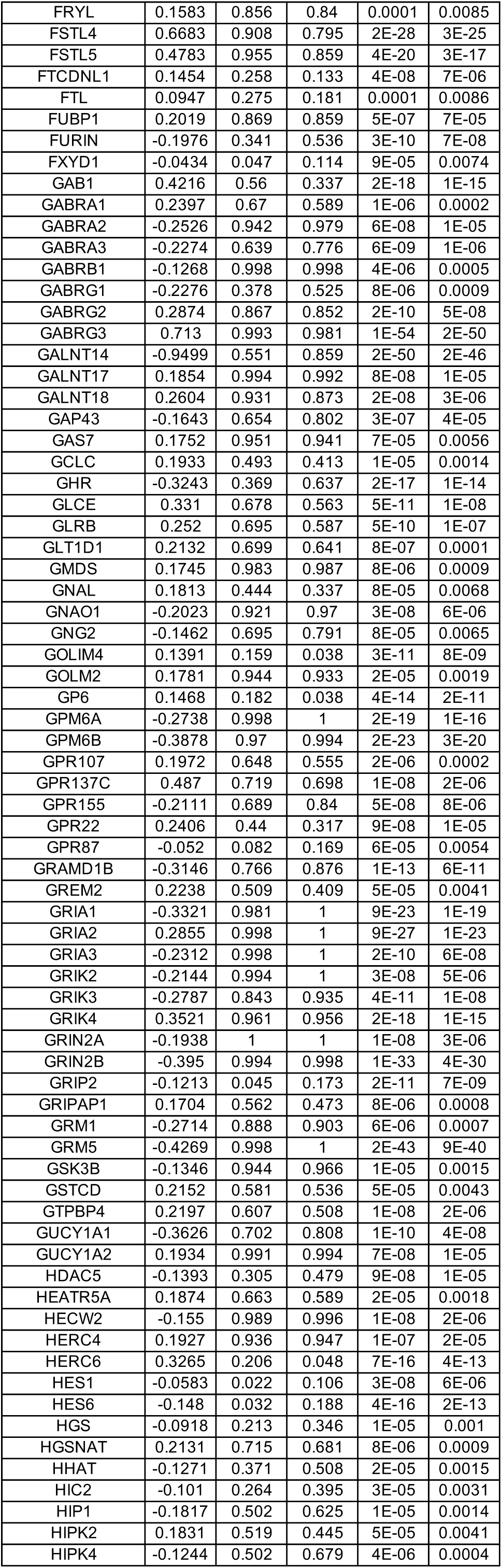
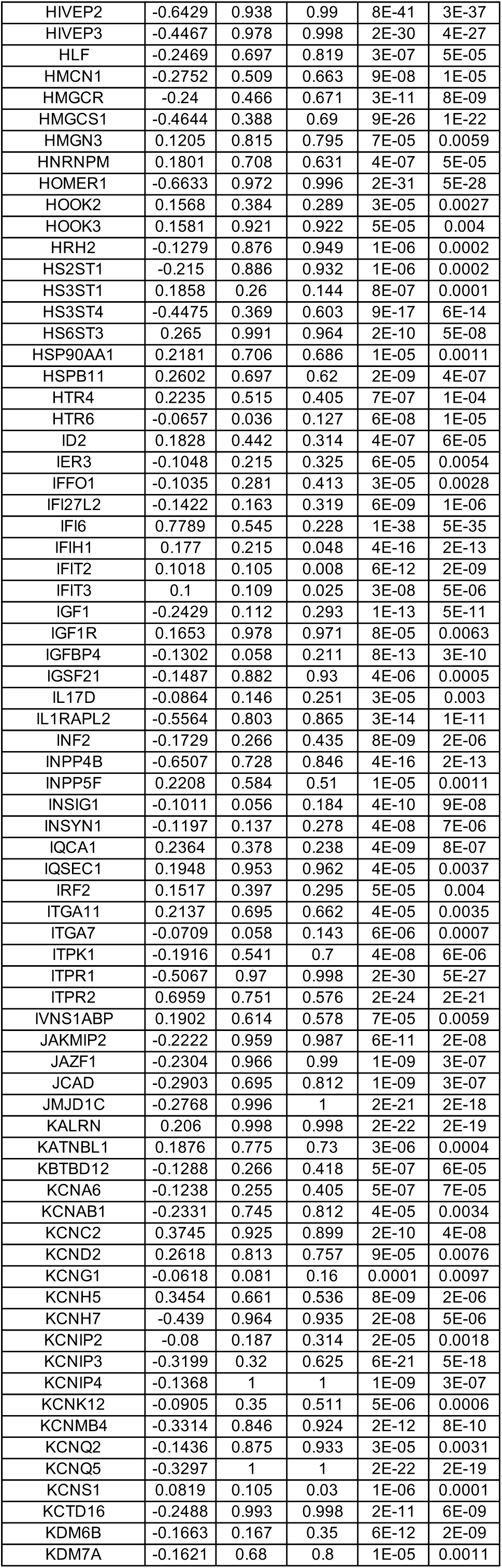
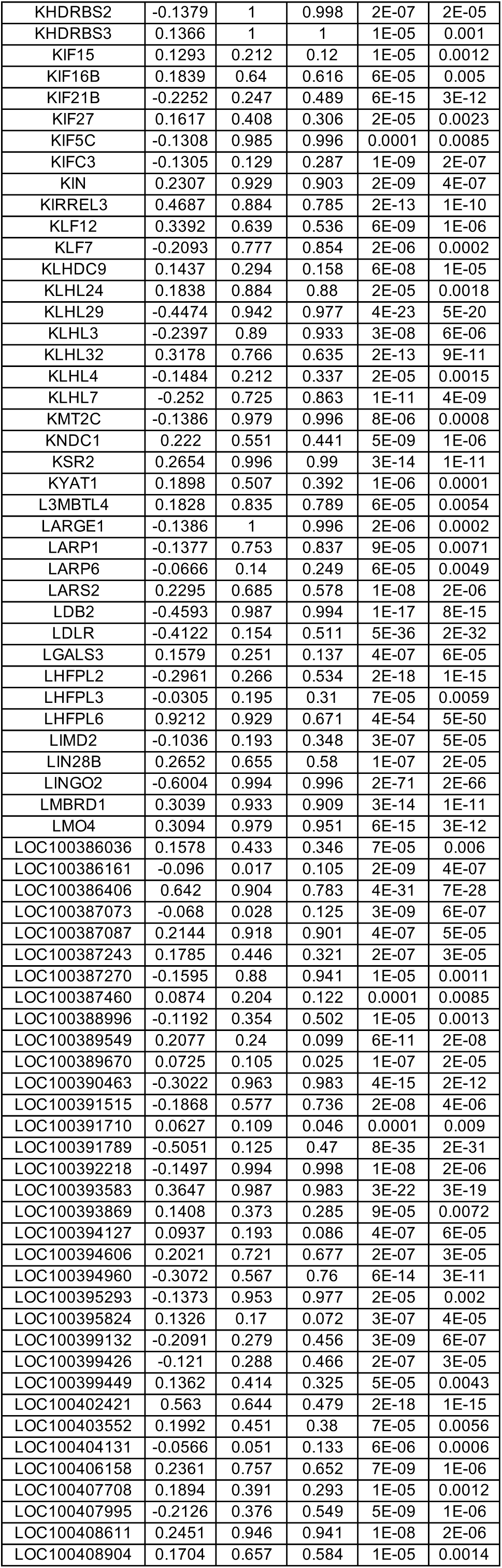
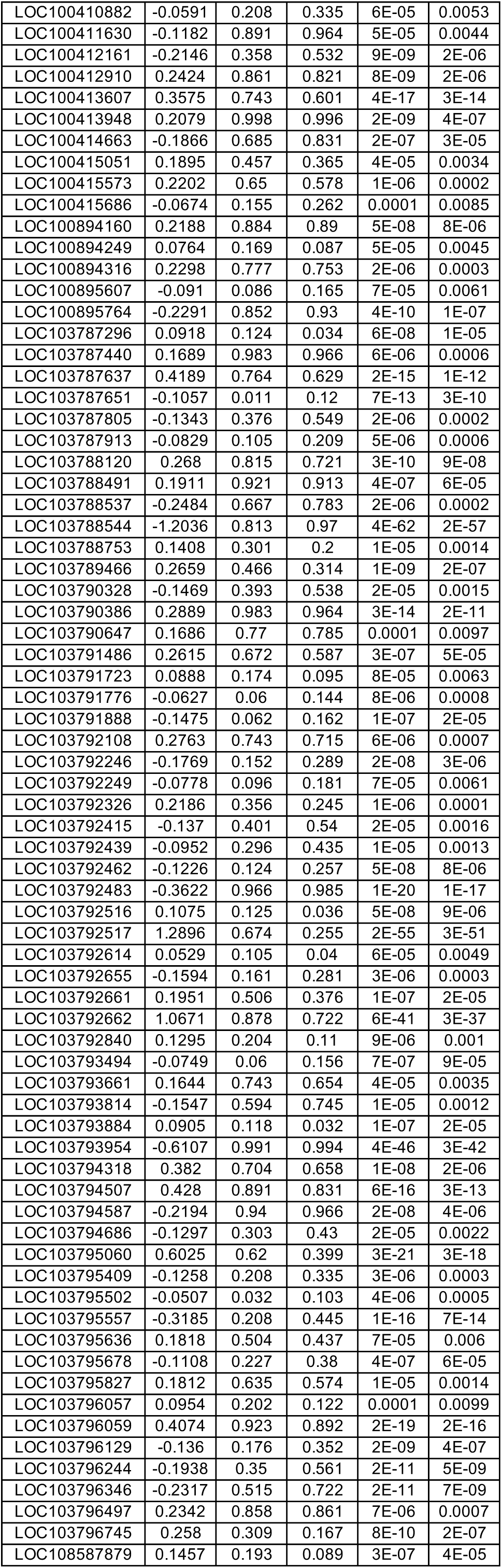
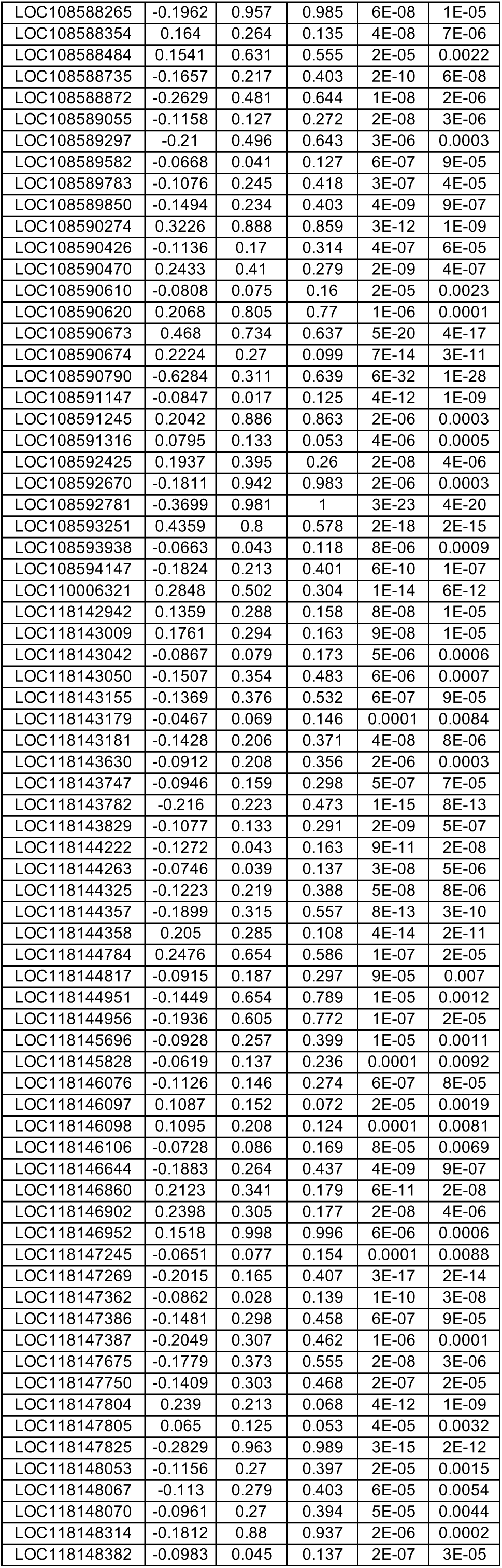
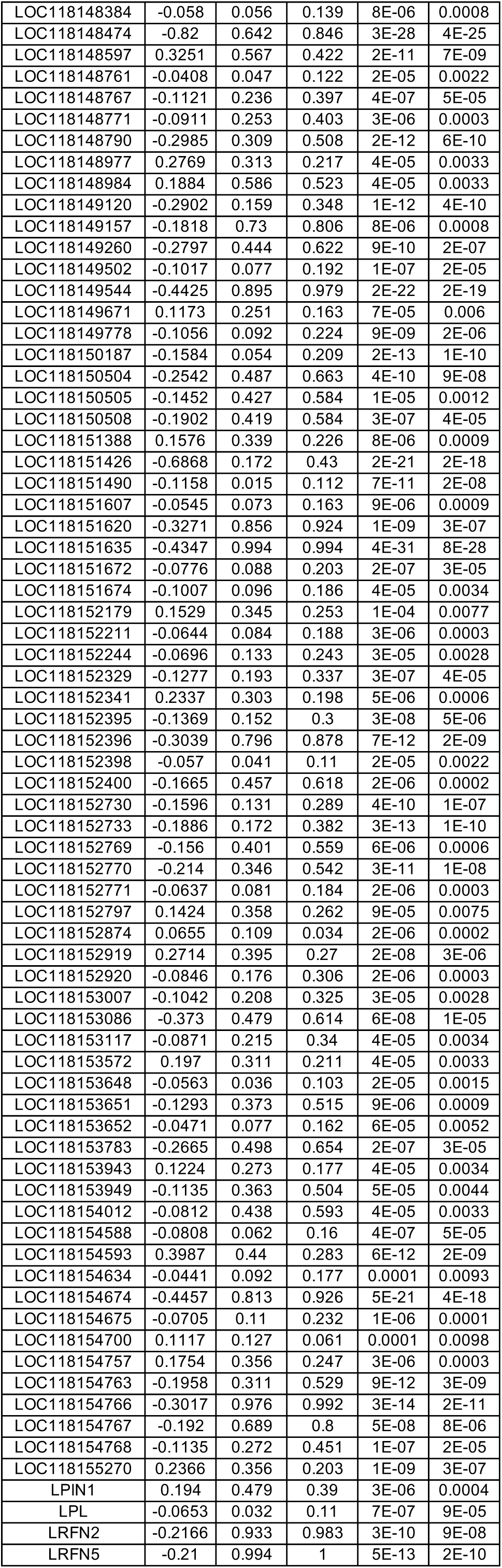
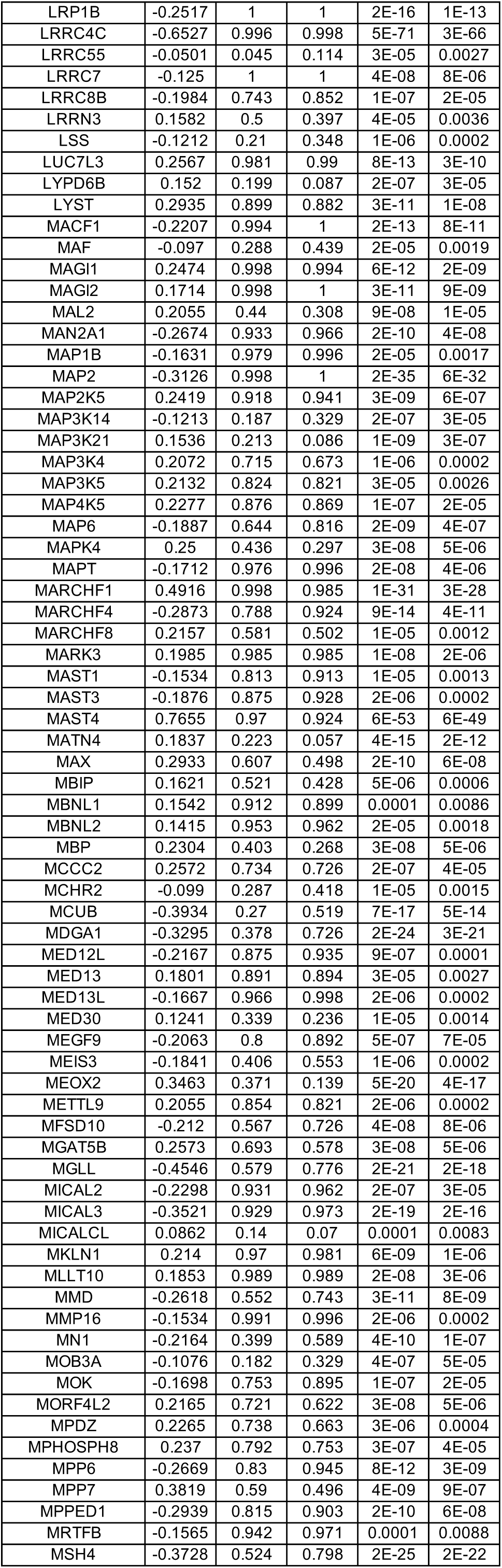
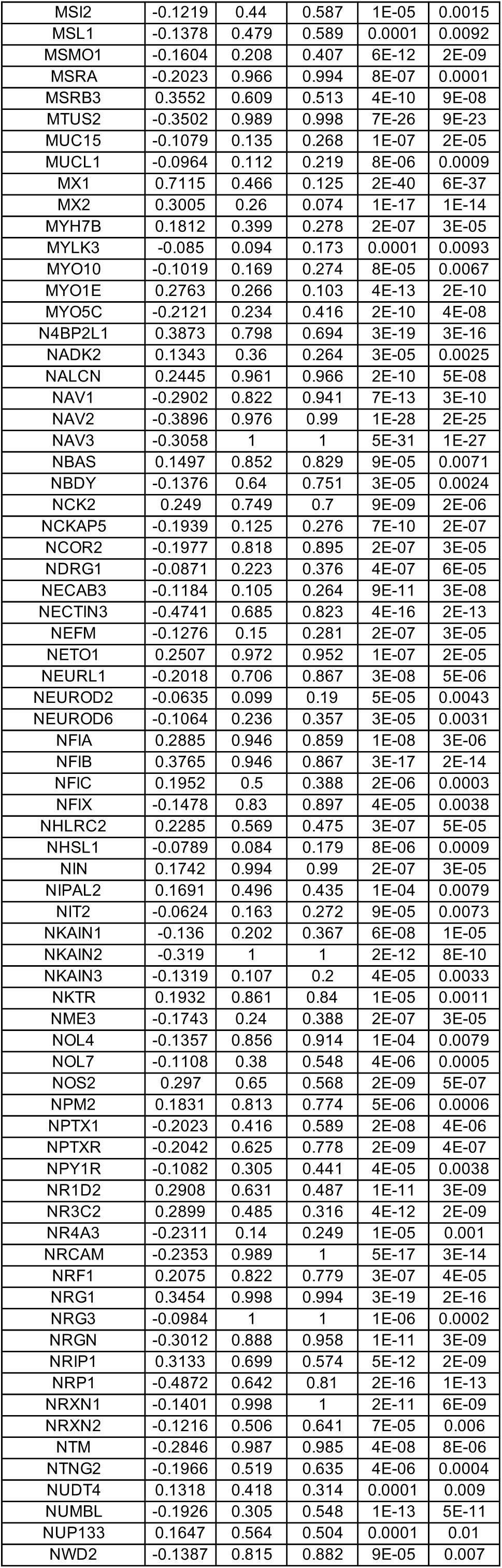
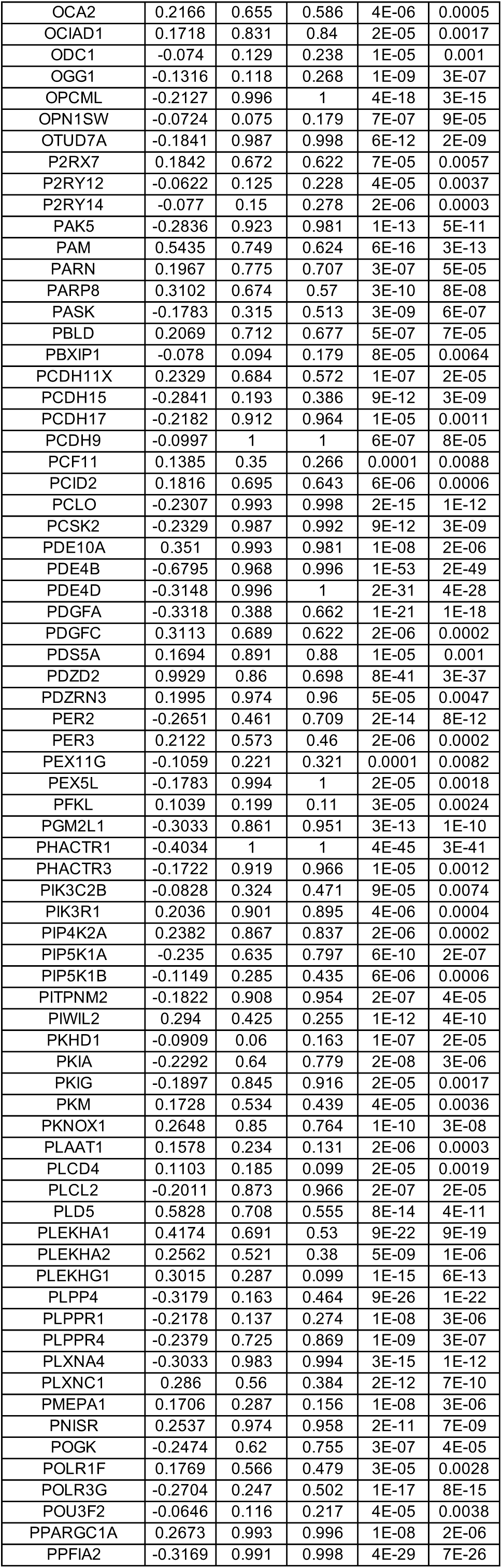
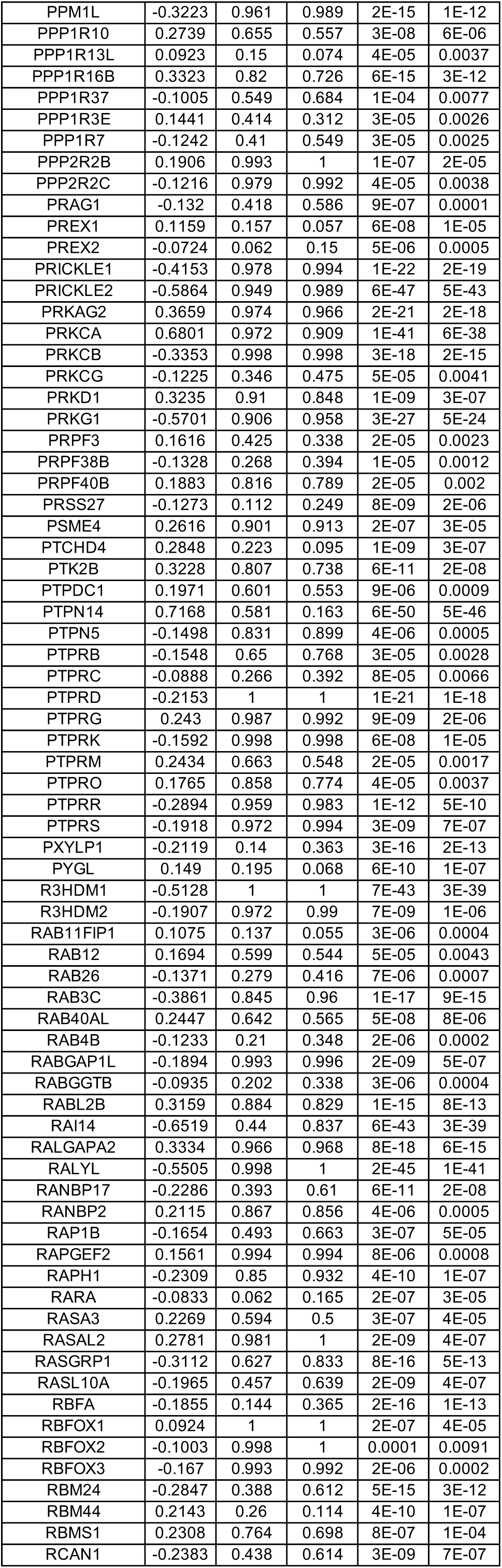
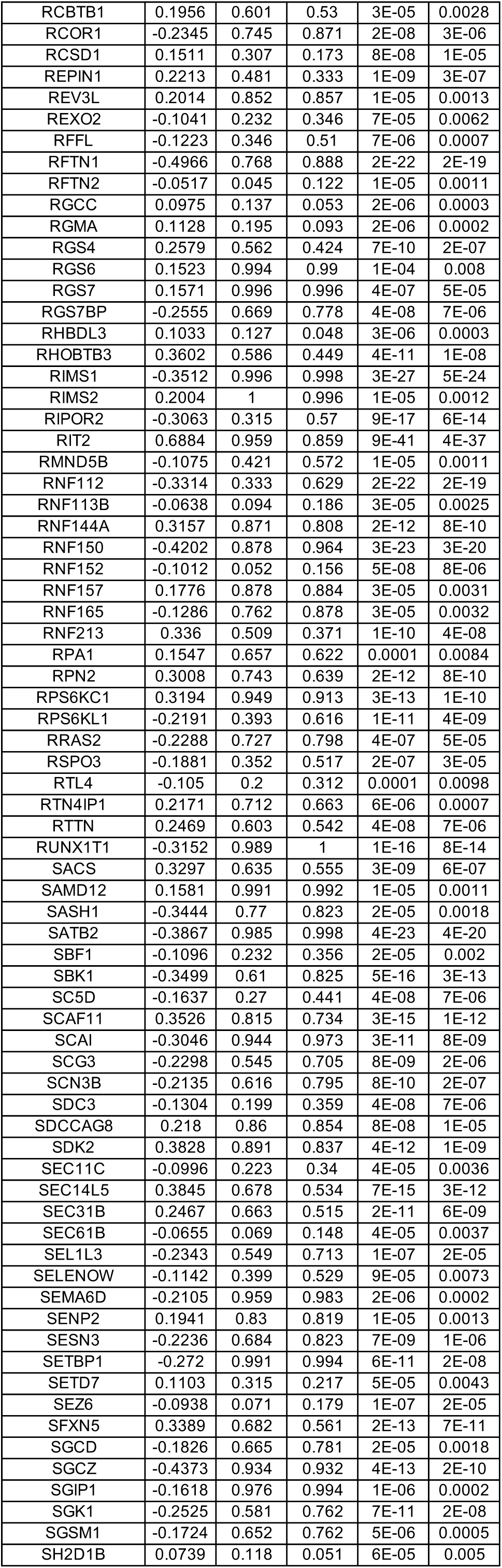
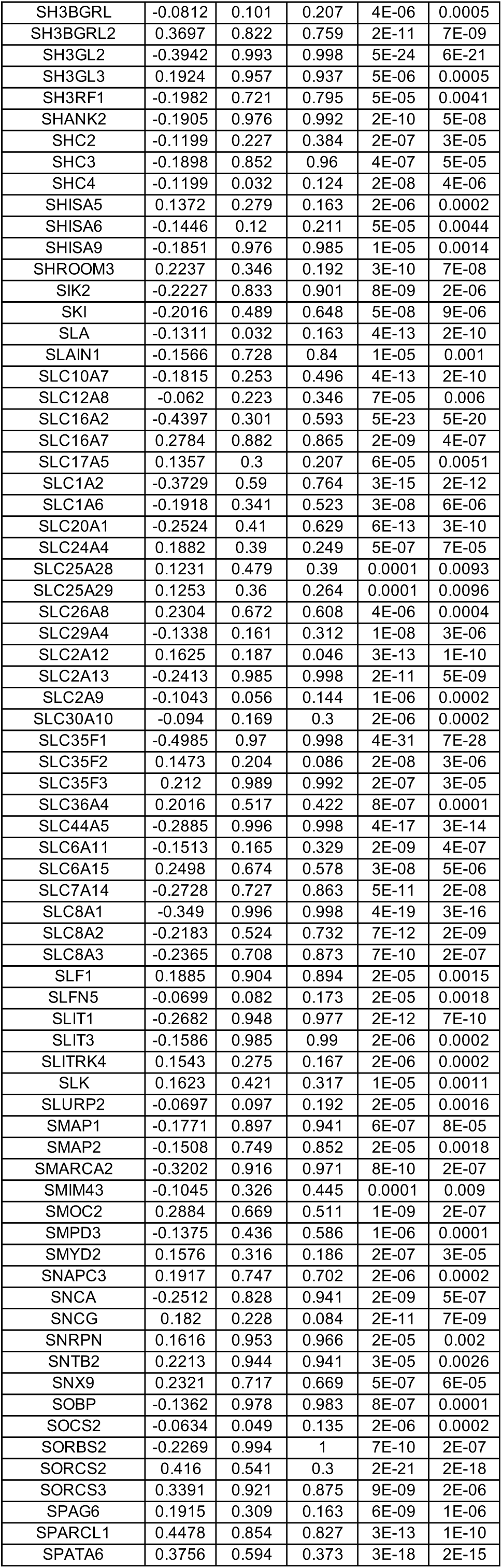
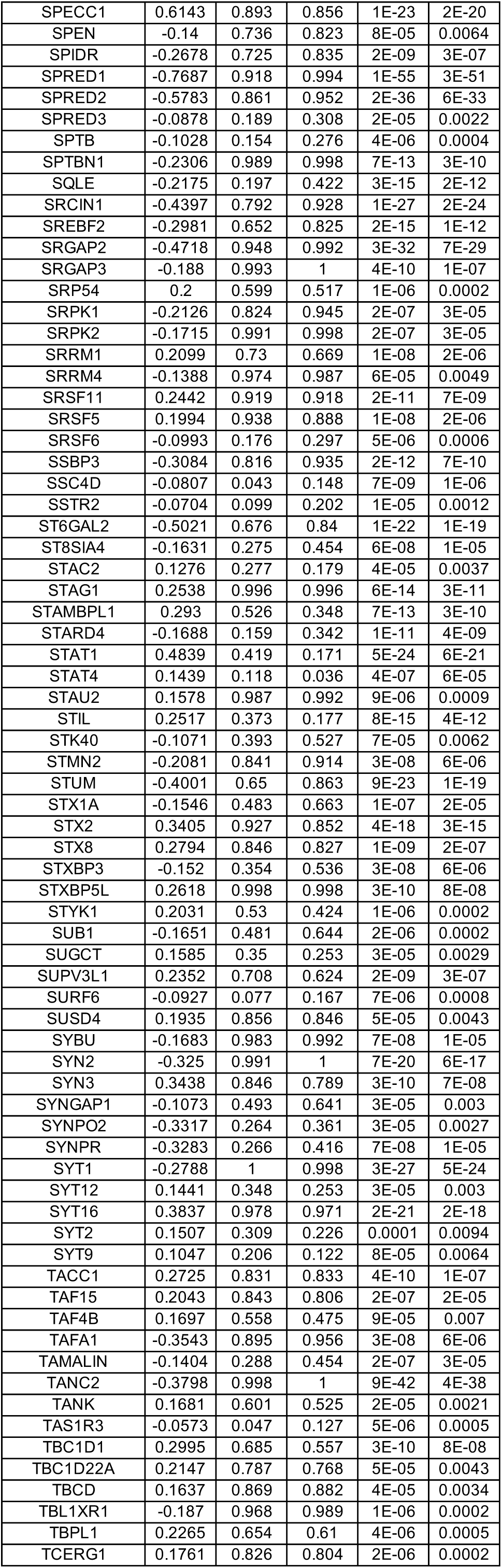
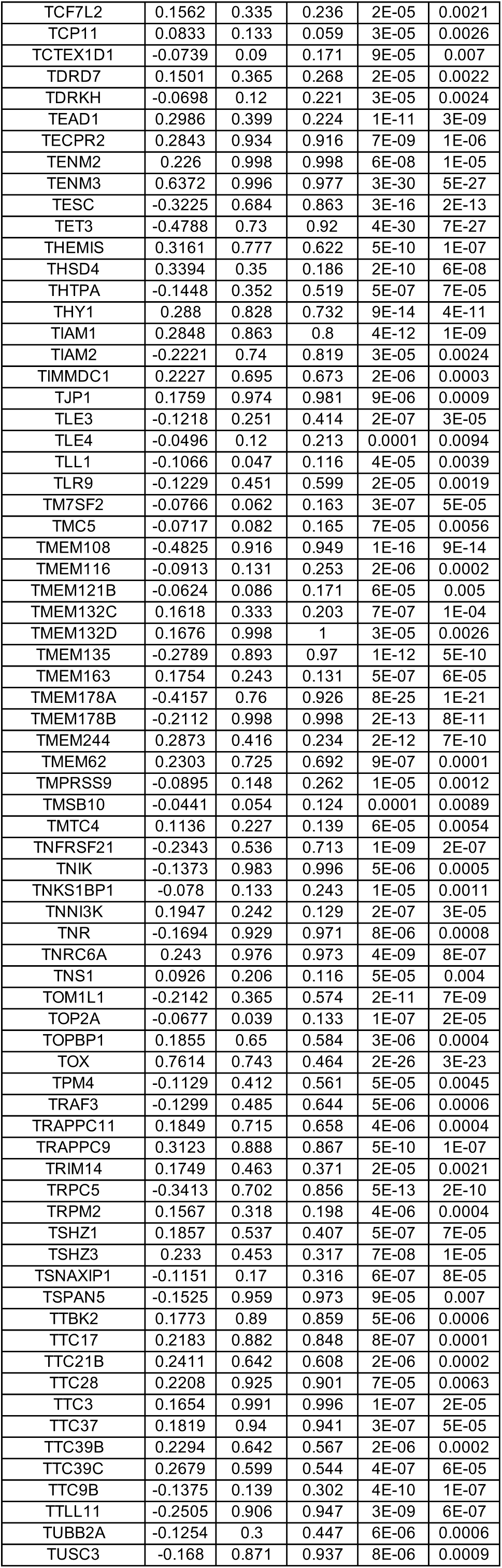
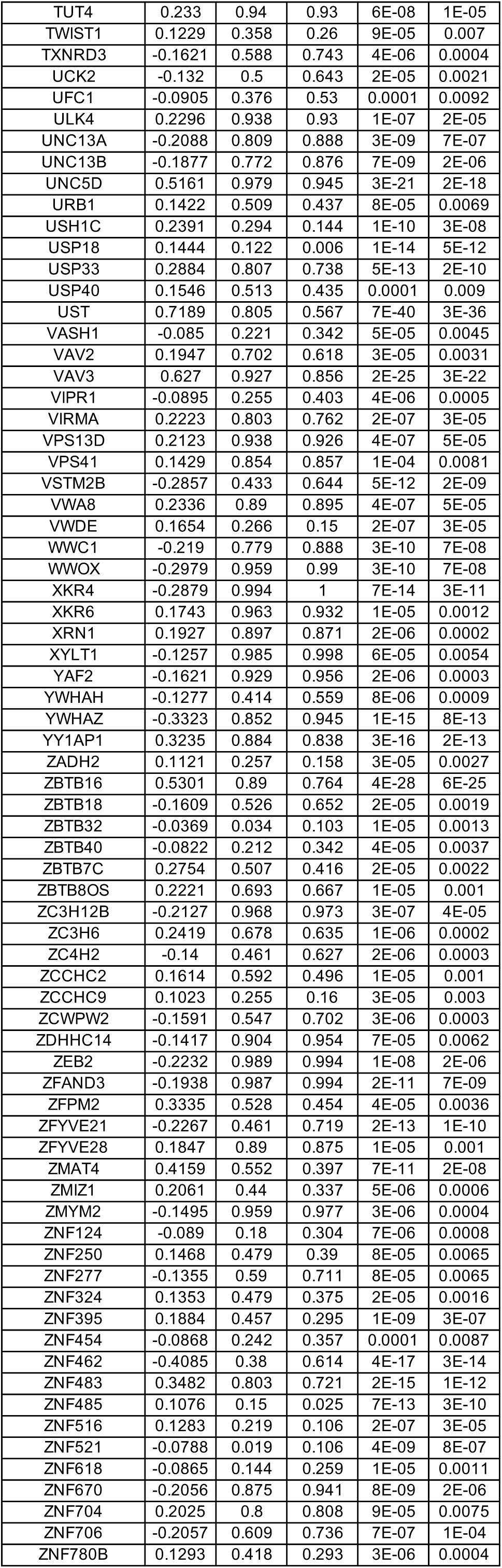
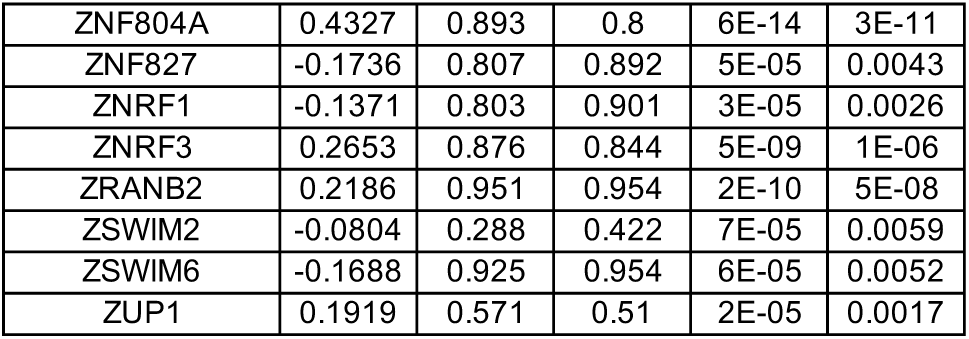
DEGs between wild-type (WT) and MECP2-null (KO) upper layer excitatory neurons (Ex_5) of PFC (adjusted p-value < 0.01)

**Table S19.**
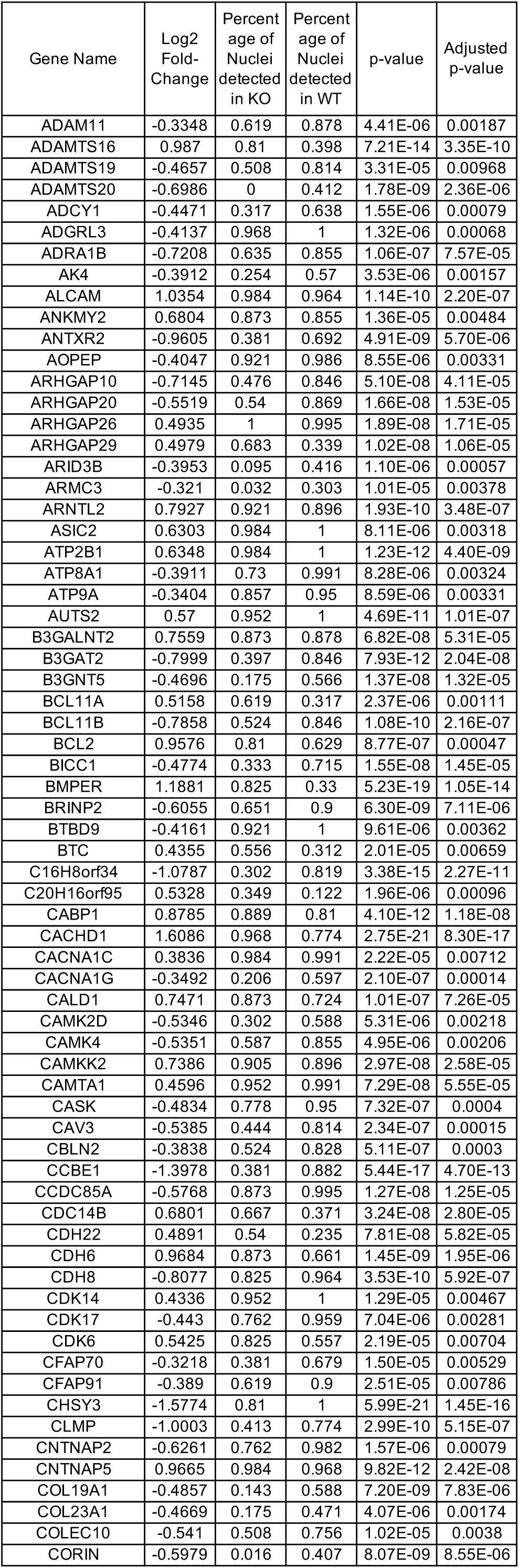
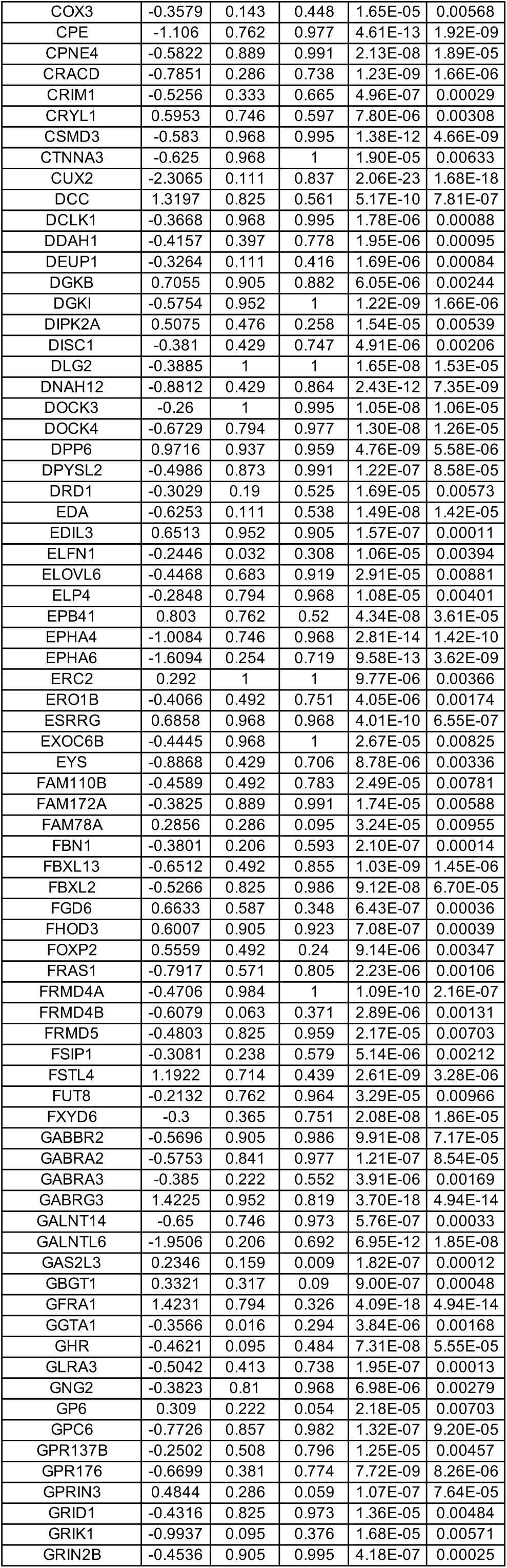
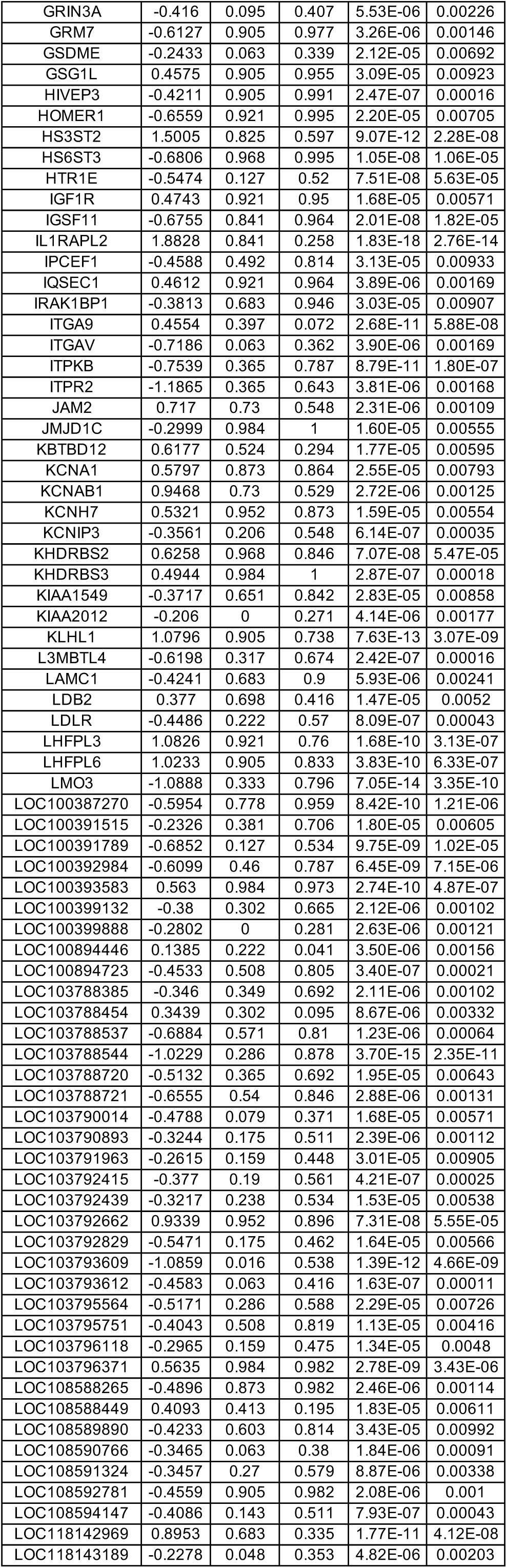
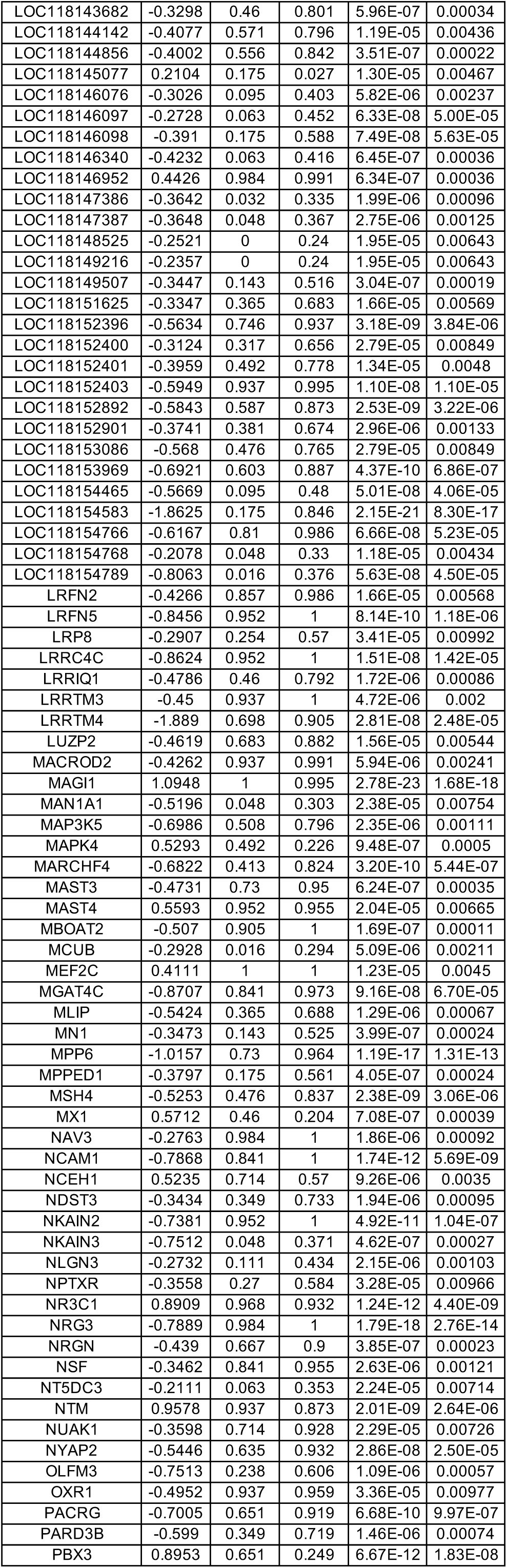
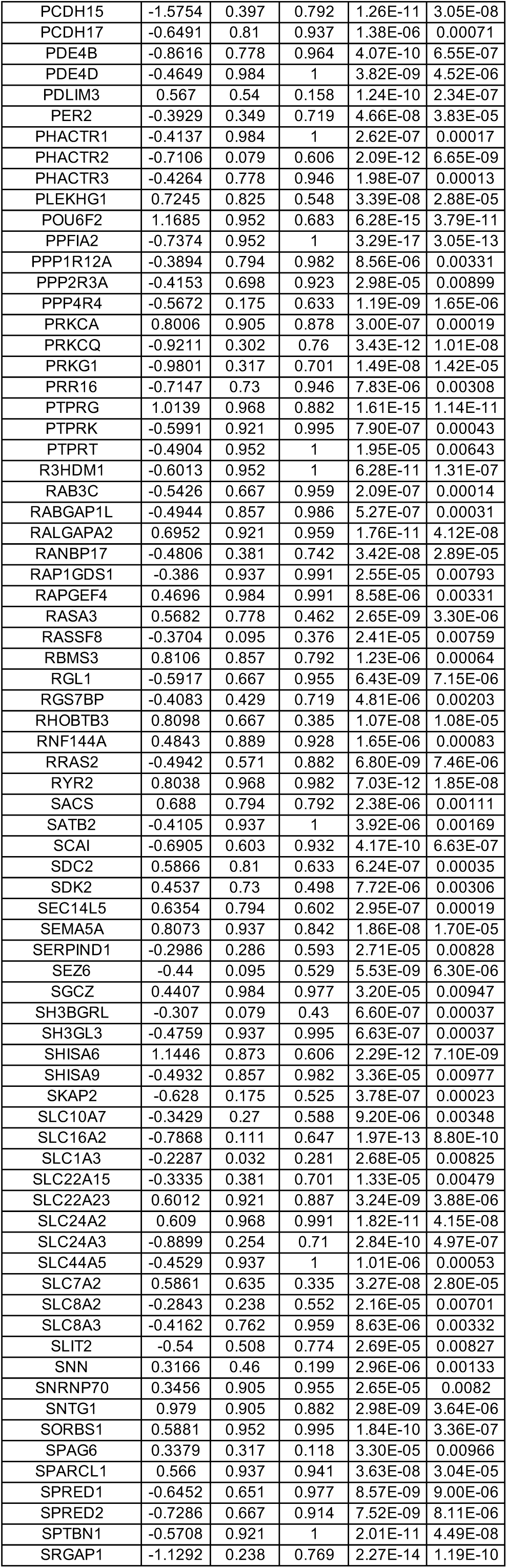
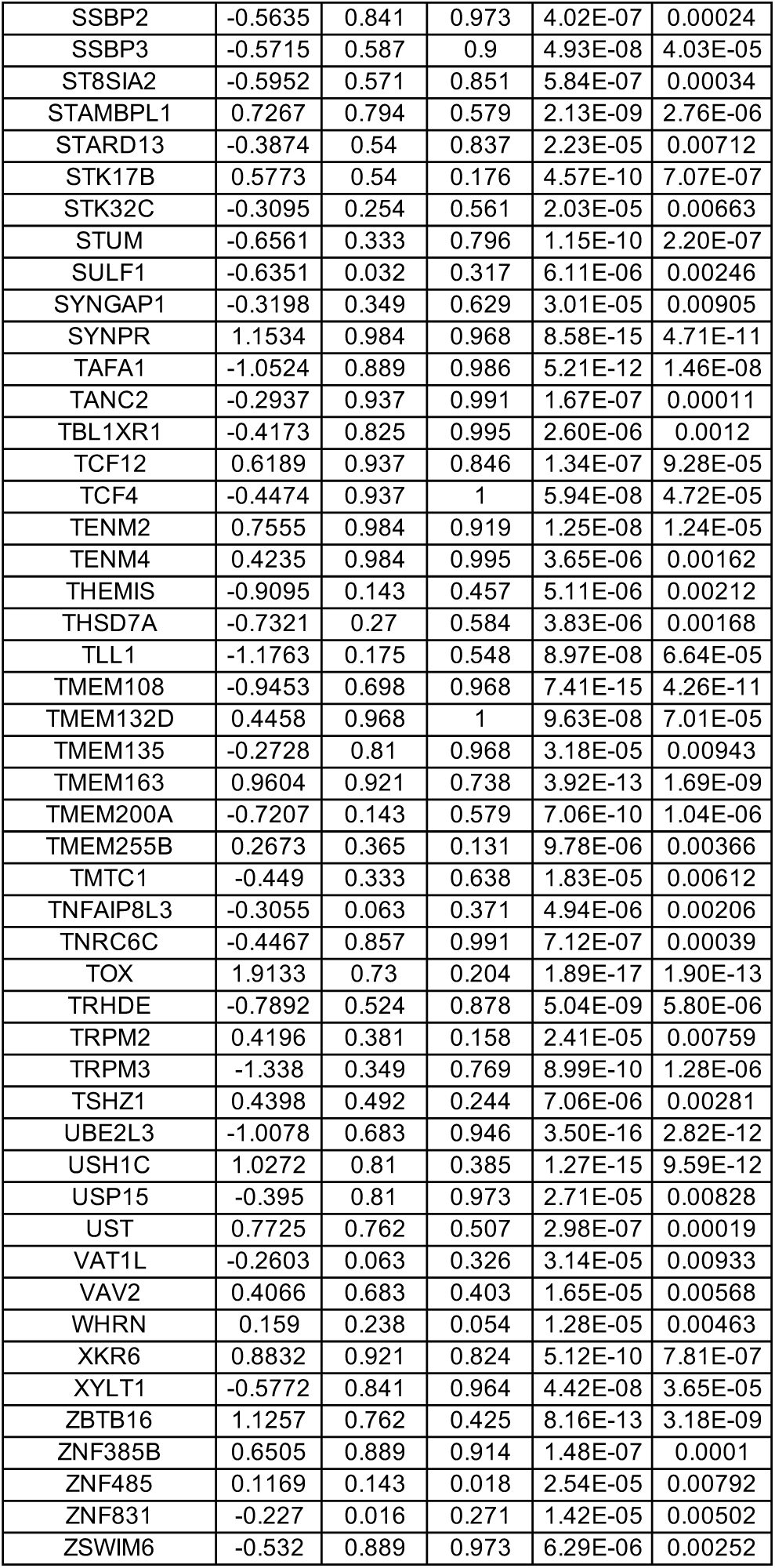
DEGs between wild-type (WT) and MECP2-null (KO) upper layer excitatory neurons (Ex_6) of PFC (adjusted p-value < 0.01)

**Table S20.**
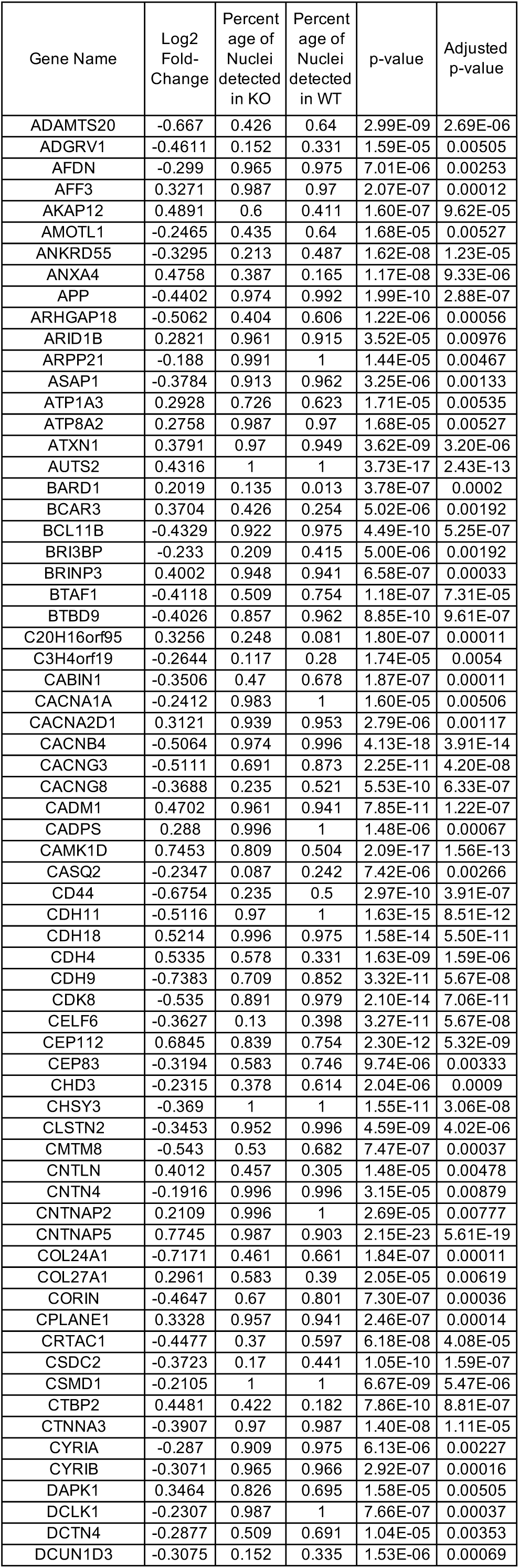
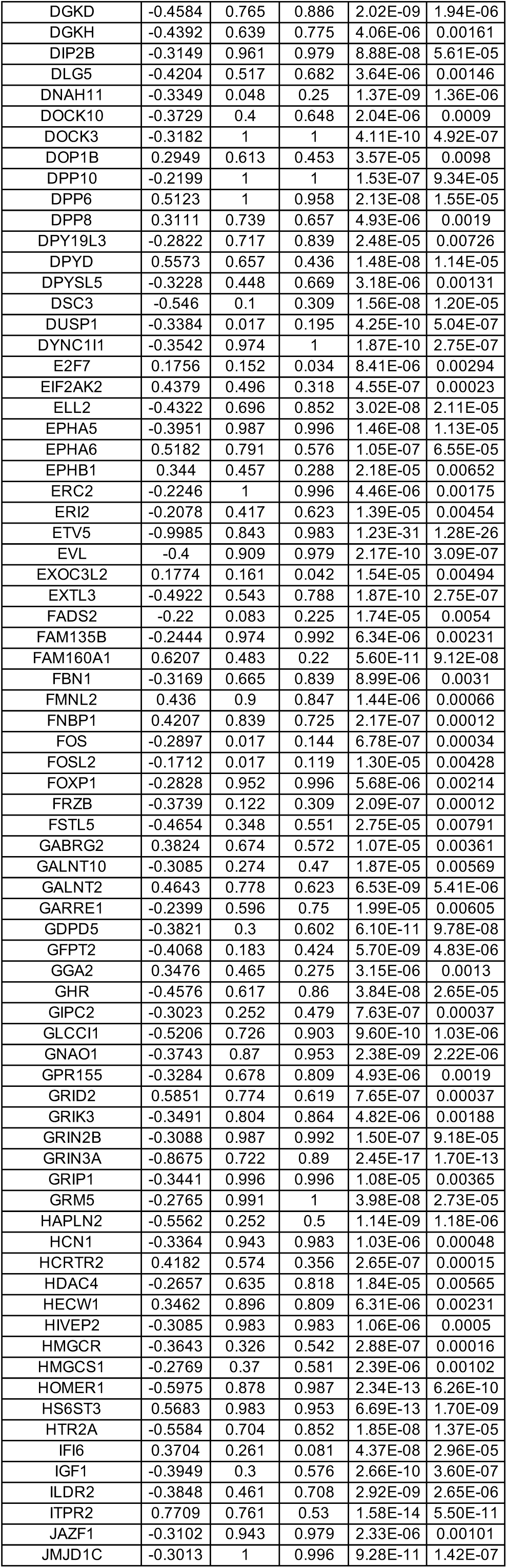
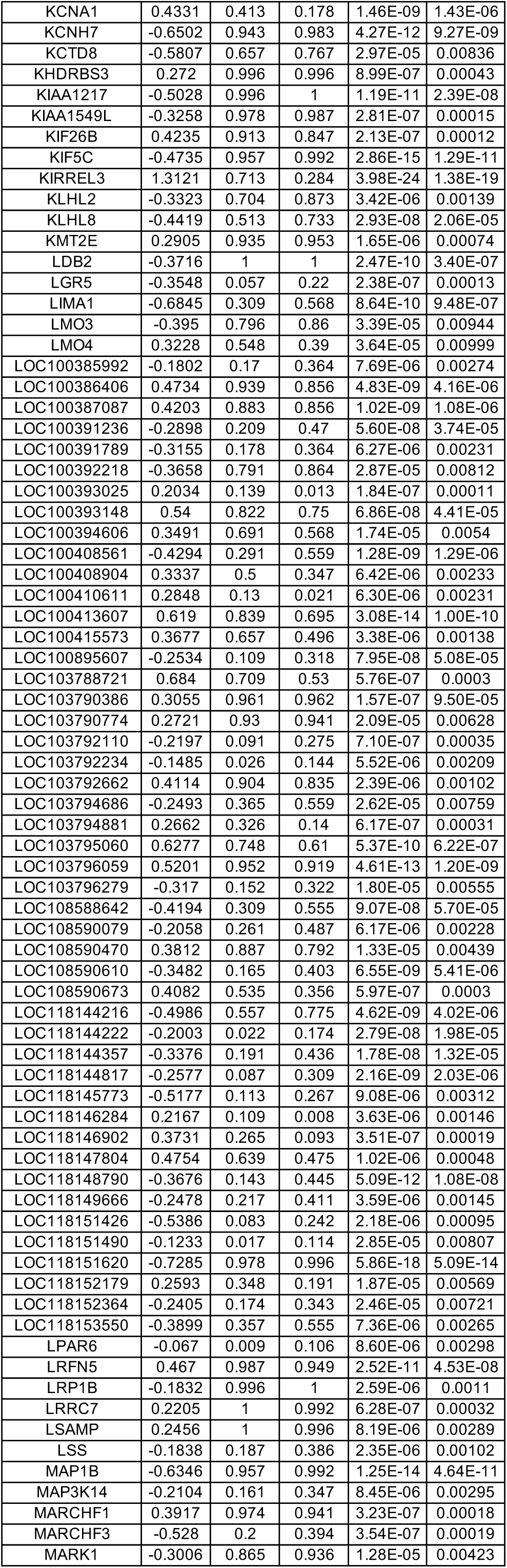
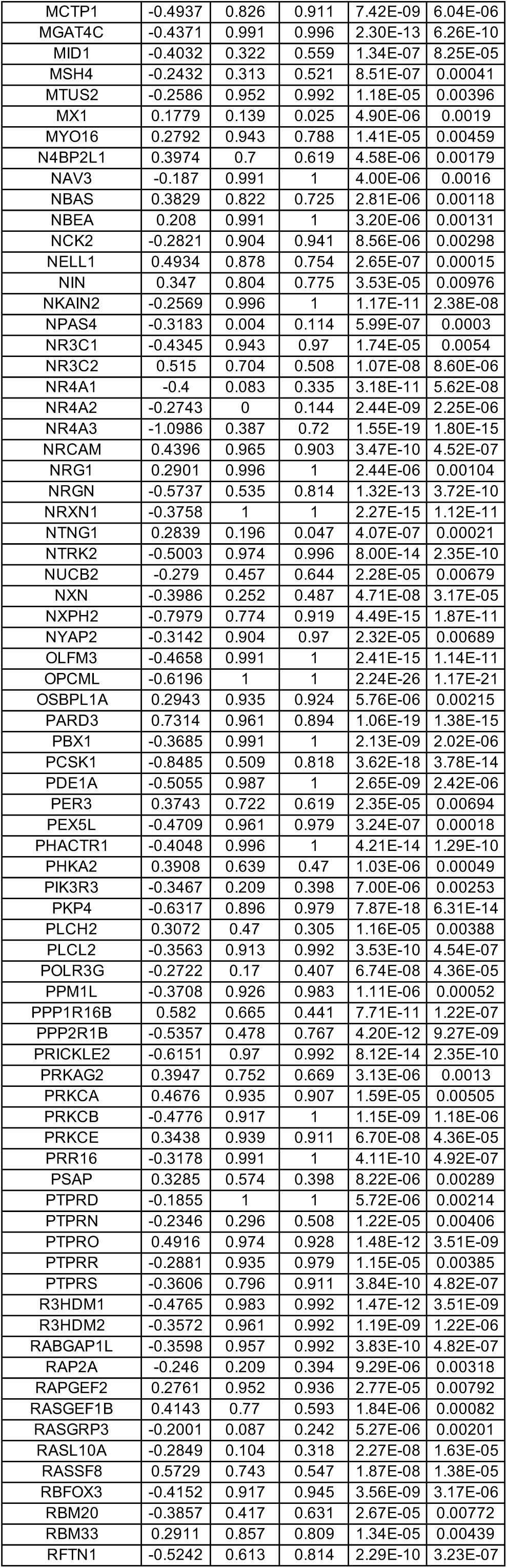
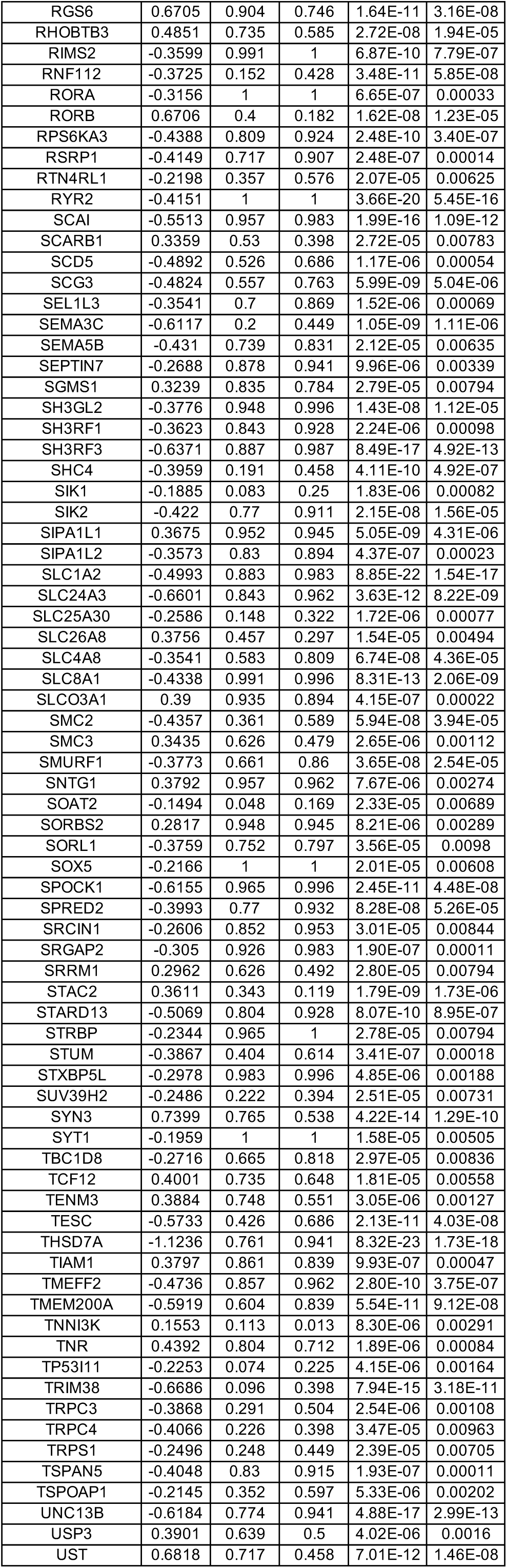
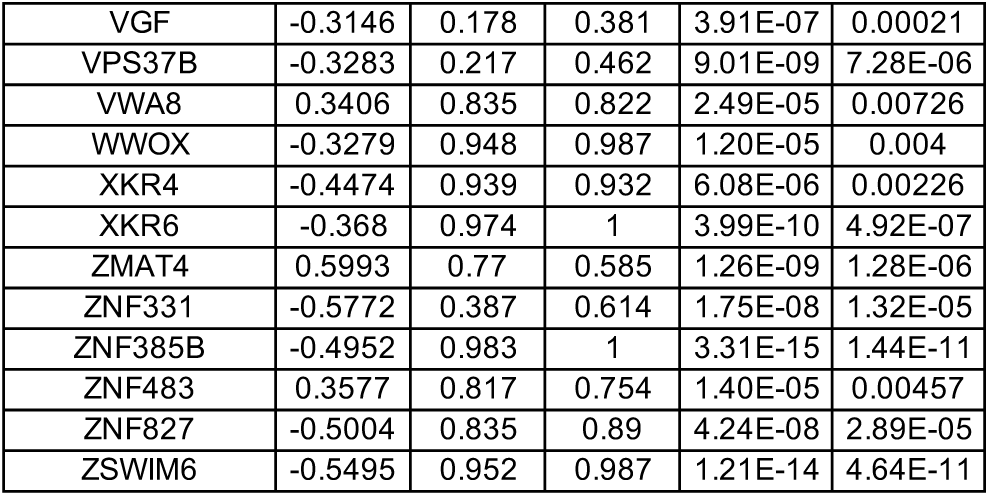
DEGs between wild-type (WT) and MECP2-null (KO) upper layer excitatory neurons (Ex_7) of PFC (adjusted p-value < 0.01)

**Table S21.**
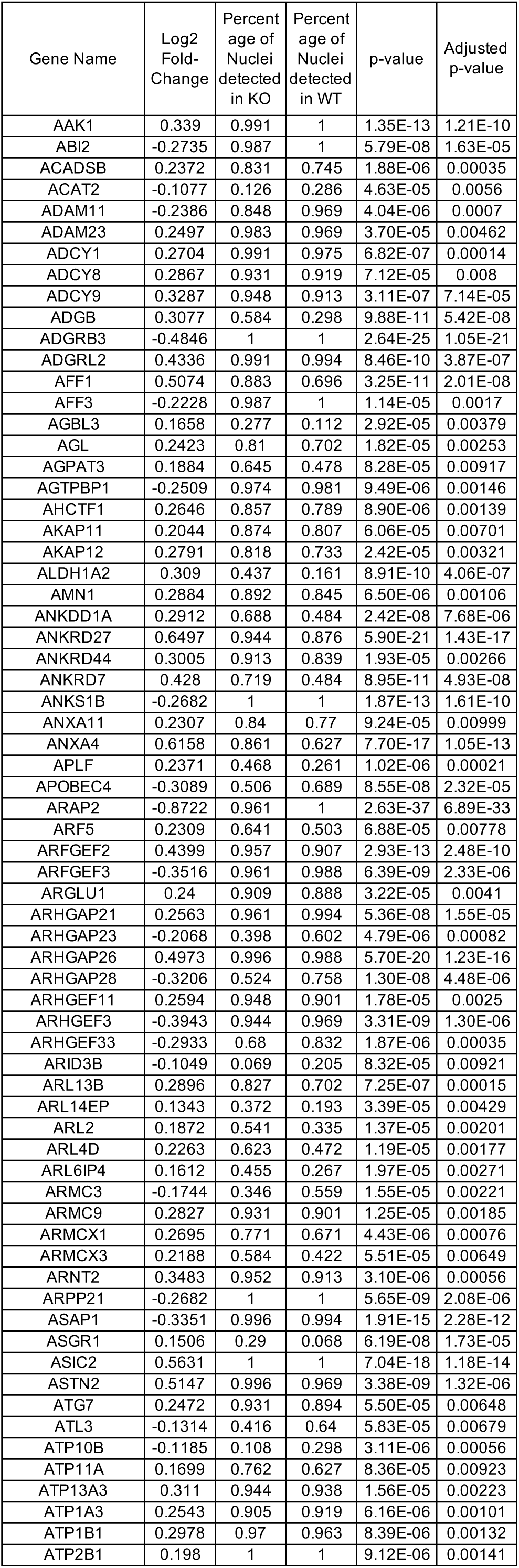
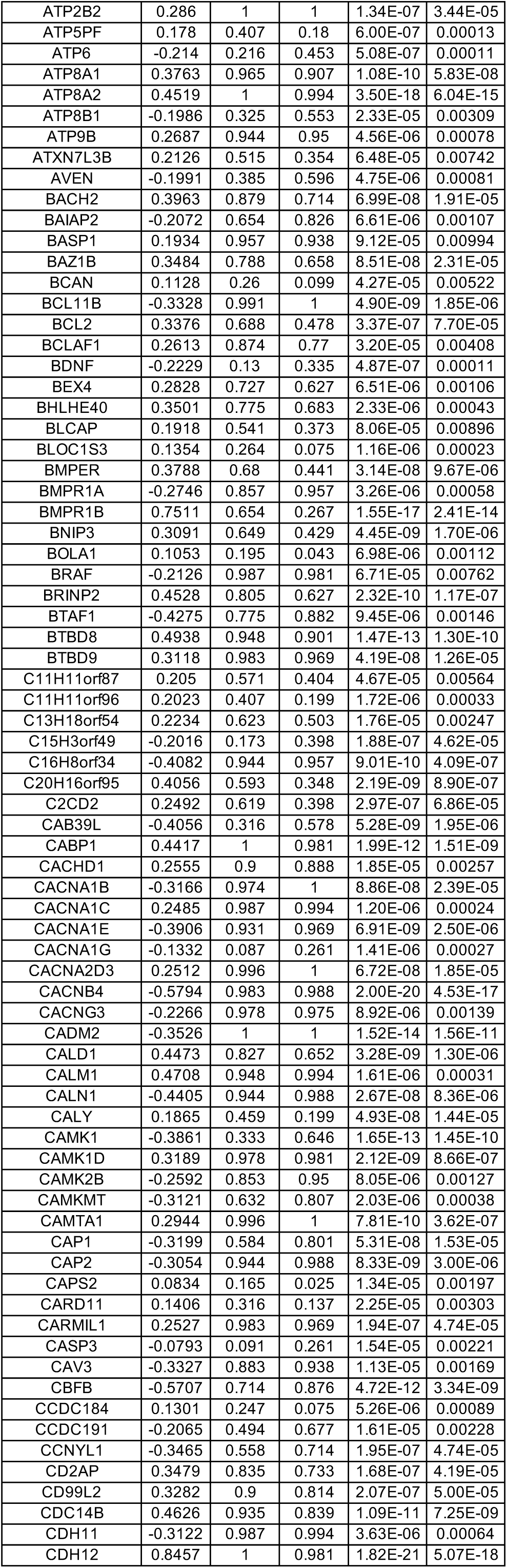
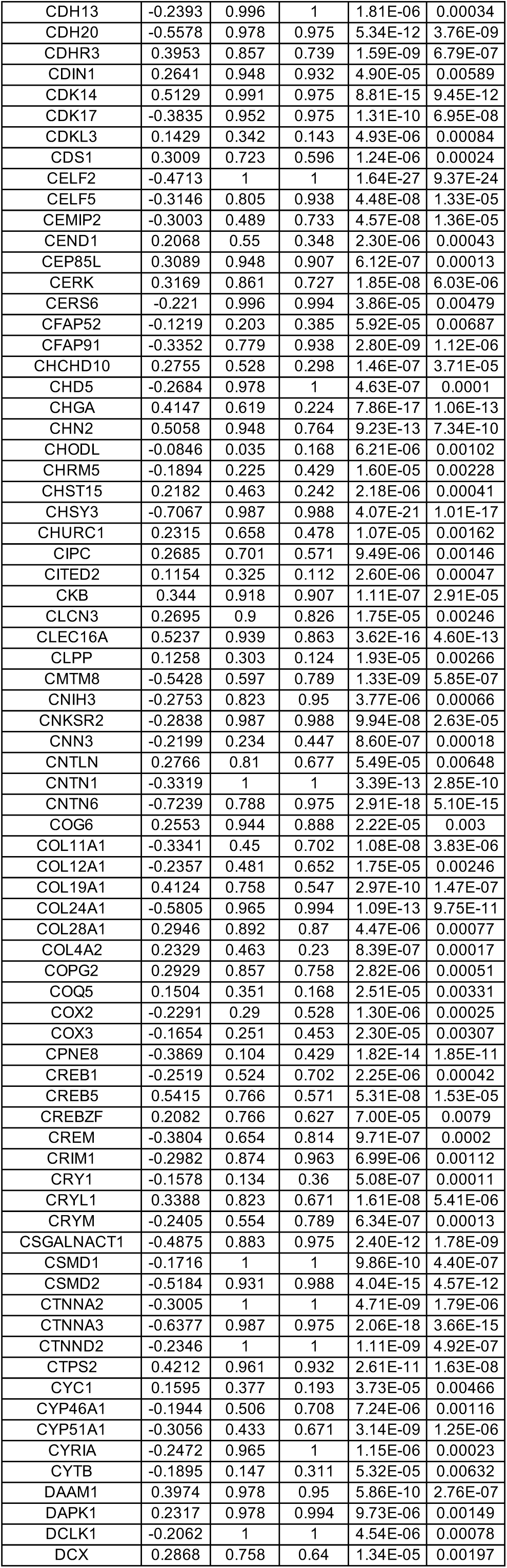
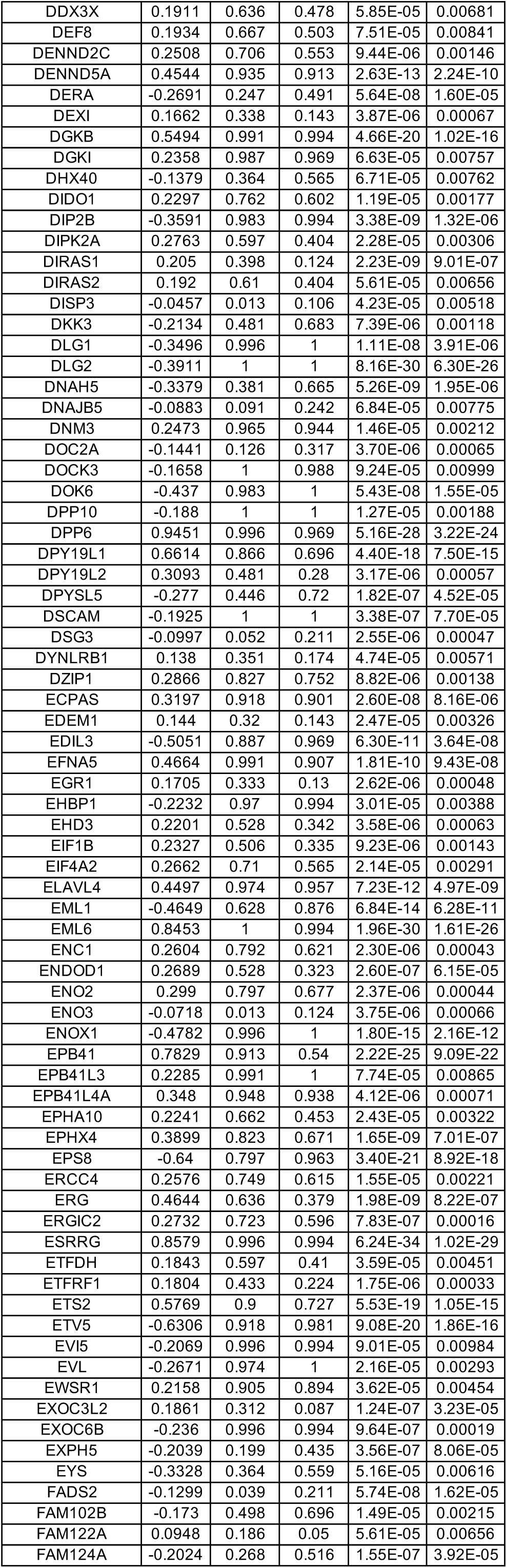
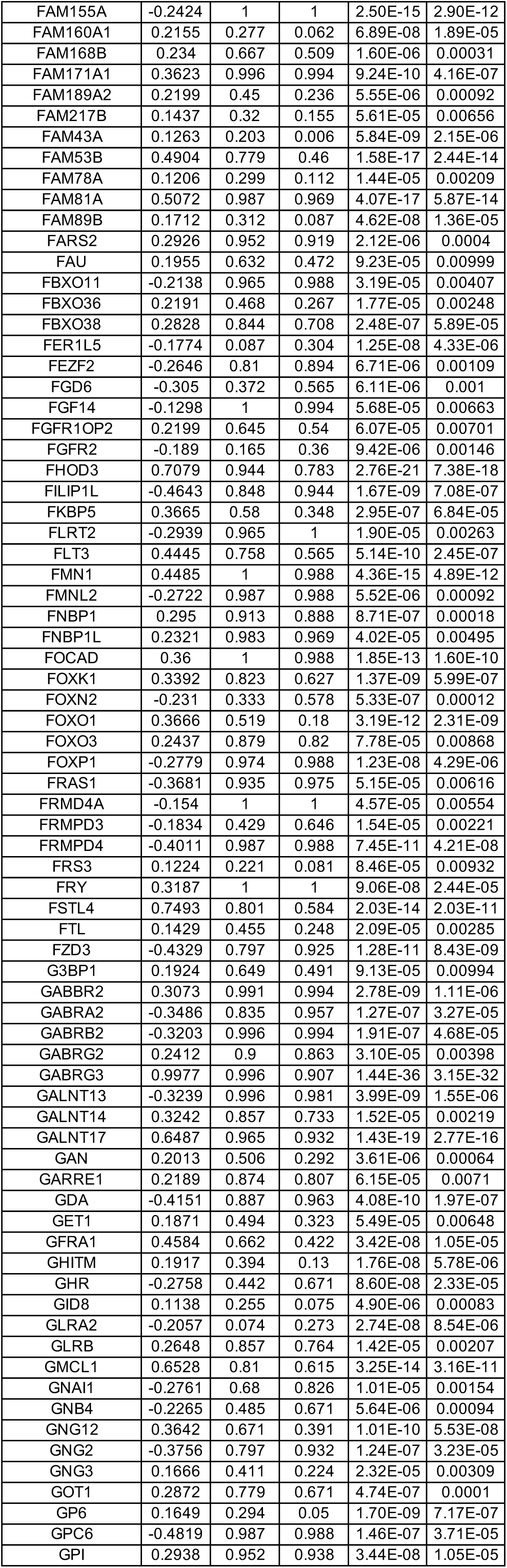
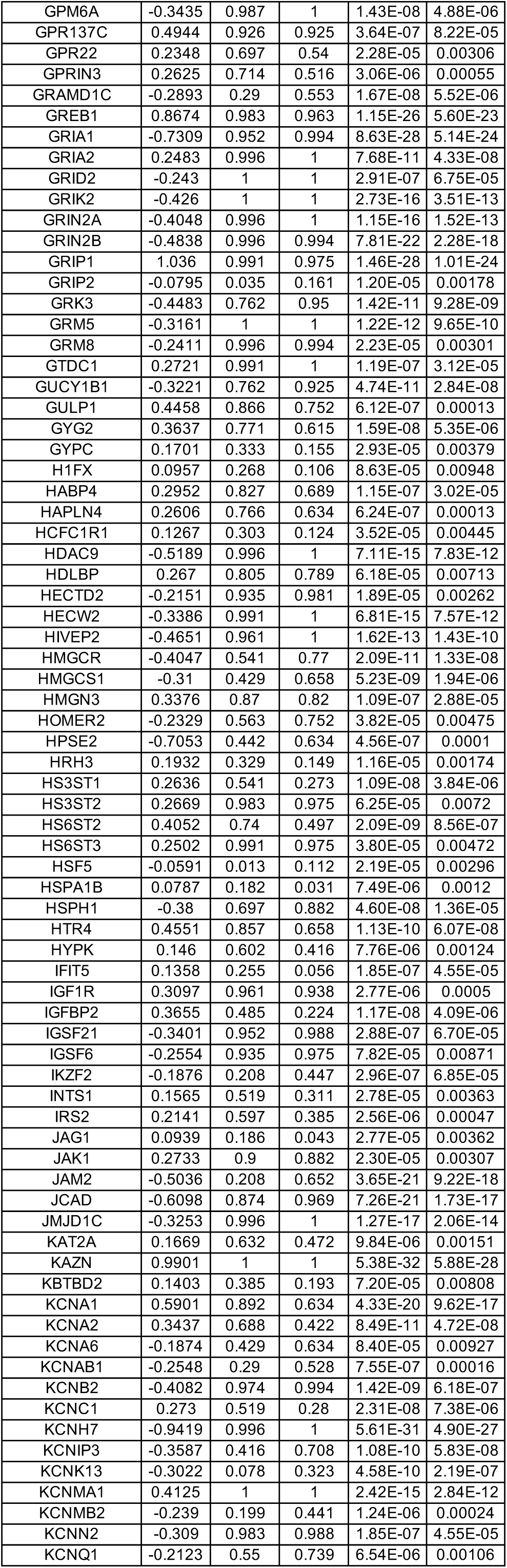
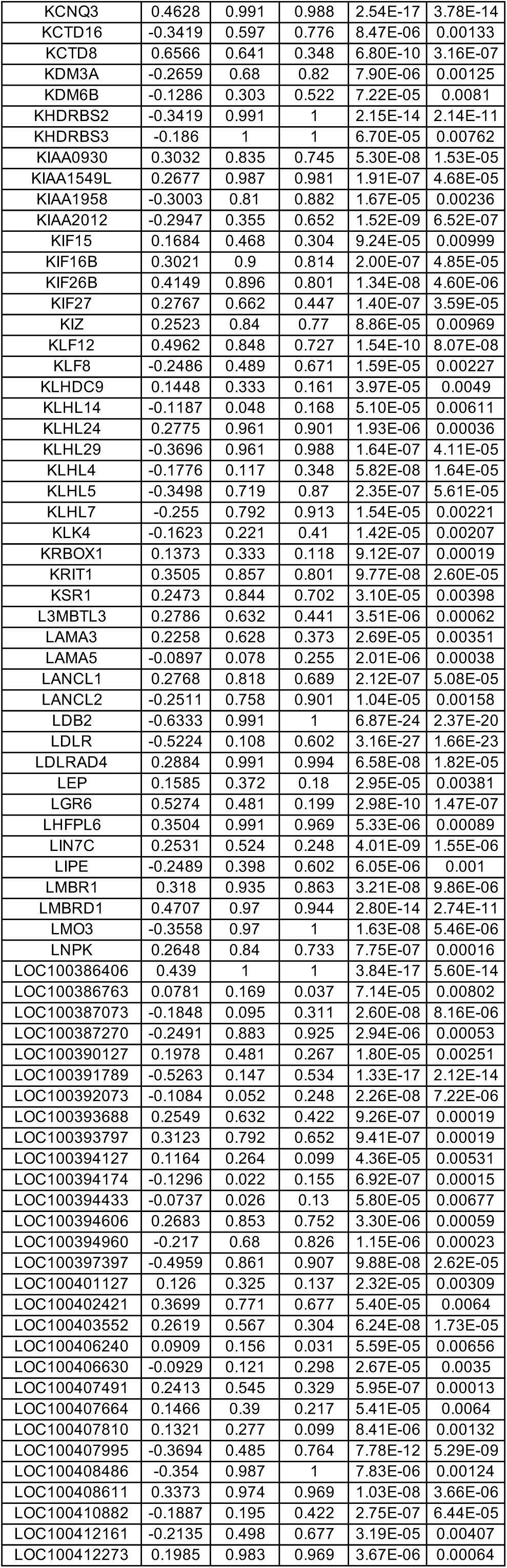
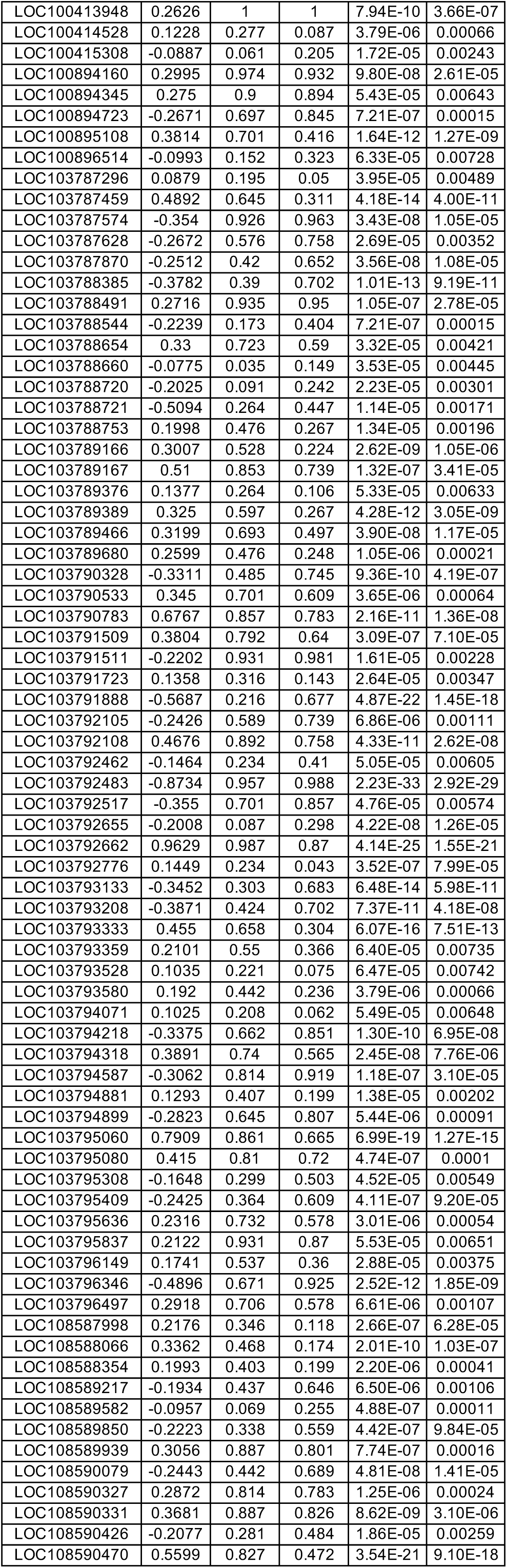
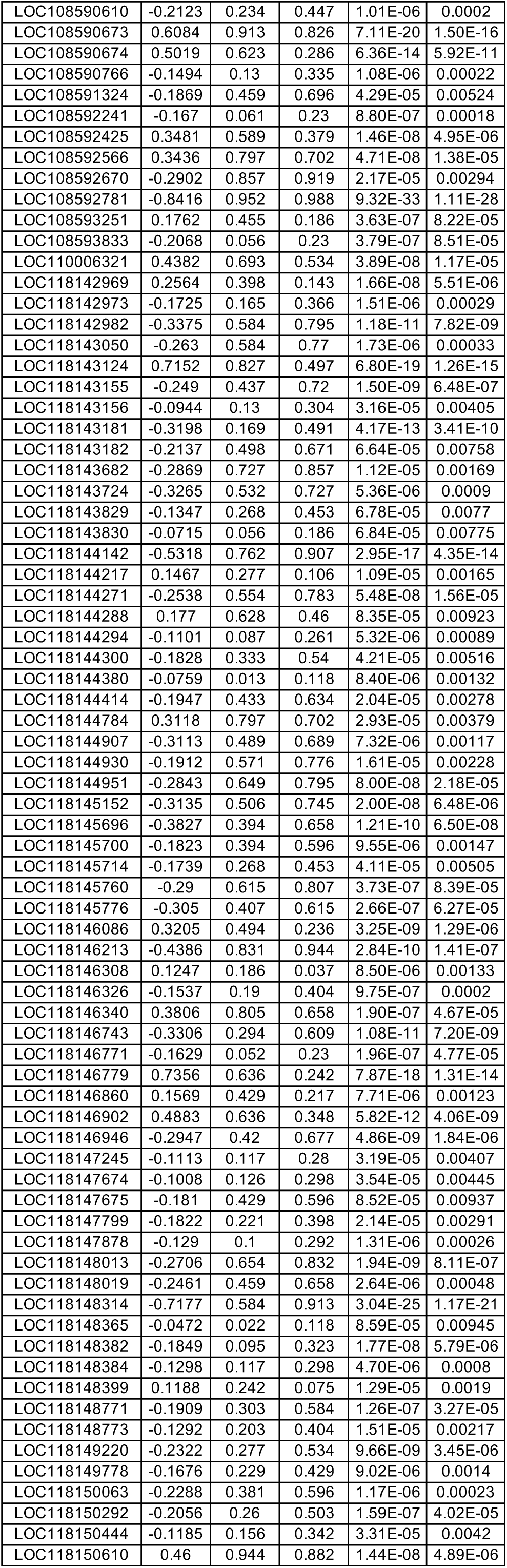
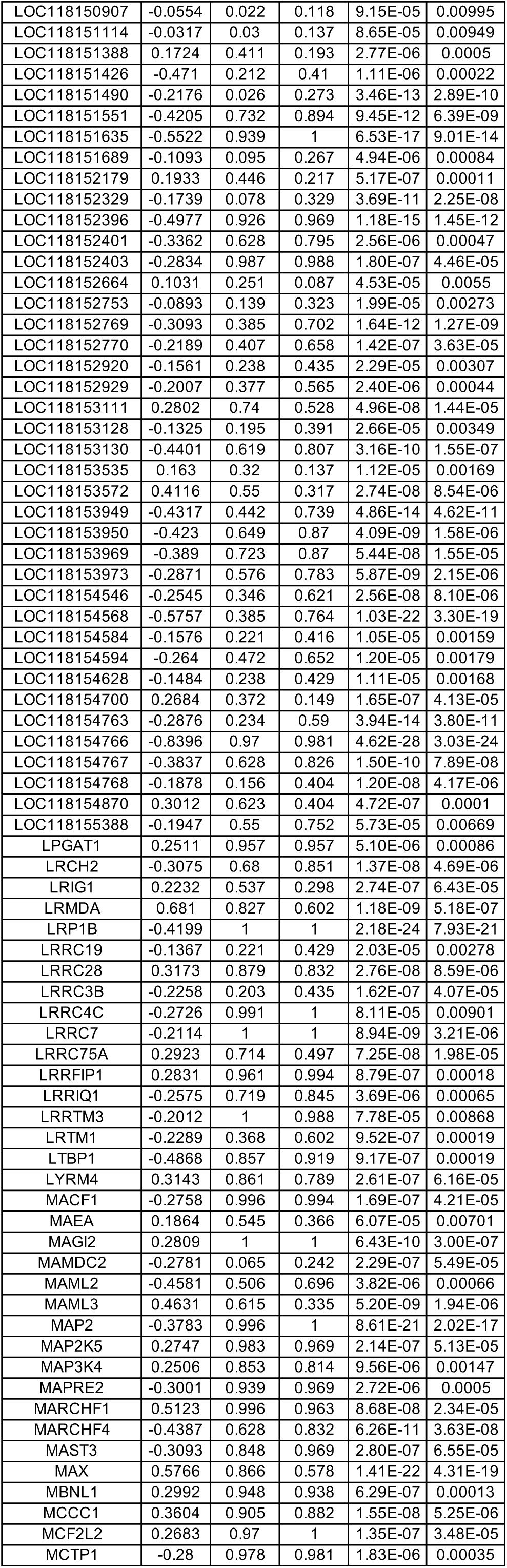
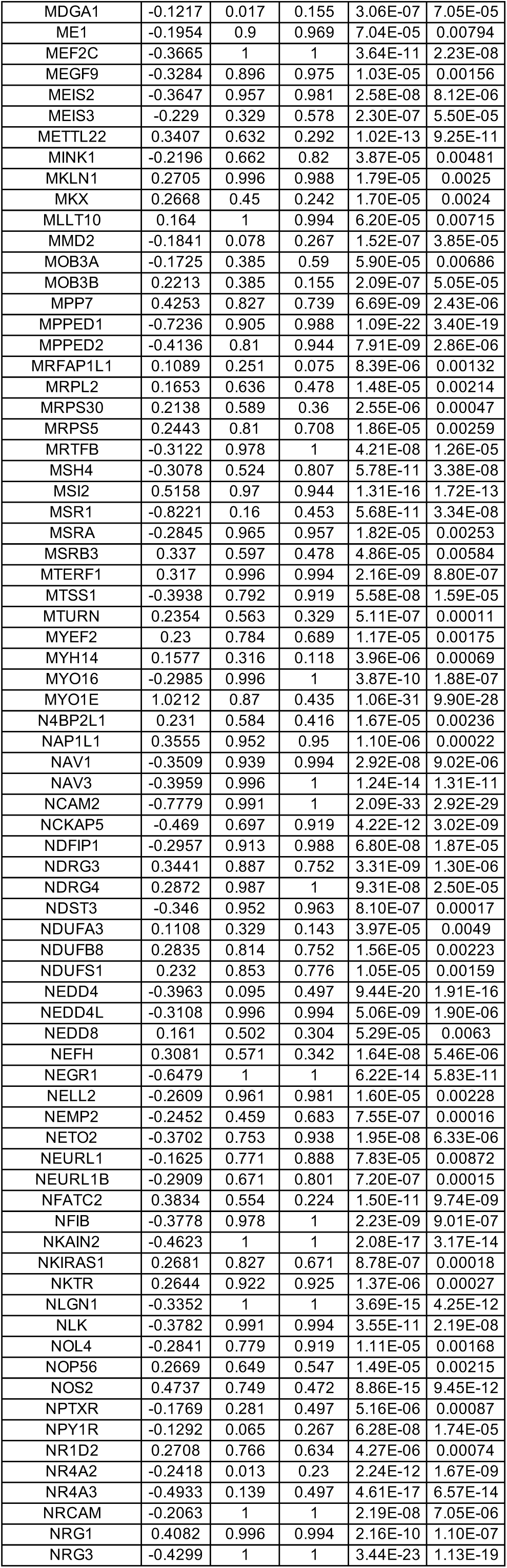
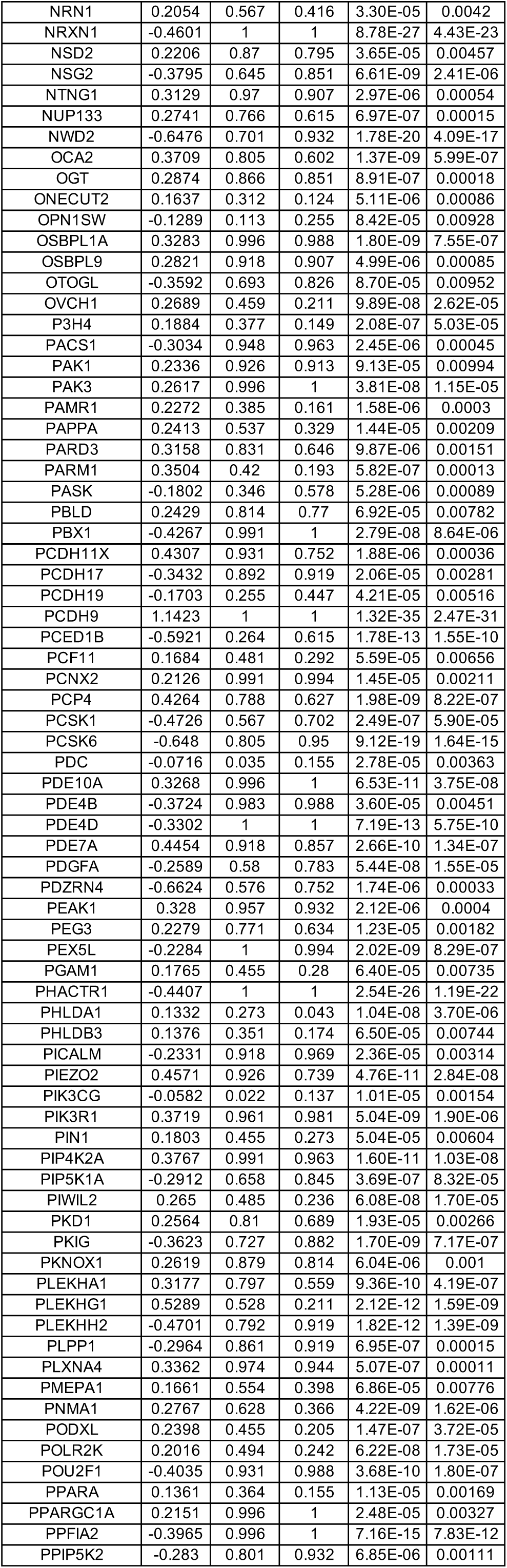
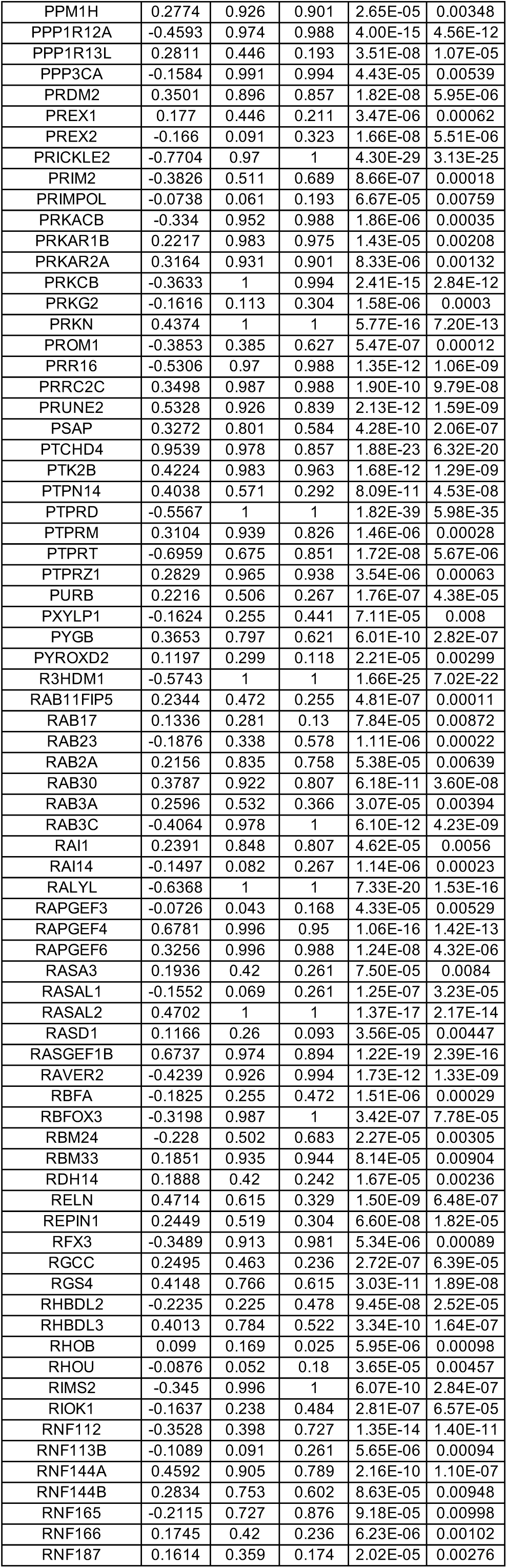
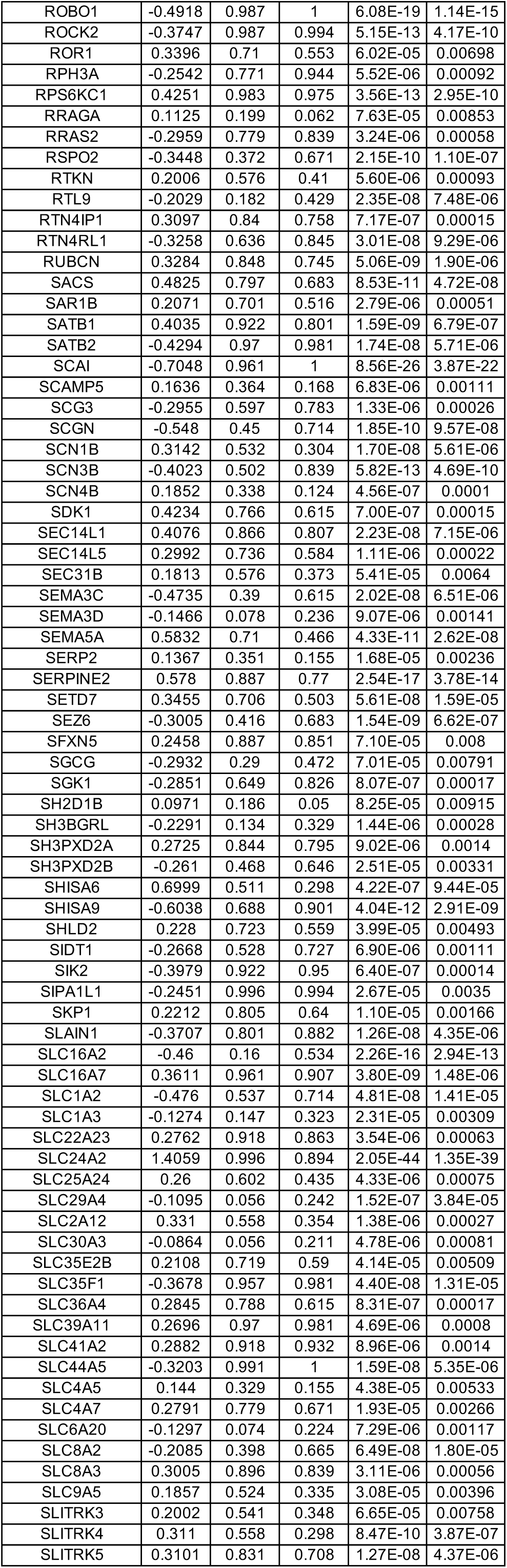
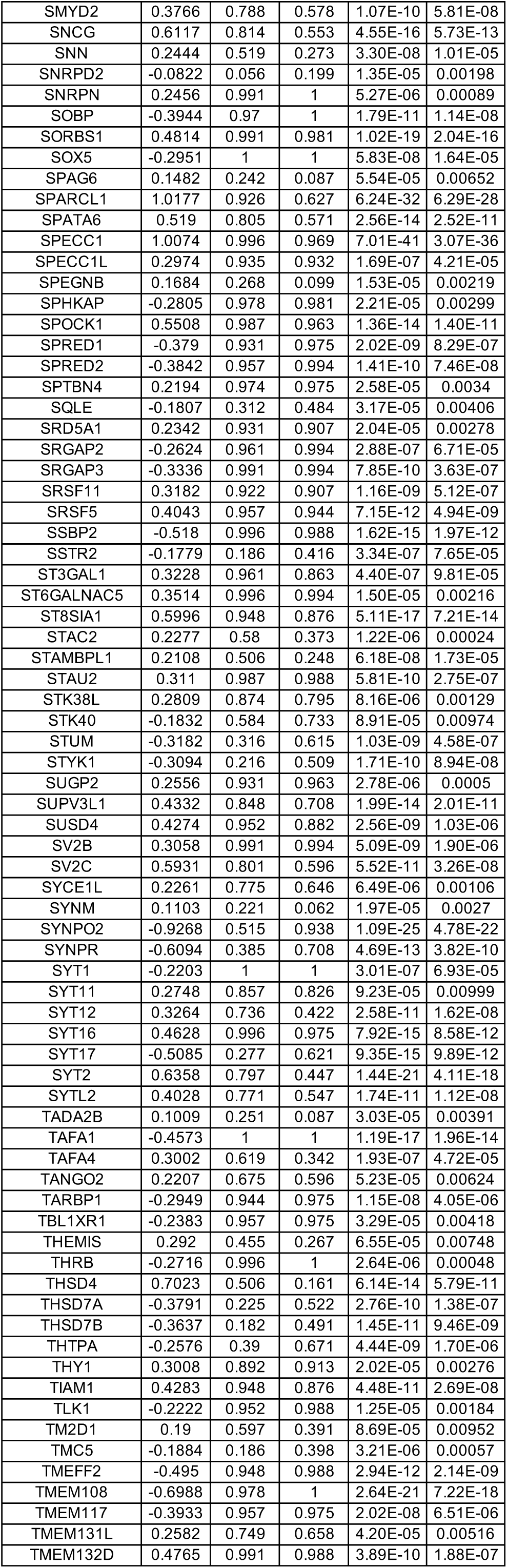
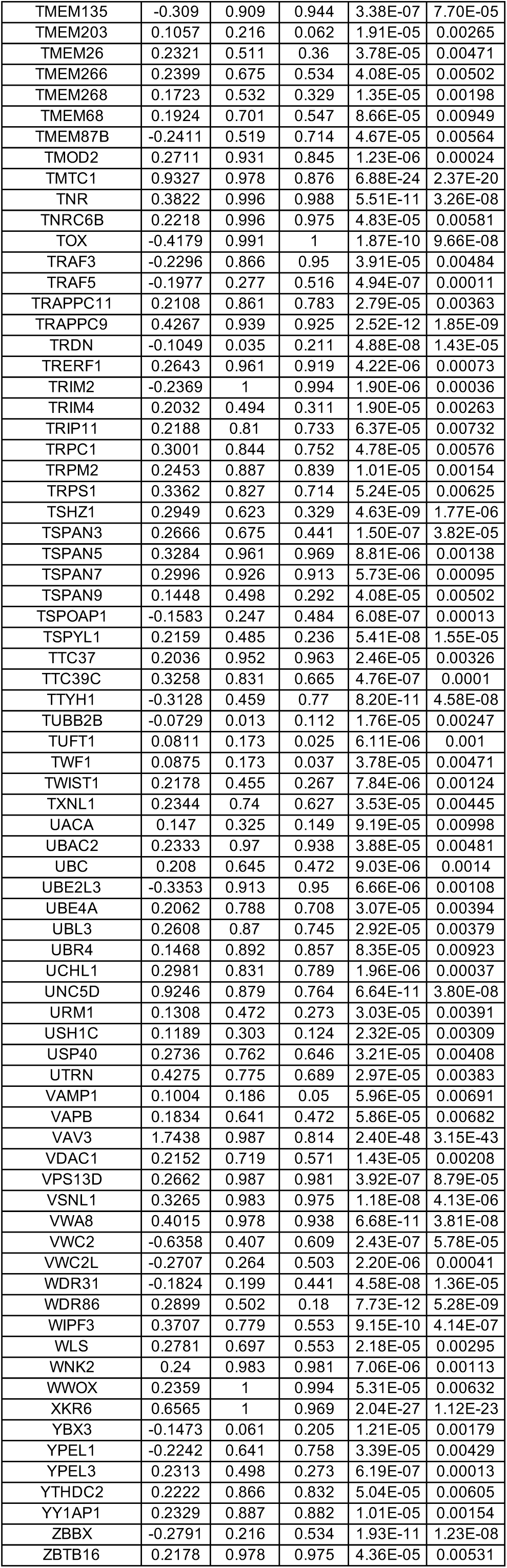
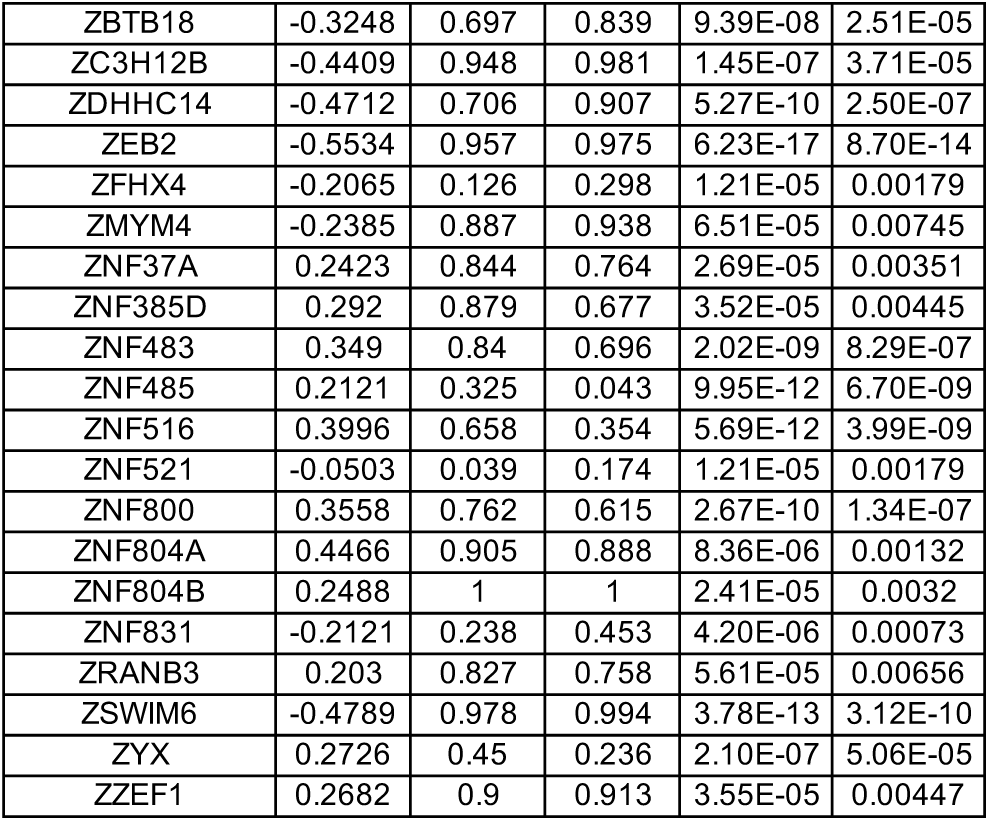
DEGs between wild-type (WT) and MECP2-null (KO) upper layer excitatory neurons (Ex_8) of PFC (adjusted p-value < 0.01)

**Table S22.**
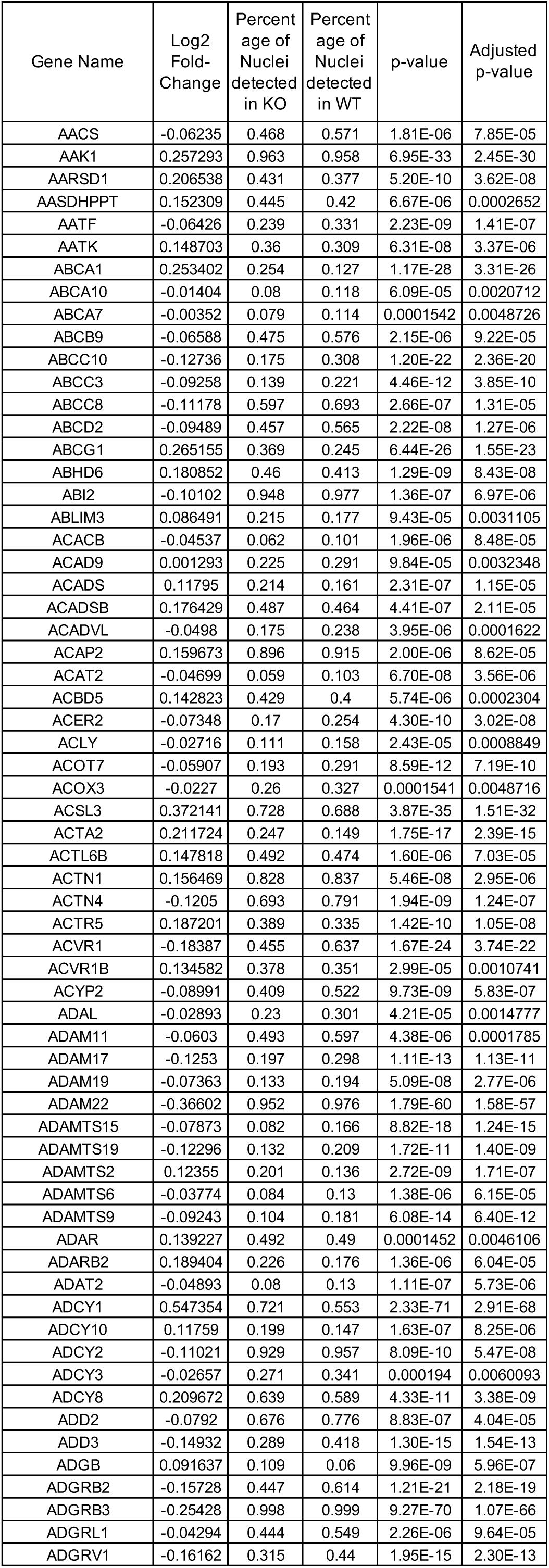
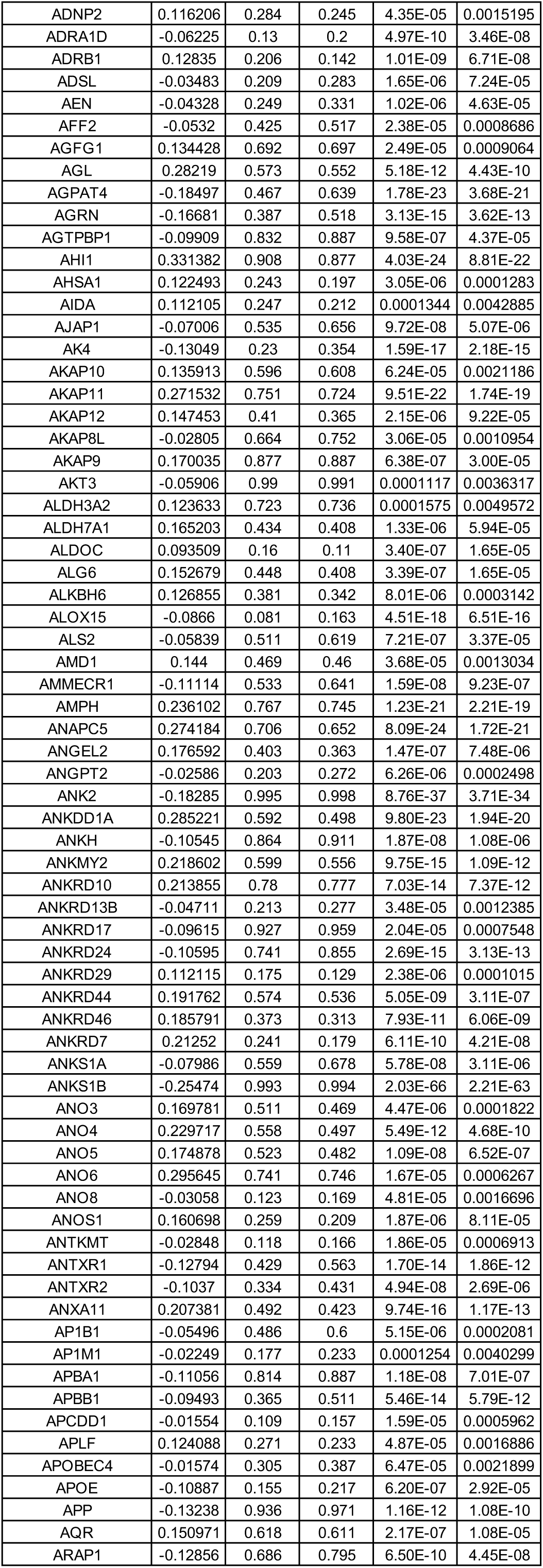
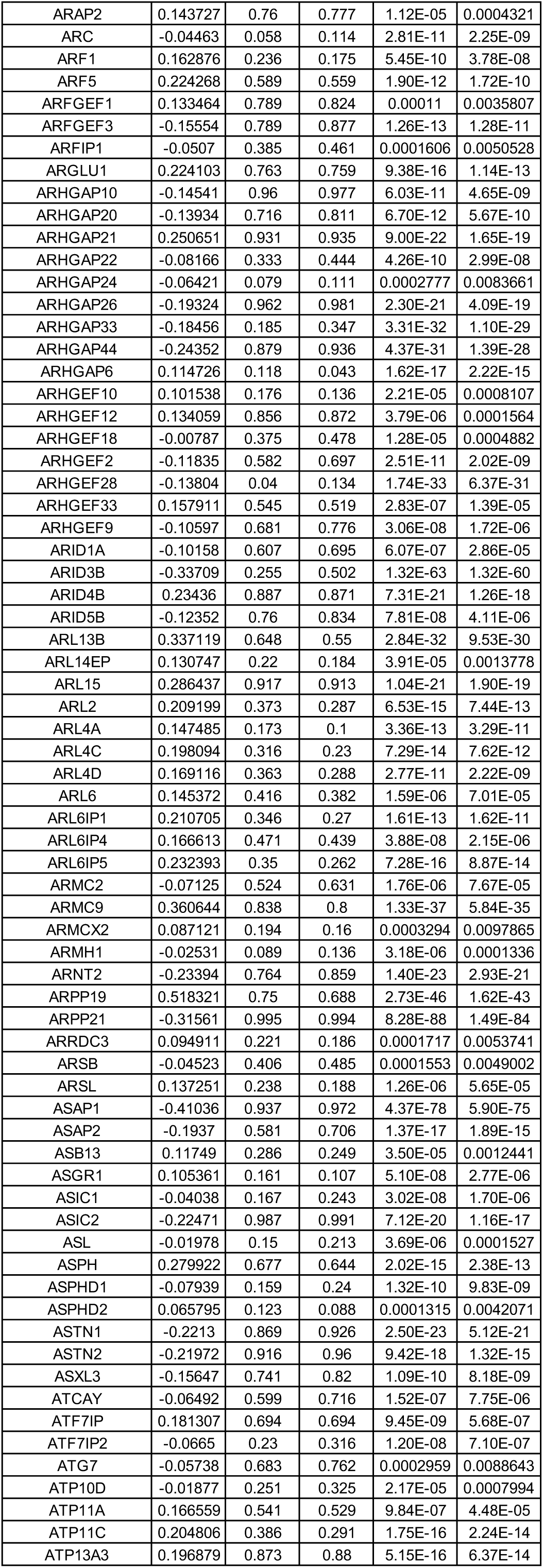
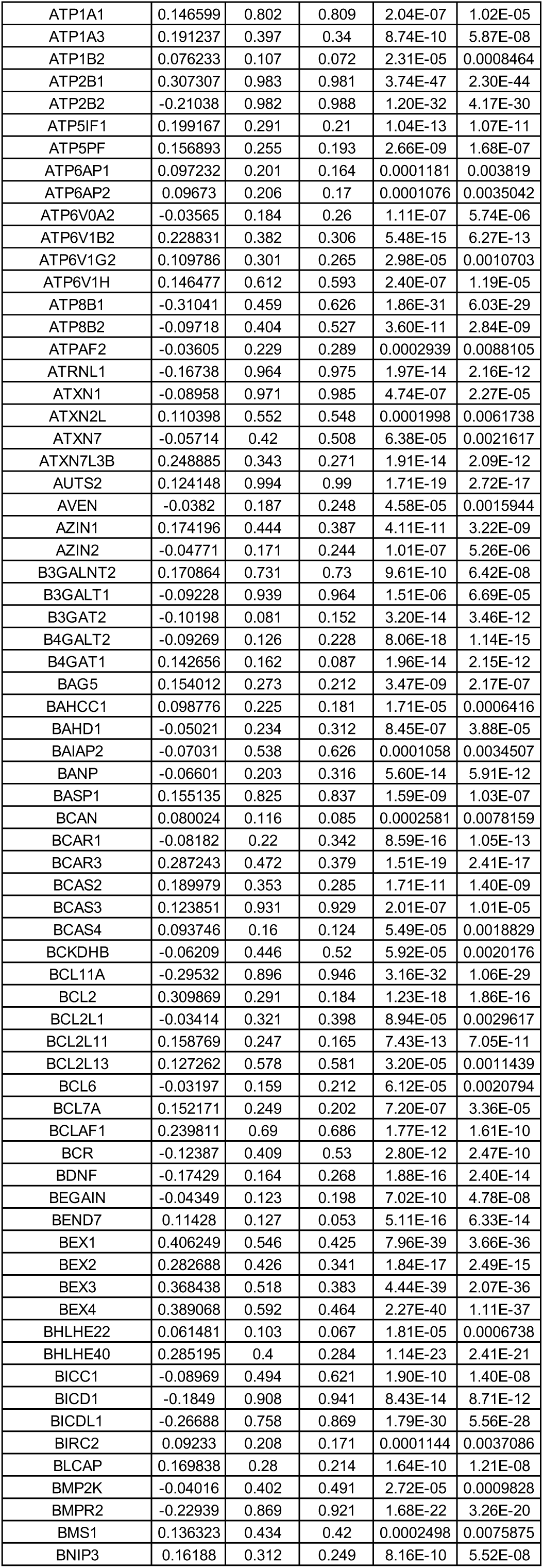
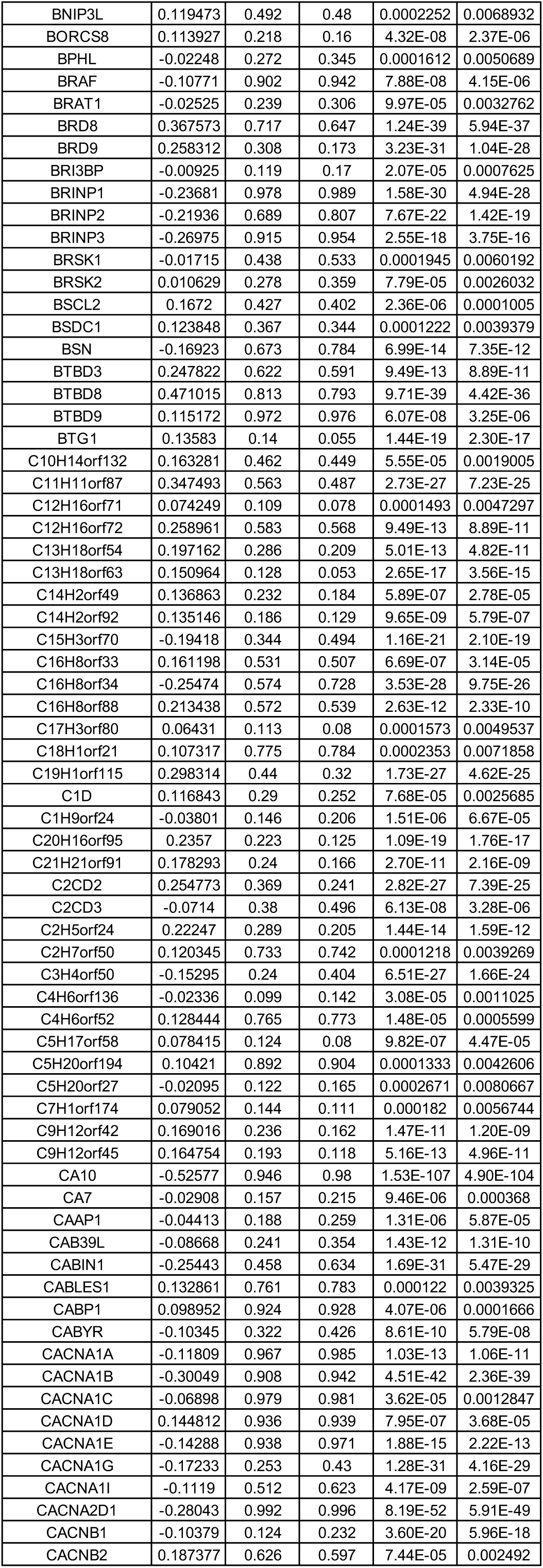
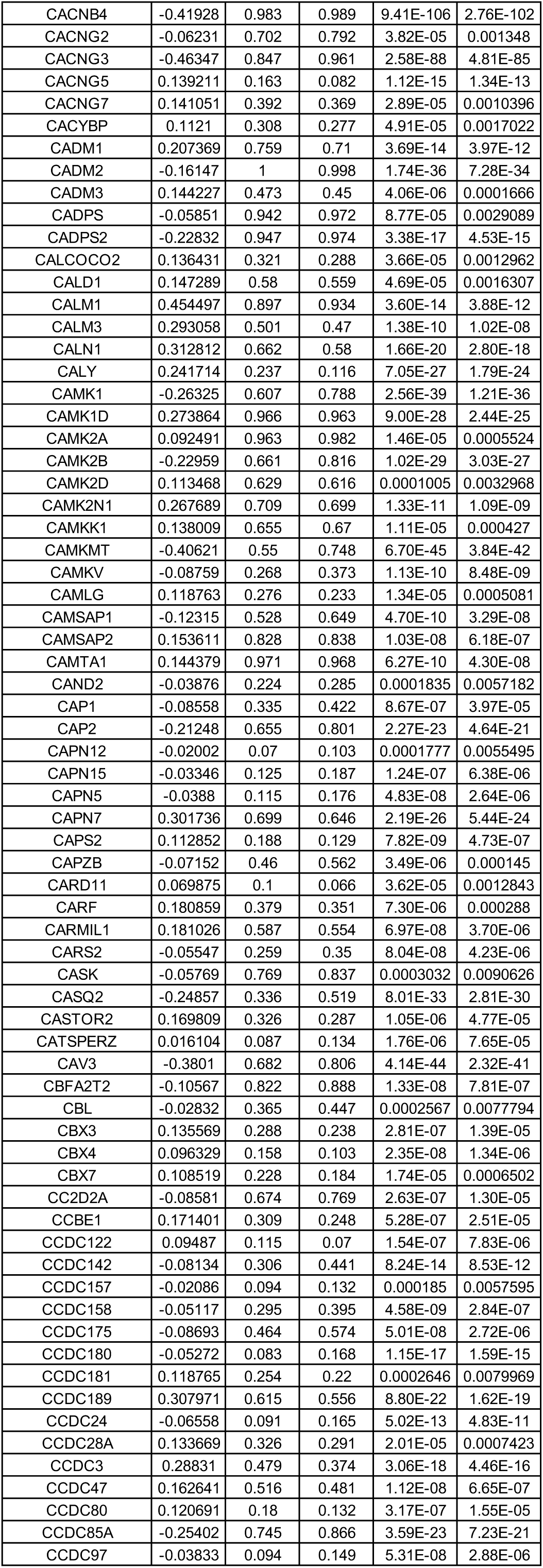
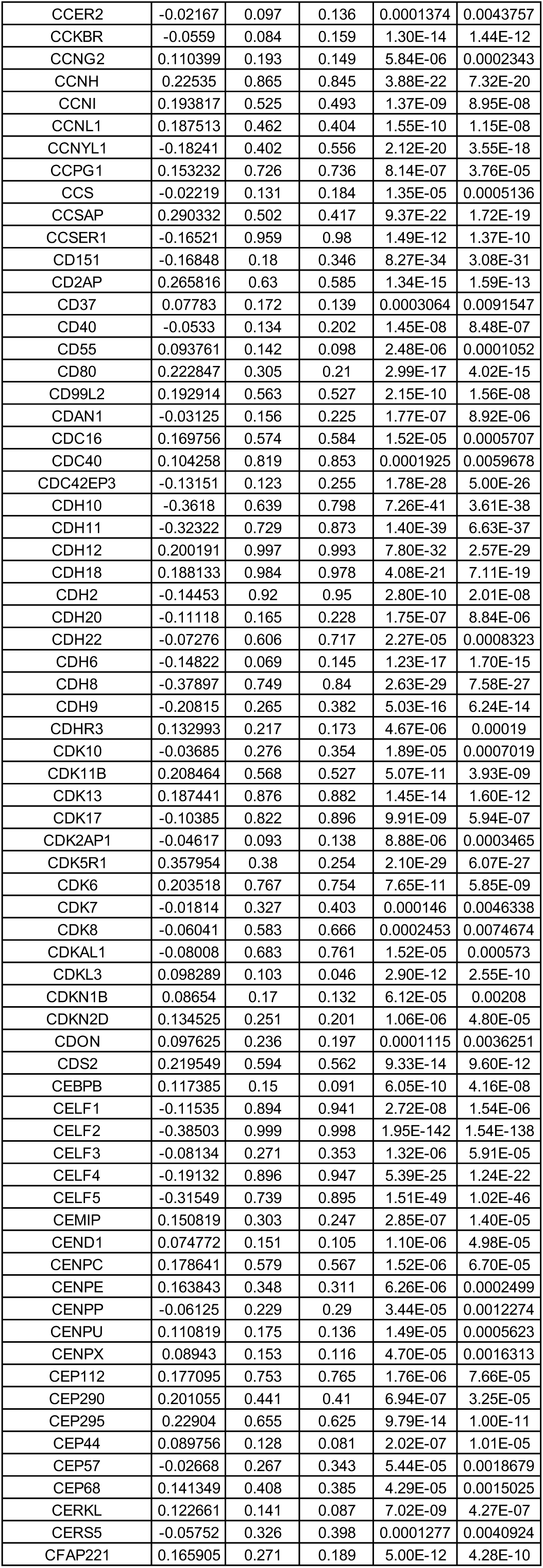
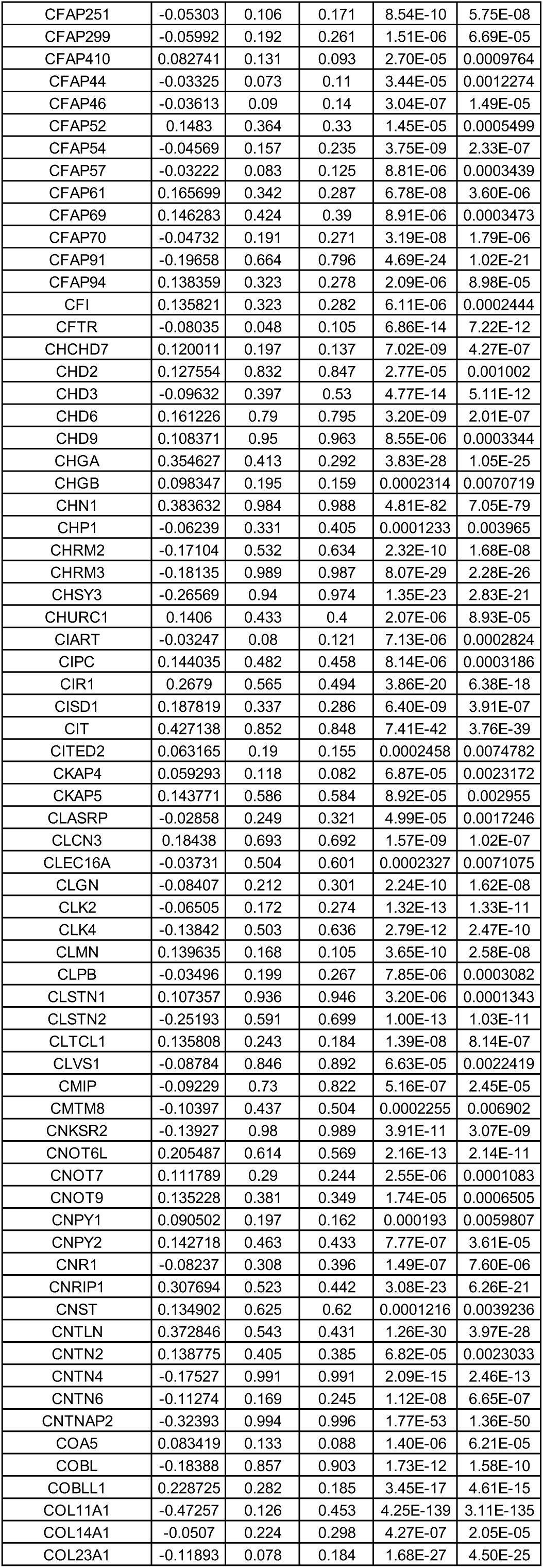
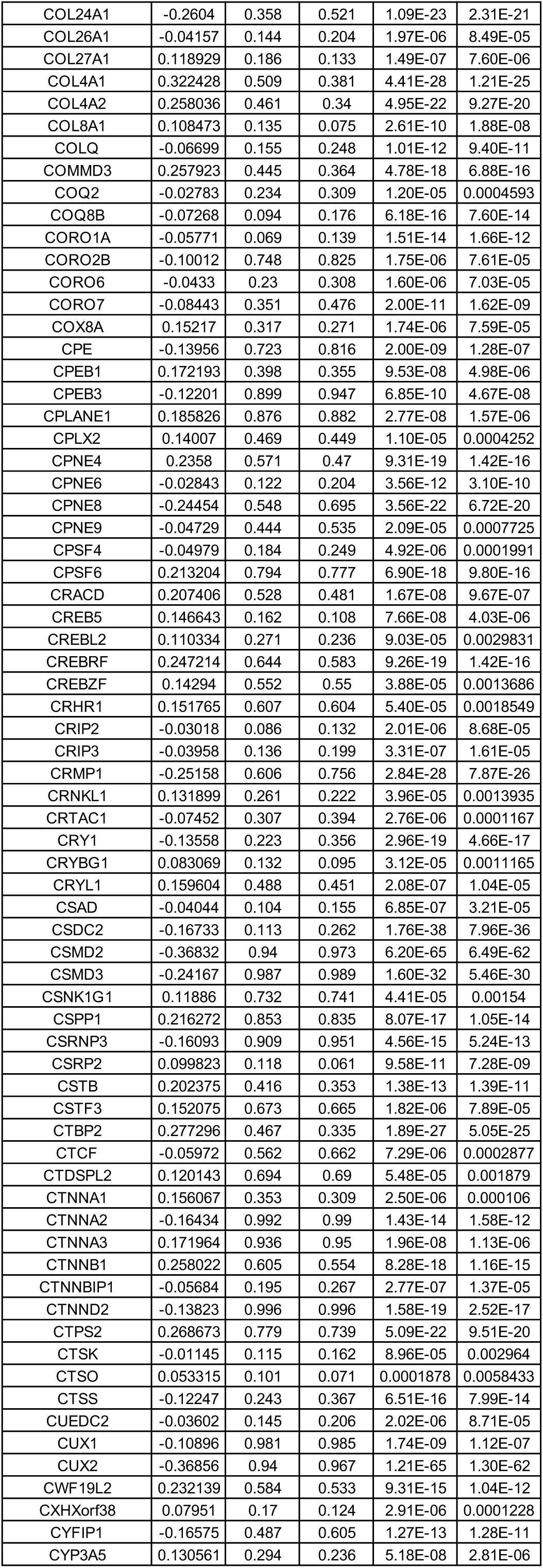
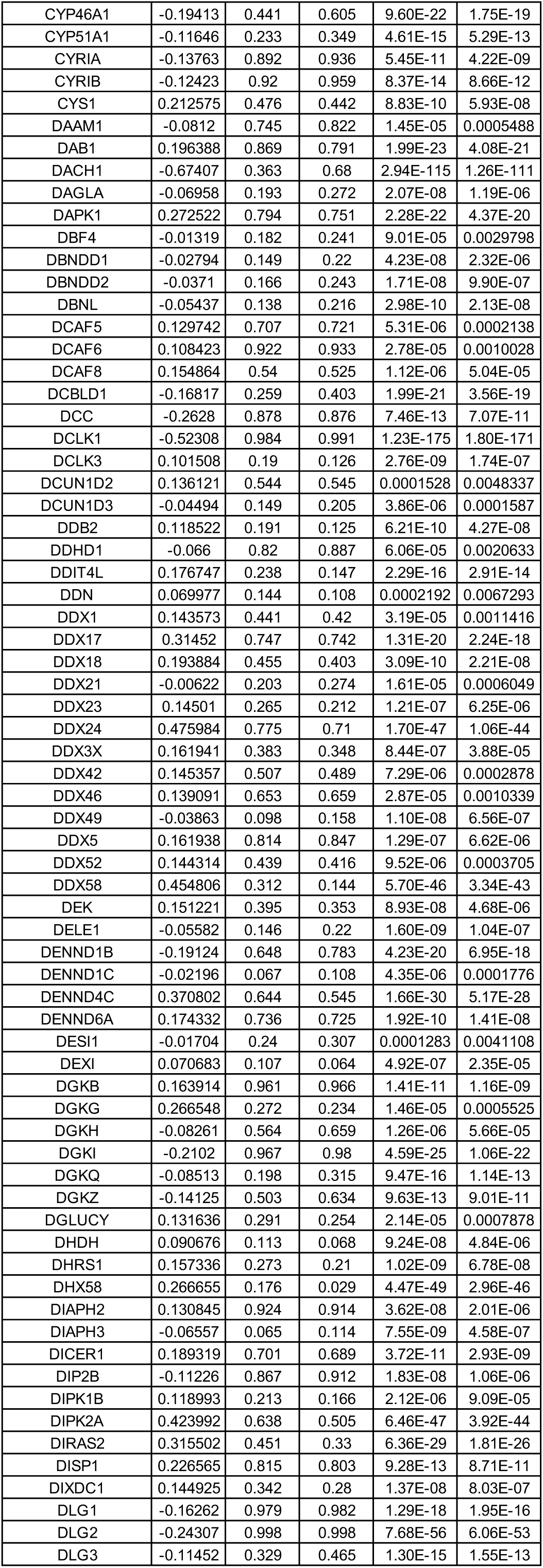
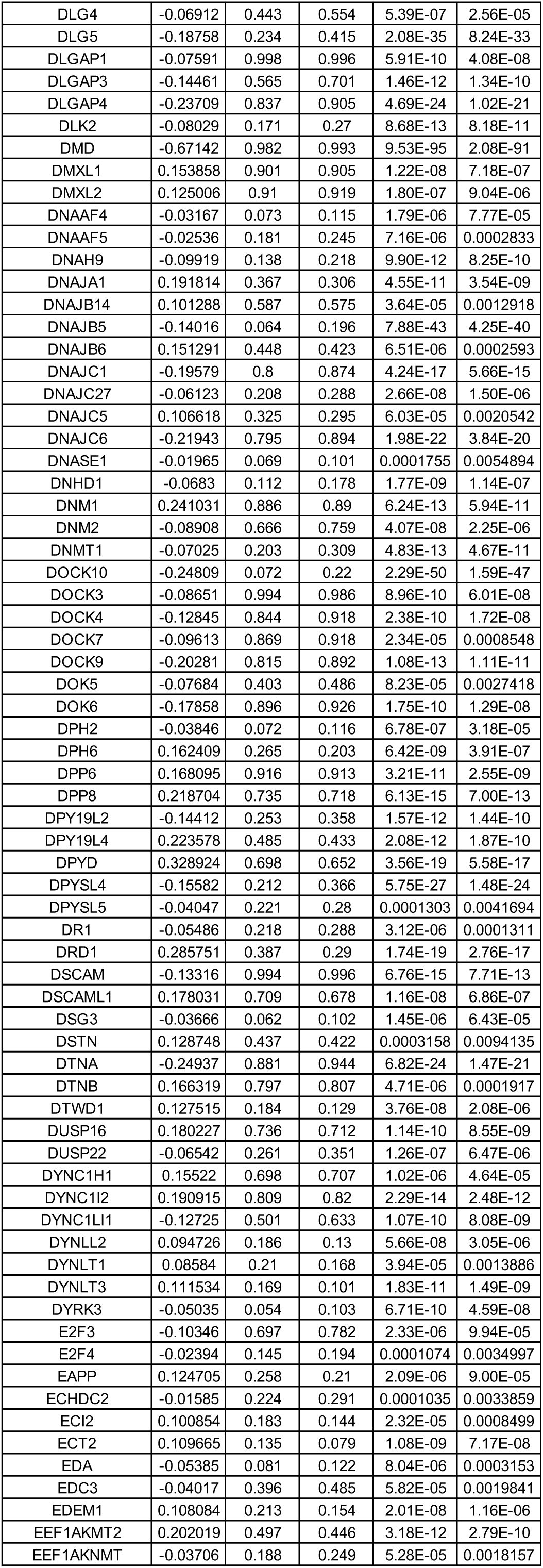
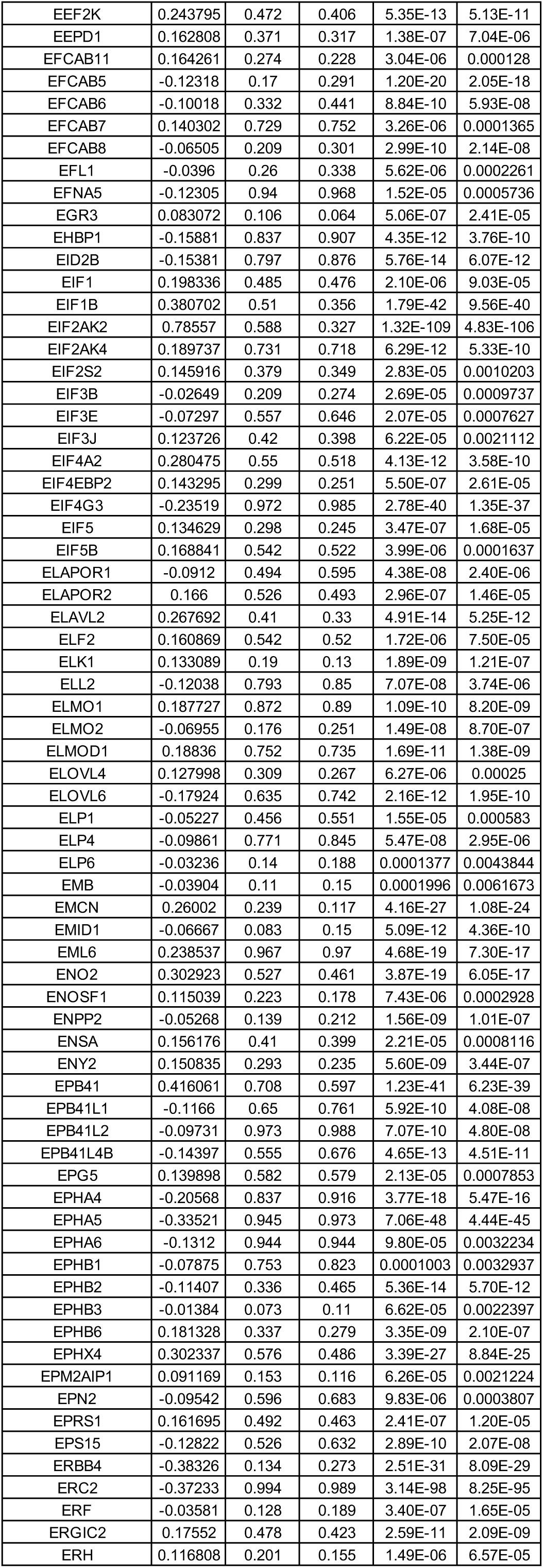
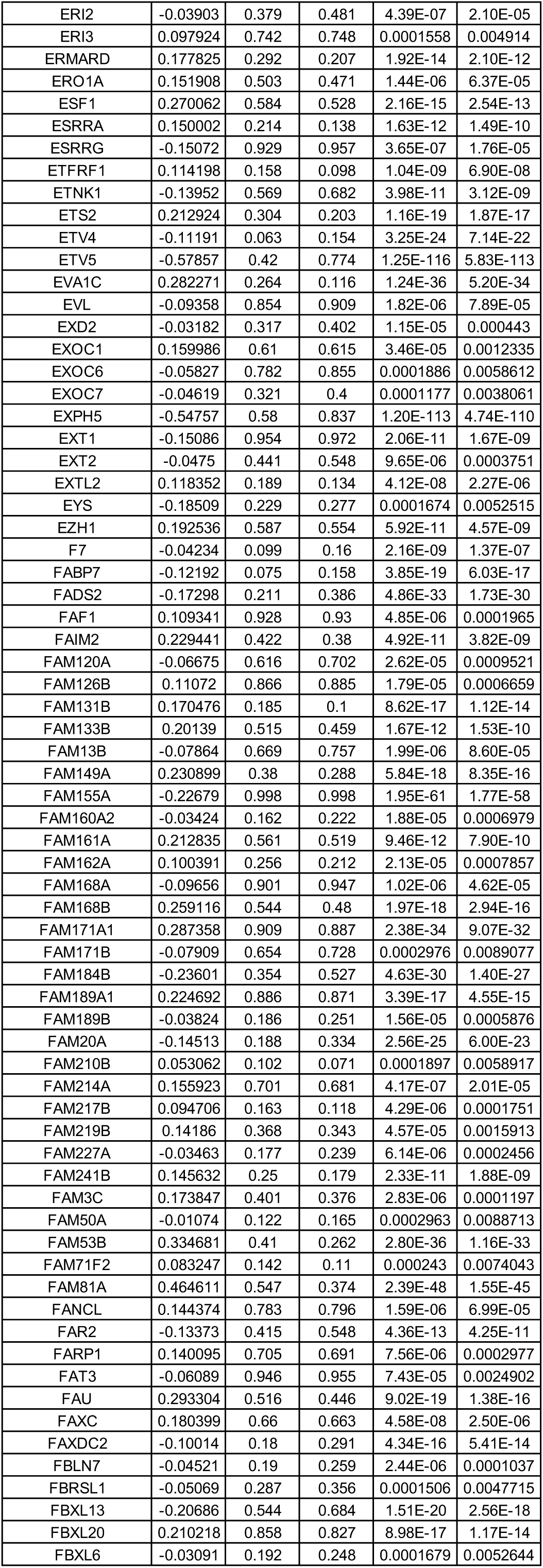
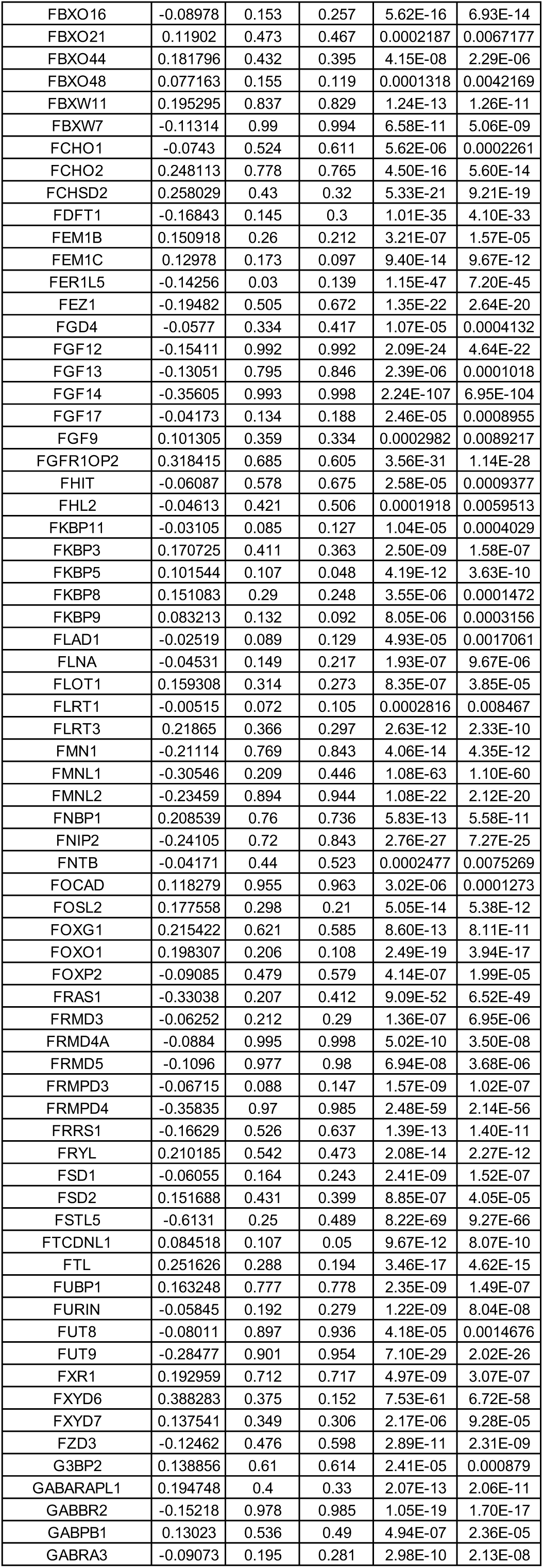
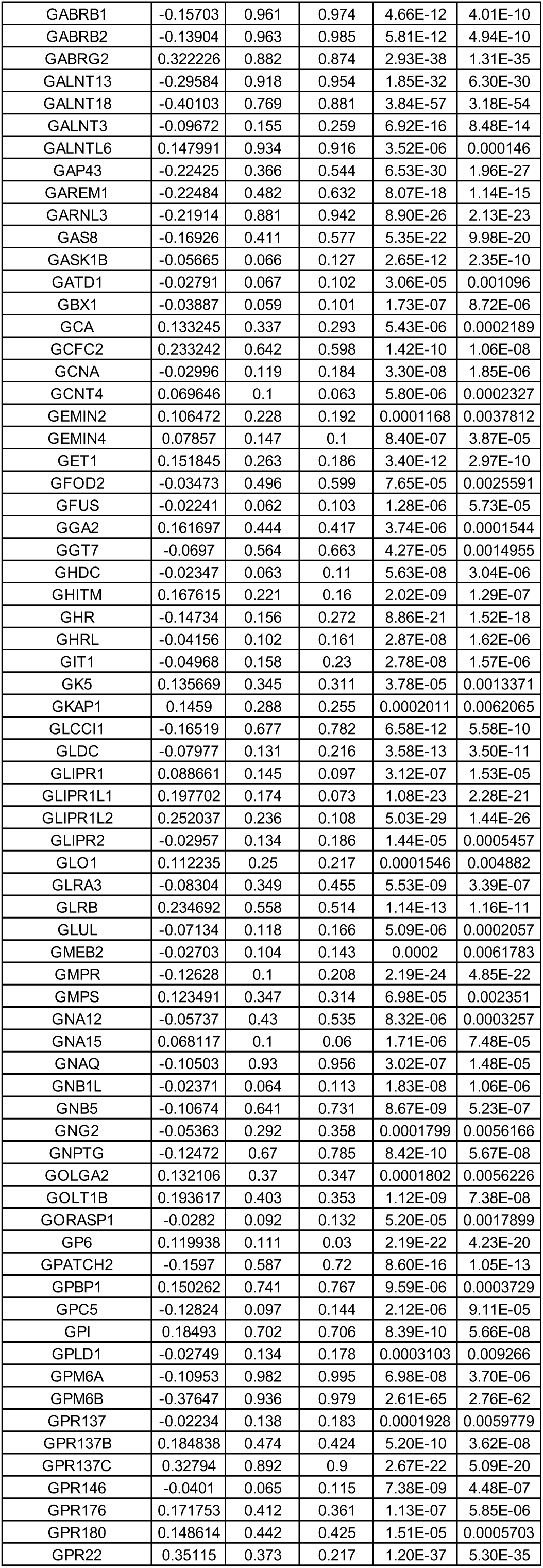
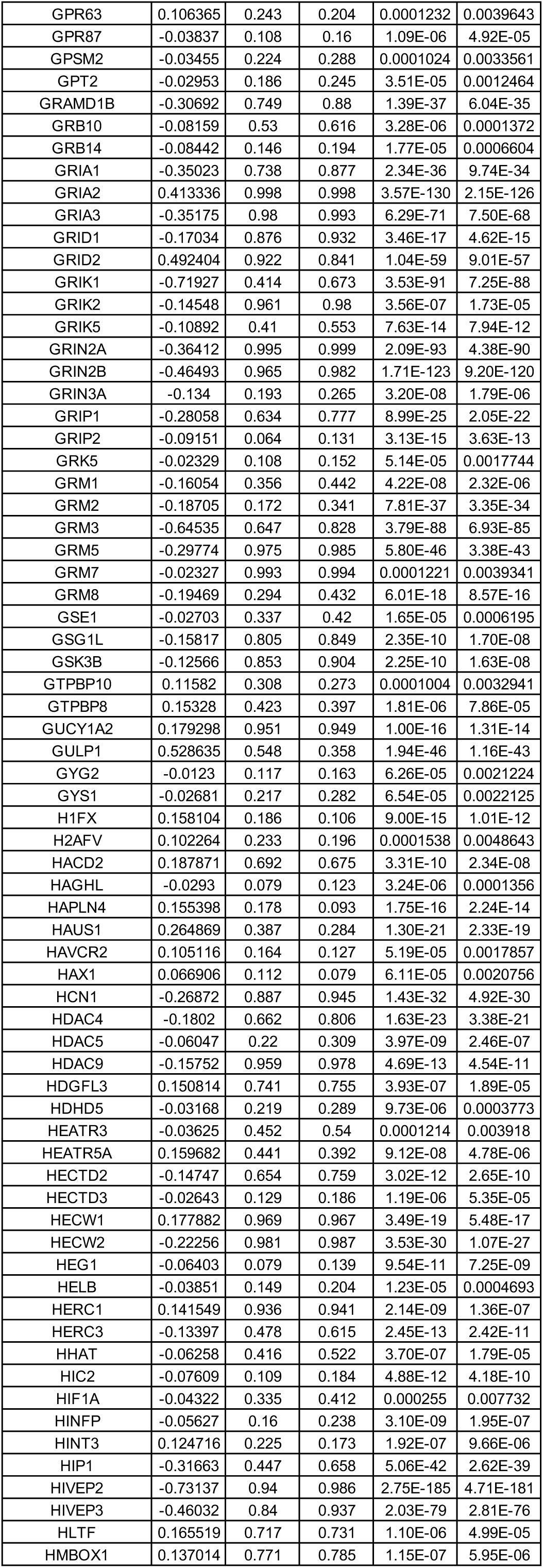
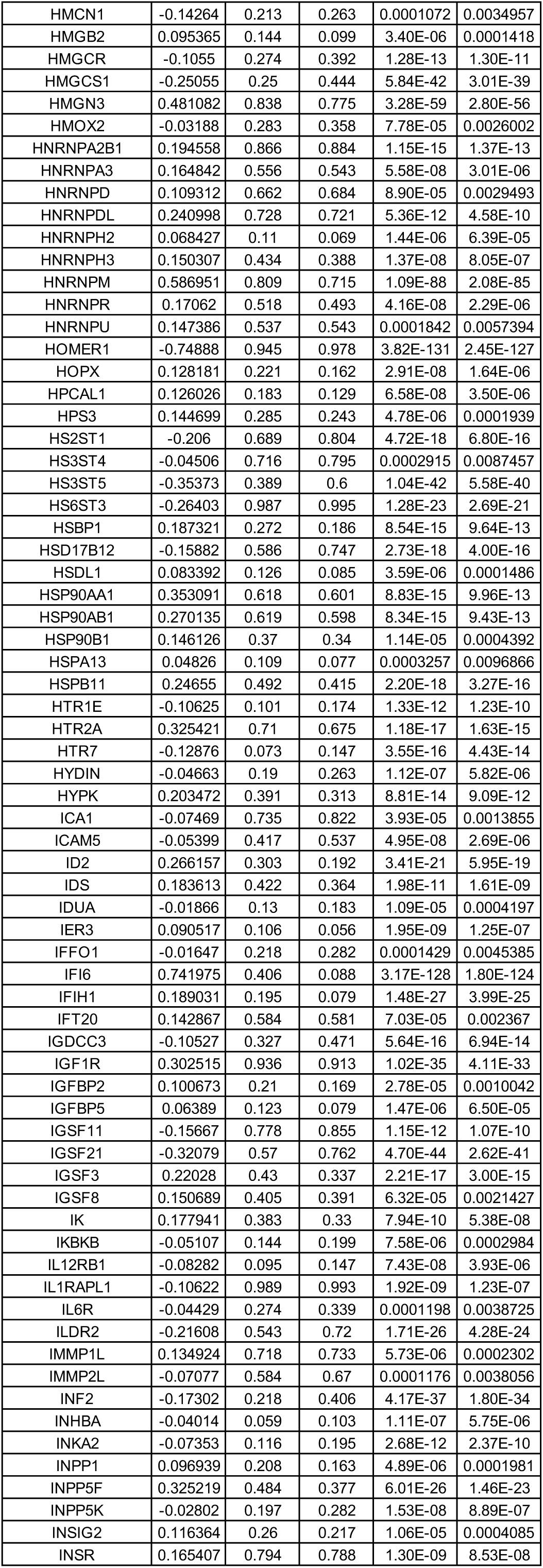
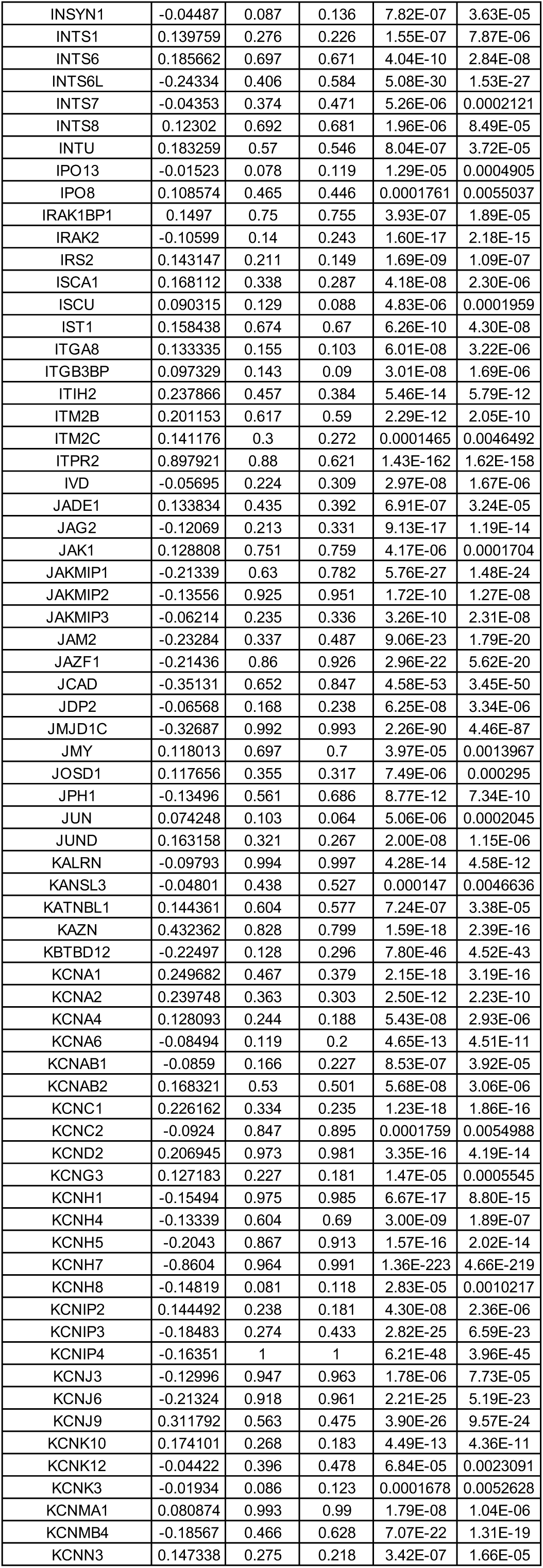
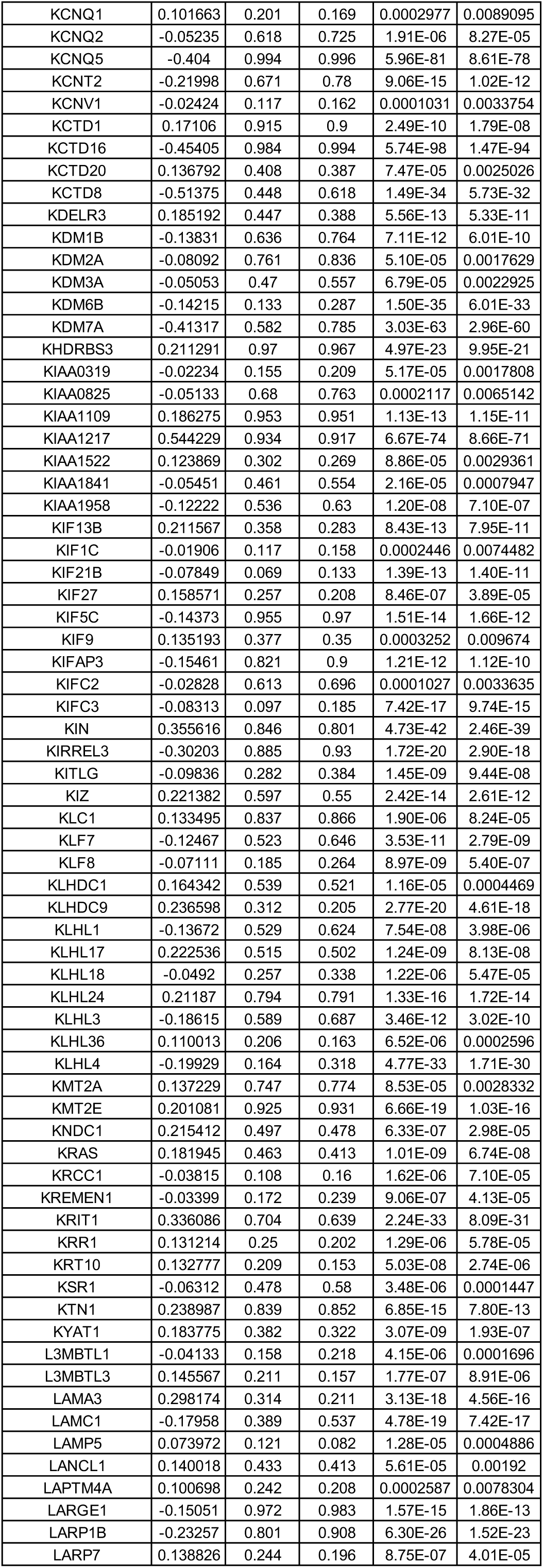
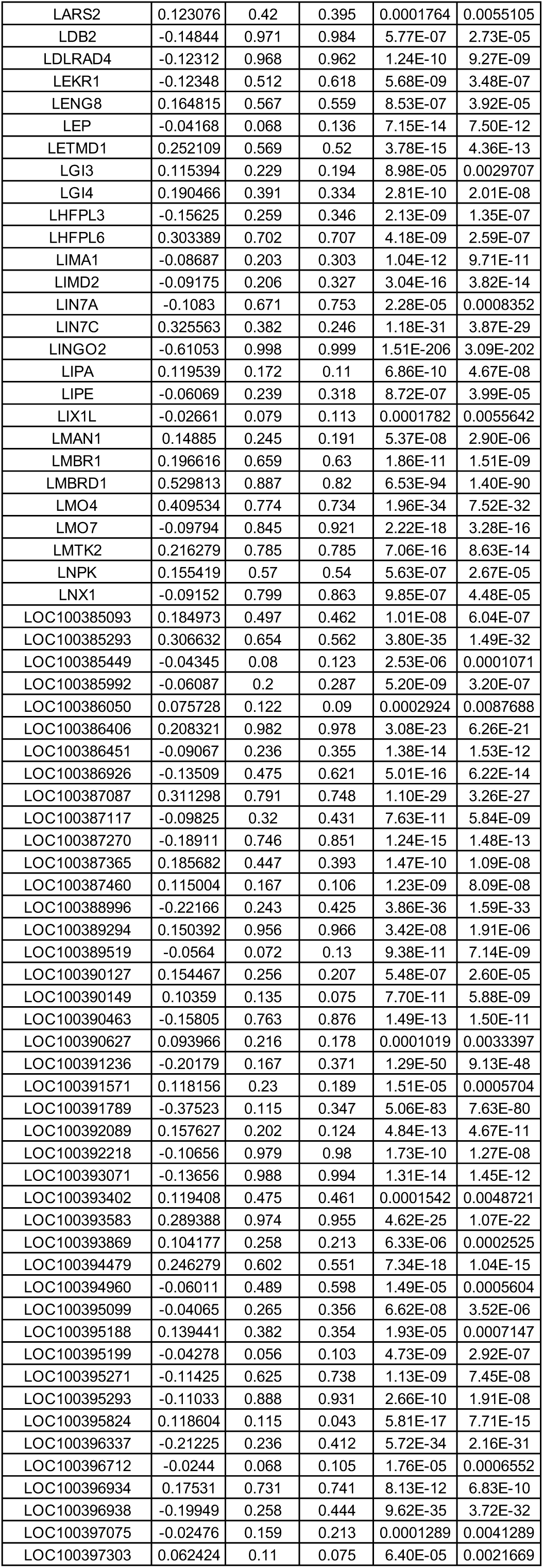
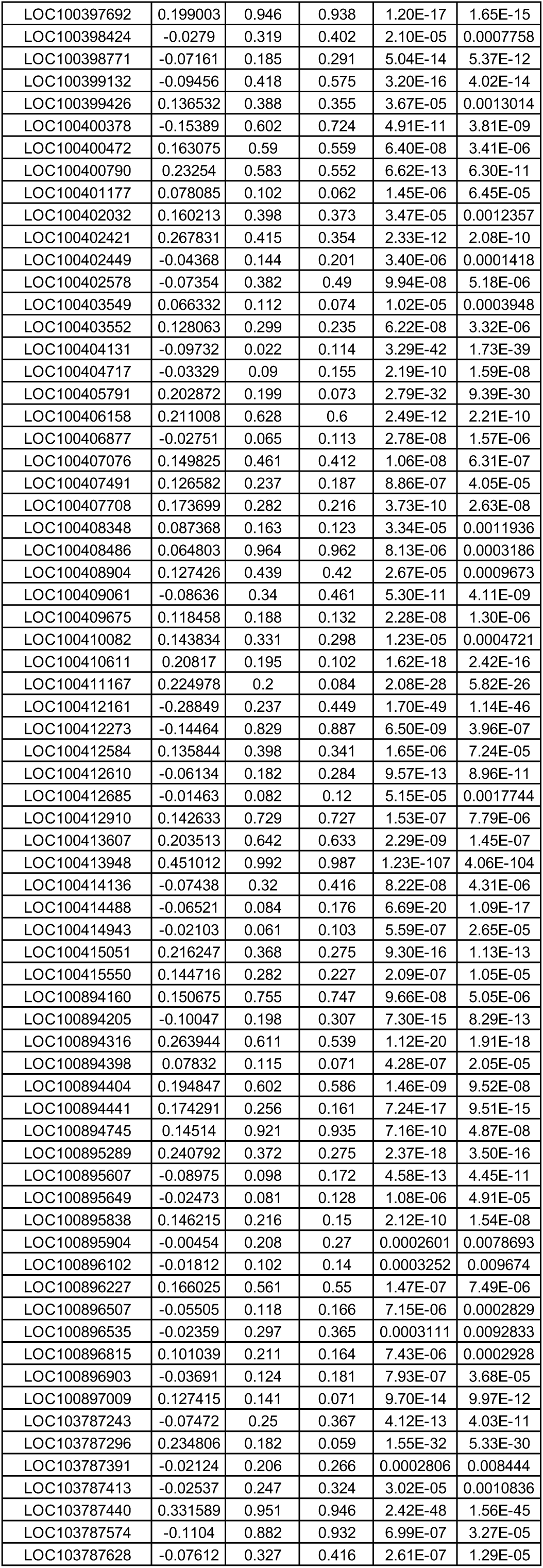
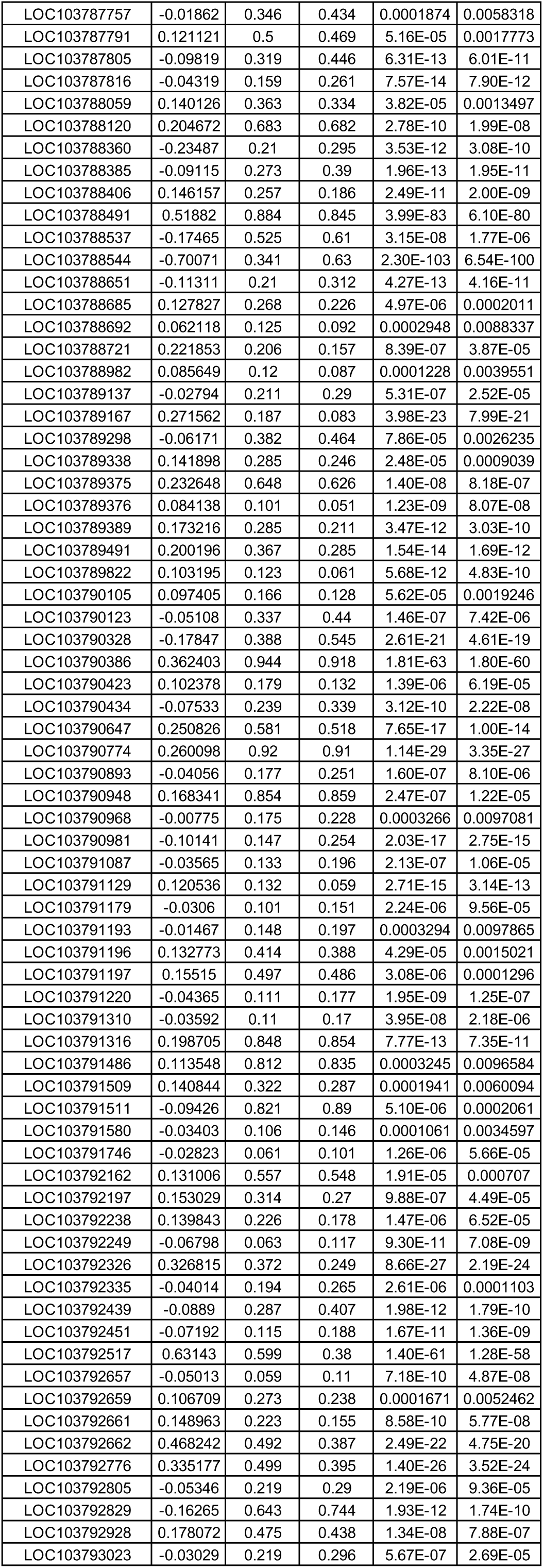
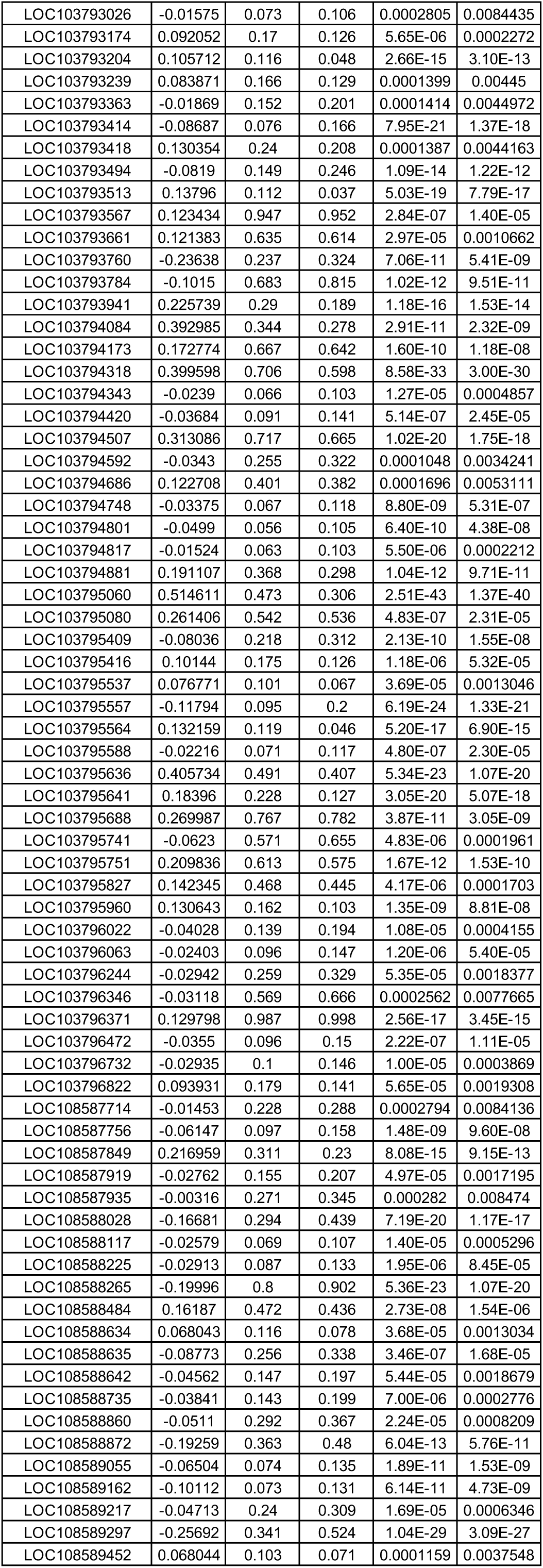
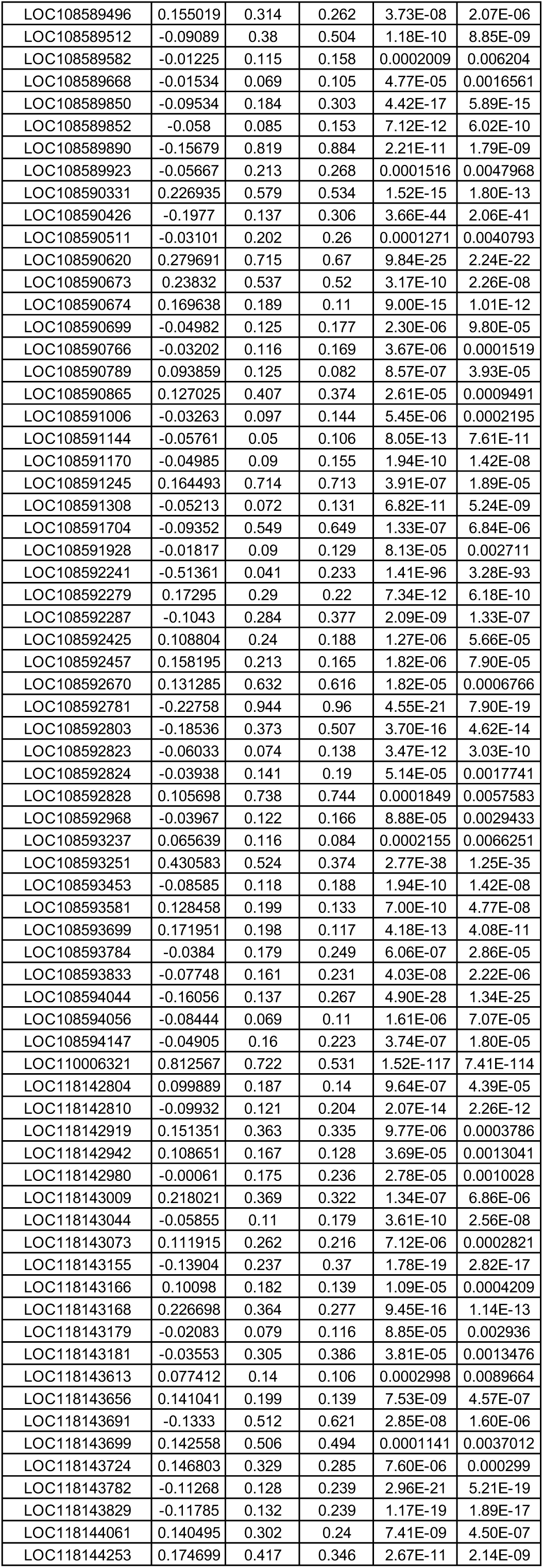
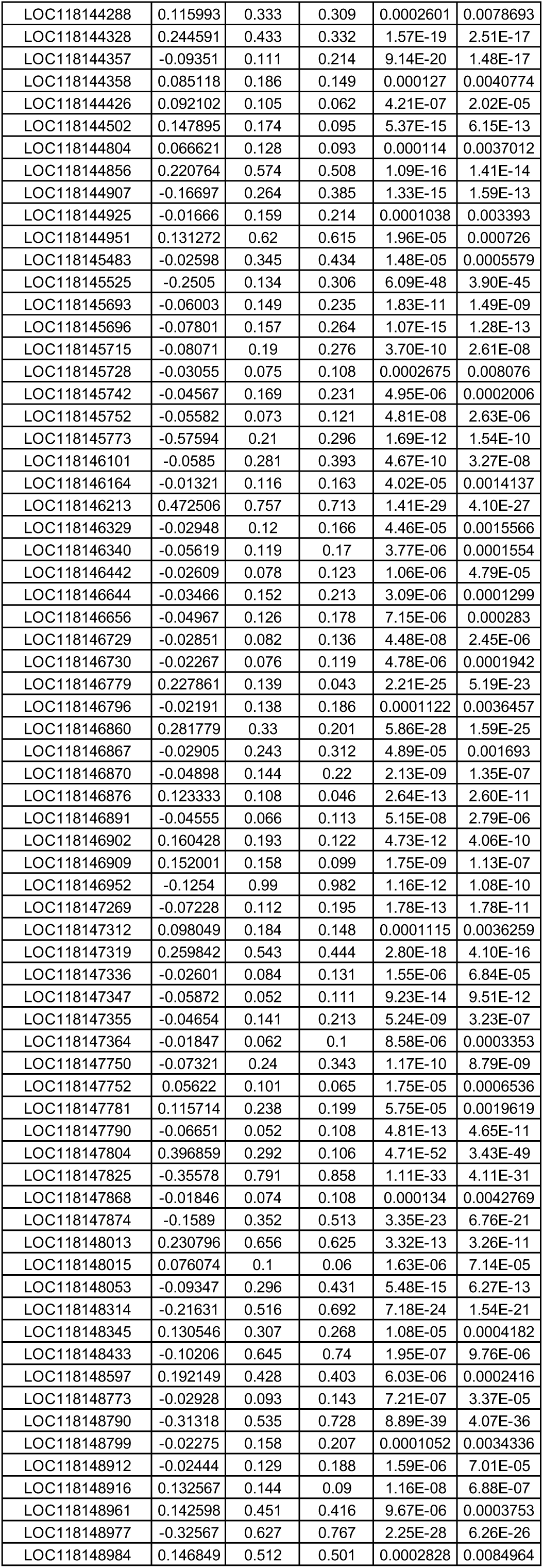
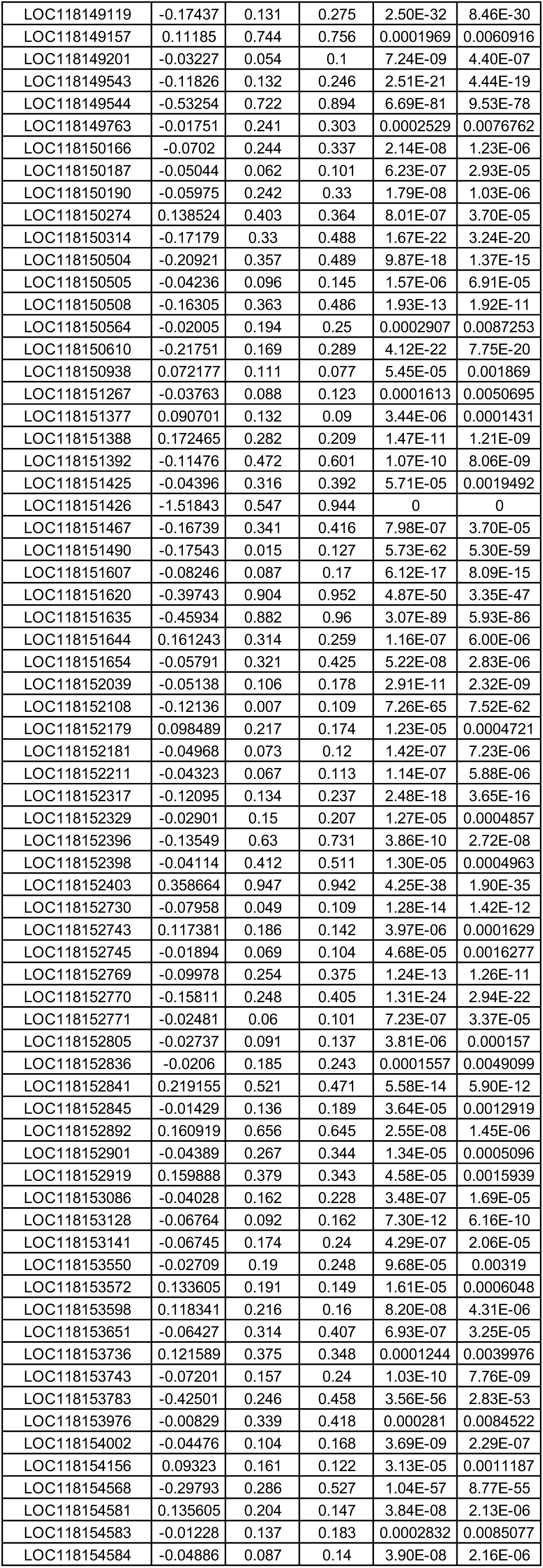
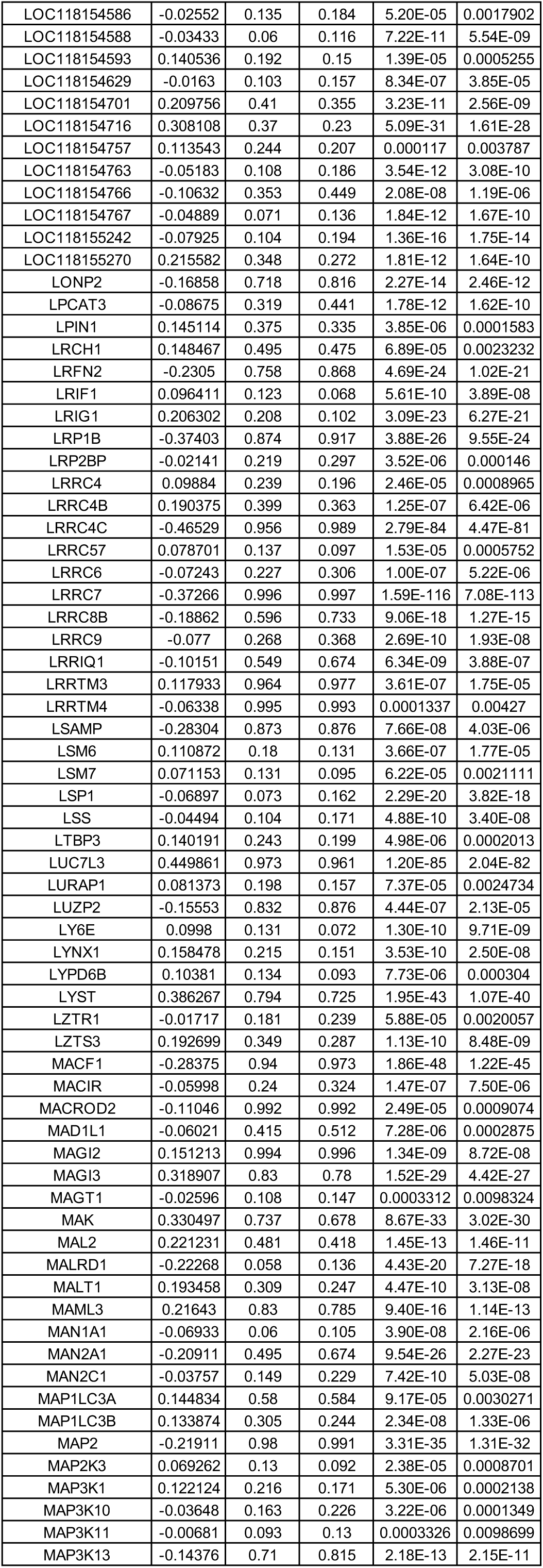
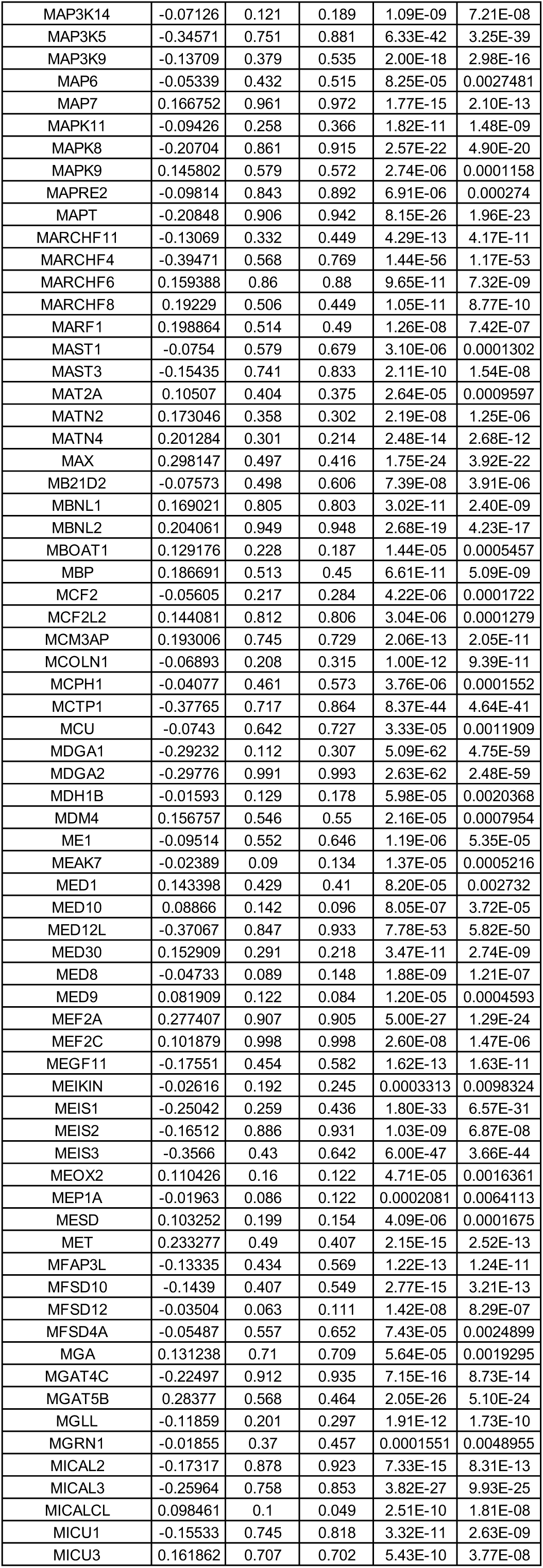
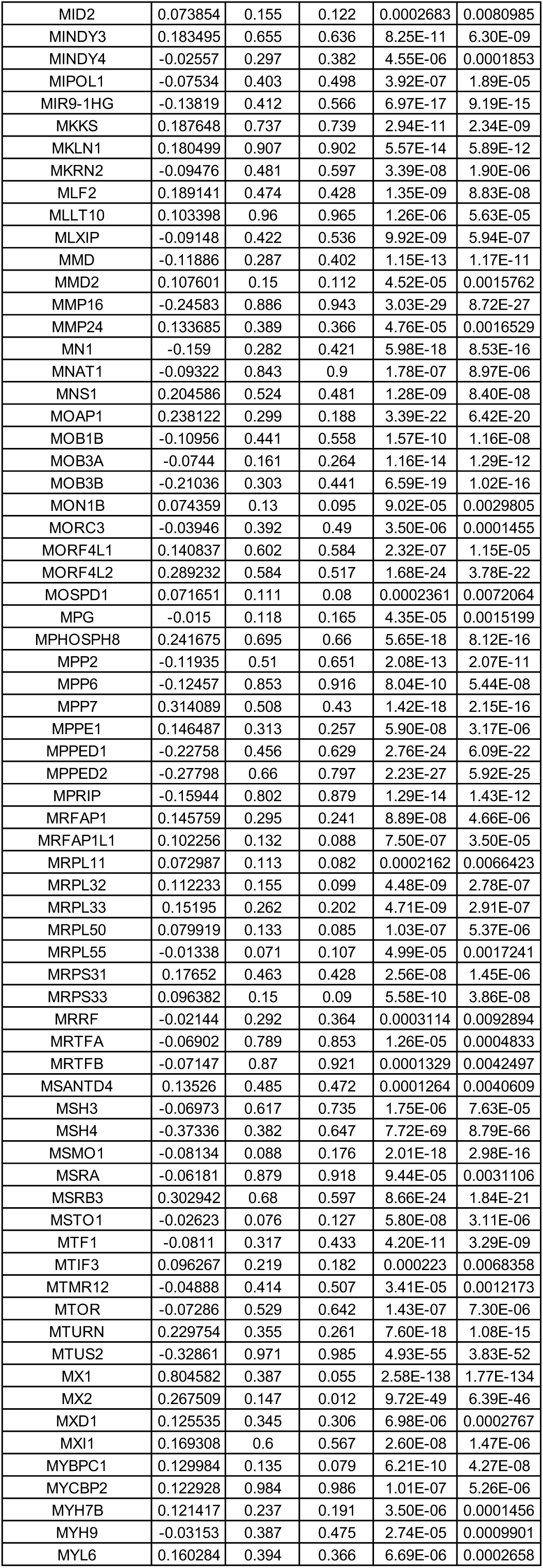
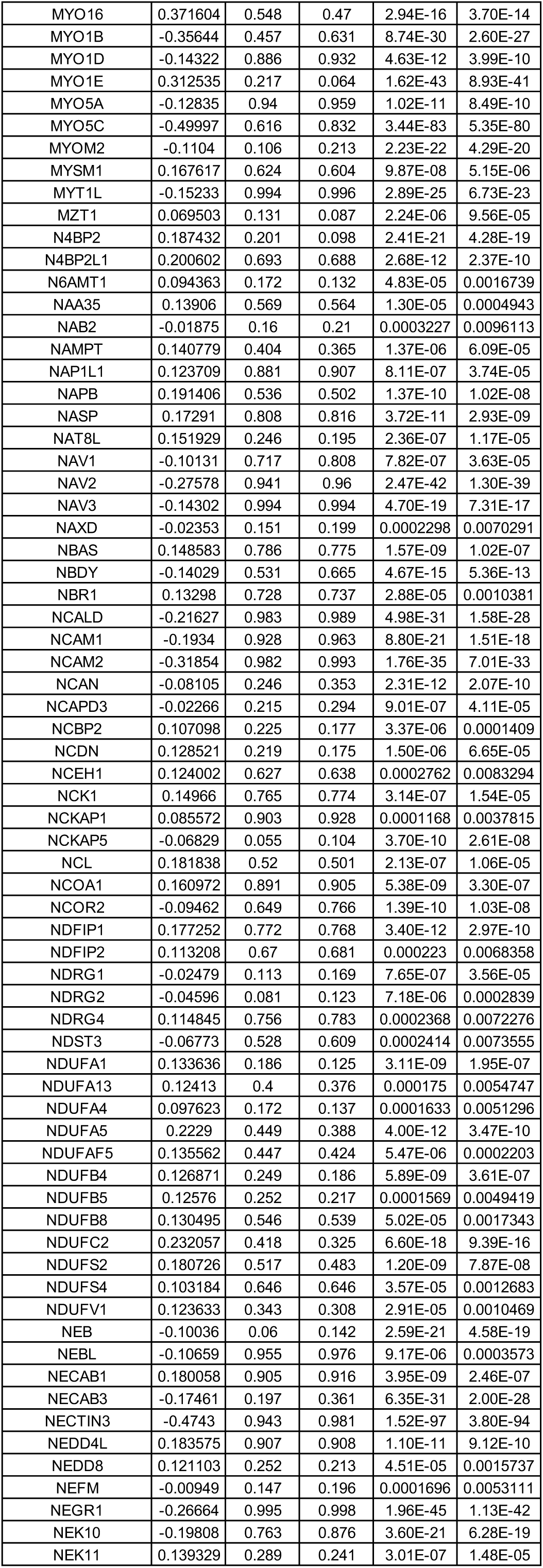
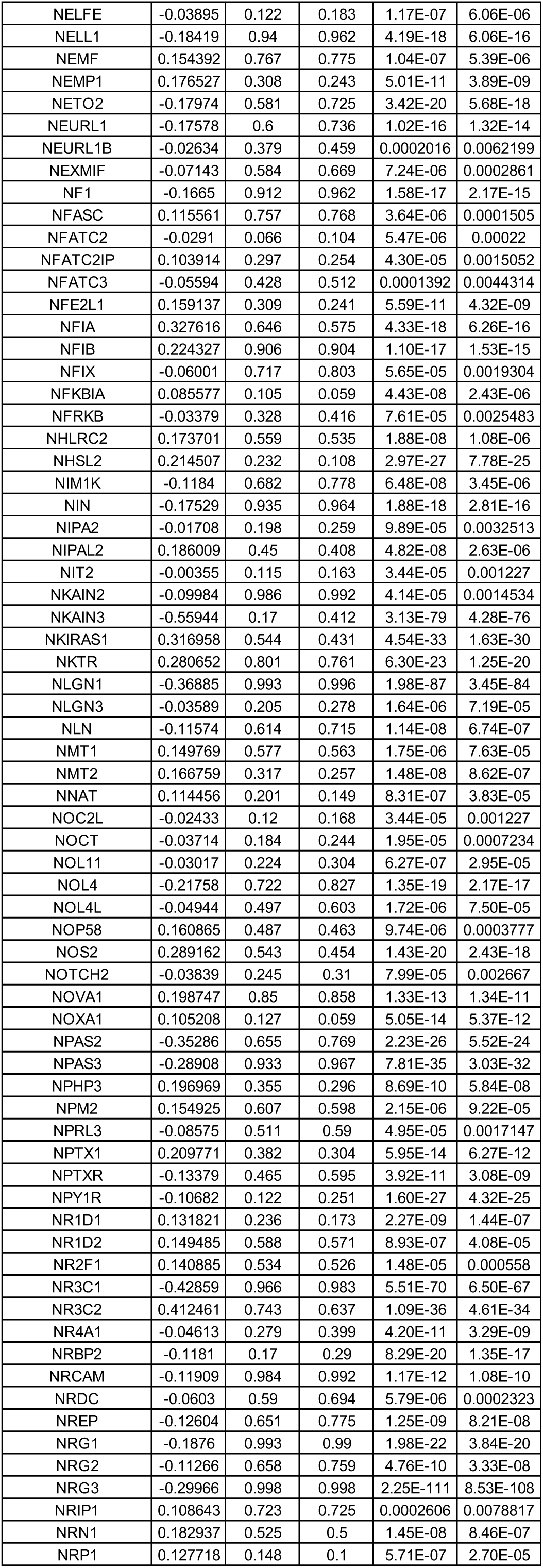
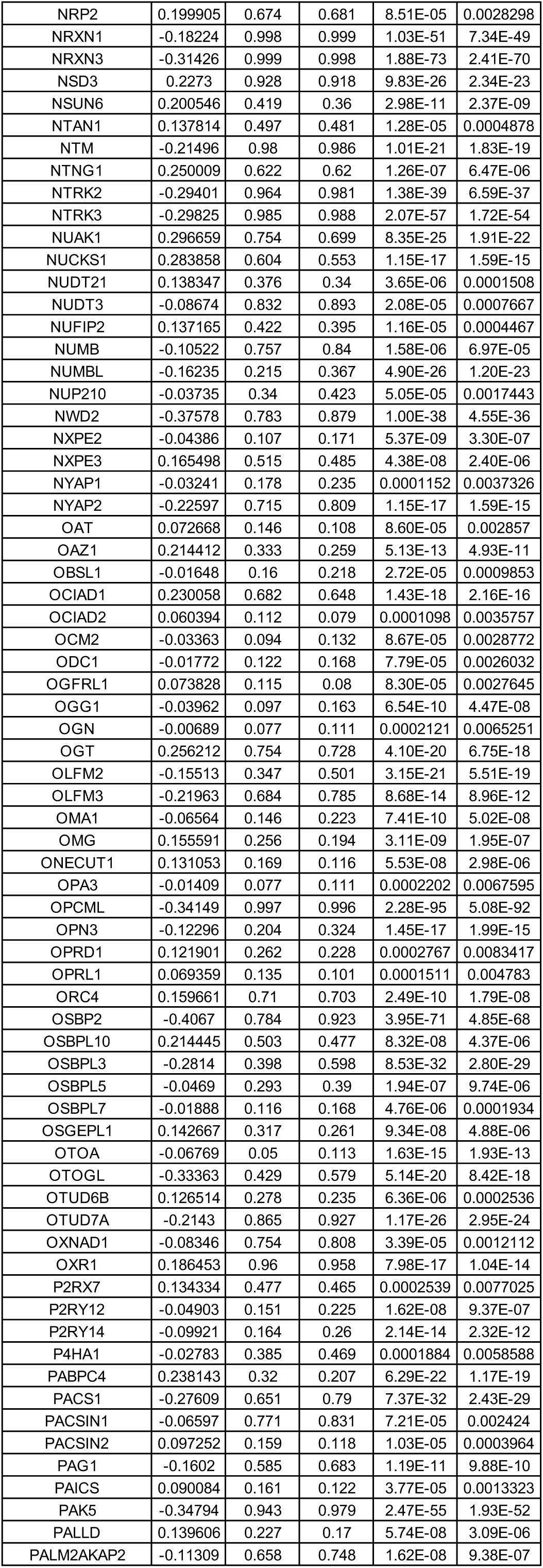
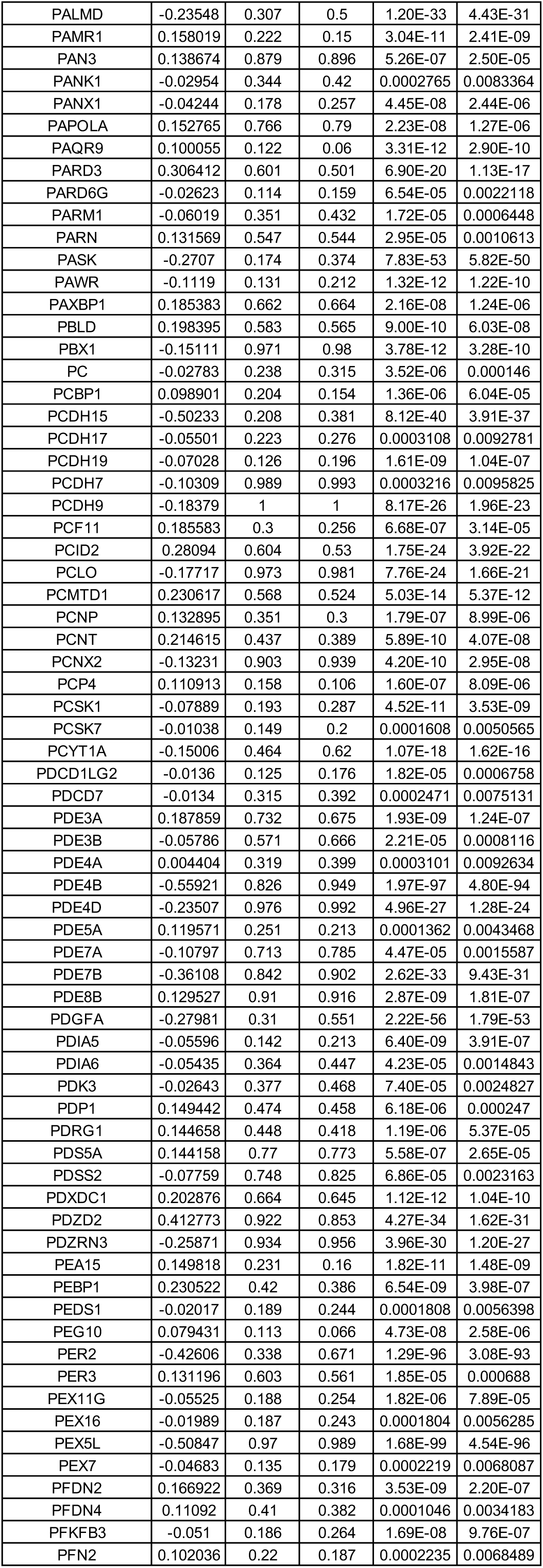
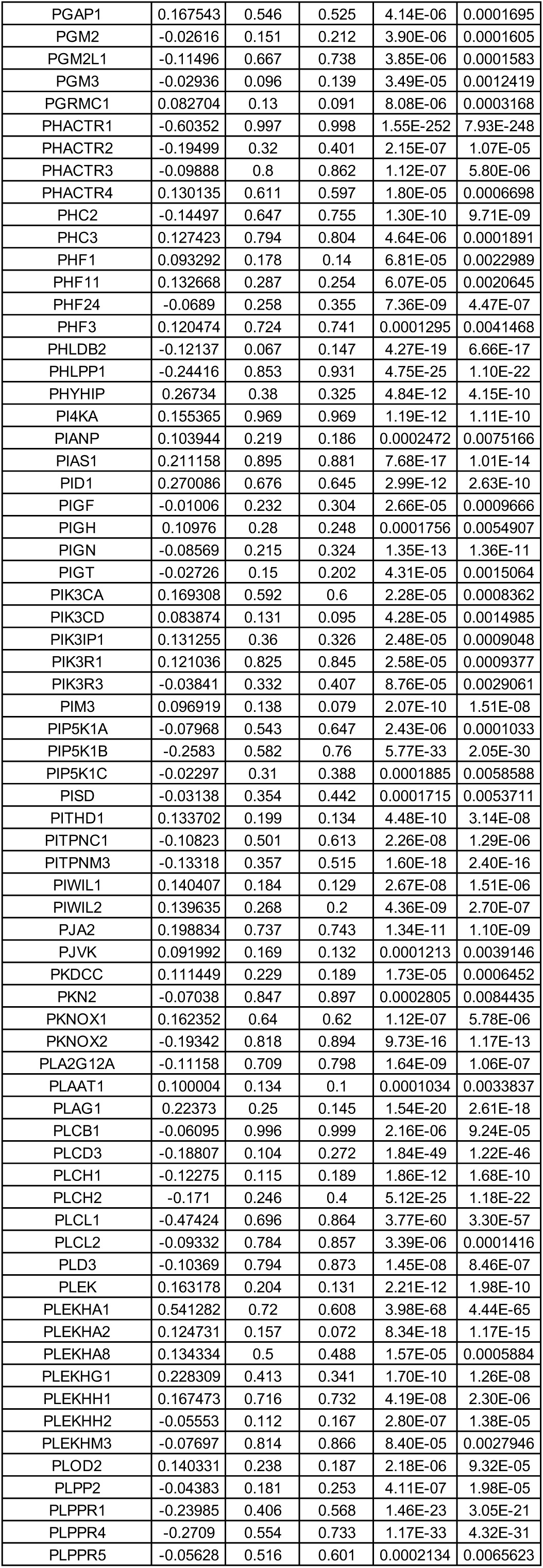
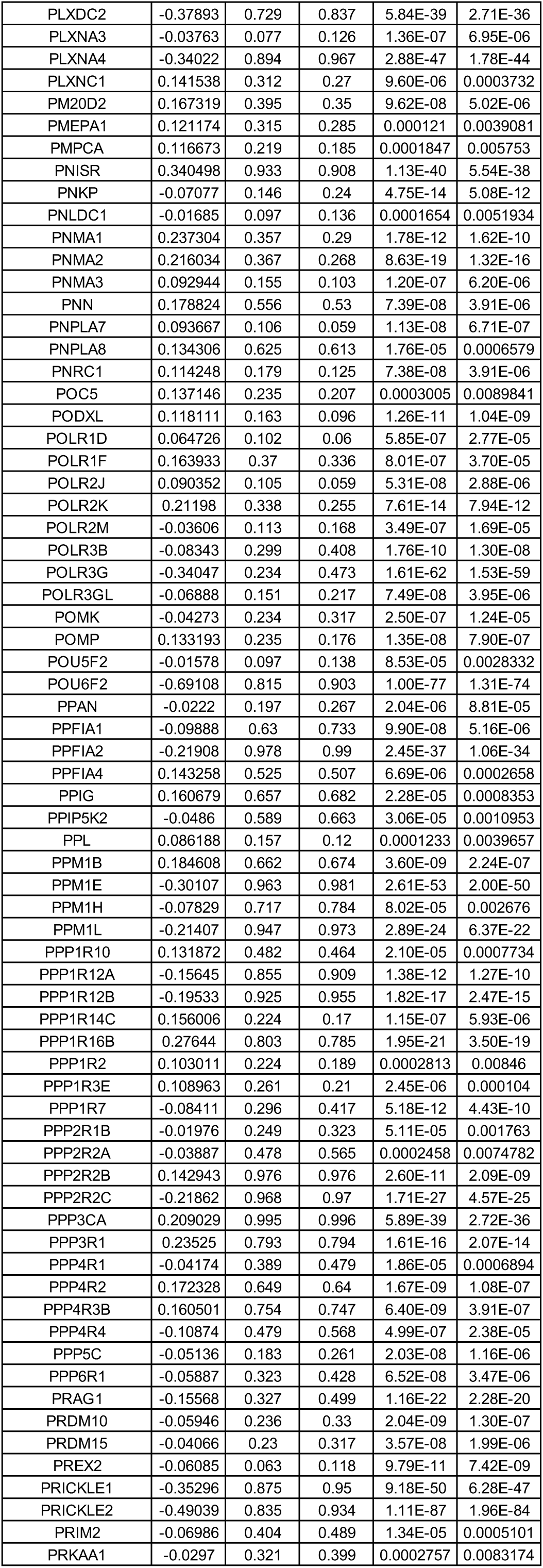
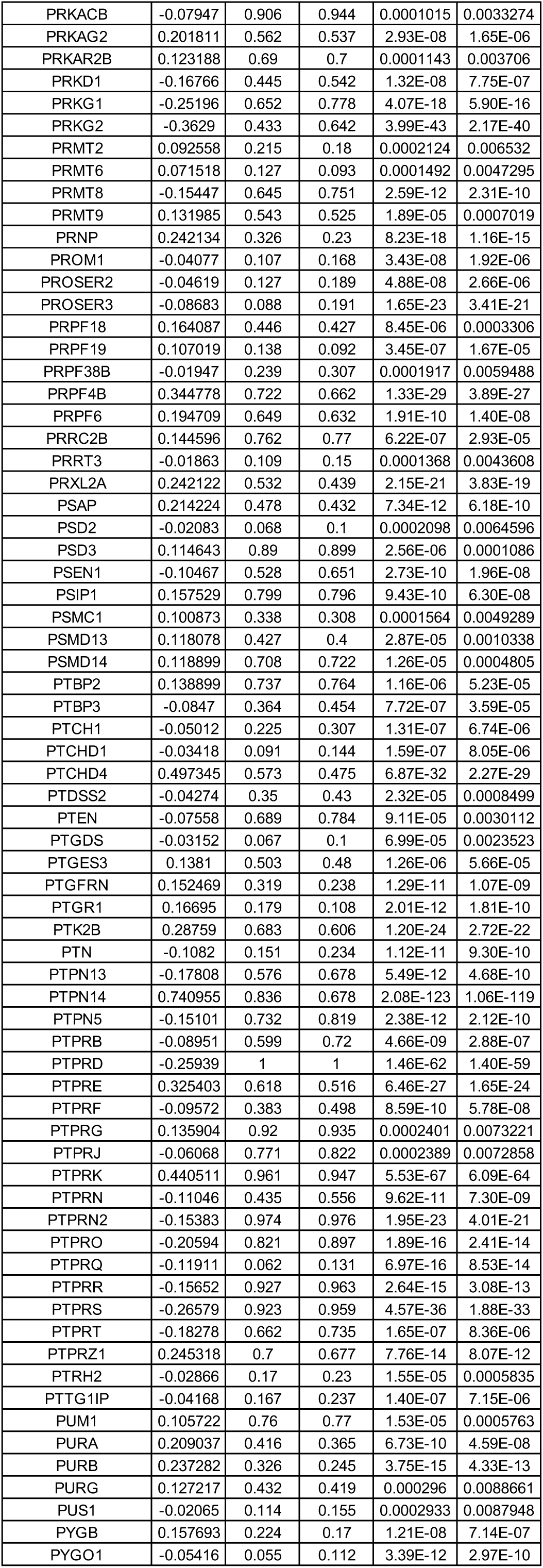
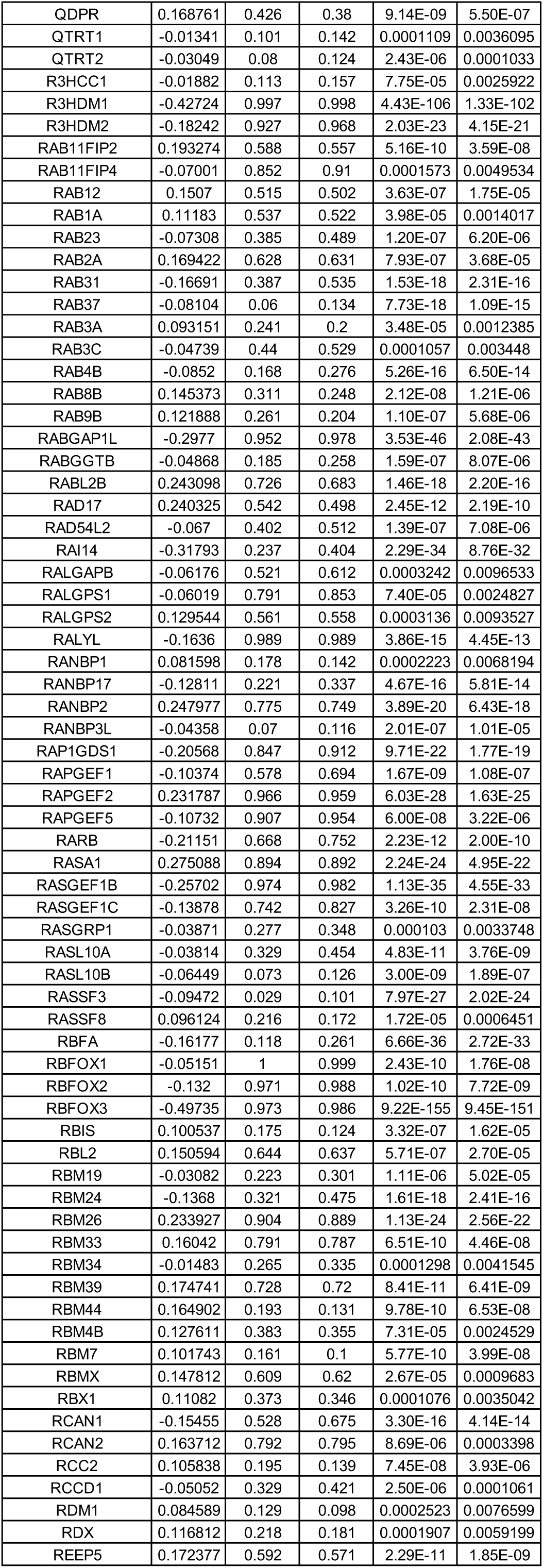
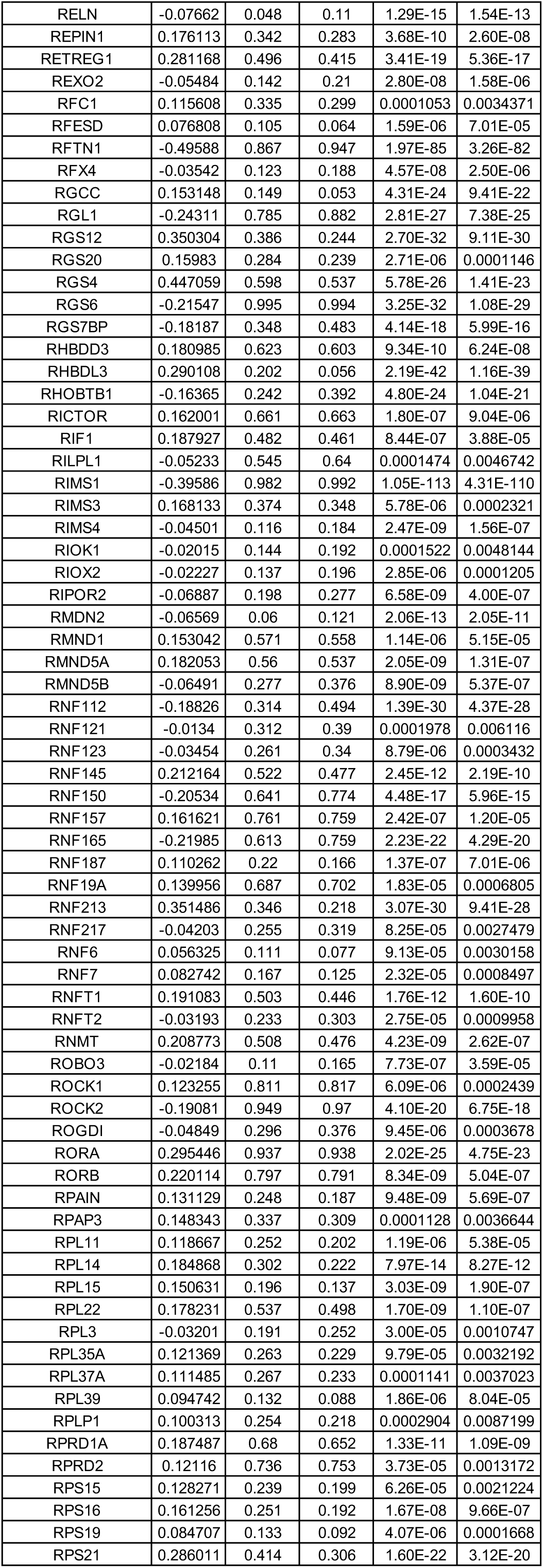
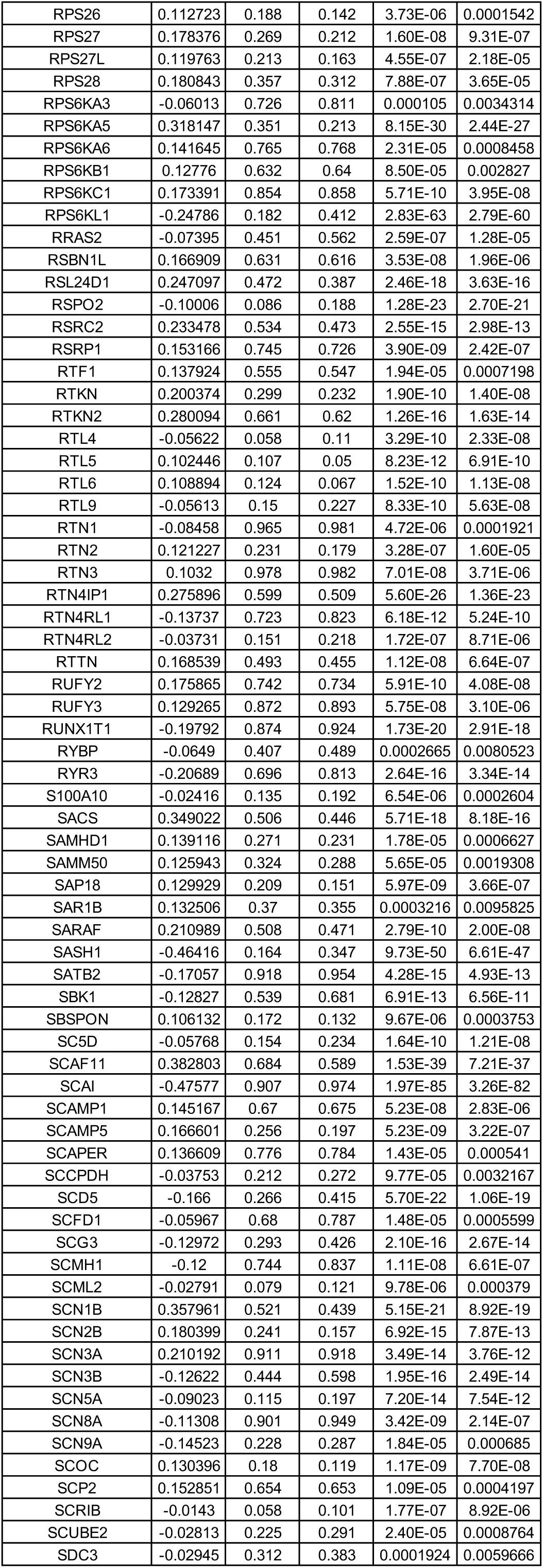
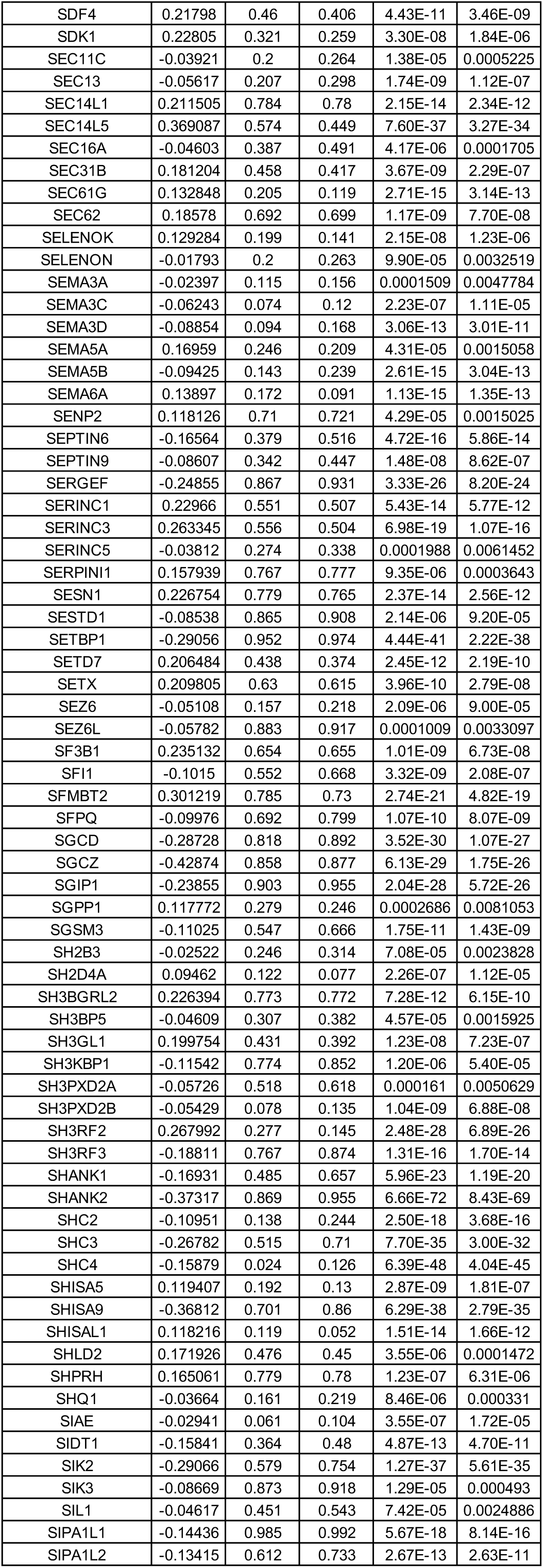
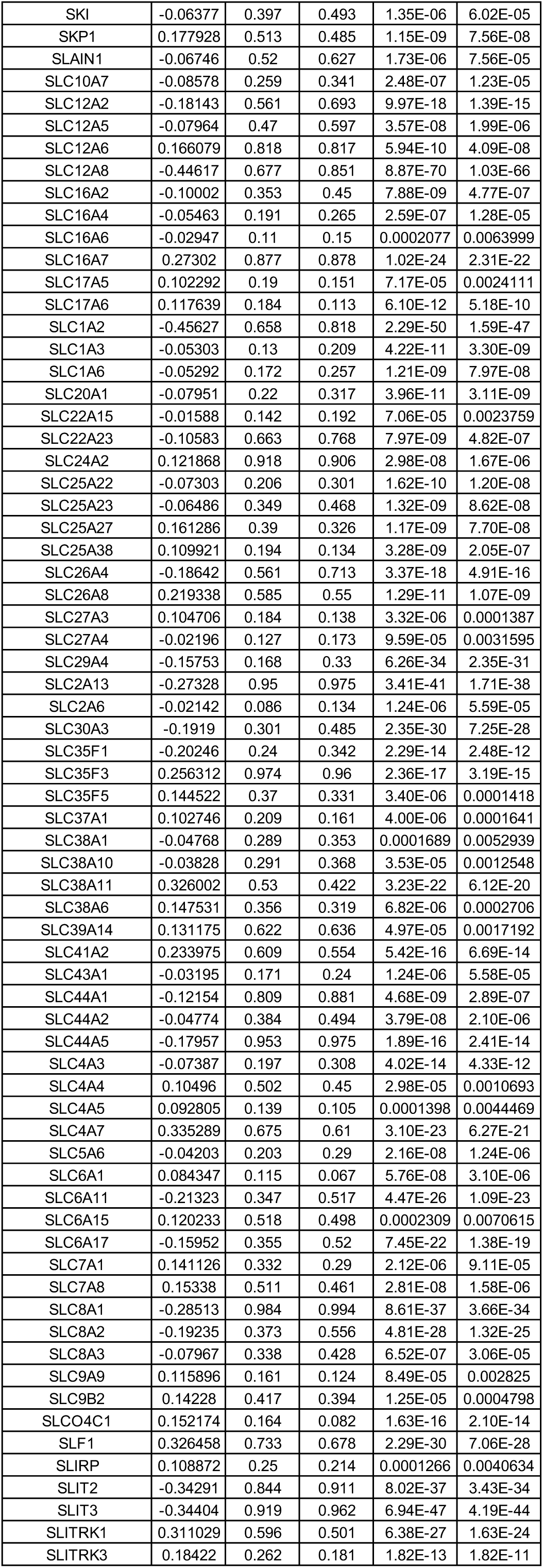
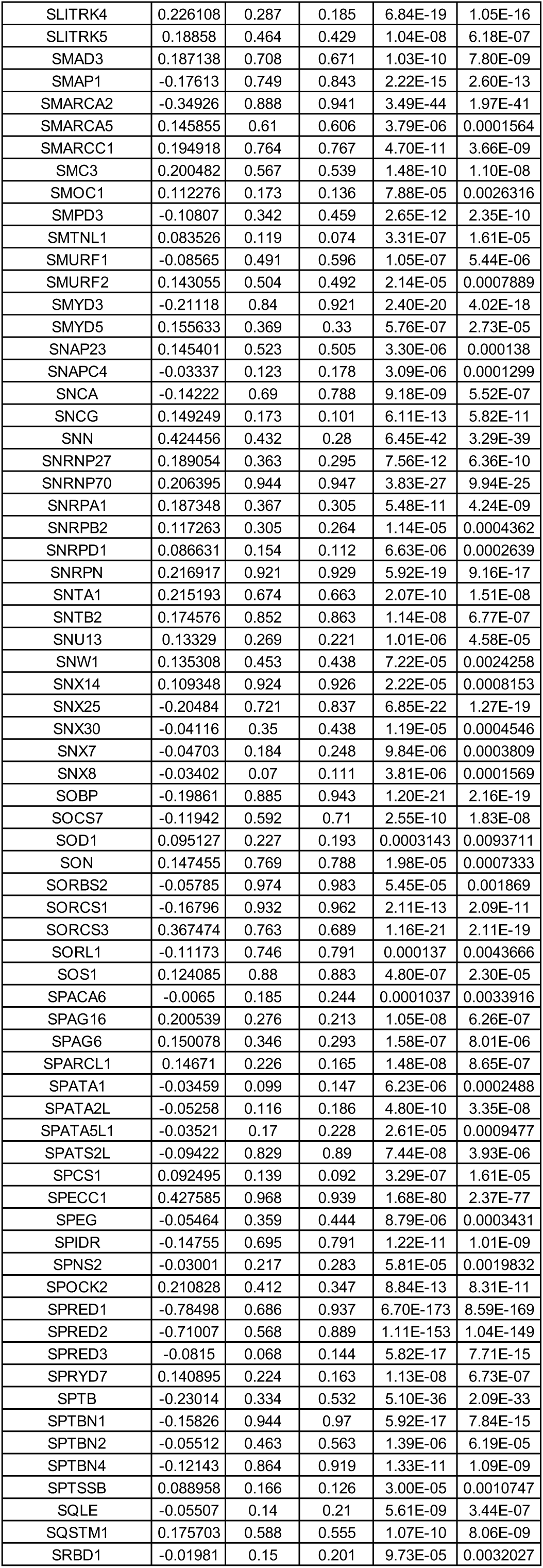
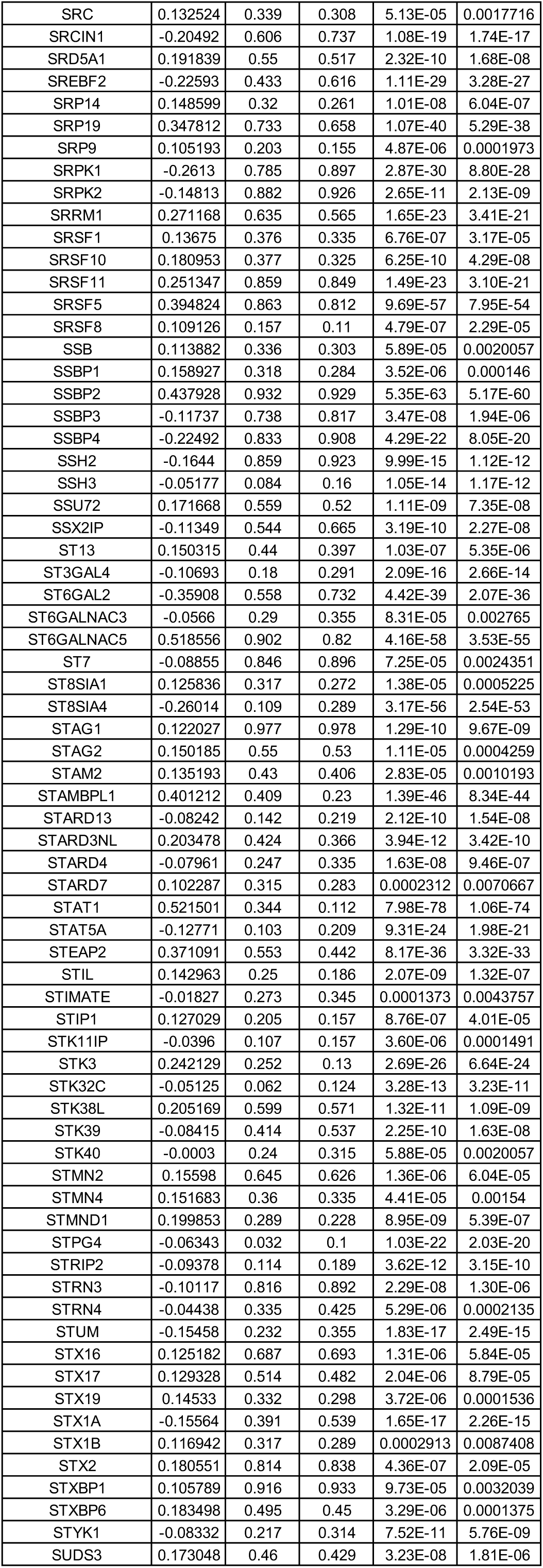
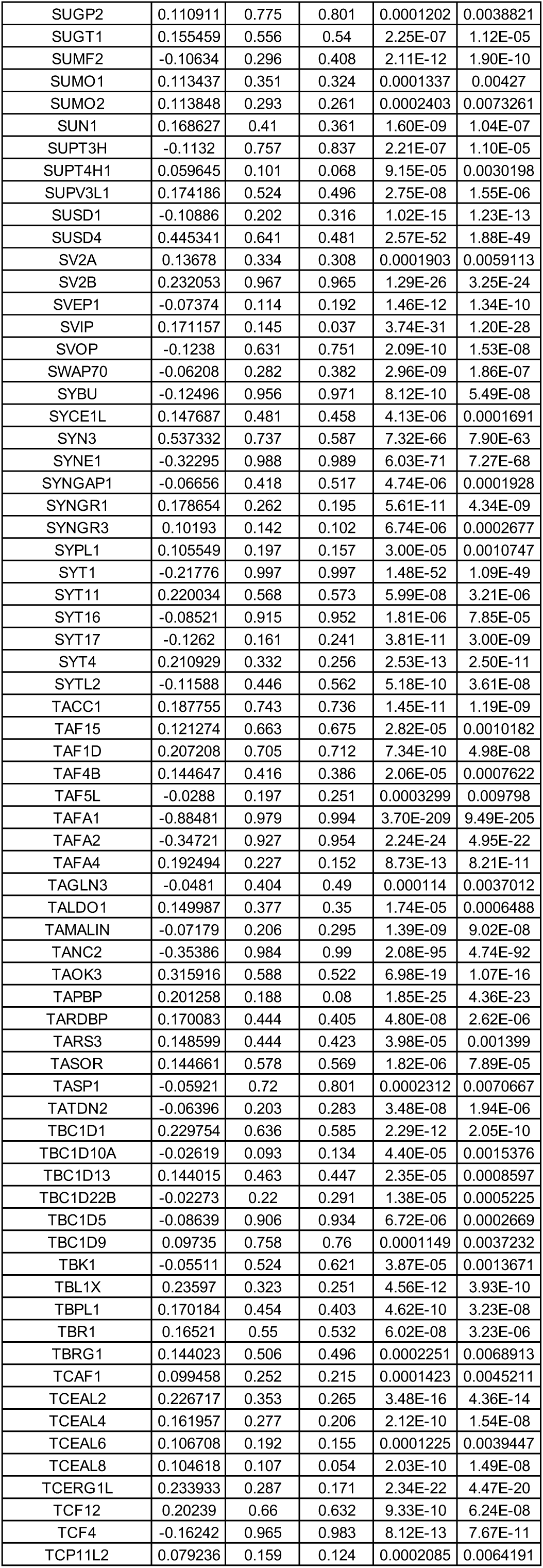
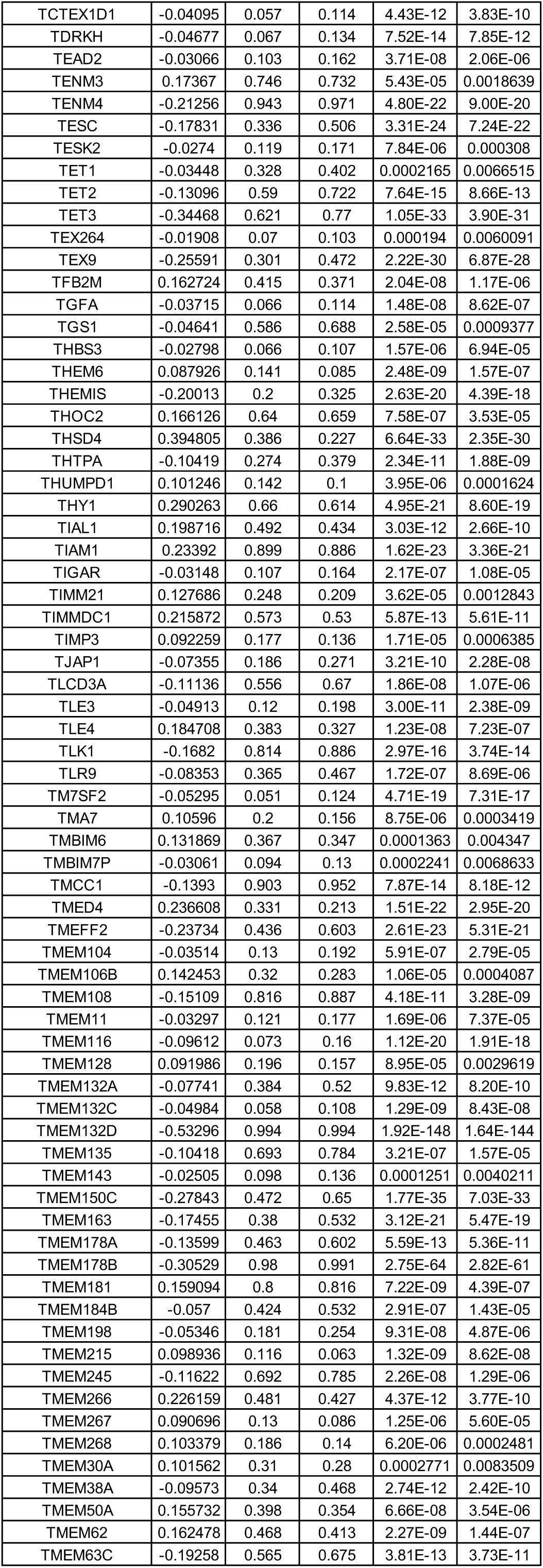
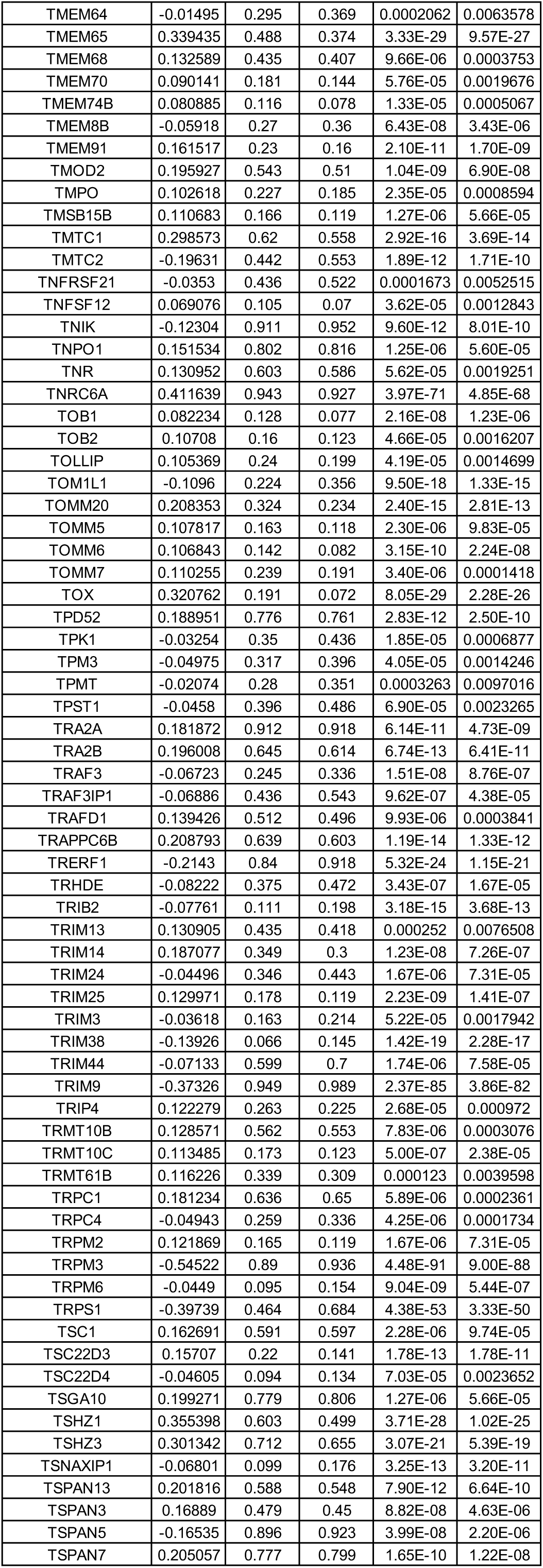
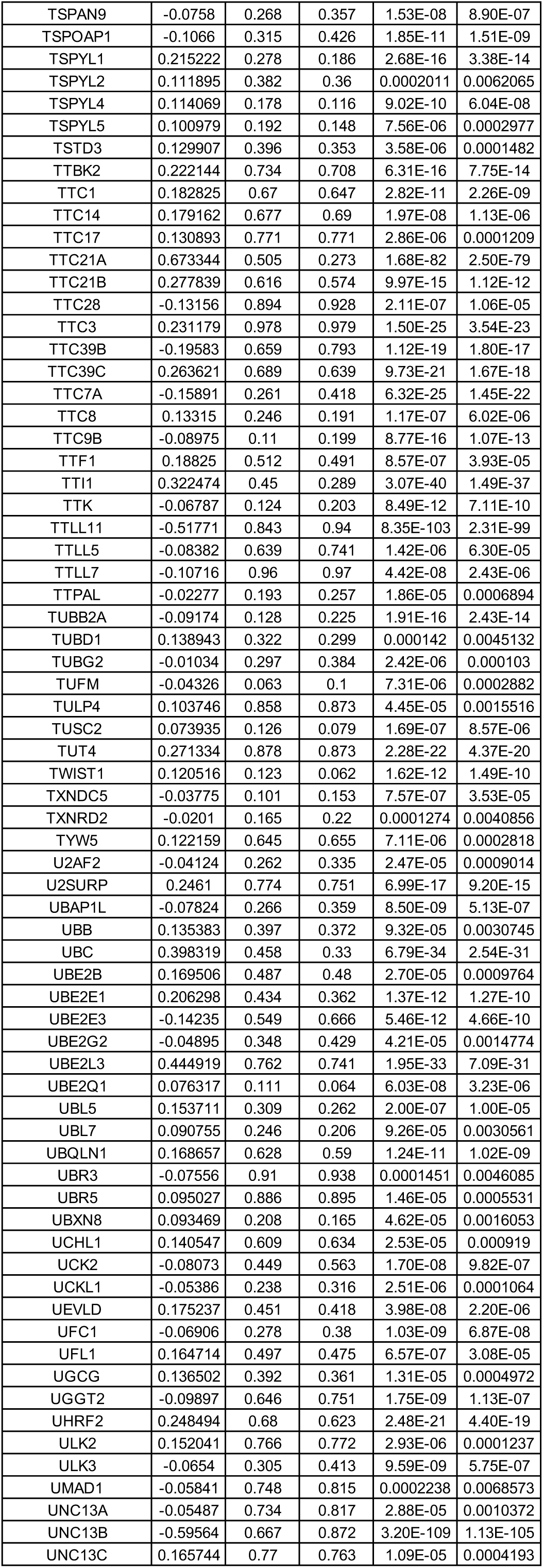
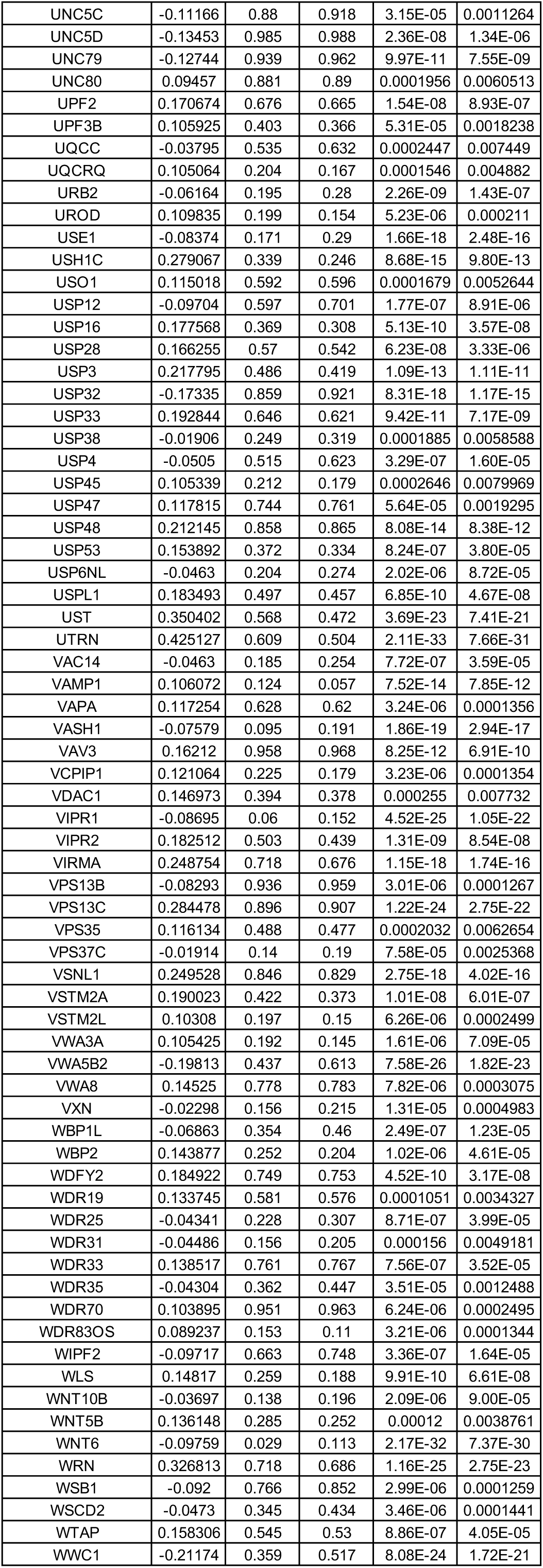
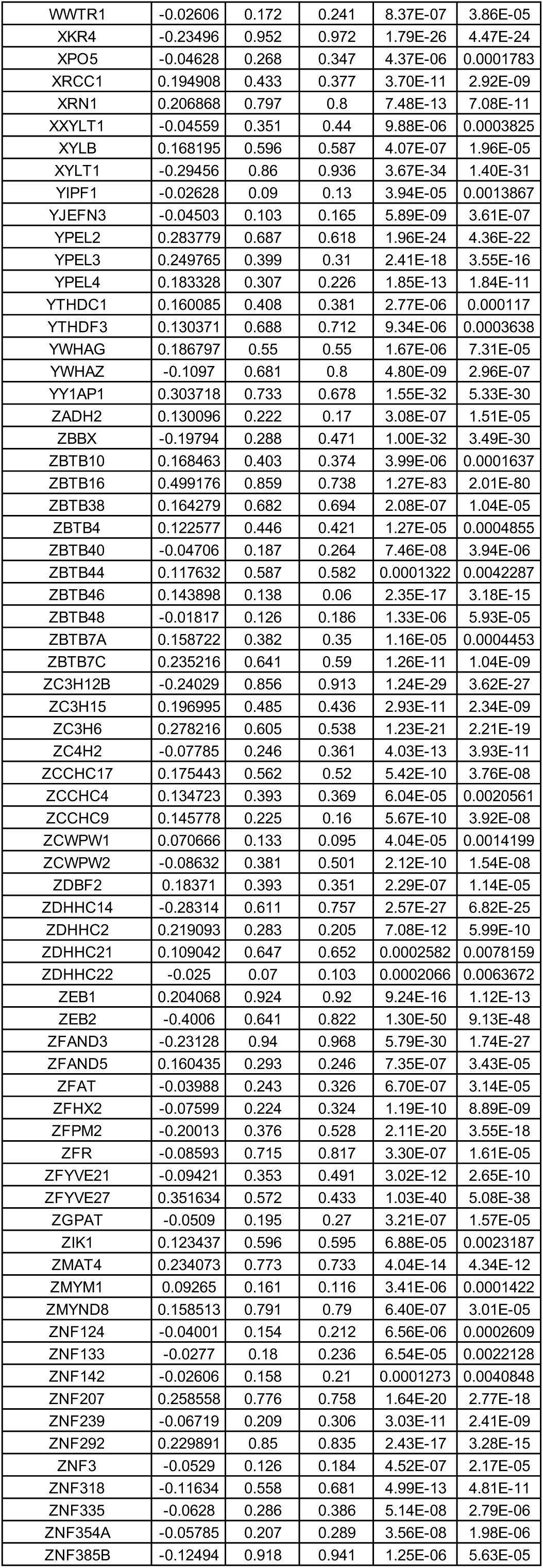
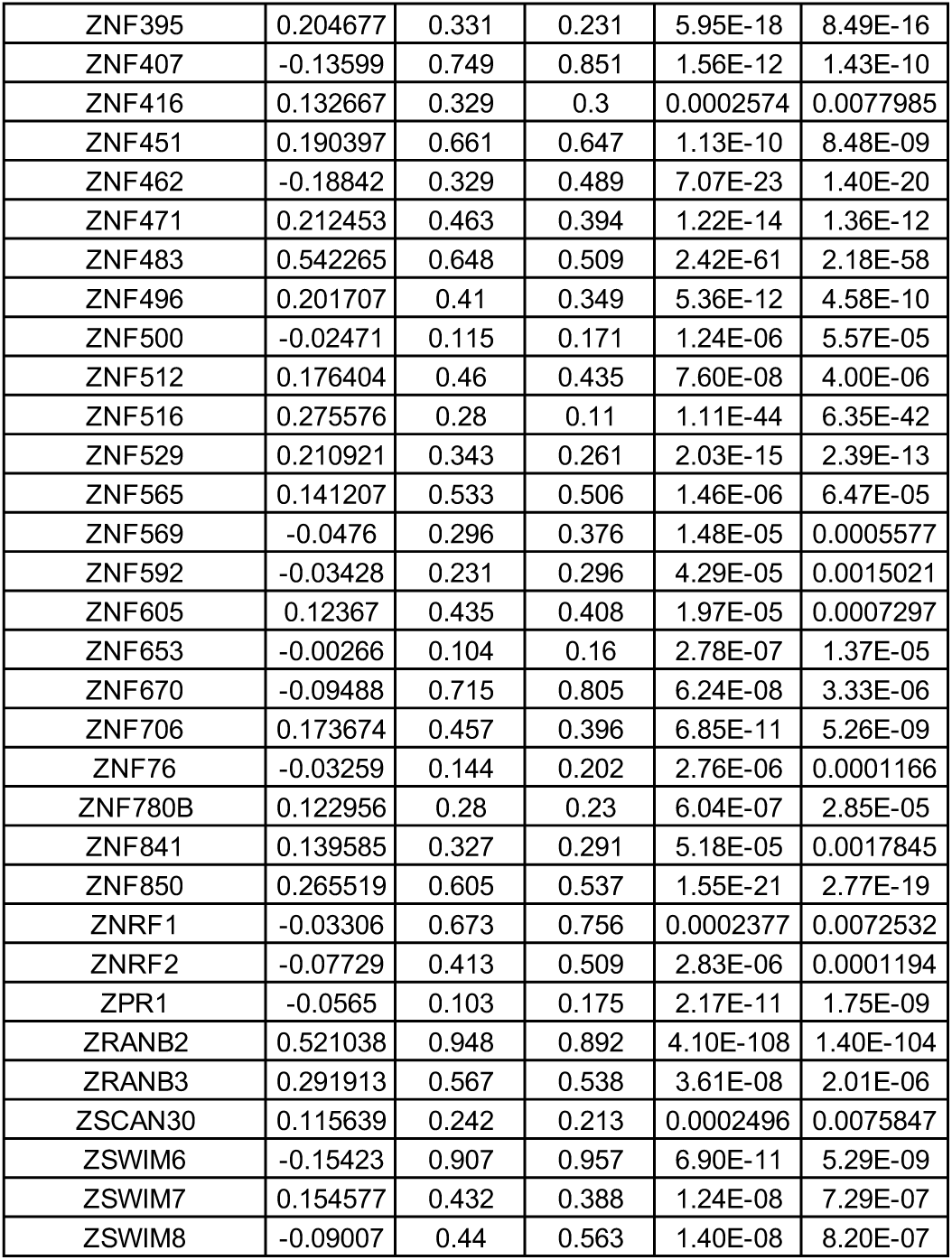
DEGs between wild-type (WT) and MECP2-null (KO) upper layer excitatory neurons (Ex_1) of OC (adjusted p-value < 0.01)

**Table S23.**
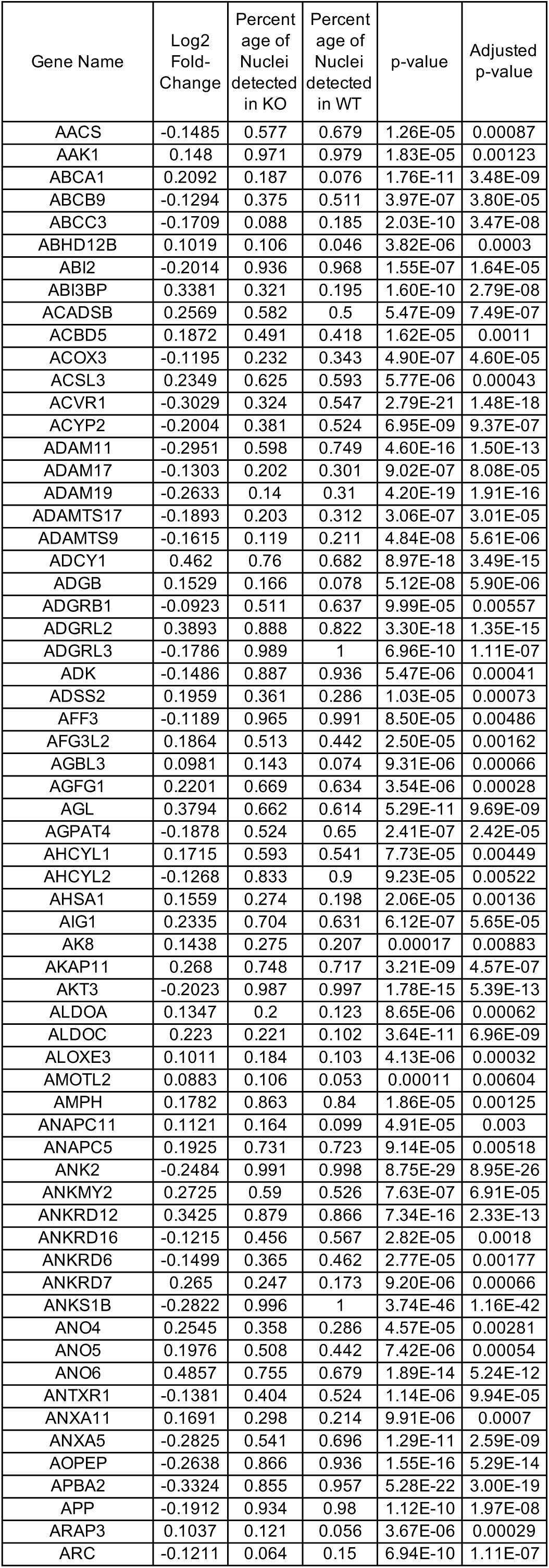
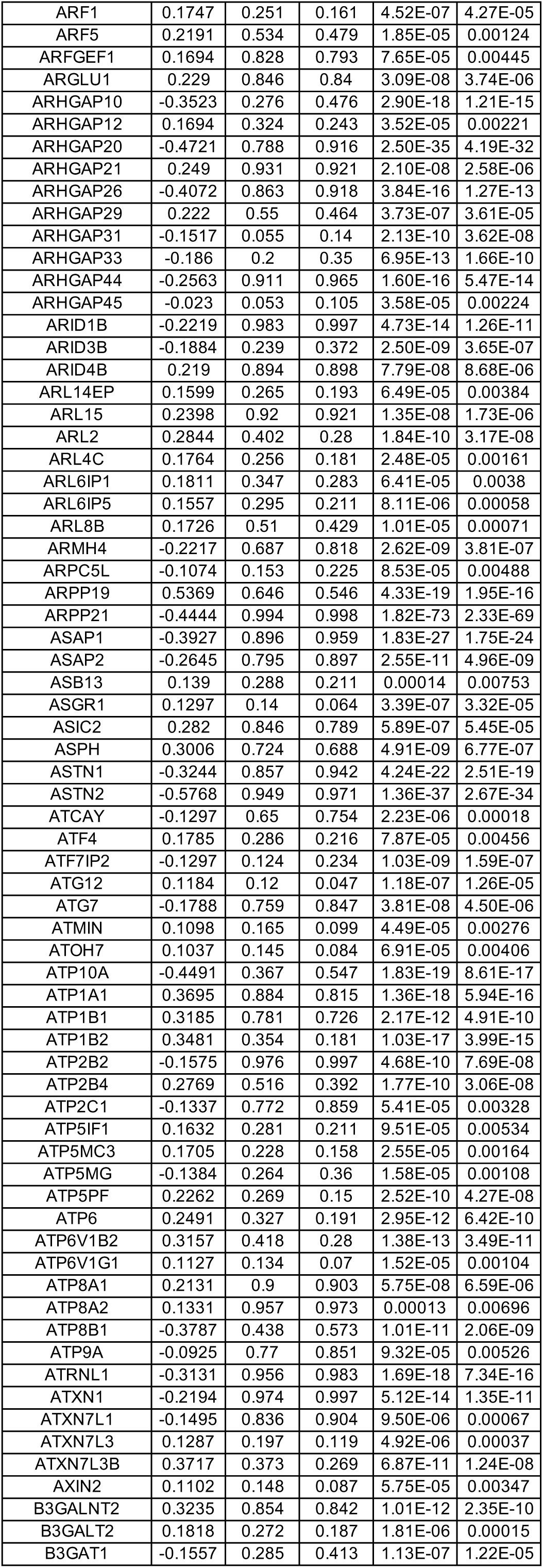
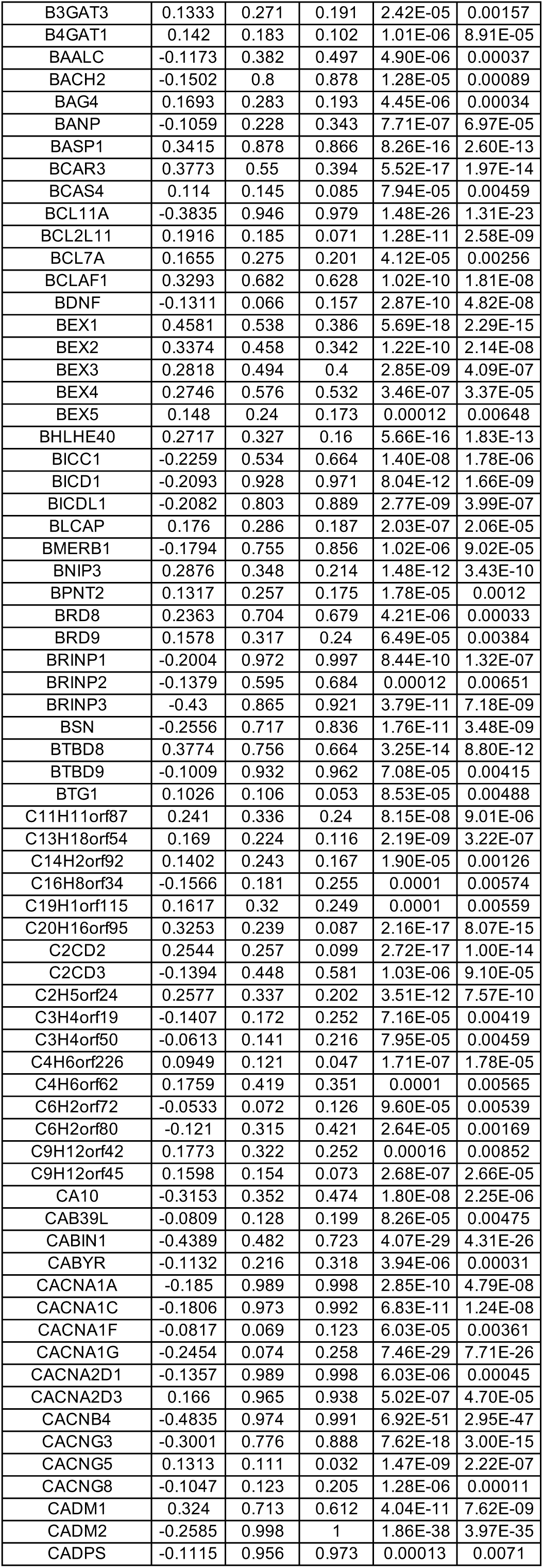
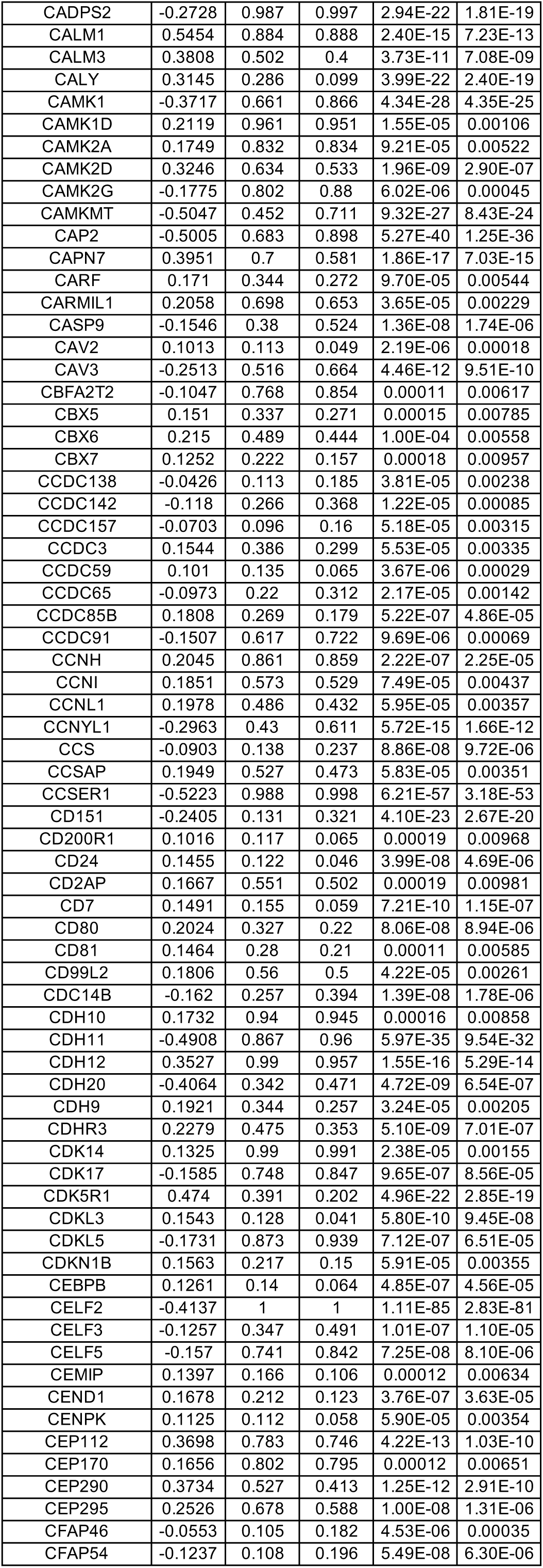
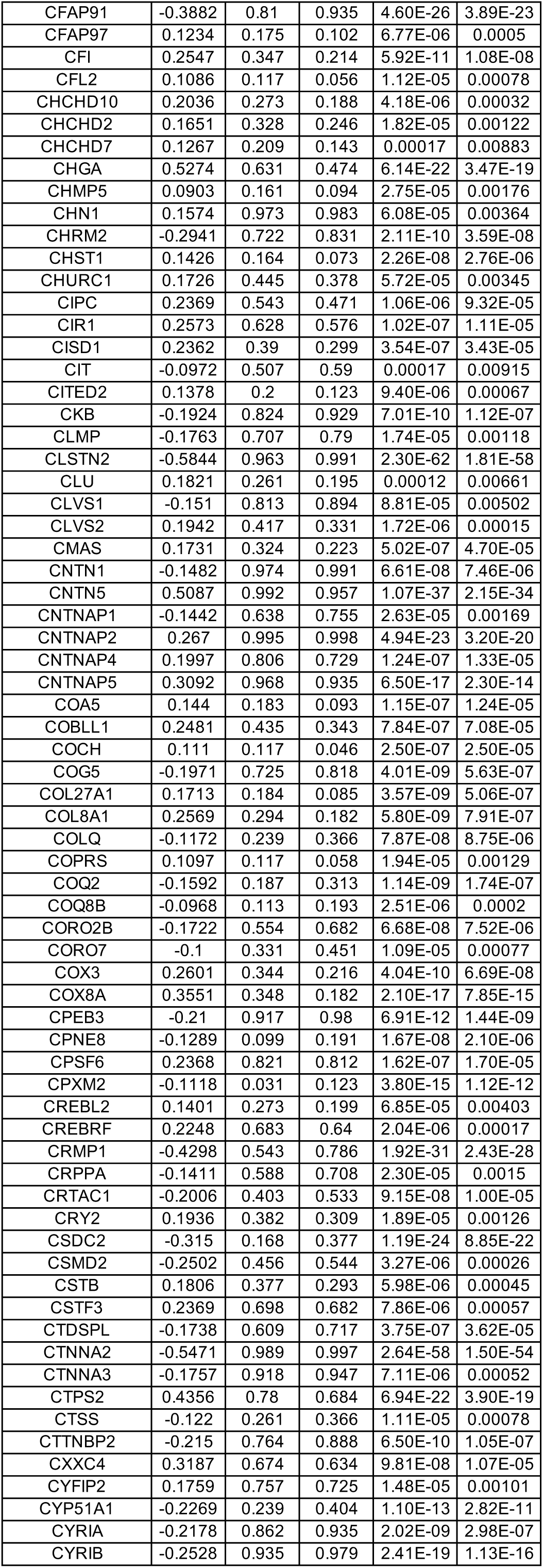
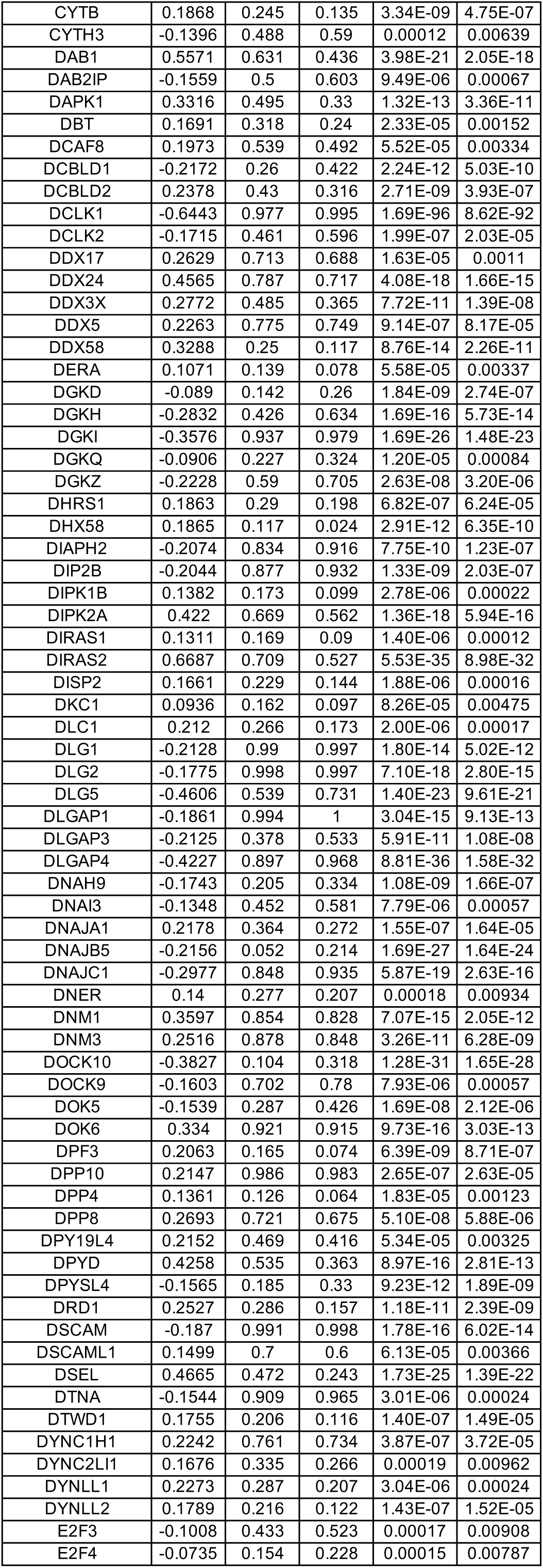
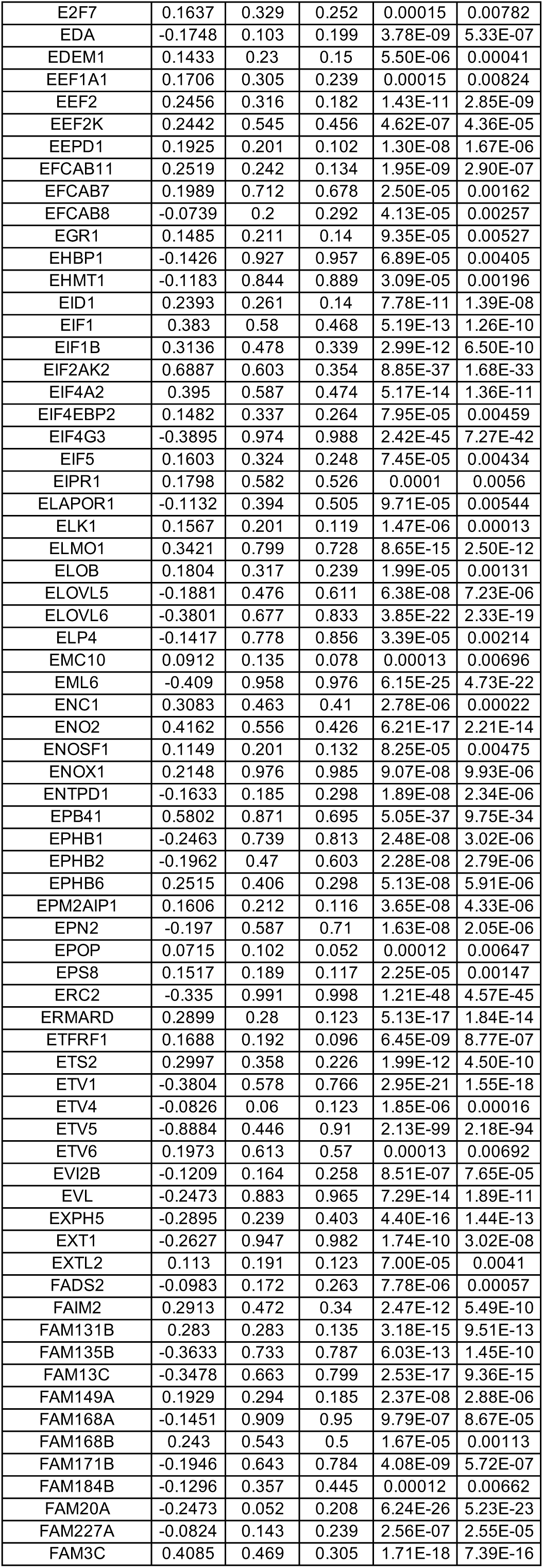
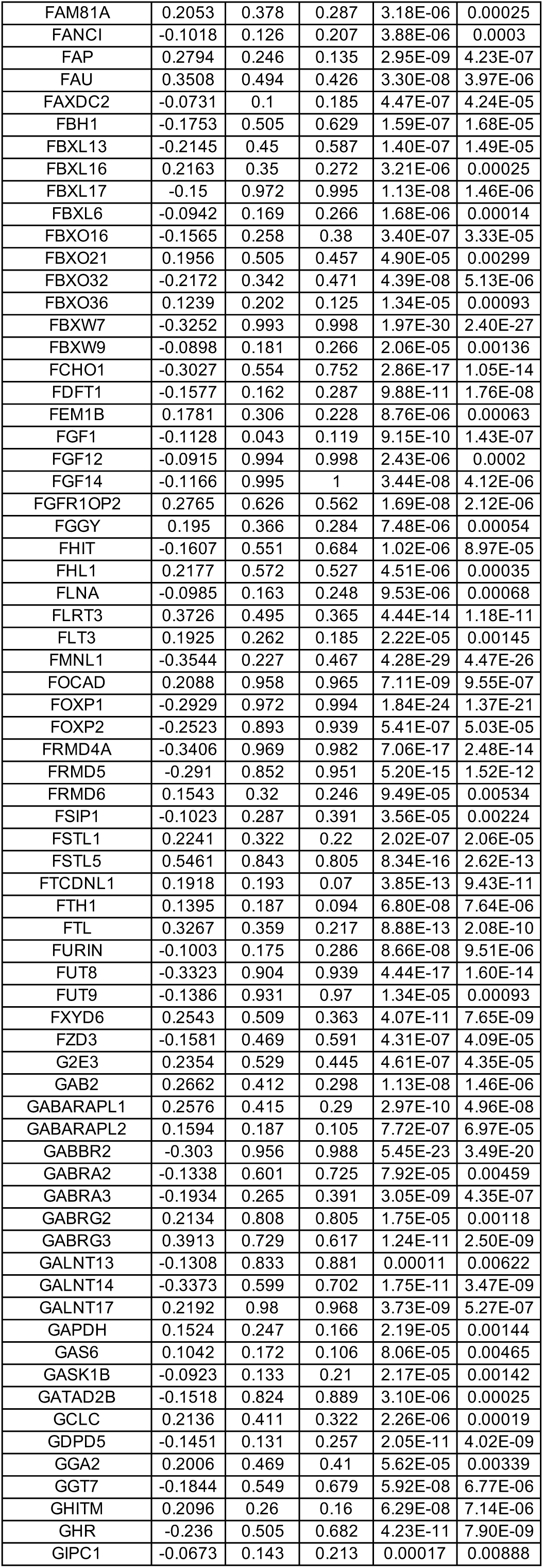
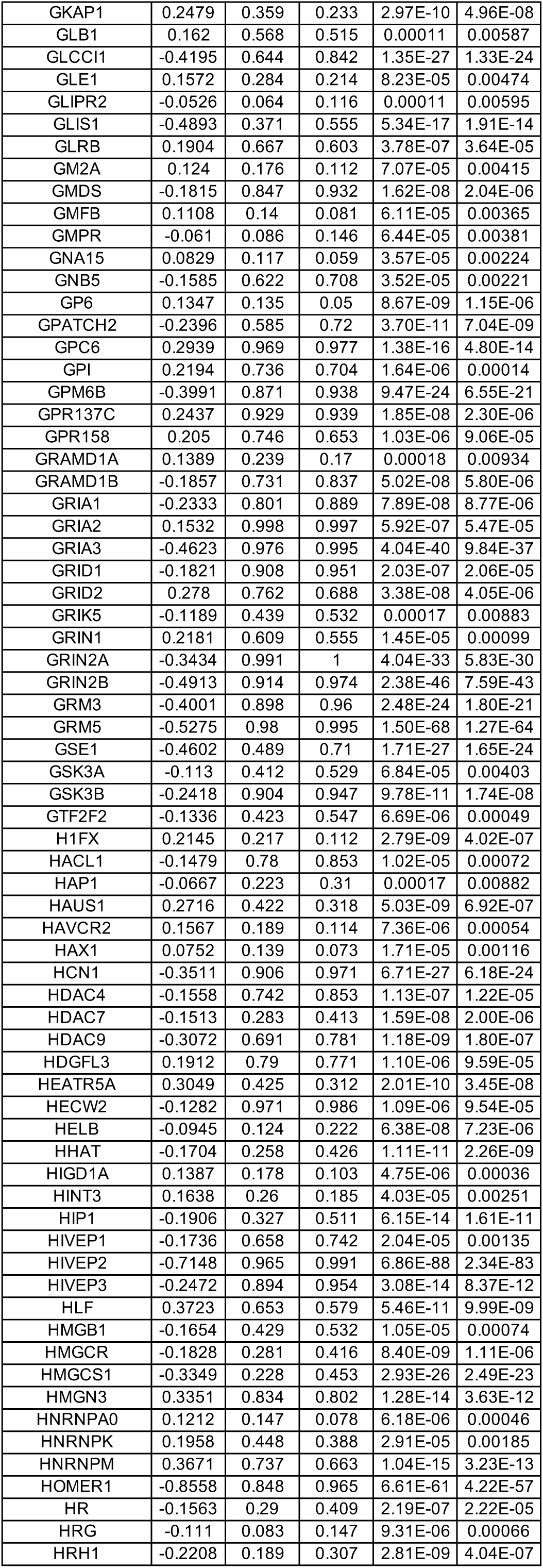
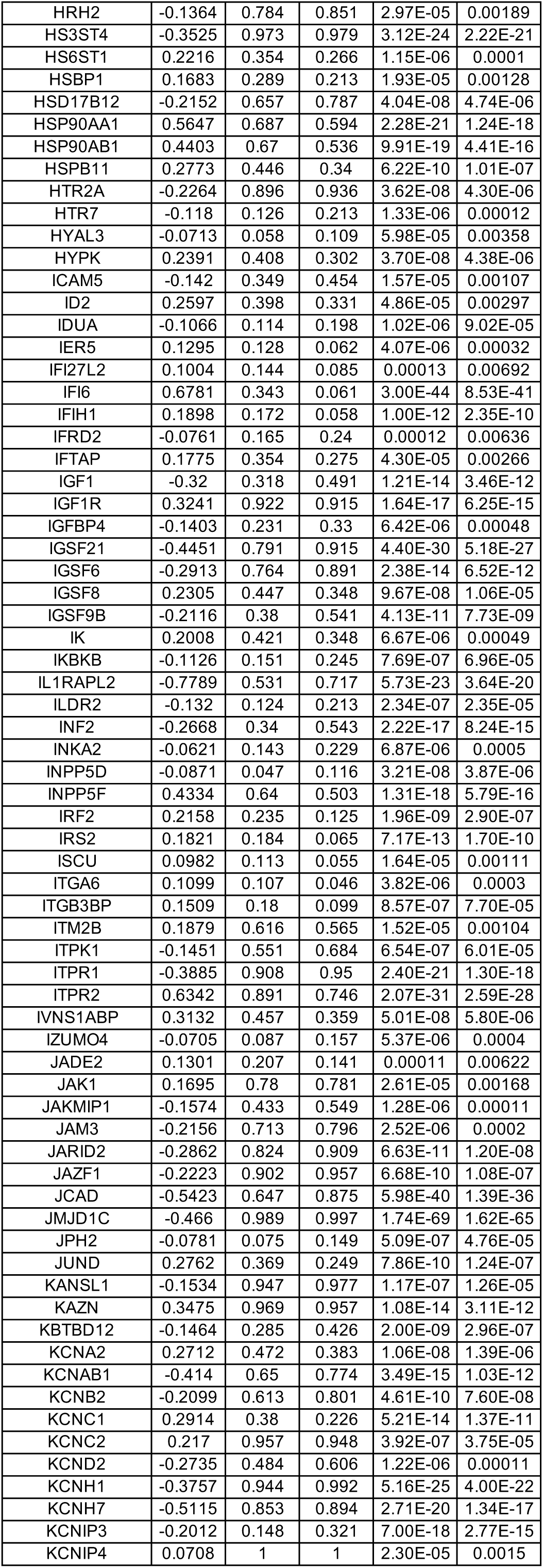
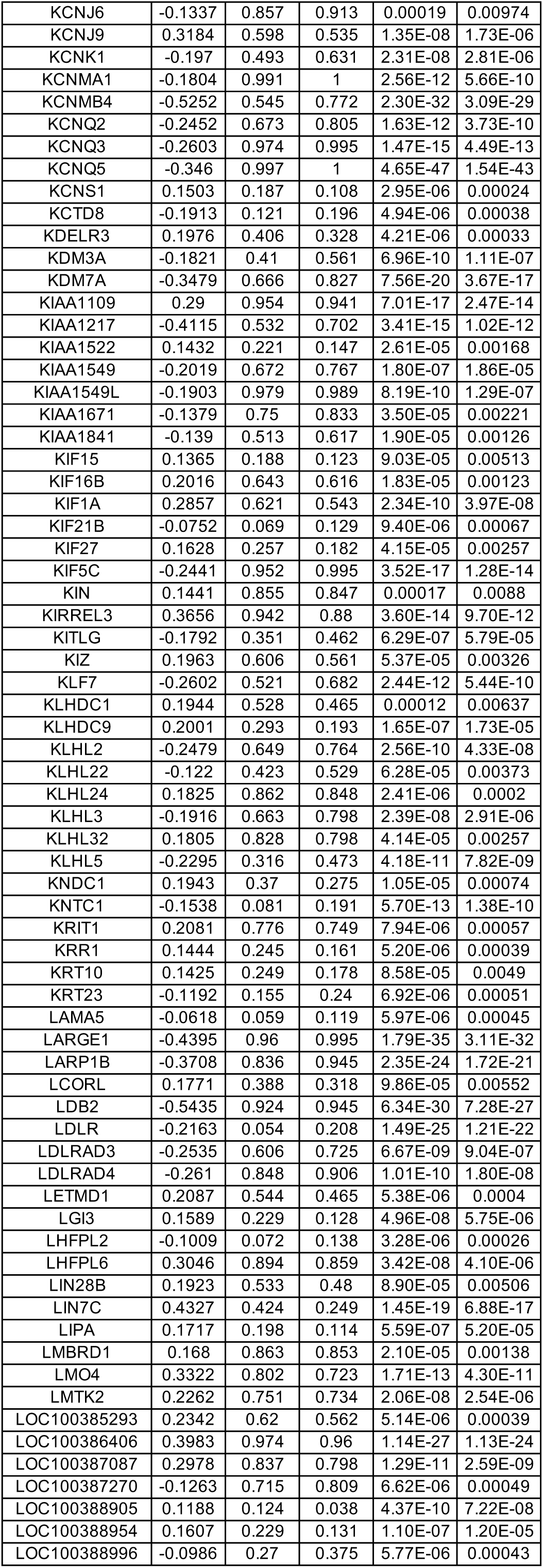
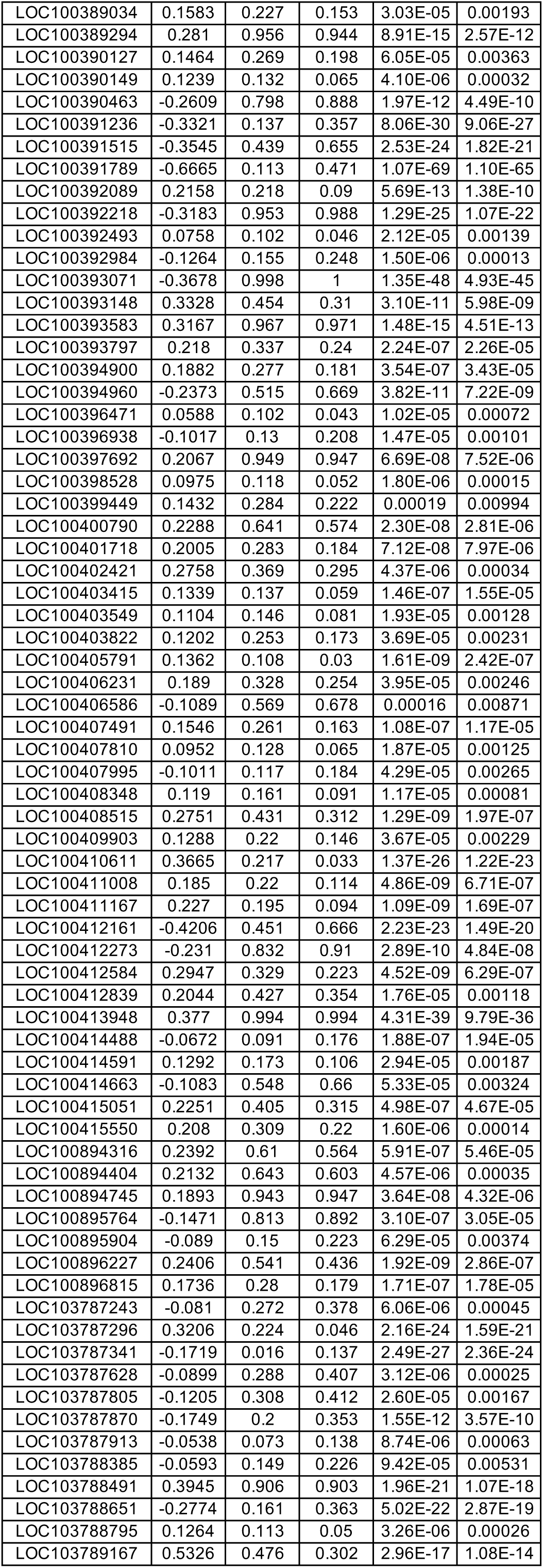
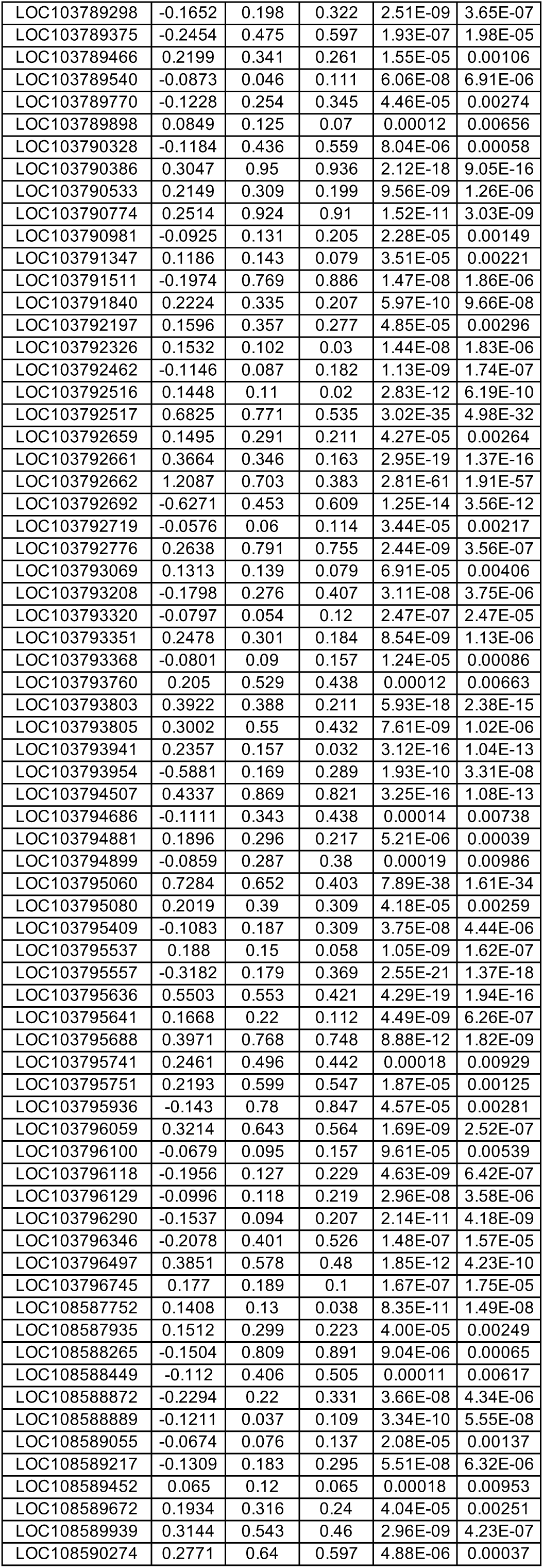
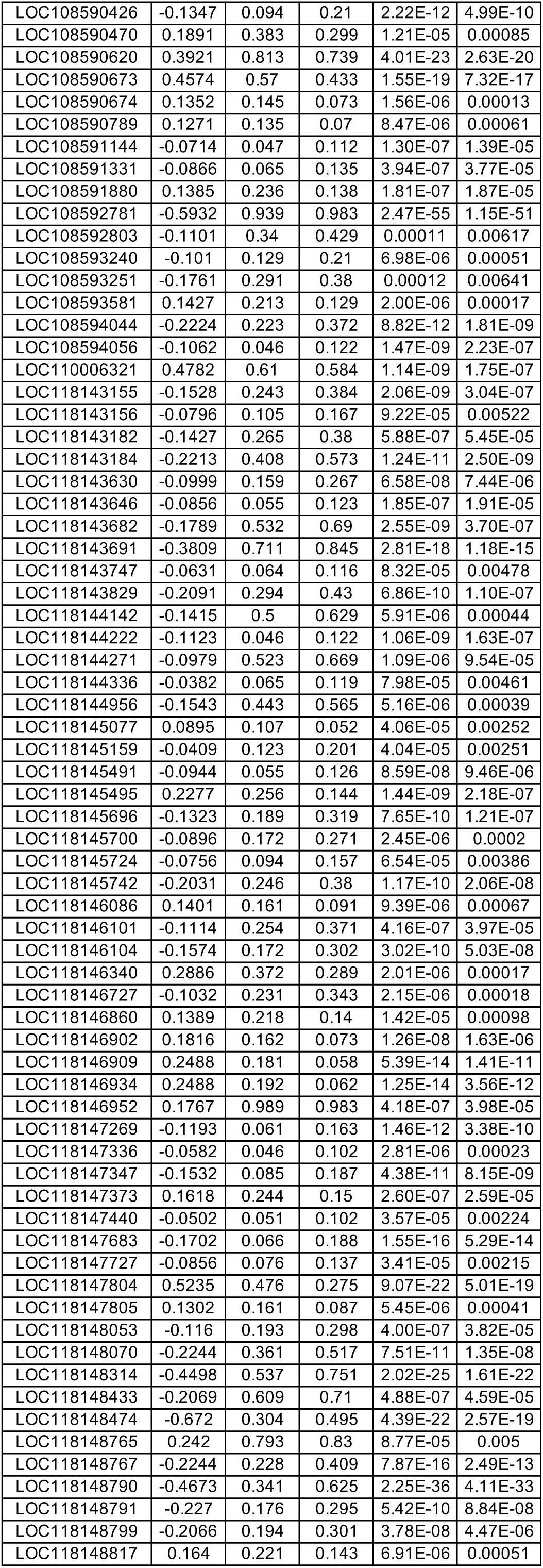
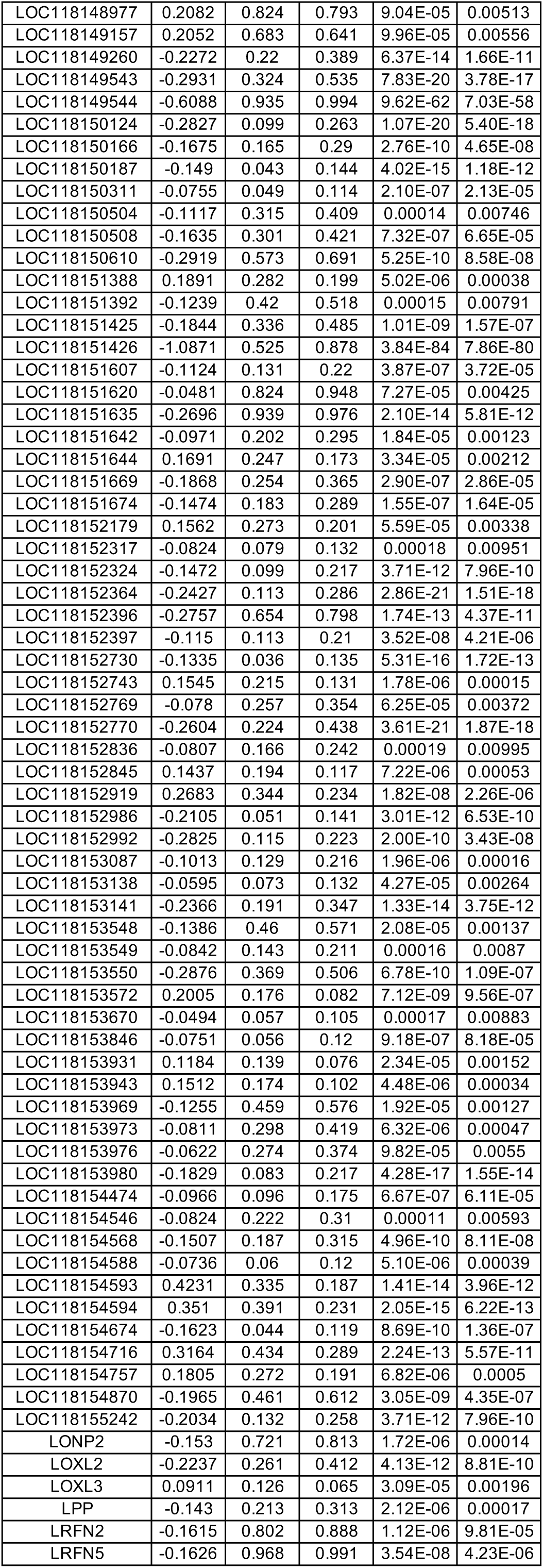
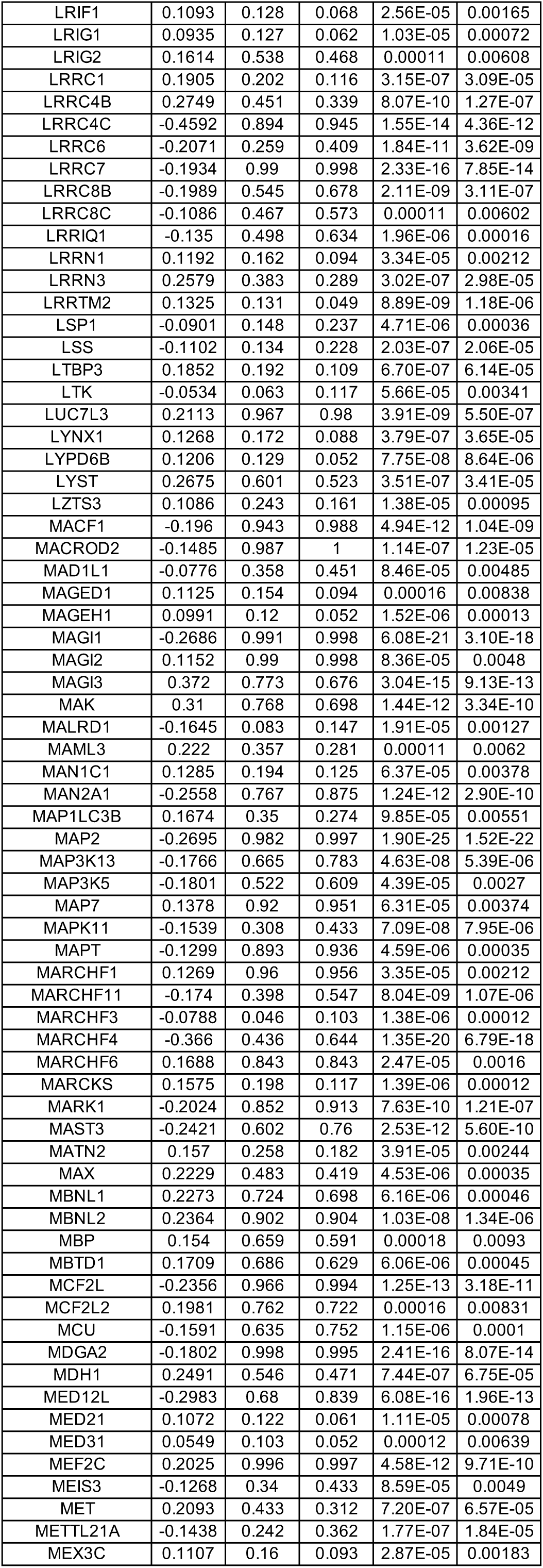
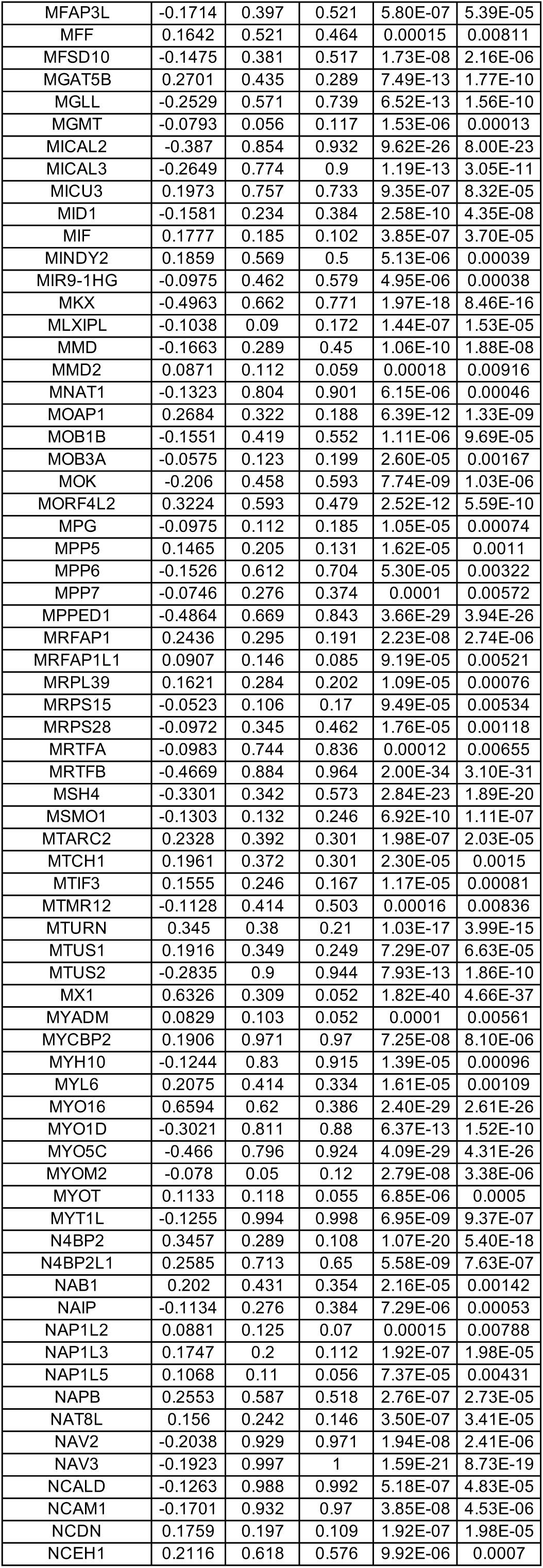
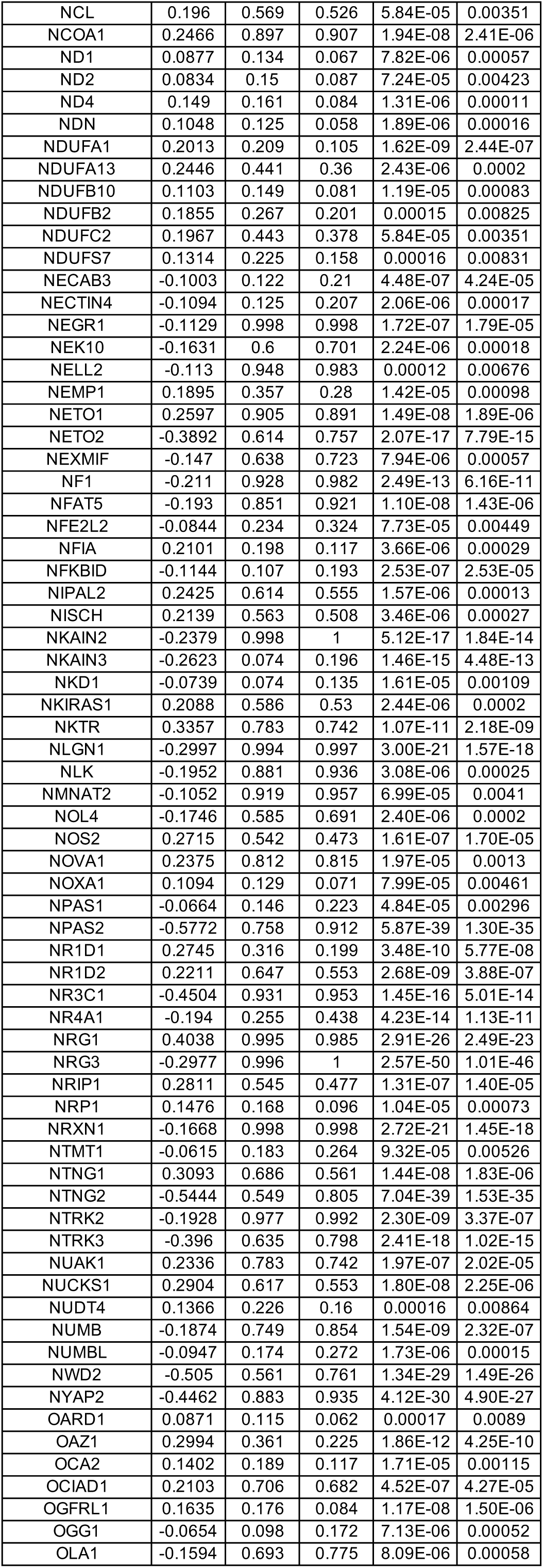
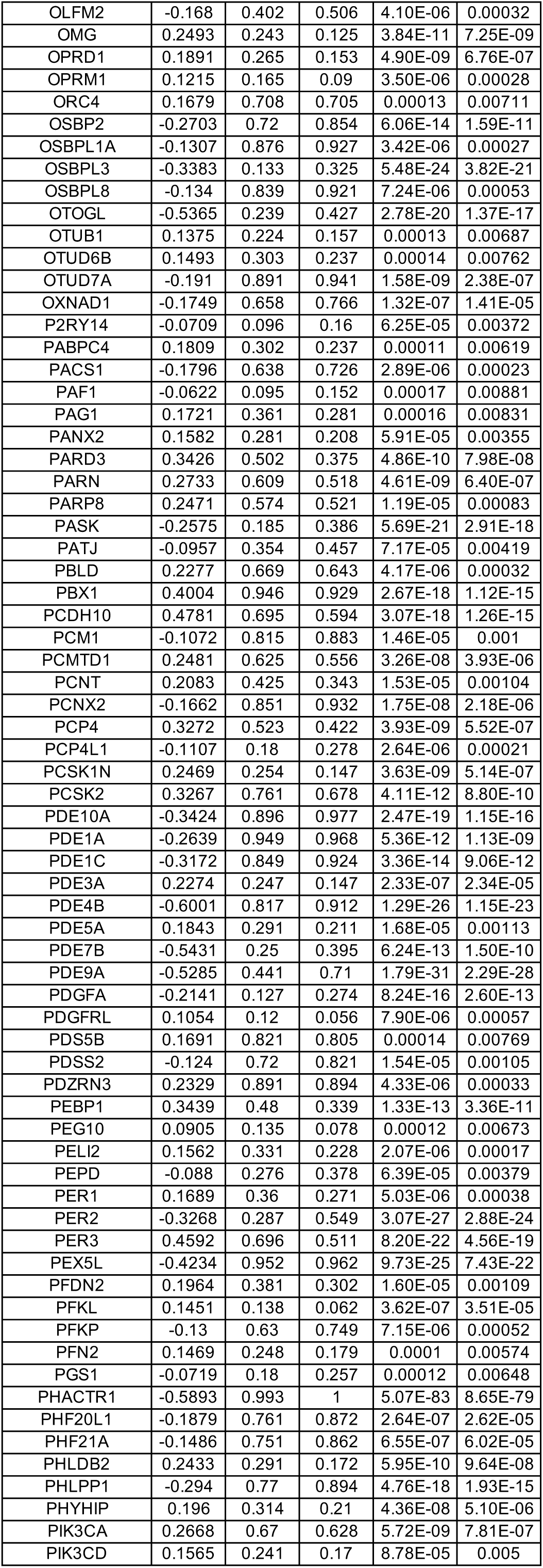
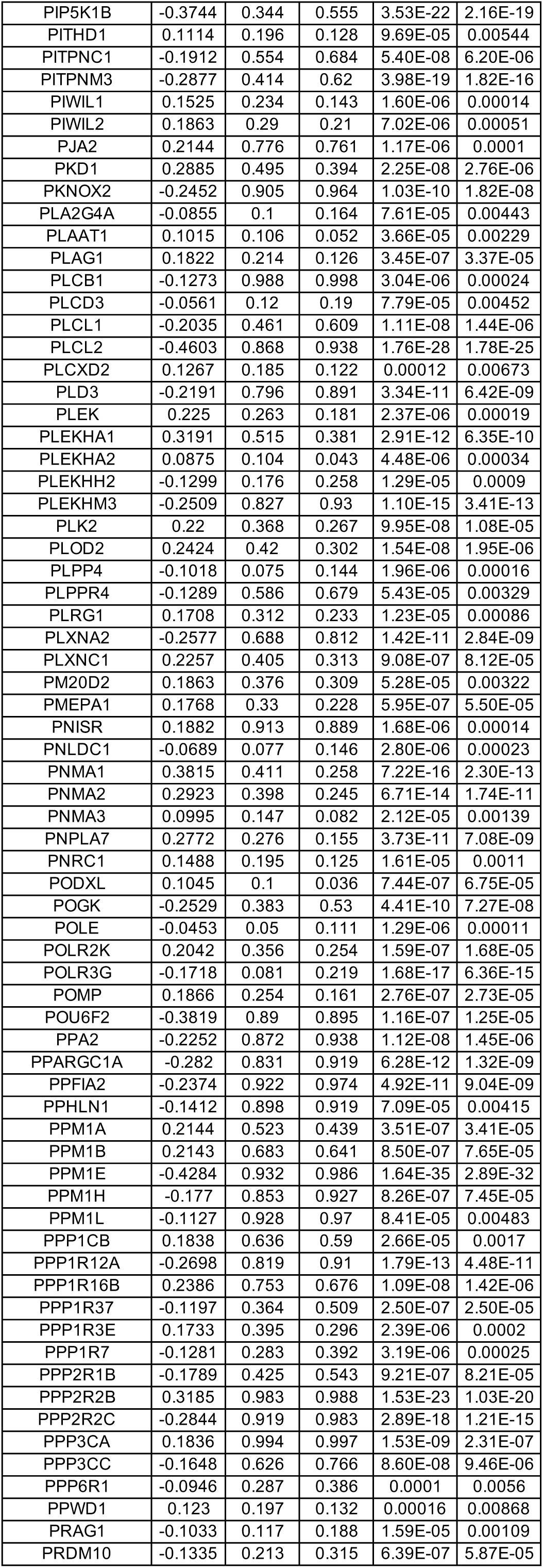
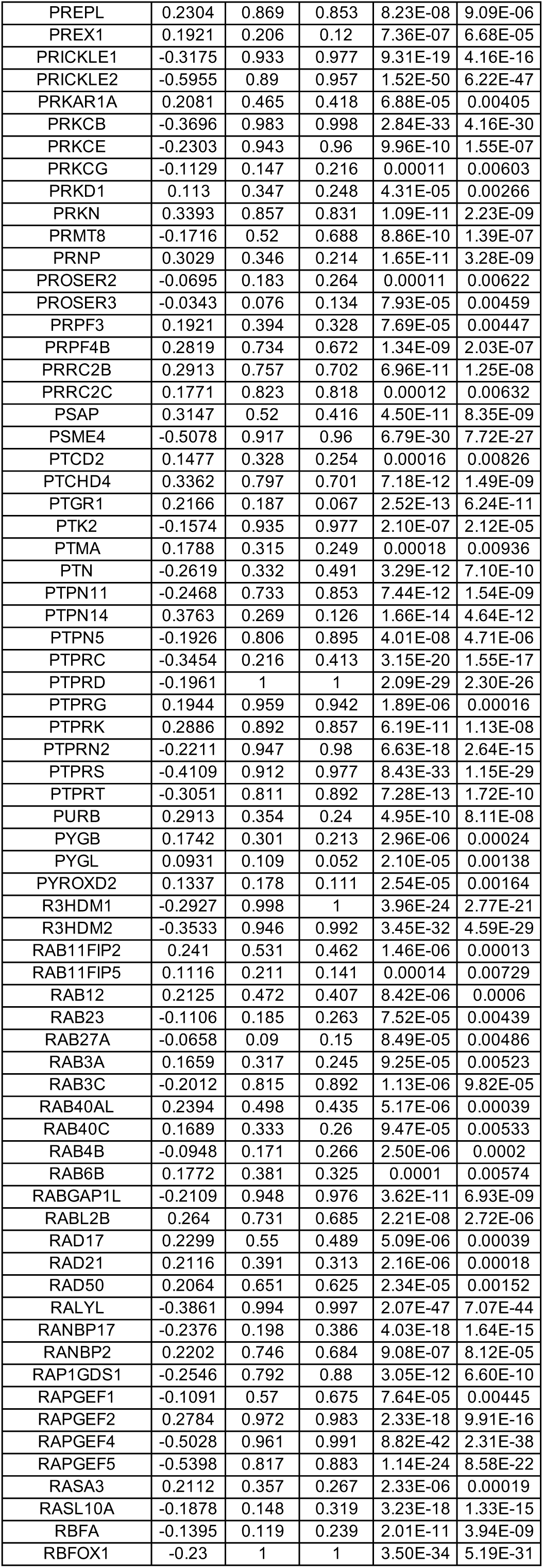
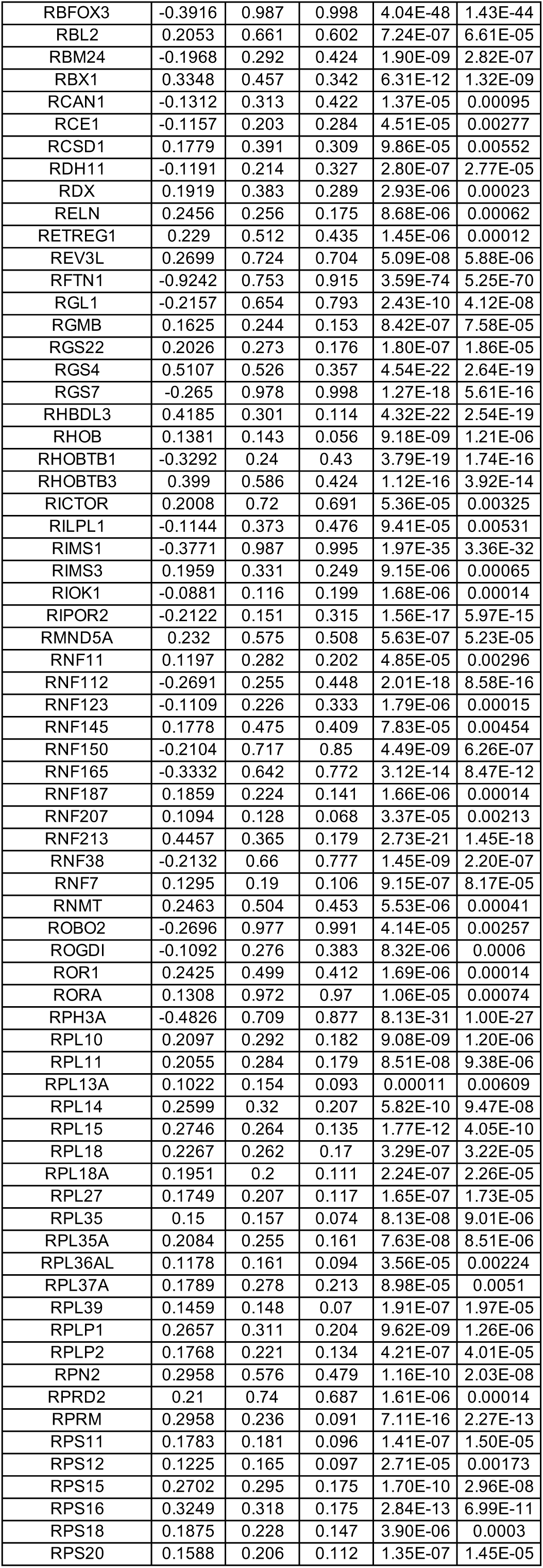
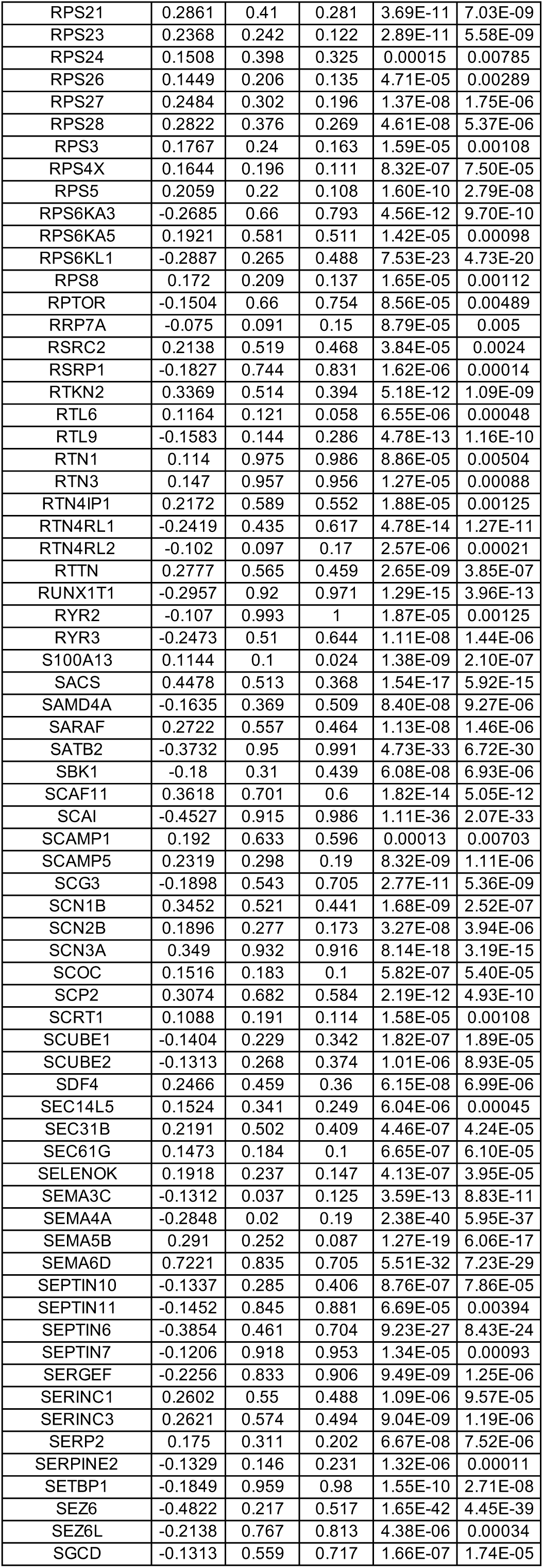
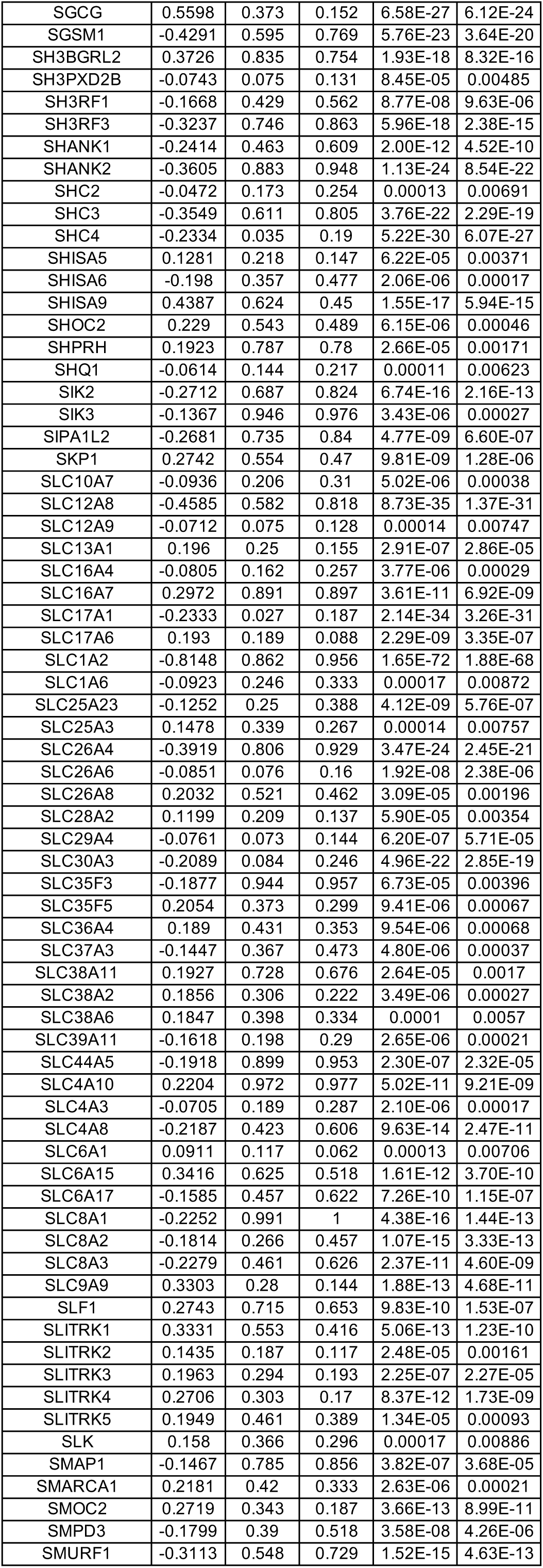
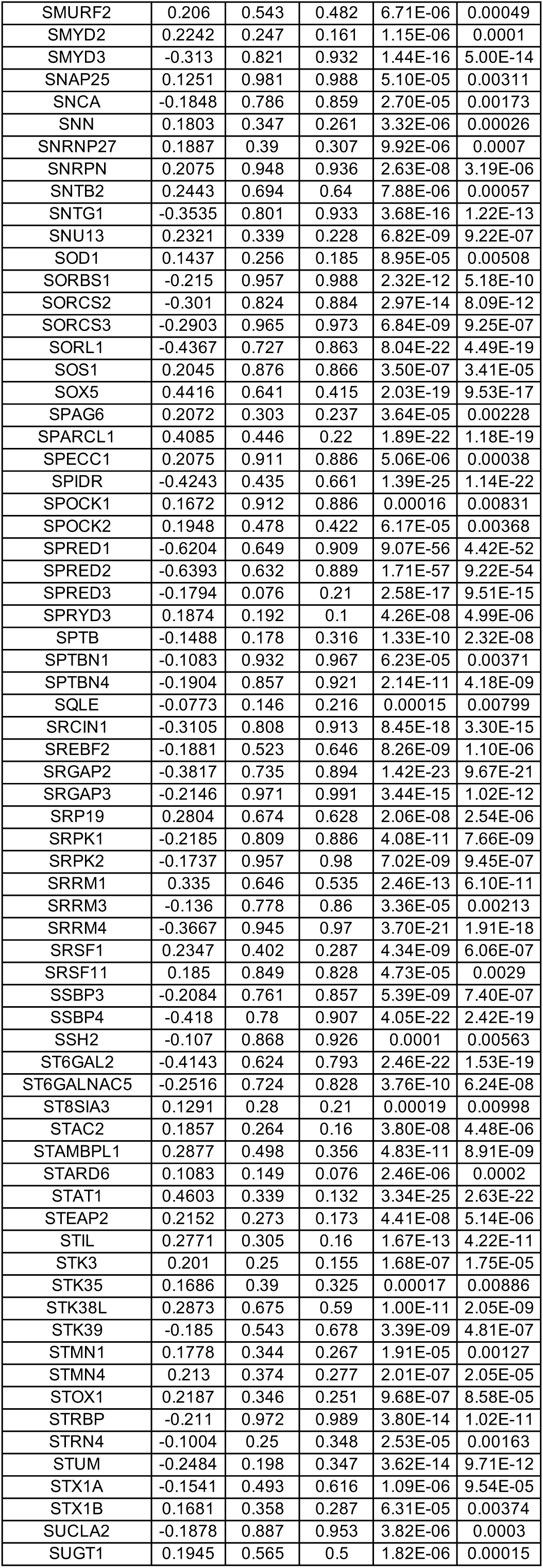
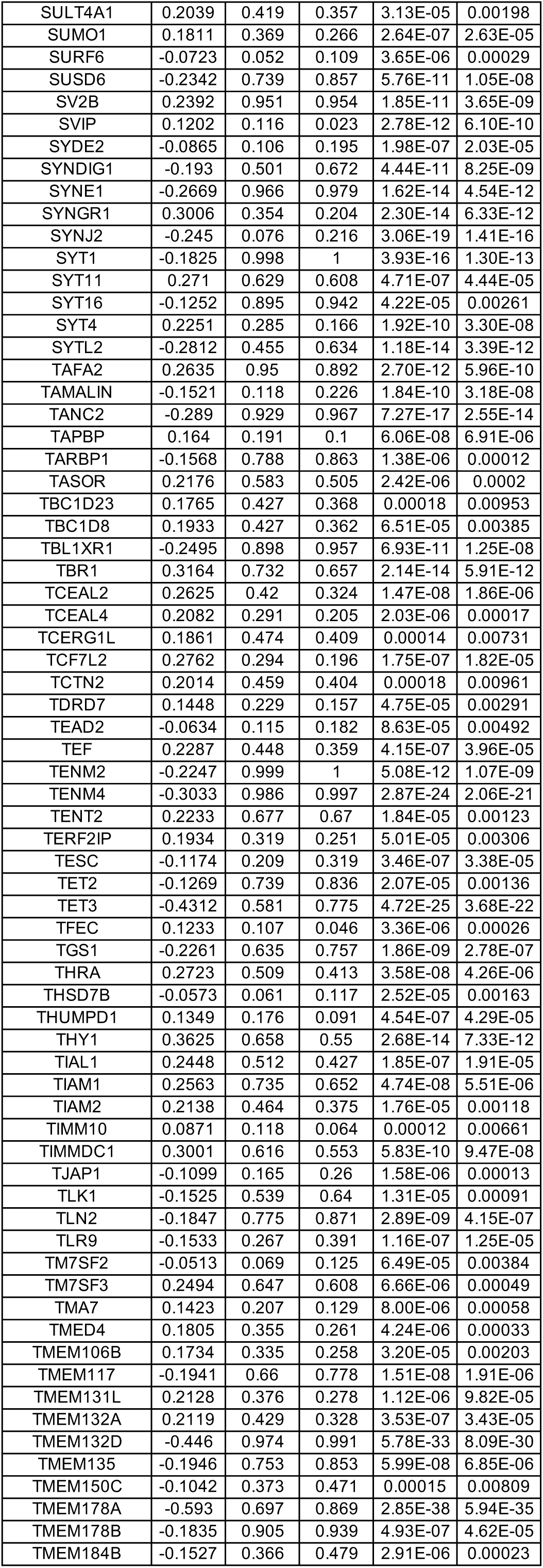
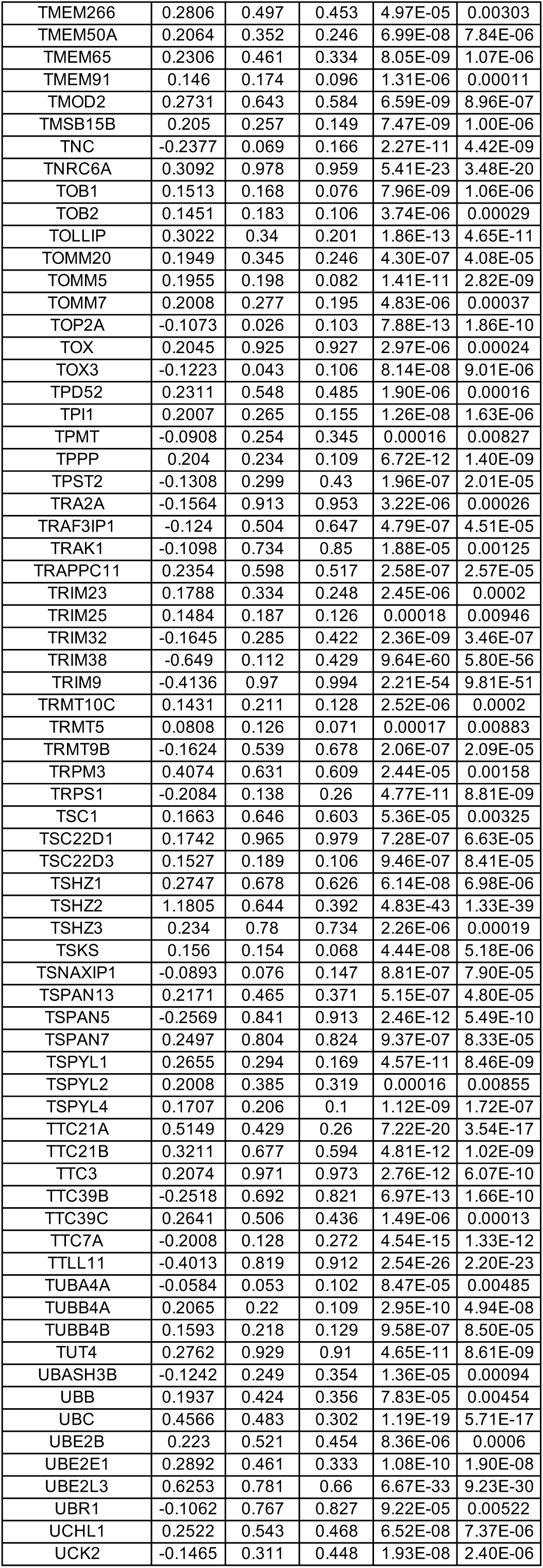
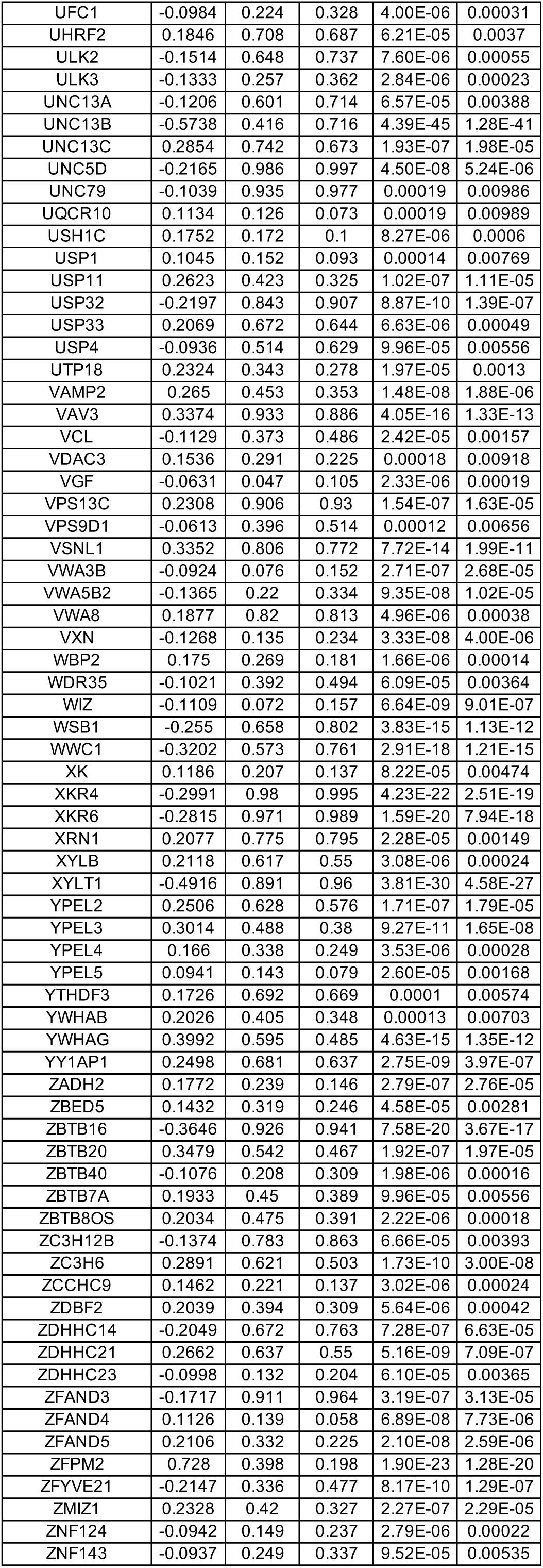
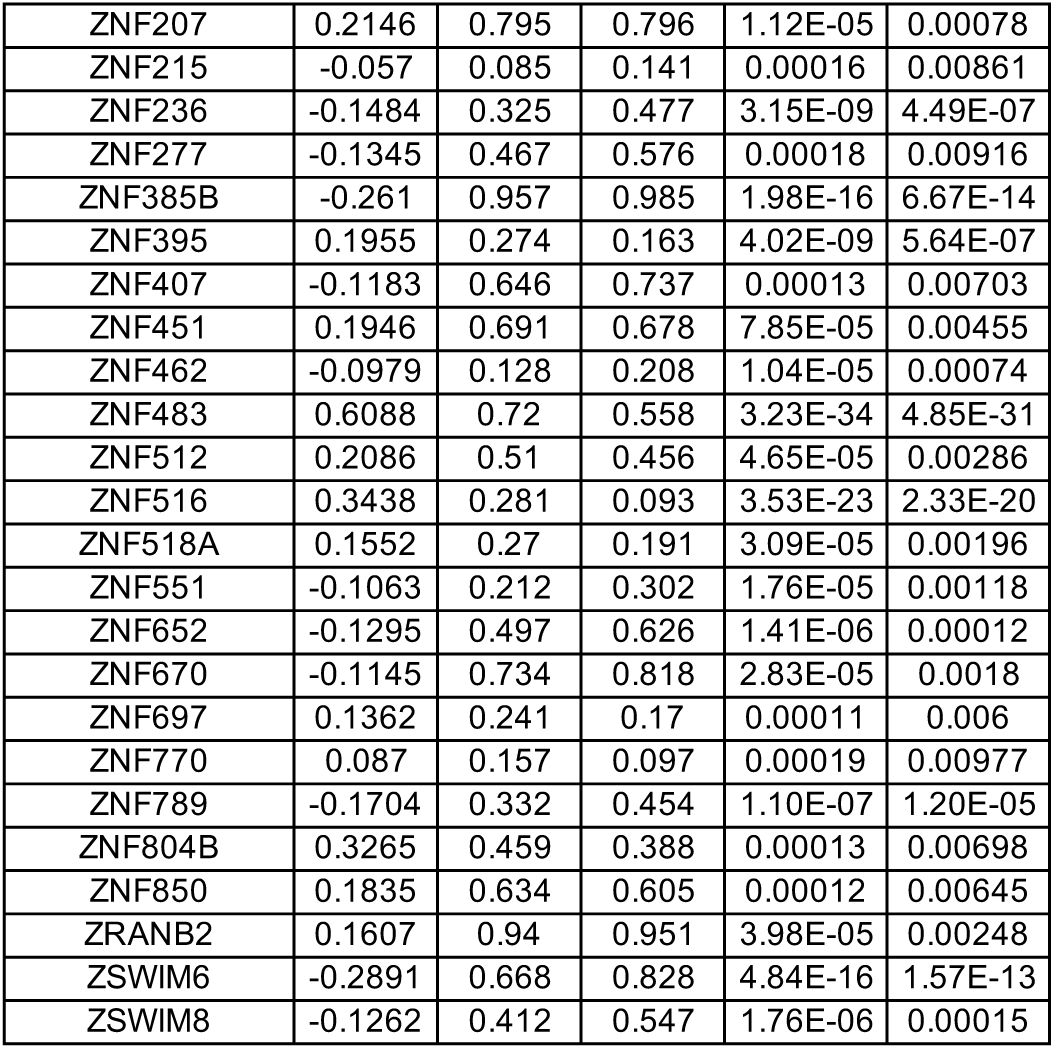
DEGs between wild-type (WT) and MECP2-null (KO) upper layer excitatory neurons (Ex_3) of OC (adjusted p-value < 0.01)

**Table S24.**
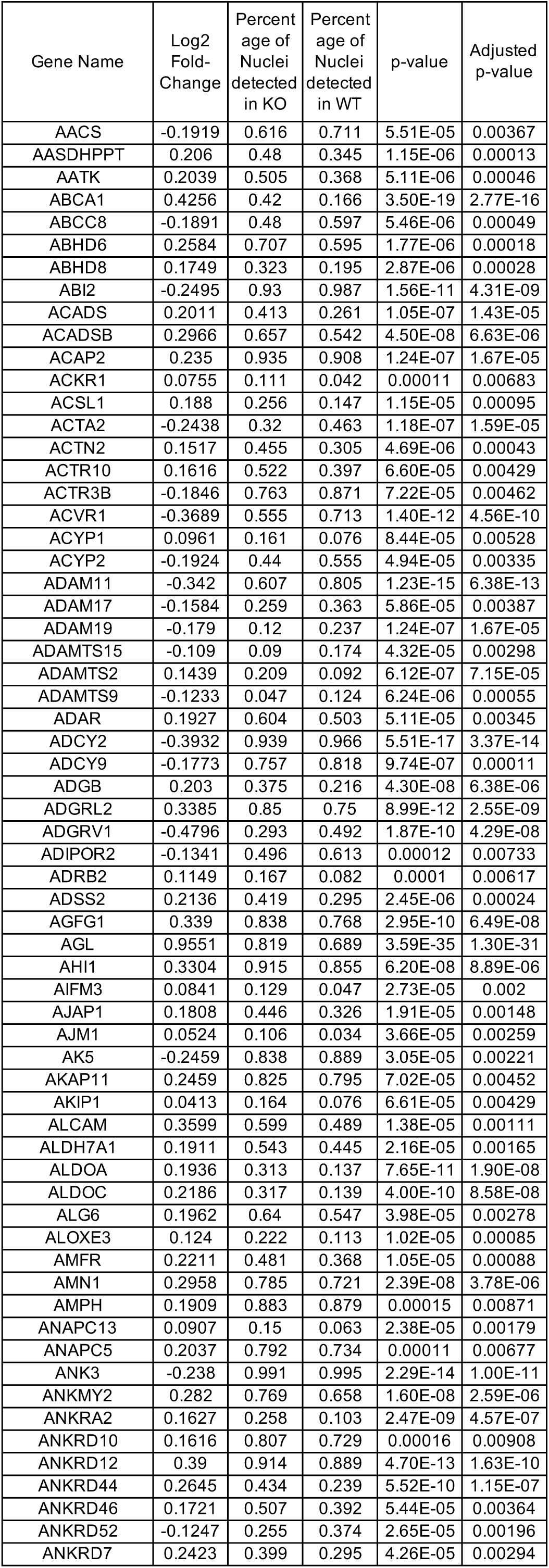
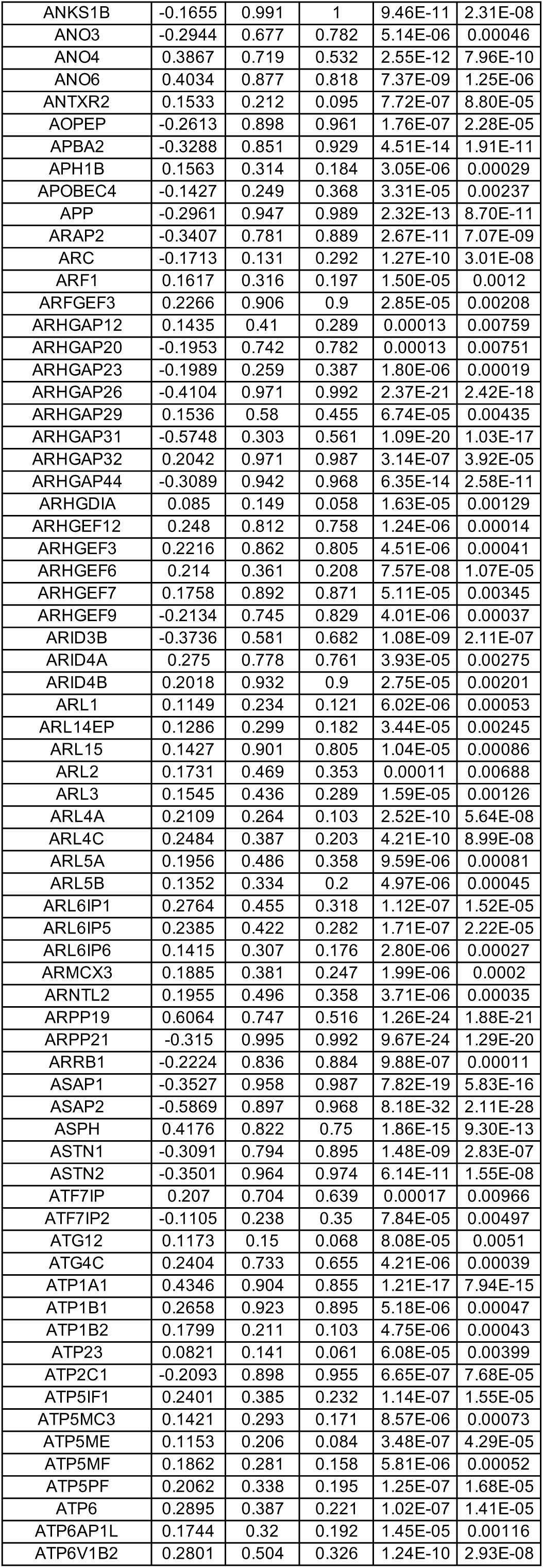
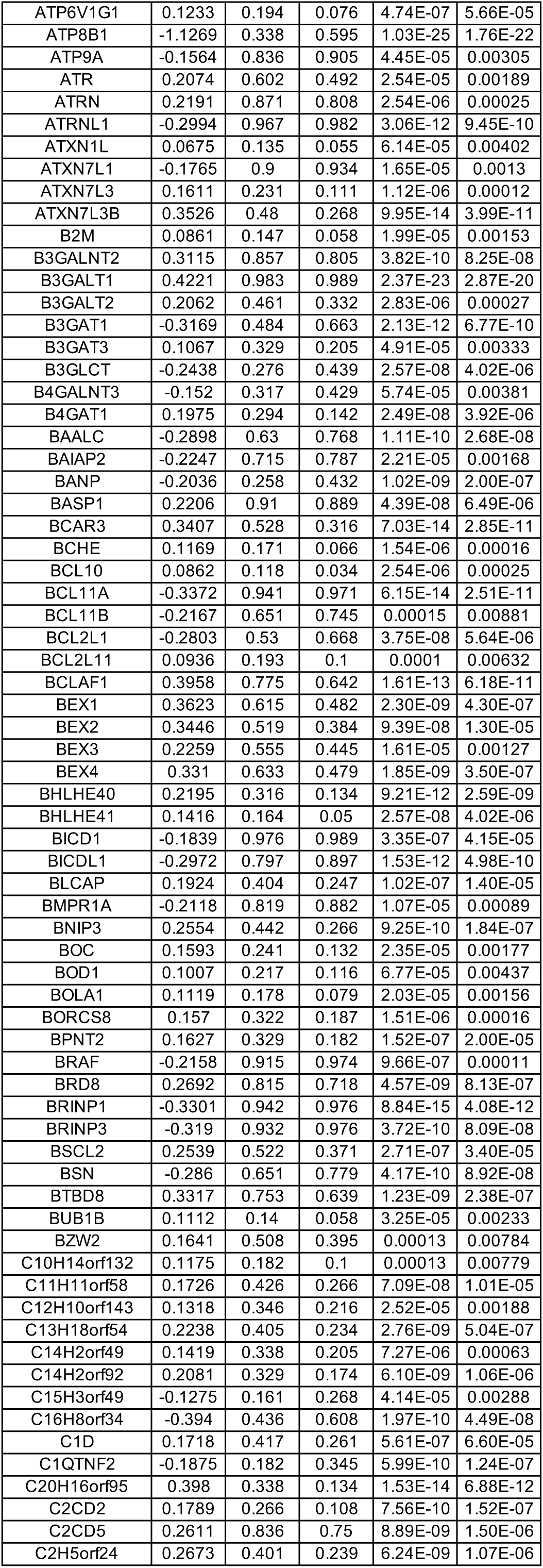
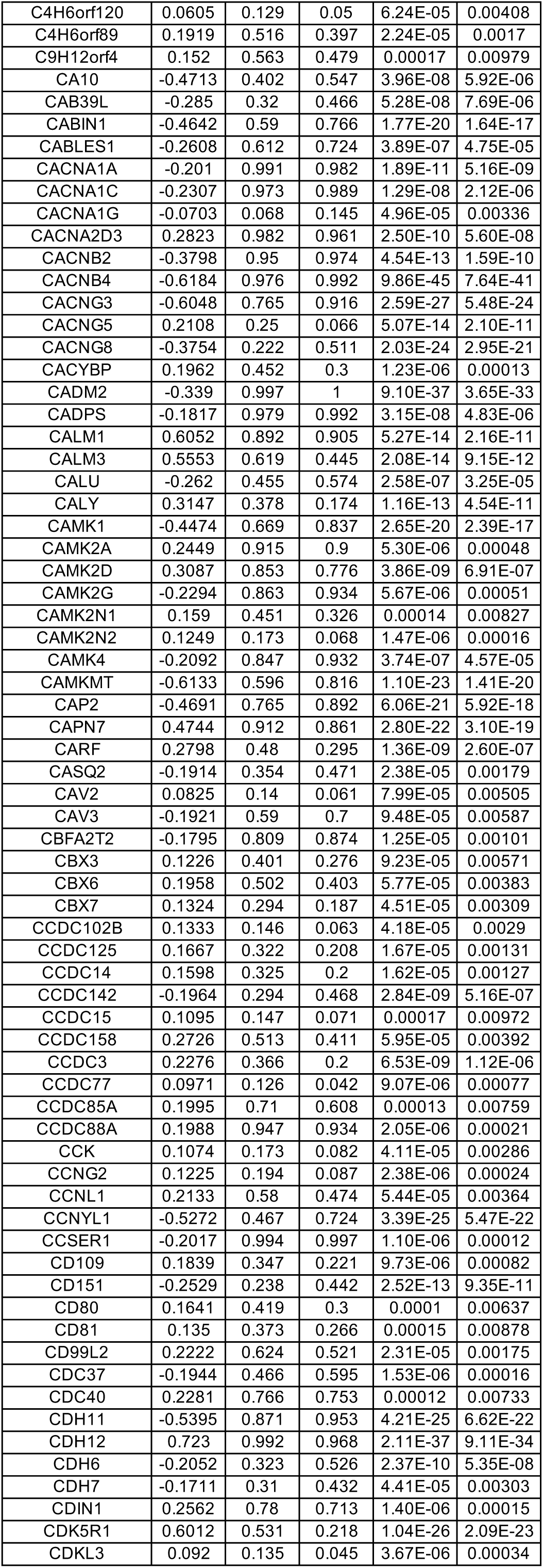
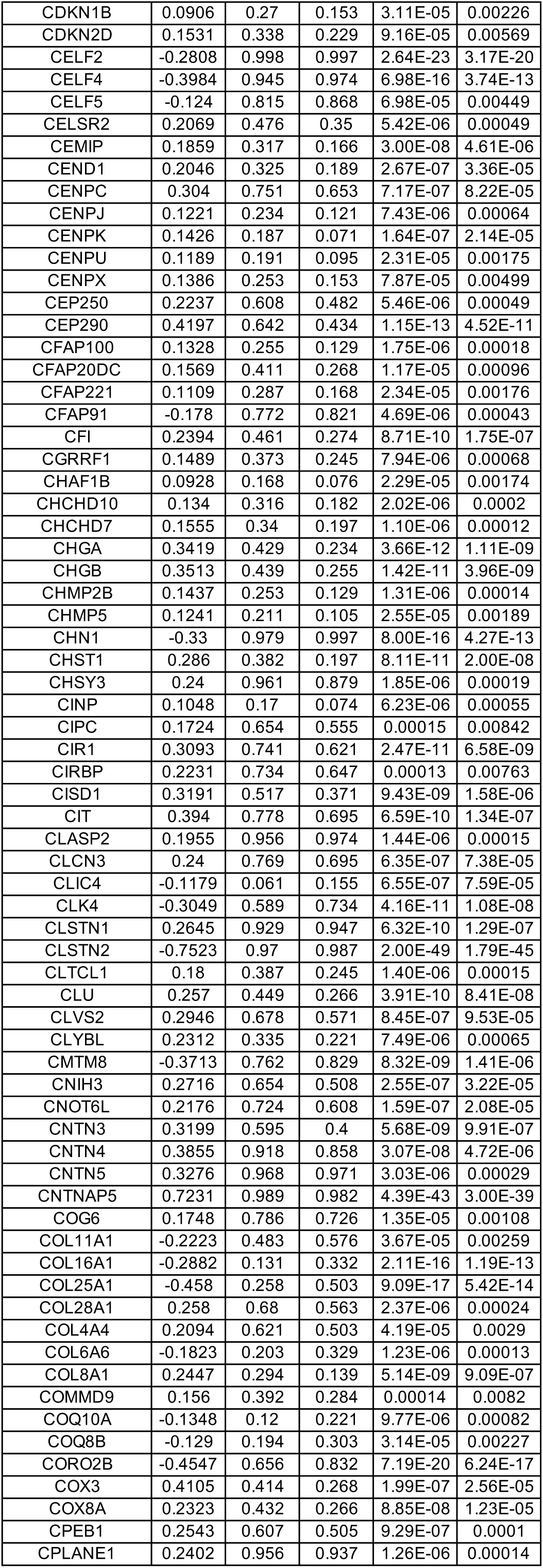
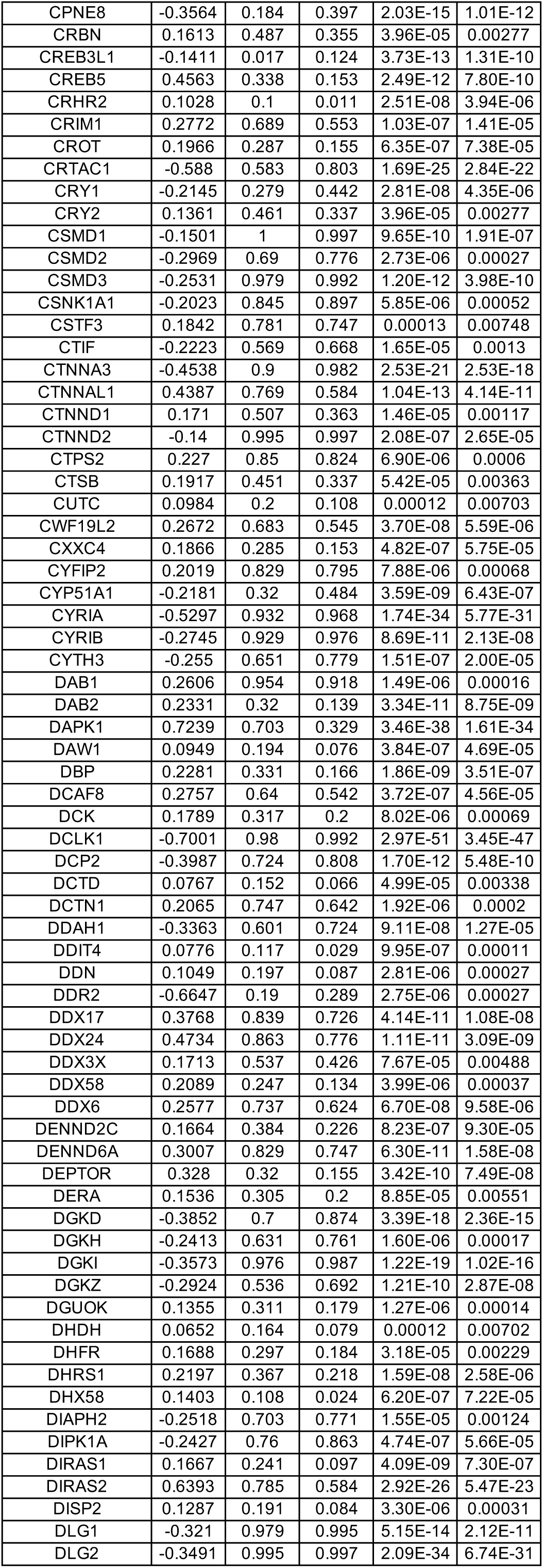
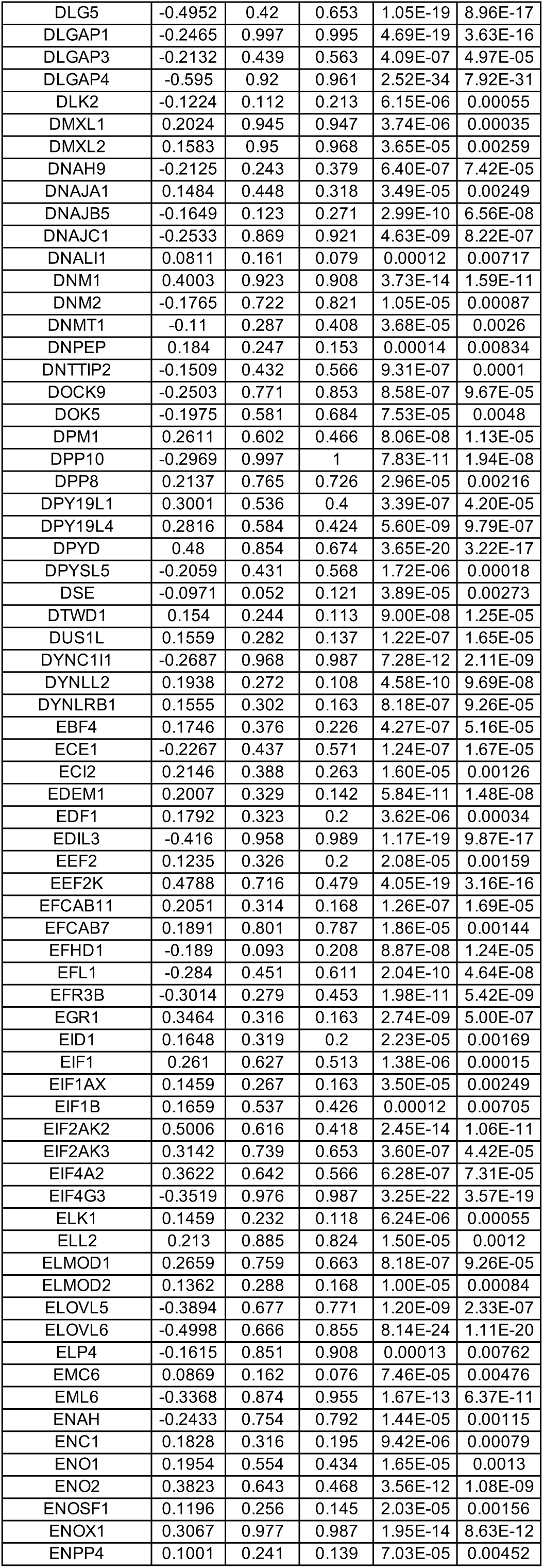
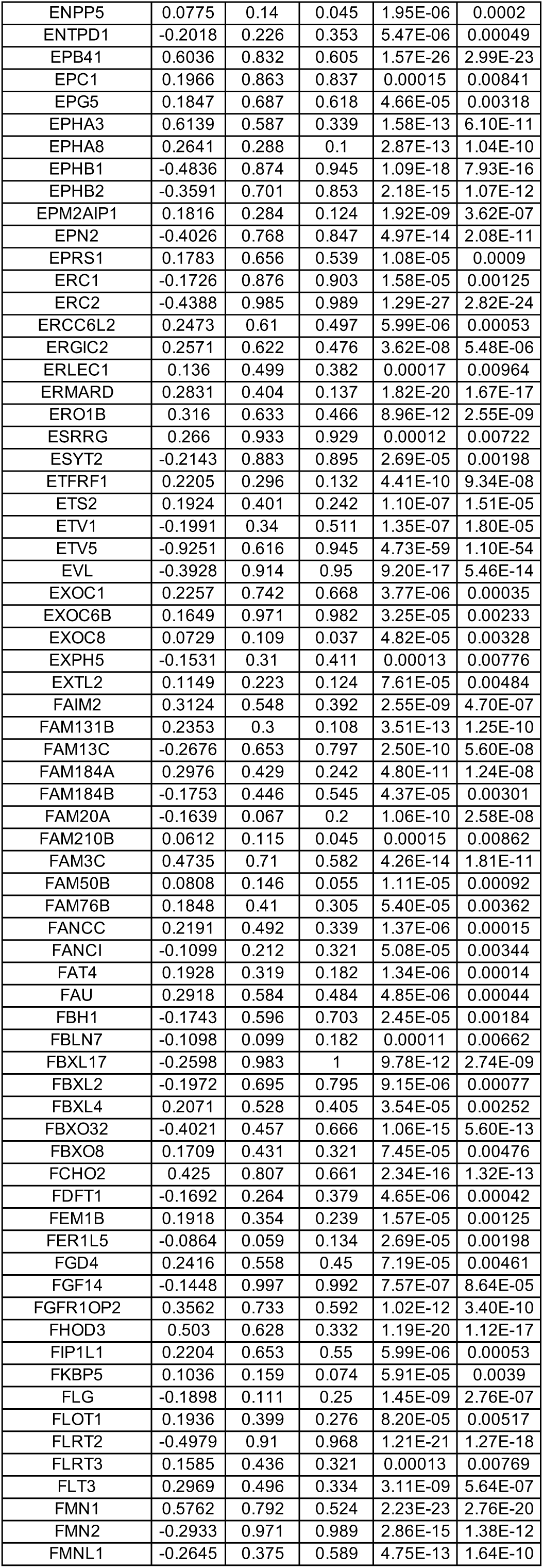
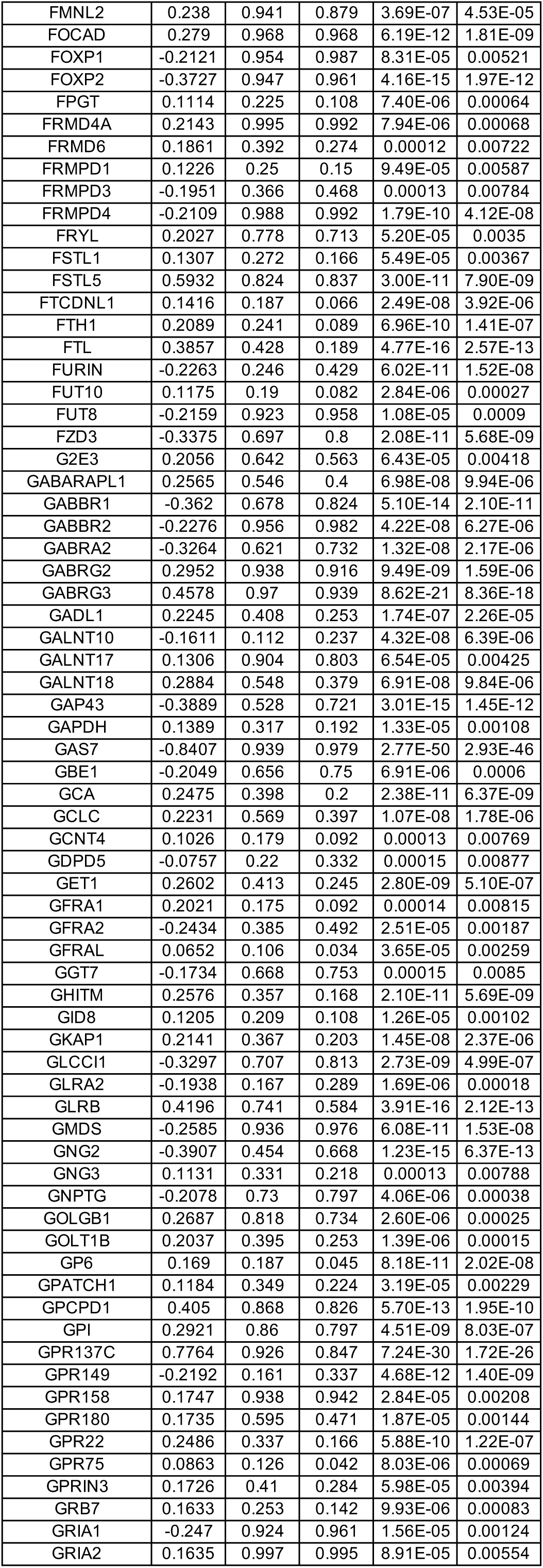
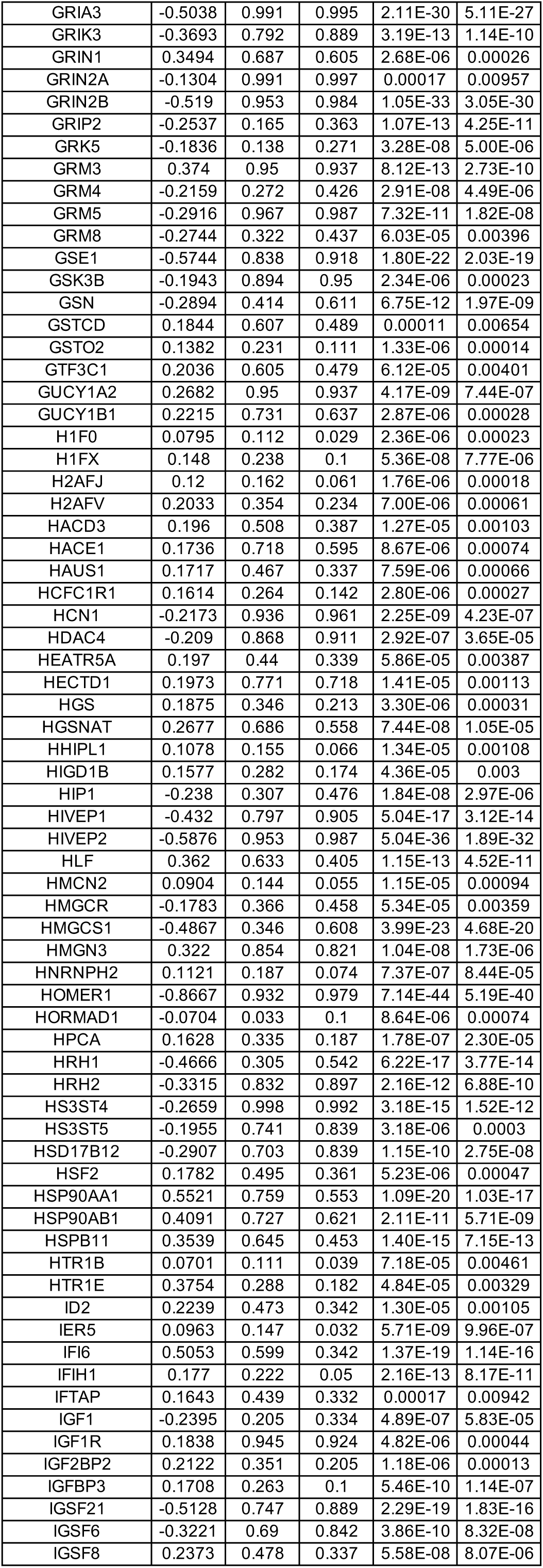
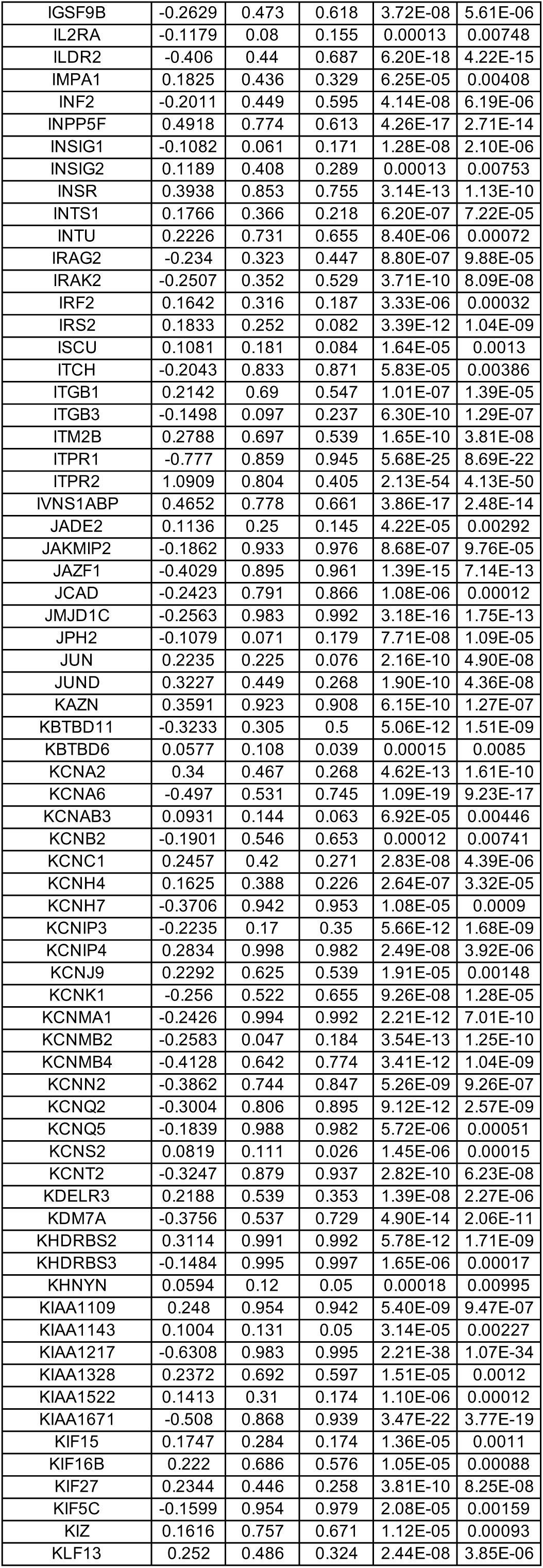
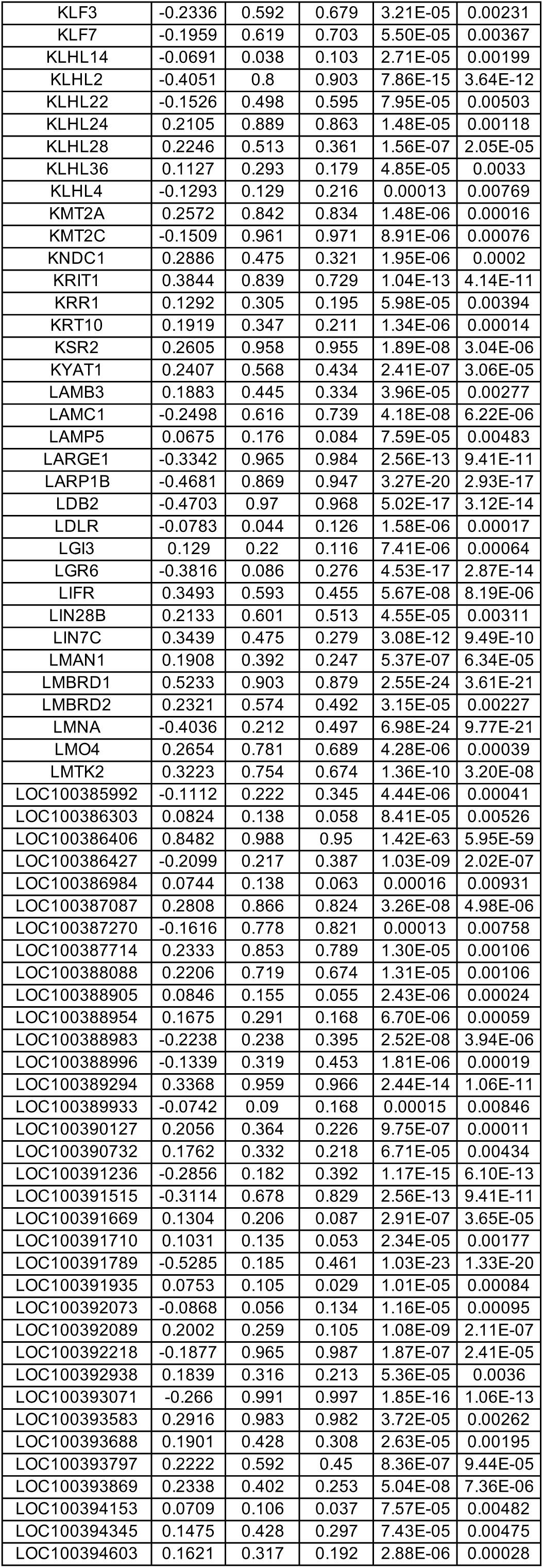
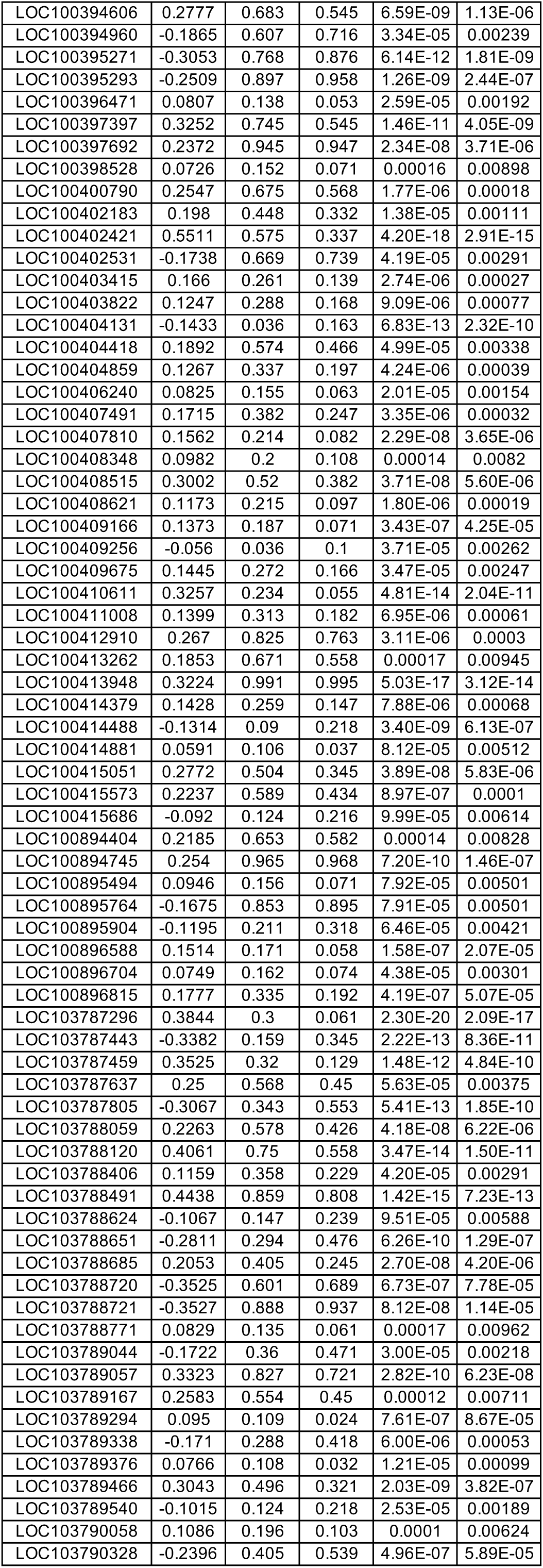
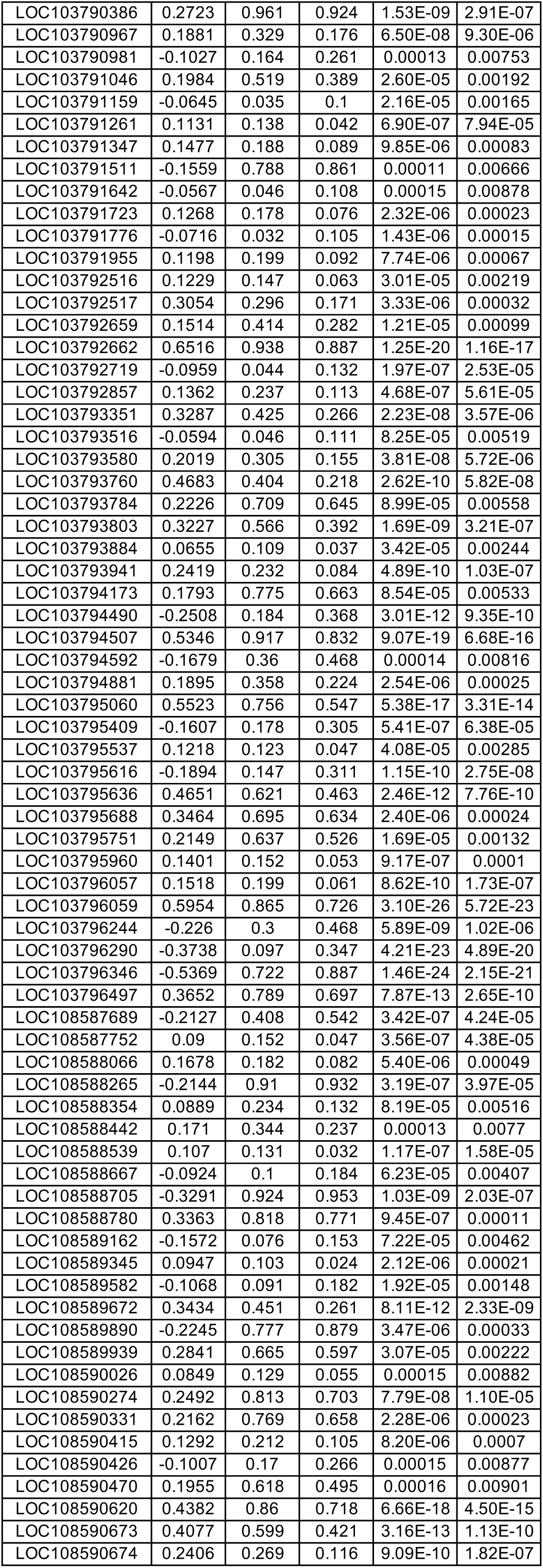
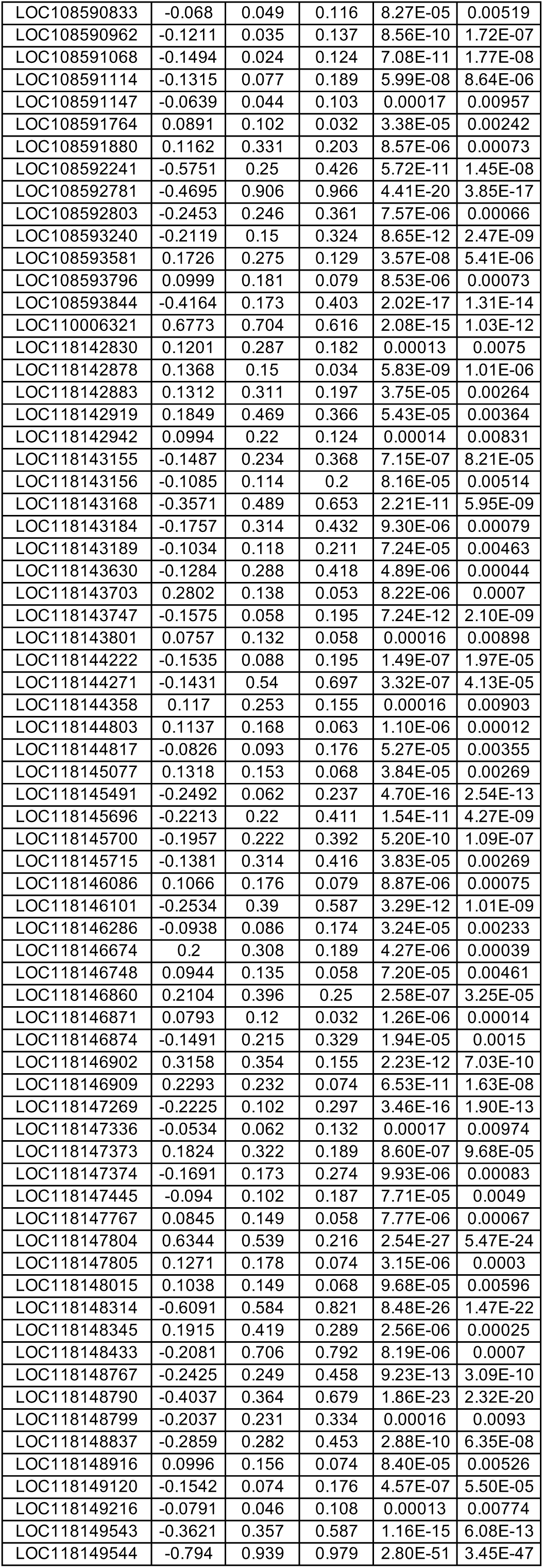
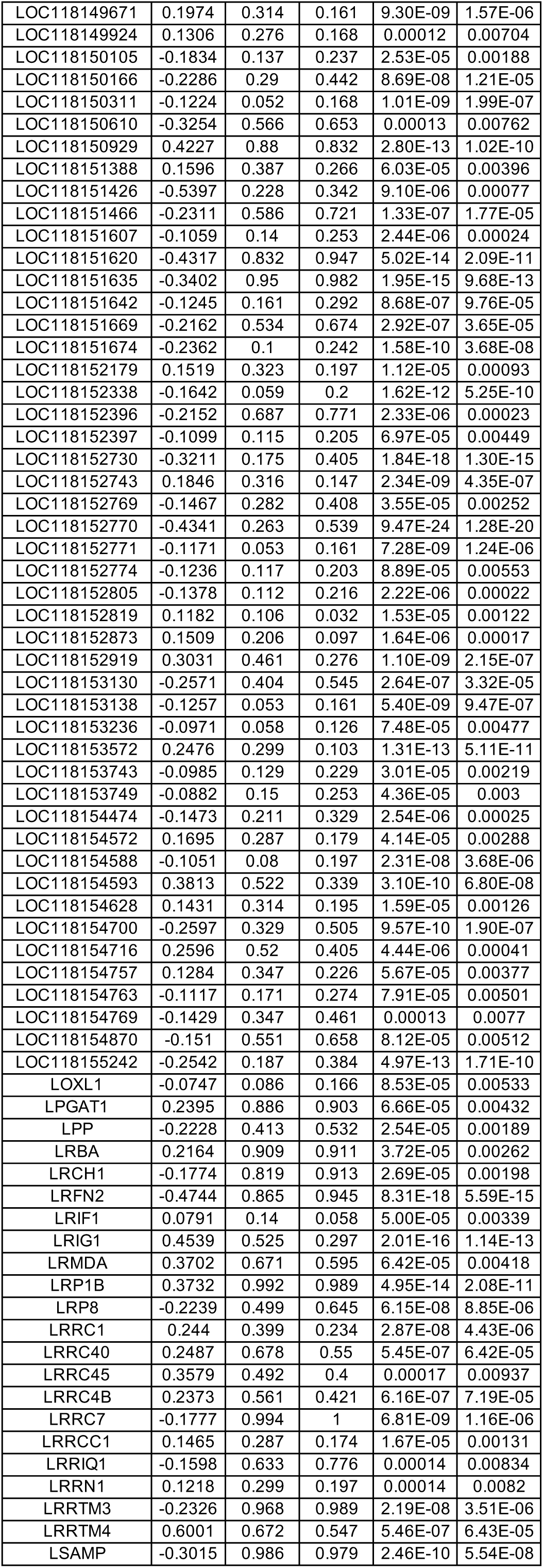
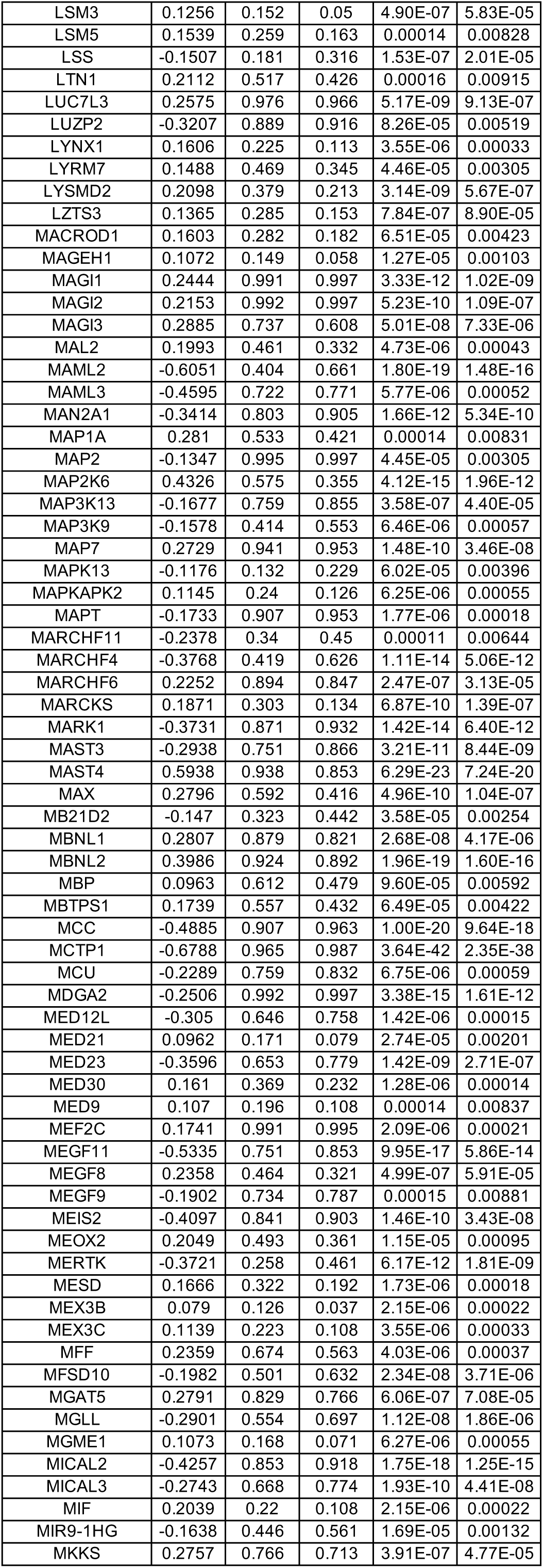
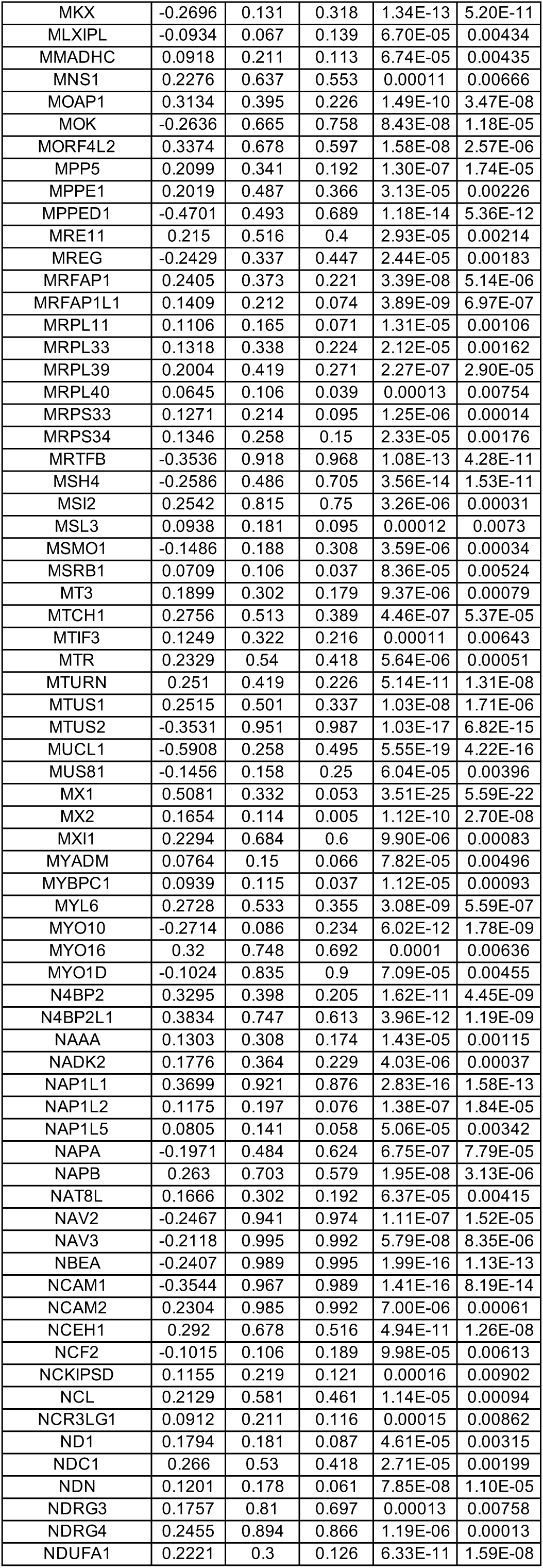
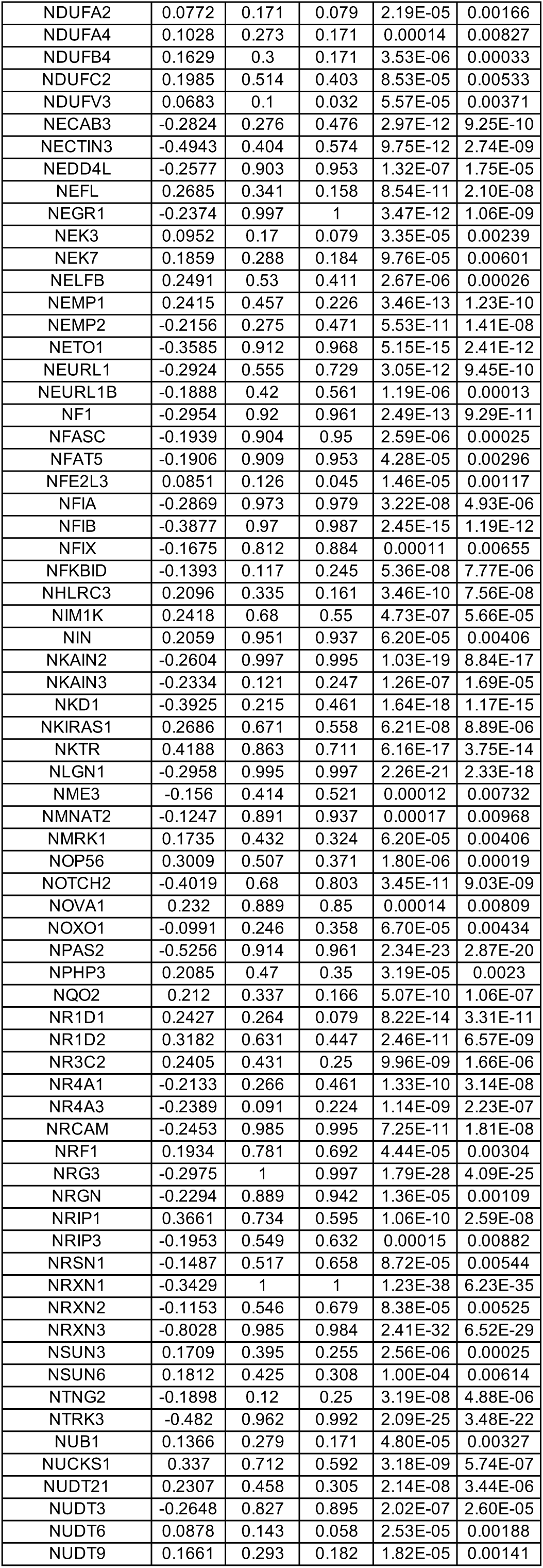
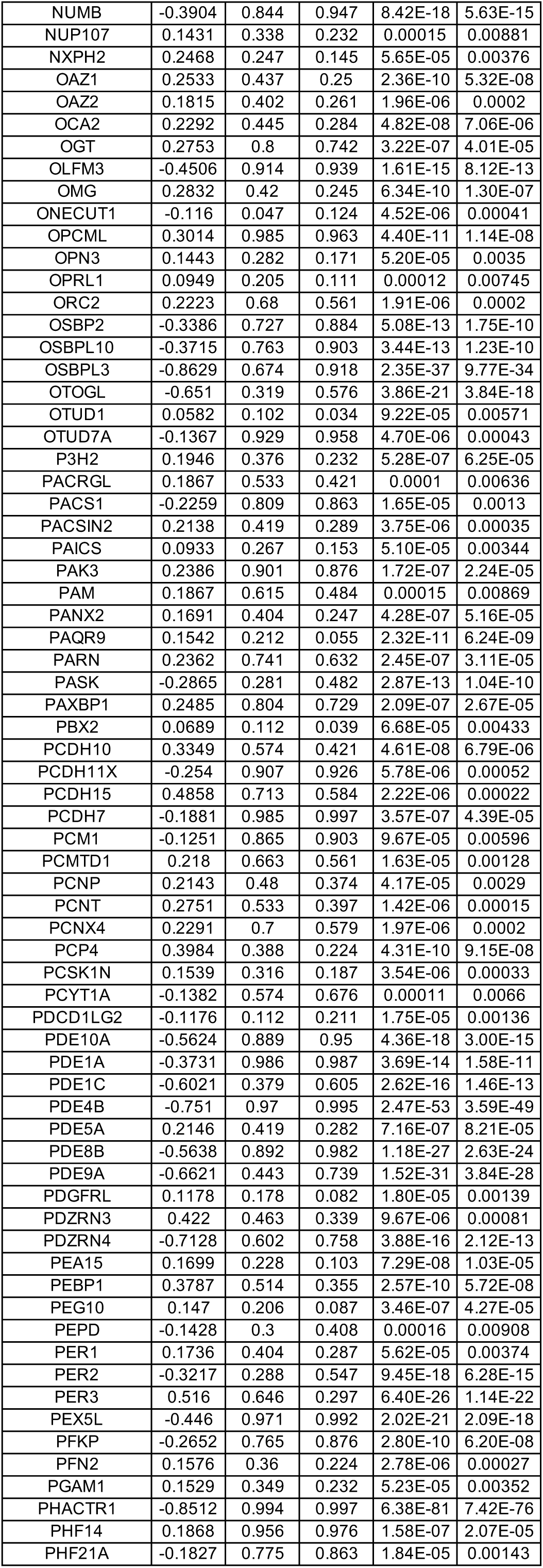
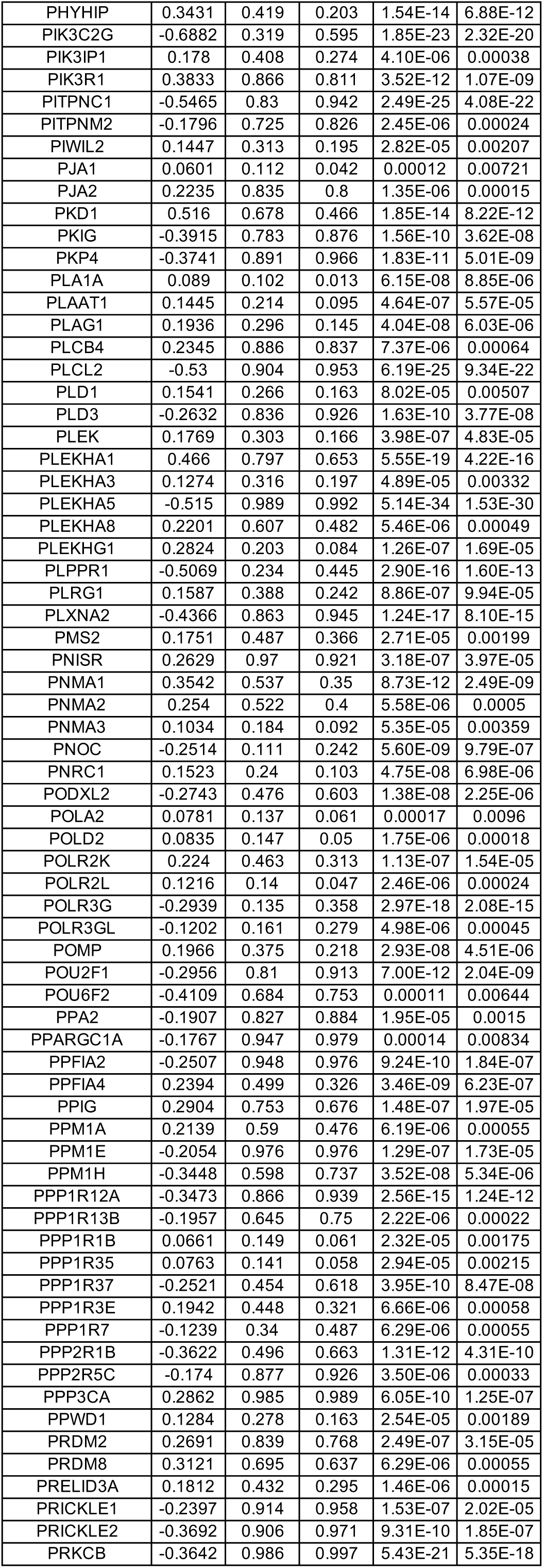
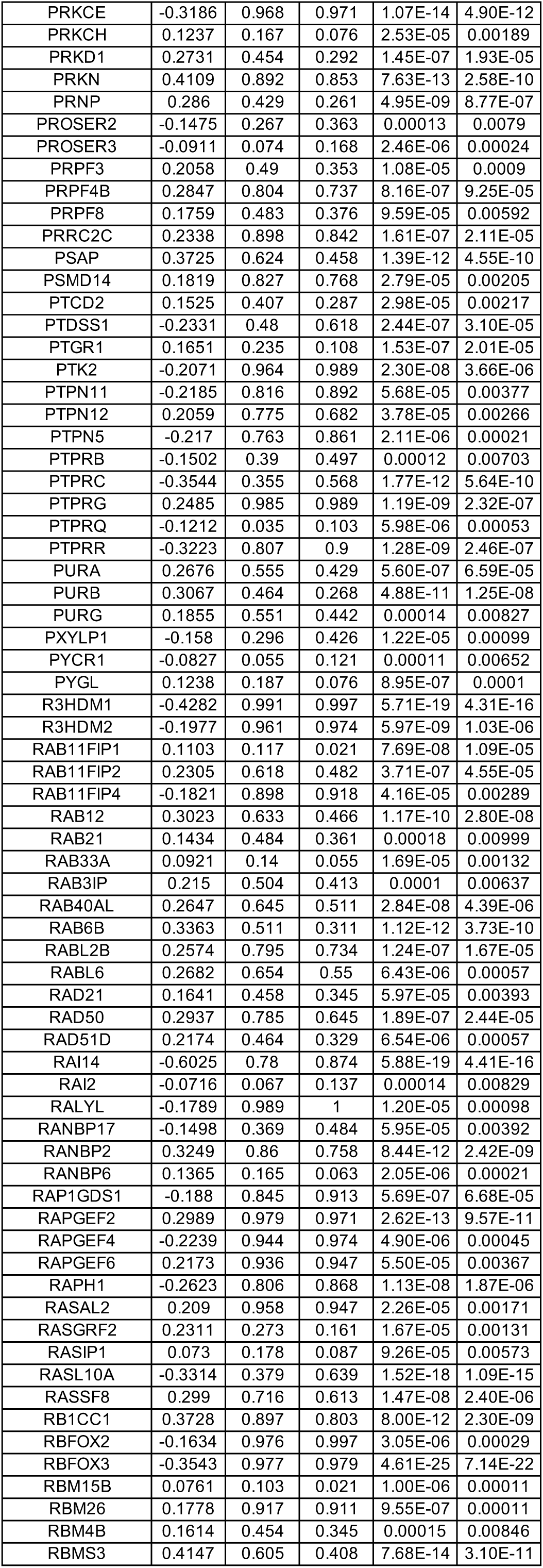
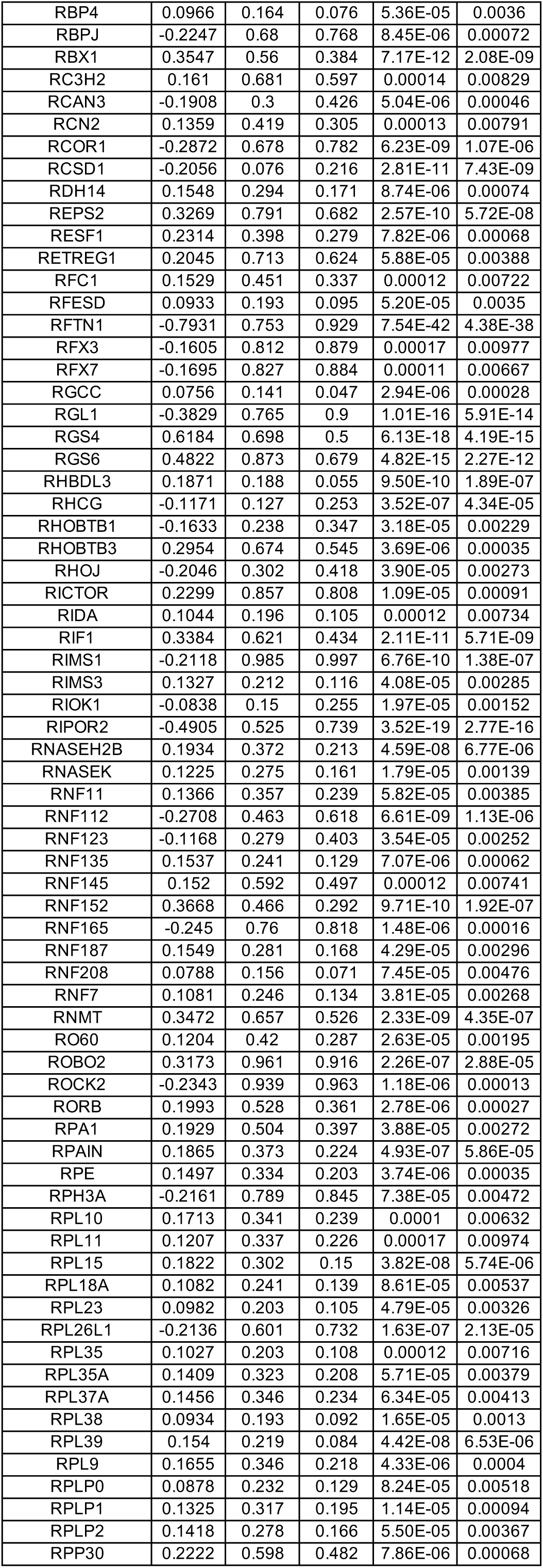
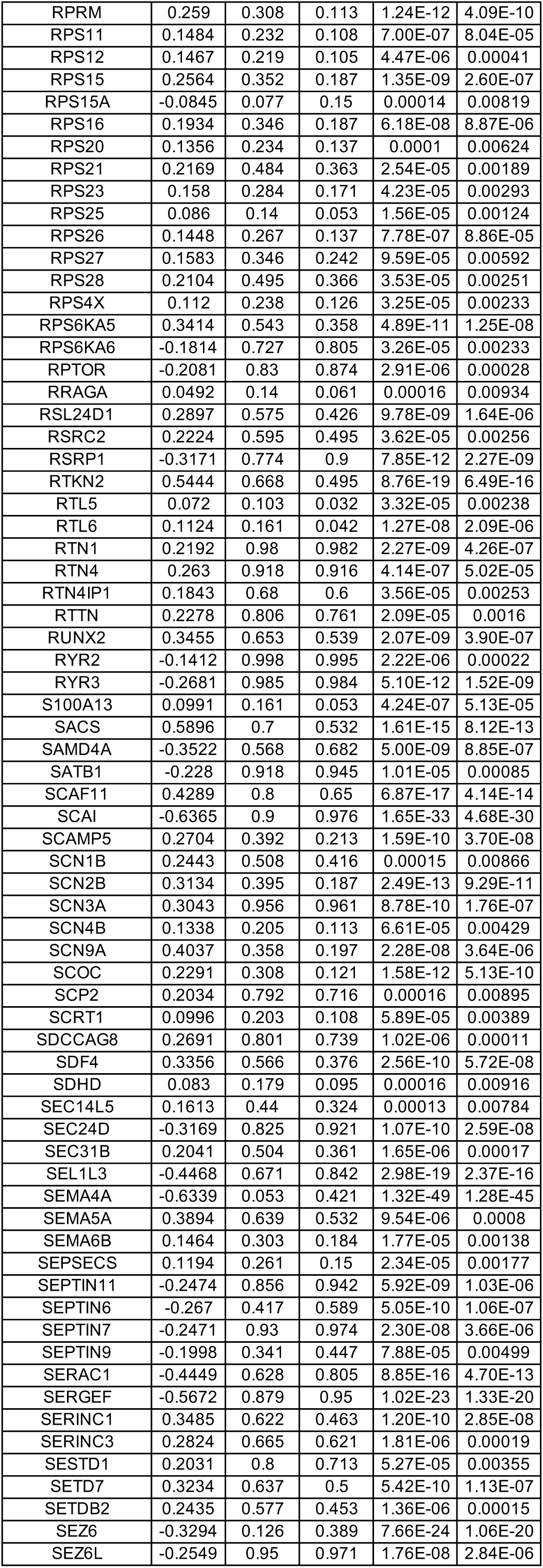
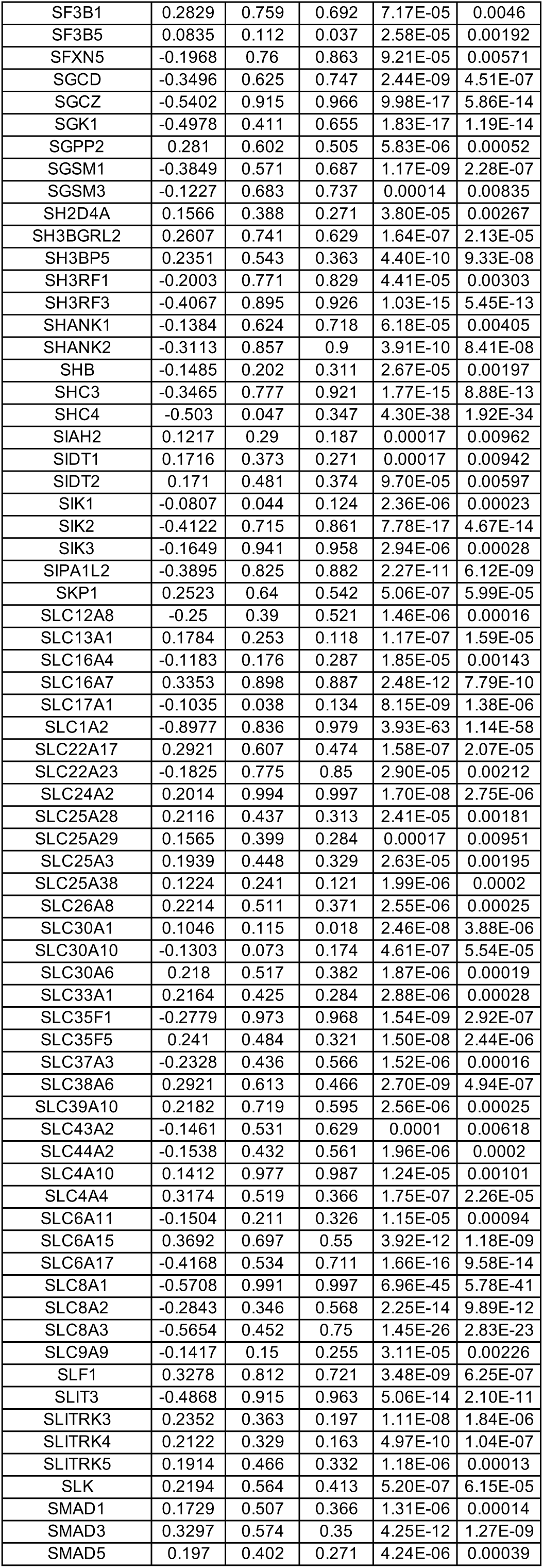
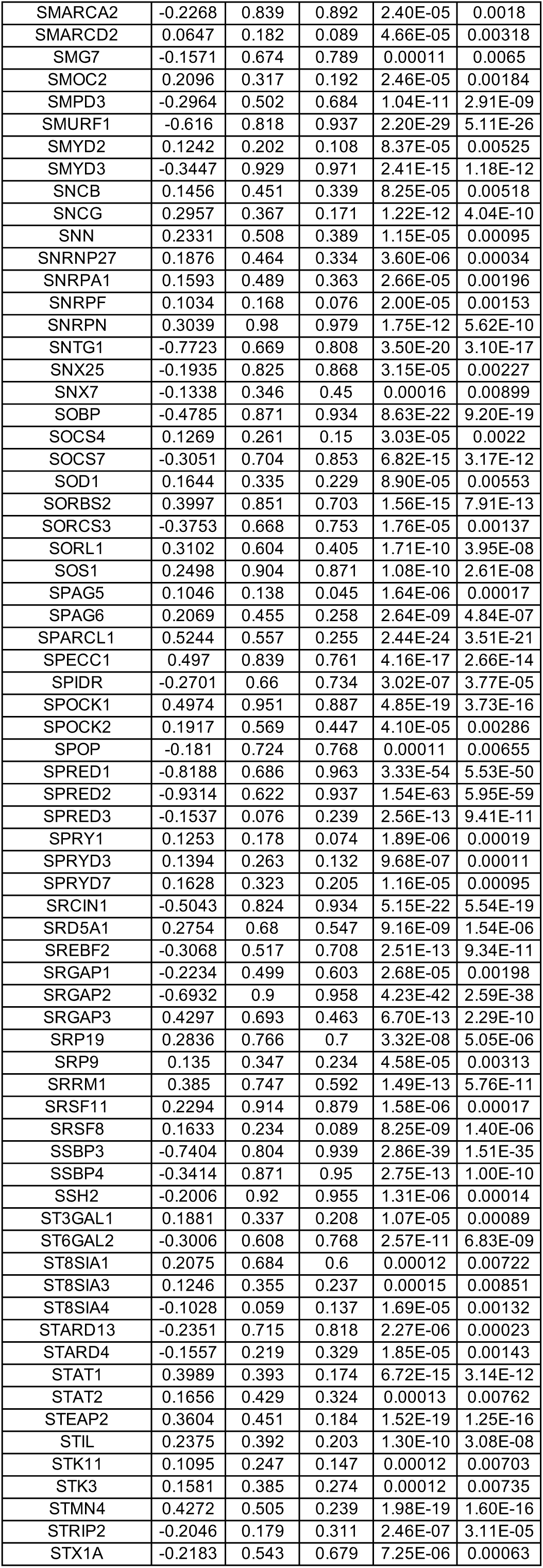
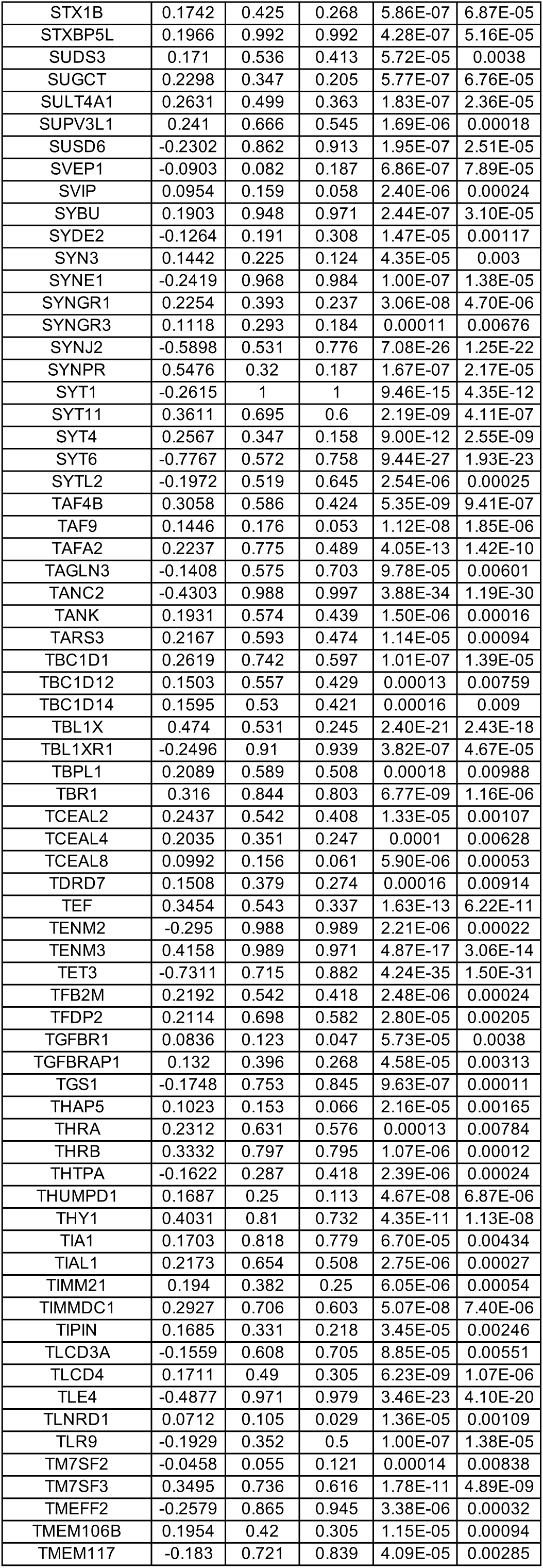
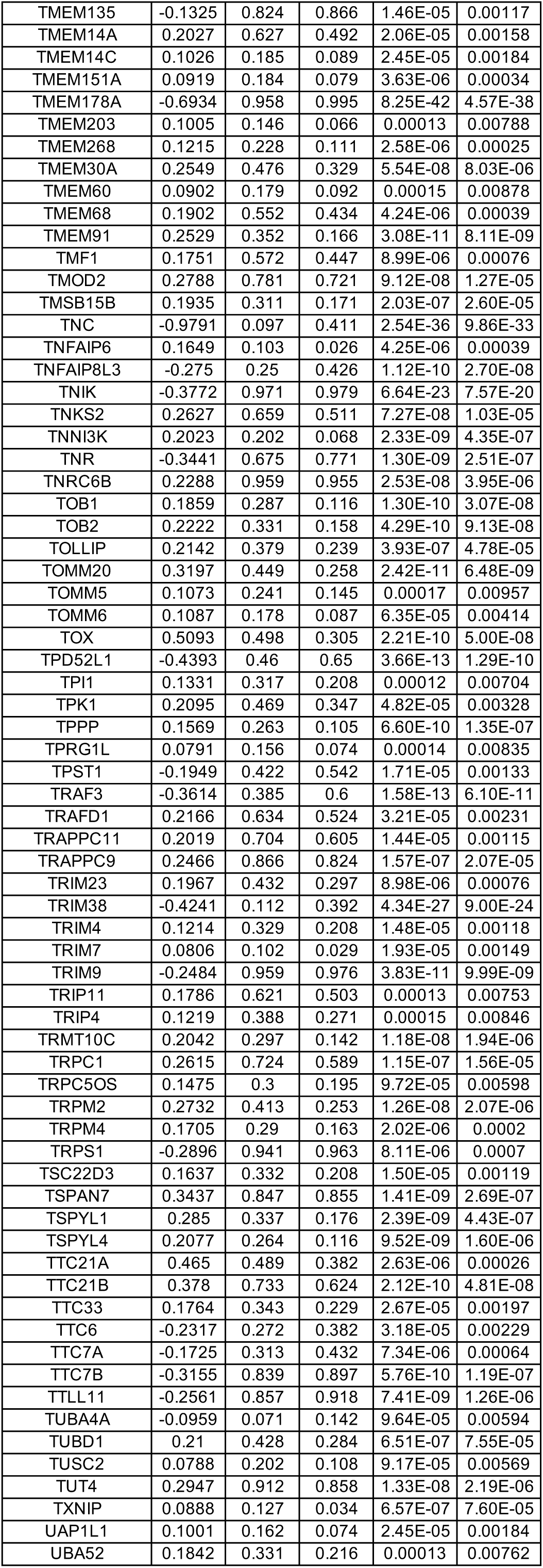
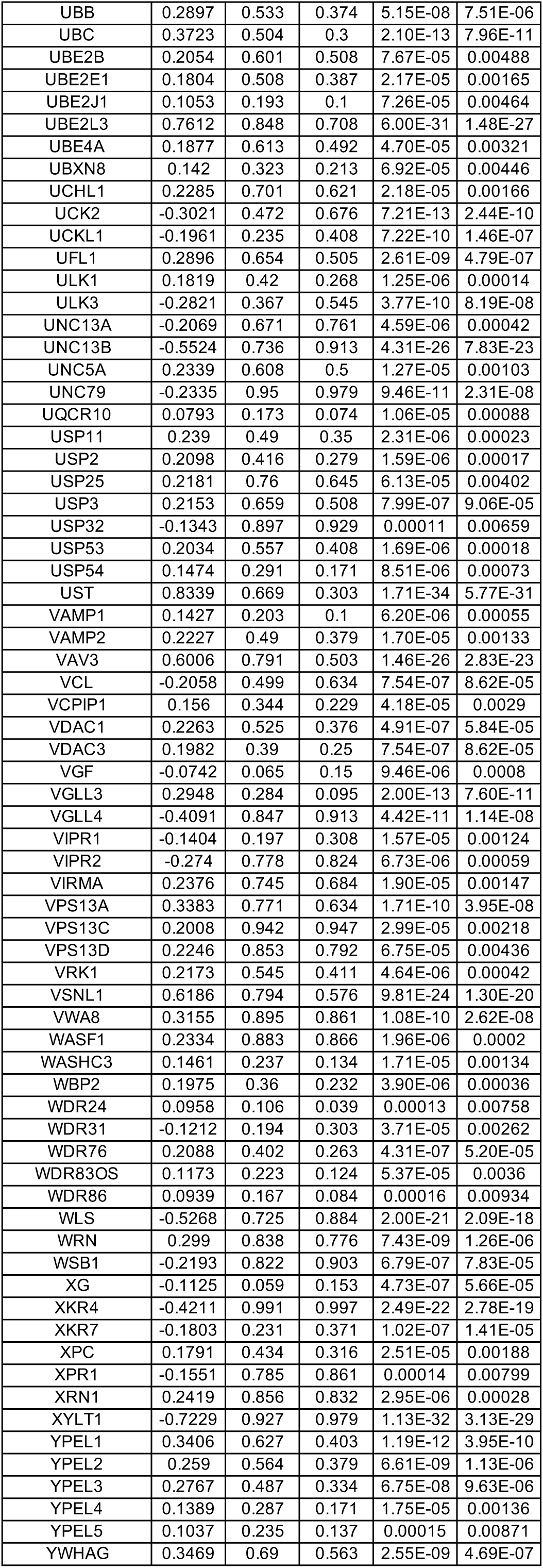
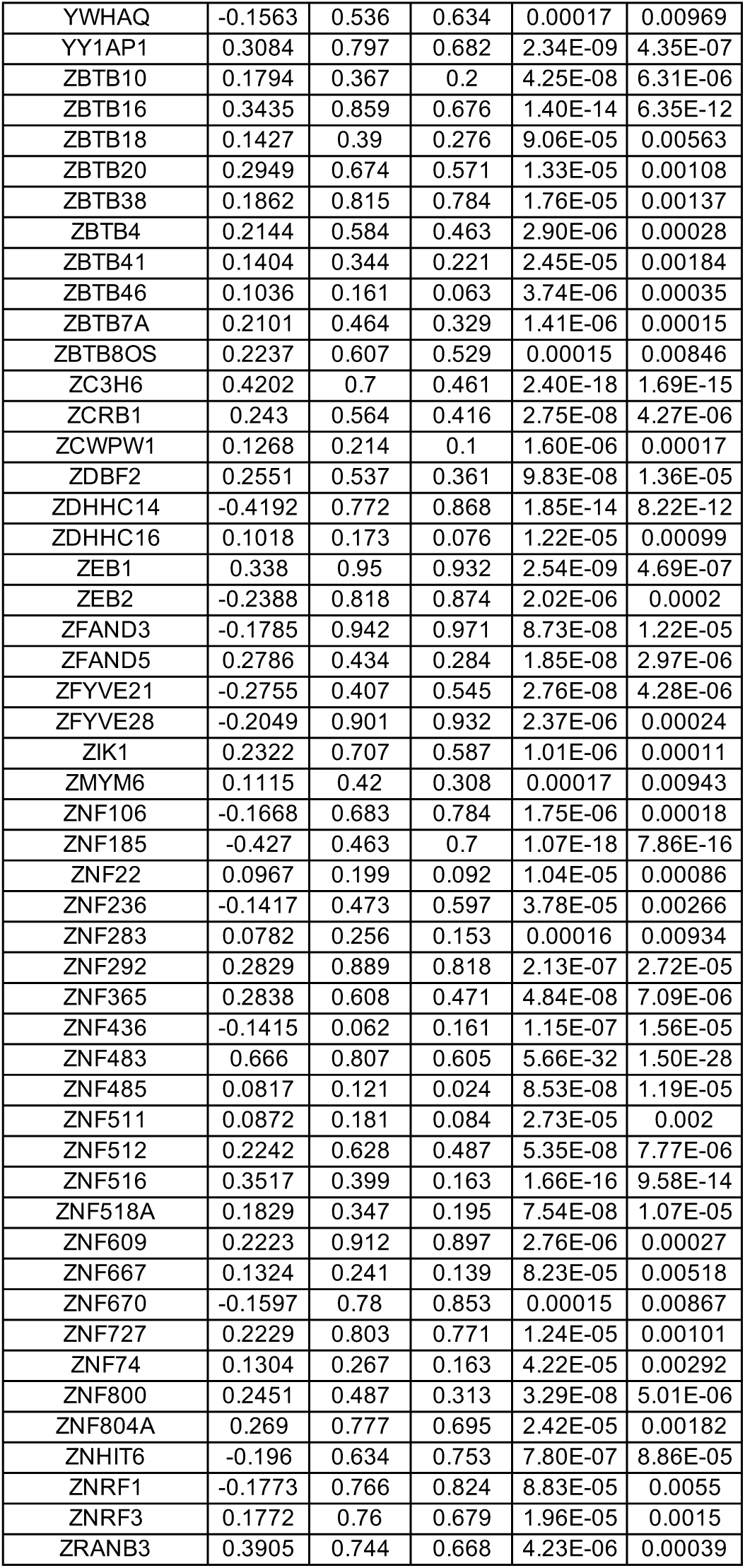
DEGs between wild-type (WT) and MECP2-null (KO) upper layer excitatory neurons (Ex_4) of OC (adjusted p-value < 0.01)

**Table S25.**
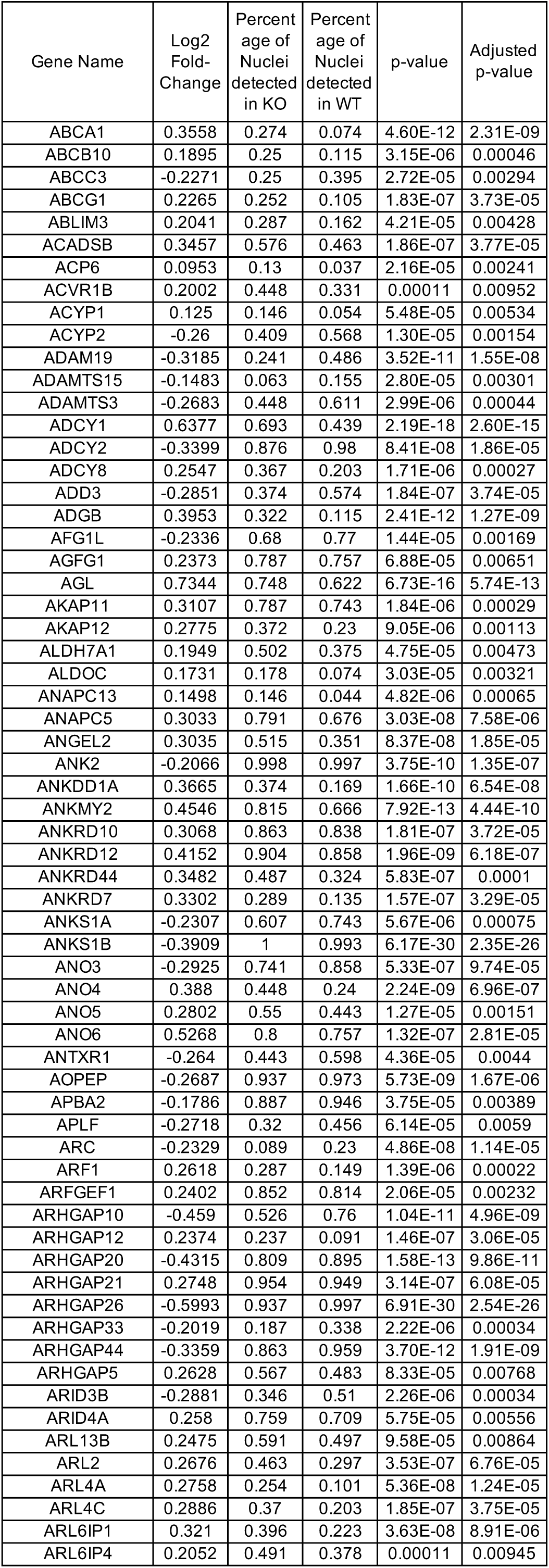
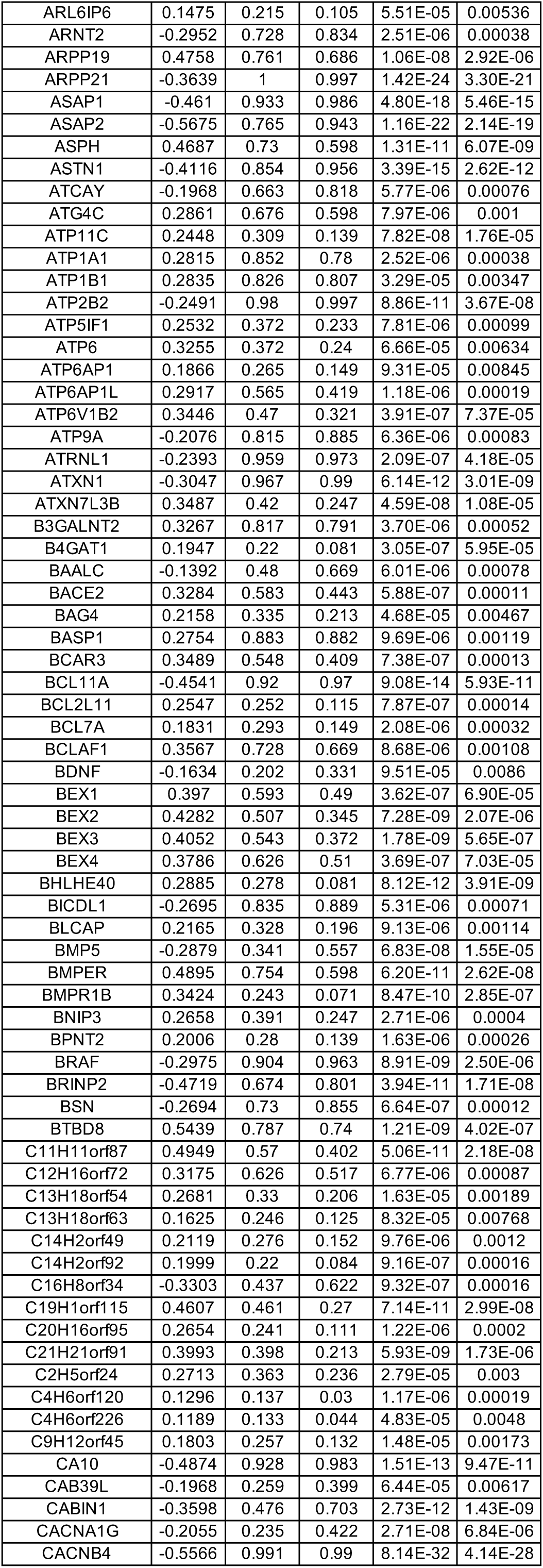
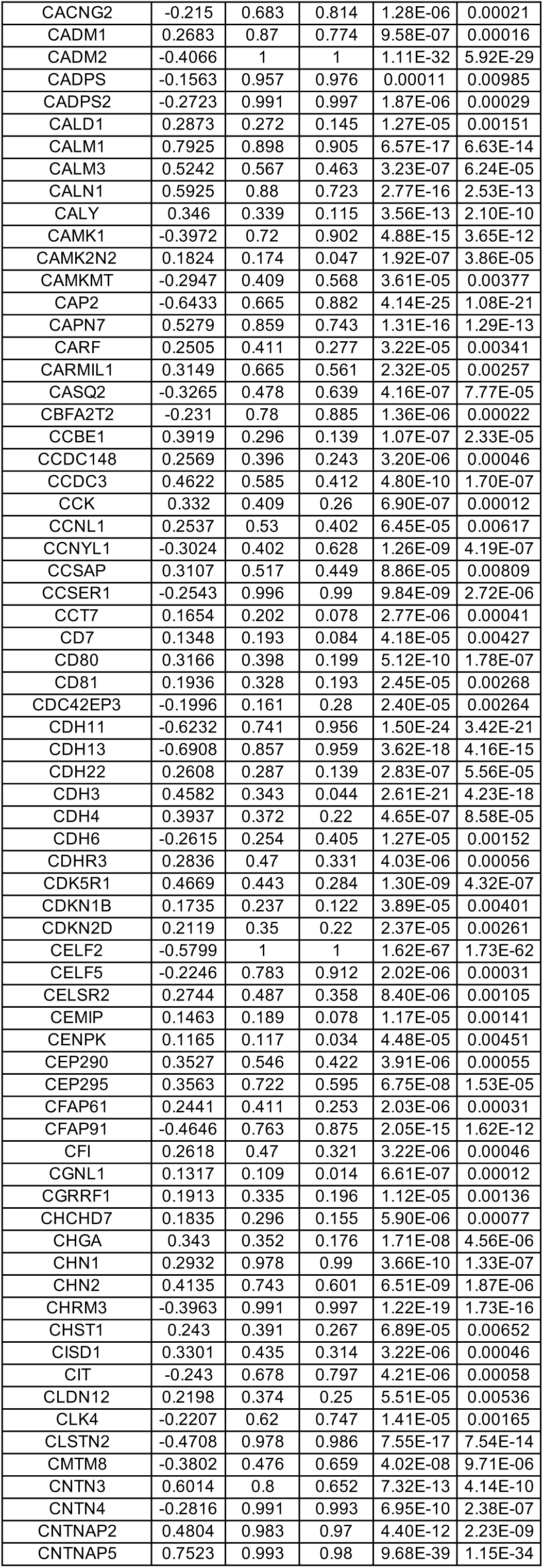
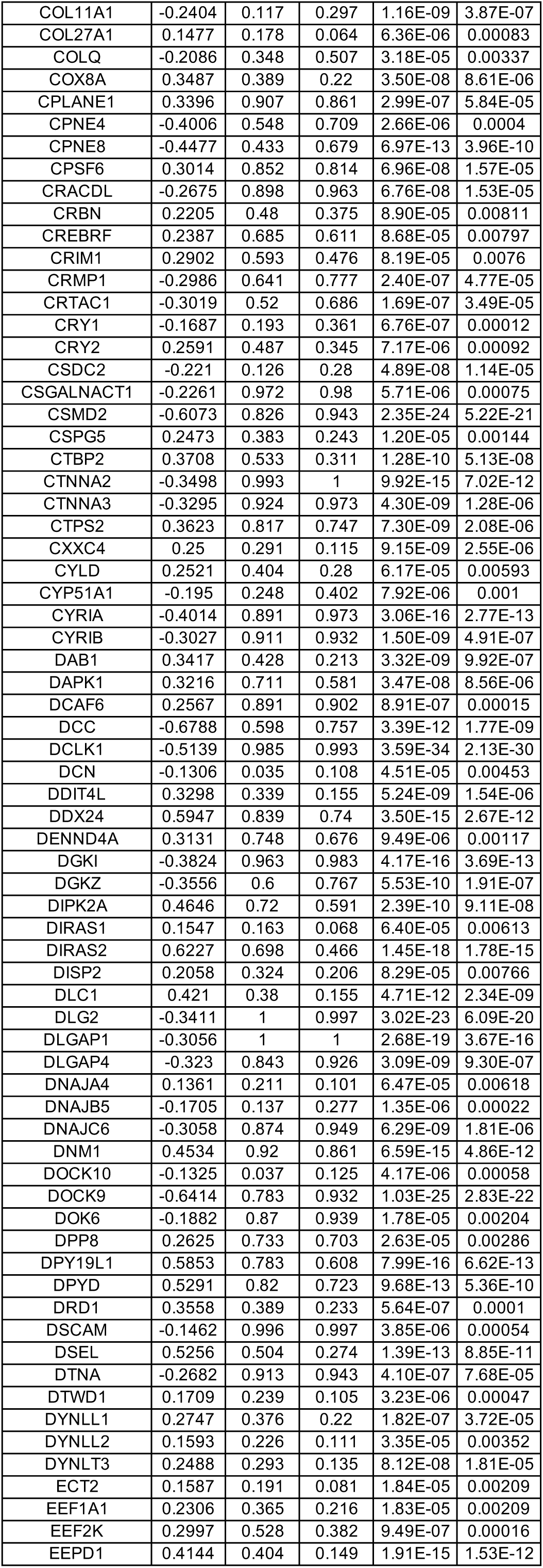
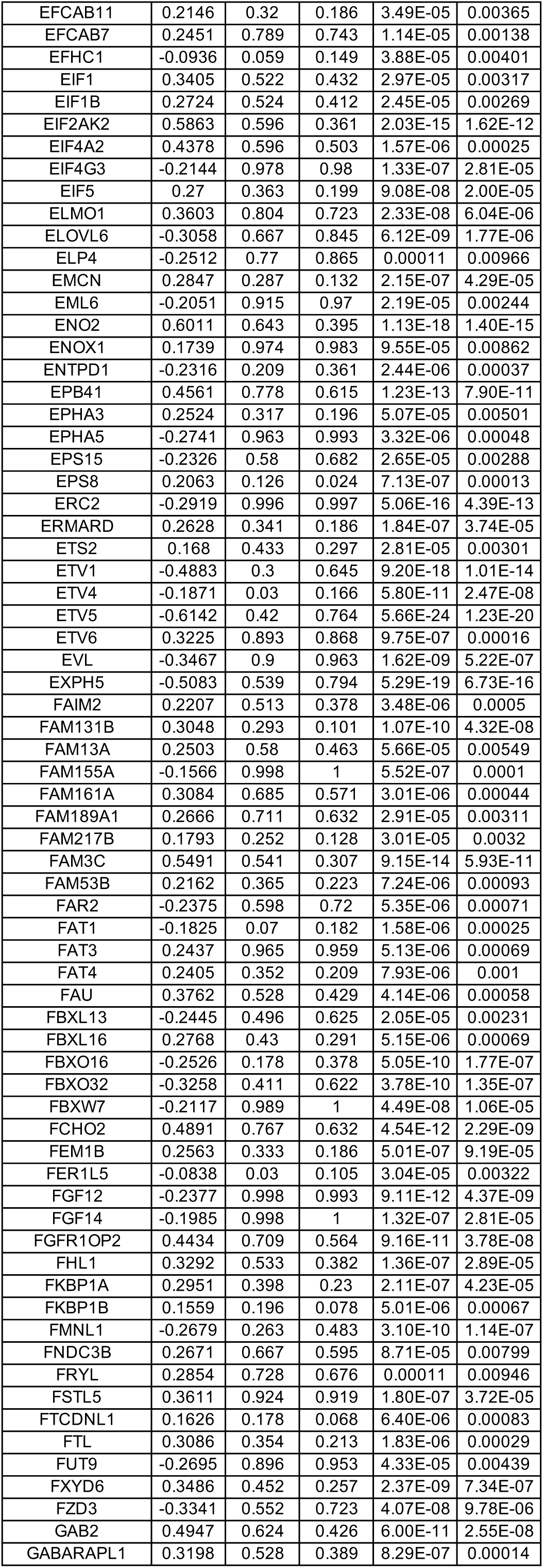
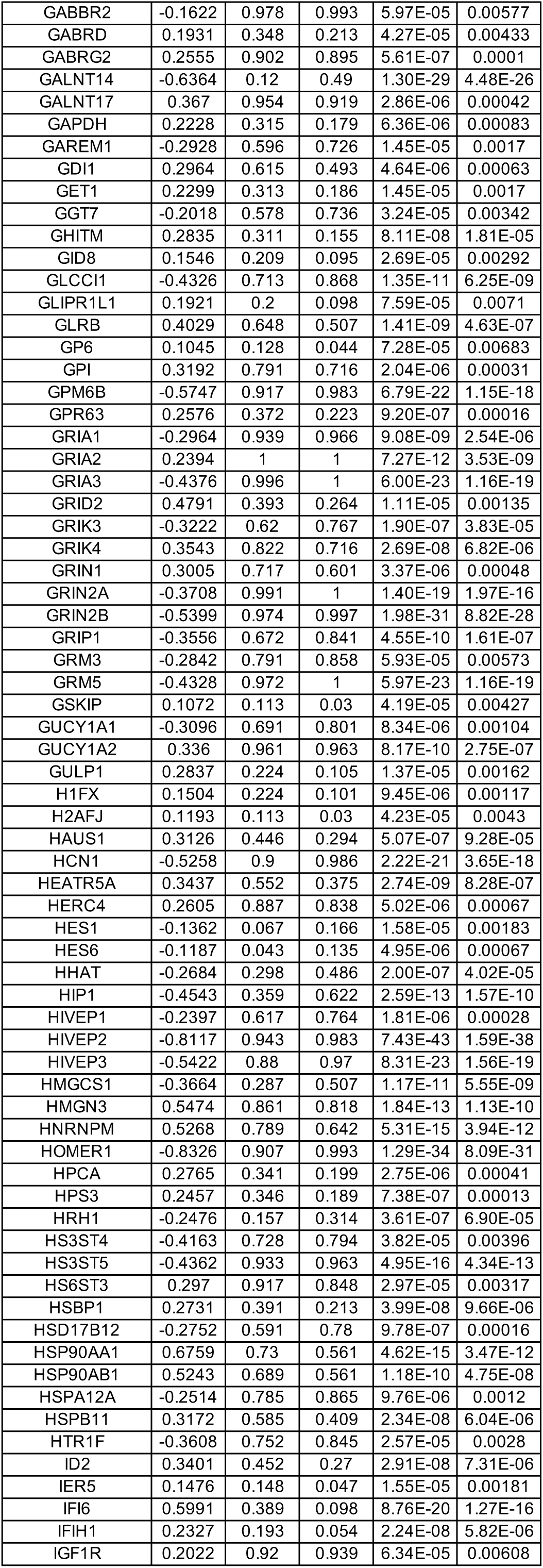
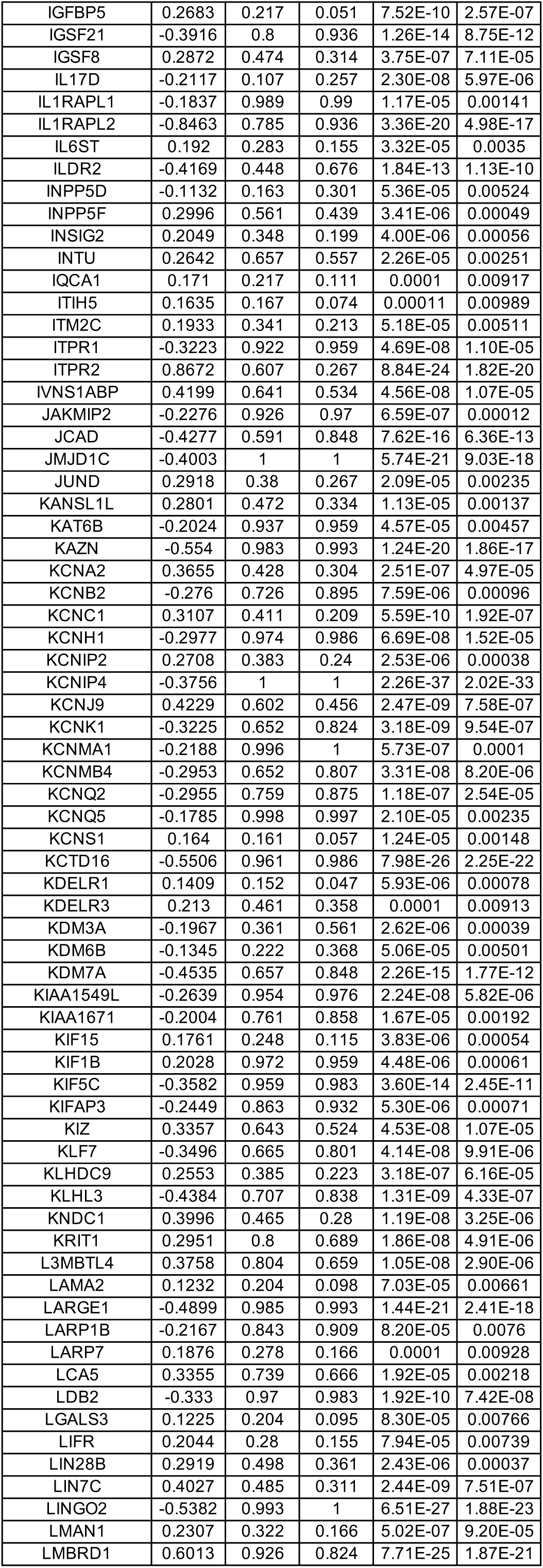
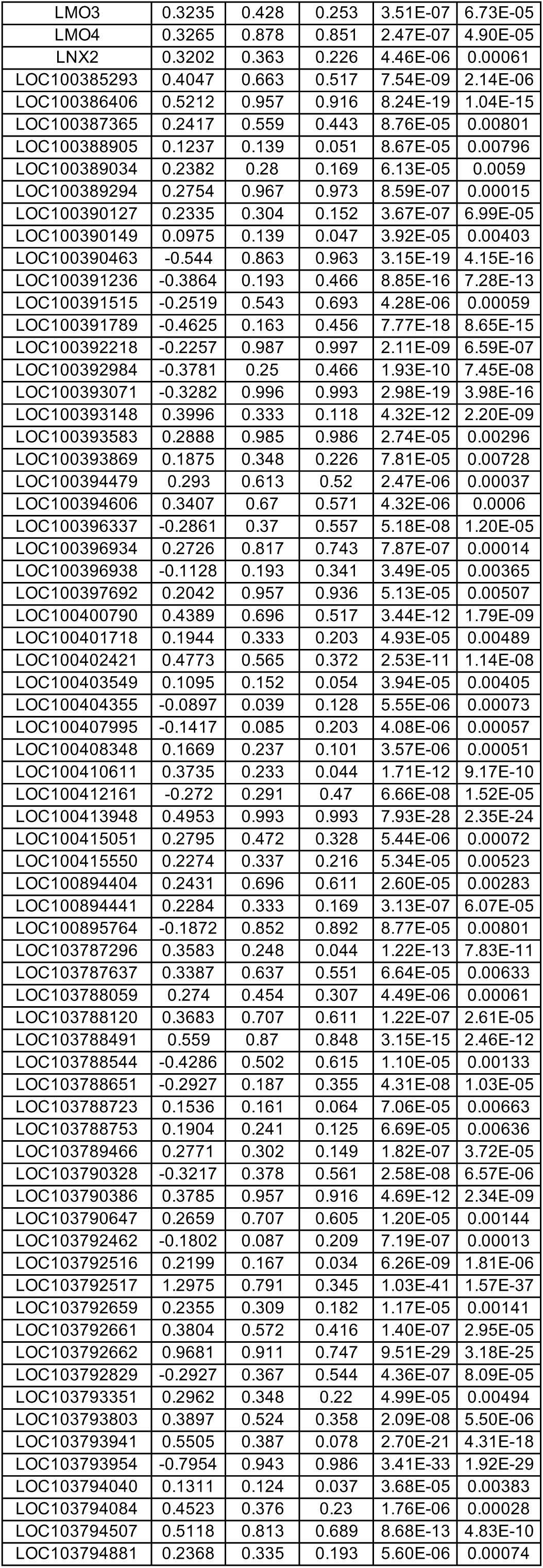
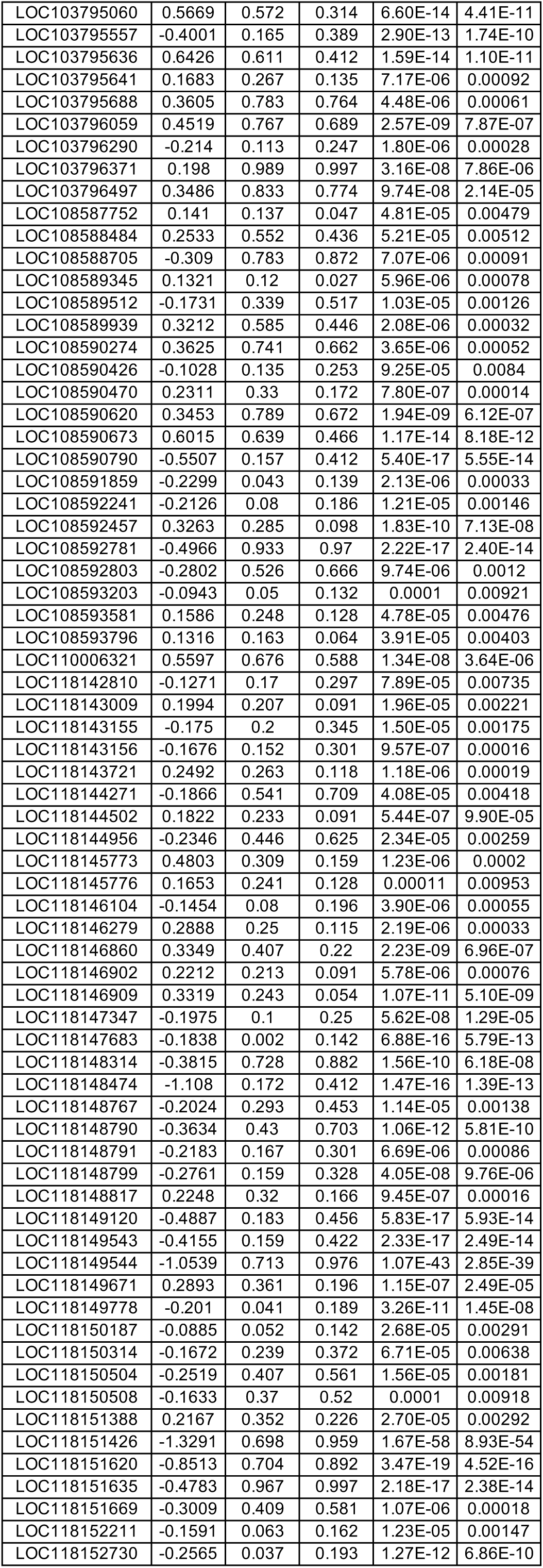
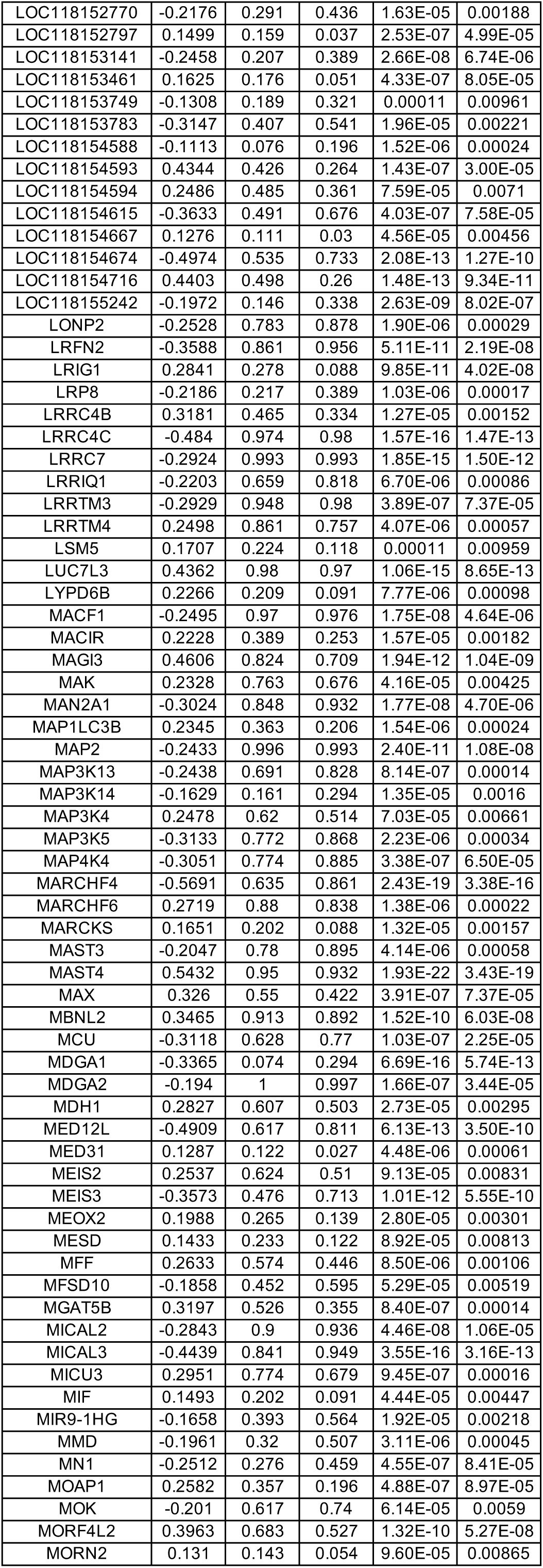
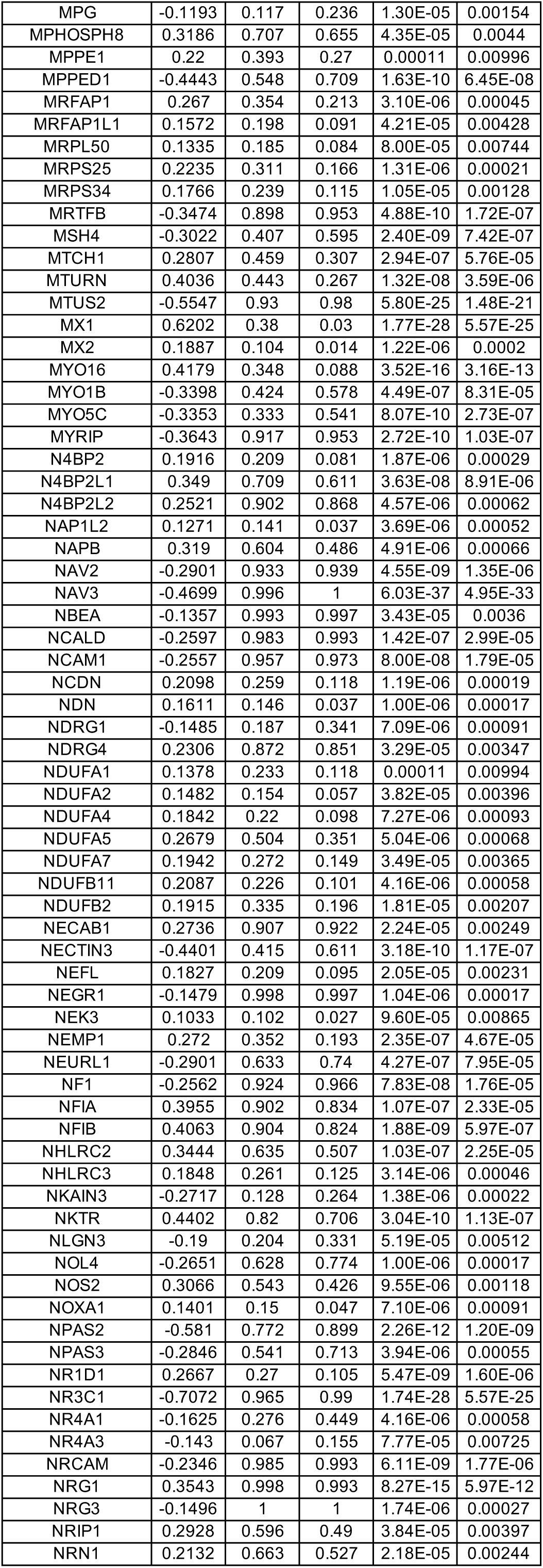
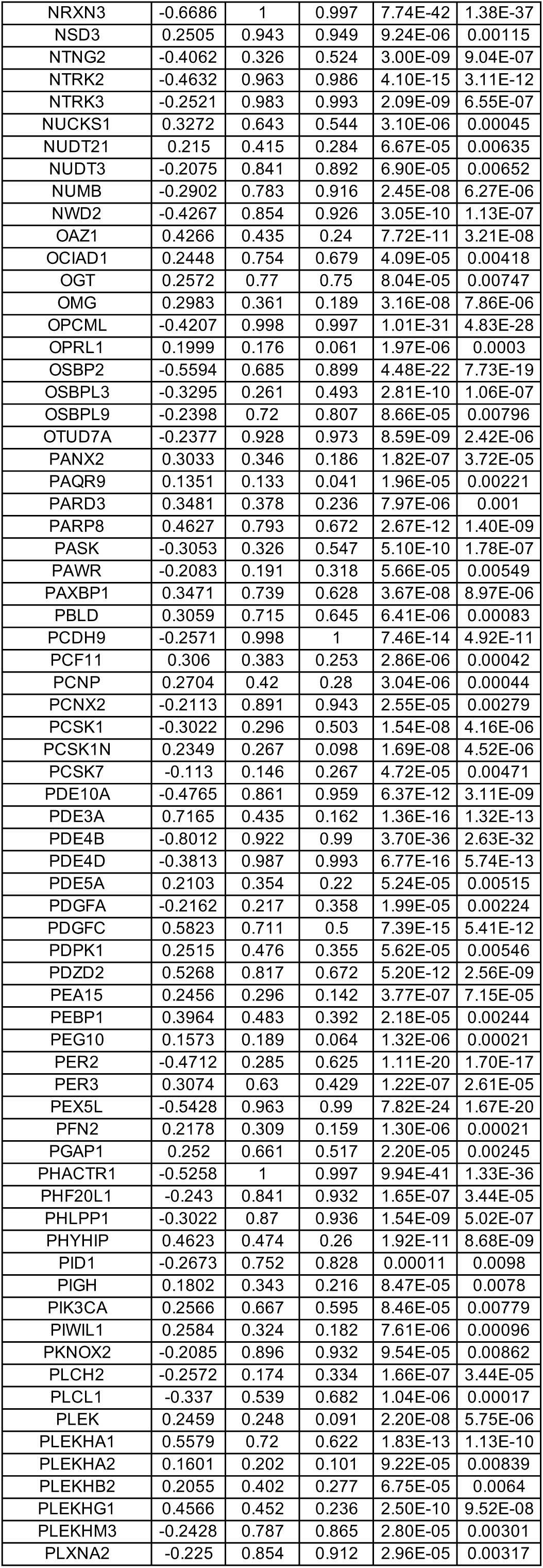
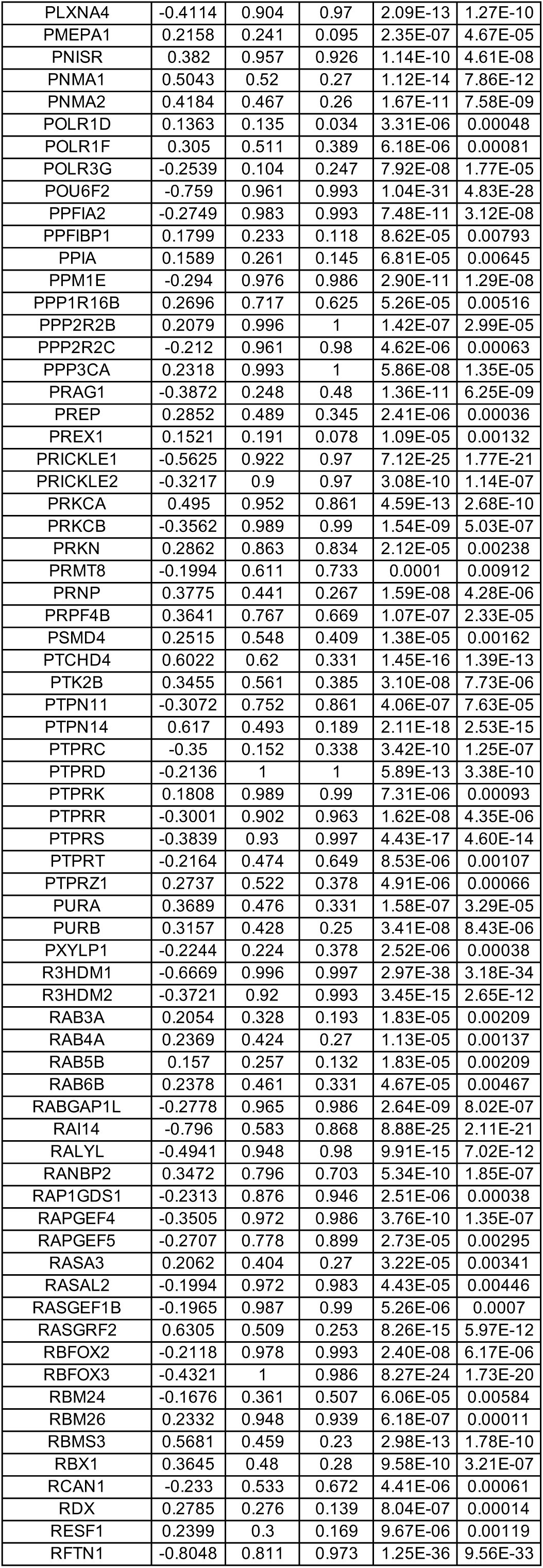
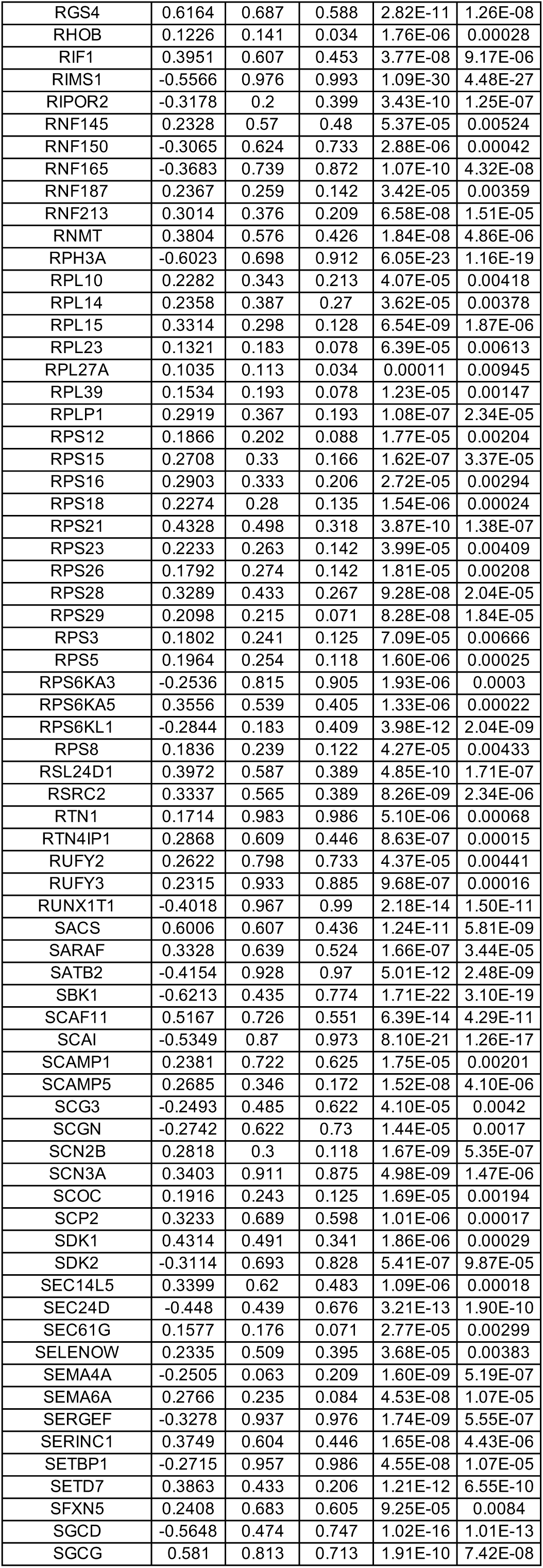
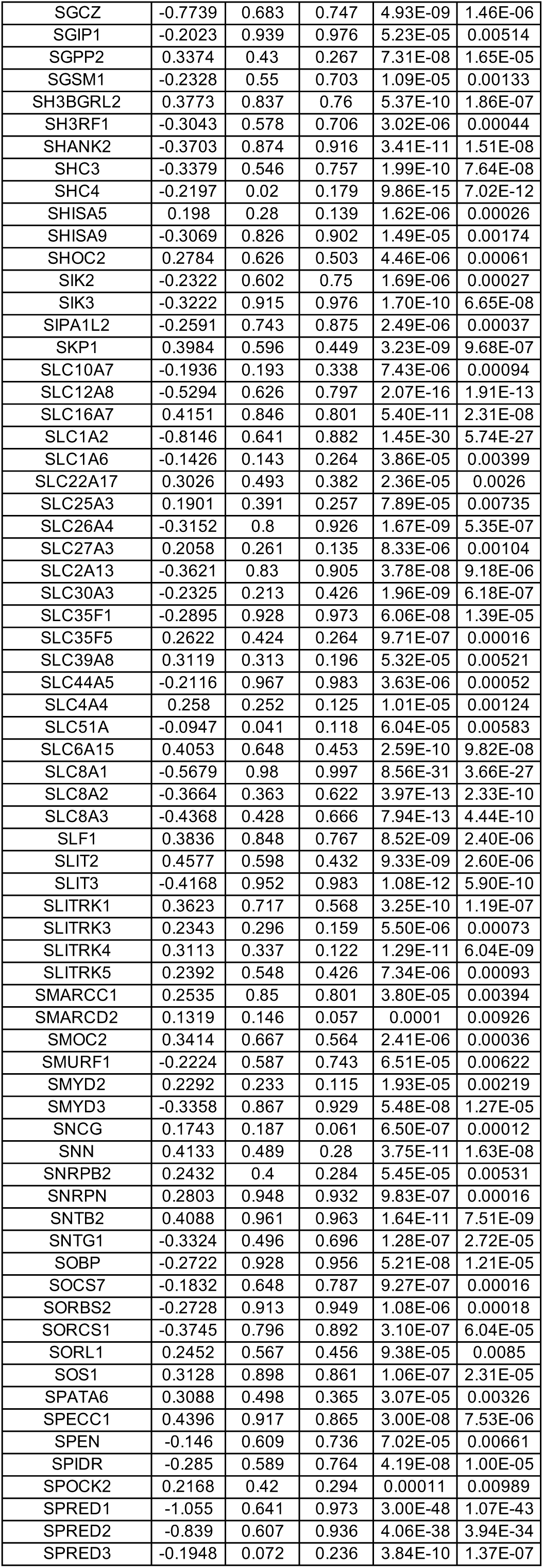
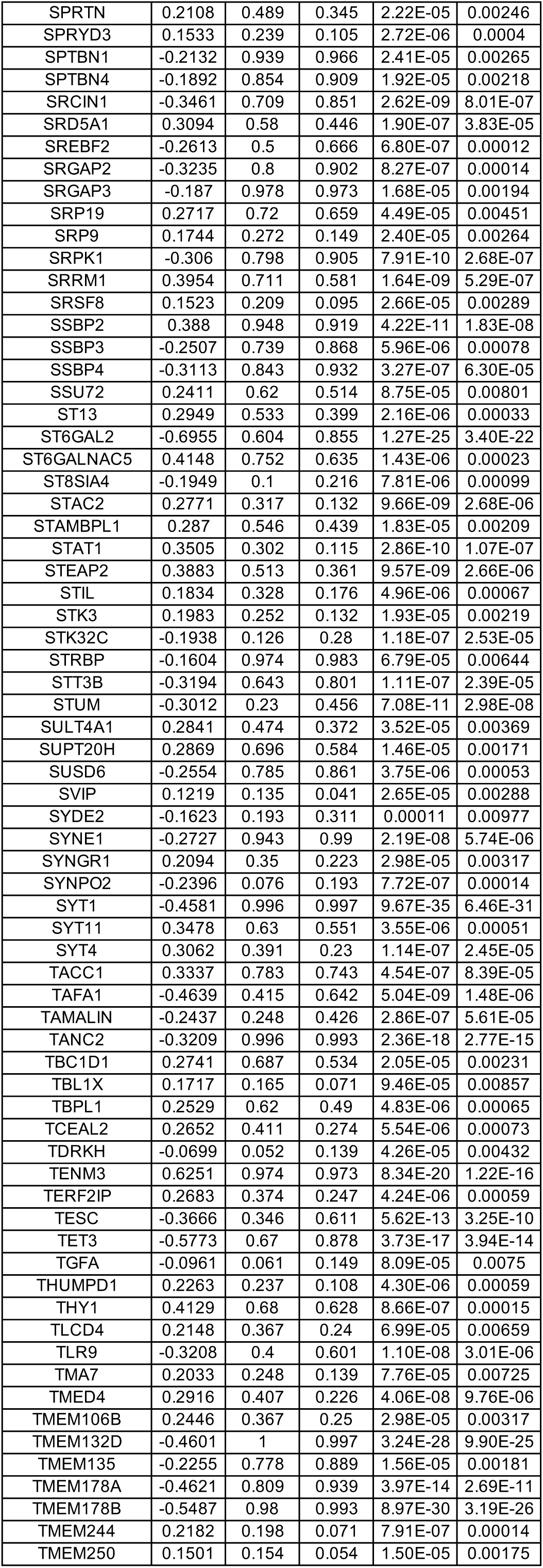
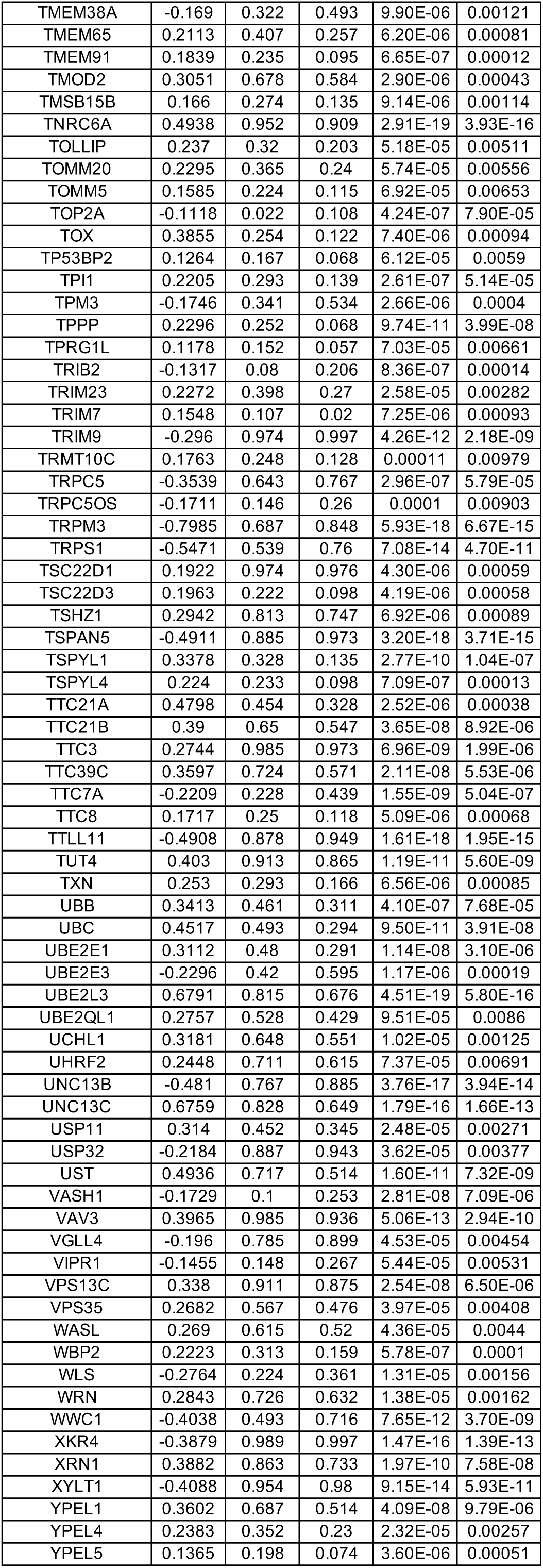
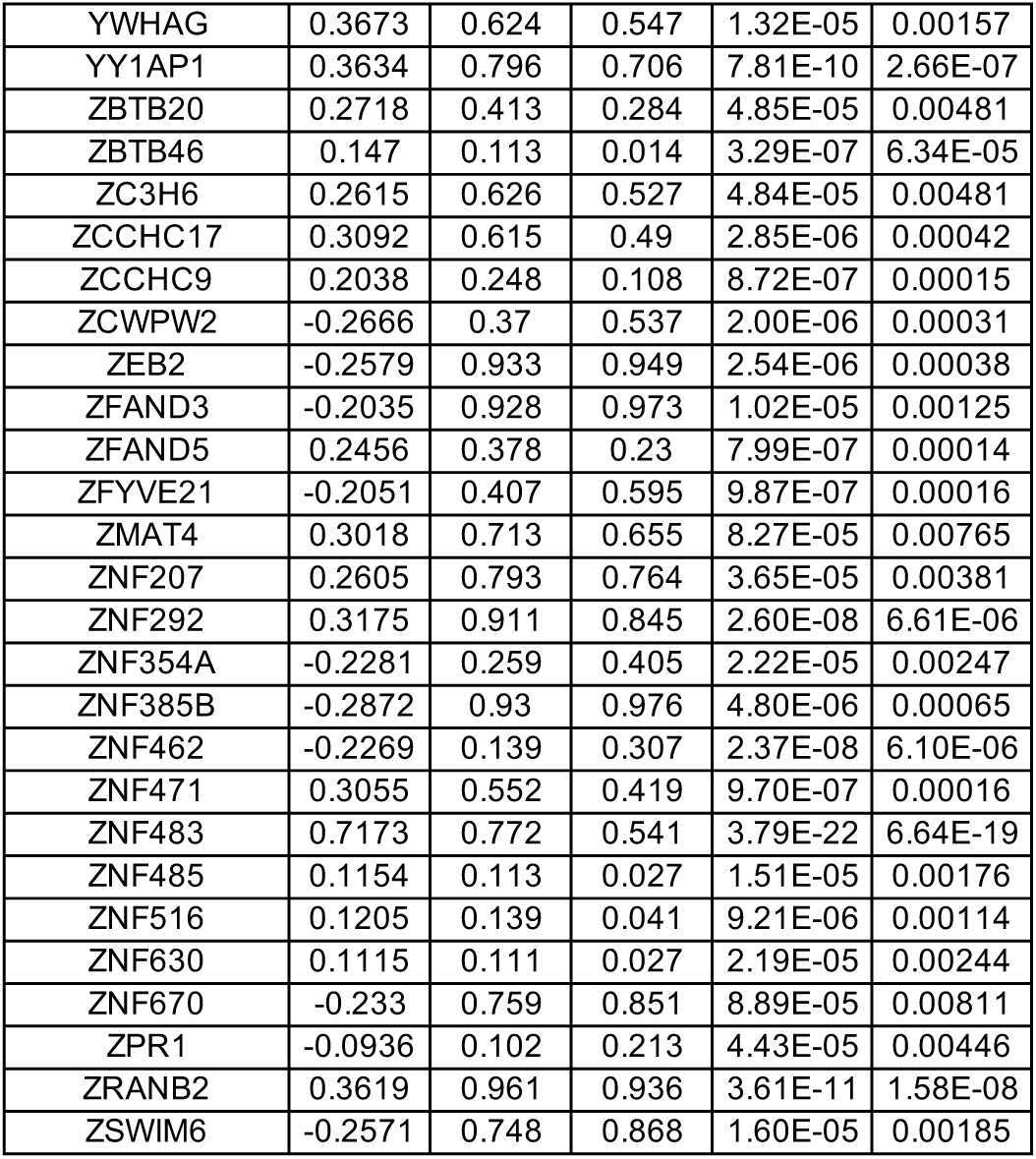
DEGs between wild-type (WT) and MECP2-null (KO) upper layer excitatory neurons (Ex_5) of OC (adjusted p-value < 0.01)

**Table S26.**
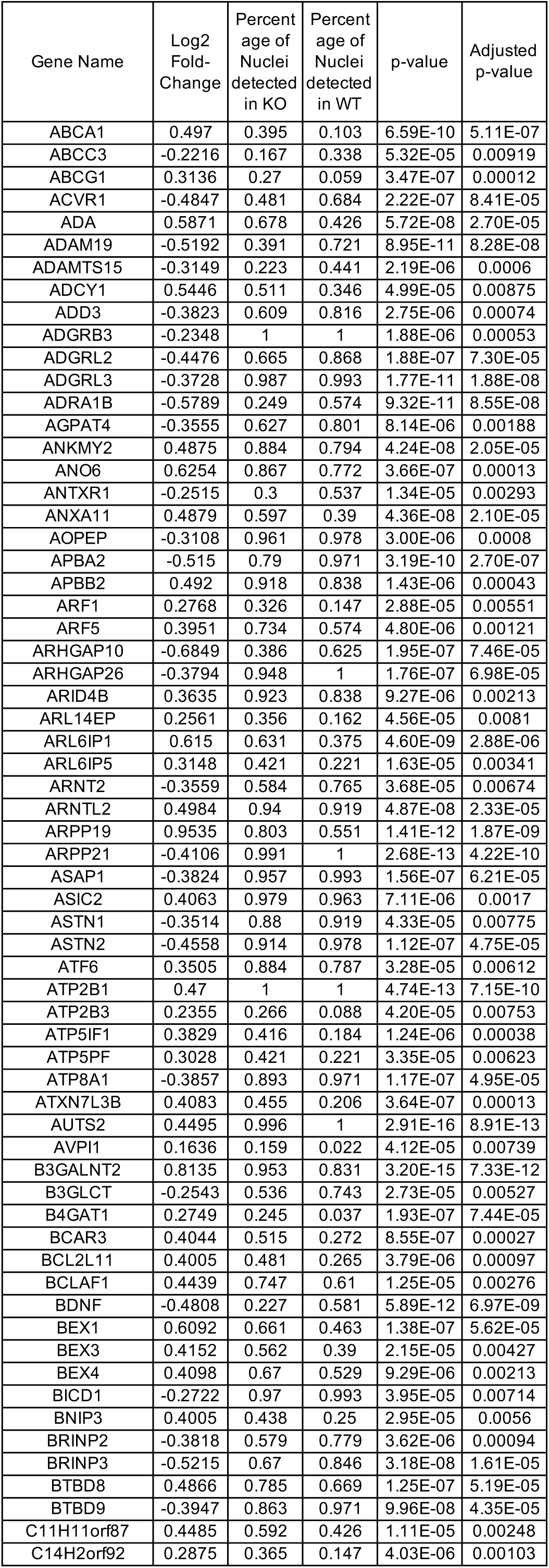
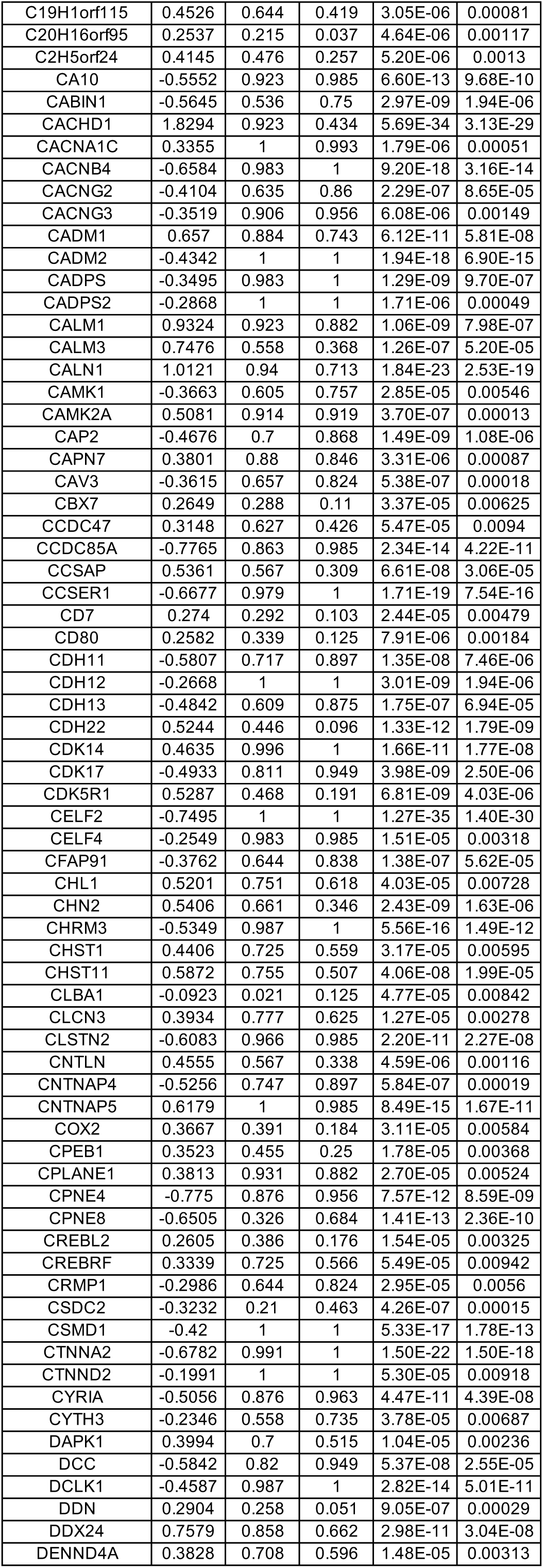
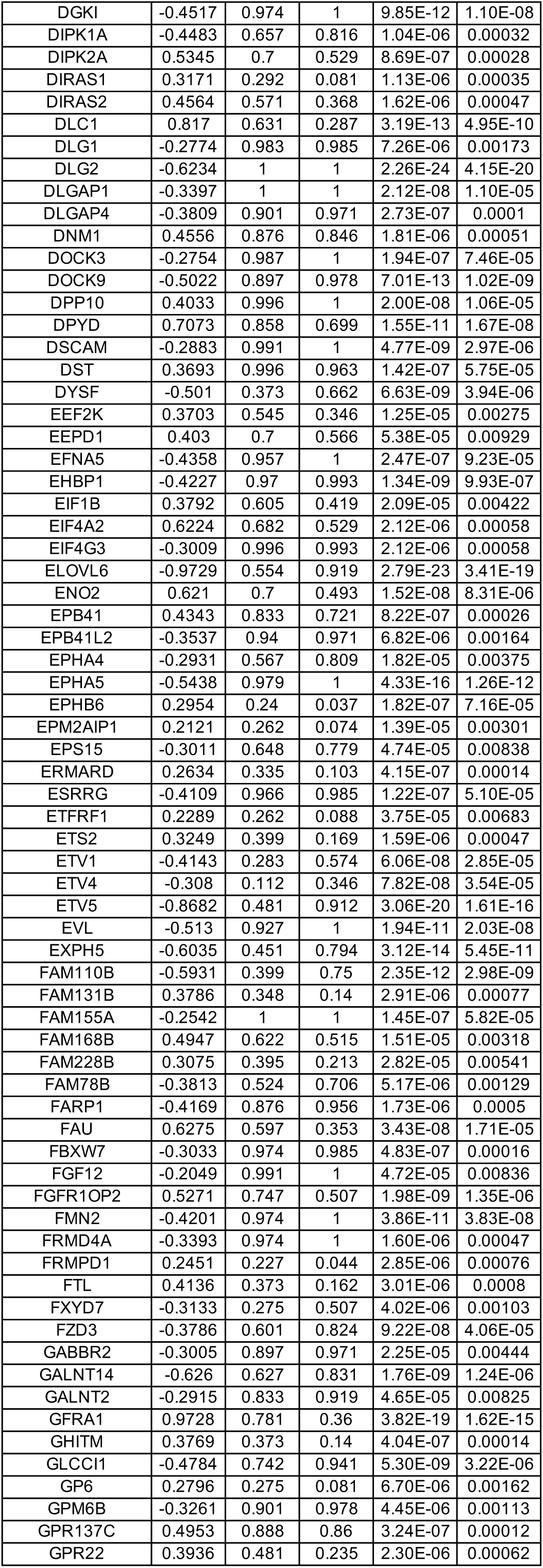
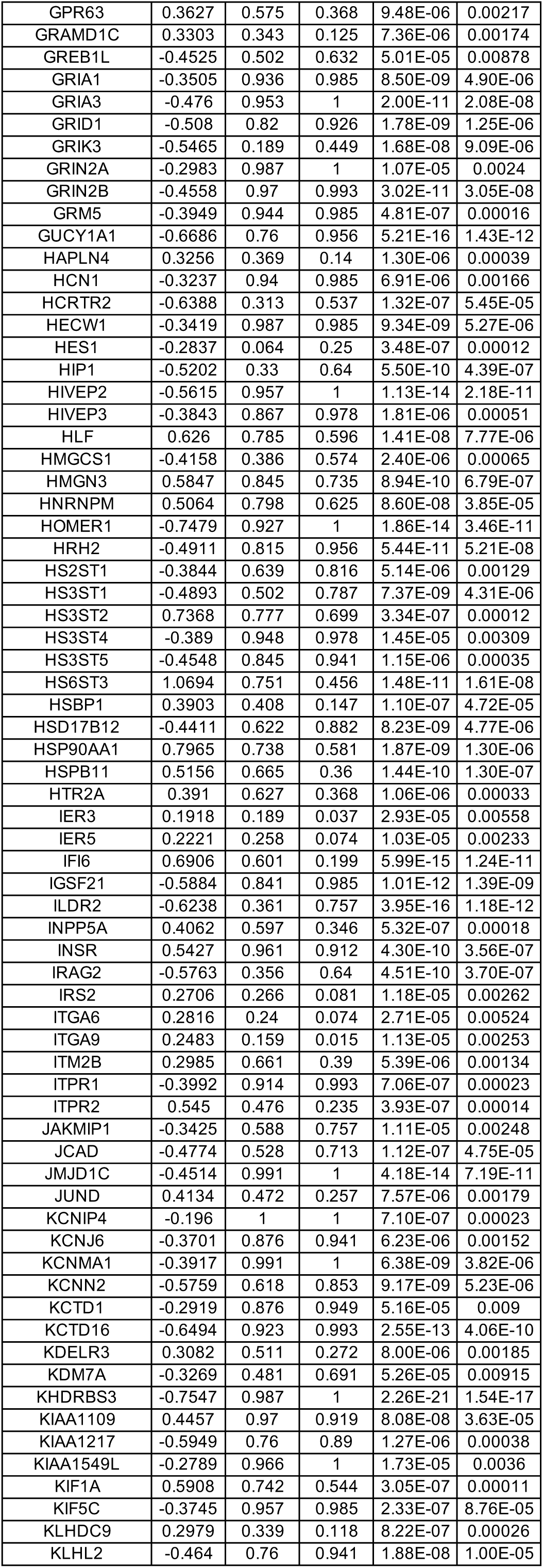
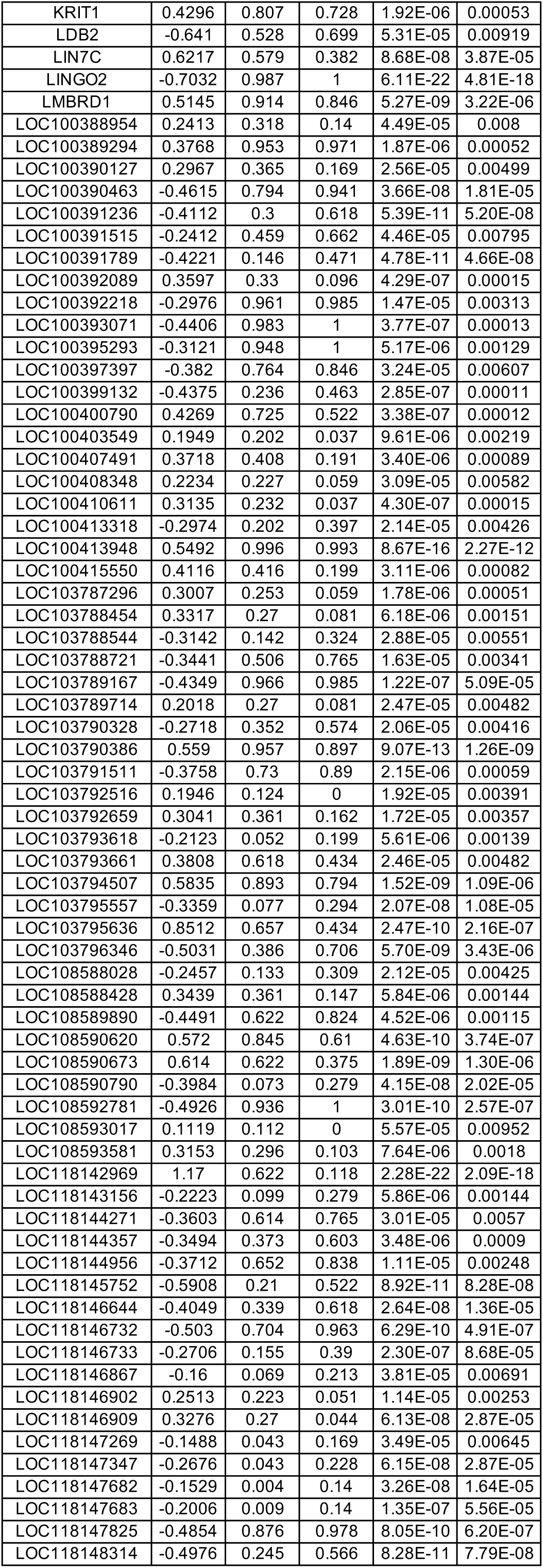
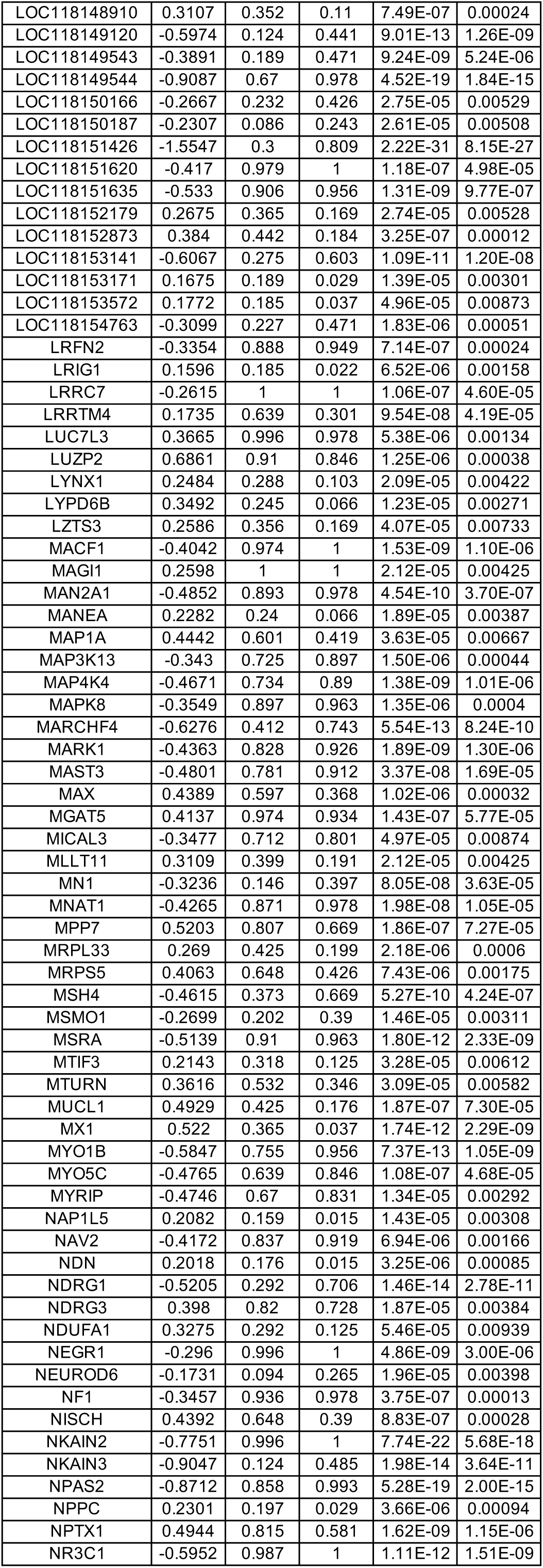
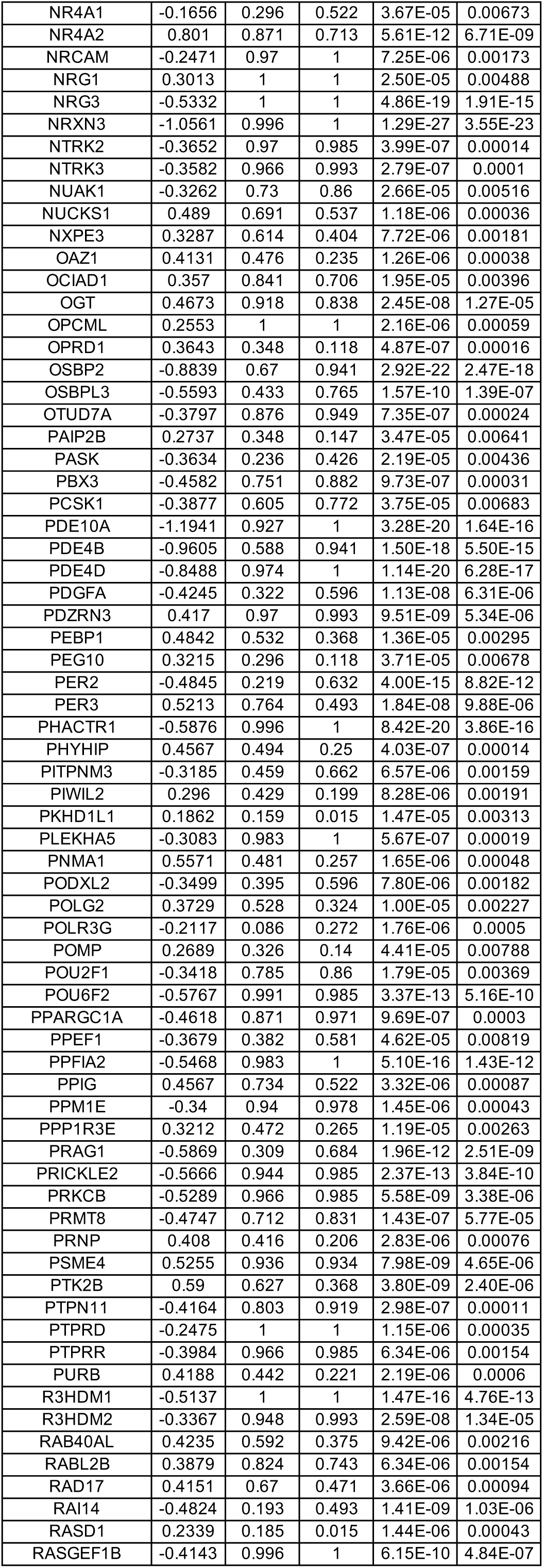
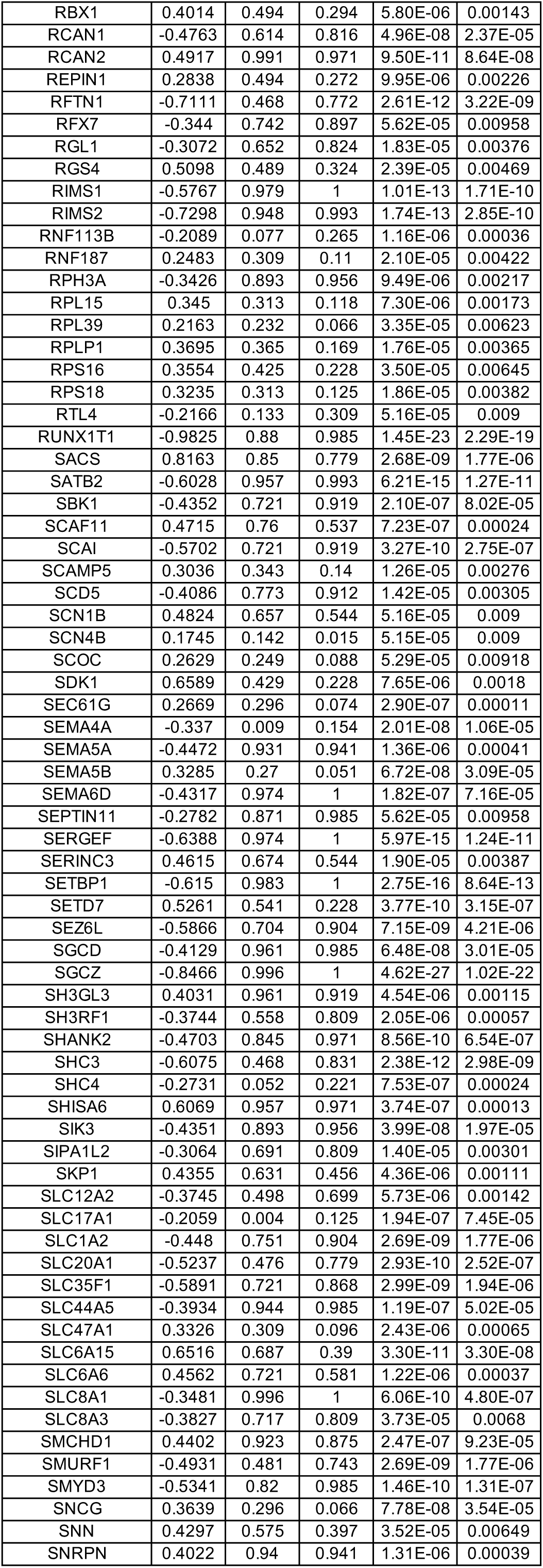
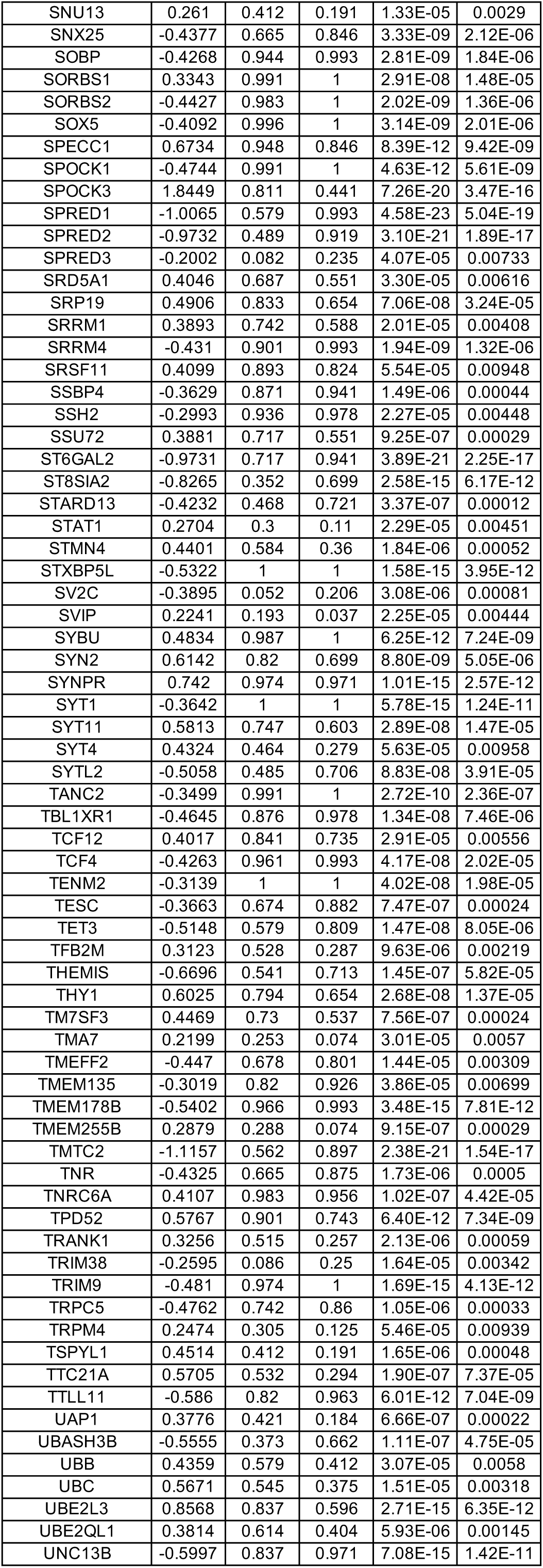
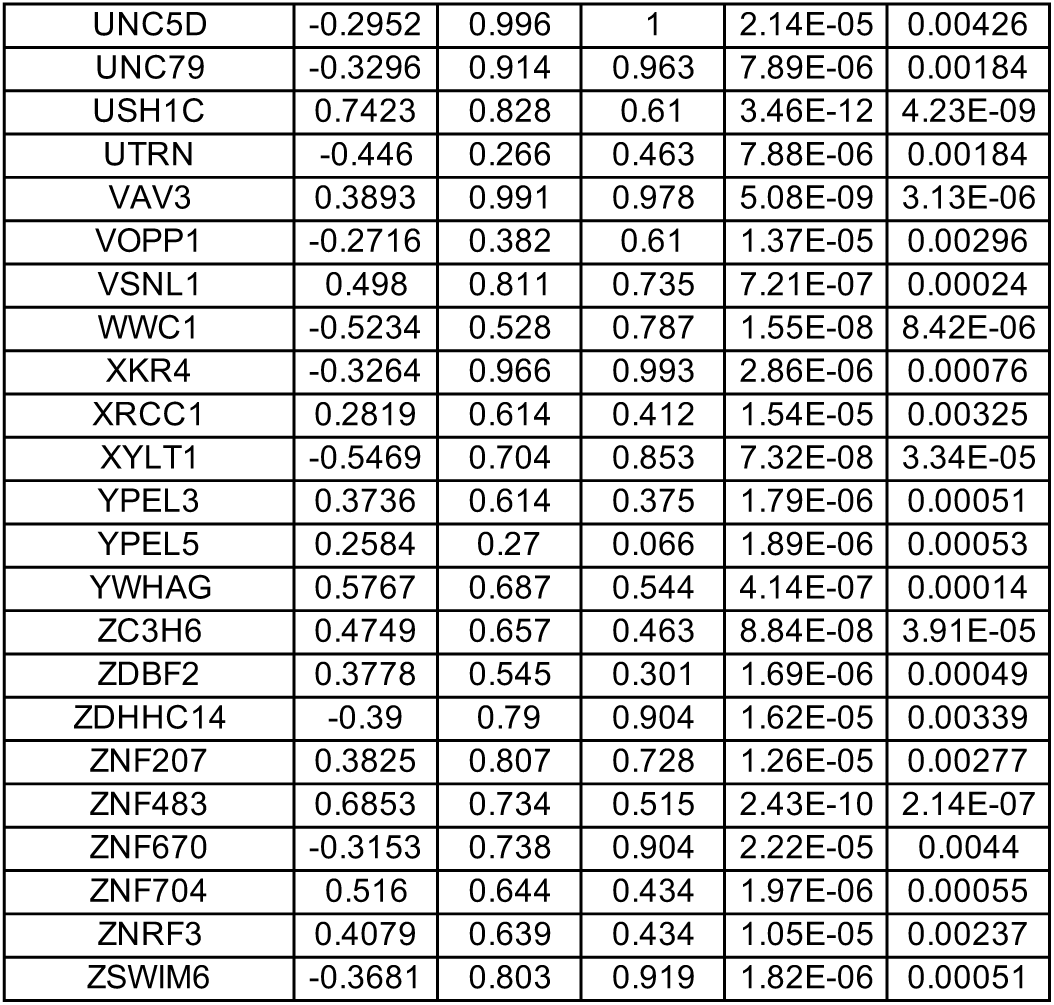
DEGs between wild-type (WT) and MECP2-null (KO) upper layer excitatory neurons (Ex_6) of OC (adjusted p-value < 0.01)

**Table S27.**
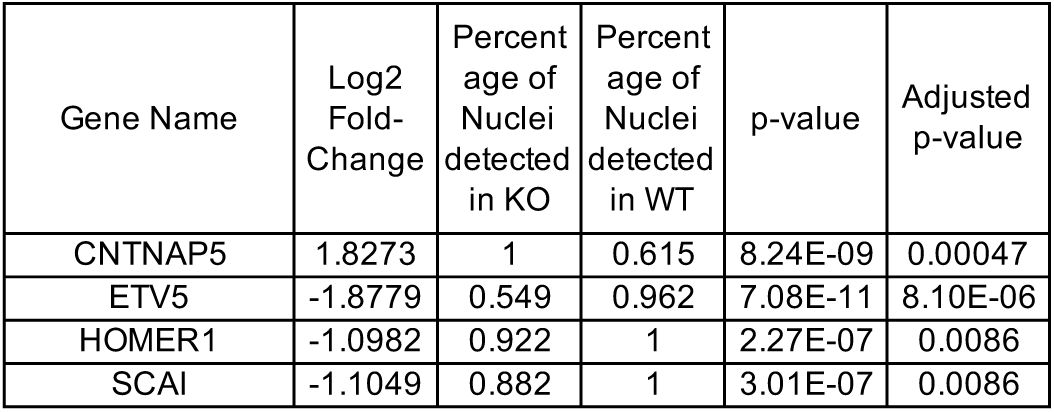
DEGs between wild-type (WT) and MECP2-null (KO) upper layer excitatory neurons (Ex_7) of OC (adjusted p-value < 0.01)

**Table S28.**
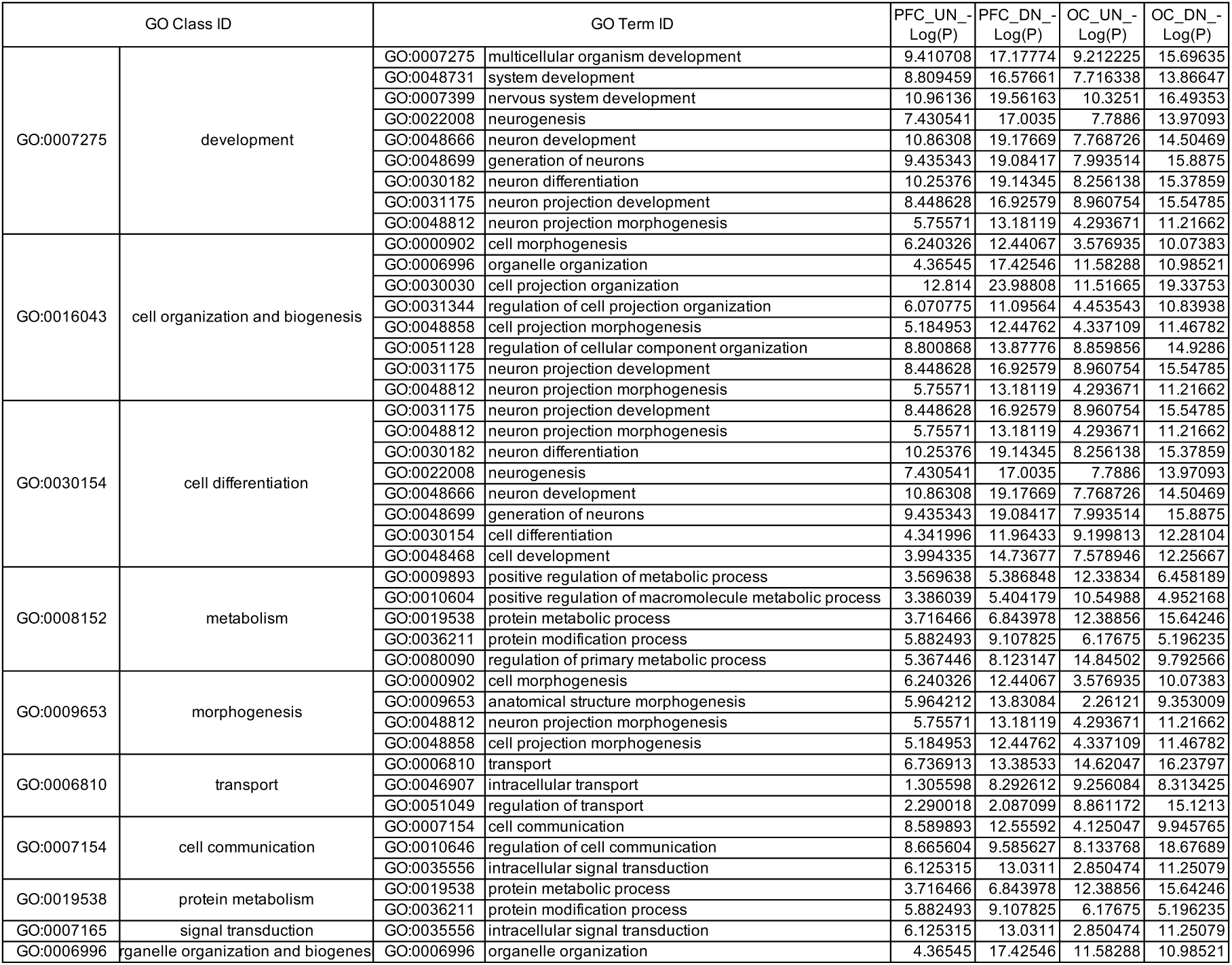
Subclassification of the top 30 GO categories of upregulated genes in MECP2-null upper layer excitatory neurons (UN) and deep layer excitatory neurons (DN) of the PFC and OC.

**Table S29.**
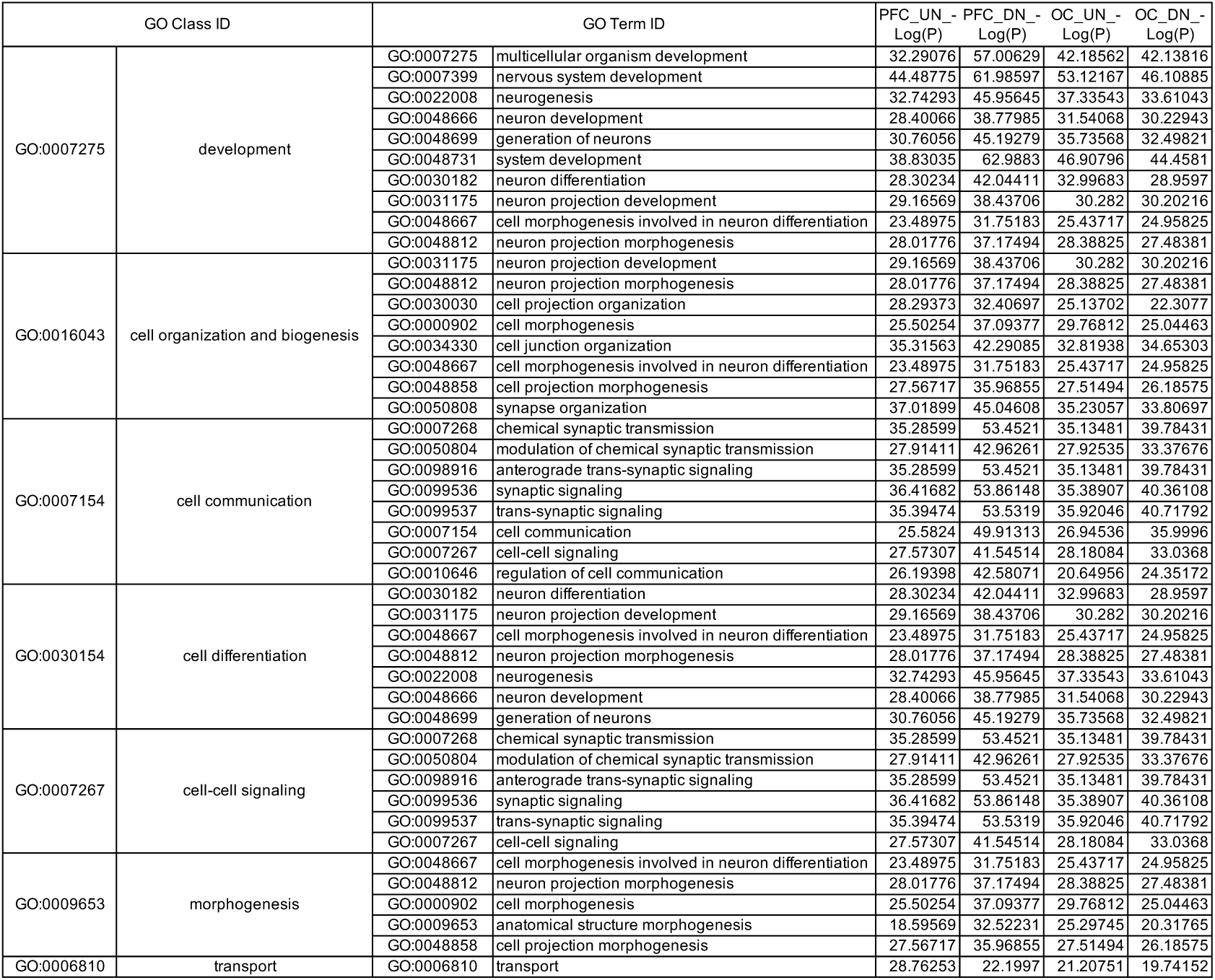
Subclassification of the top 30 GO categories of downregulated genes in MECP2-null upper layer excitatory neurons (UN) and deep layer excitatory neurons (DN) of the PFC and OC.

**Table S30.**
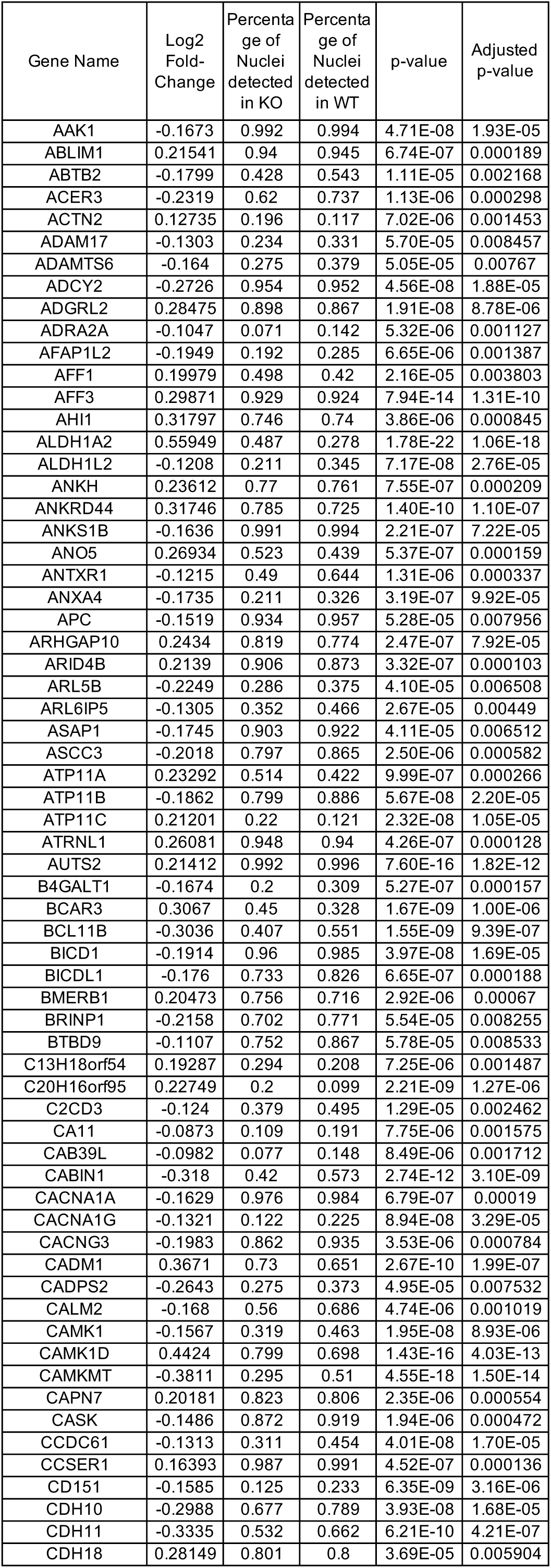
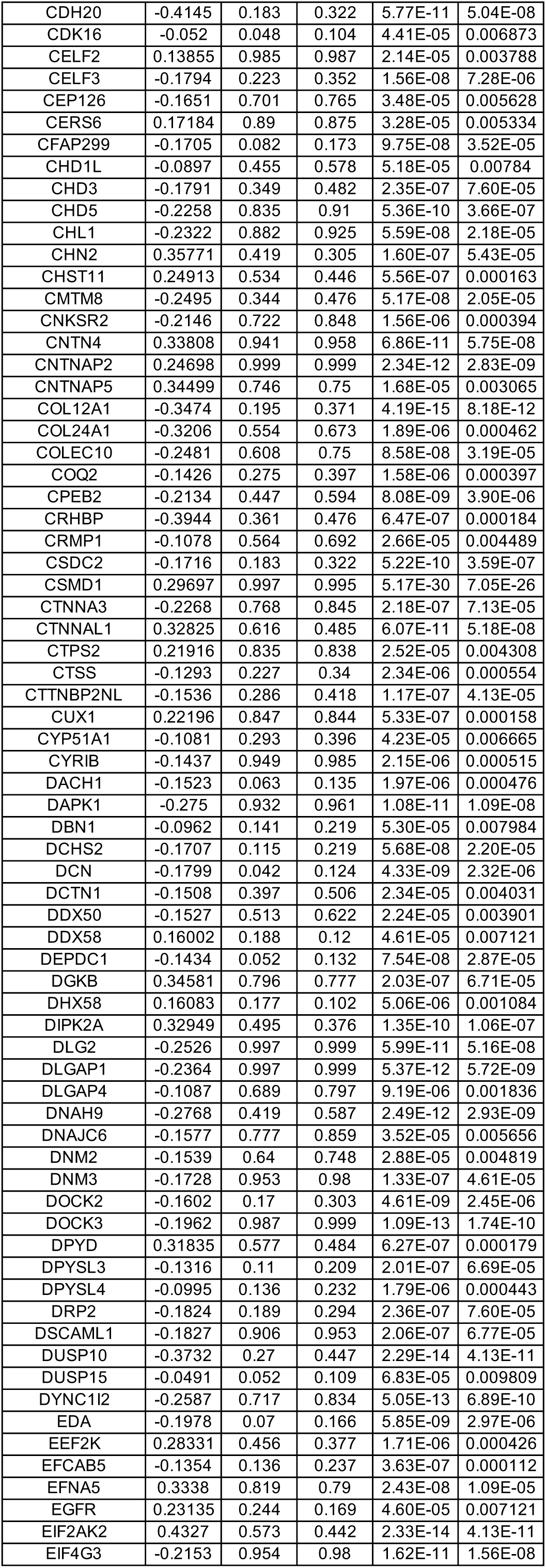
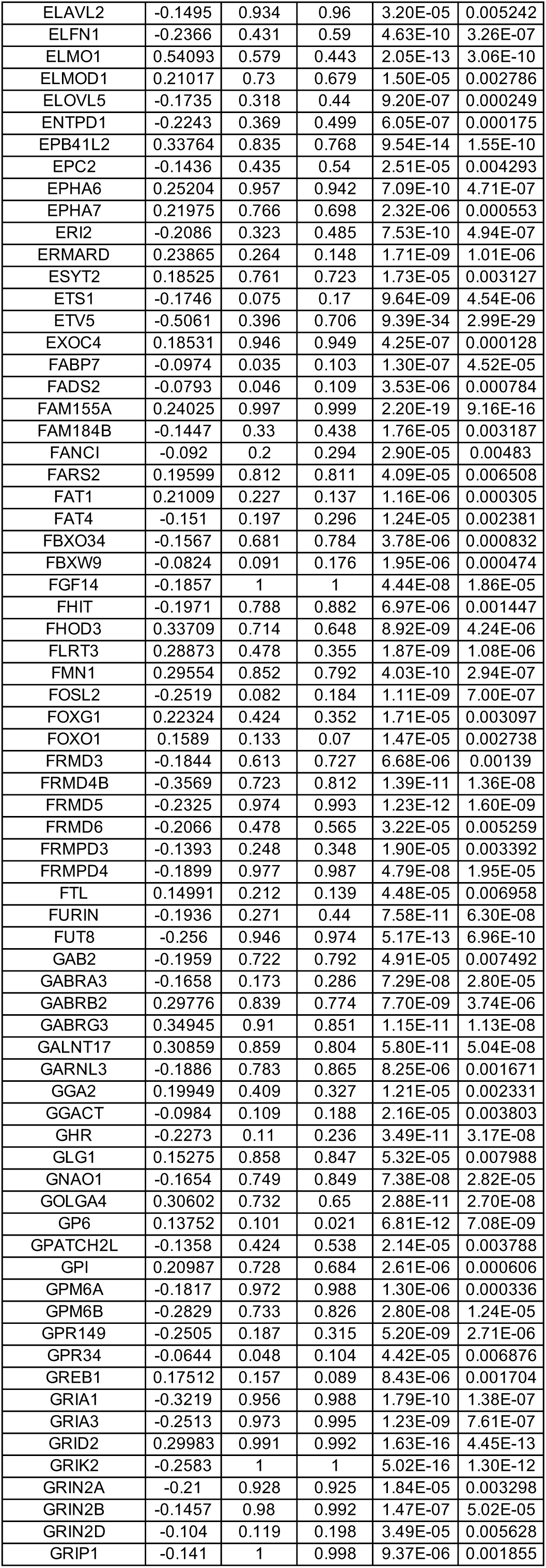
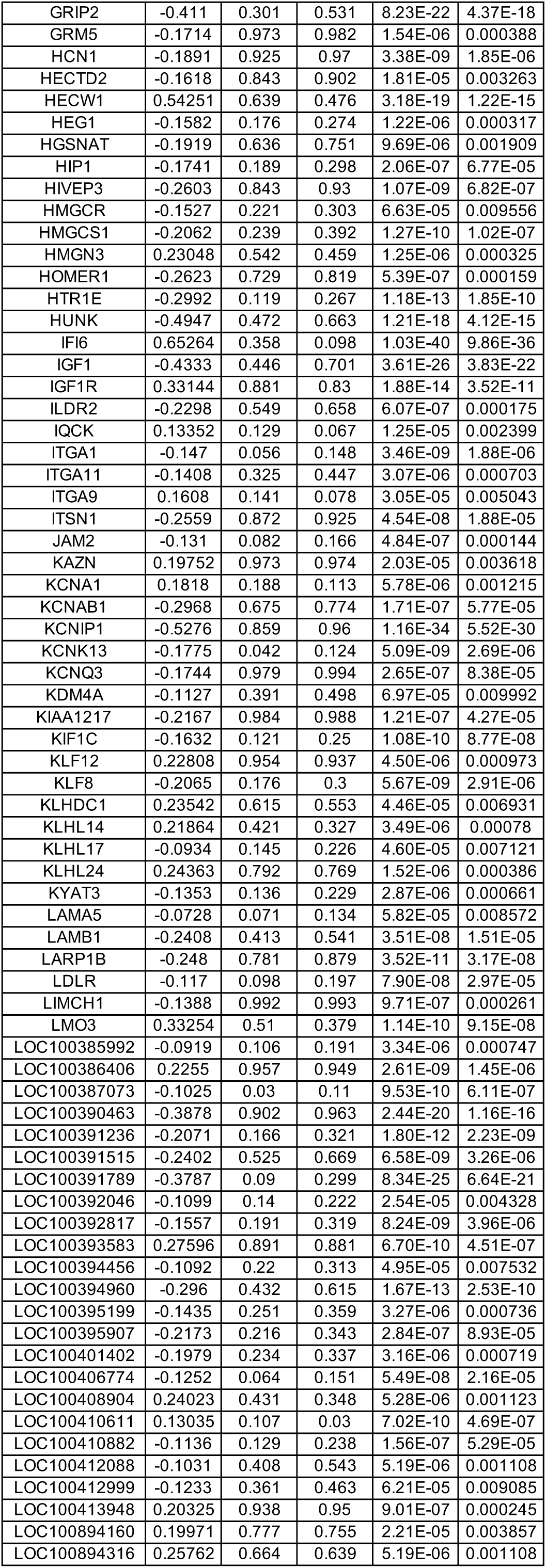
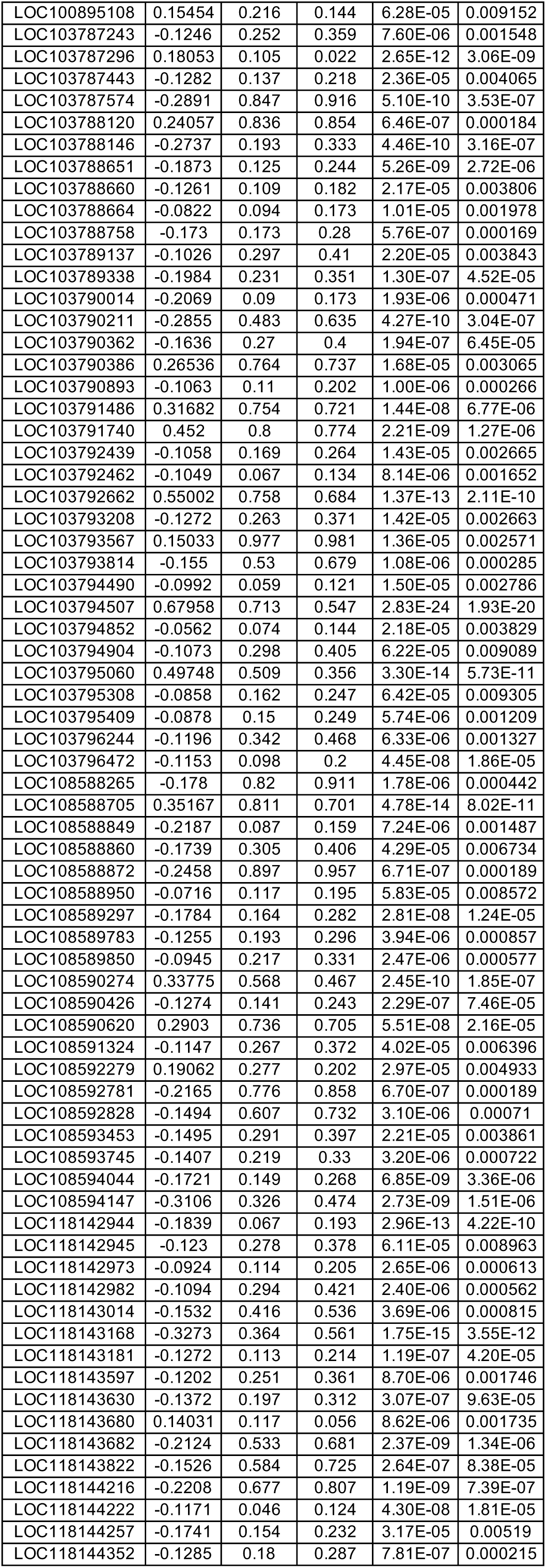
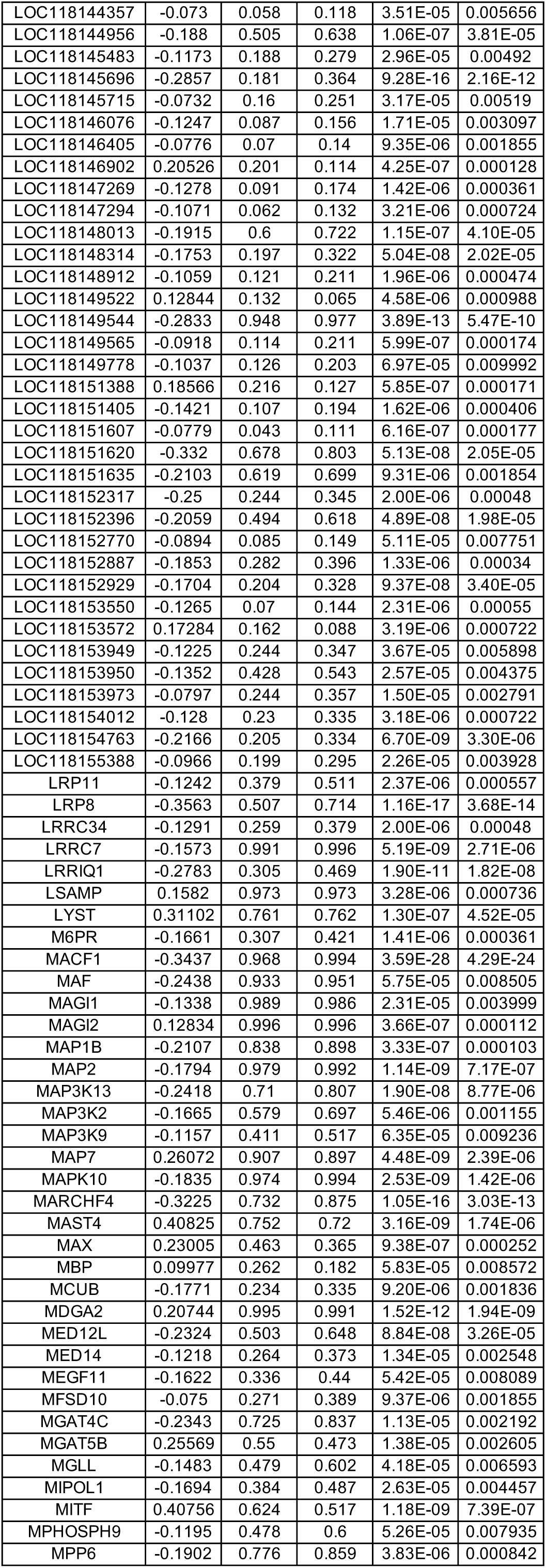
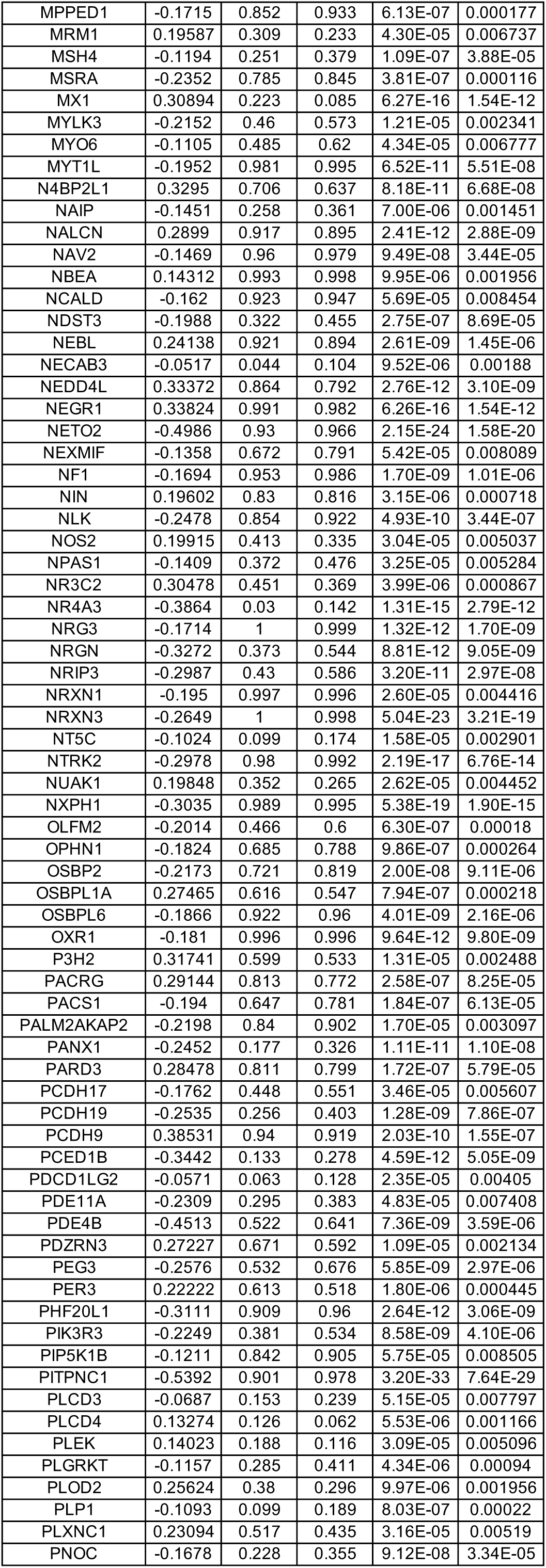
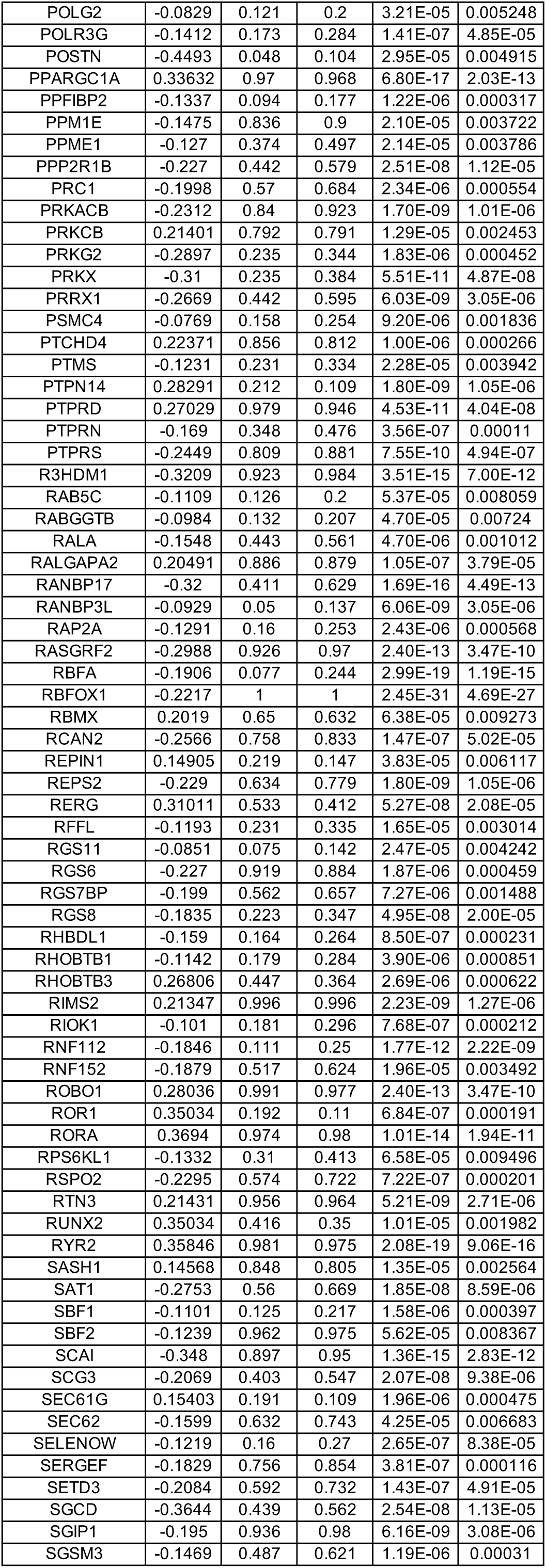
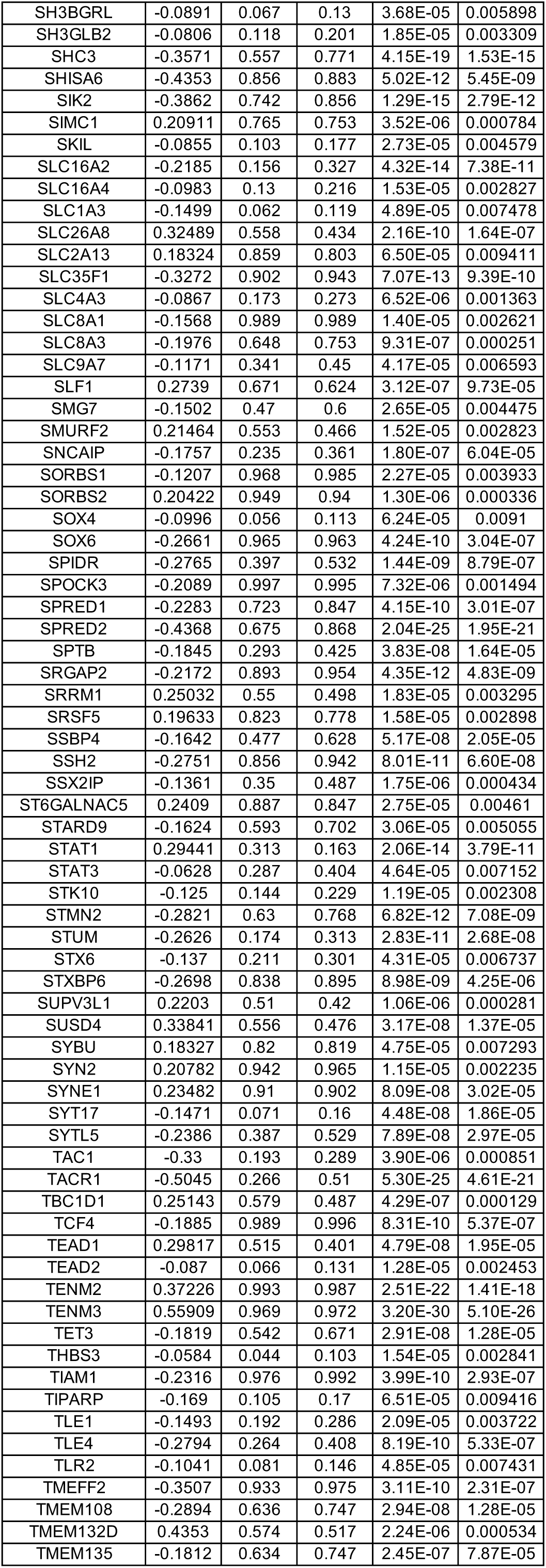
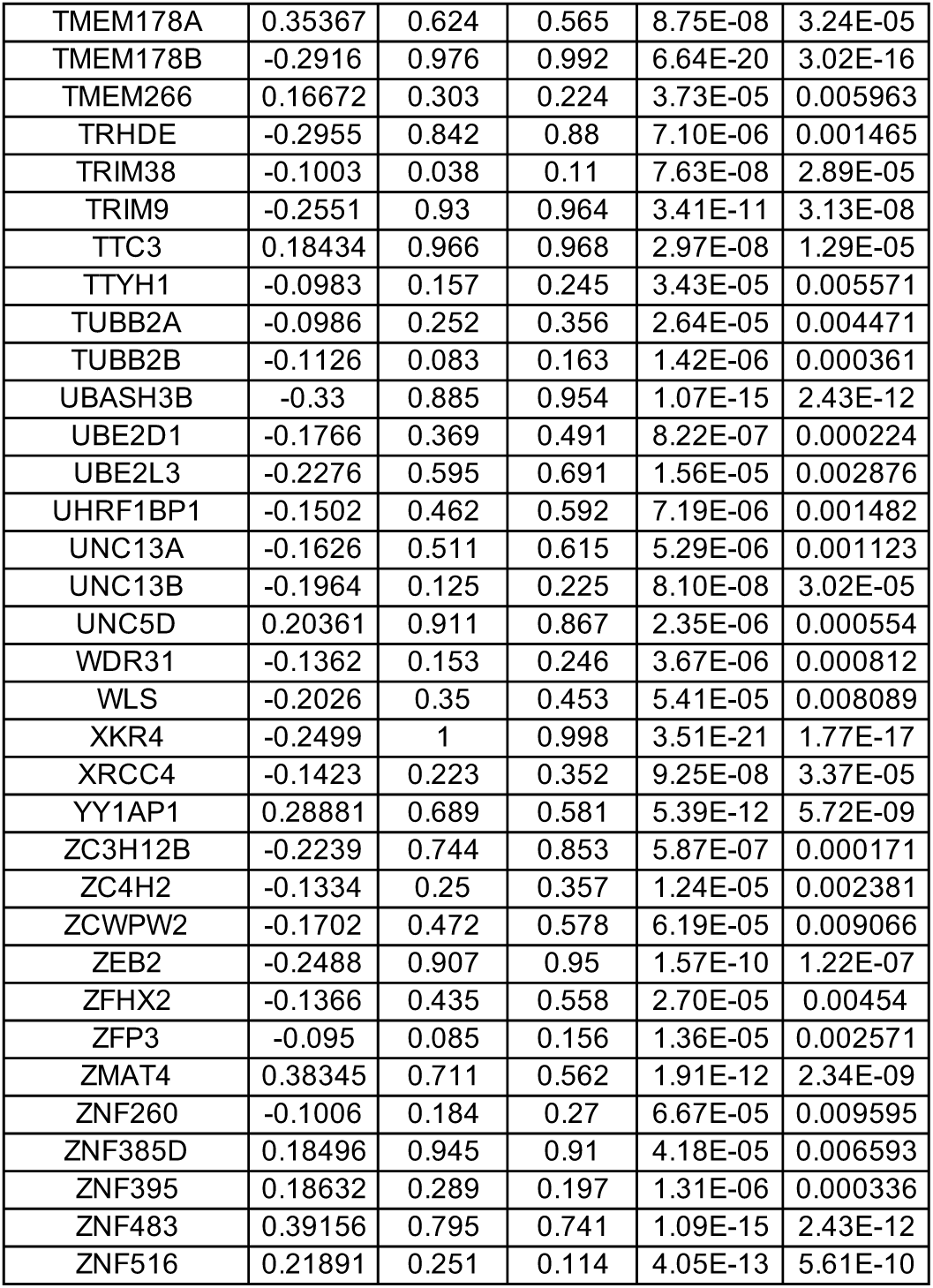
DEGs between wild-type (WT) and MECP2-null (KO) SST-positive inhibitory neurons of PFC (adjusted p-value < 0.01)

**Table S31.**
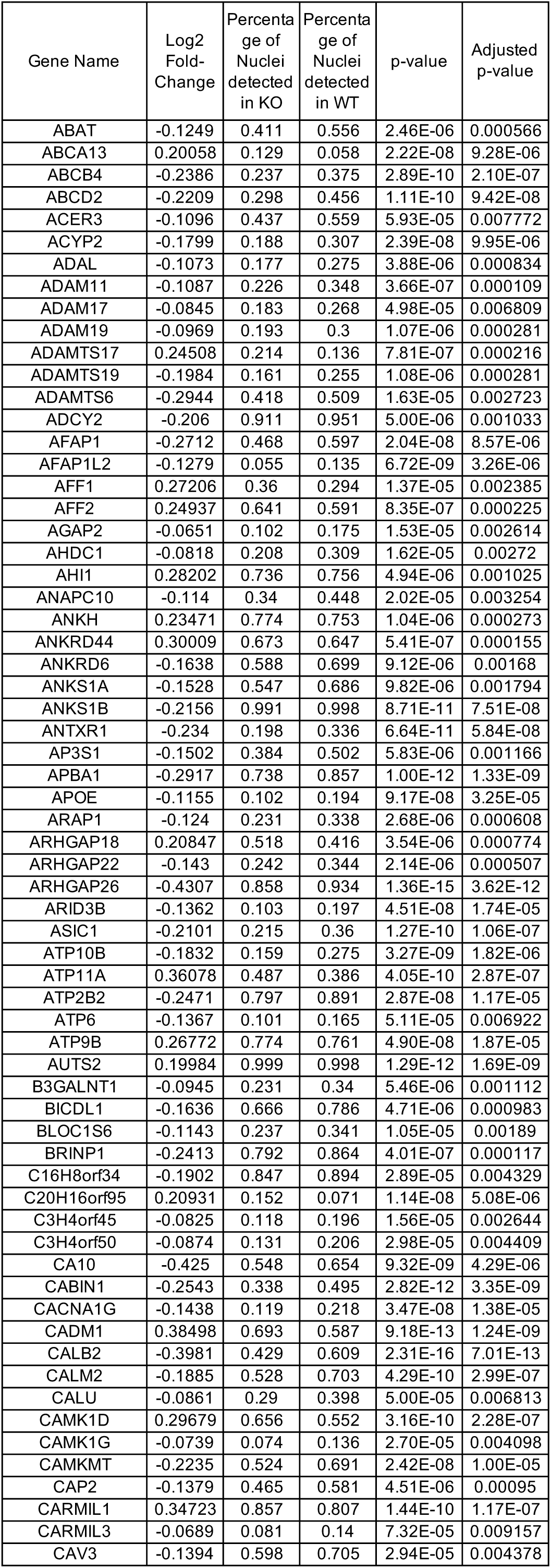
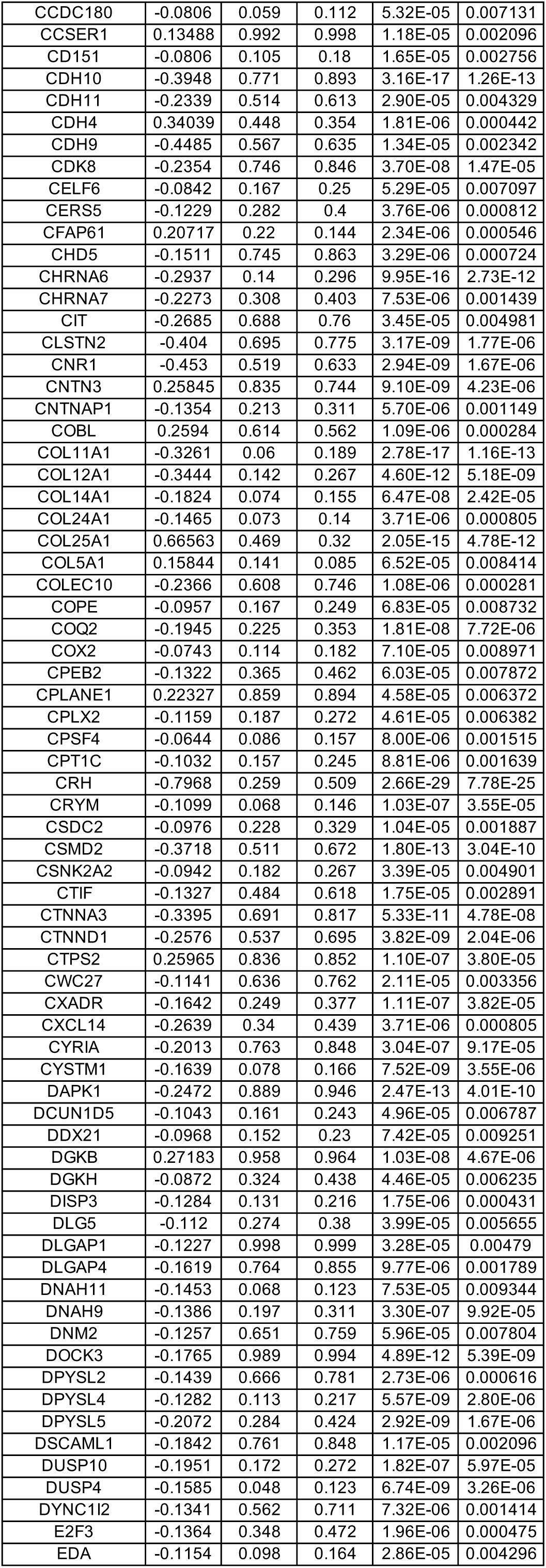
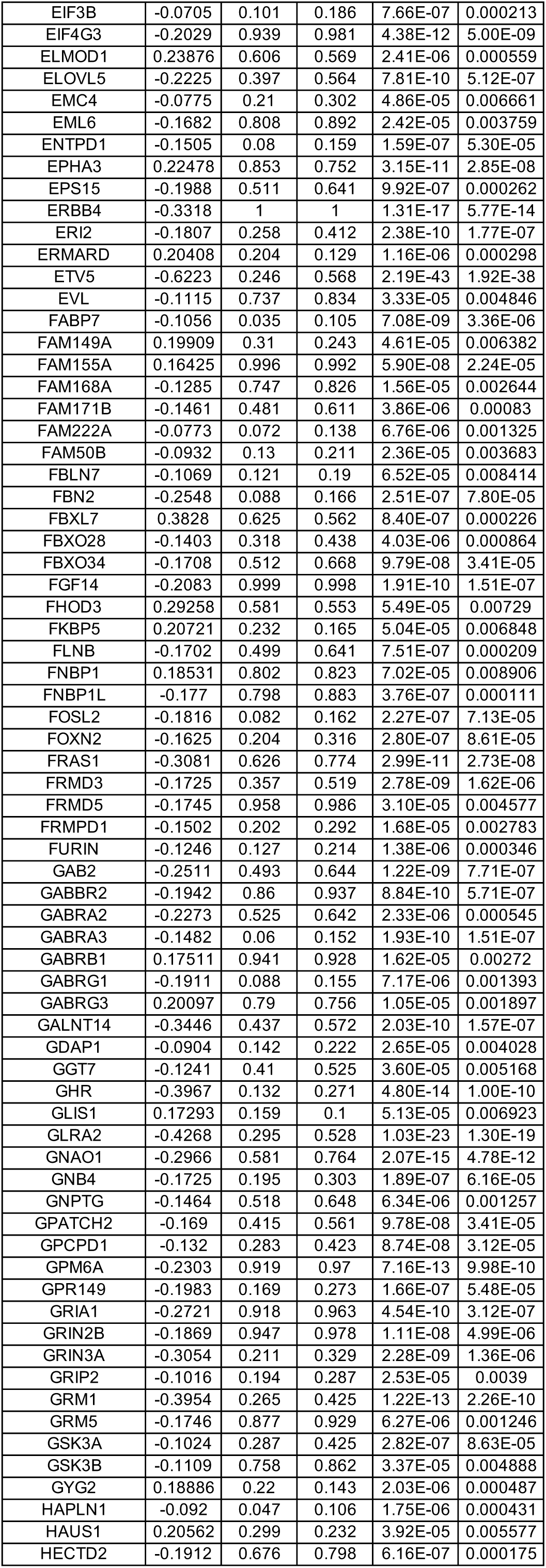
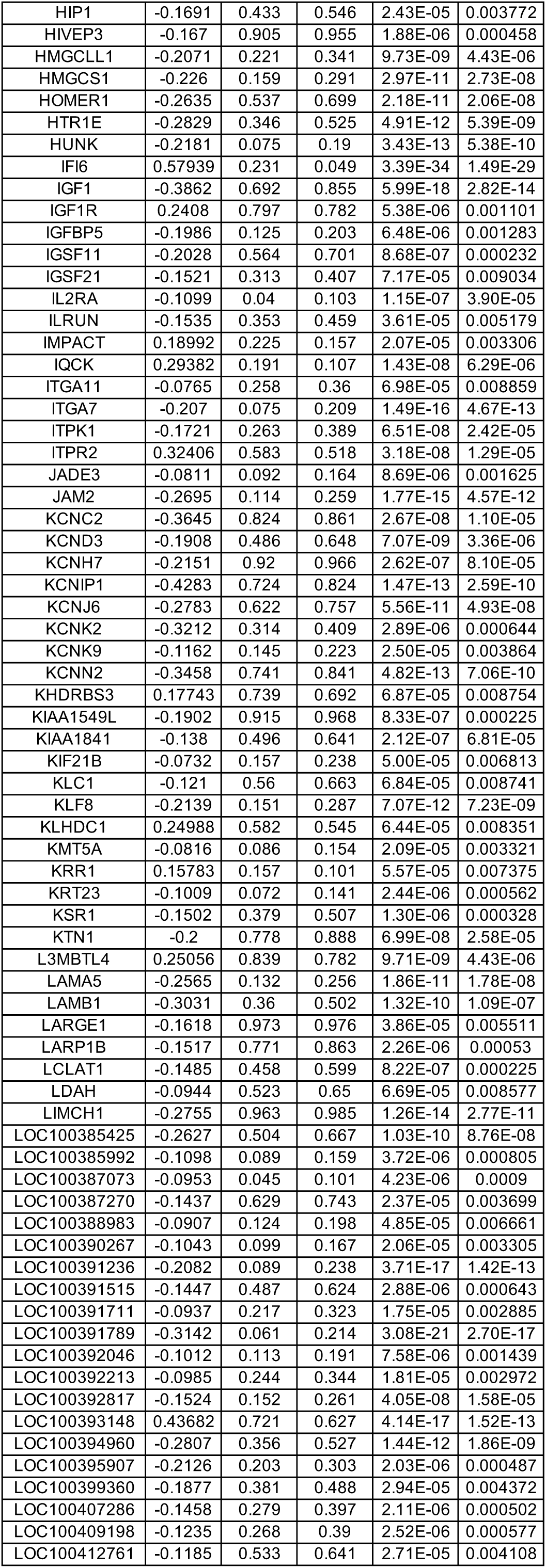
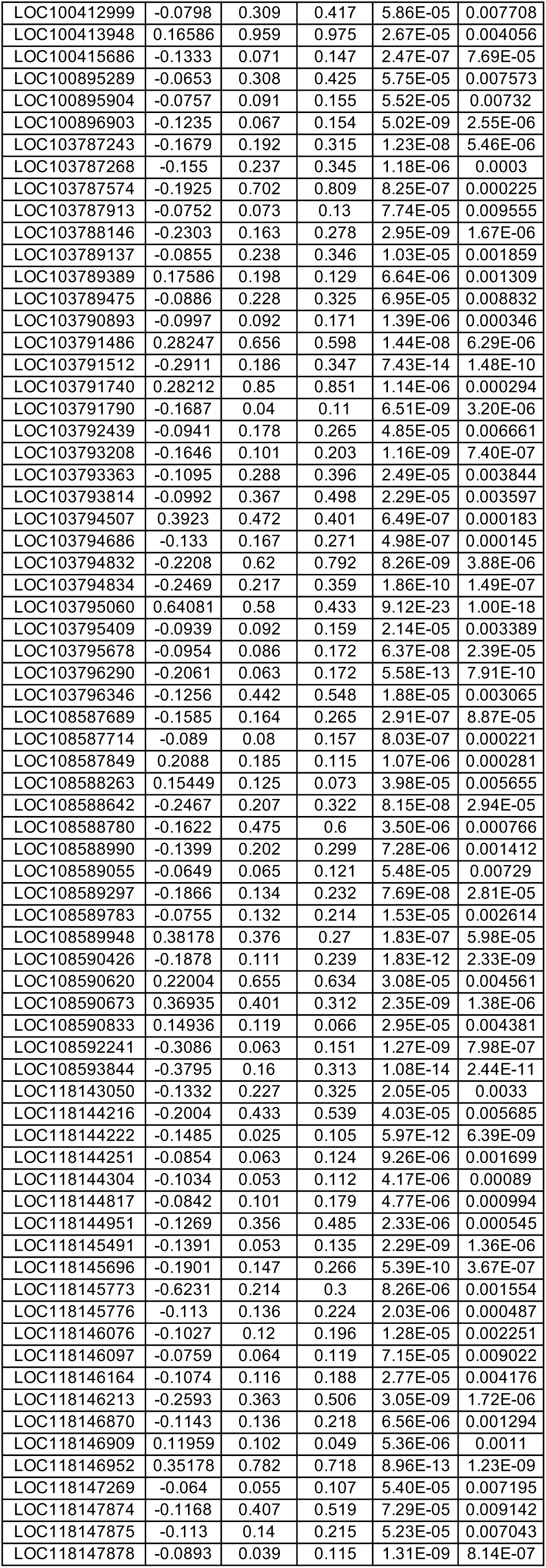
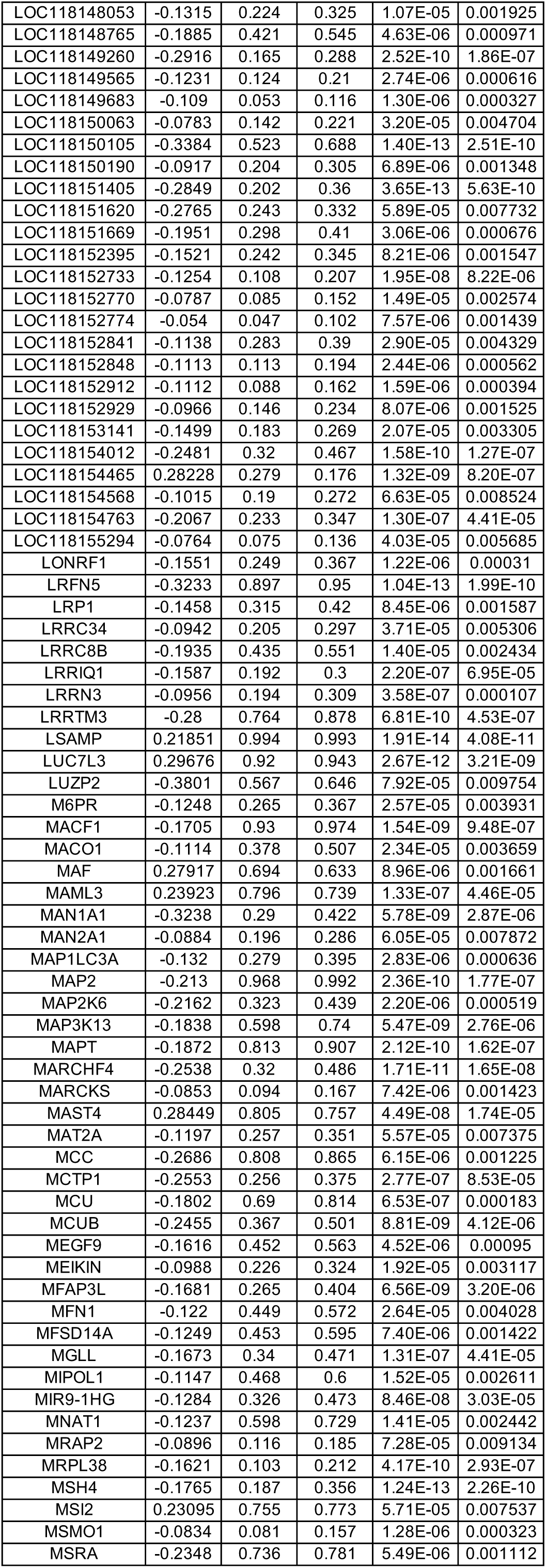
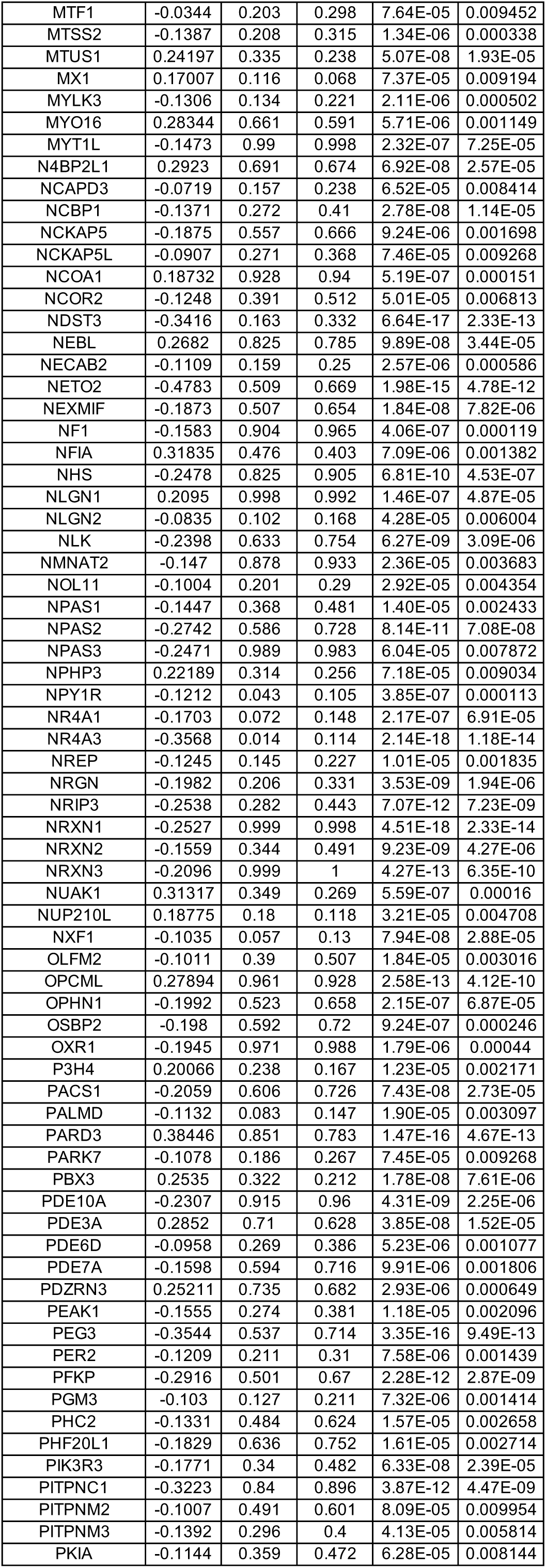
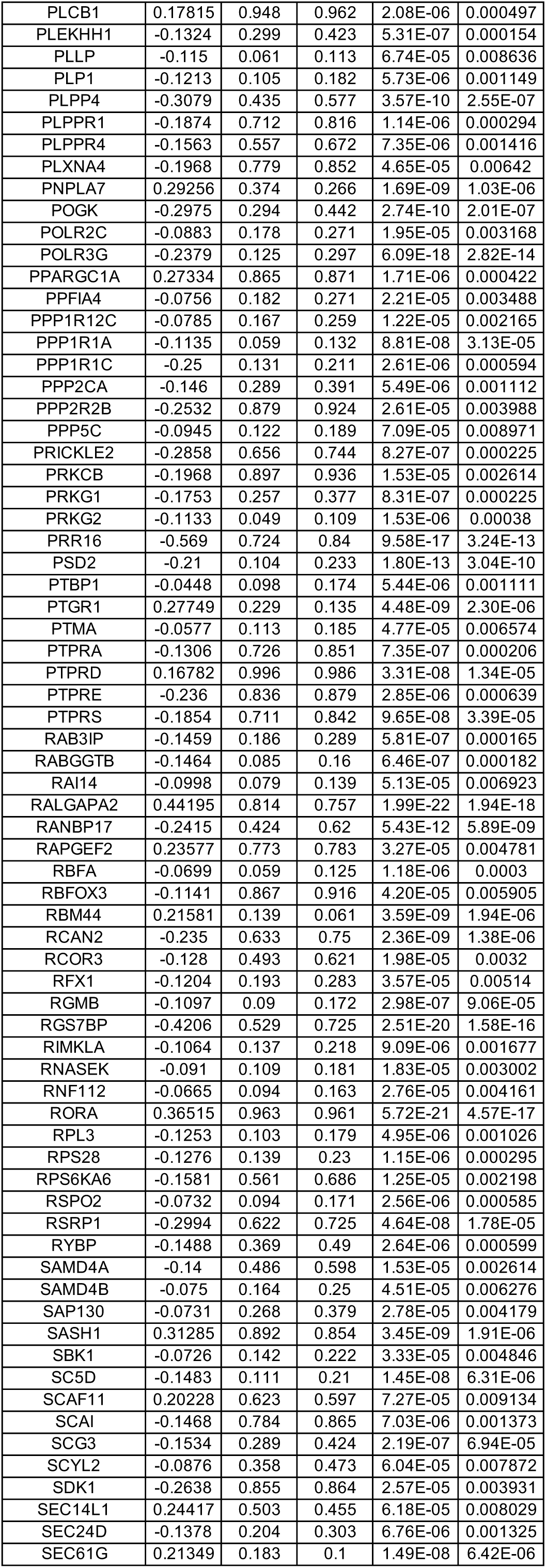
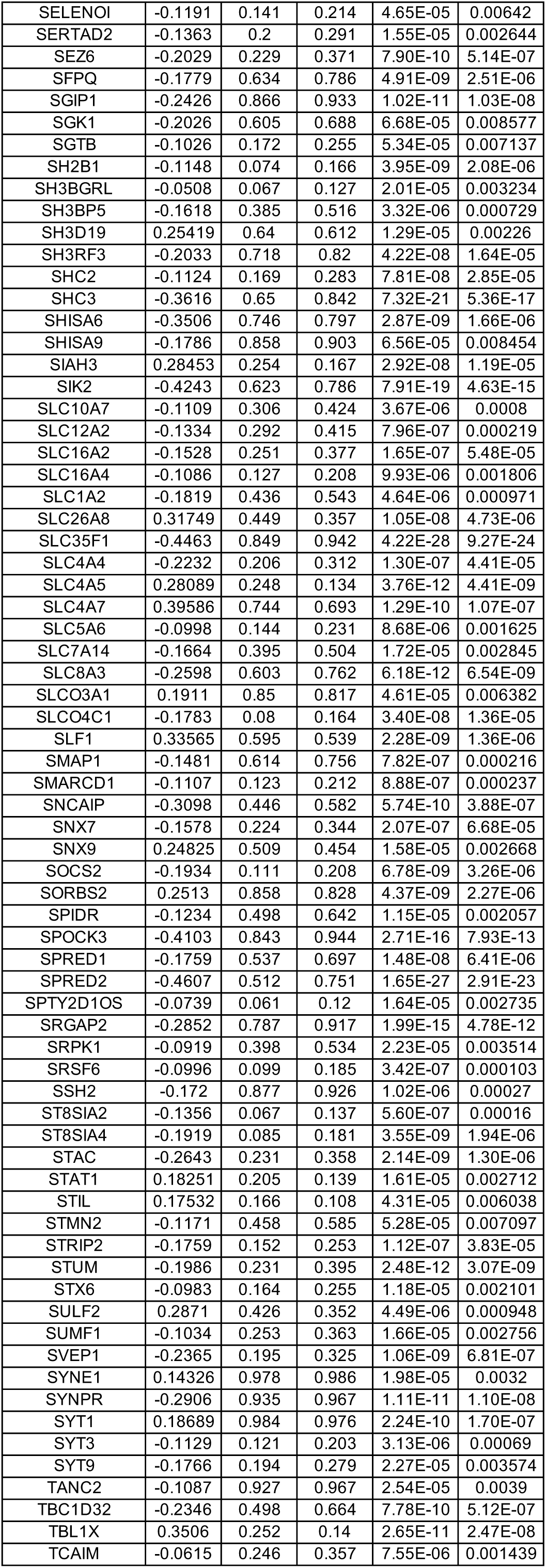
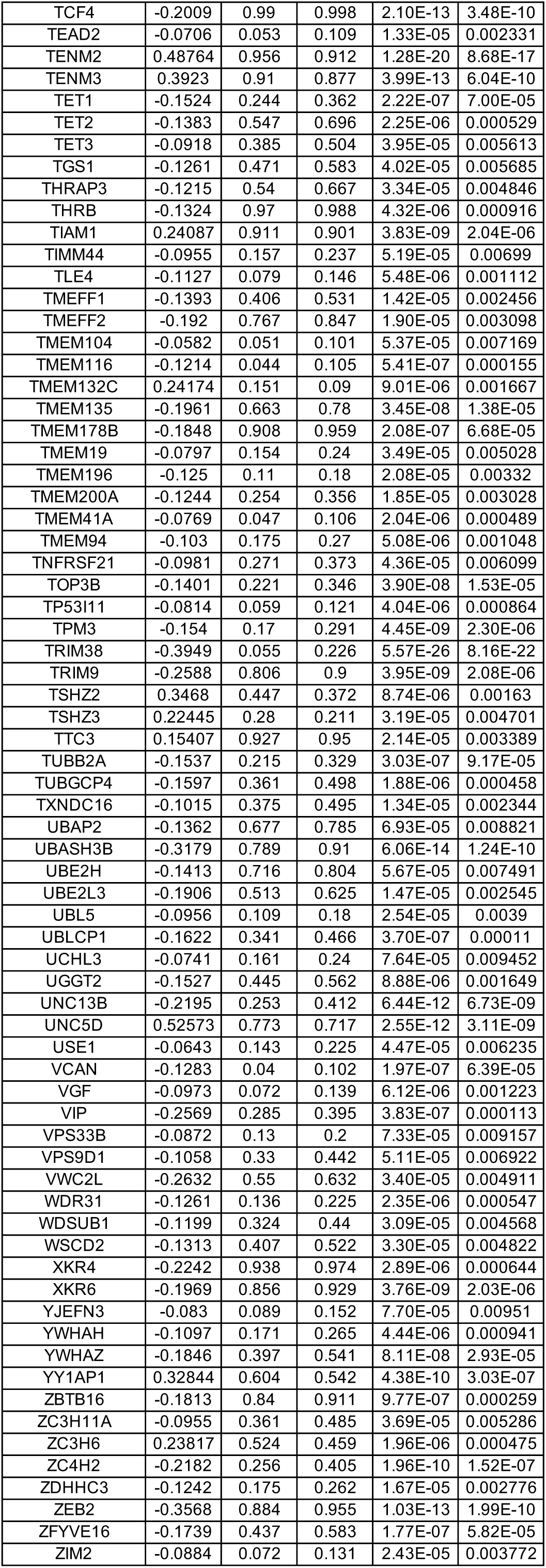
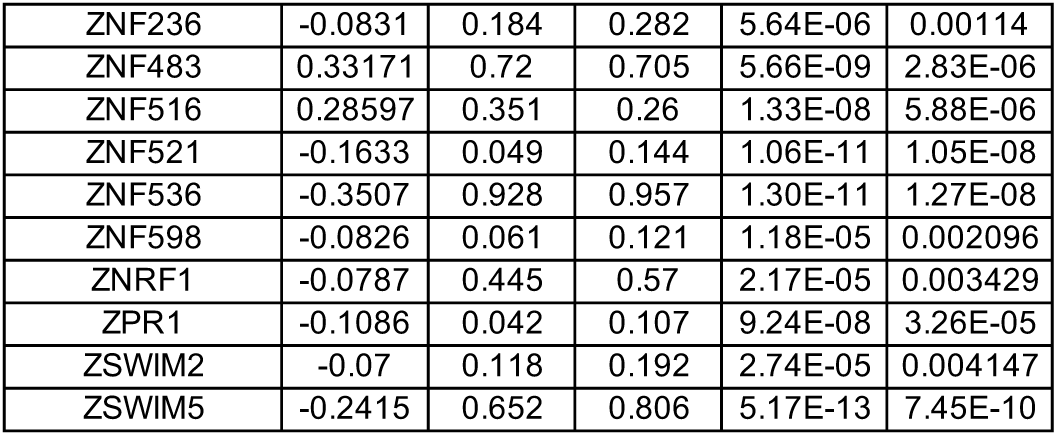
DEGs between wild-type (WT) and MECP2-null (KO) VIP-positive inhibitory neurons of PFC (adjusted p-value < 0.01)

**Table S32.**
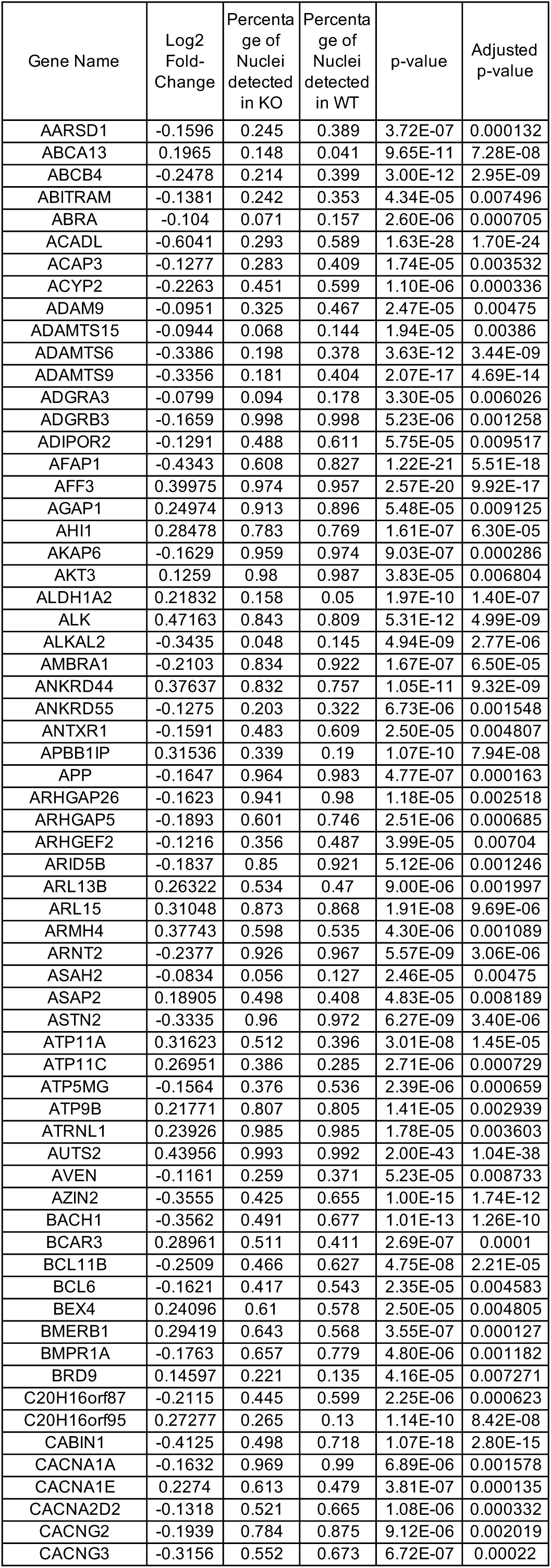
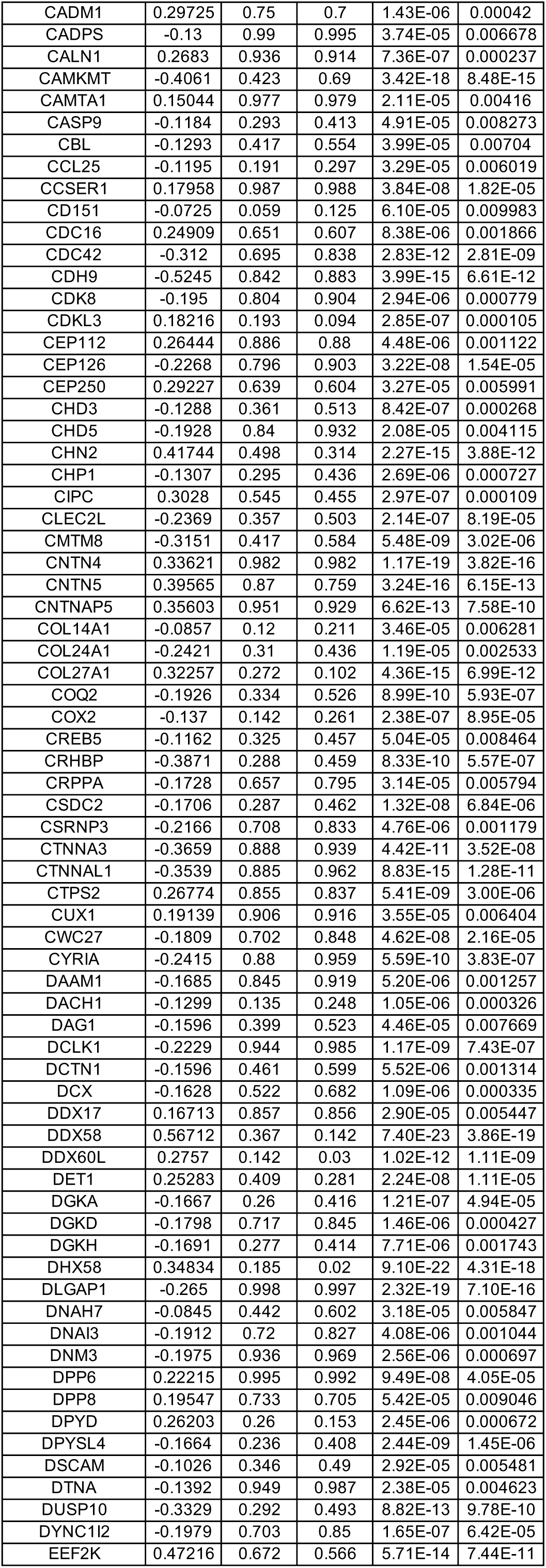
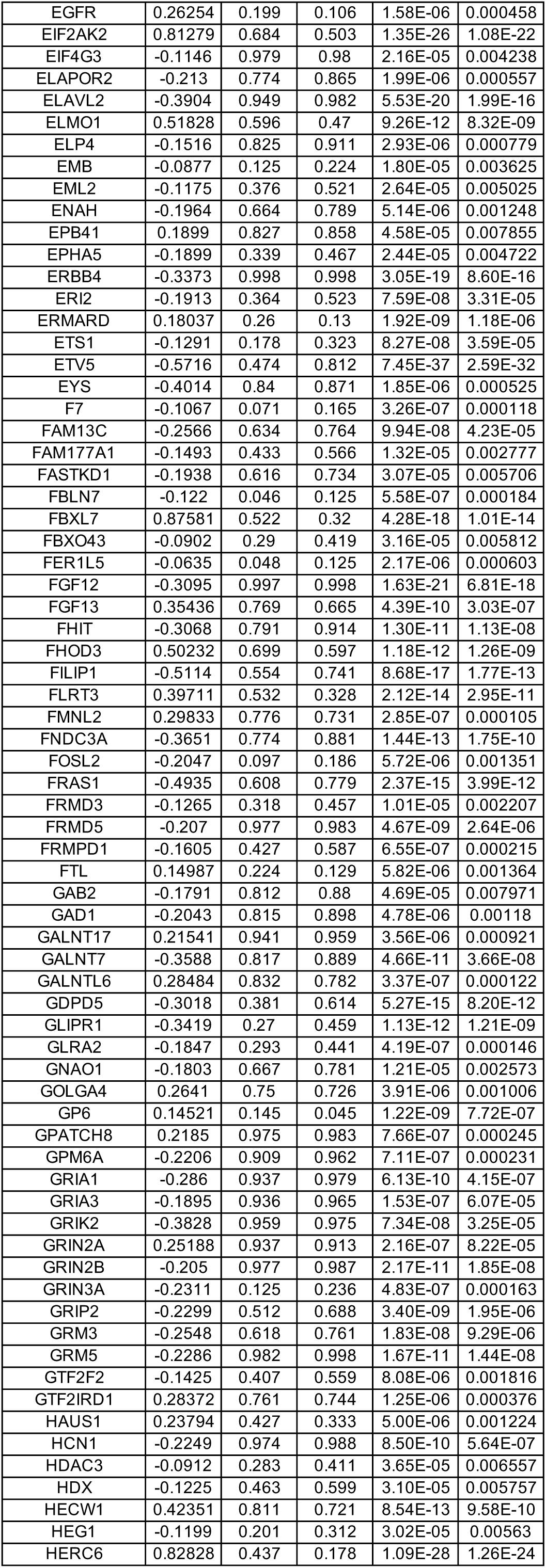
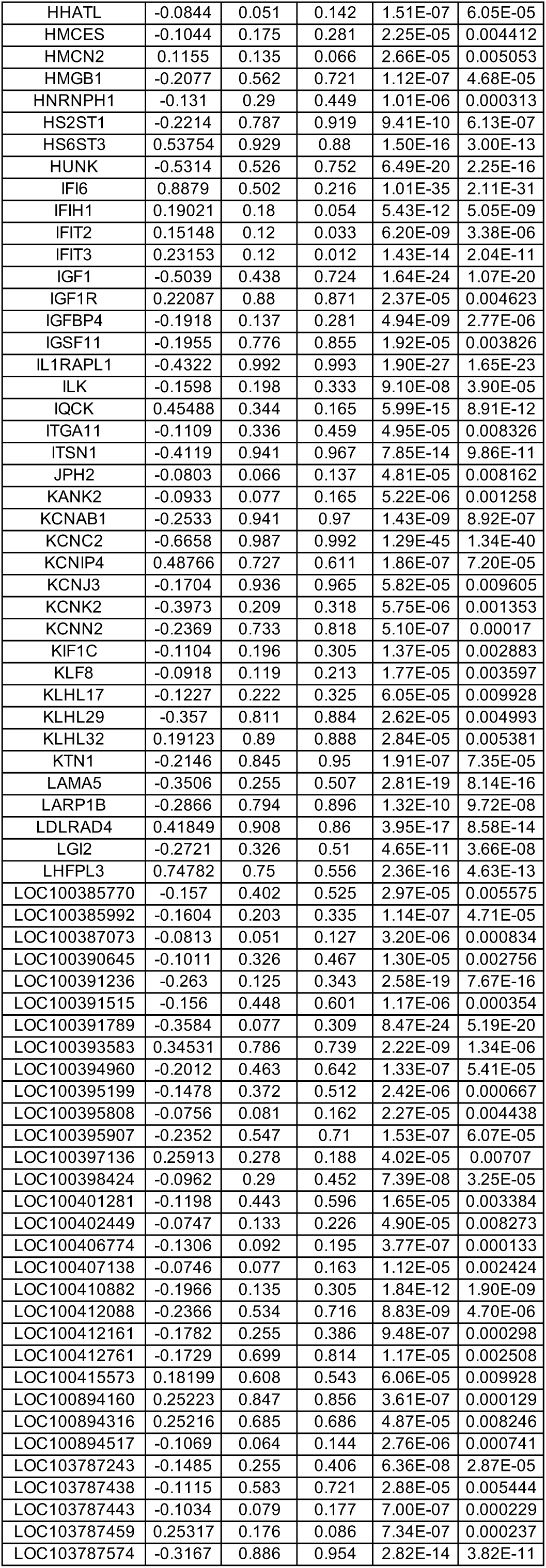
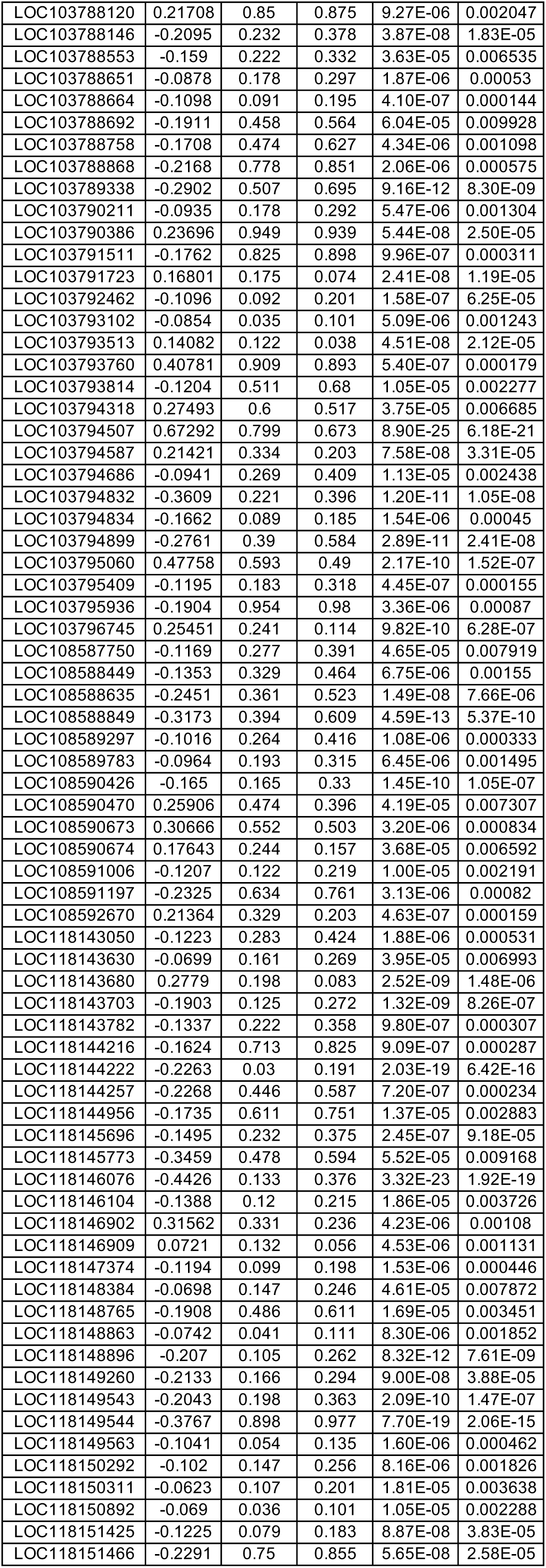
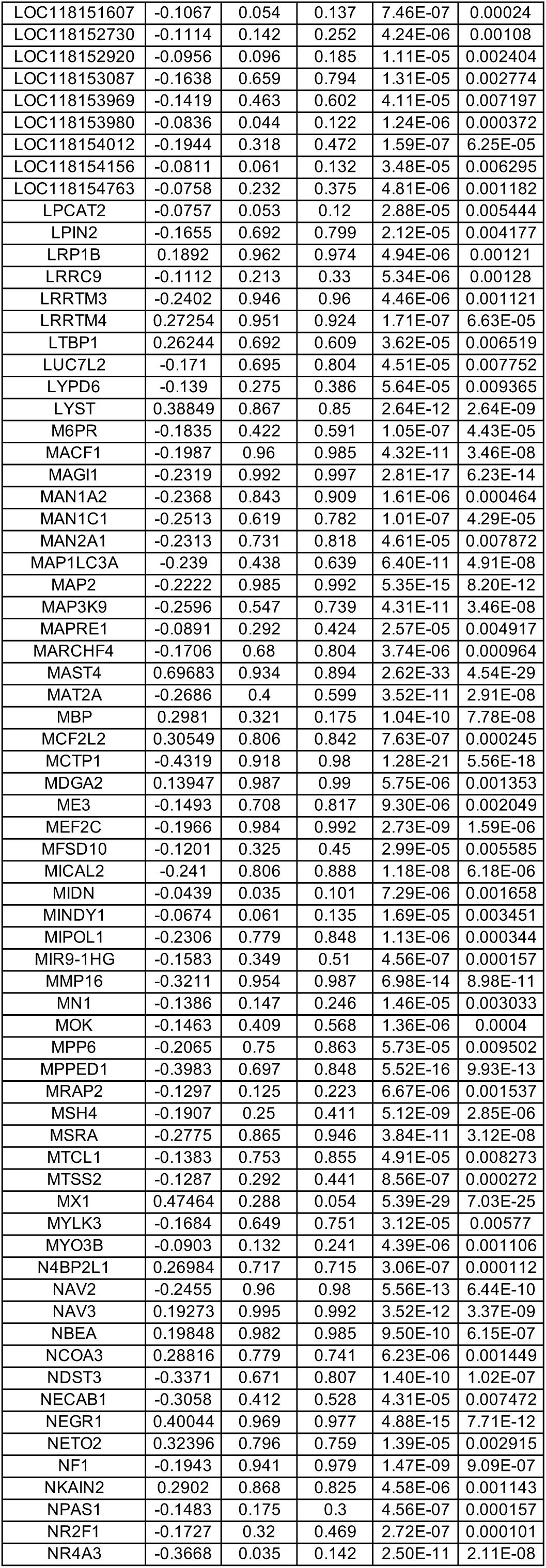
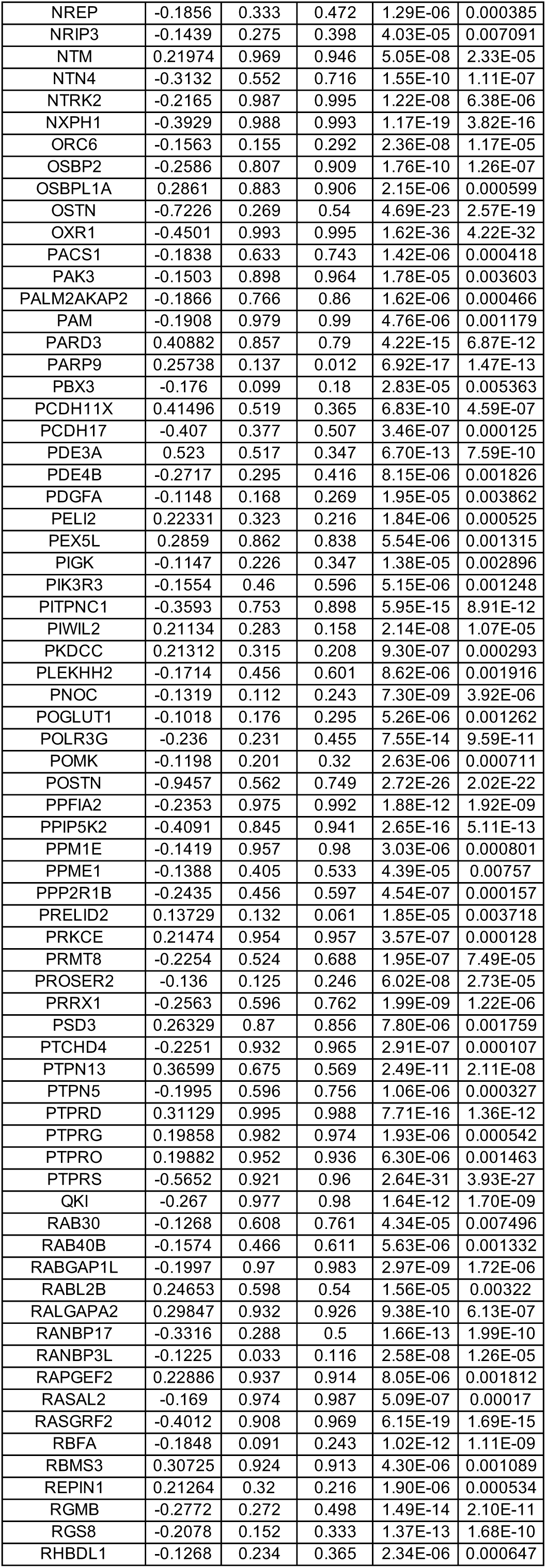
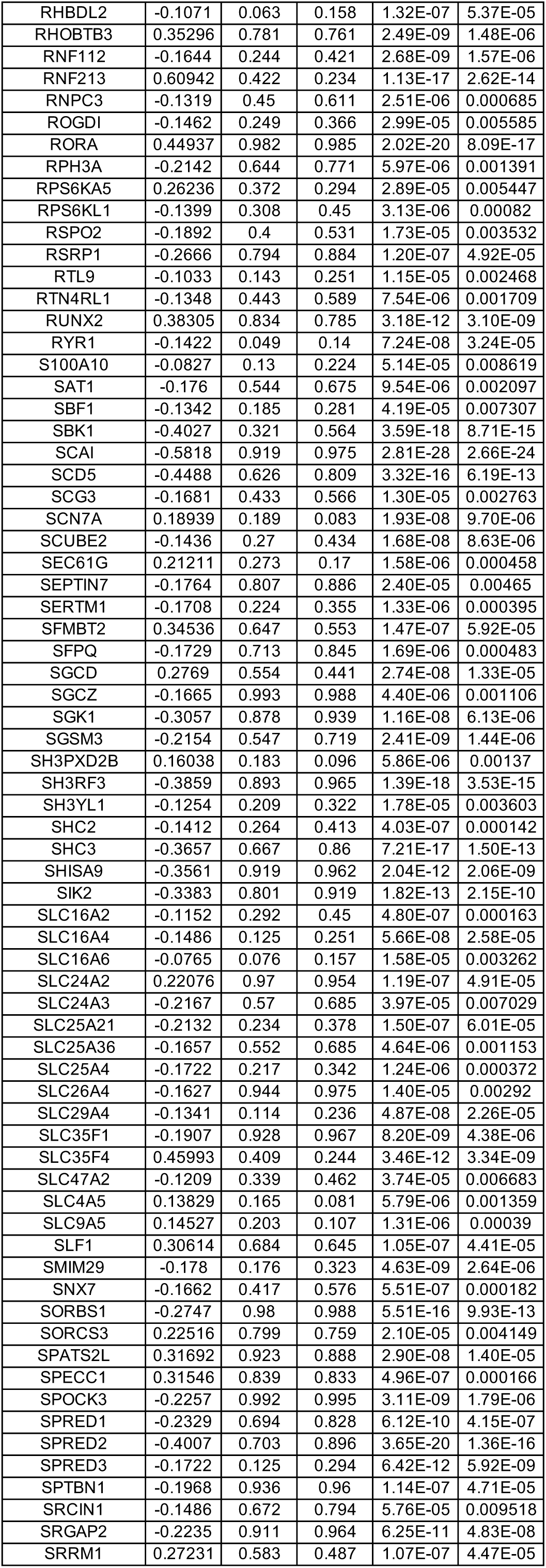
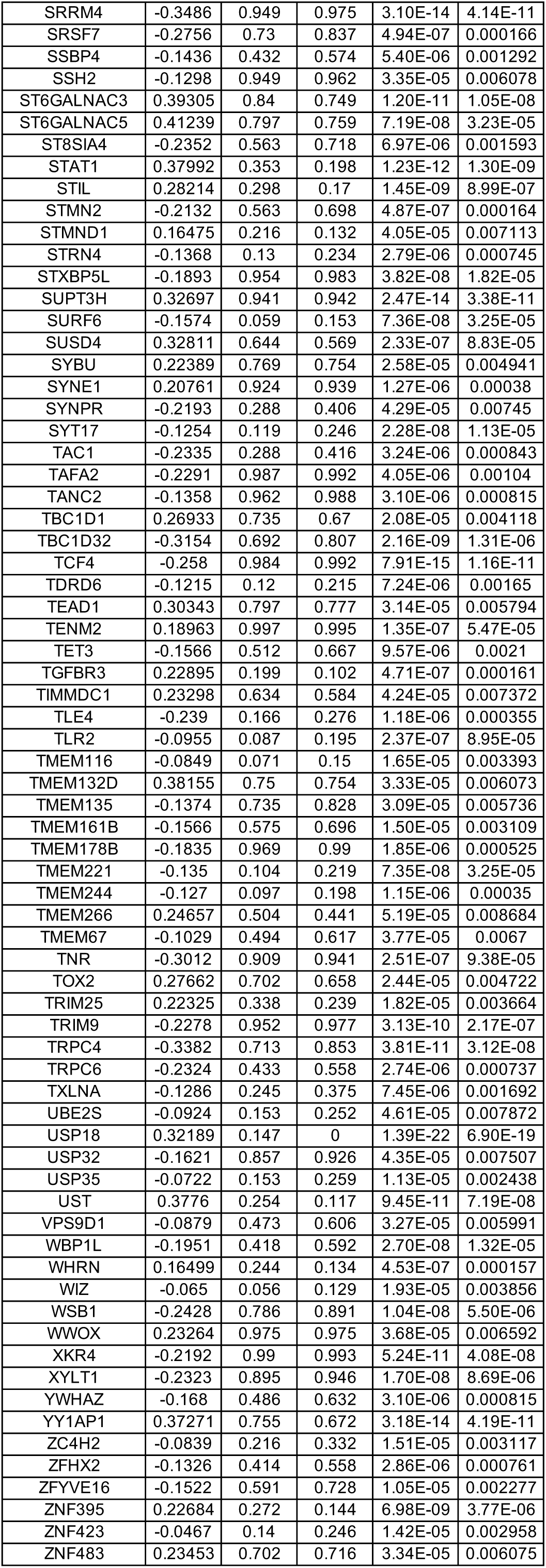
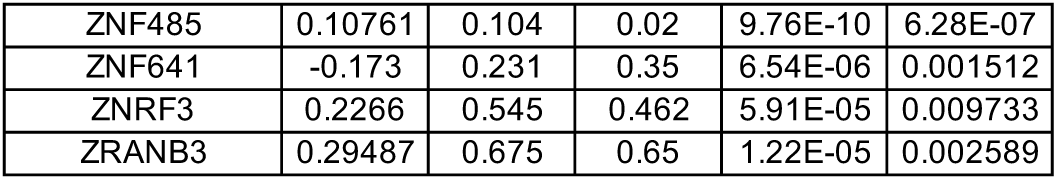
DEGs between wild-type (WT) and MECP2-null (KO) PVALB-positive inhibitory neurons of PFC (adjusted p-value < 0.01)

**Table S33.**
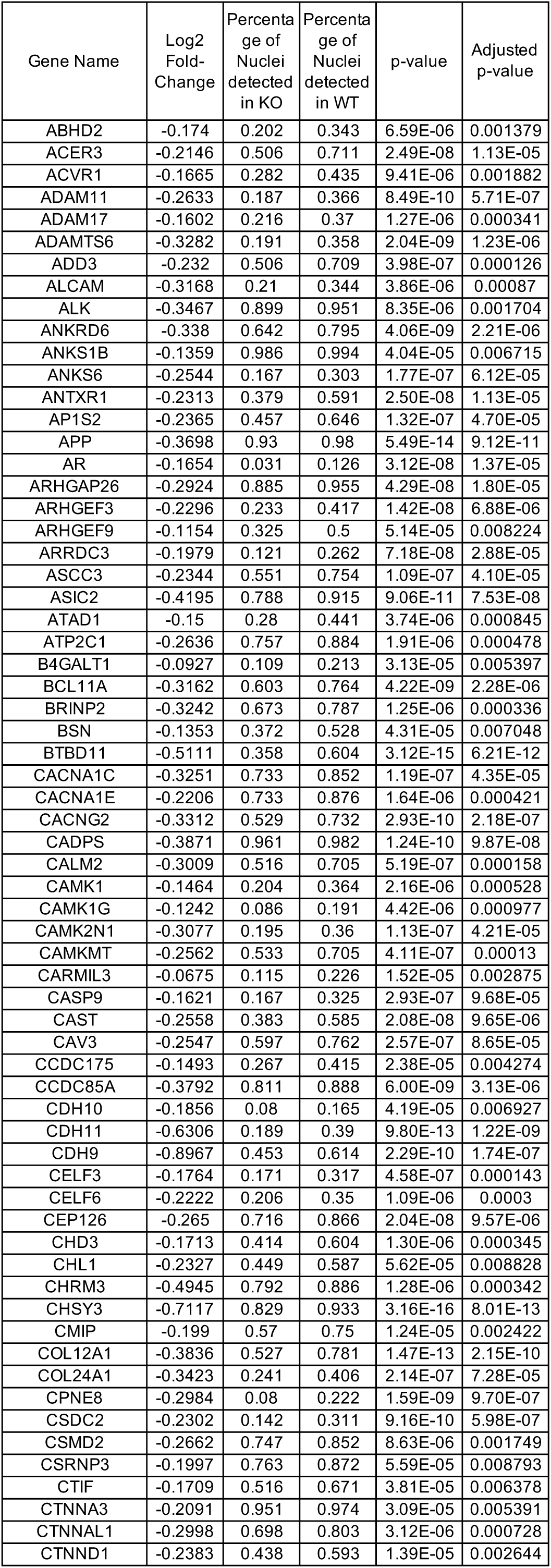
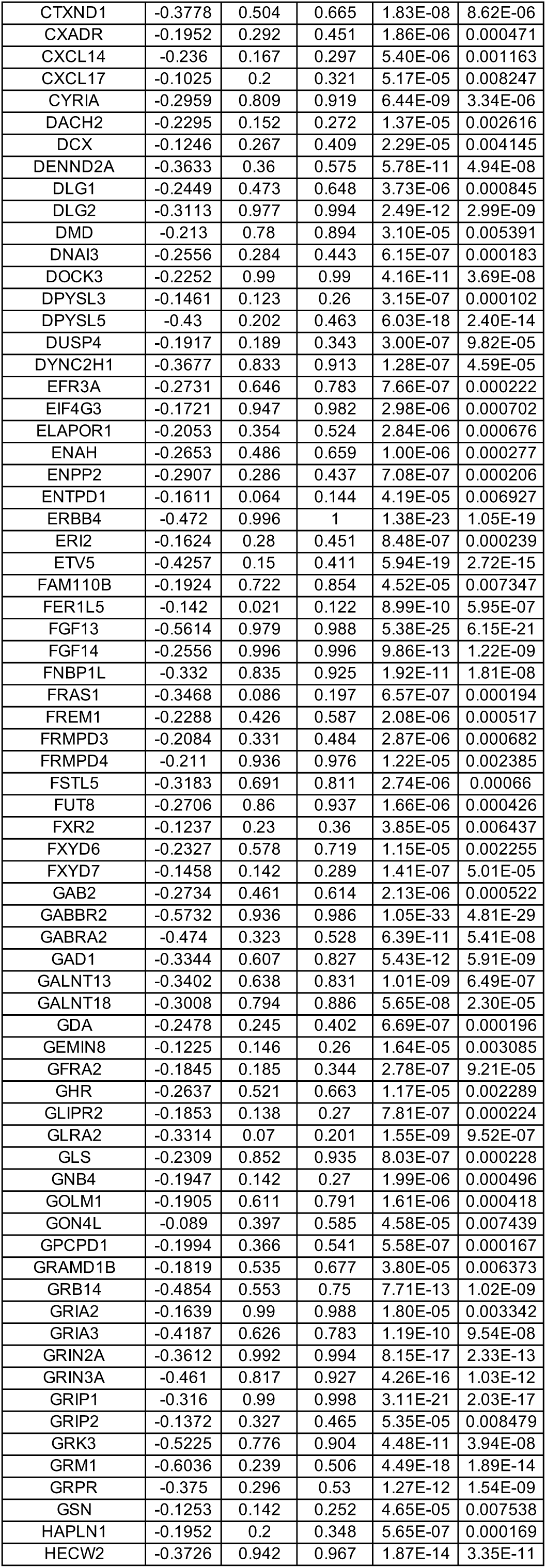
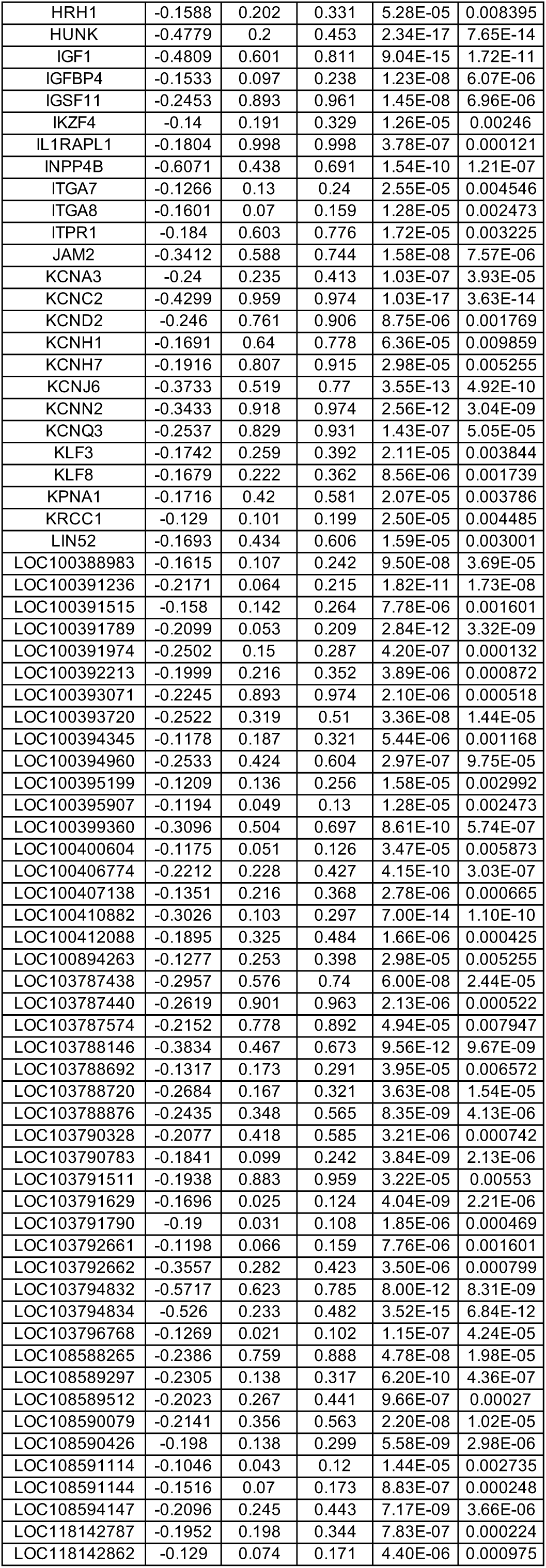
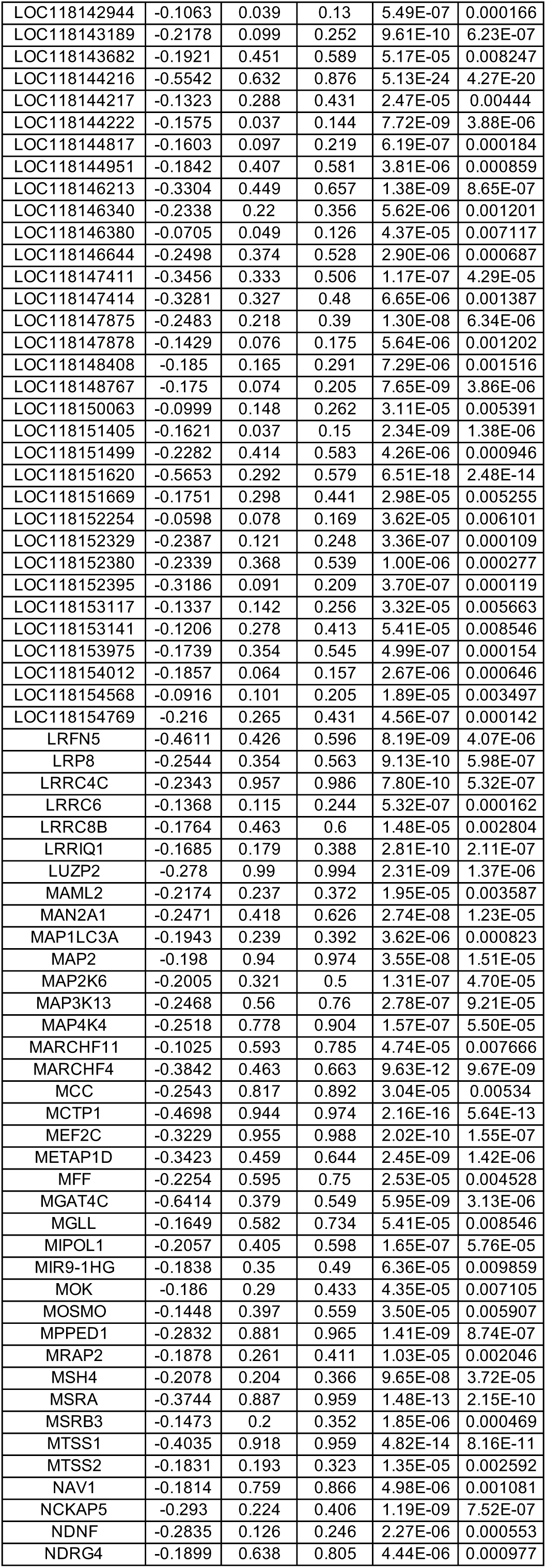
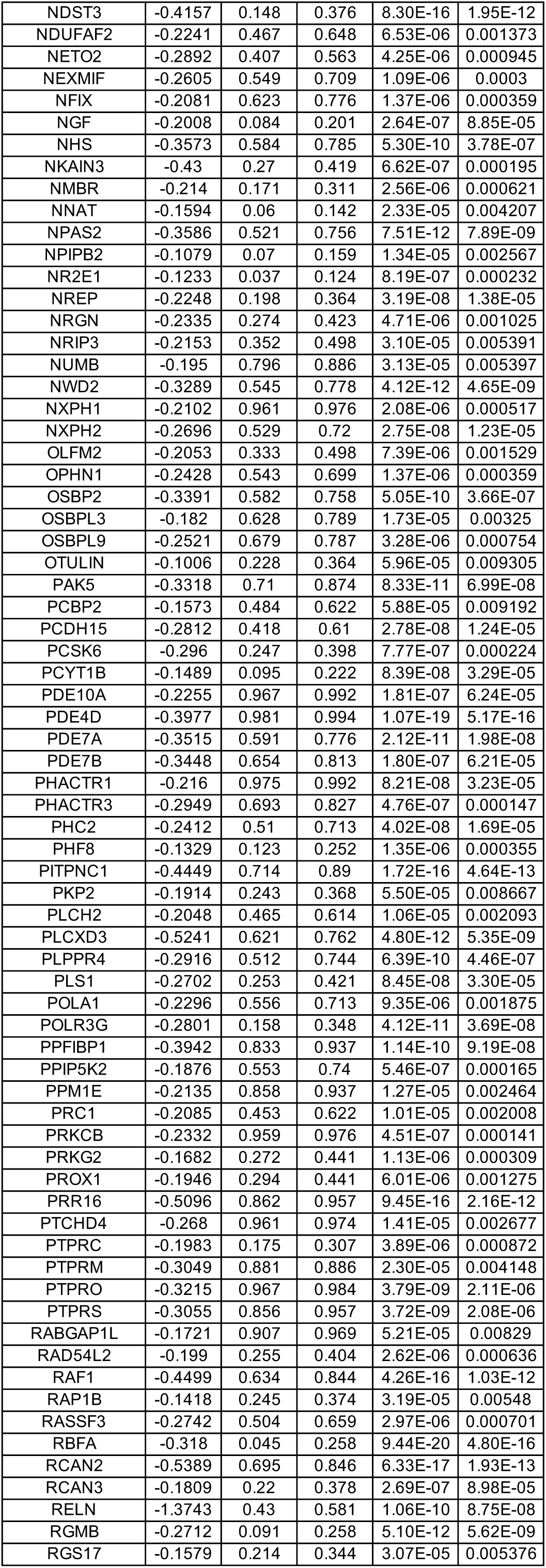
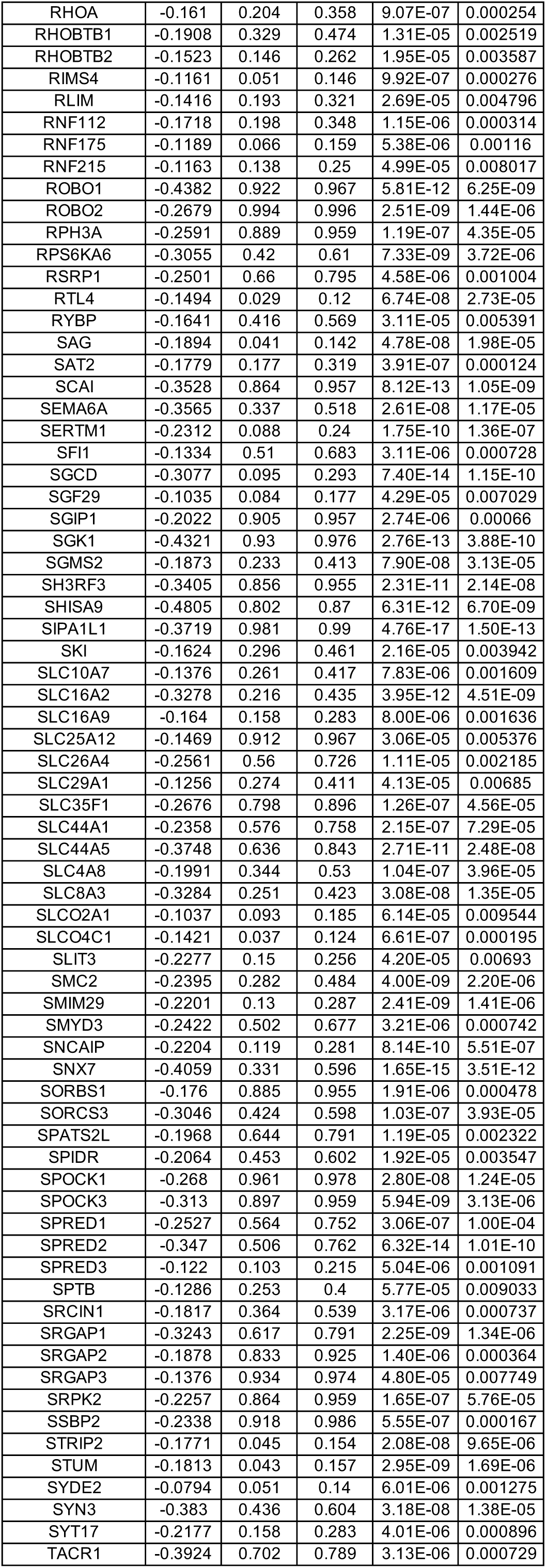
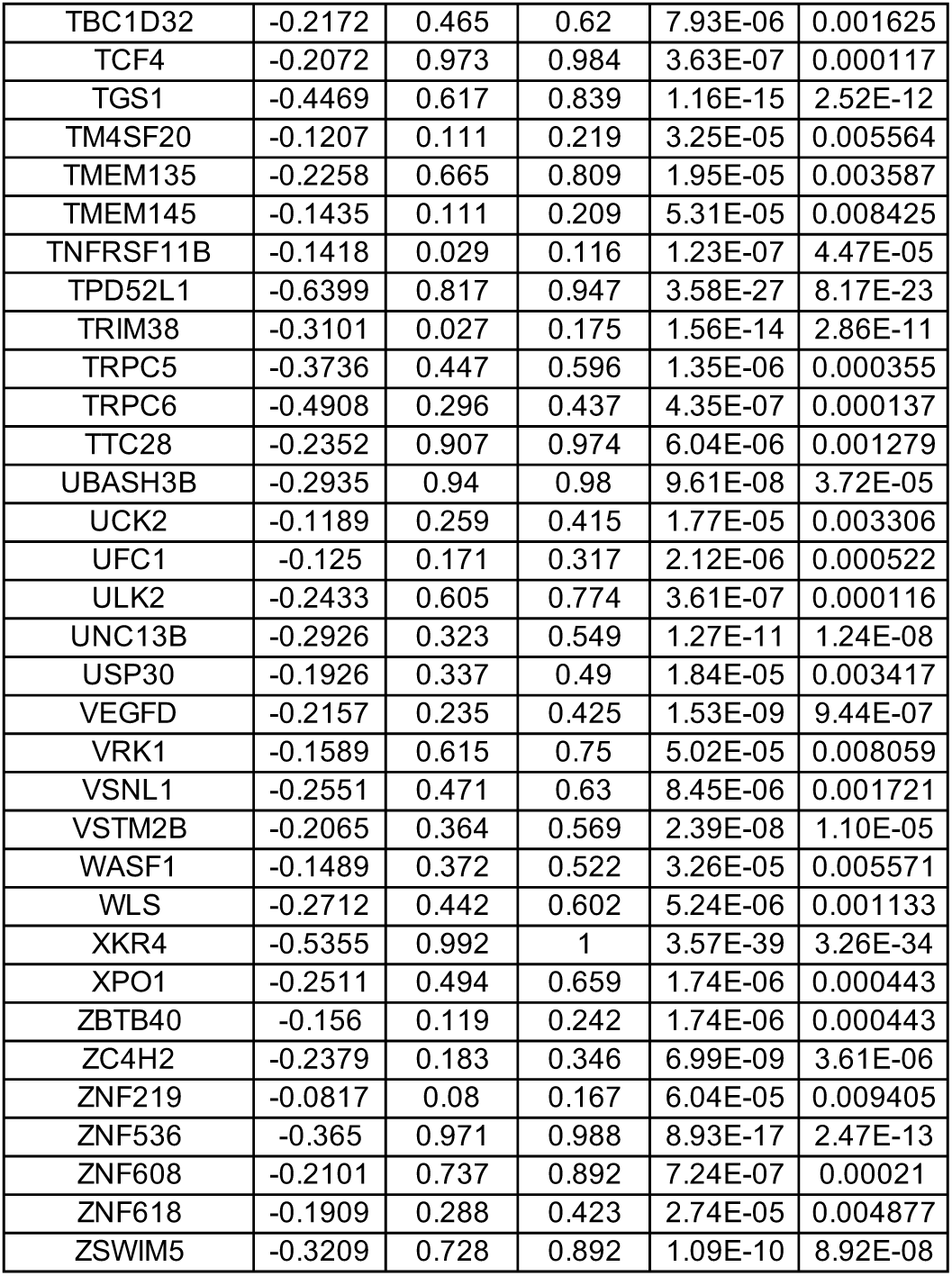
DEGs between wild-type (WT) and MECP2-null (KO) LAMP5-positive inhibitory neurons of PFC (adjusted p-value < 0.01)

**Table S34.**
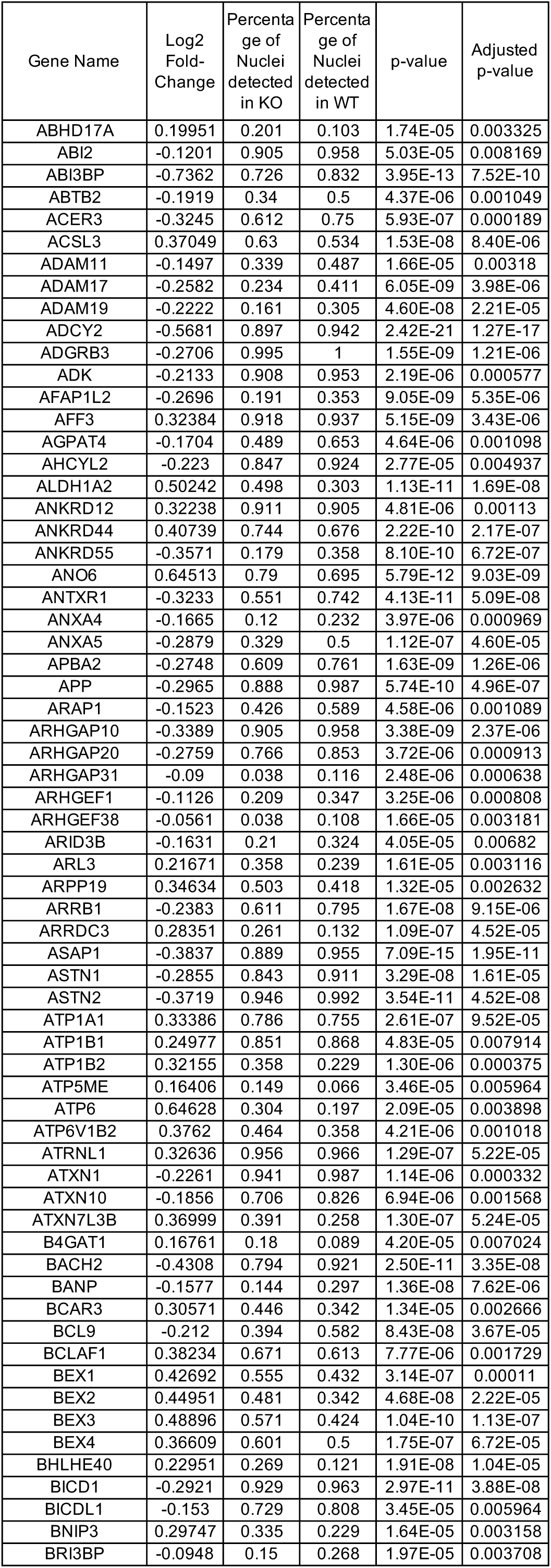
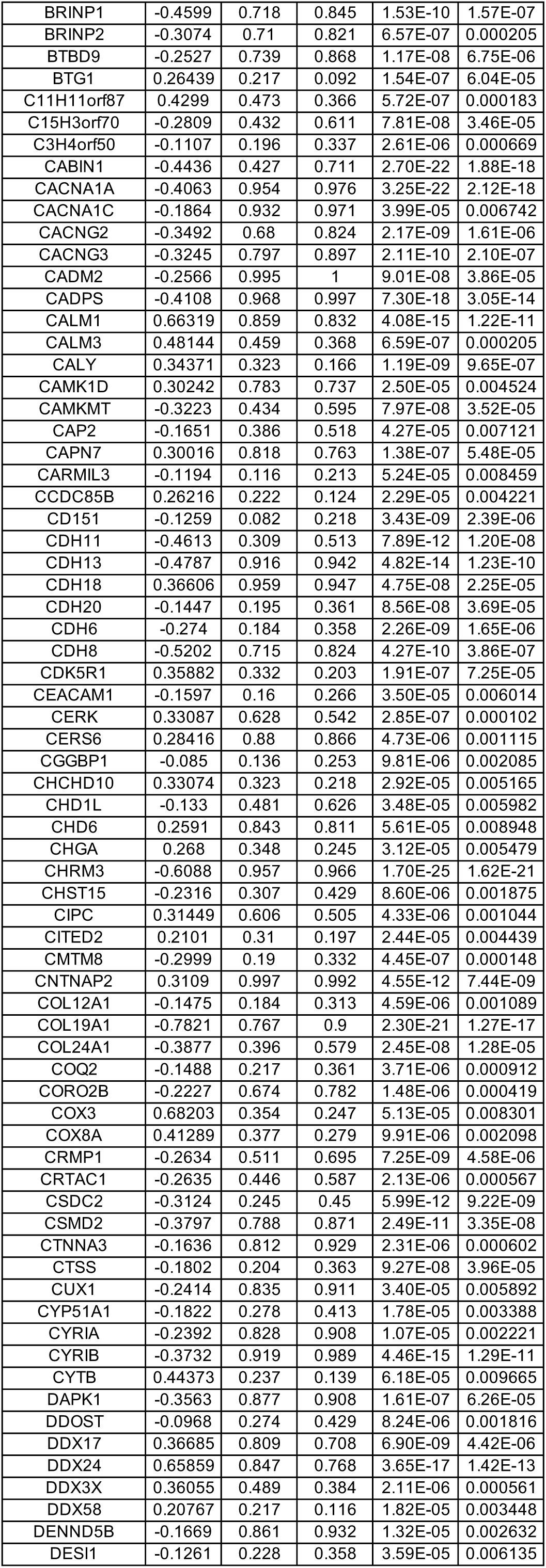
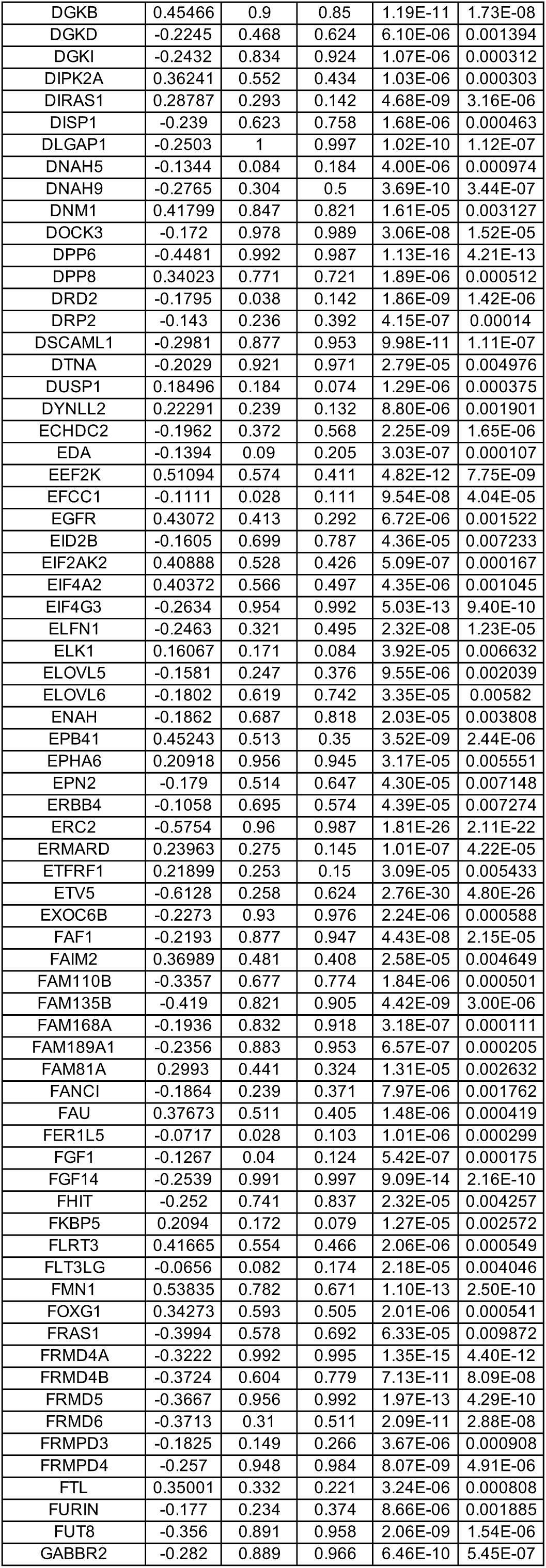
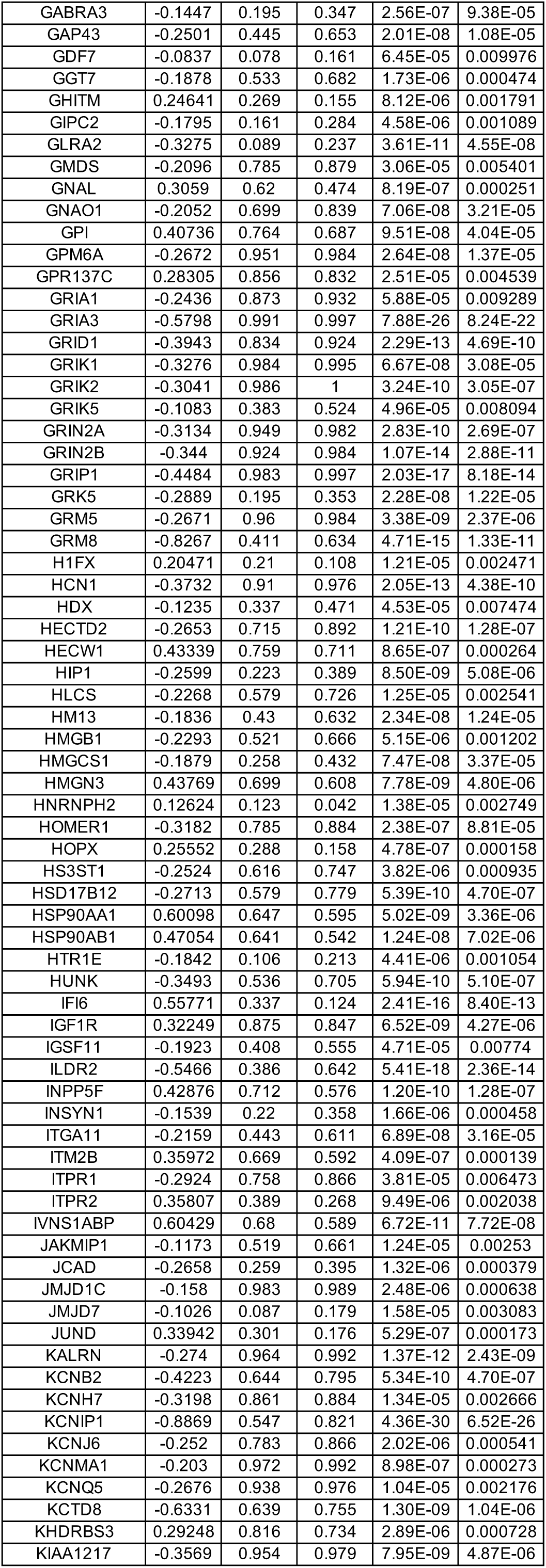
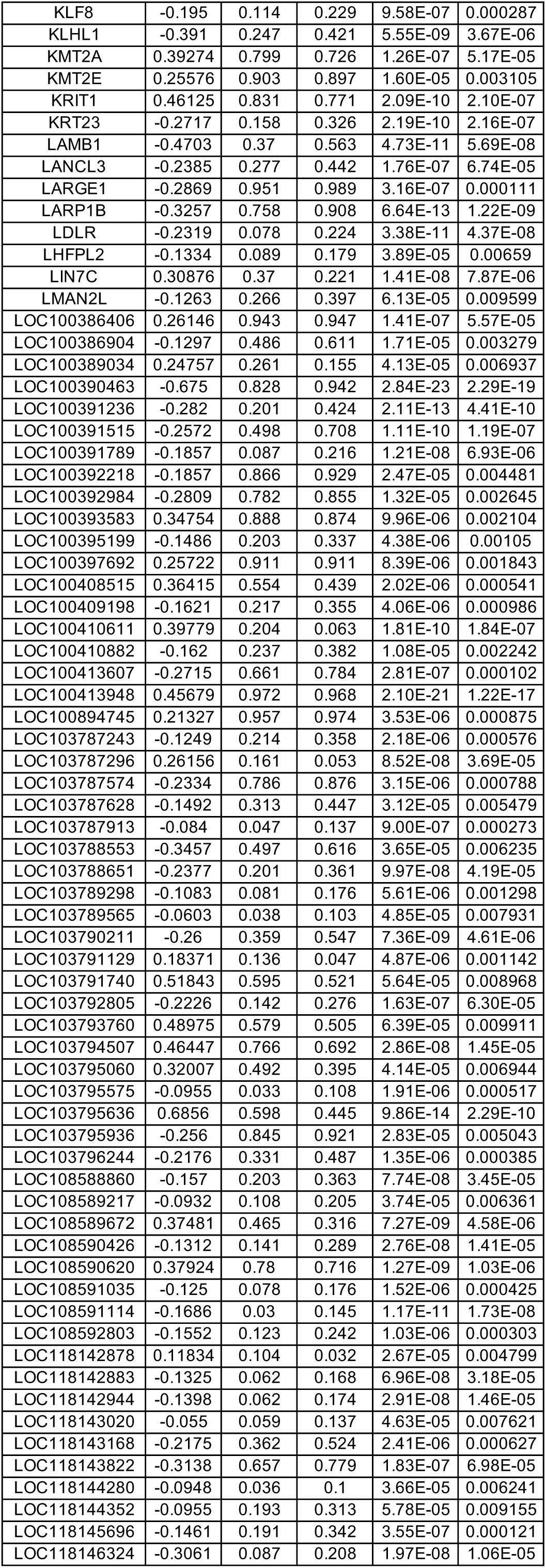
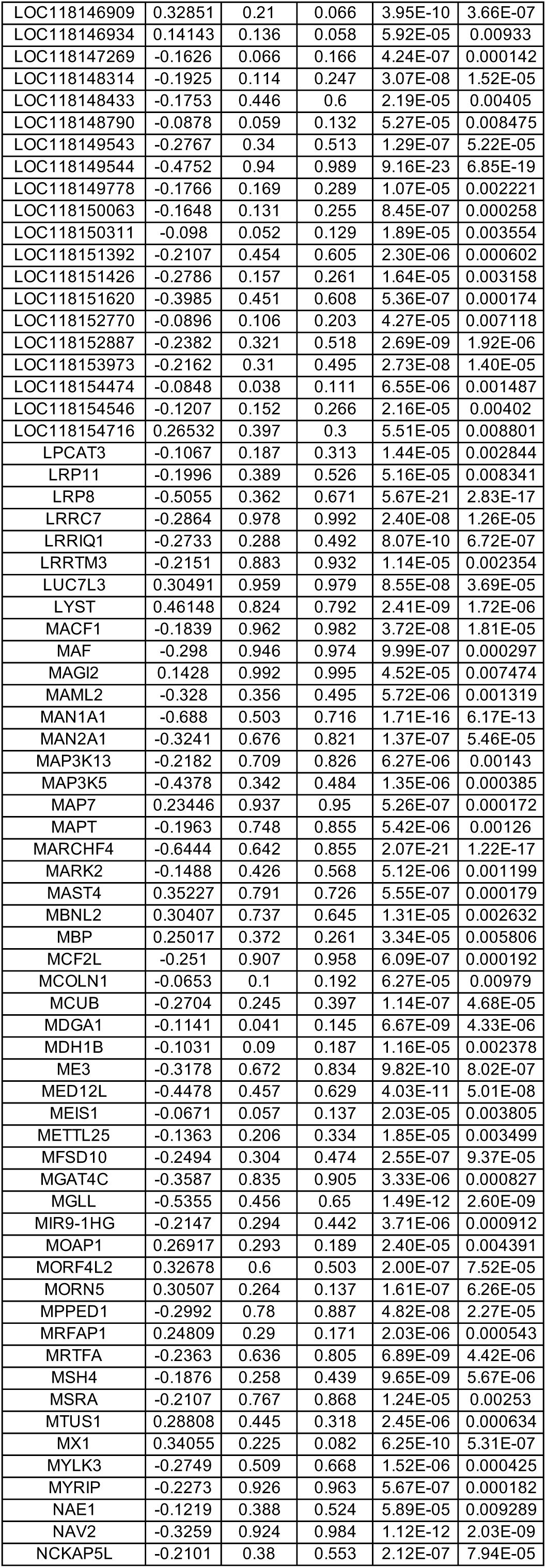
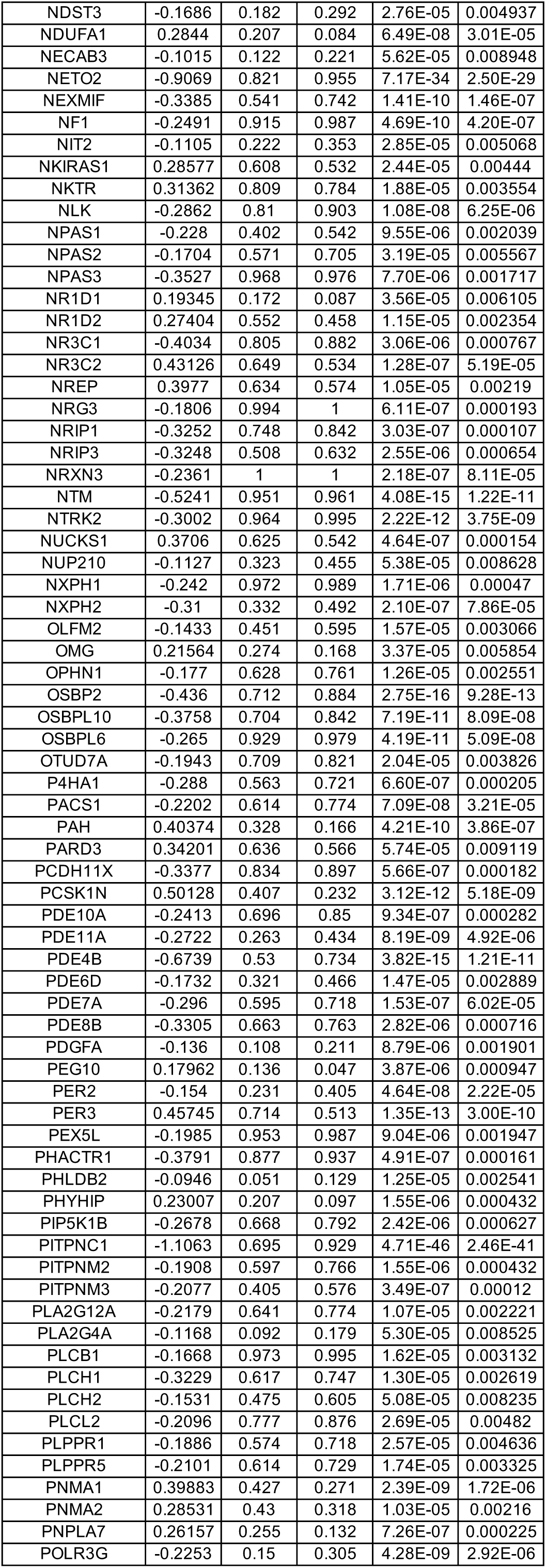
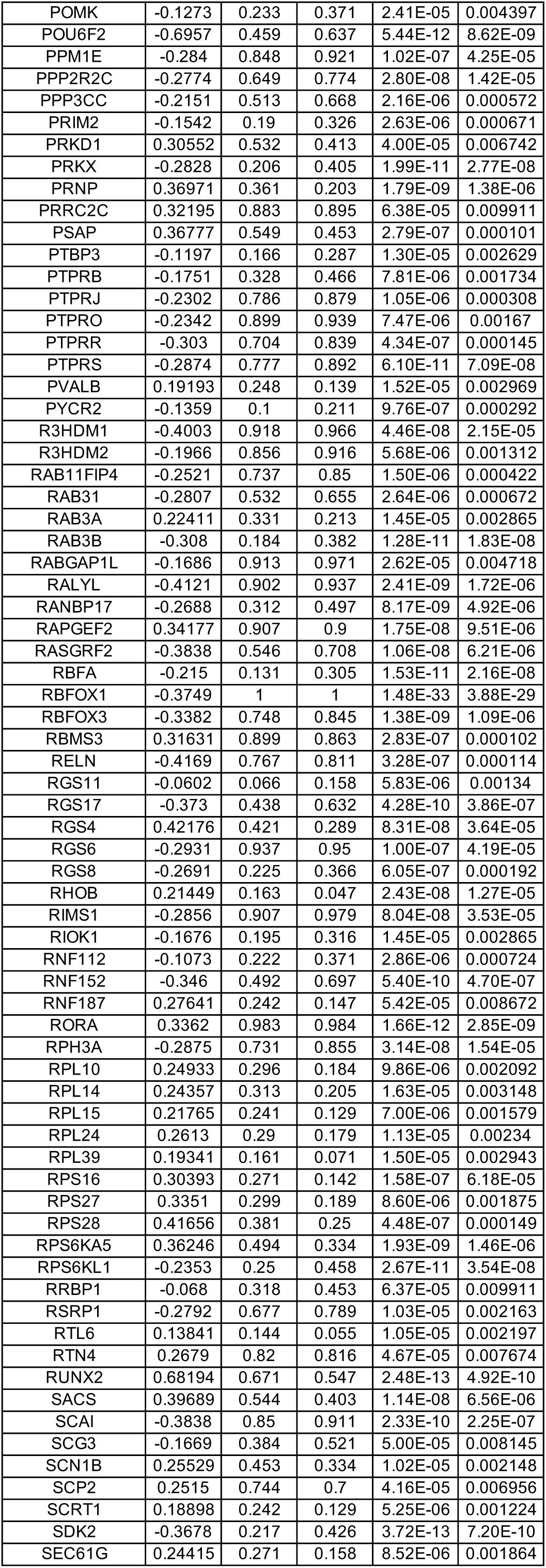
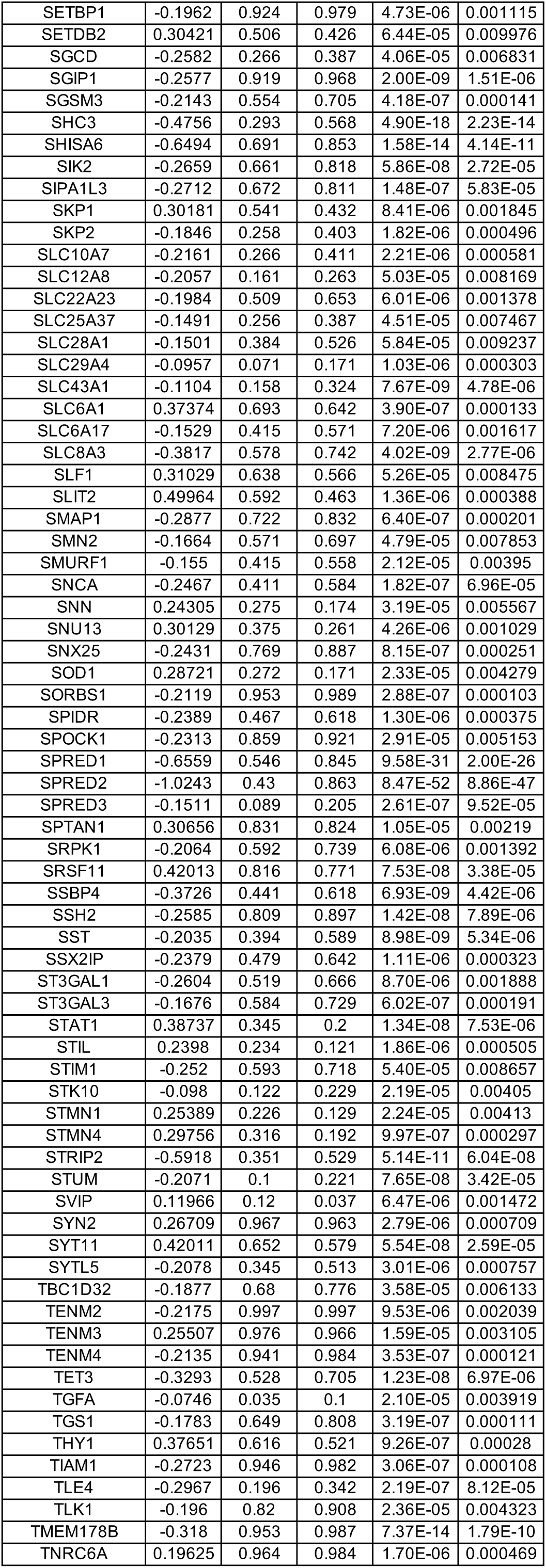
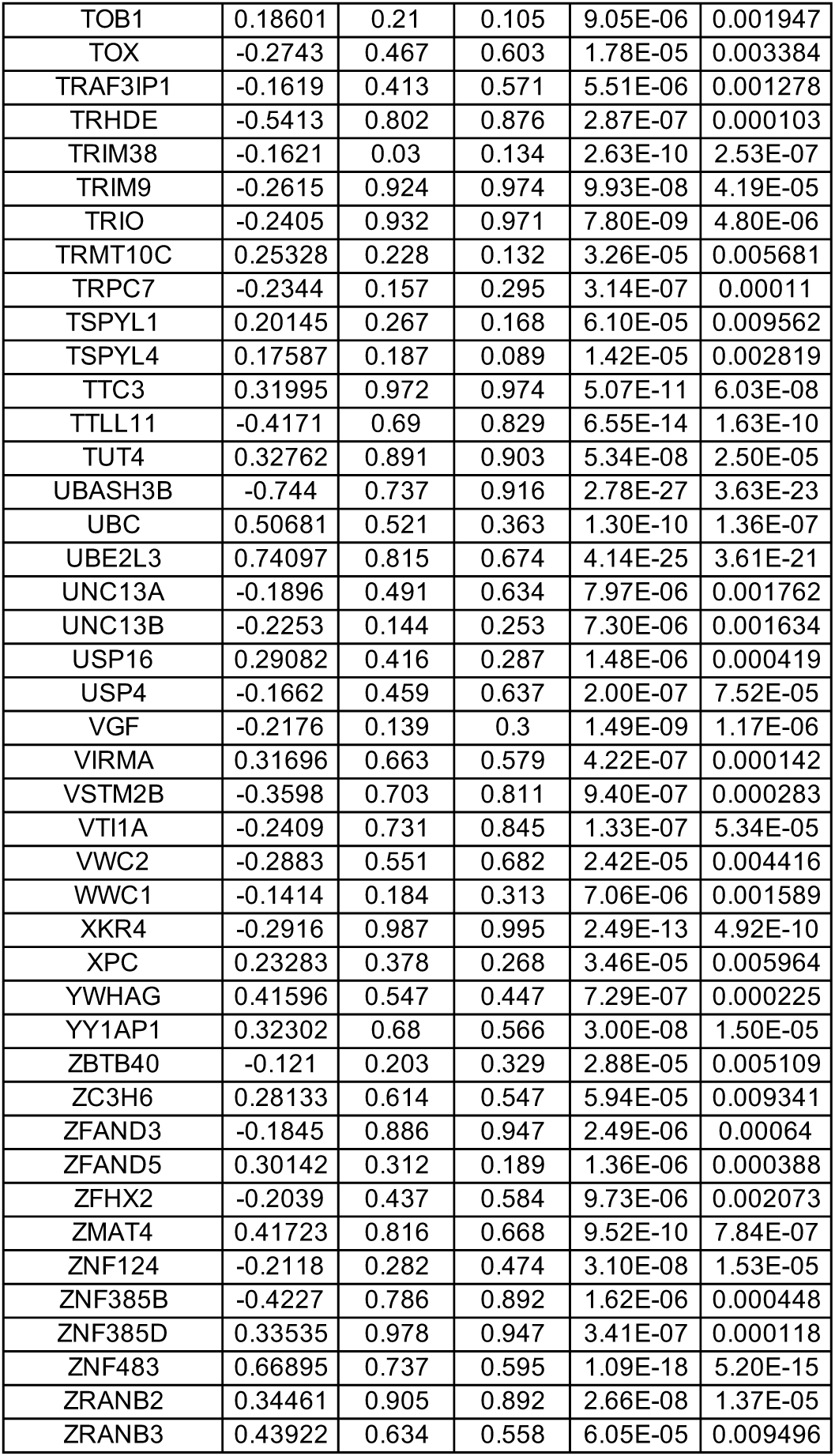
DEGs between wild-type (WT) and MECP2-null (KO) SST-positive inhibitory neurons of OC (adjusted p-value < 0.01)

**Table S35.**
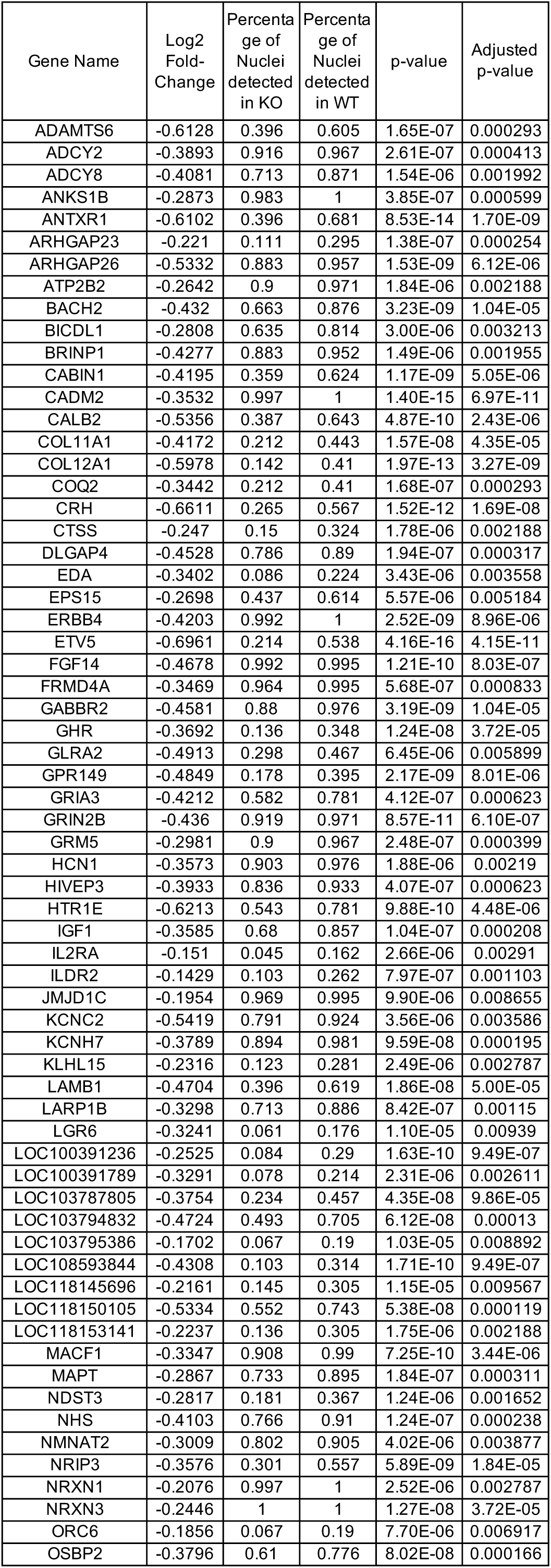
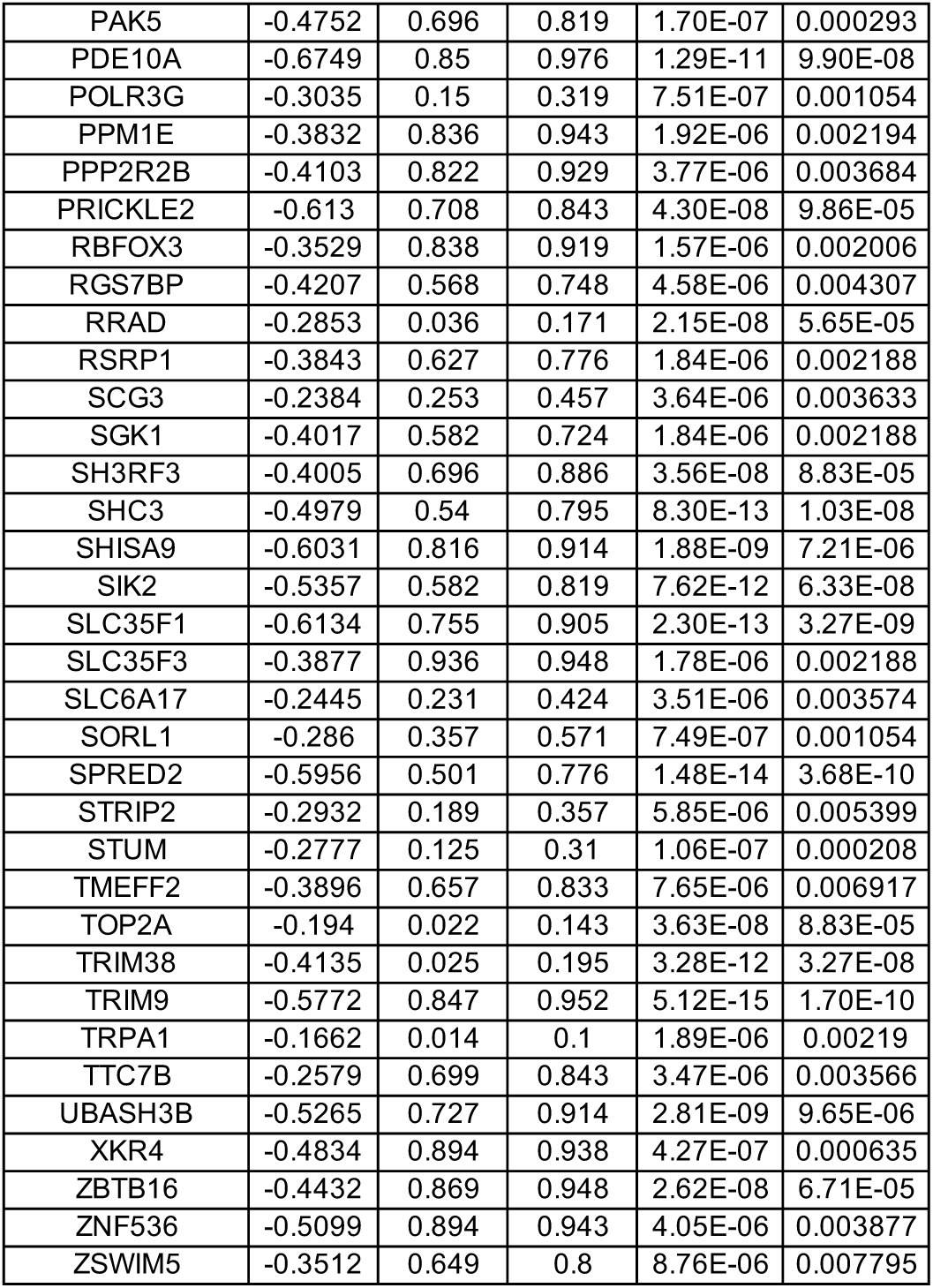
DEGs between wild-type (WT) and MECP2-null (KO) VIP-positive inhibitory neurons of OC (adjusted p-value < 0.01)

**Table S36.**
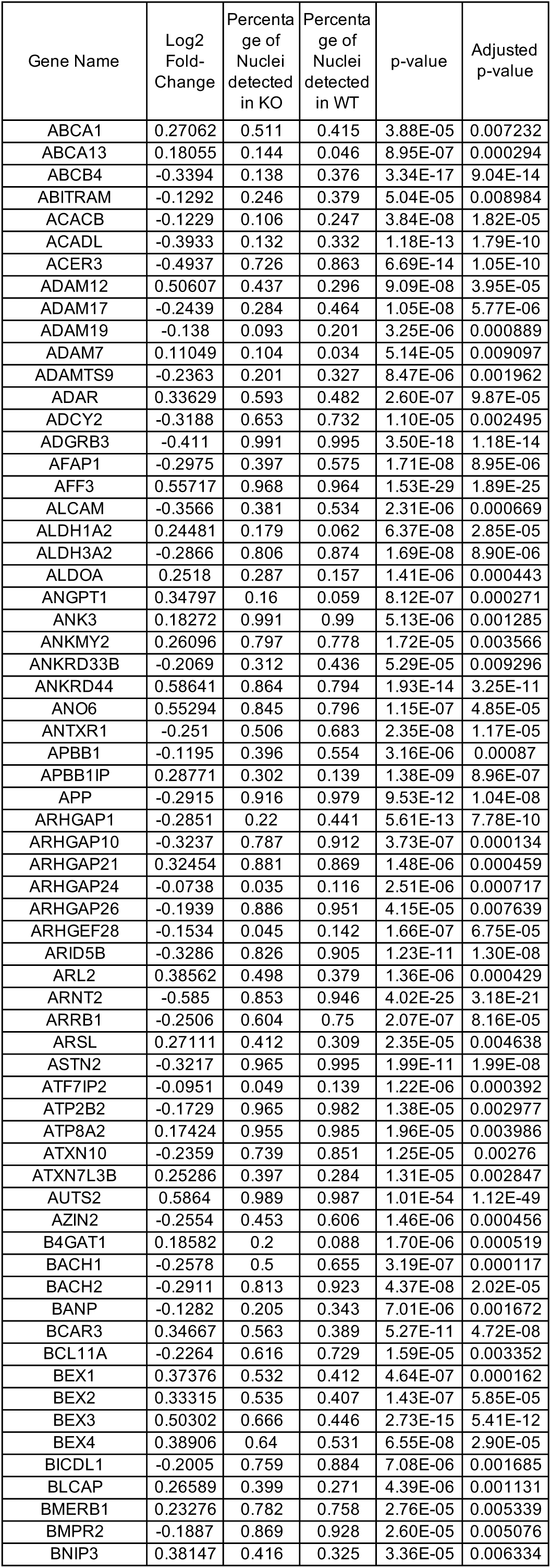
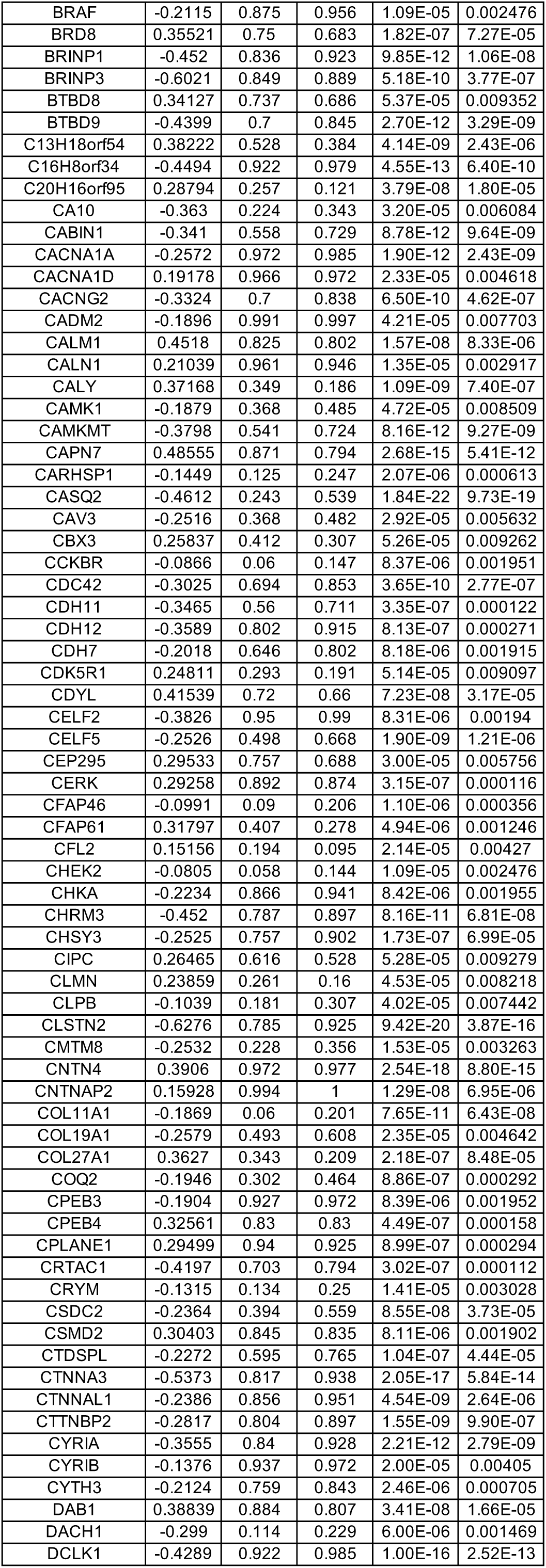
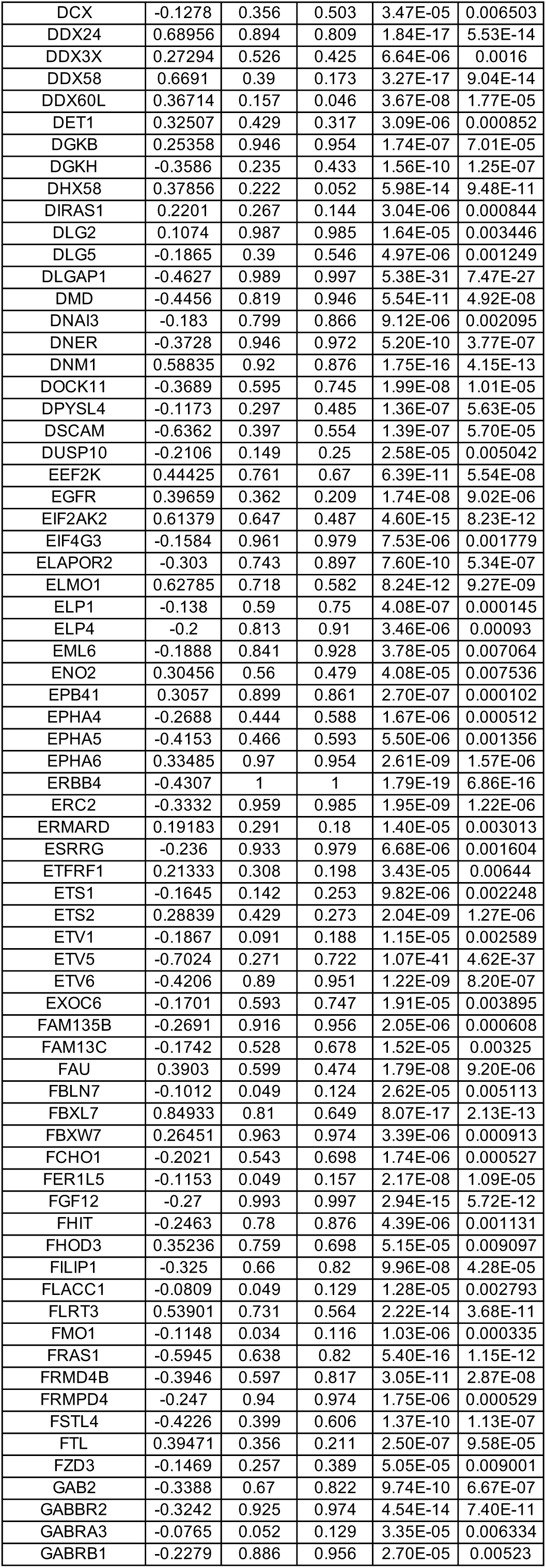
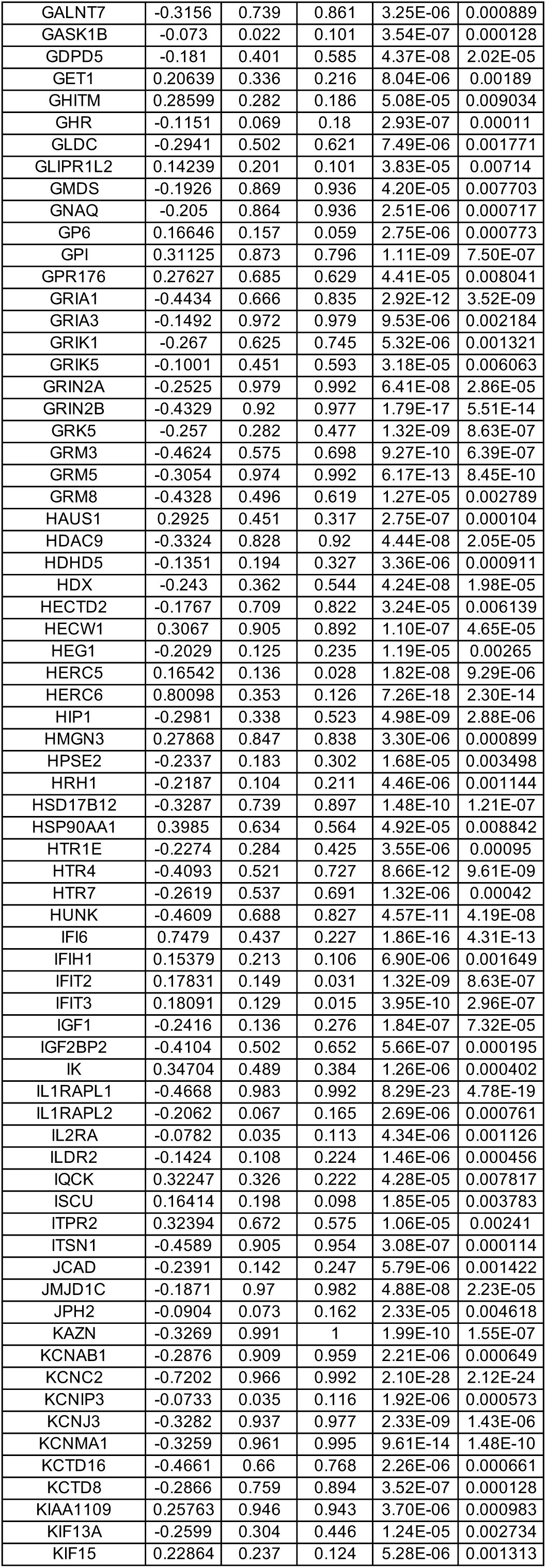
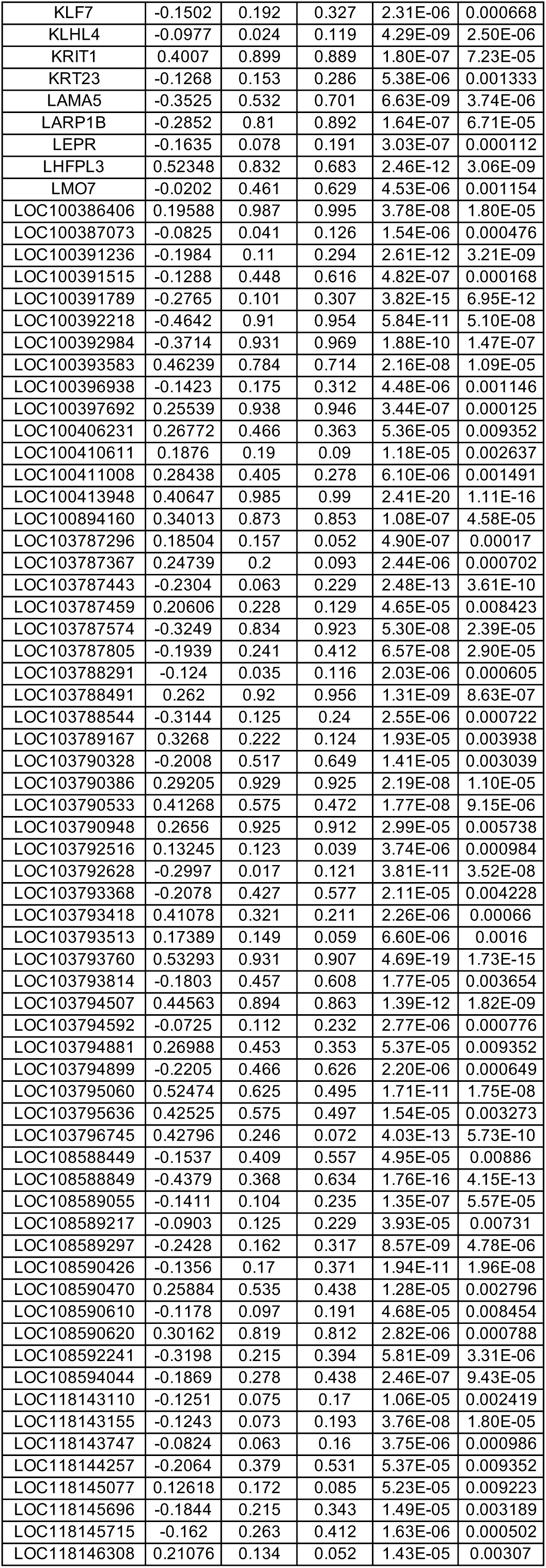
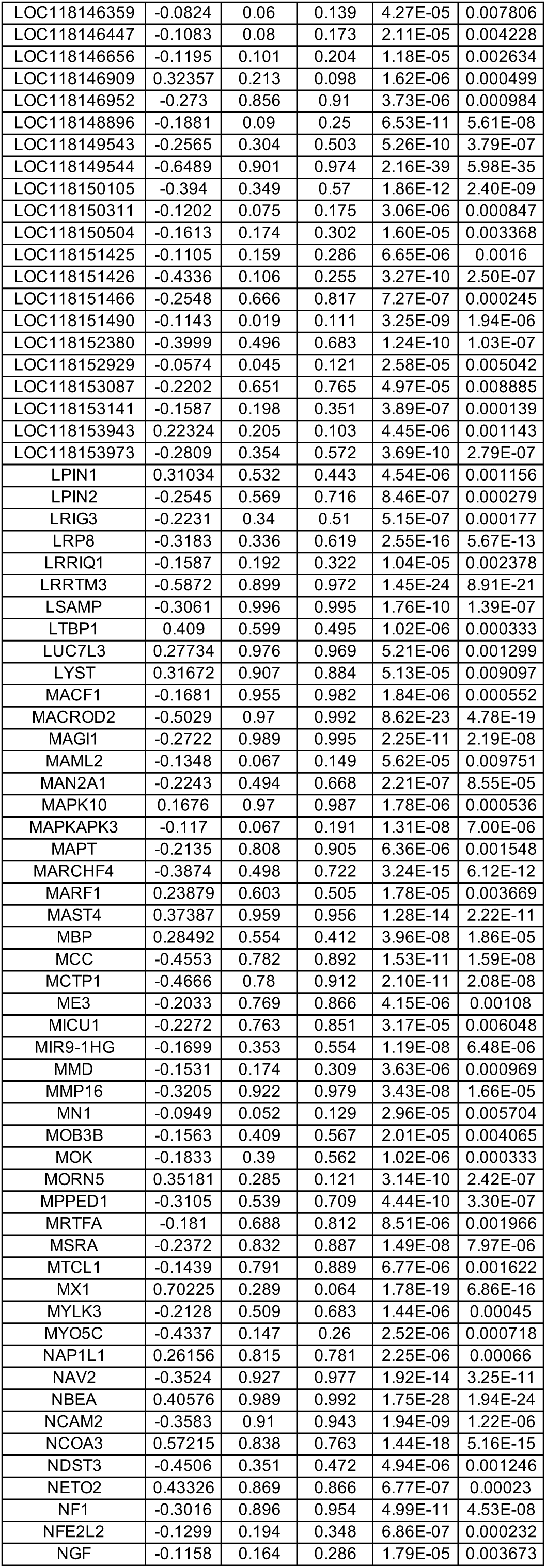
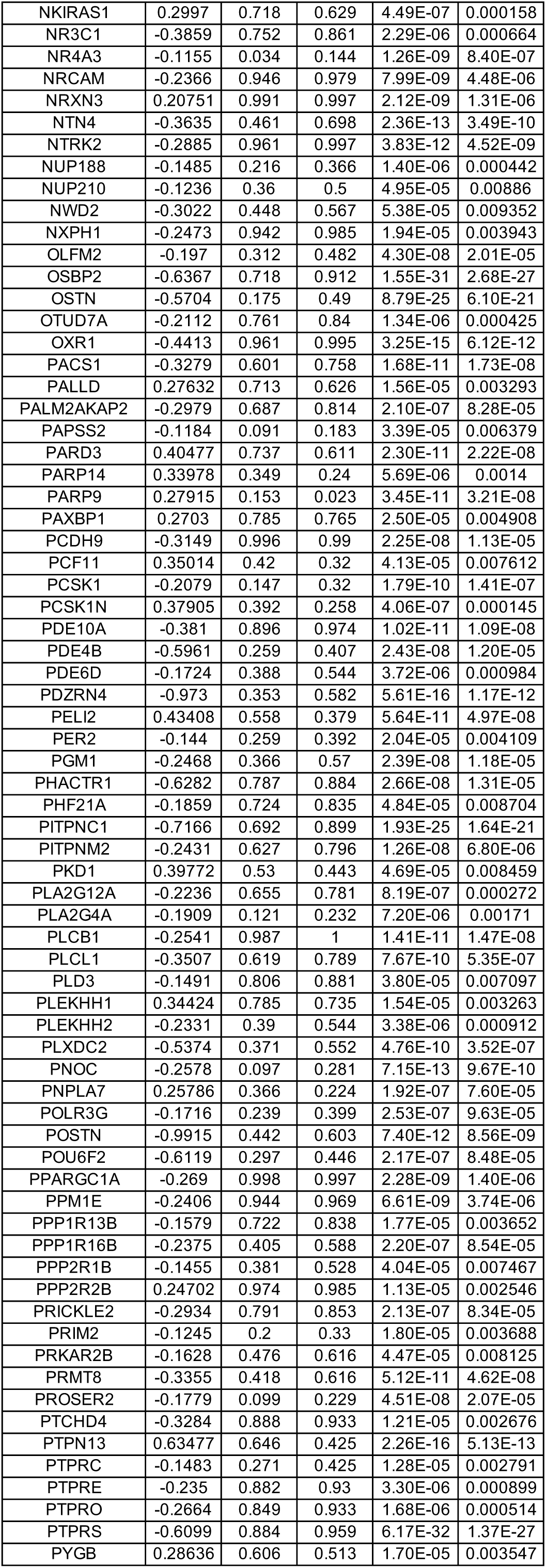
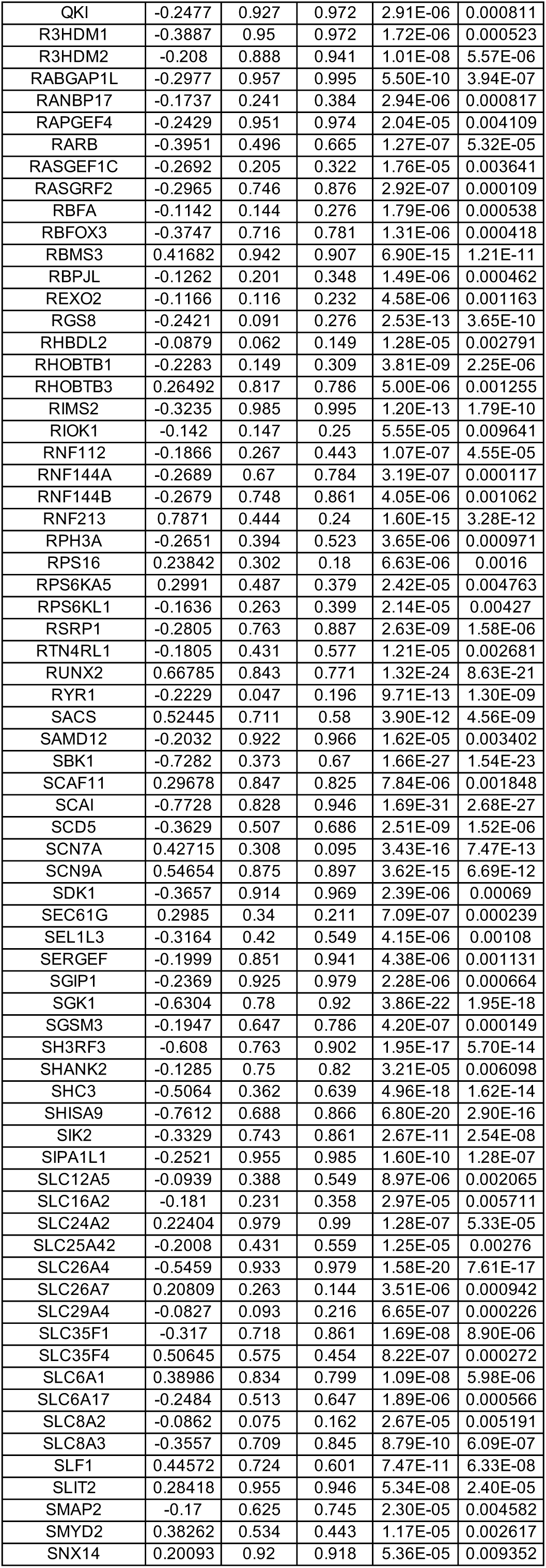
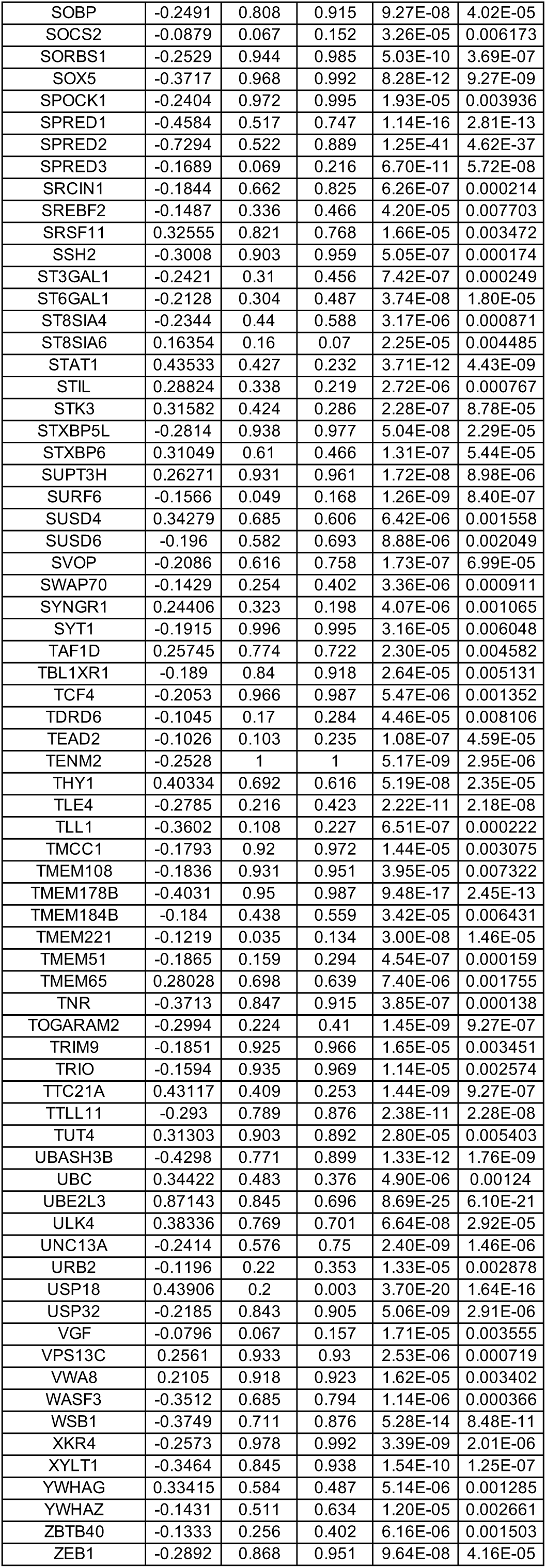
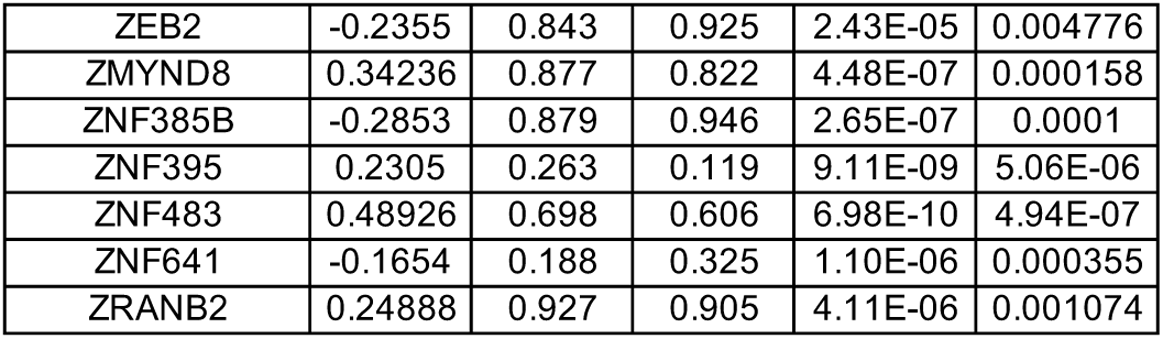
DEGs between wild-type (WT) and MECP2-null (KO) PVALB-positive inhibitory neurons of OC (adjusted p-value < 0.01)

**Table S37.**
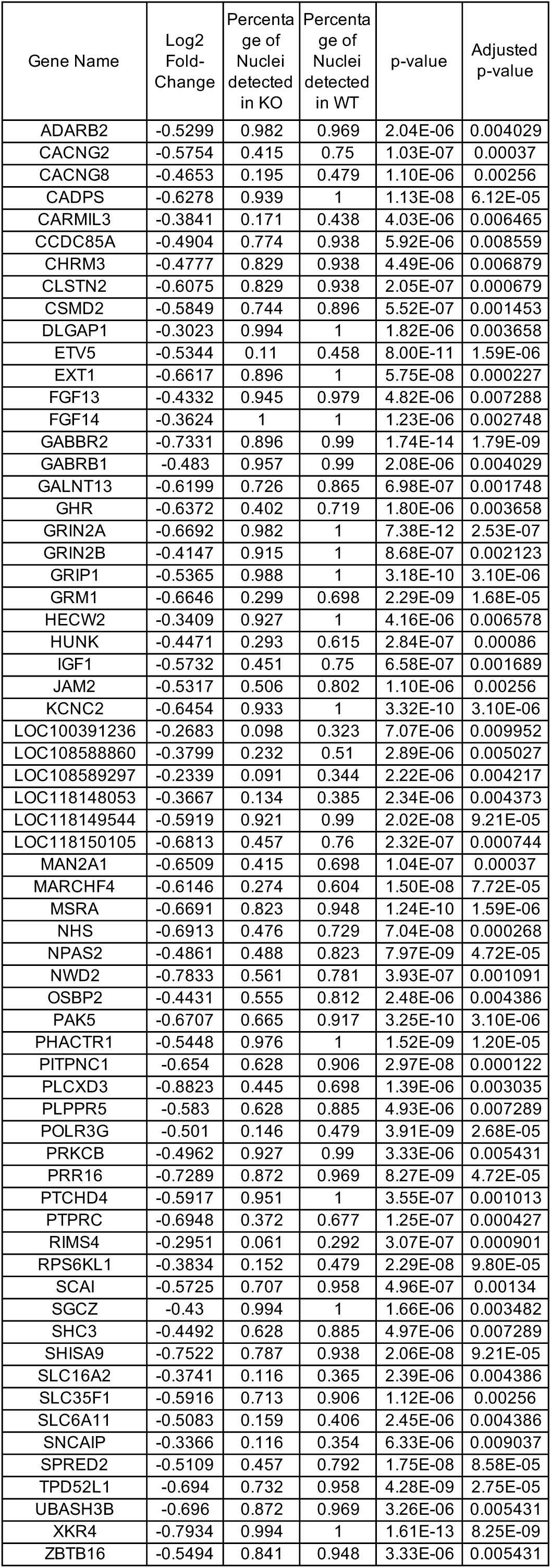
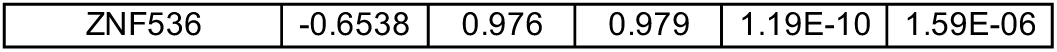
DEGs between wild-type (WT) and MECP2-null (KO) LAMP5-positive inhibitory neurons of OC (adjusted p-value < 0.01)

**Table S38.**
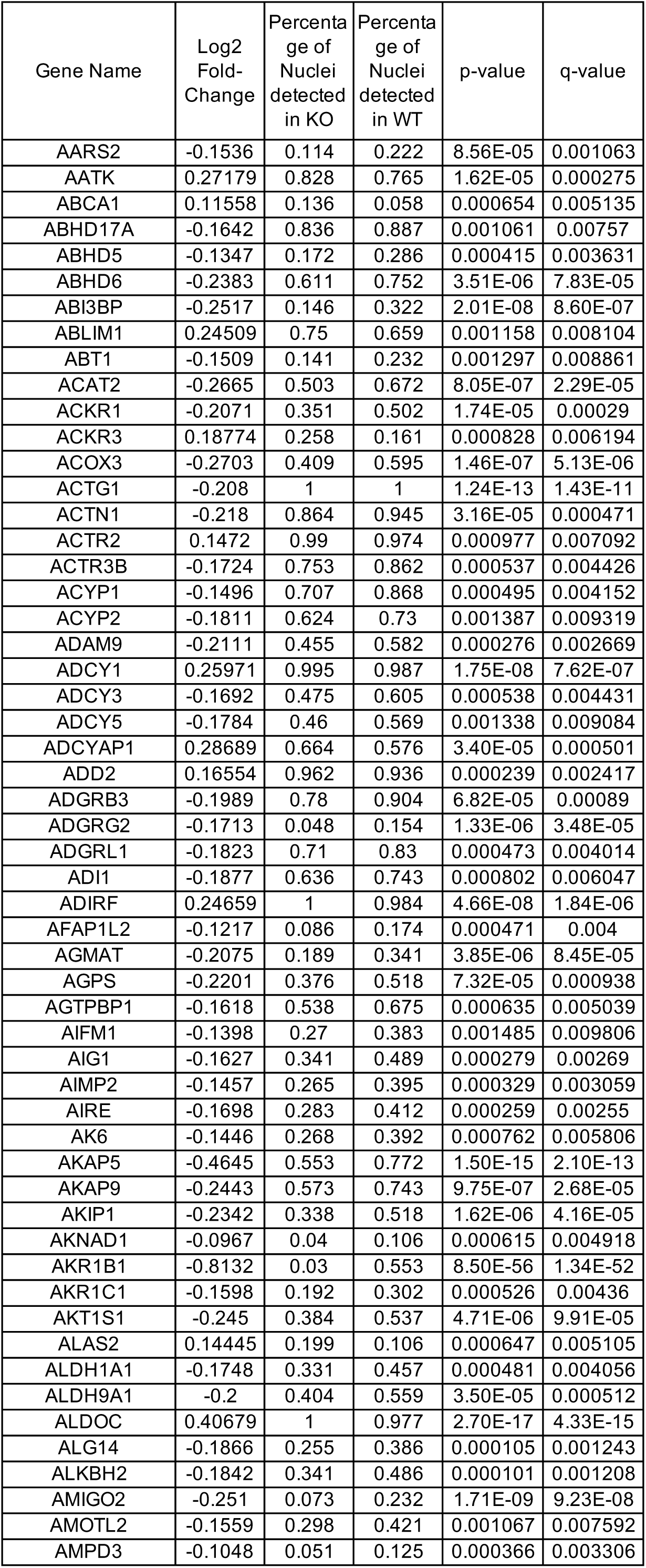
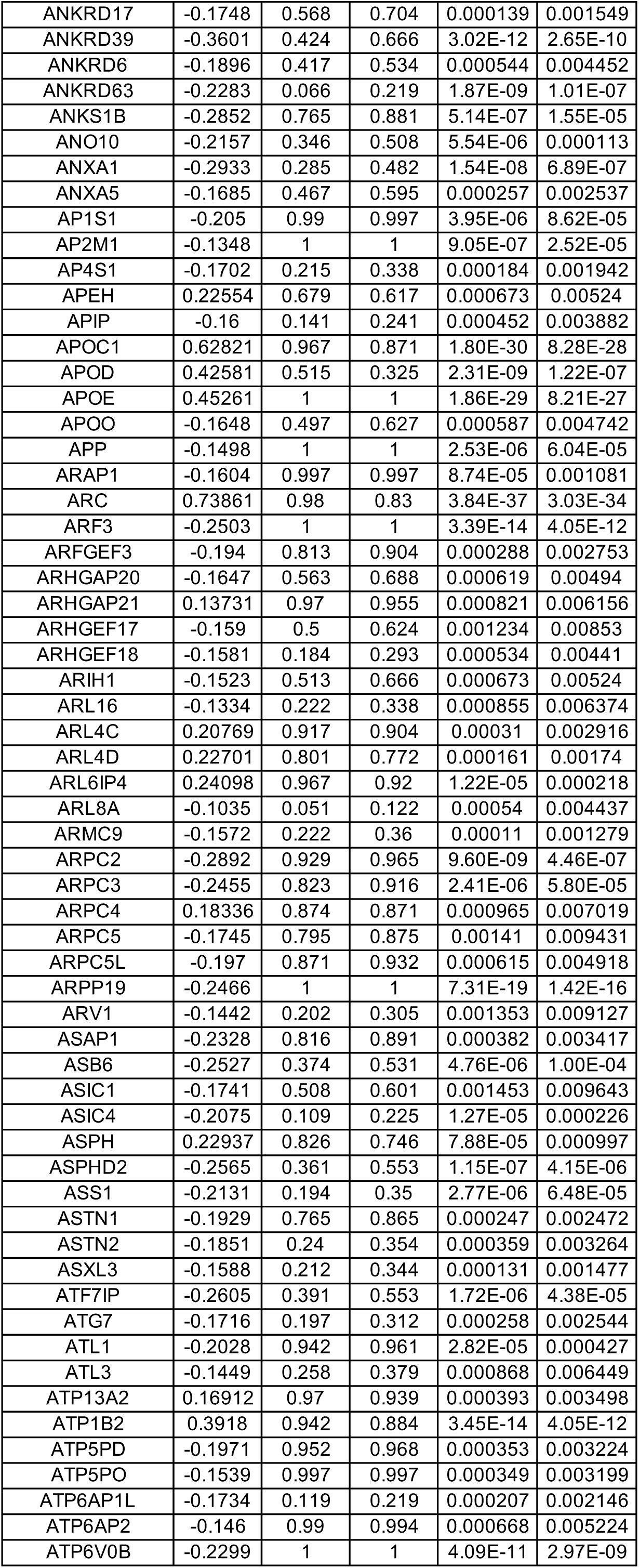
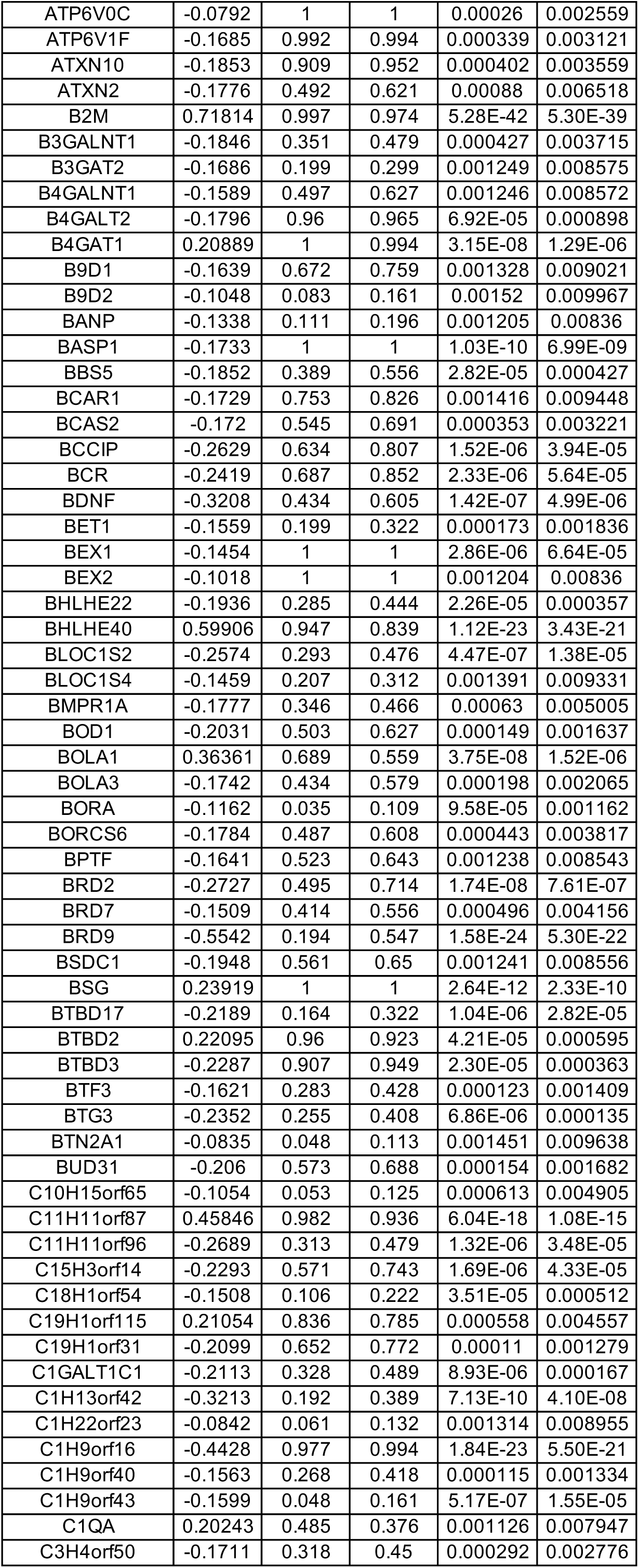
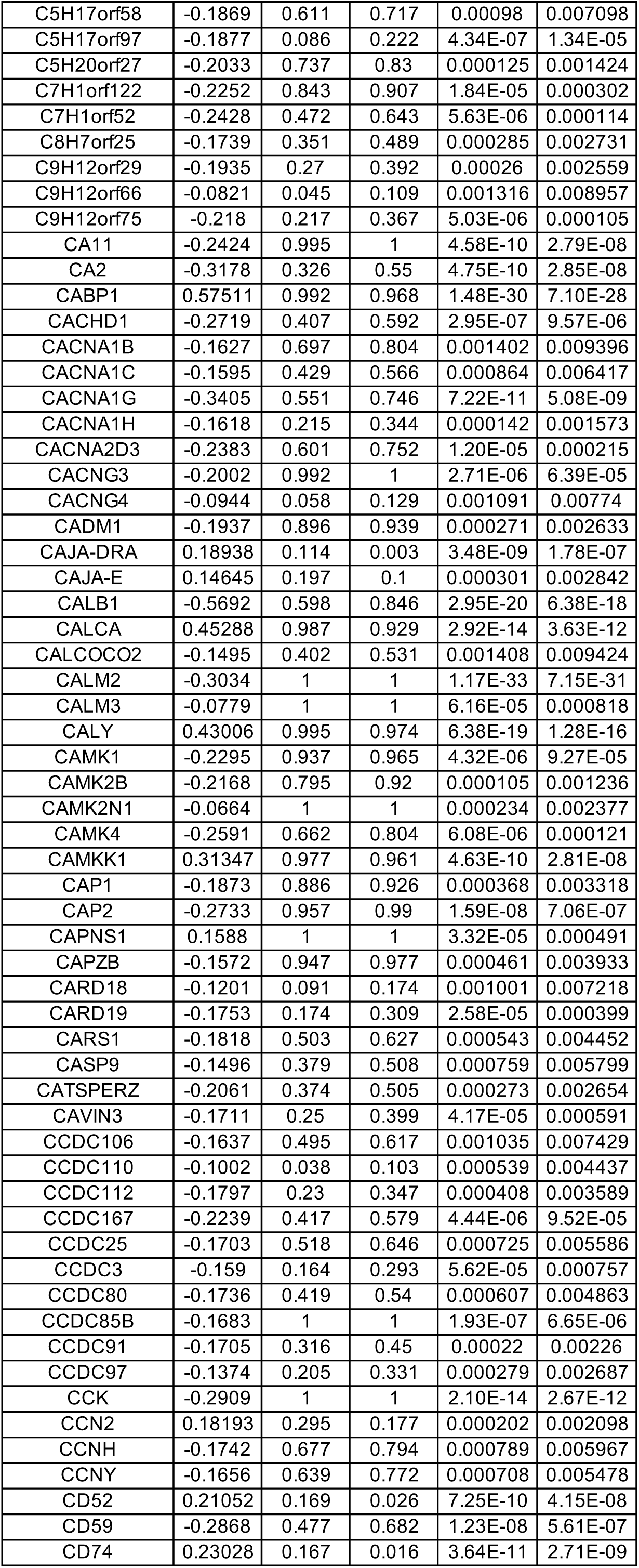
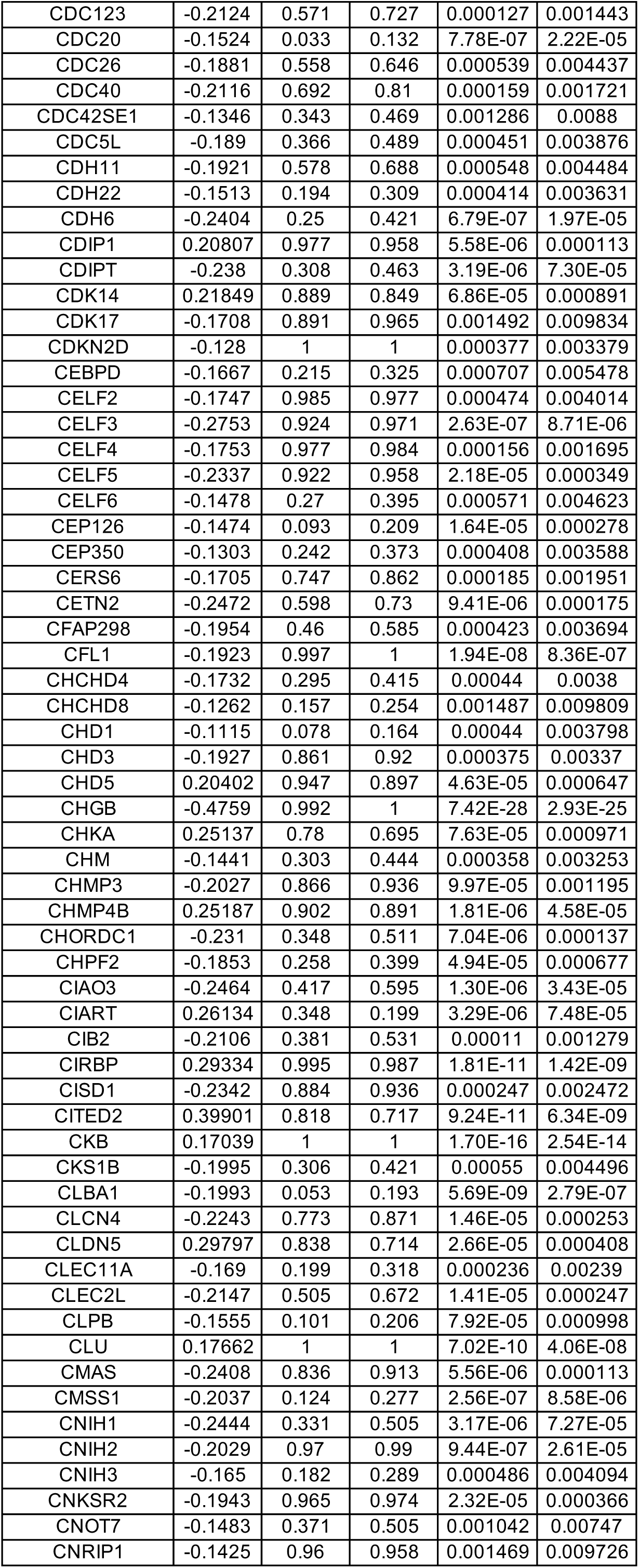
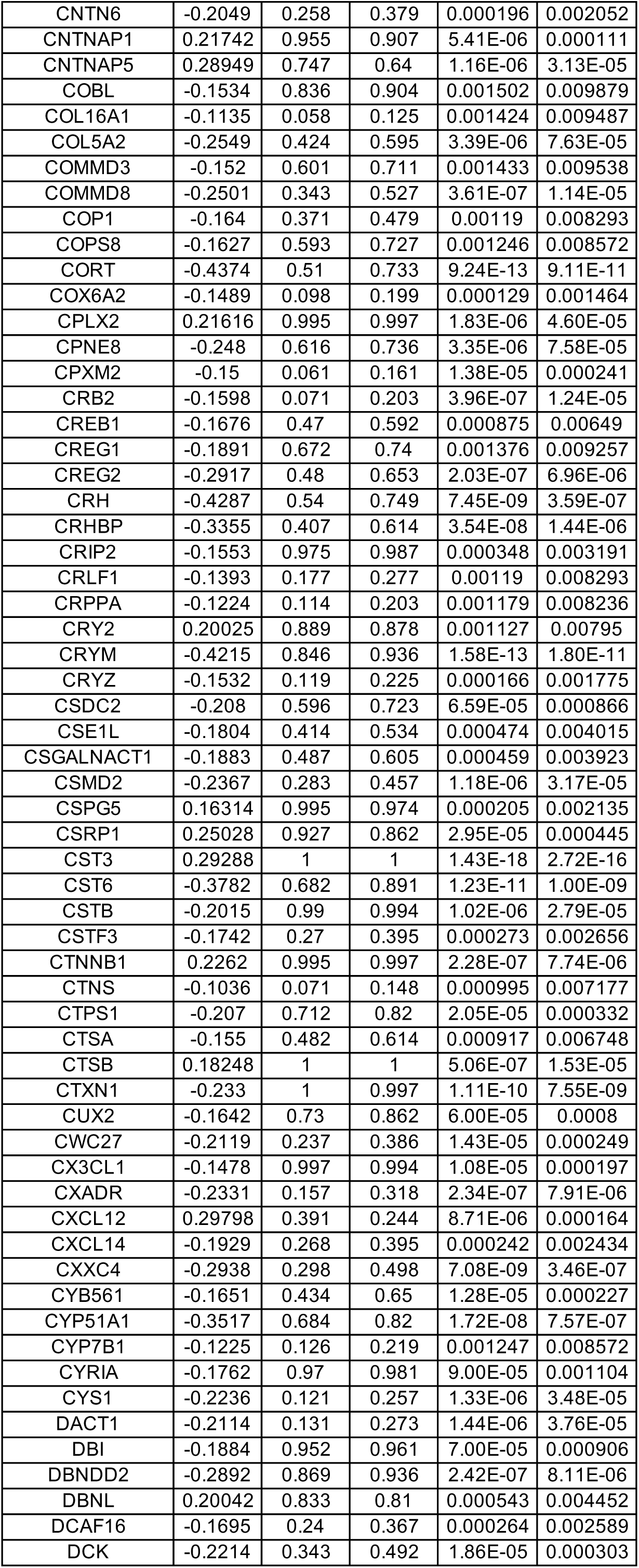
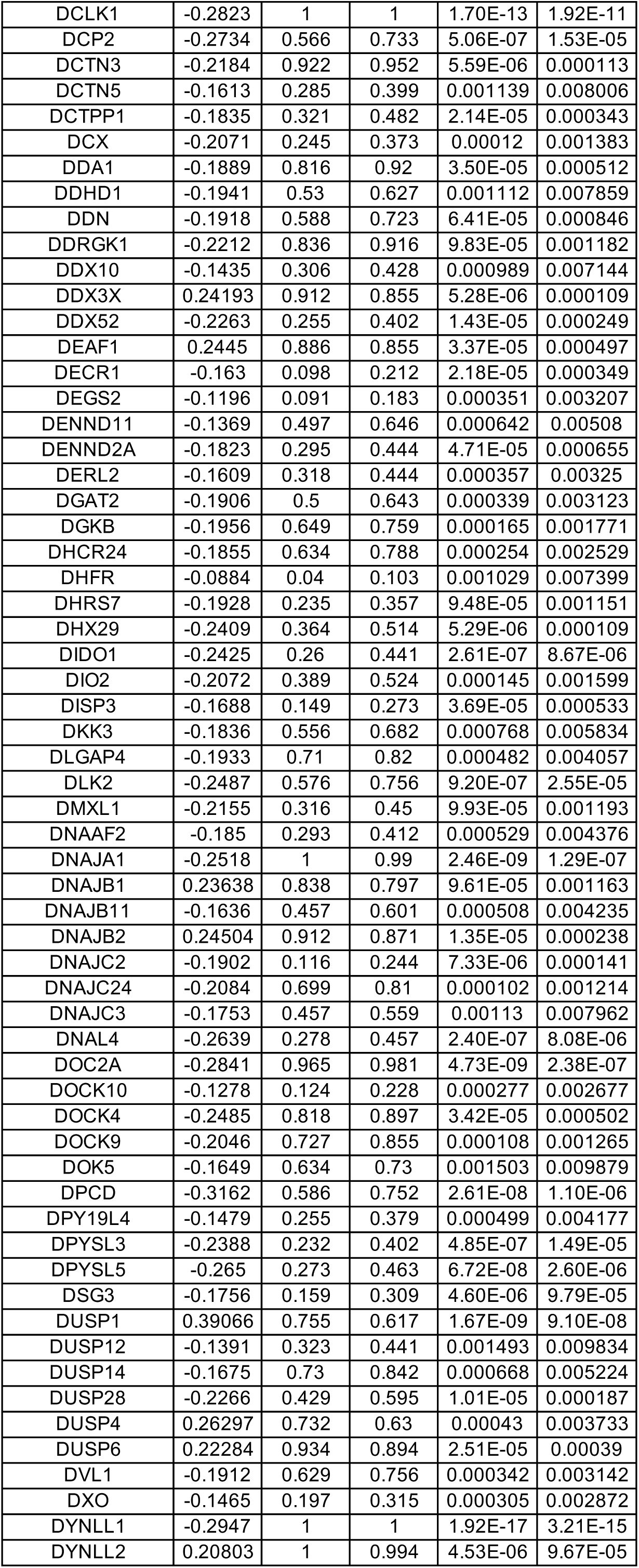
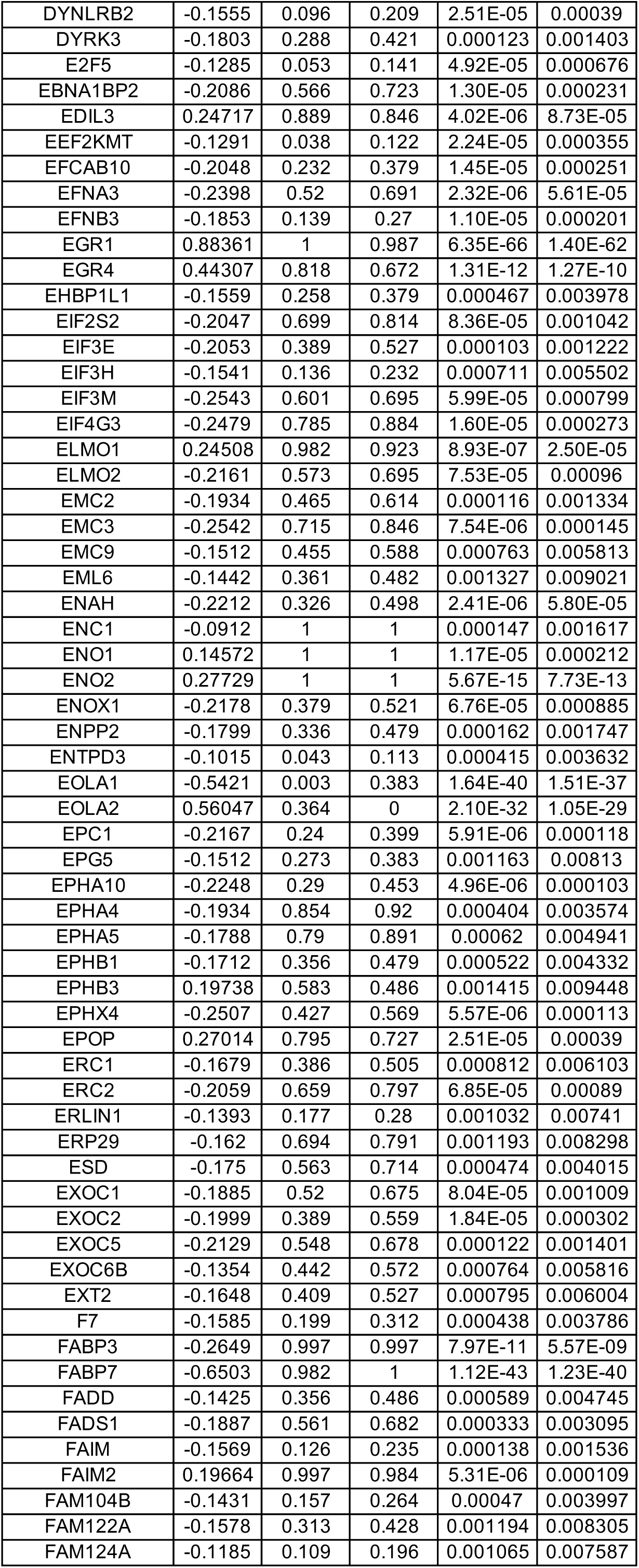
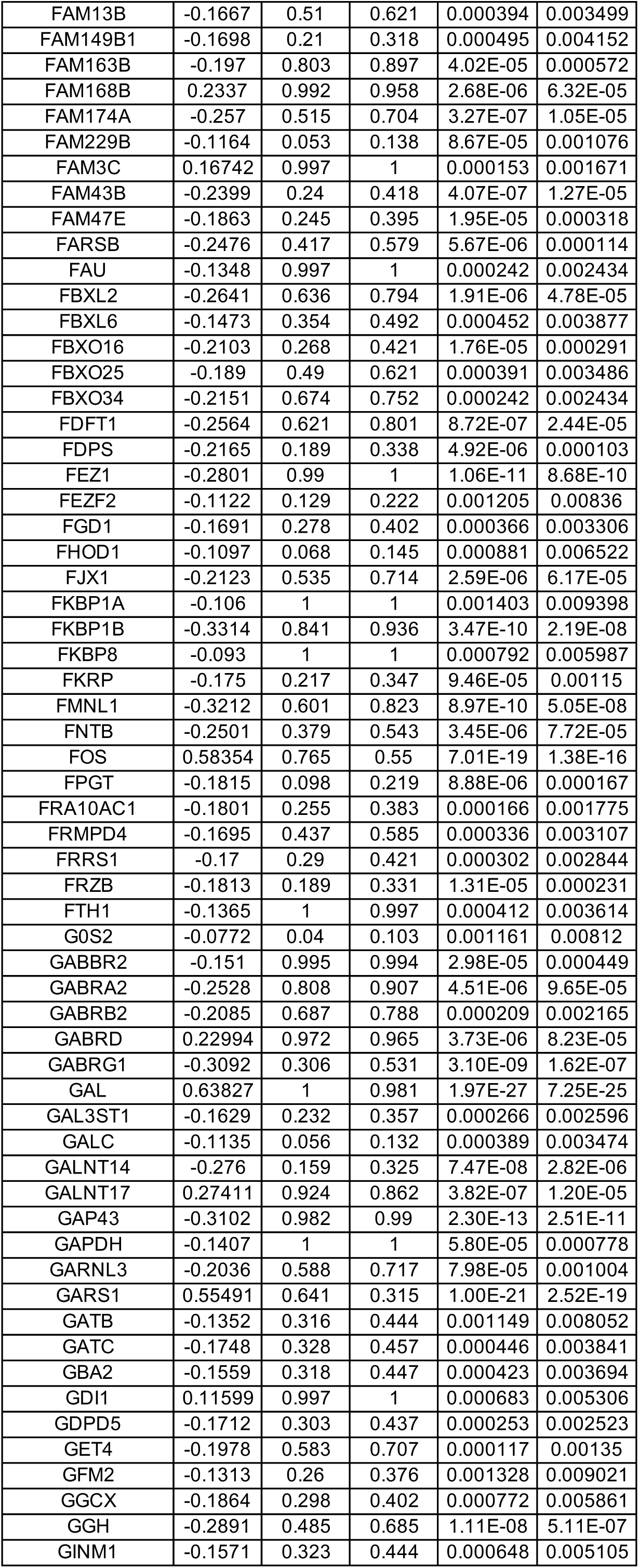
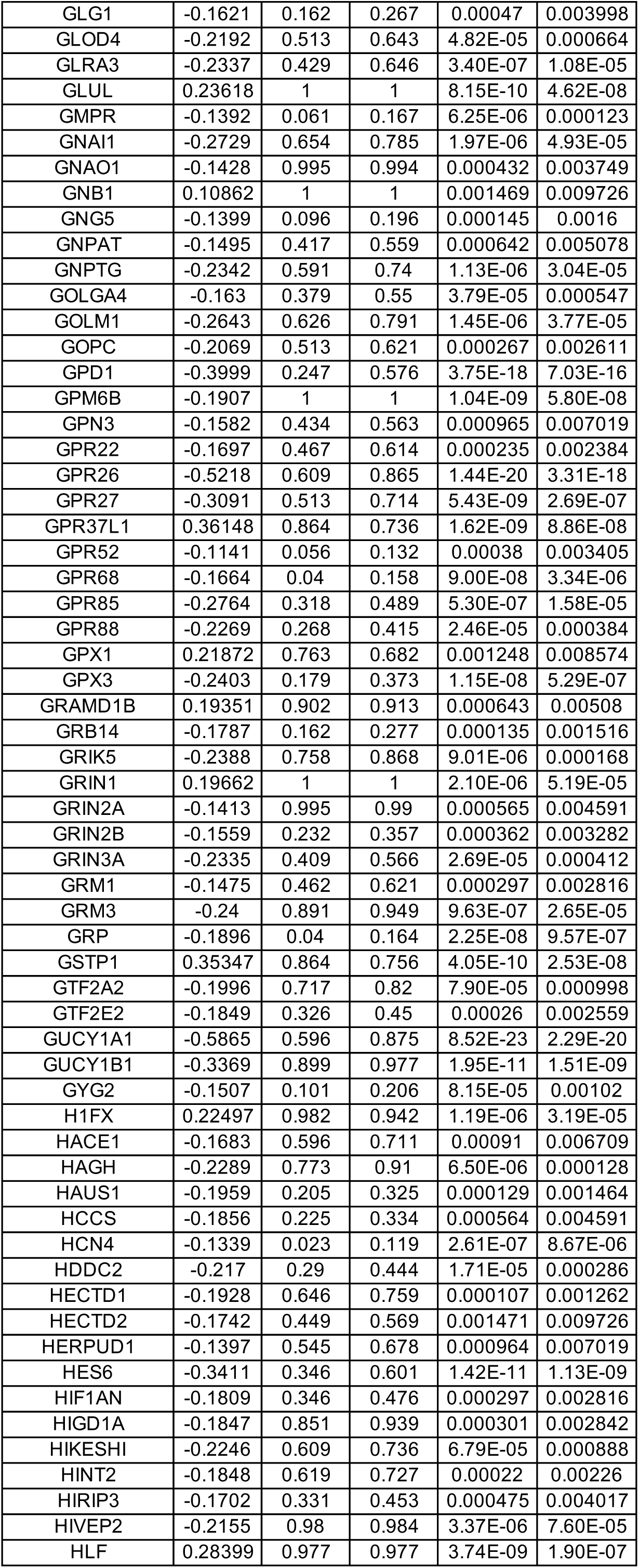
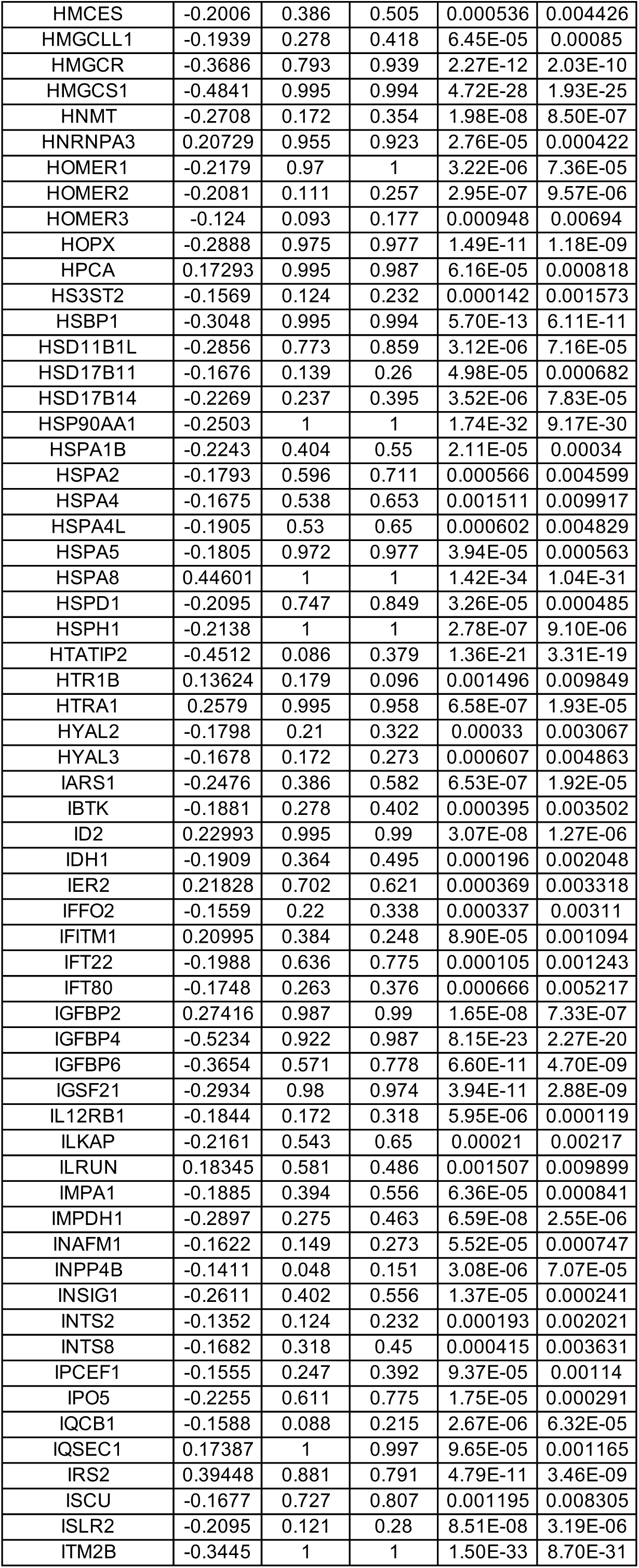
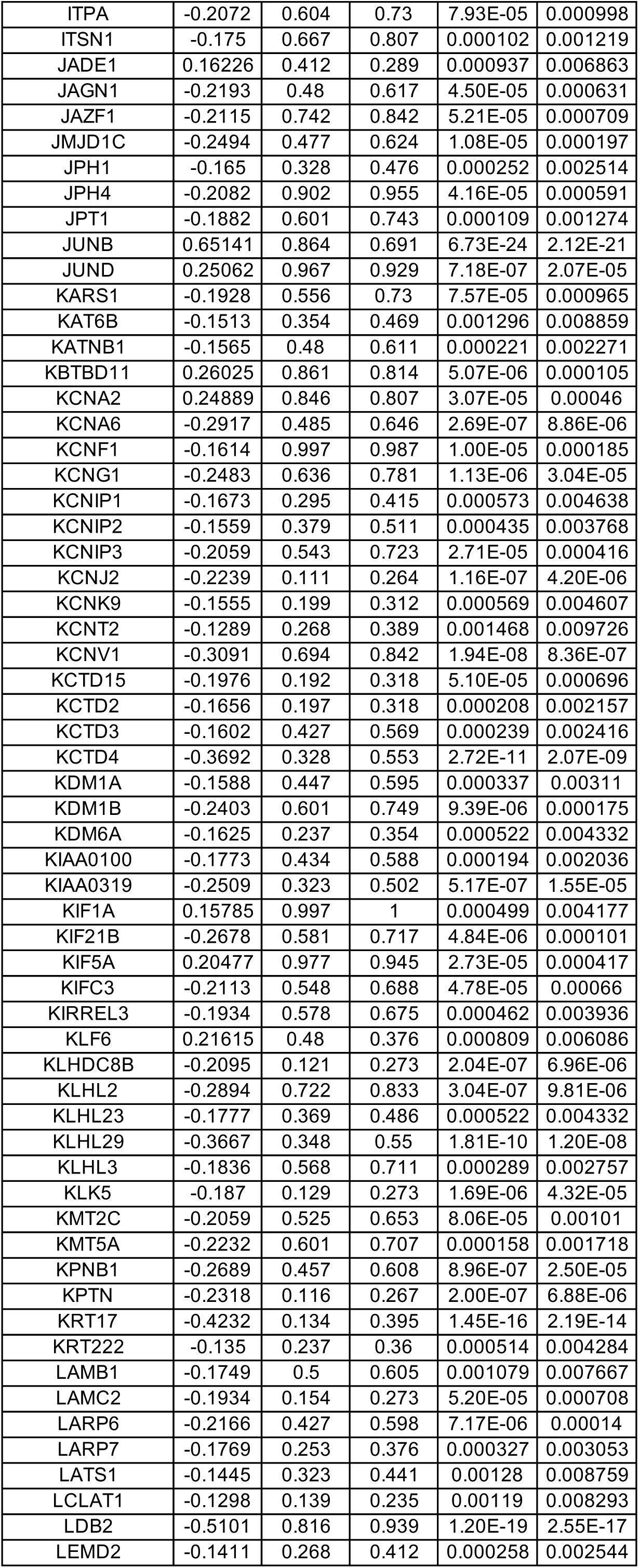
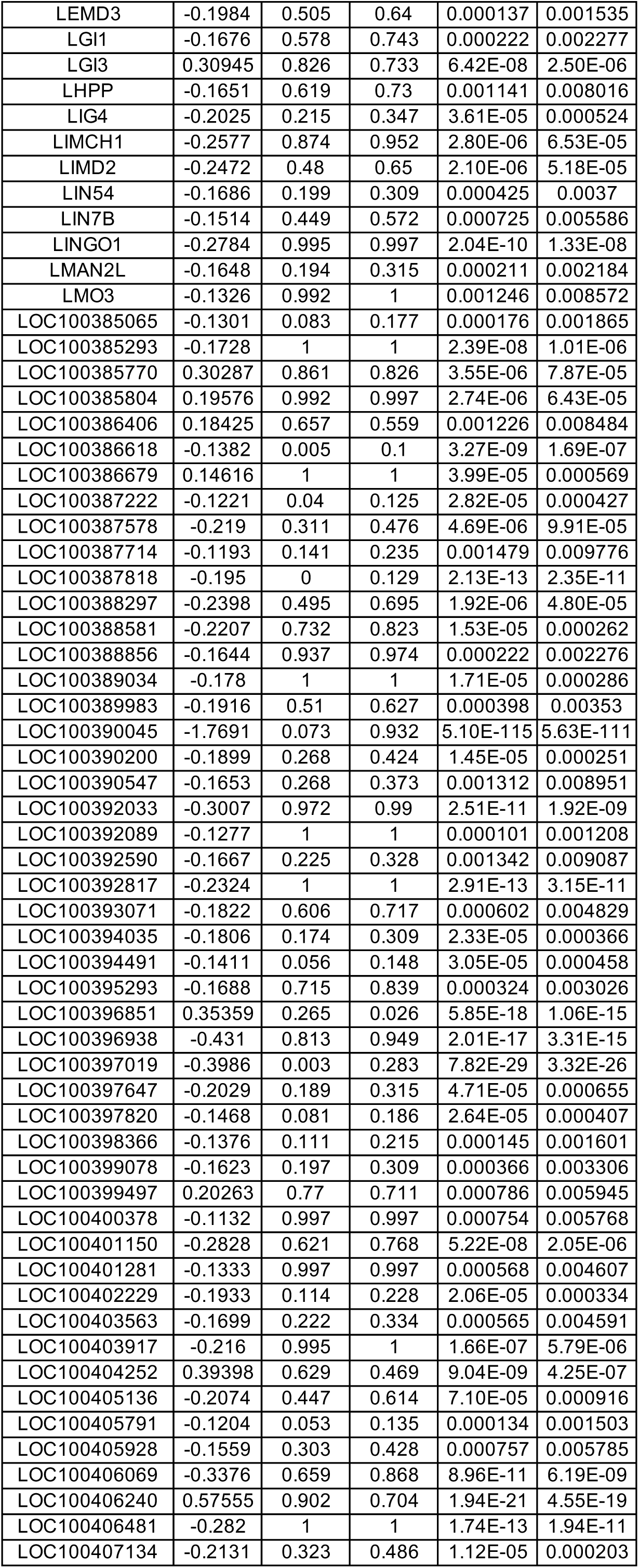
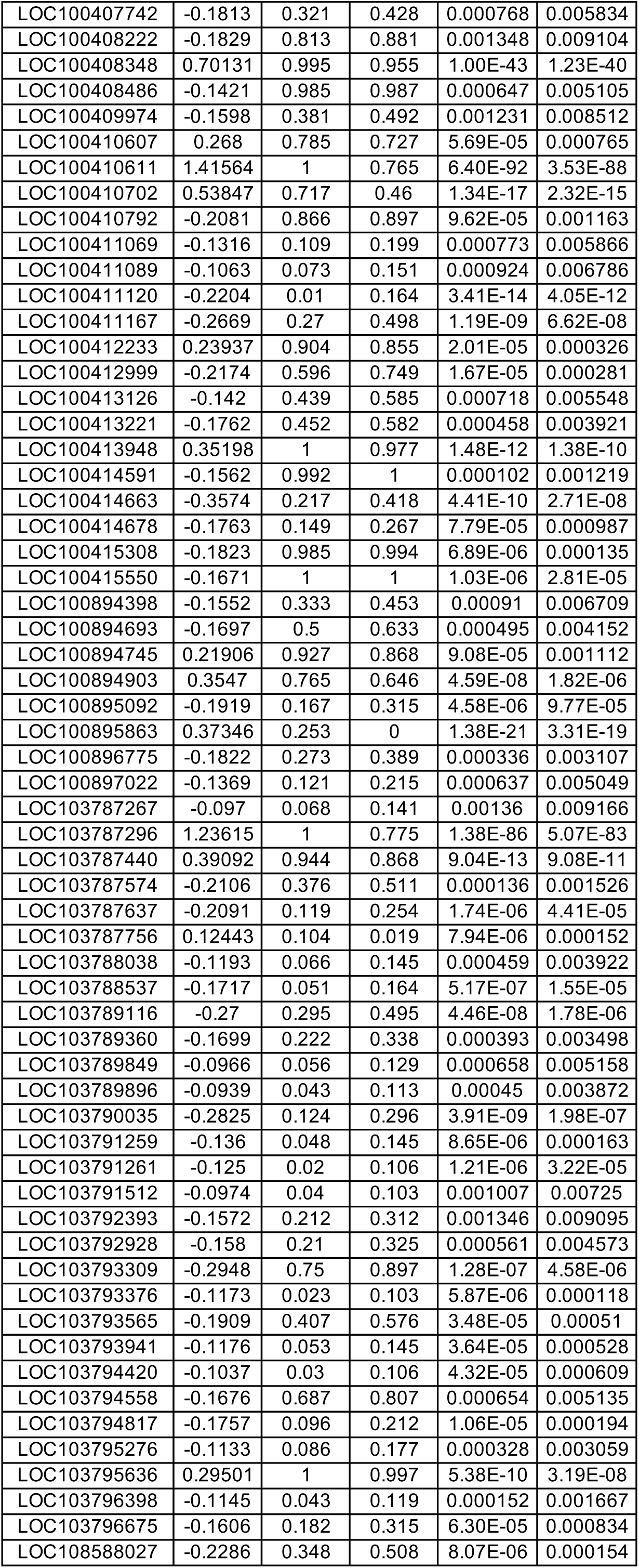
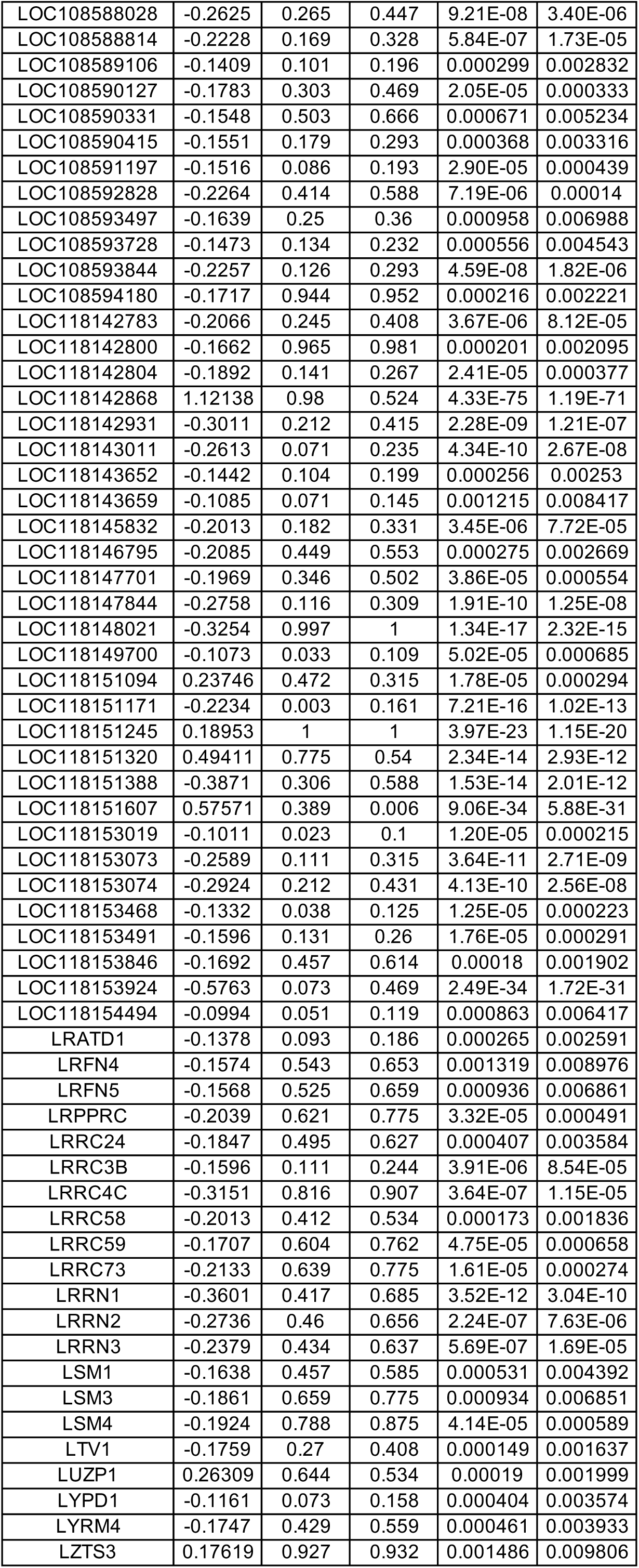
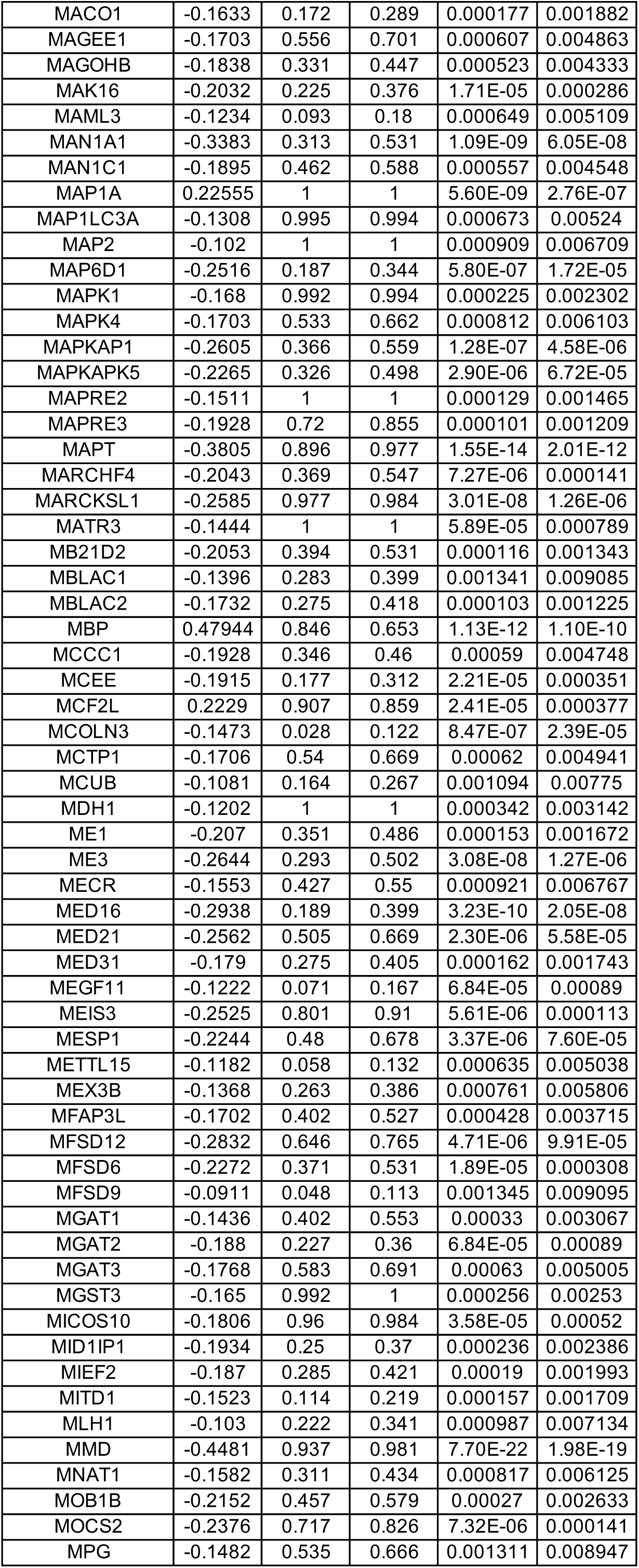
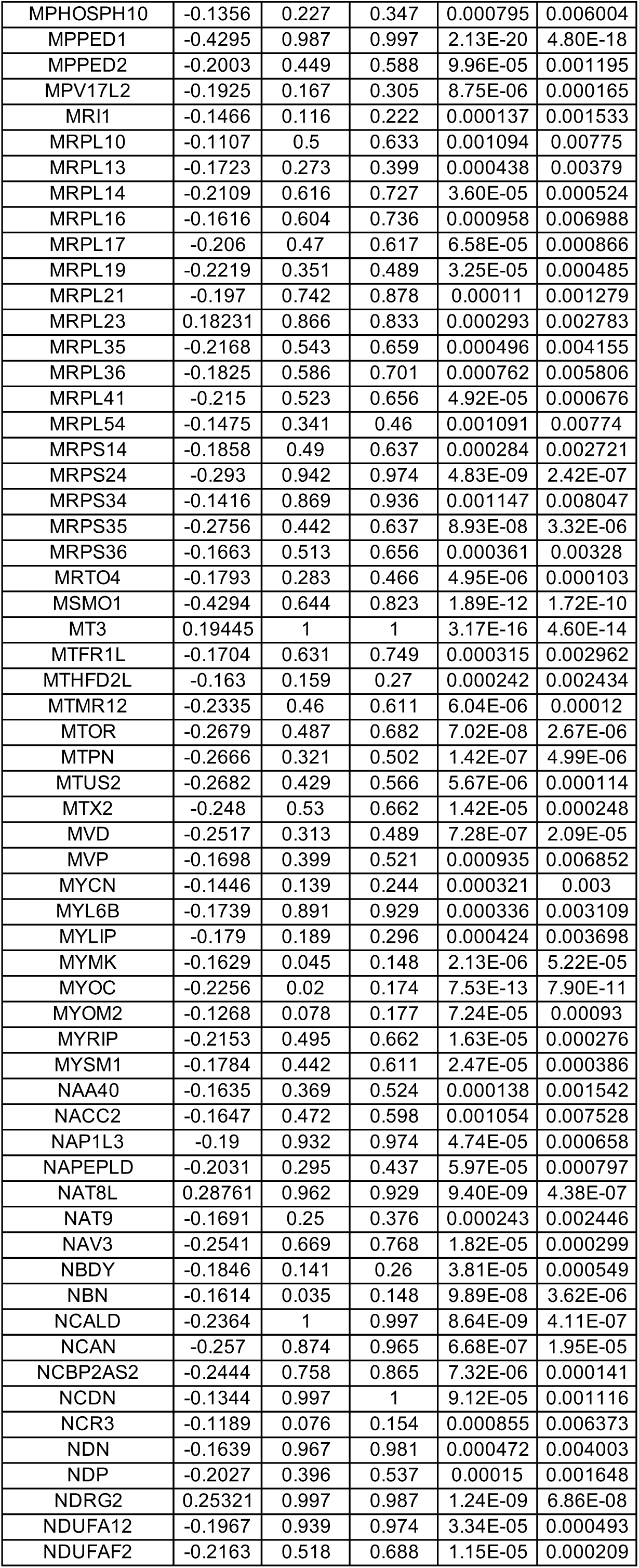
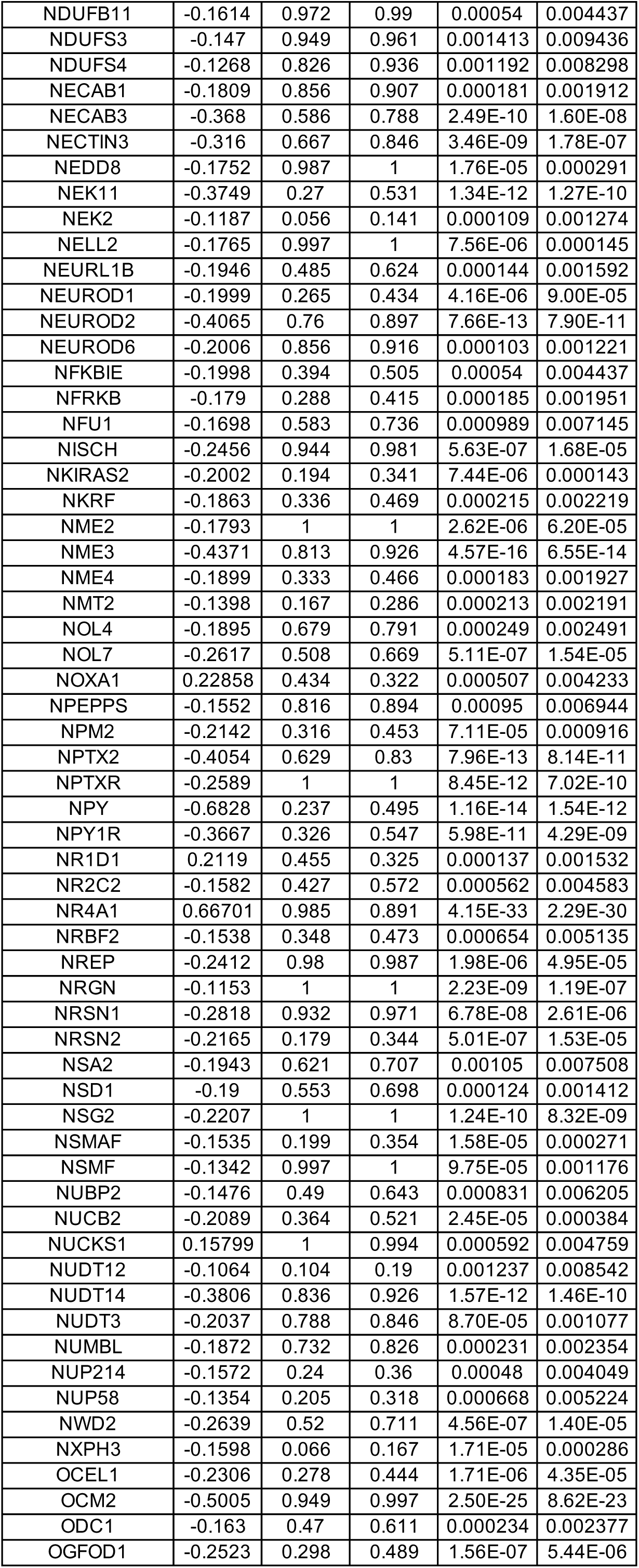
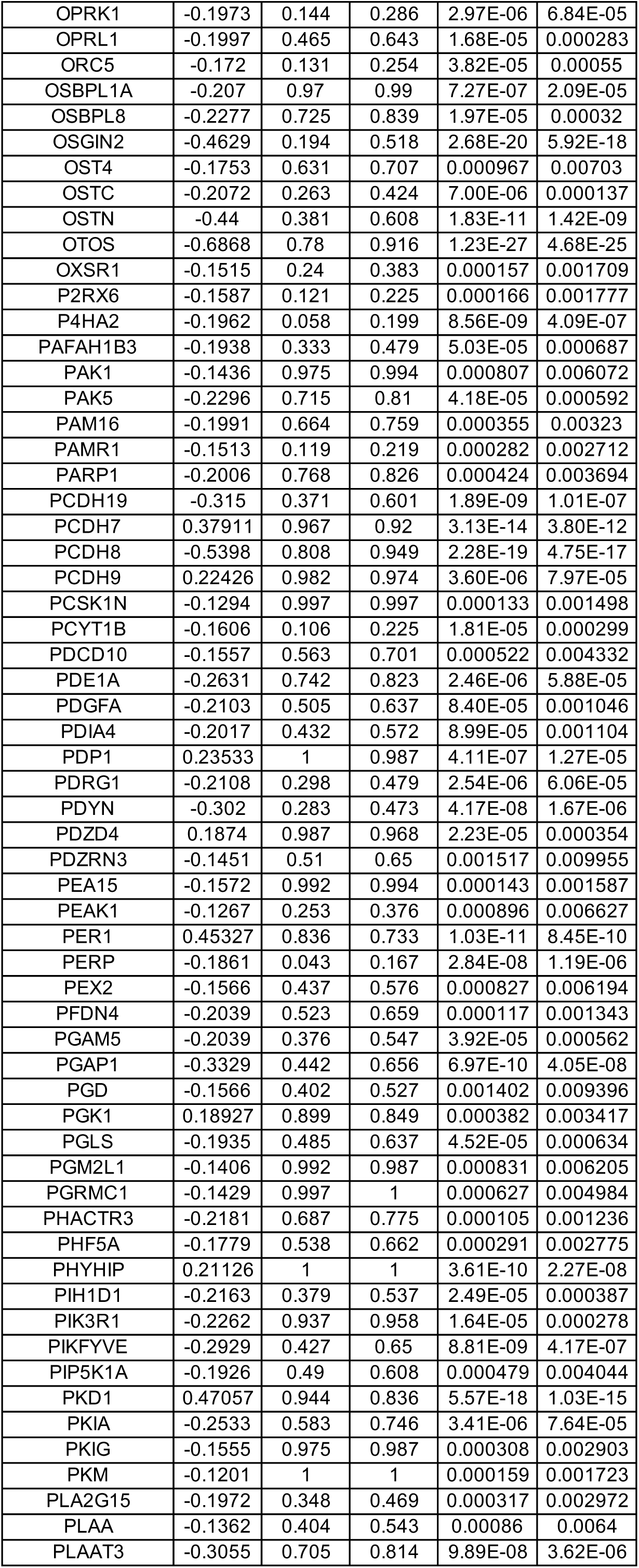
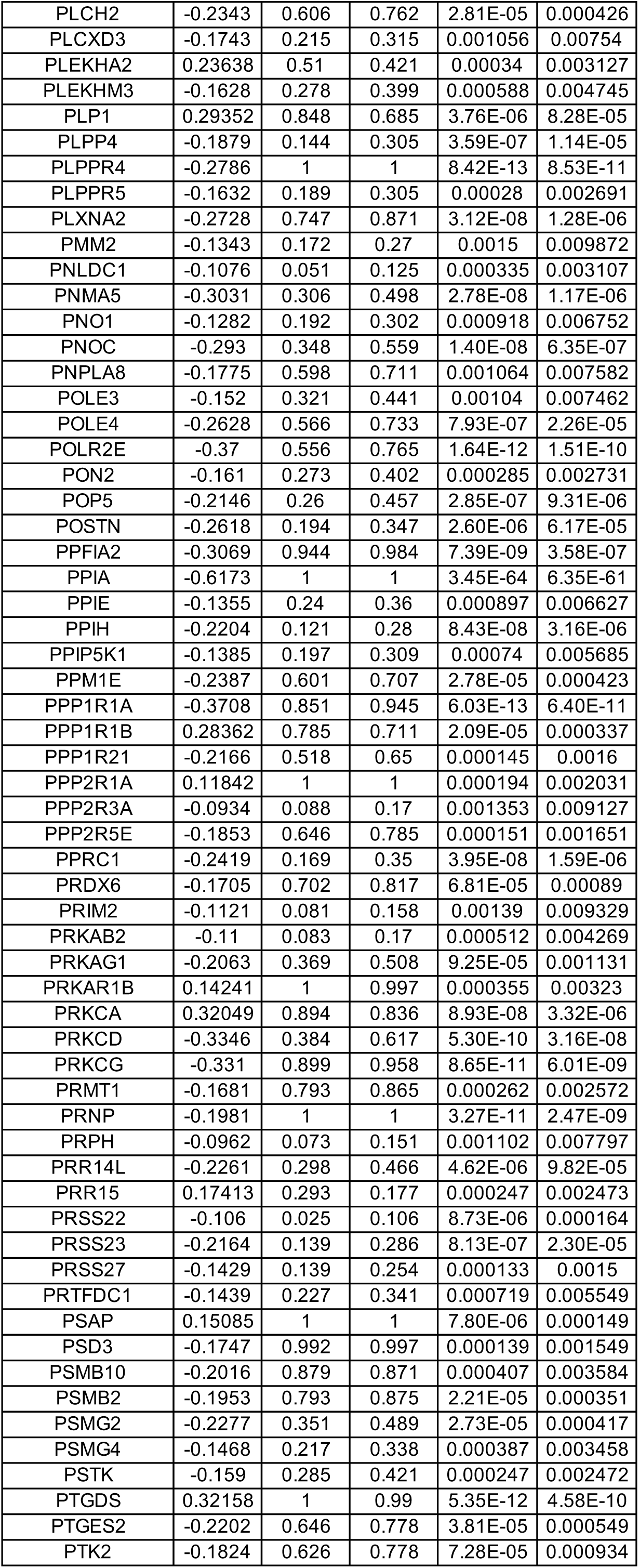
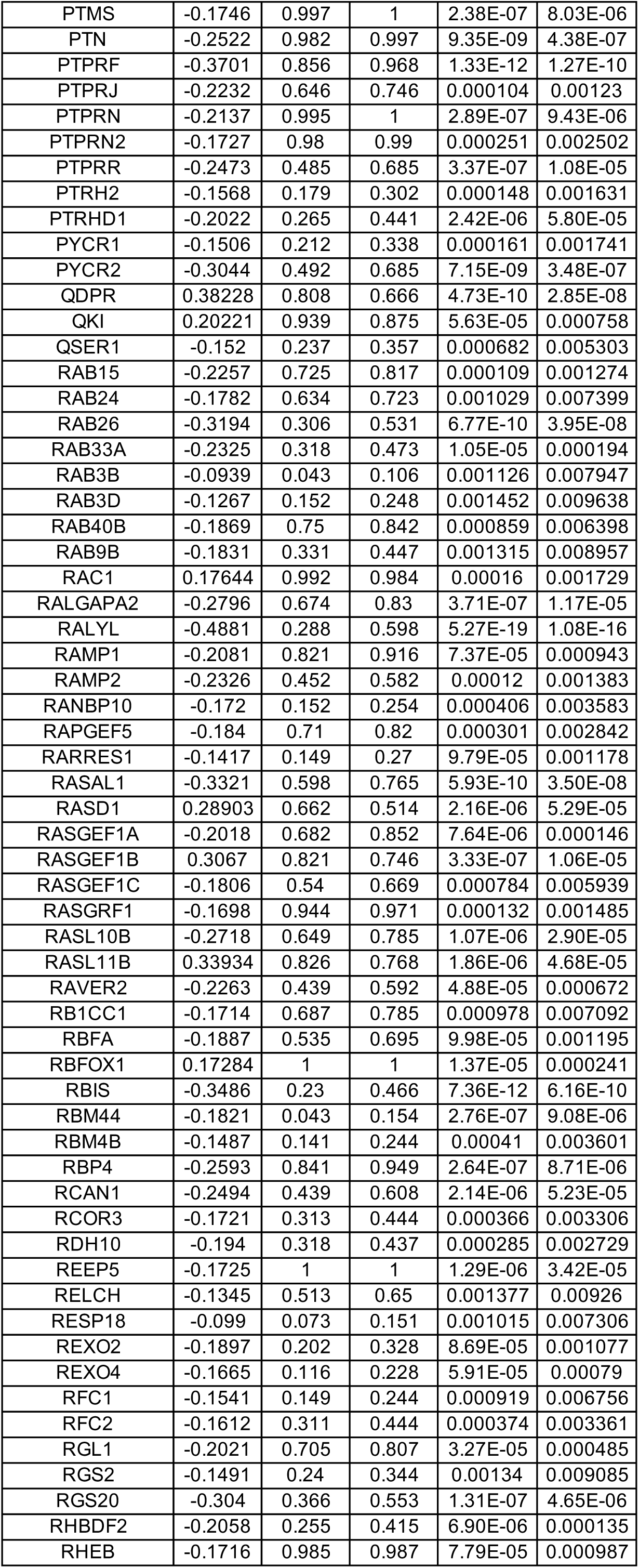
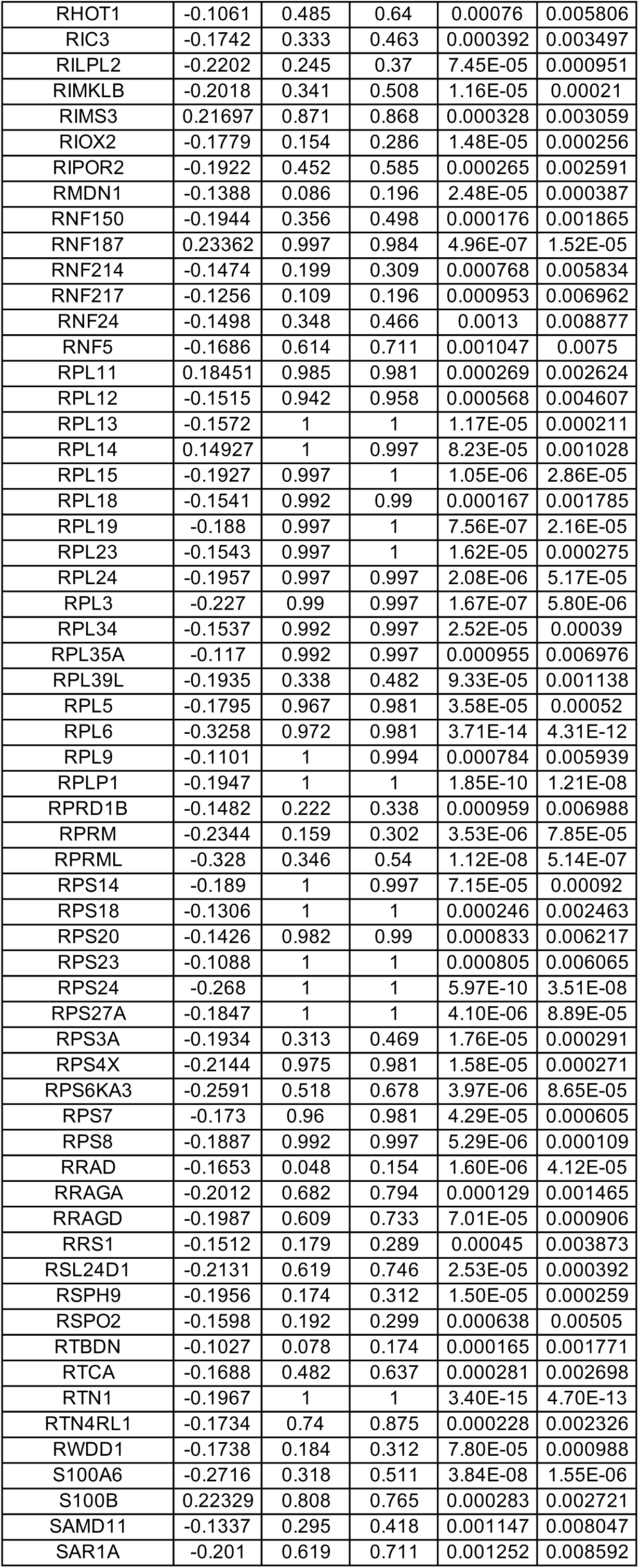
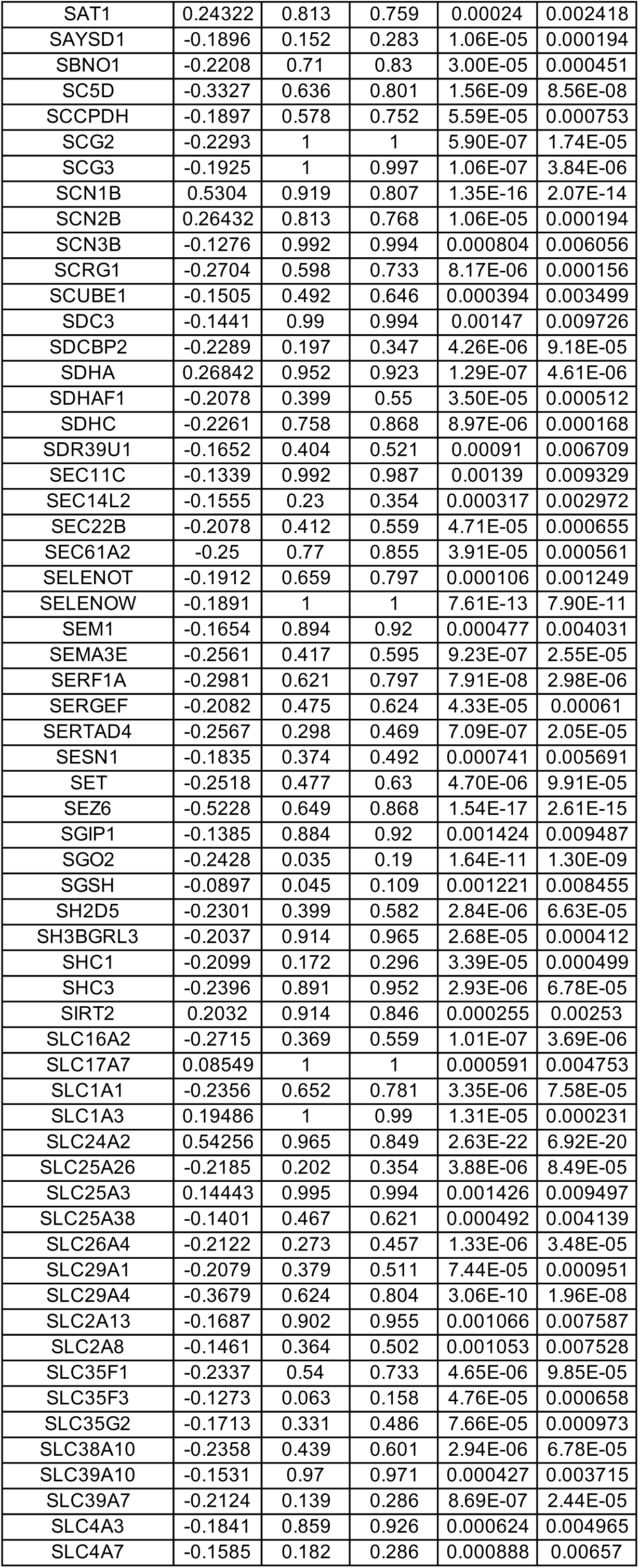
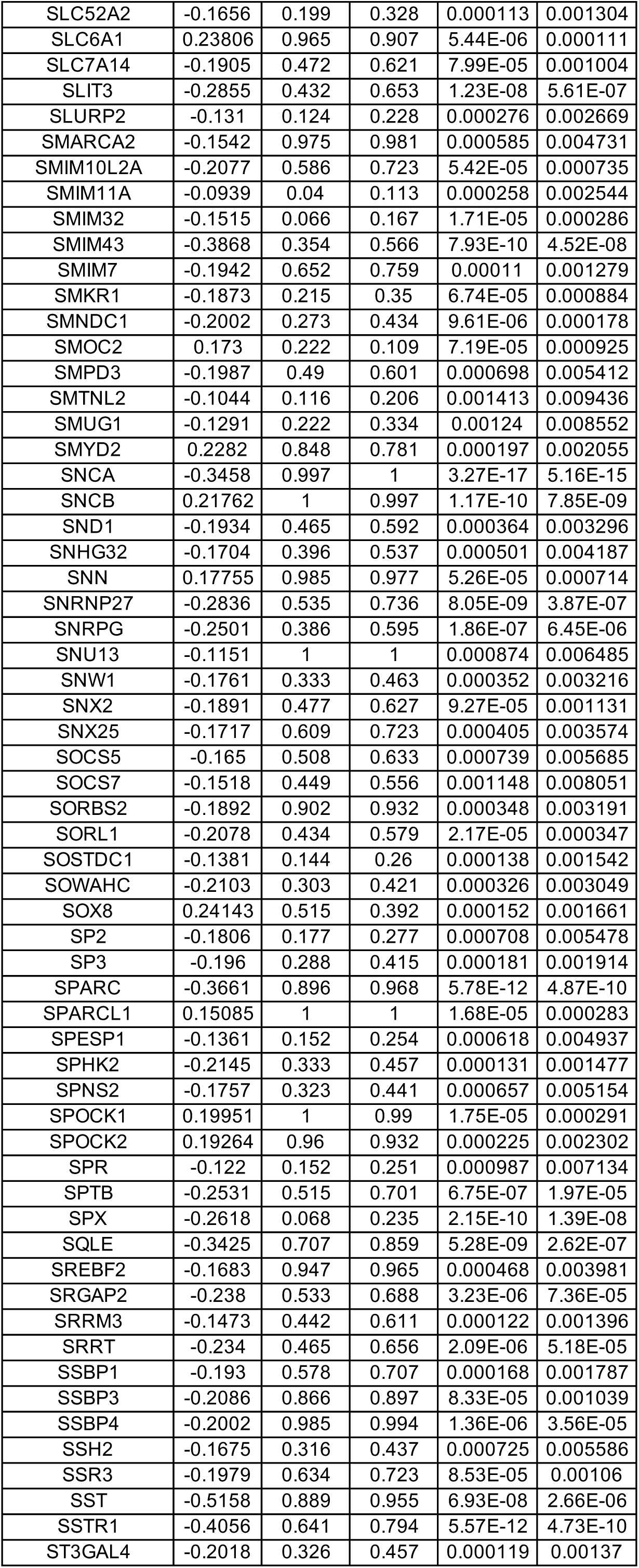
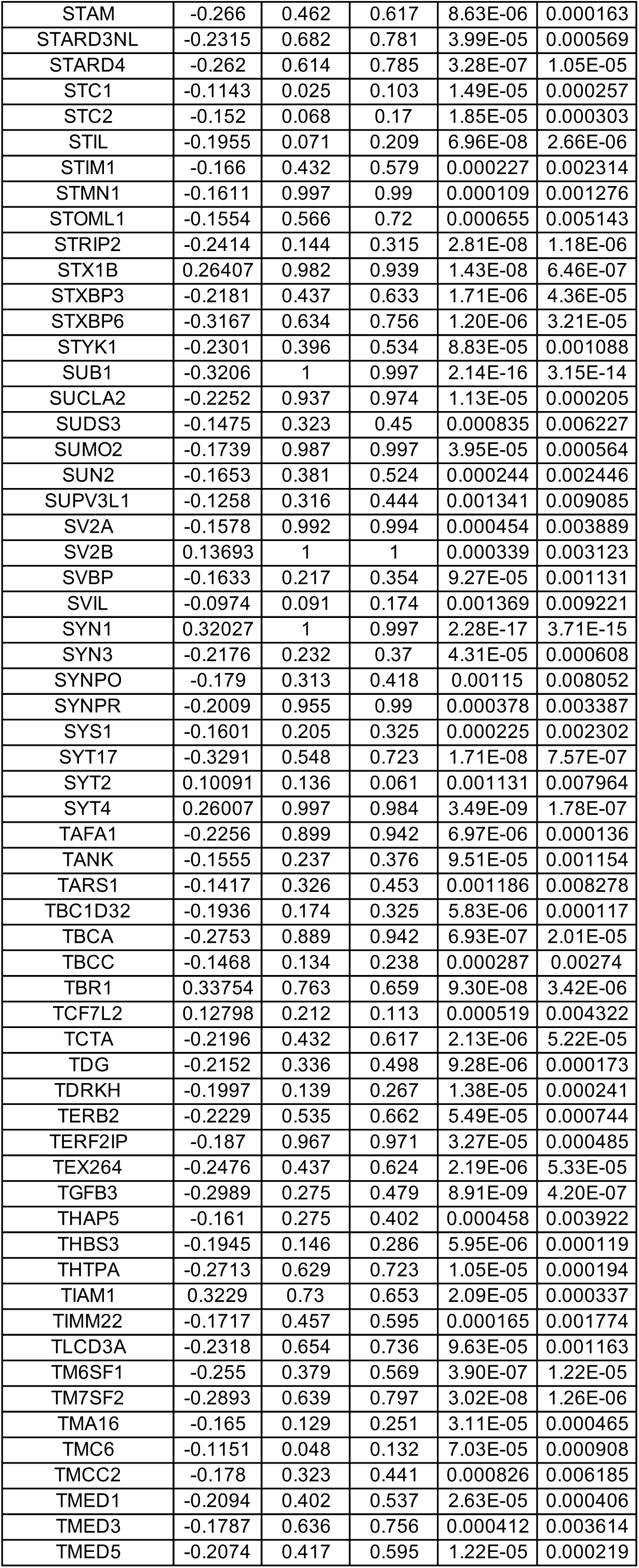
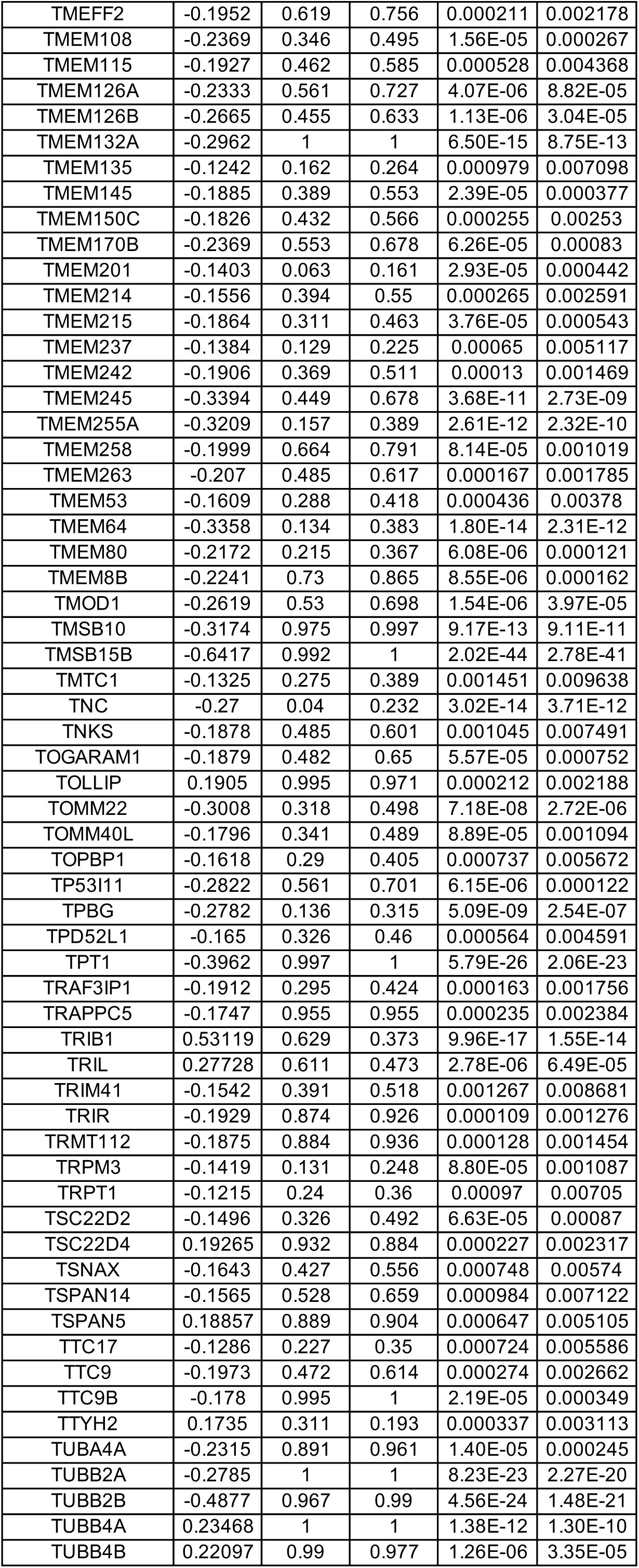
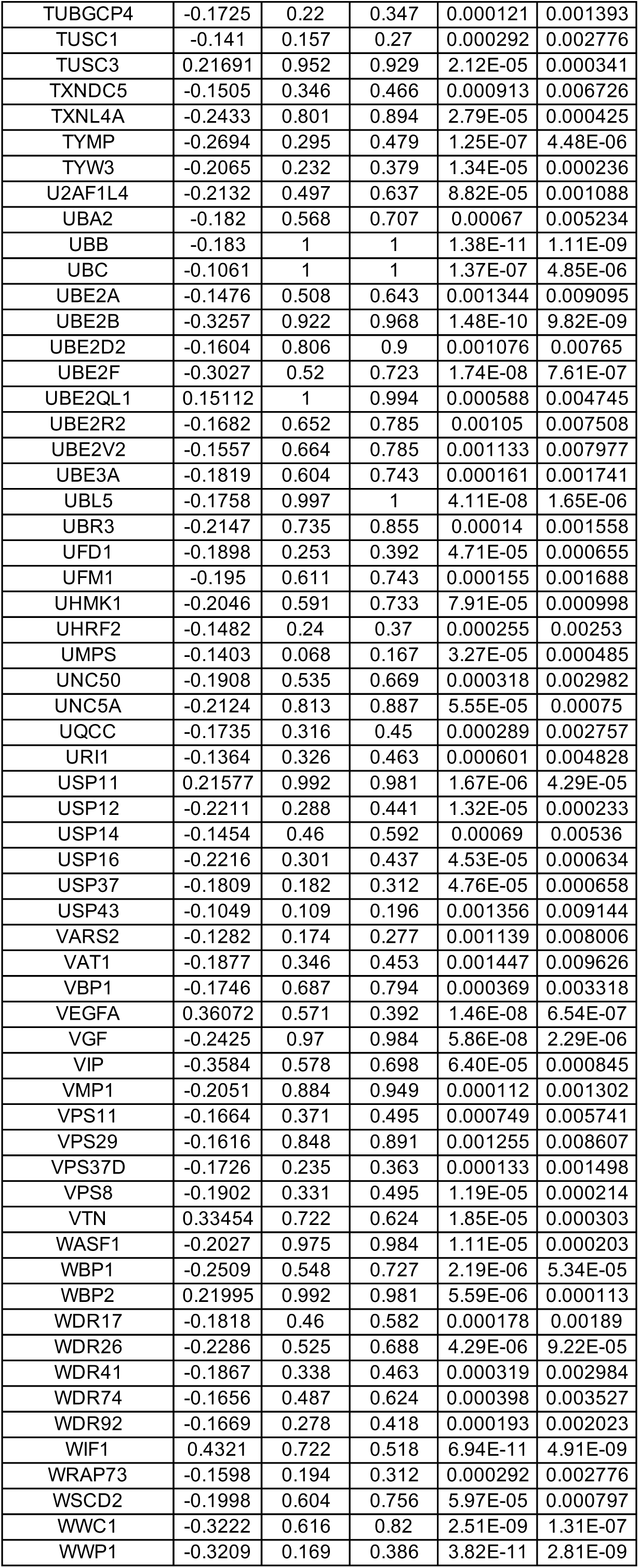
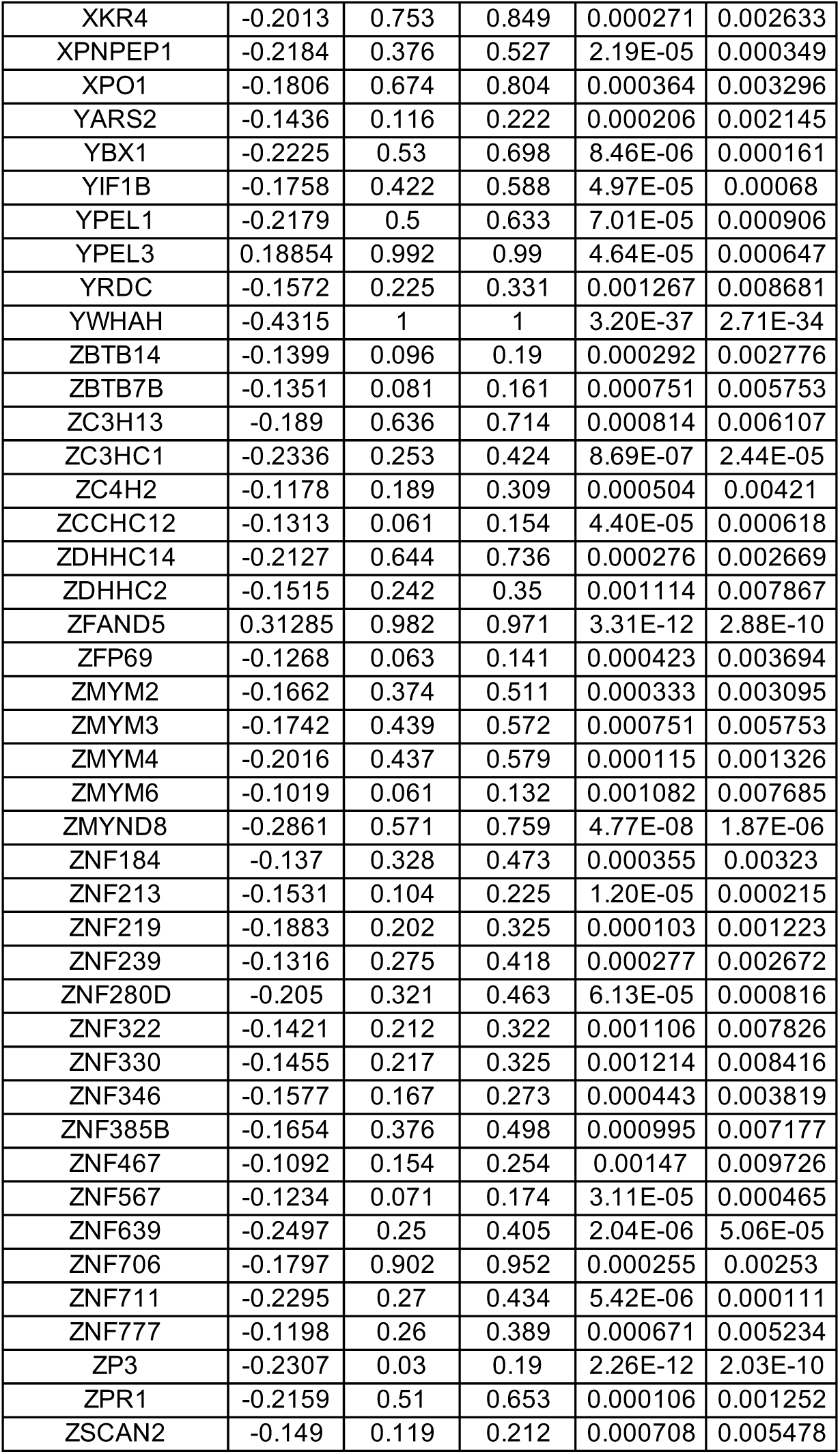
DEGs between wild-type (WT) and MECP2-null (KO) spots in Layer 2/3 in dPFC (adjusted p-value < 0.01)

**Table S39.**
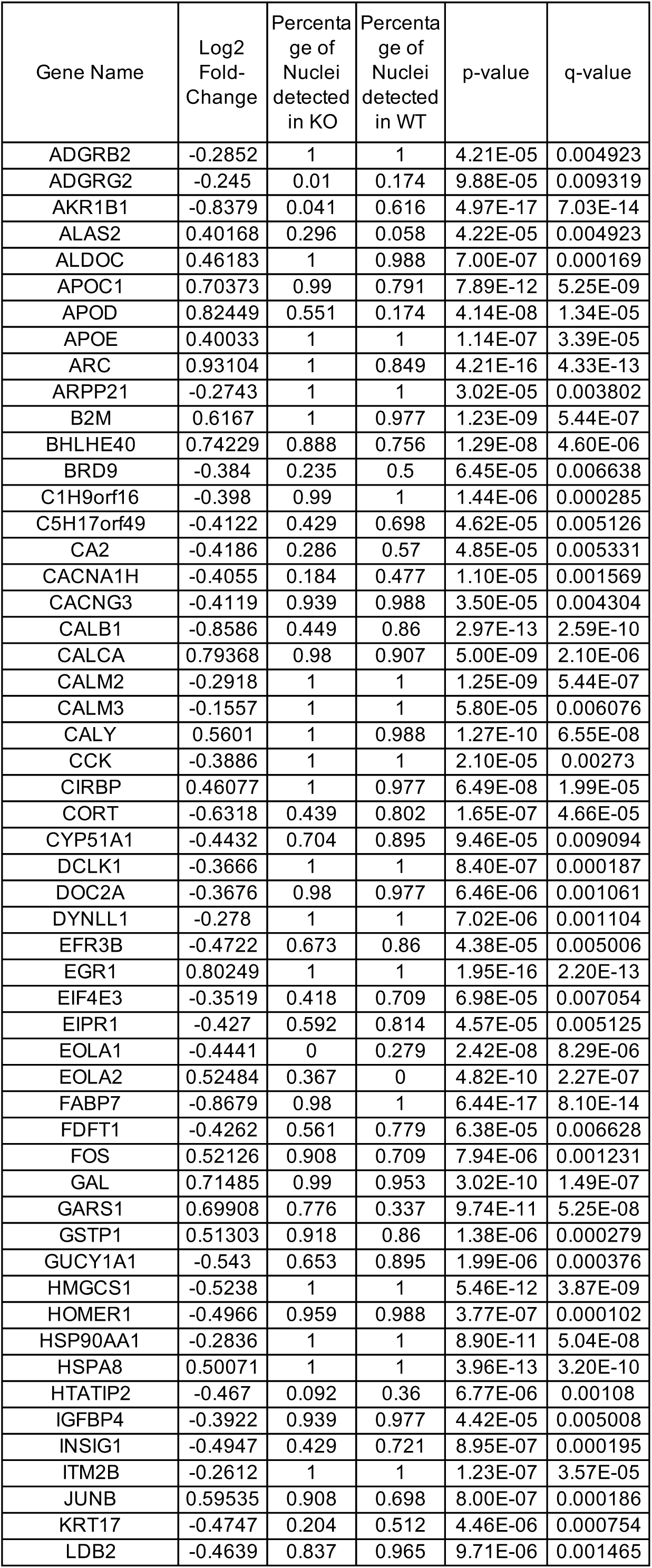
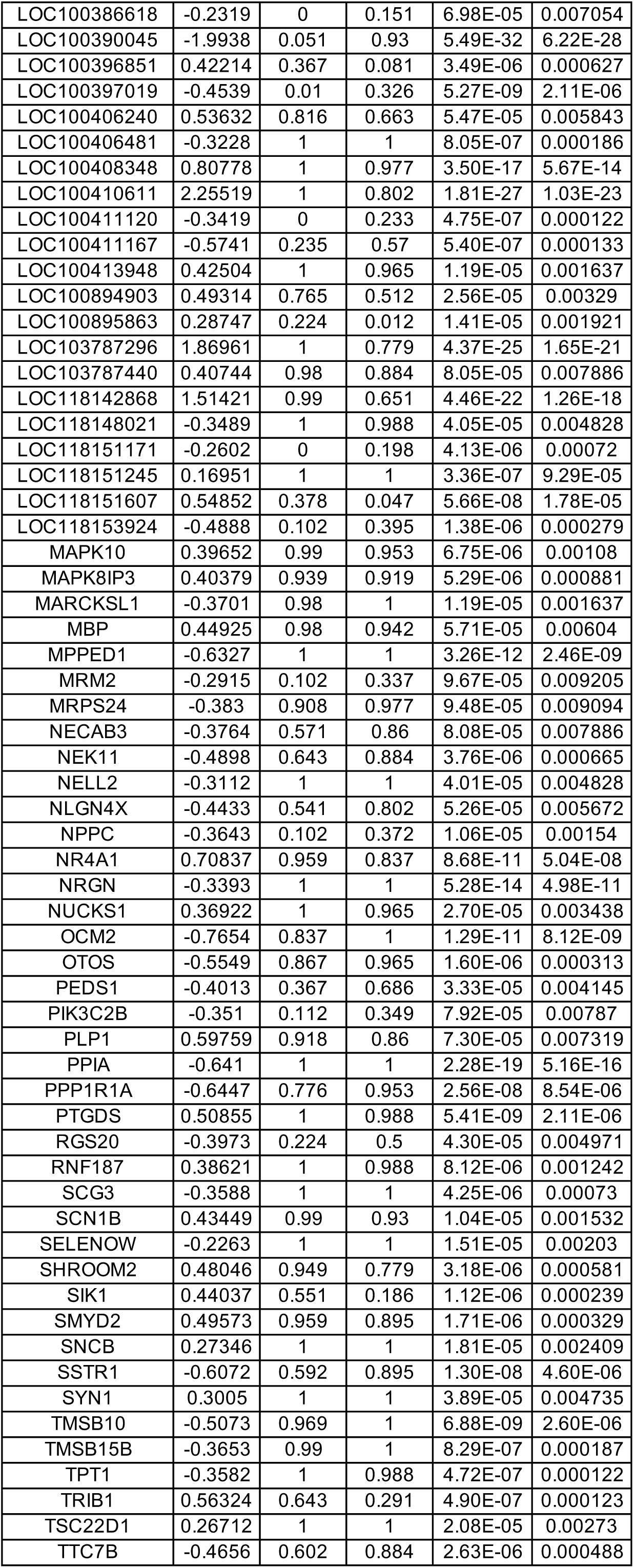
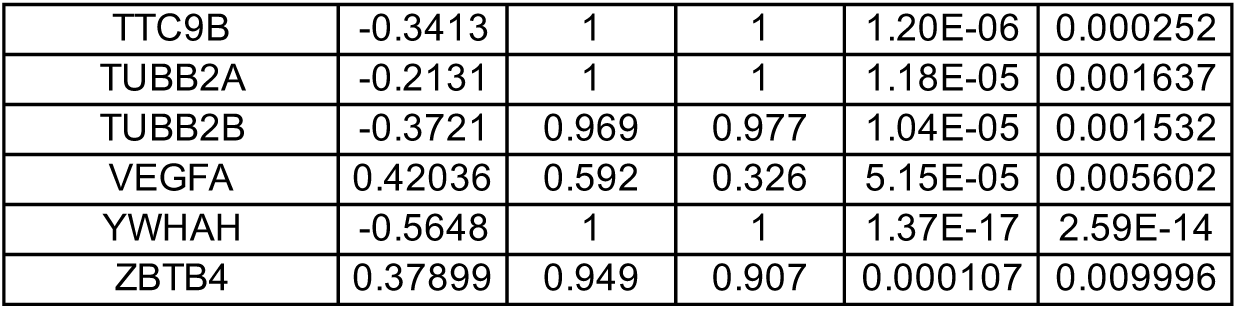
DEGs between wild-type (WT) and MECP2-null (KO) spots in Layer 4 in dPFC (adjusted p-value < 0.01)

**Table S40.**
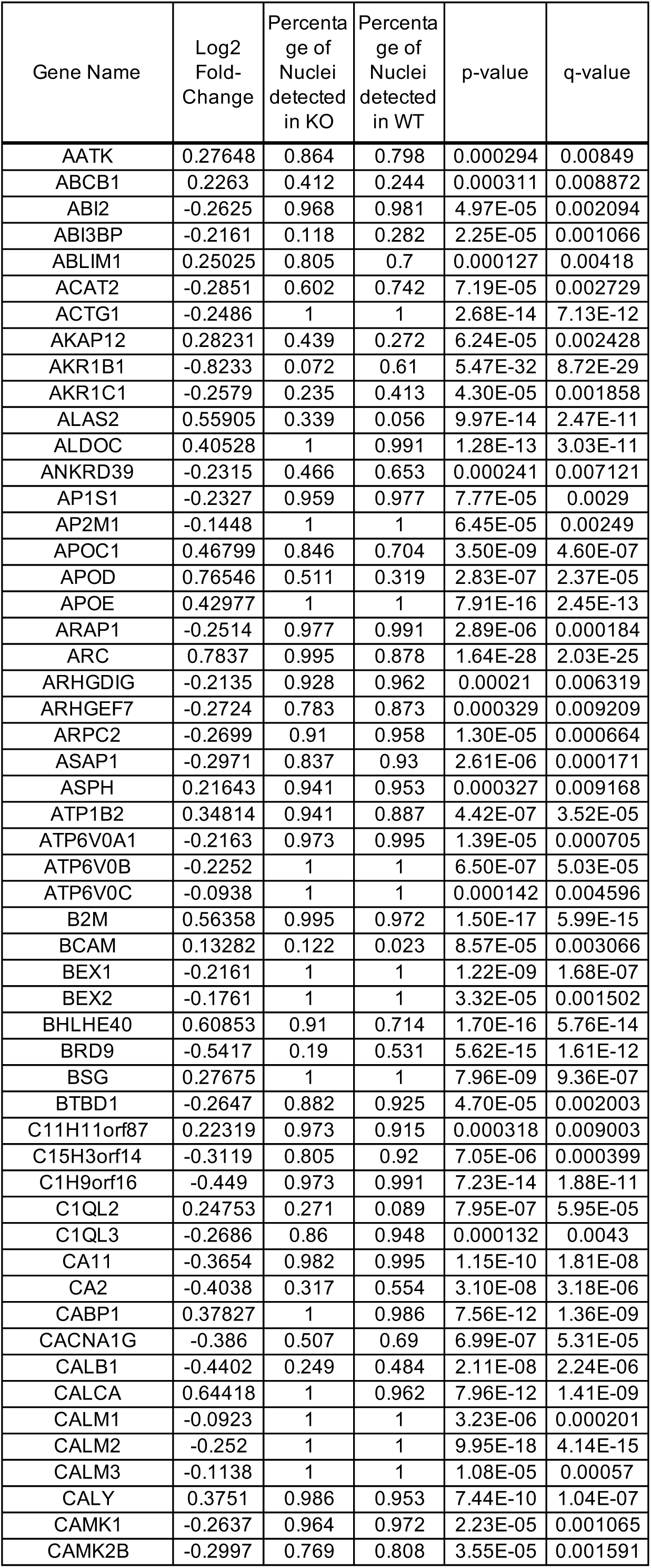
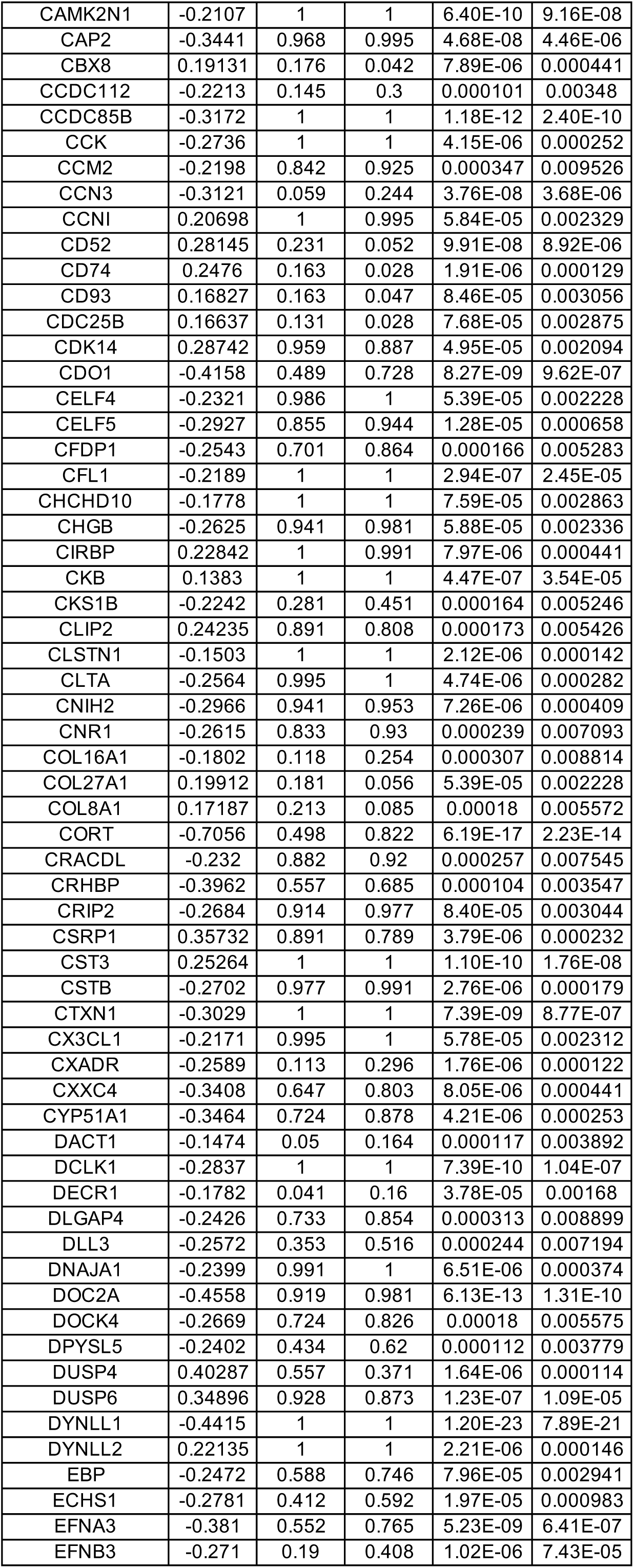
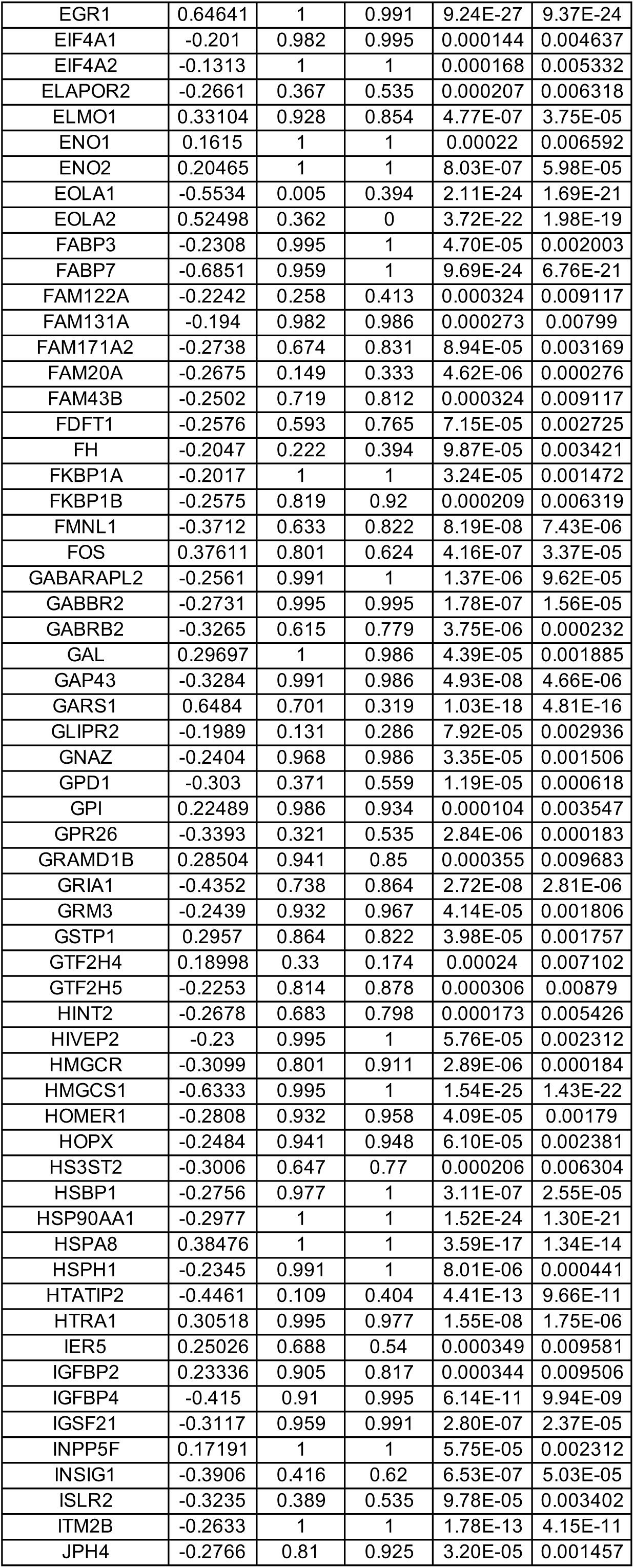
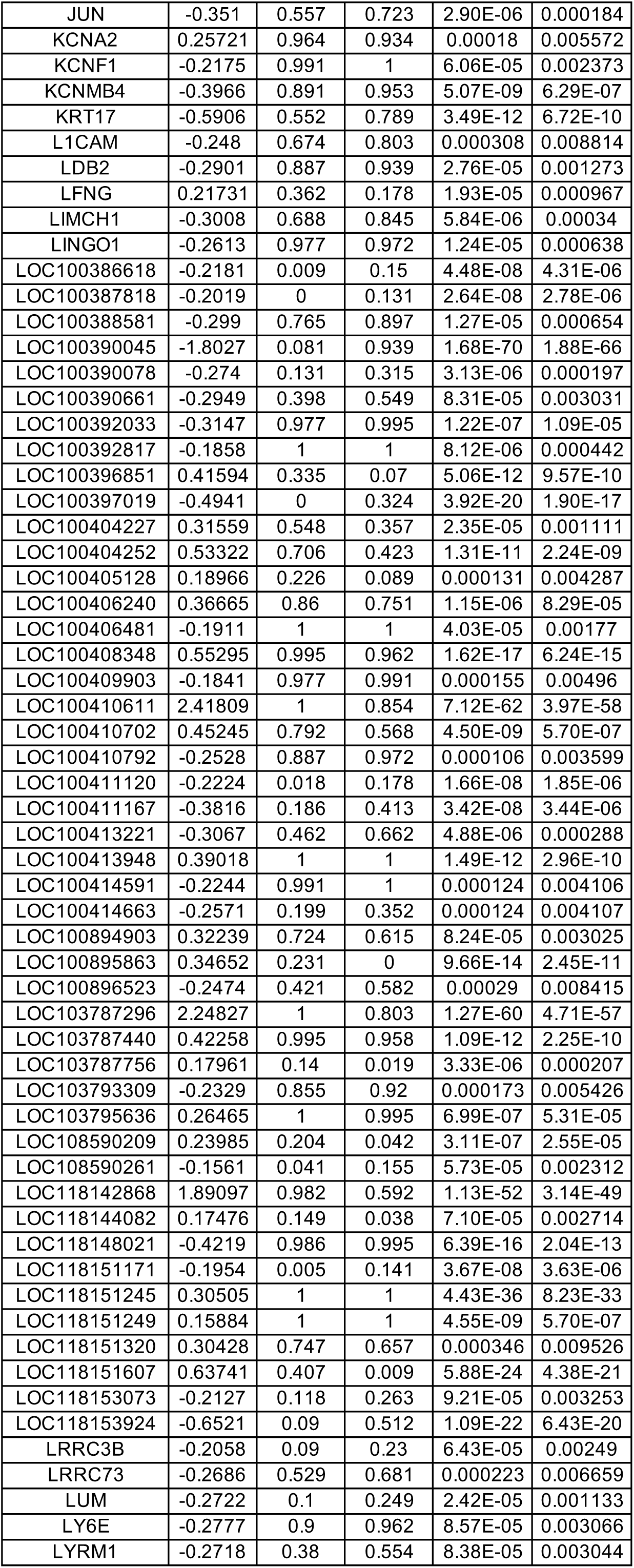
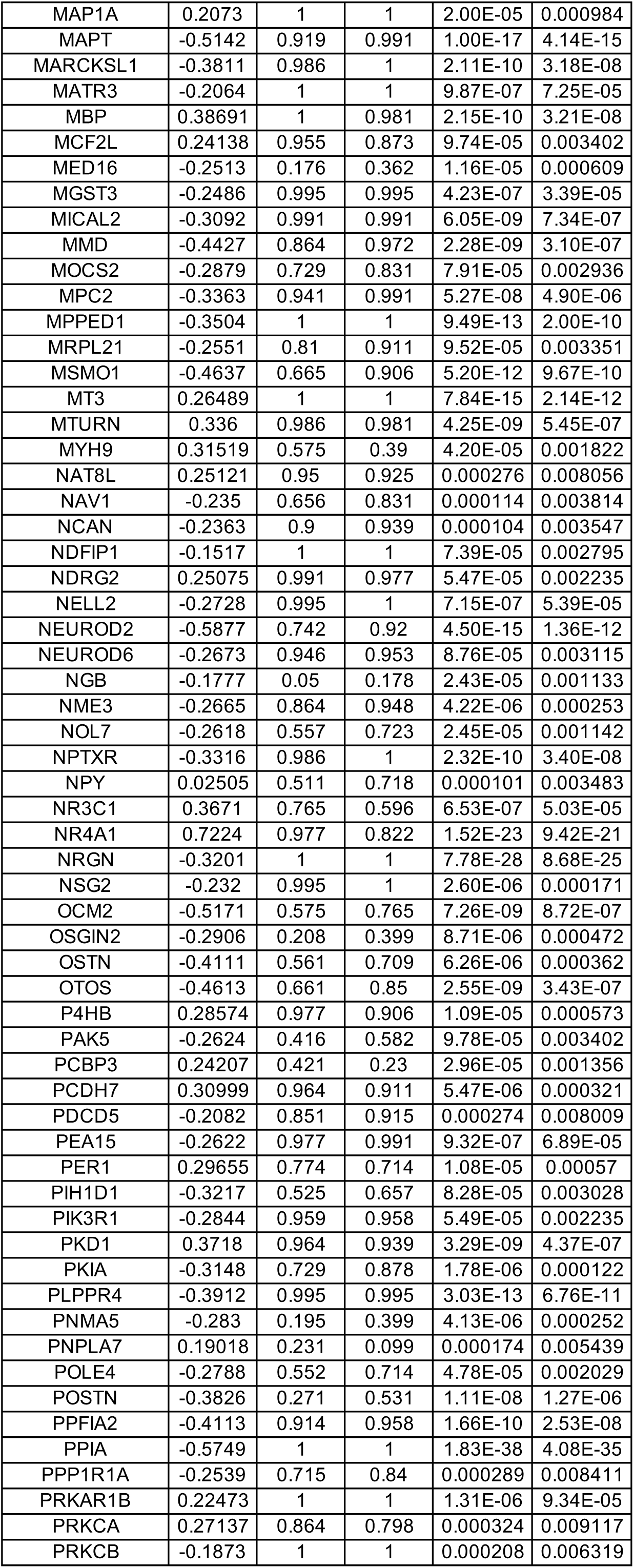
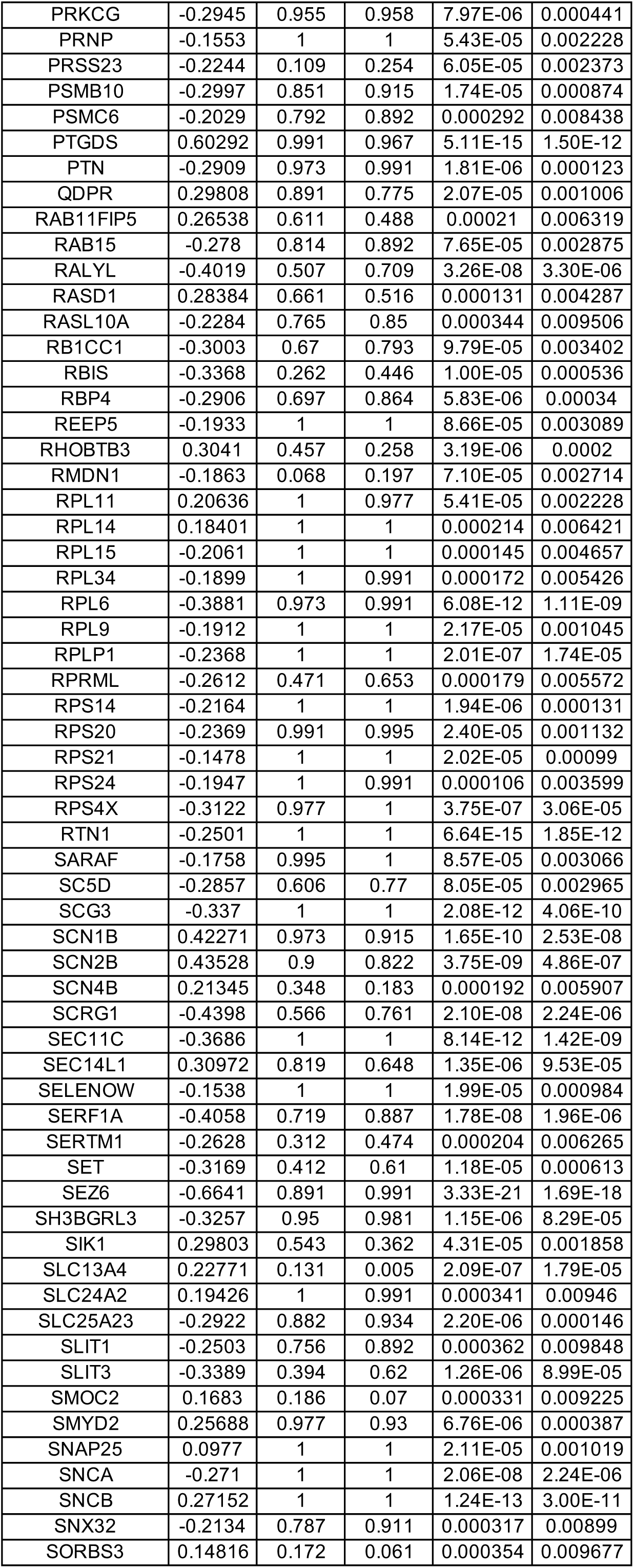
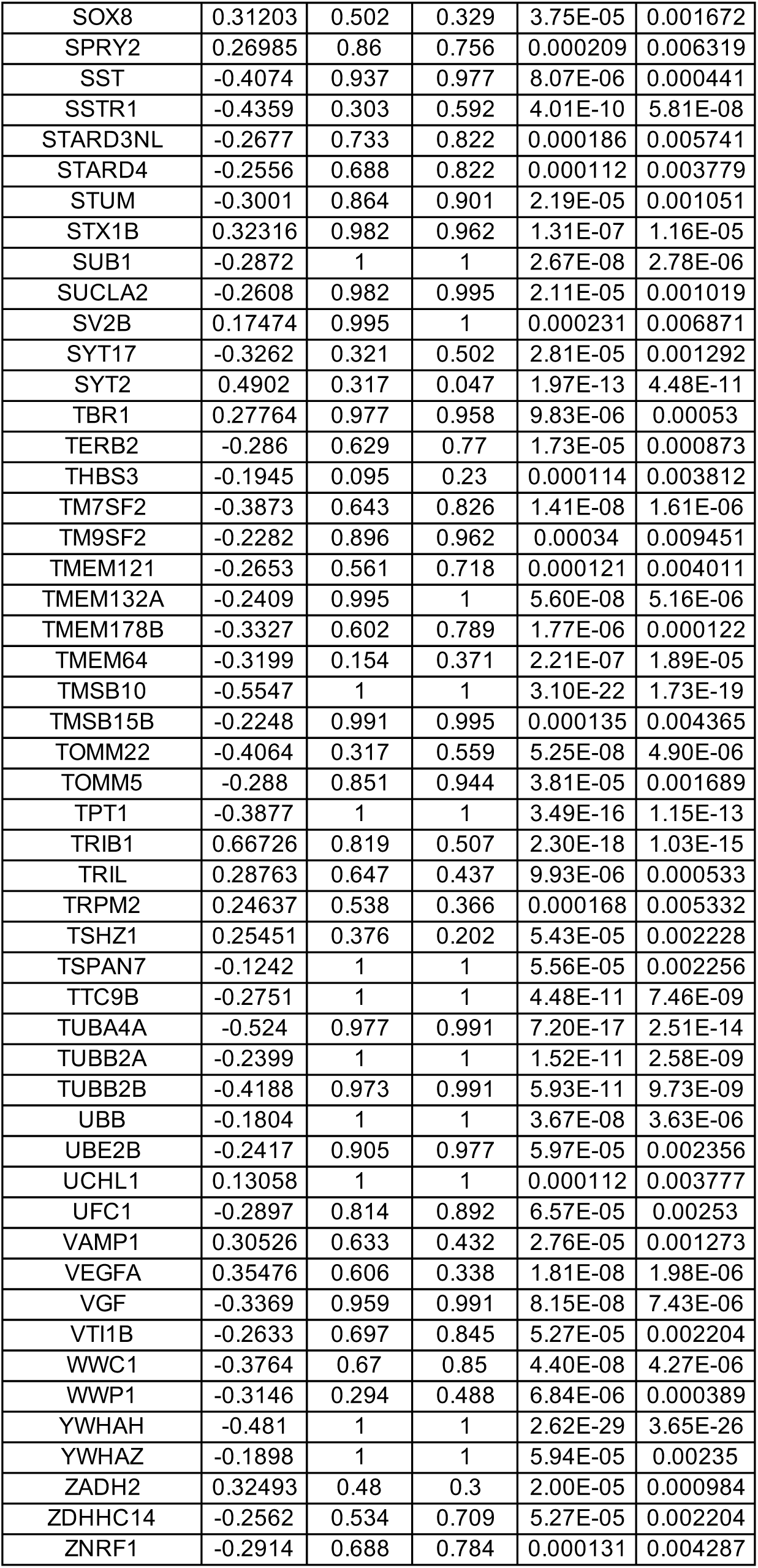
DEGs between wild-type (WT) and MECP2-null (KO) spots in Layer 5 in dPFC (adjusted p-value < 0.01)

**Table S41.**
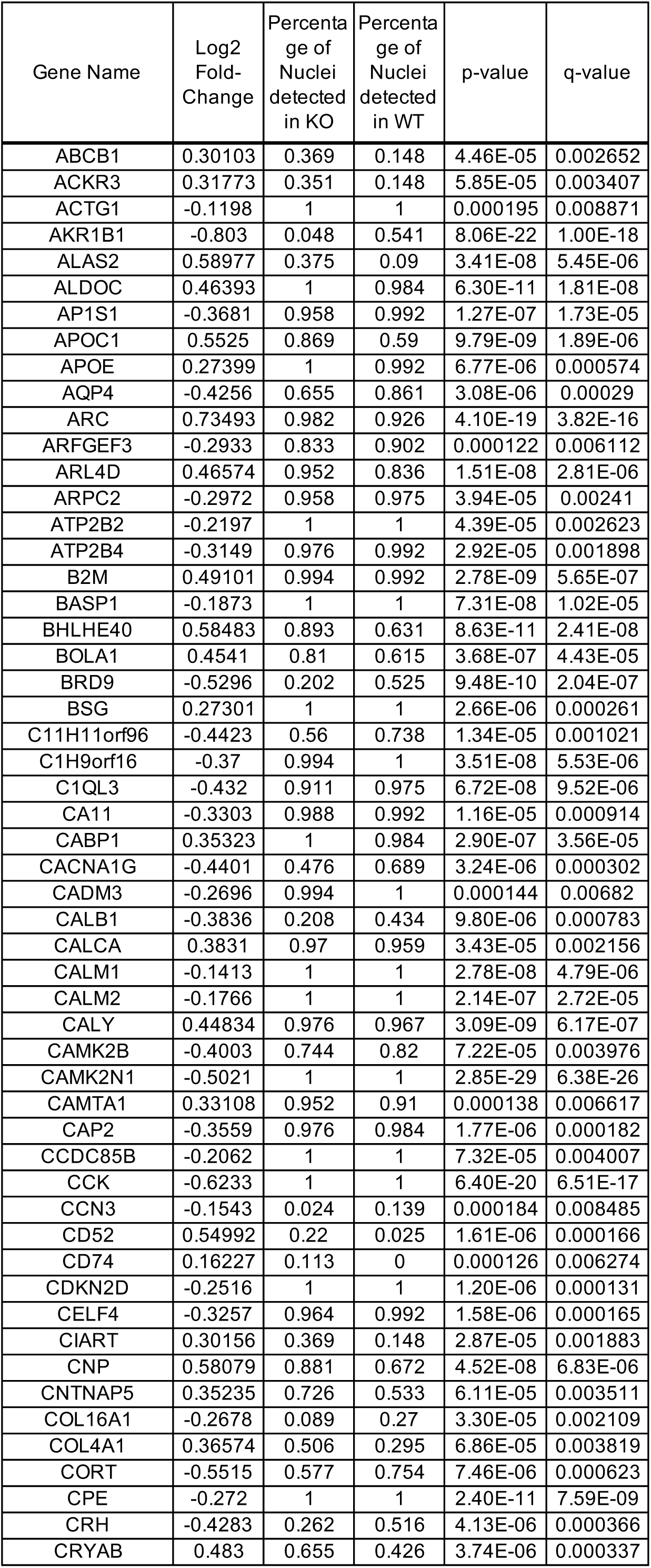
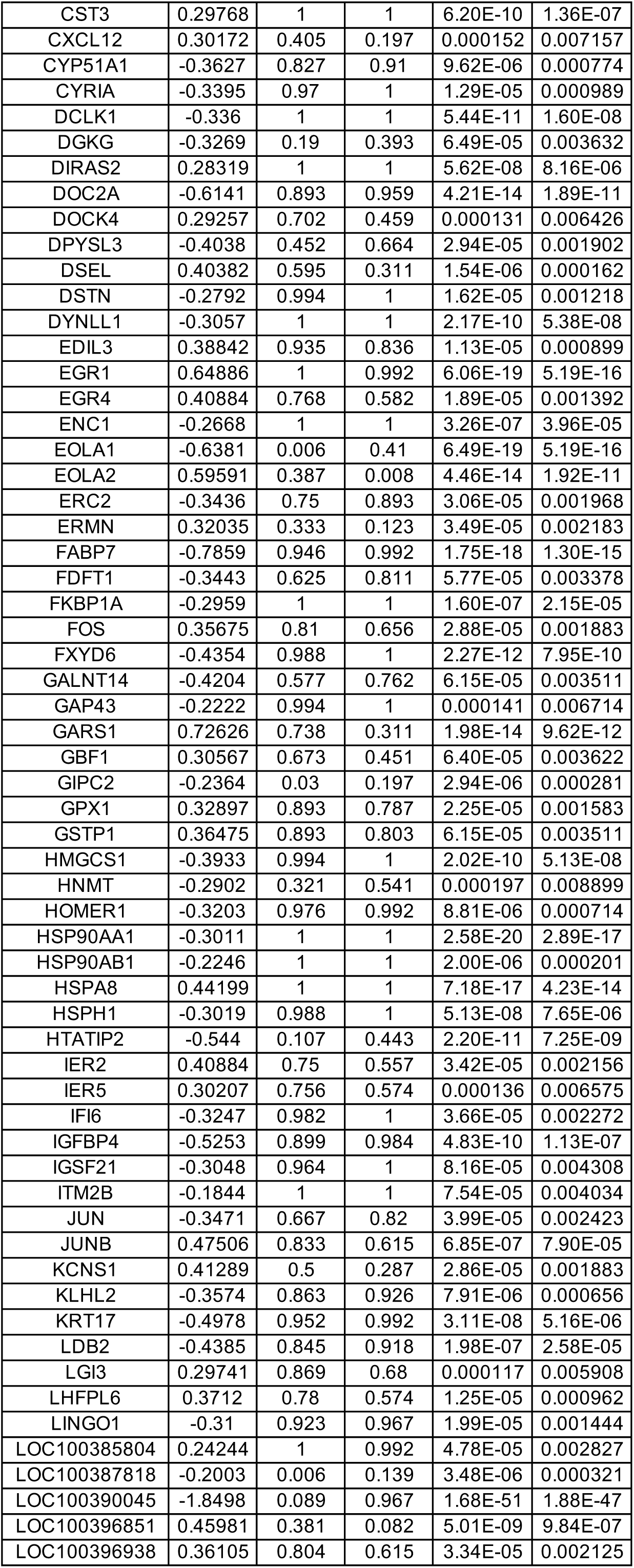
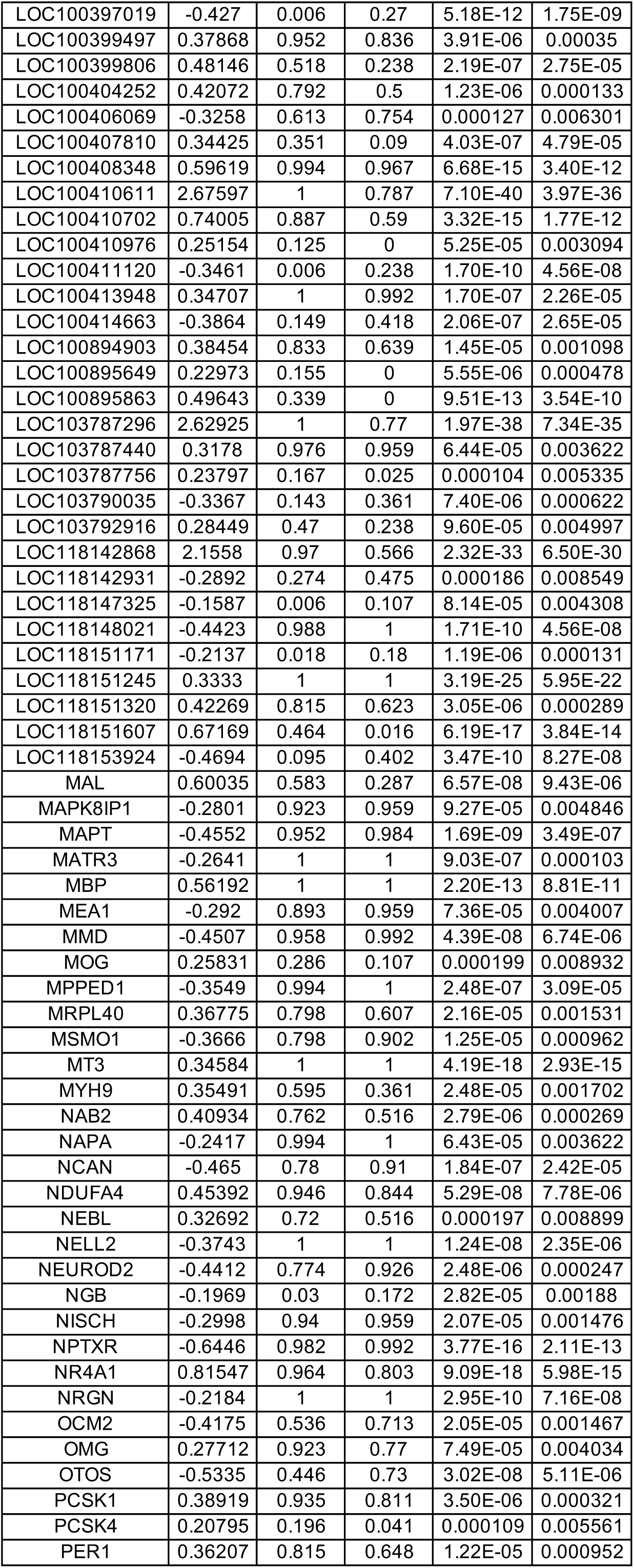
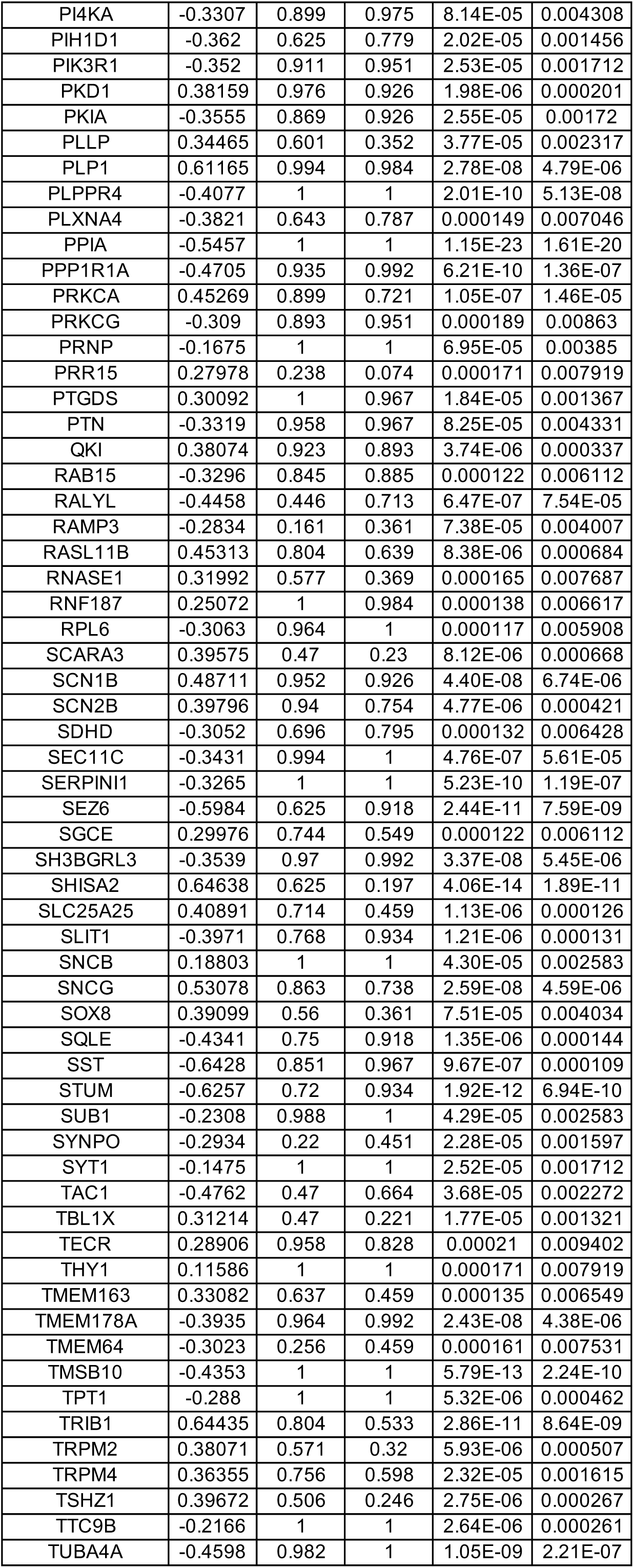
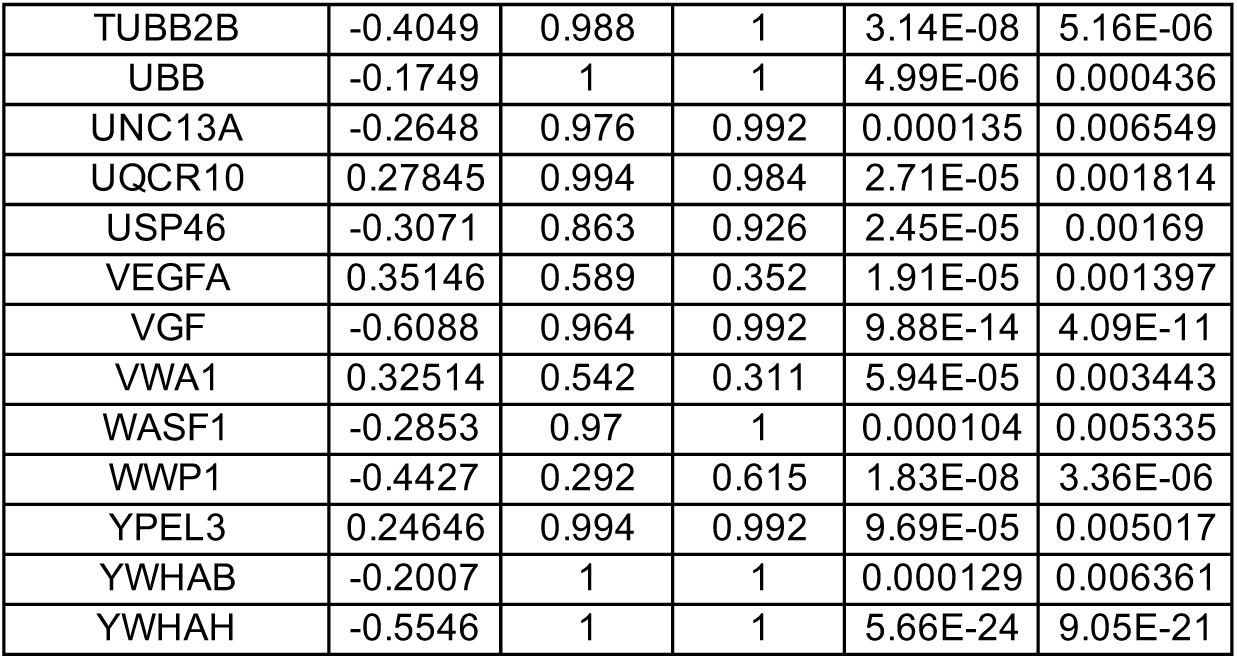
DEGs between wild-type (WT) and MECP2-null (KO) spots in Layer 6 in dPFC (adjusted p-value < 0.01)

**Table S42.**
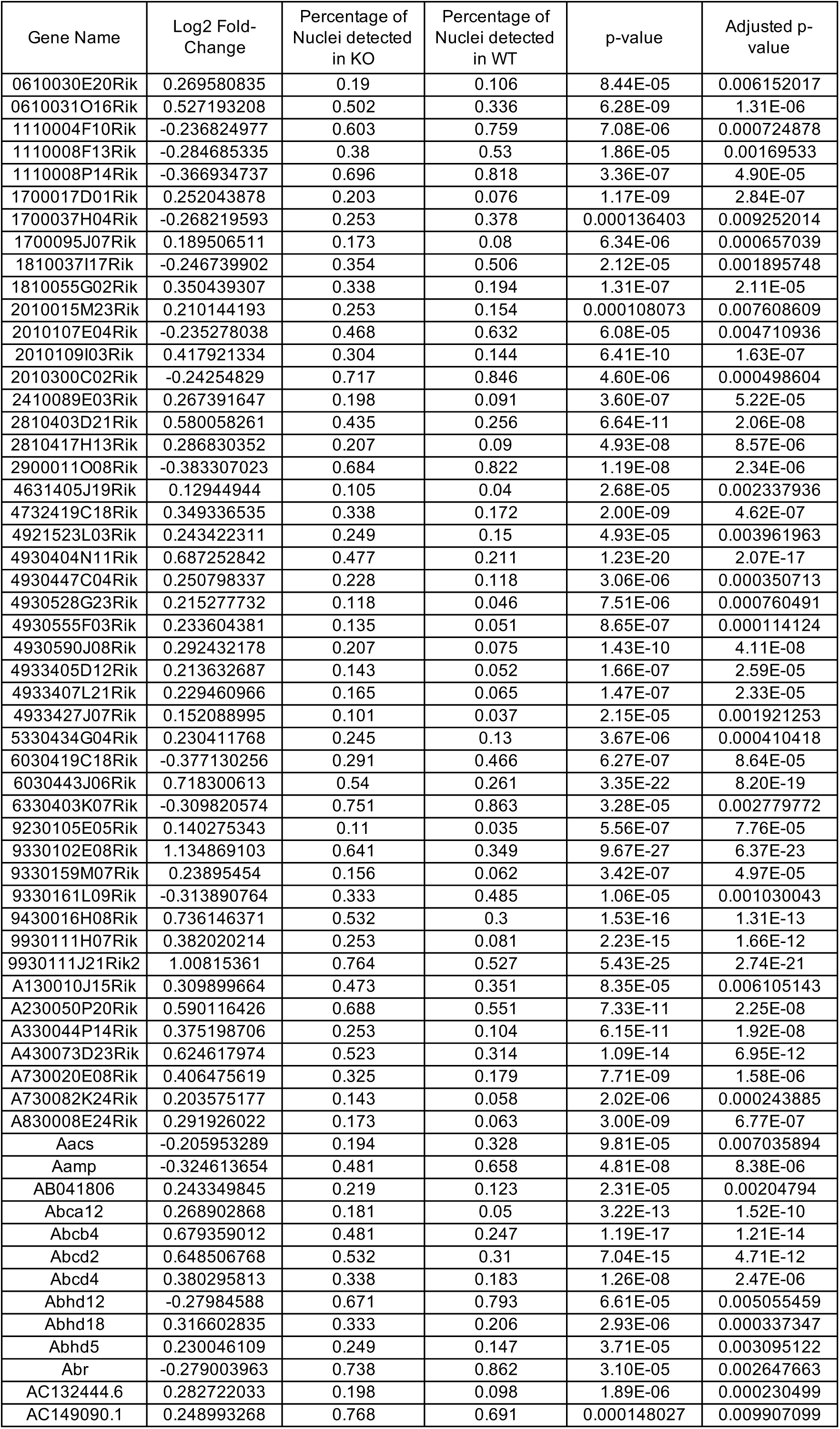
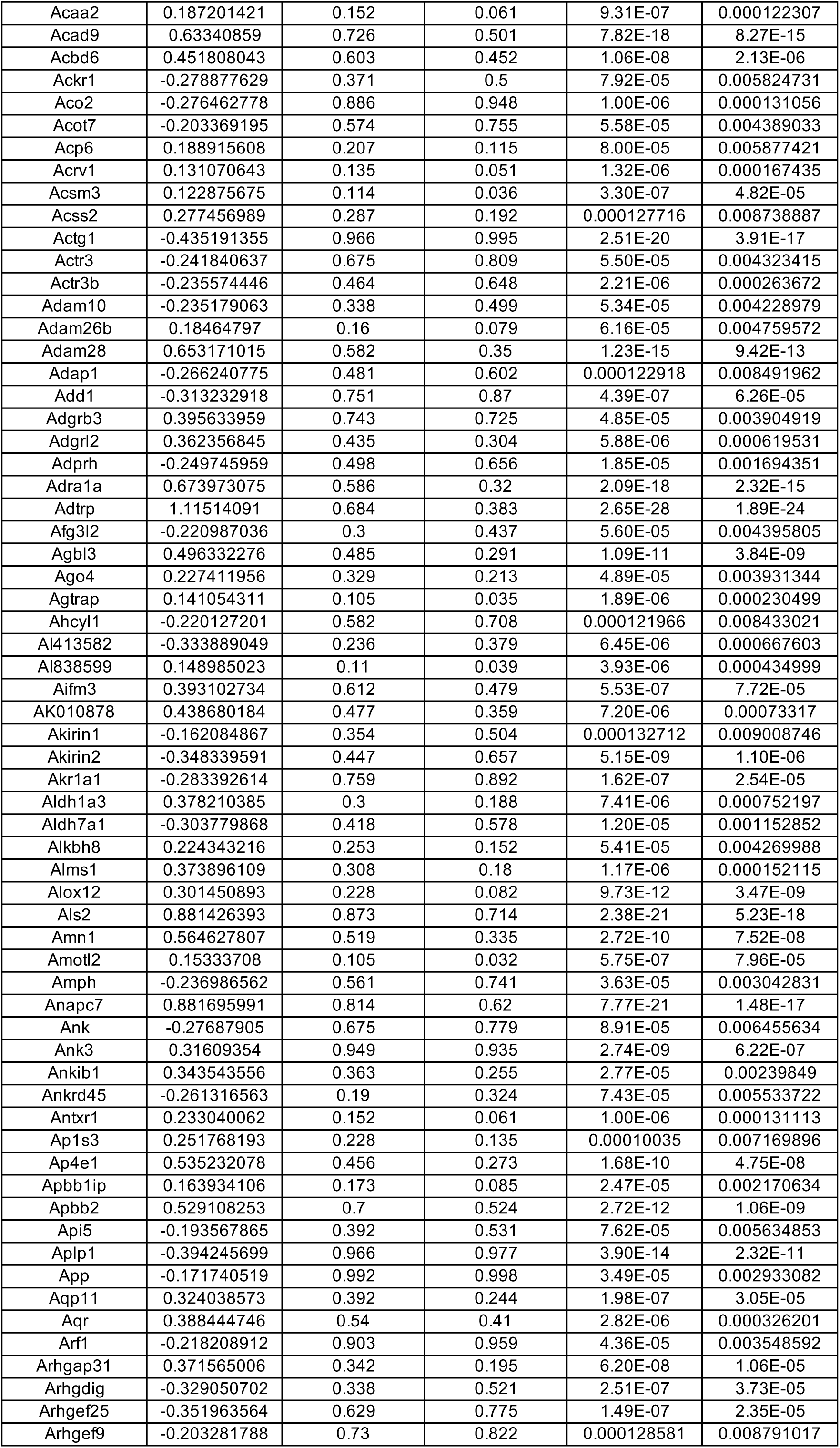
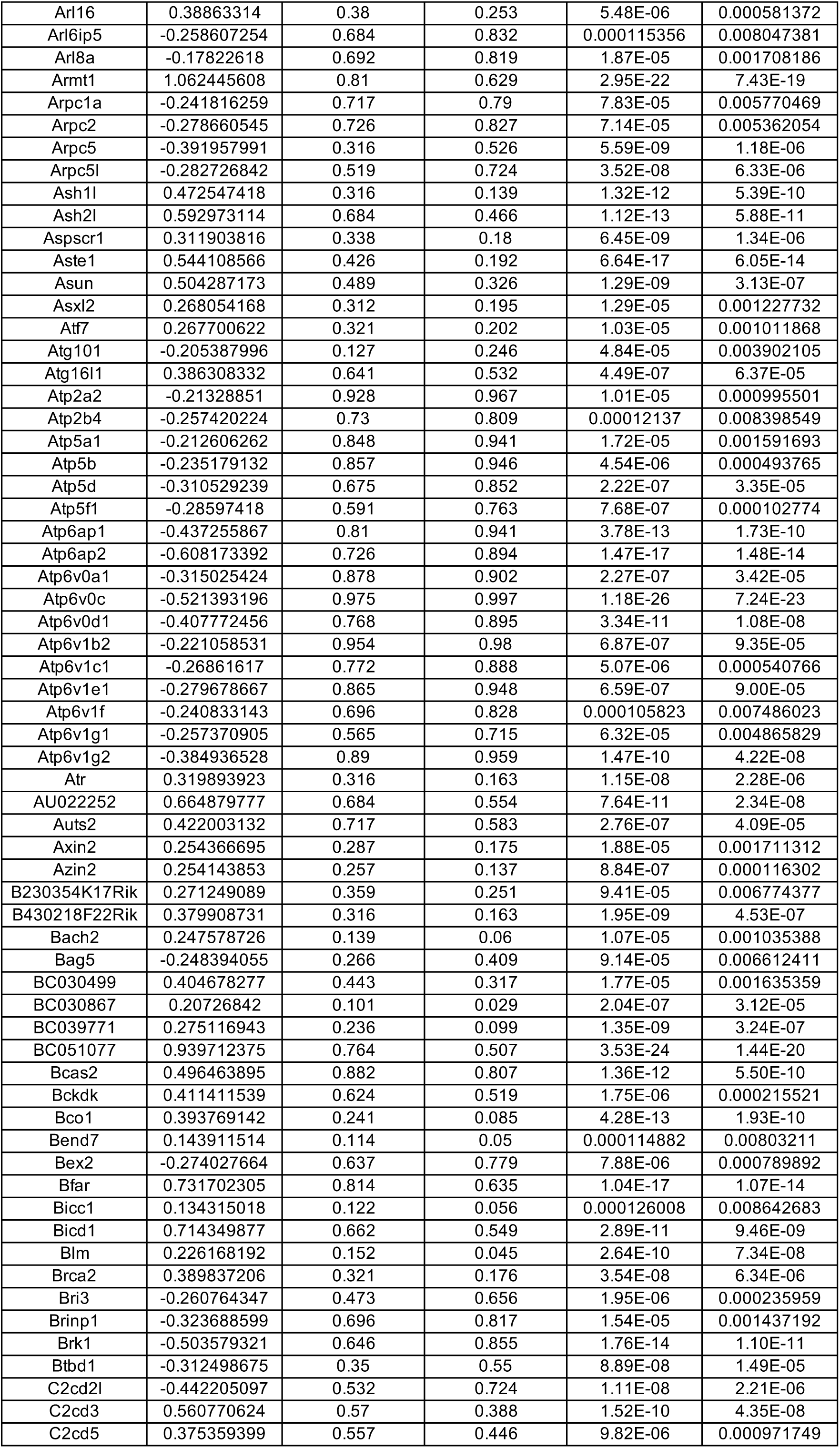
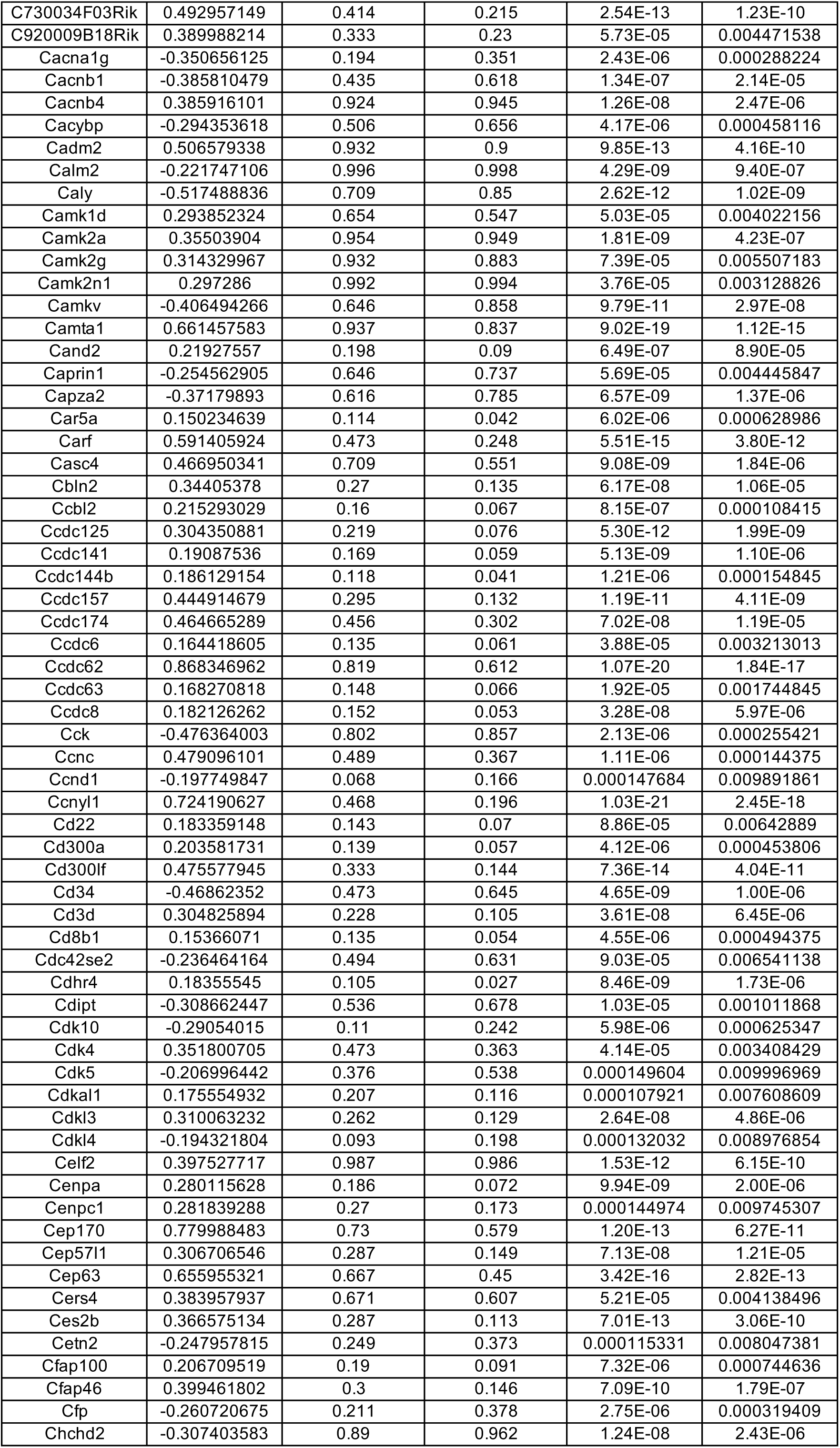
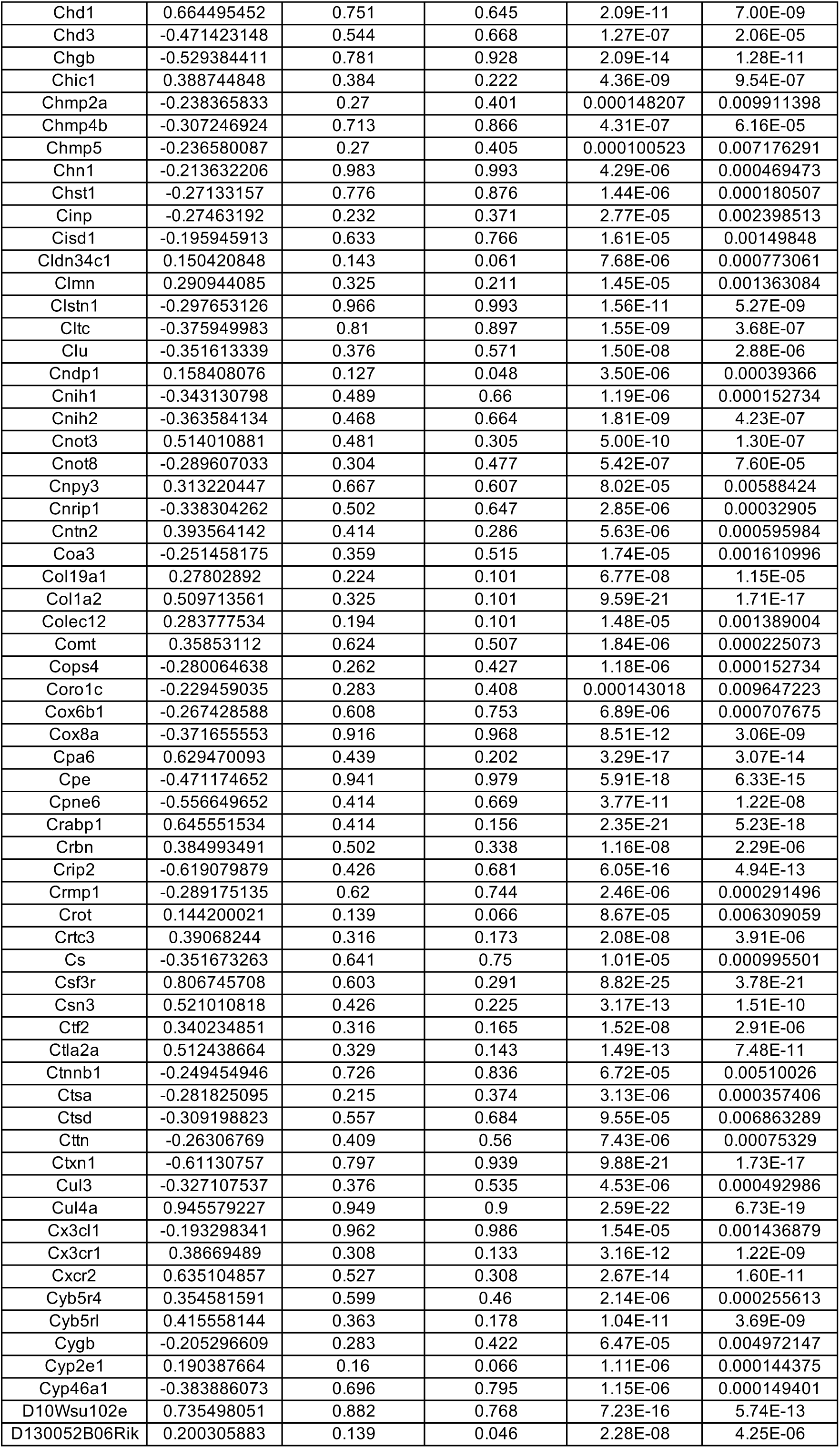
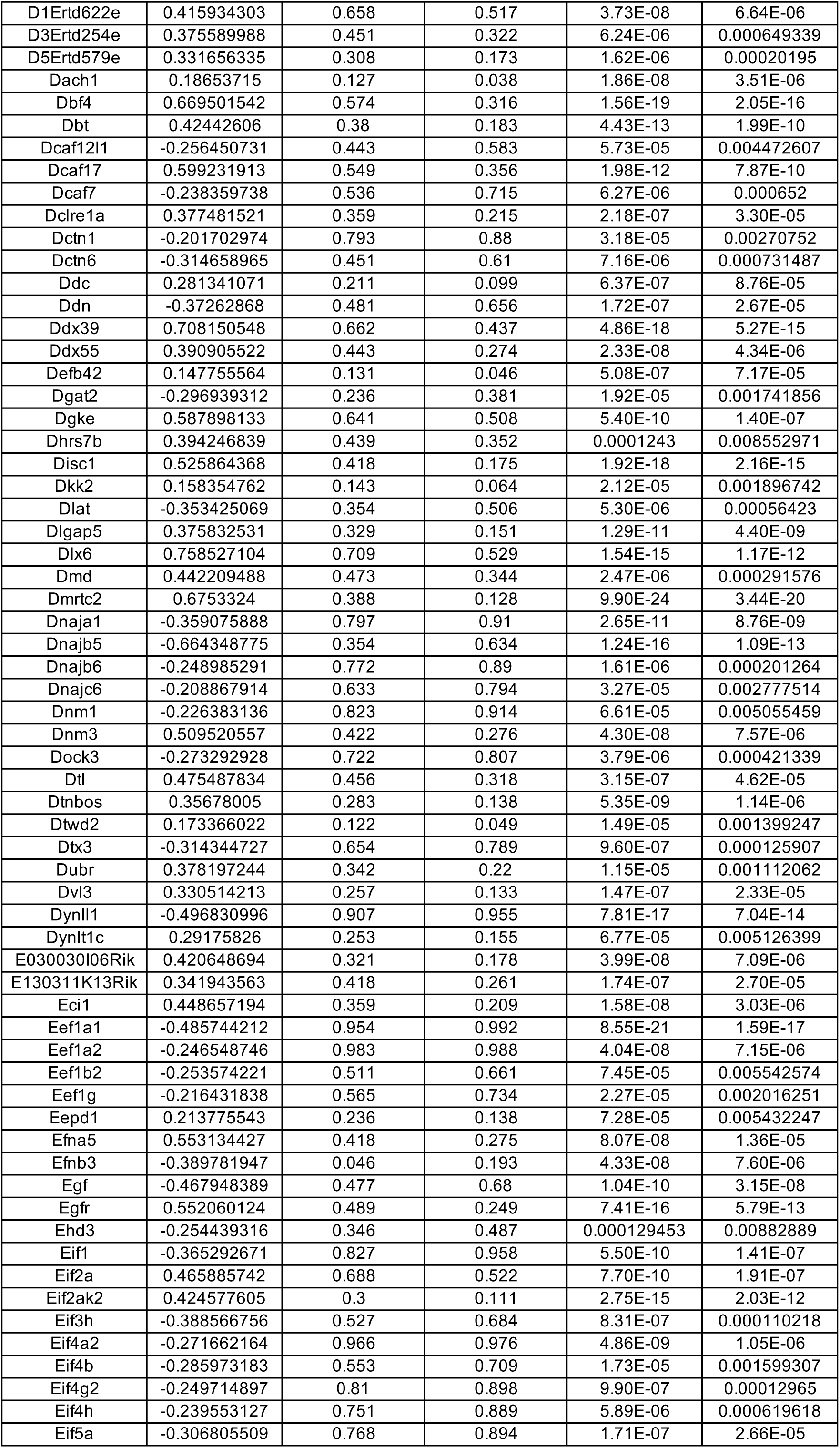
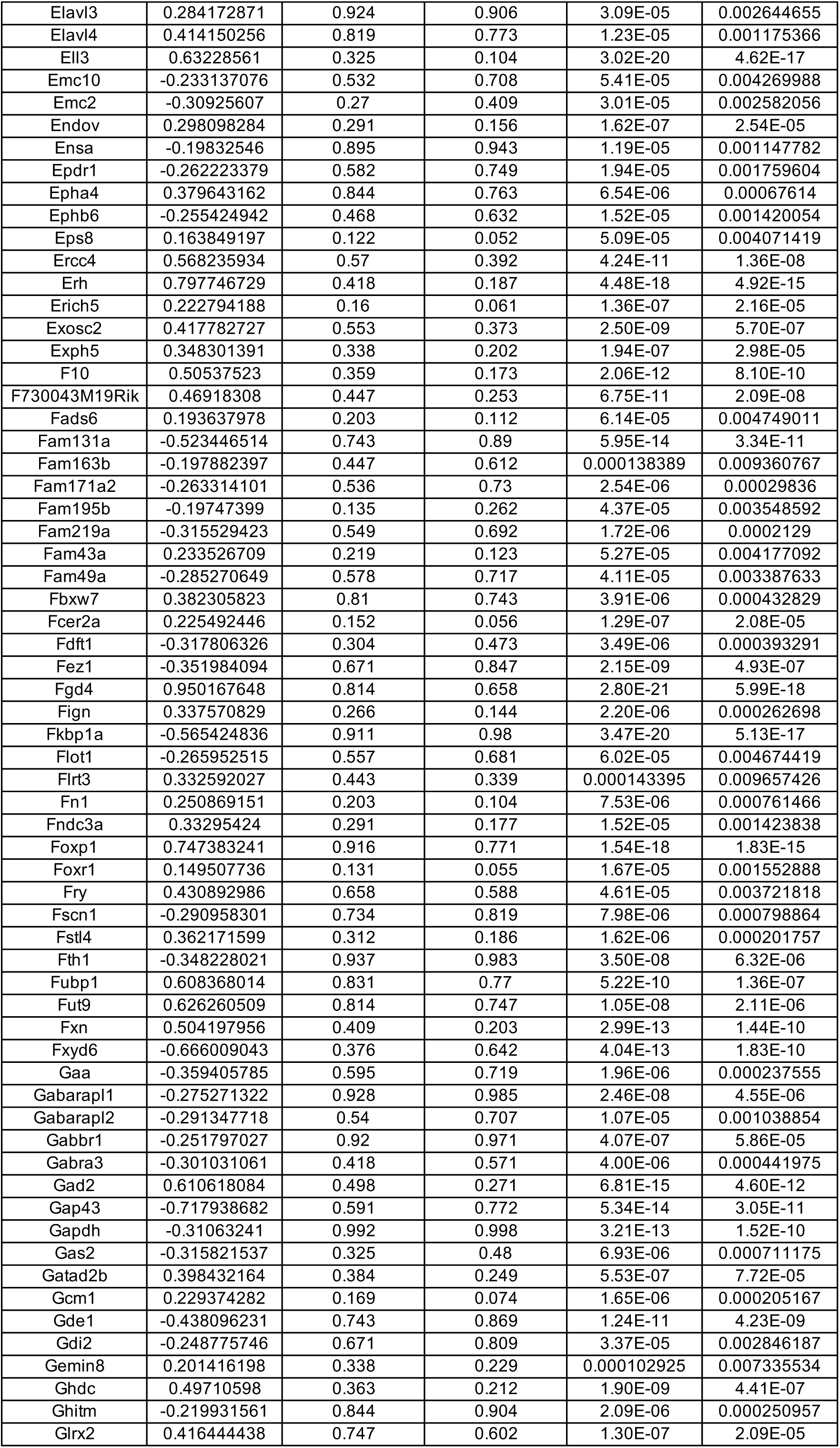
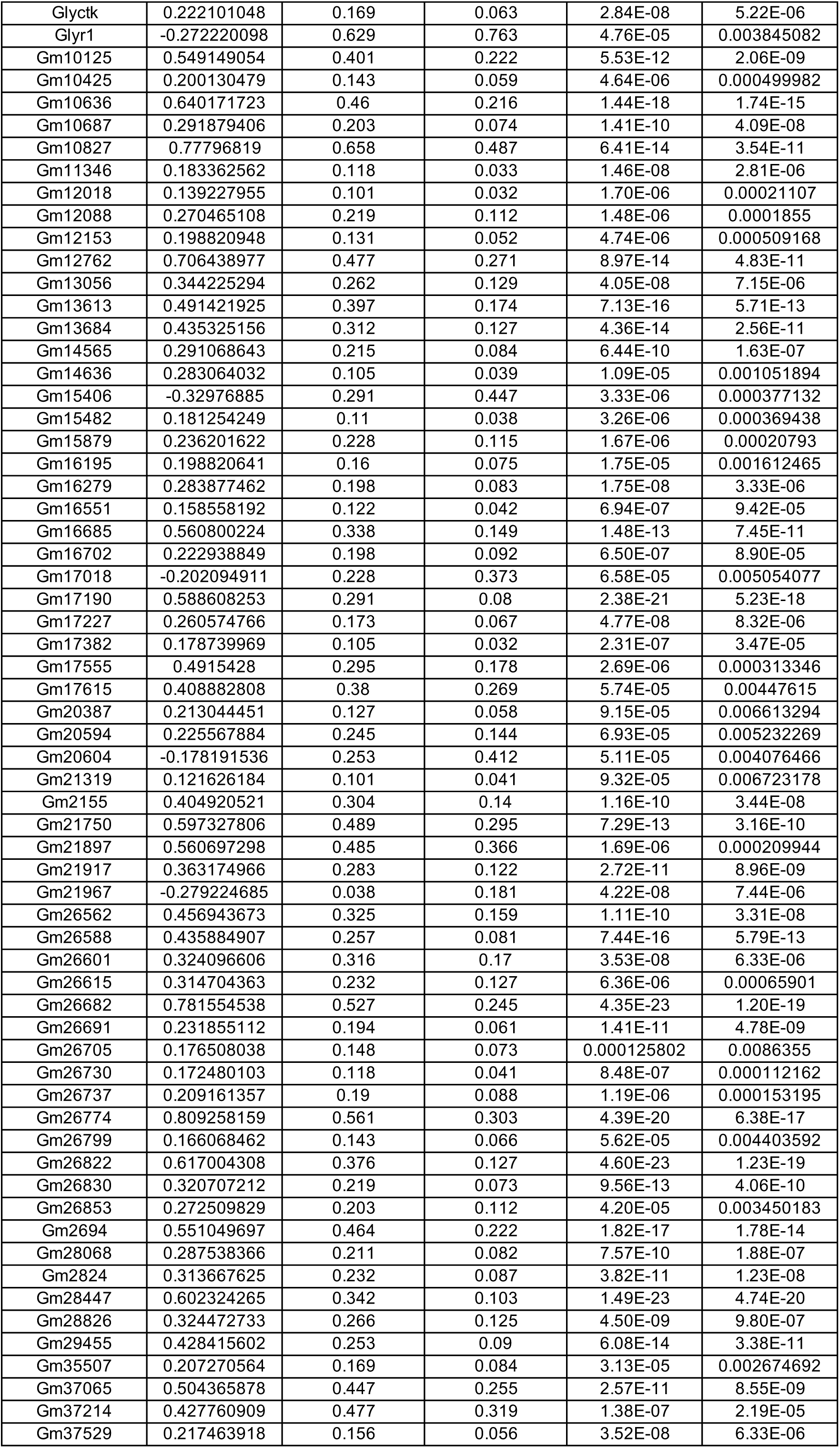
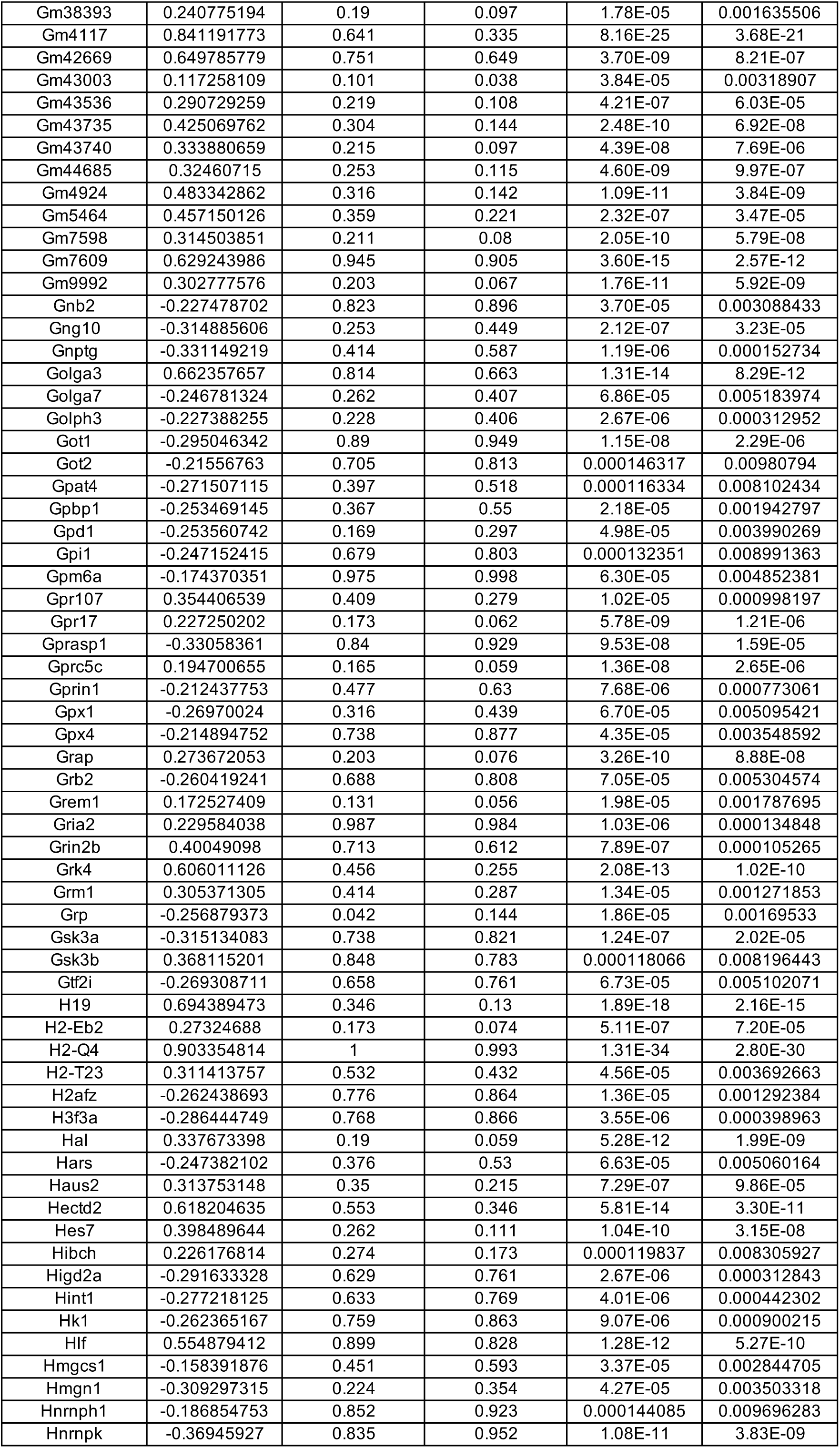
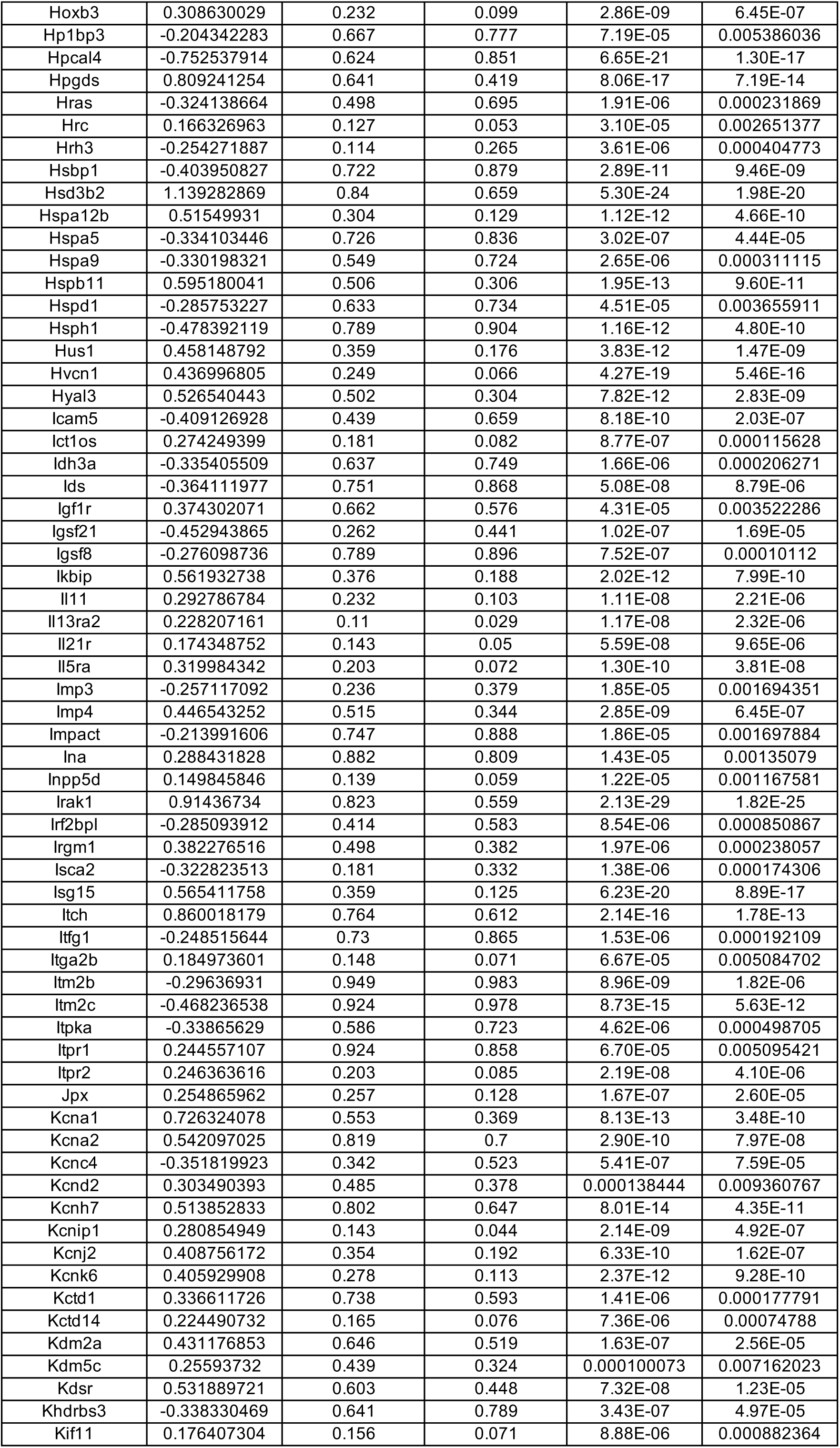
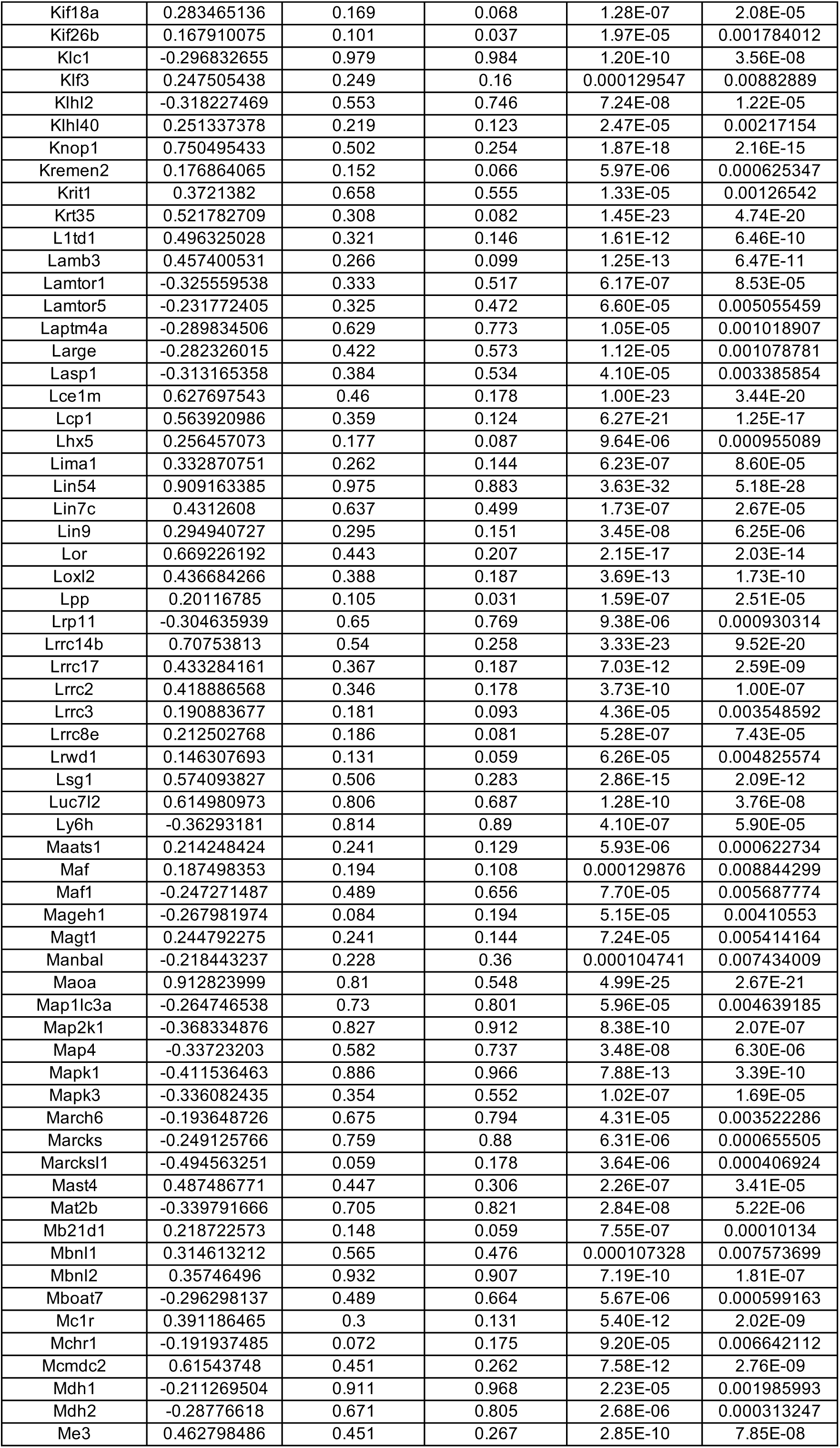
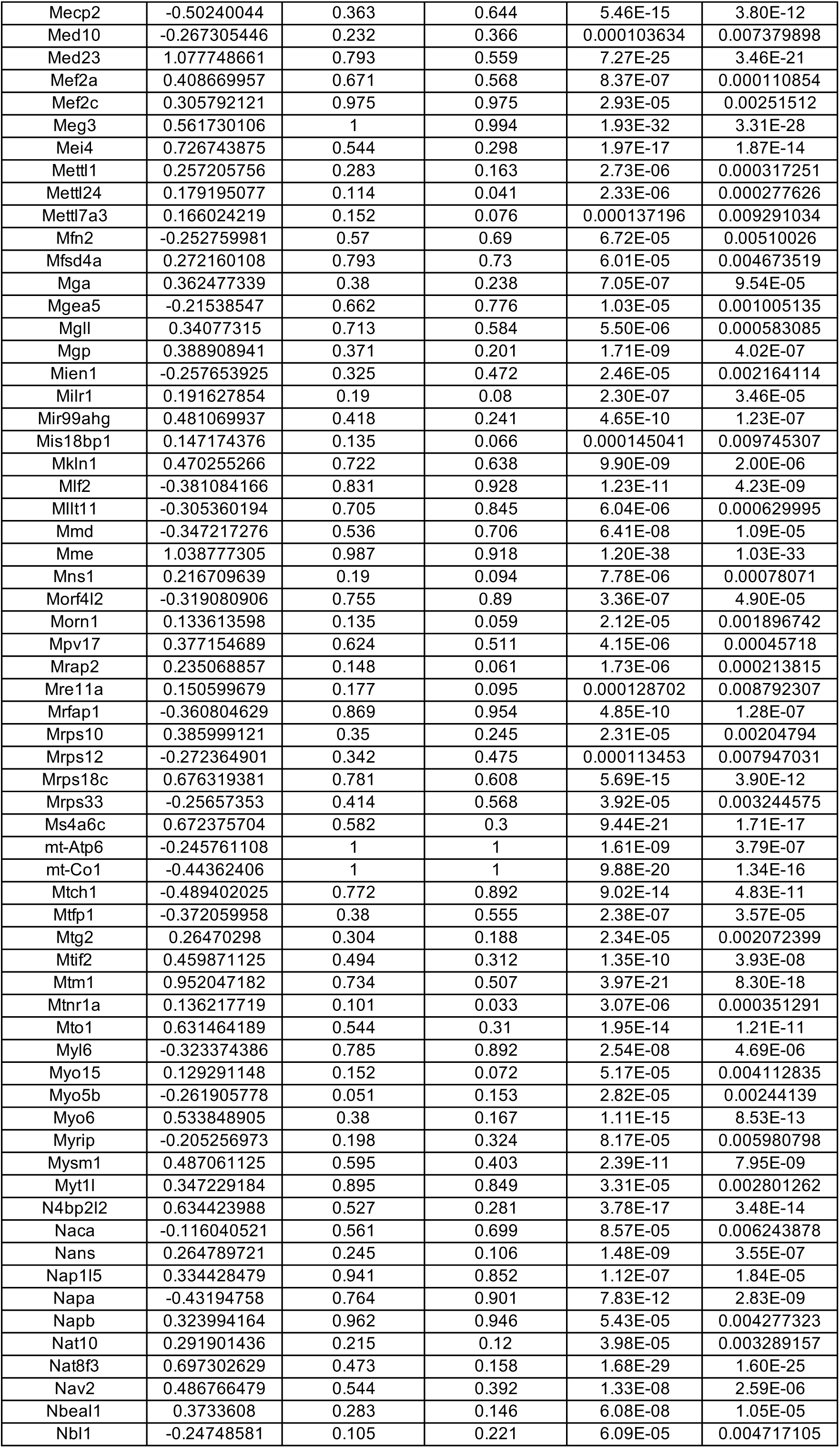
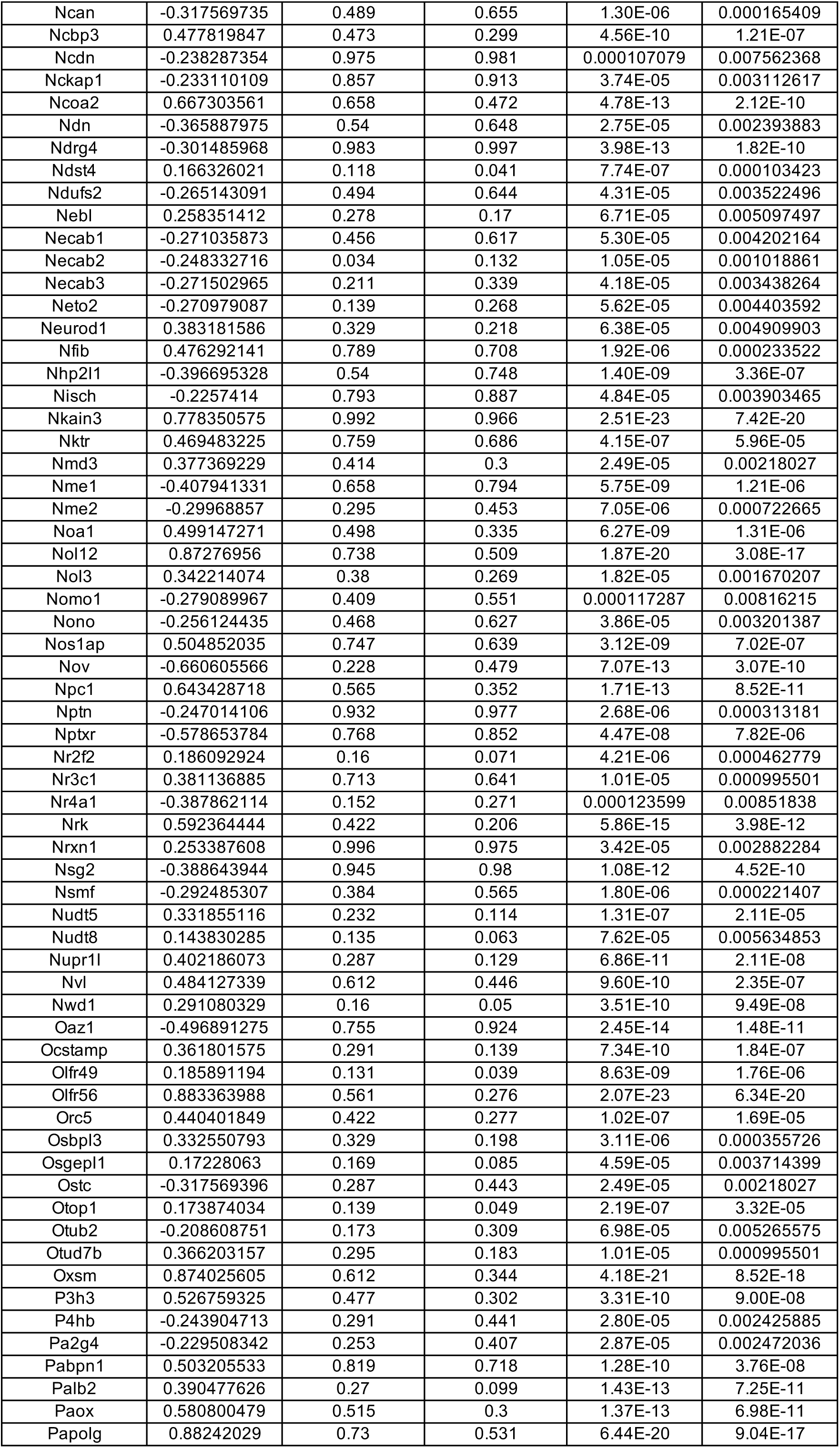
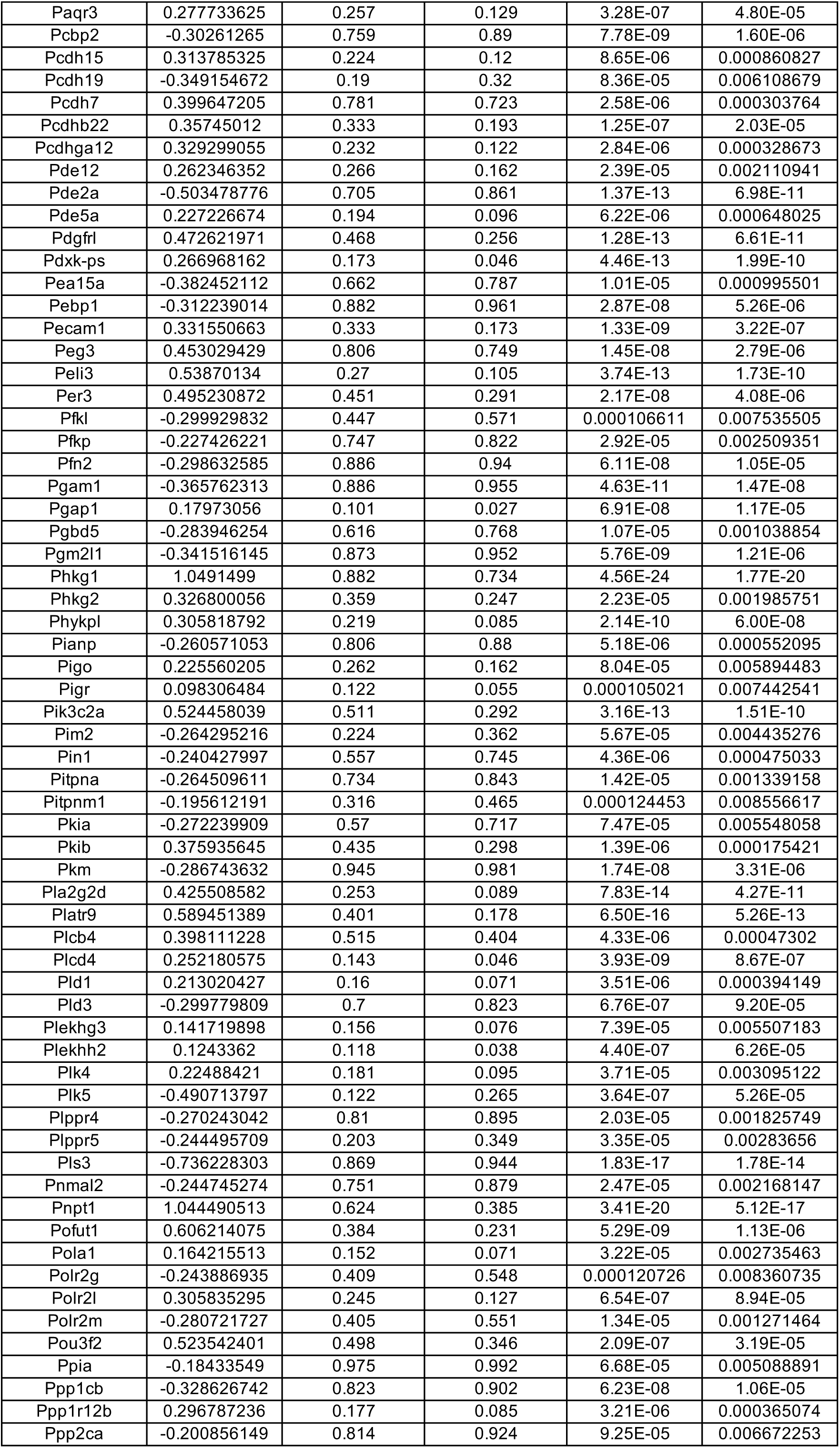
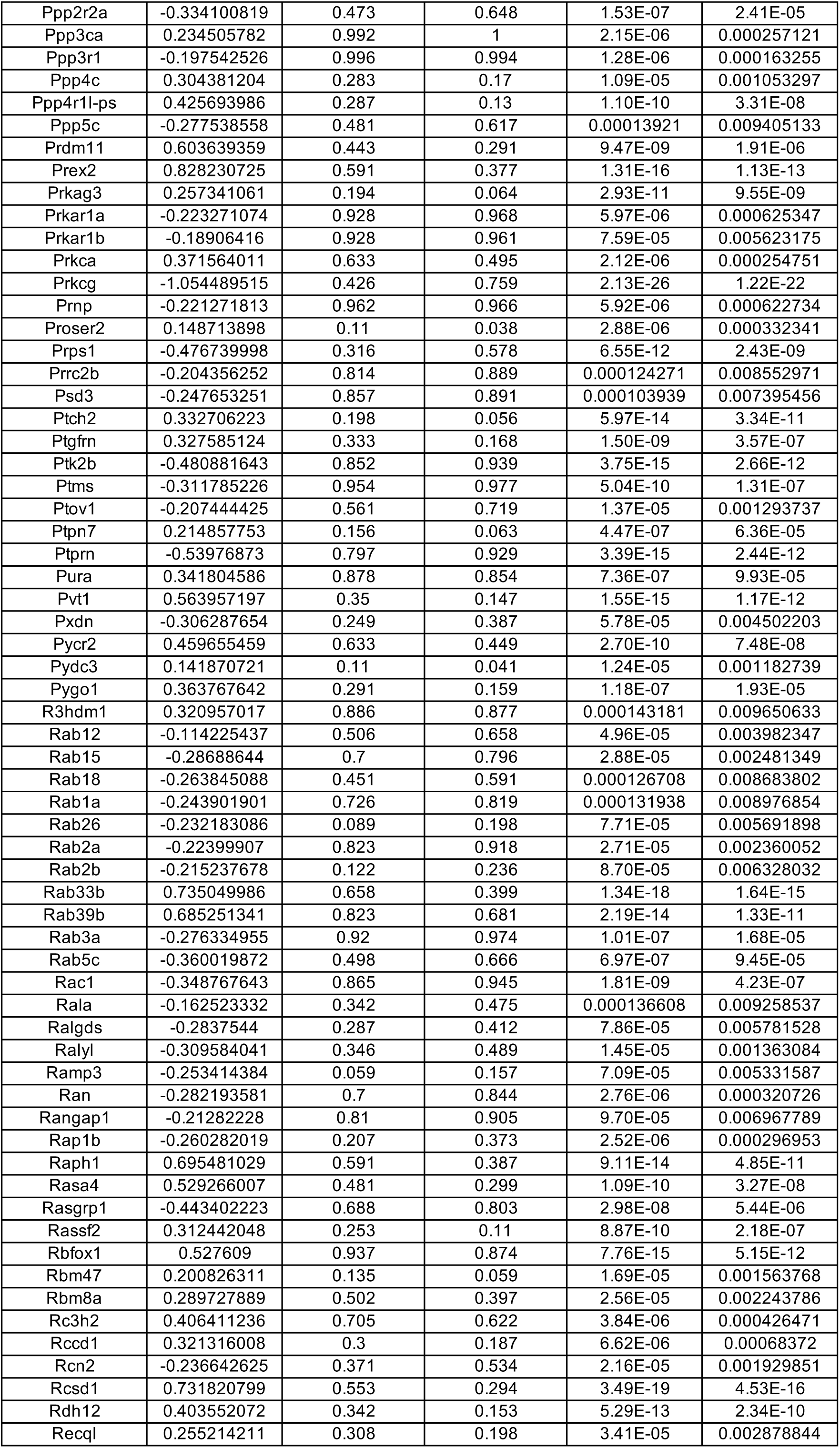
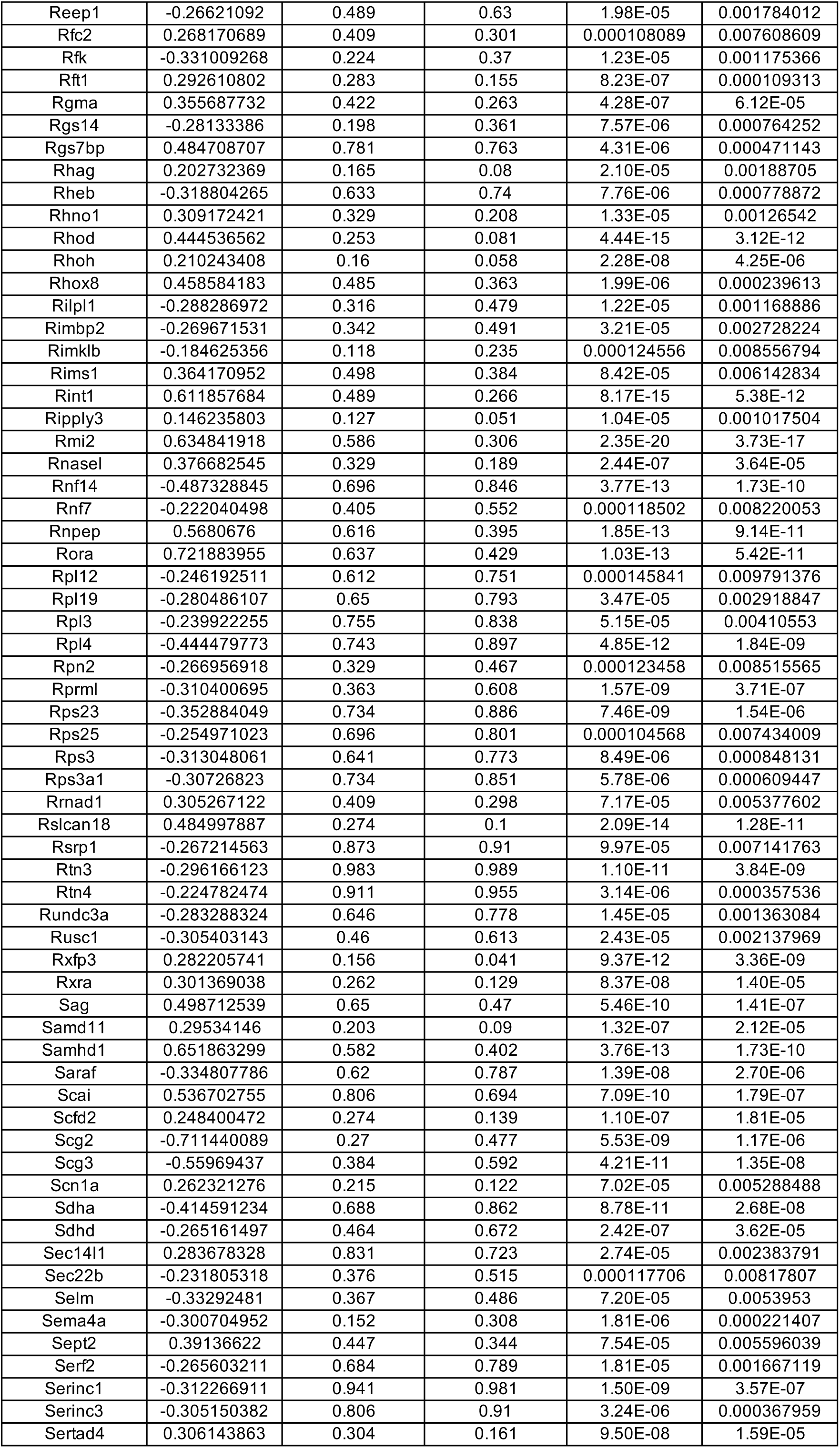
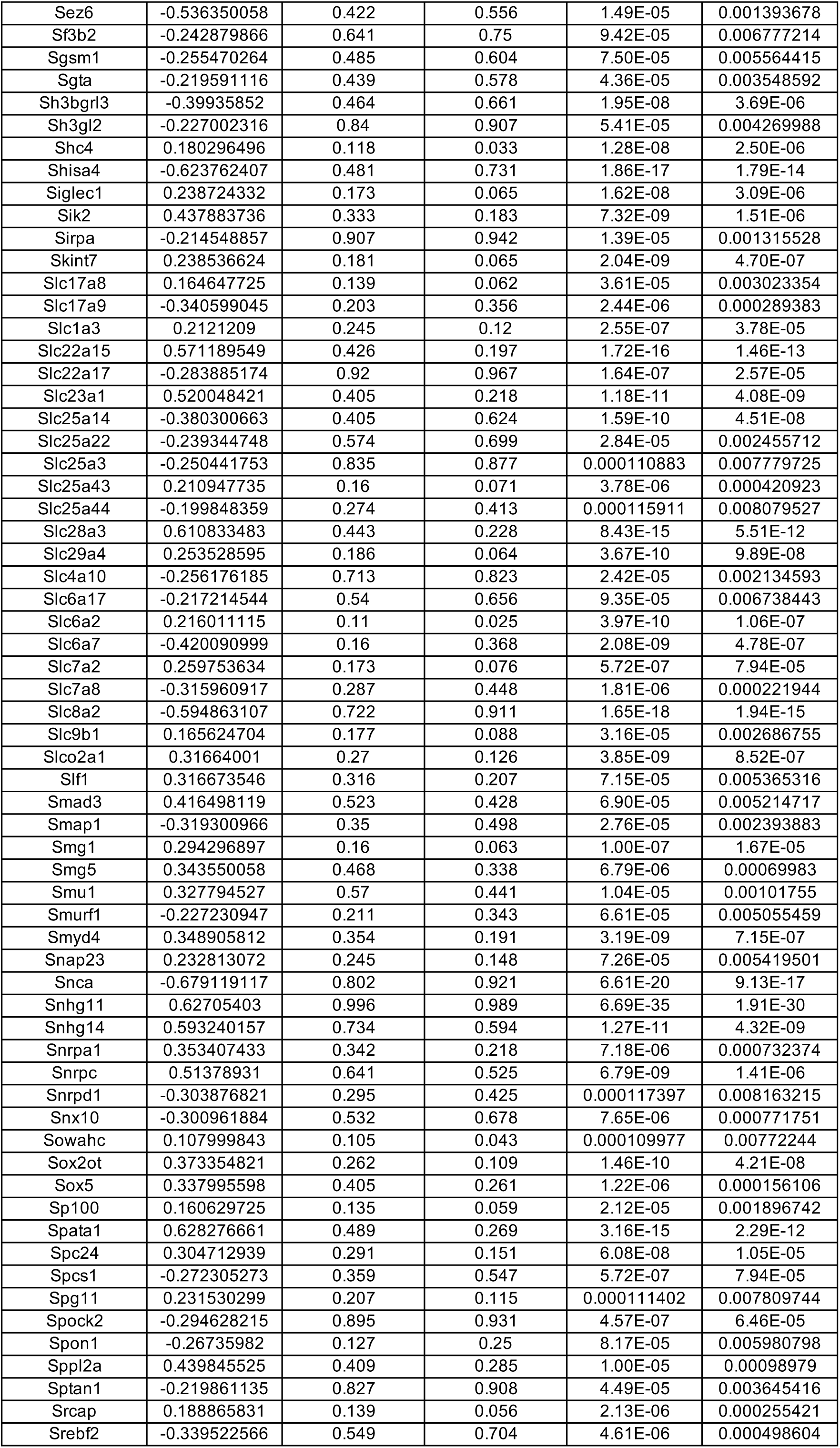
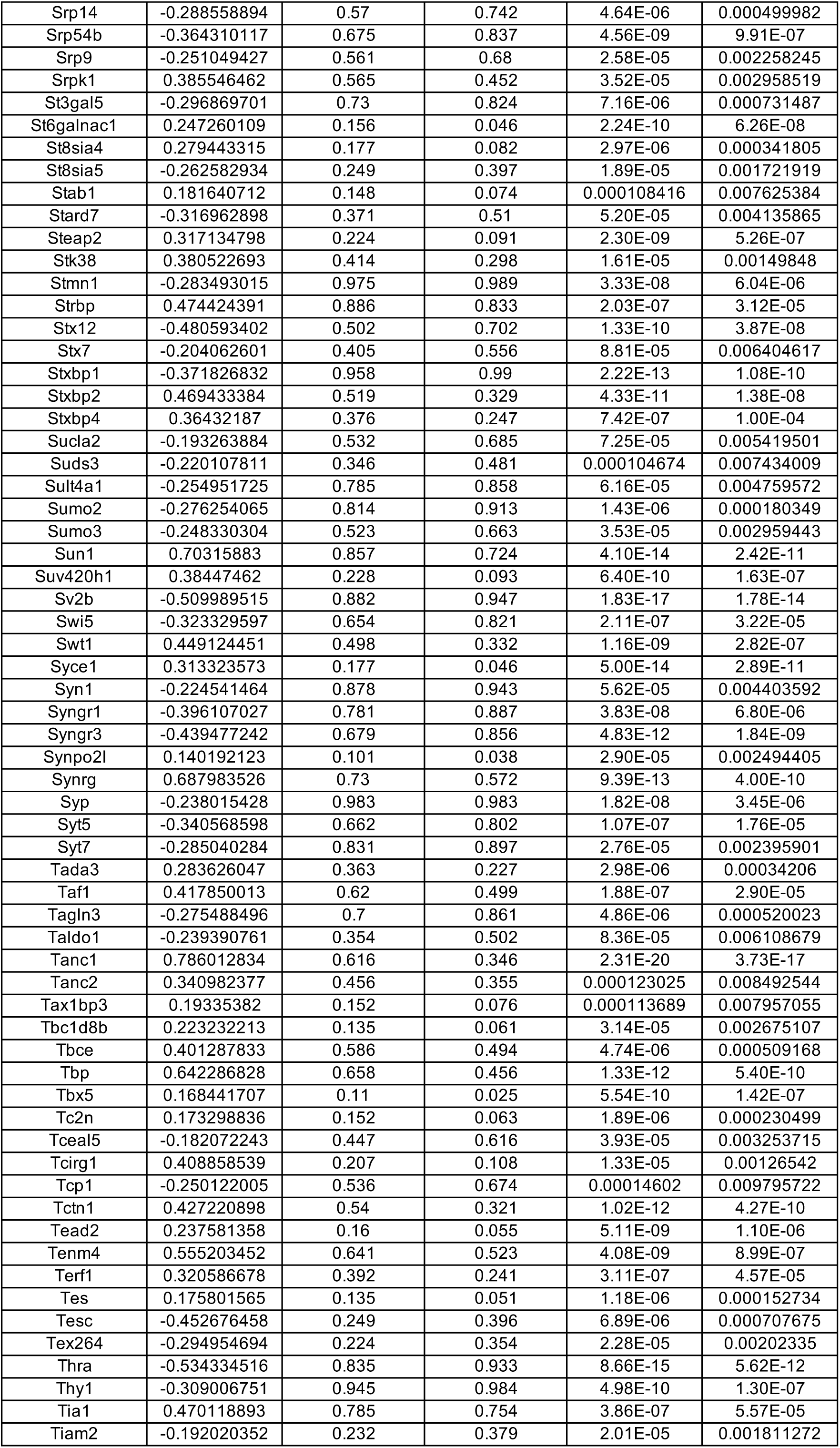
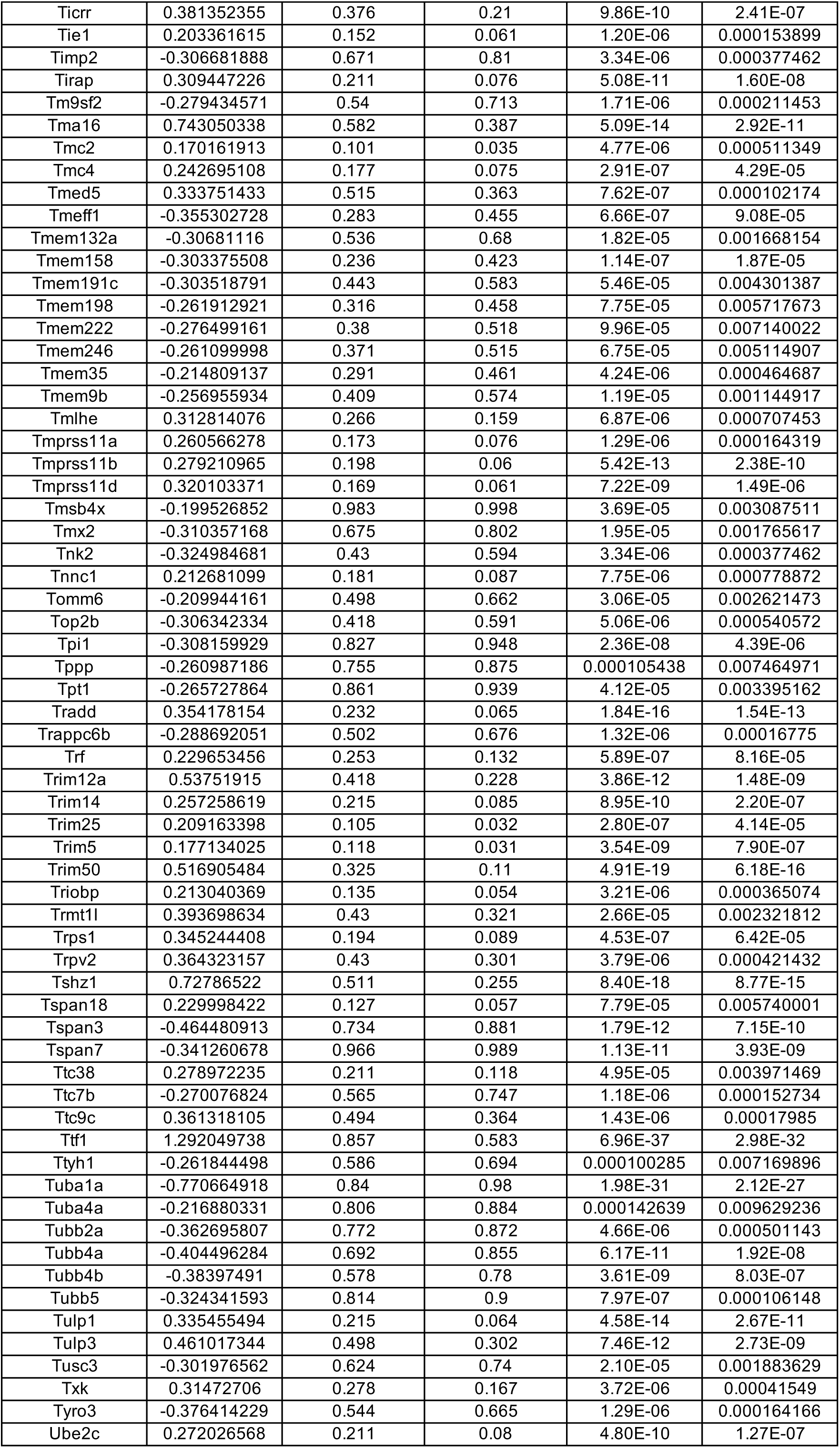
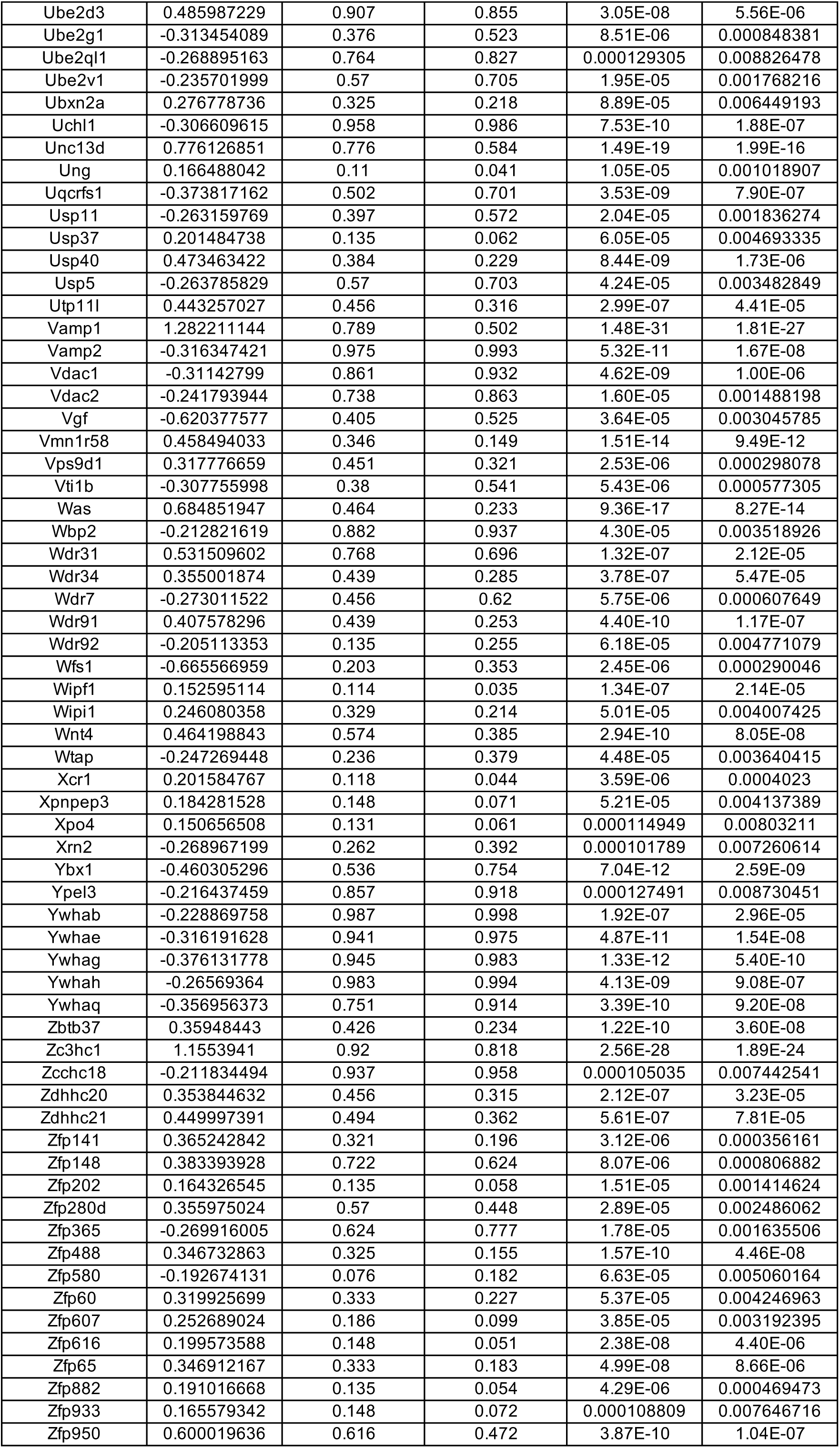
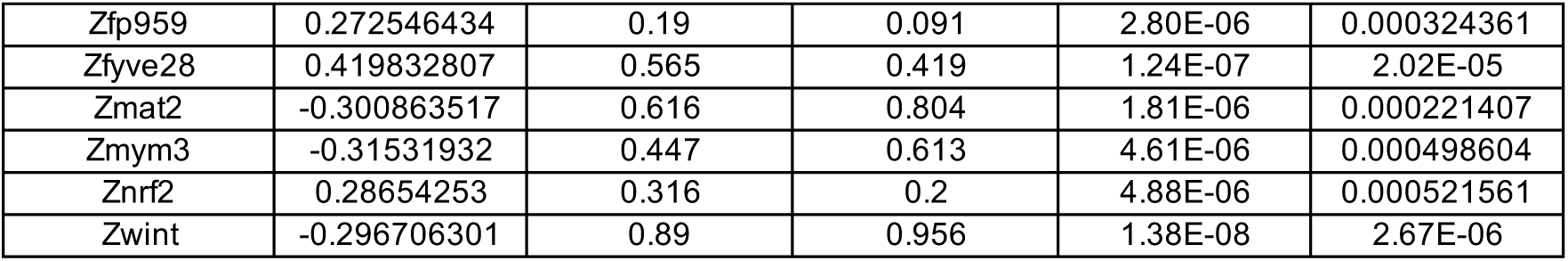
DEGs between wild-type (WT) and Mecp2-null (KO) excitatory neurons of mouse OC (adjusted p-value < 0.01)

**Table S43.**
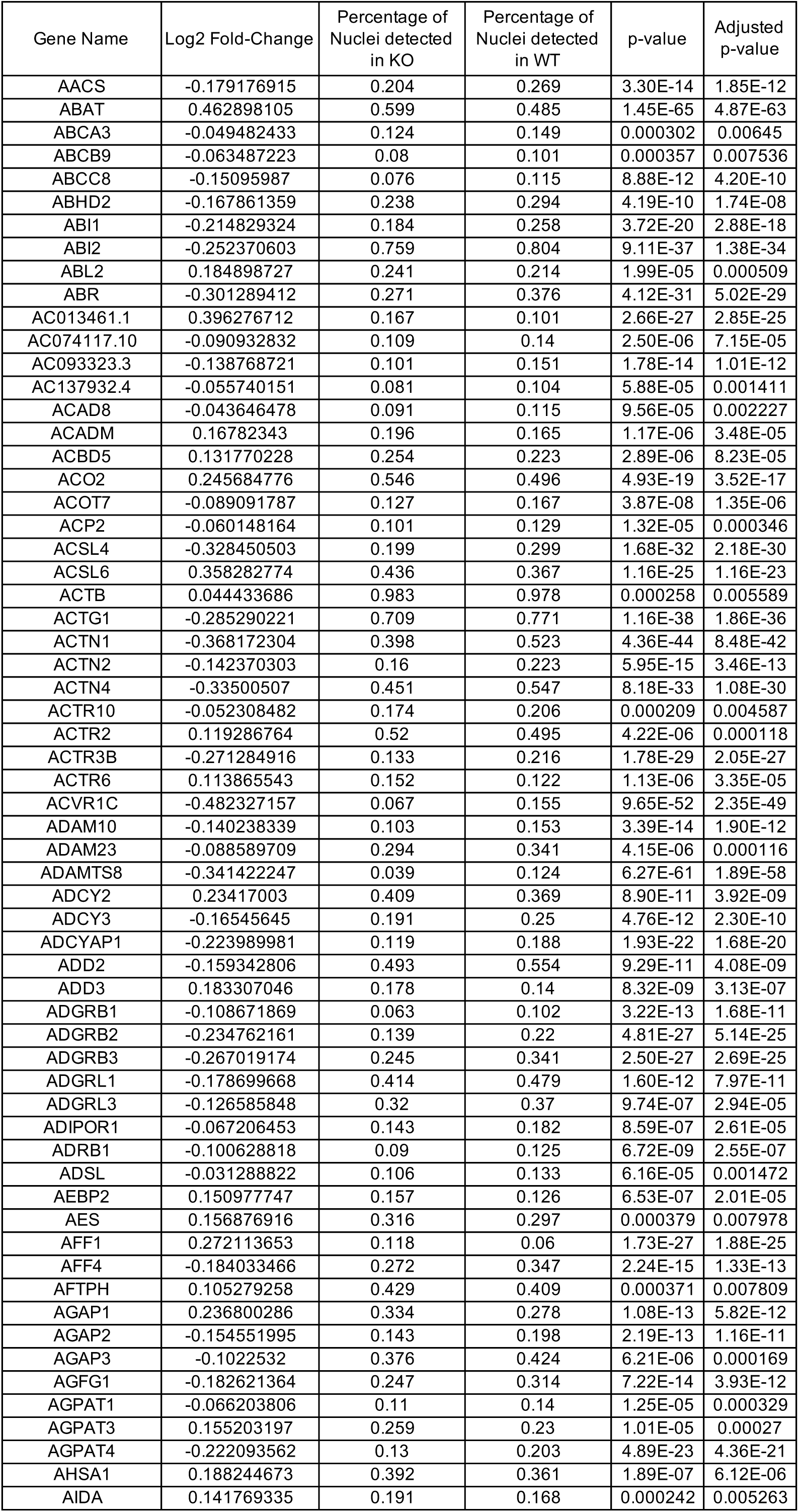
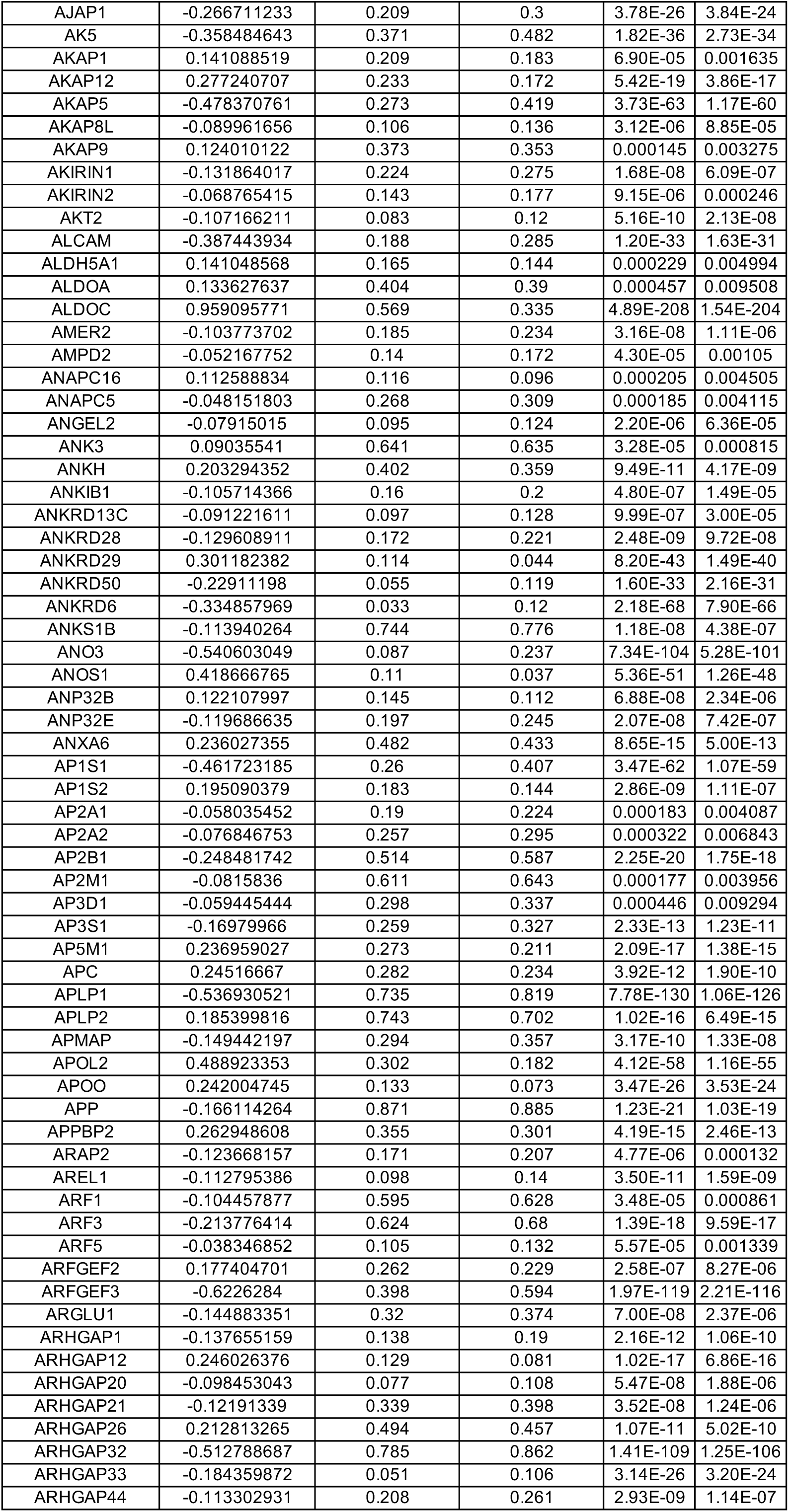
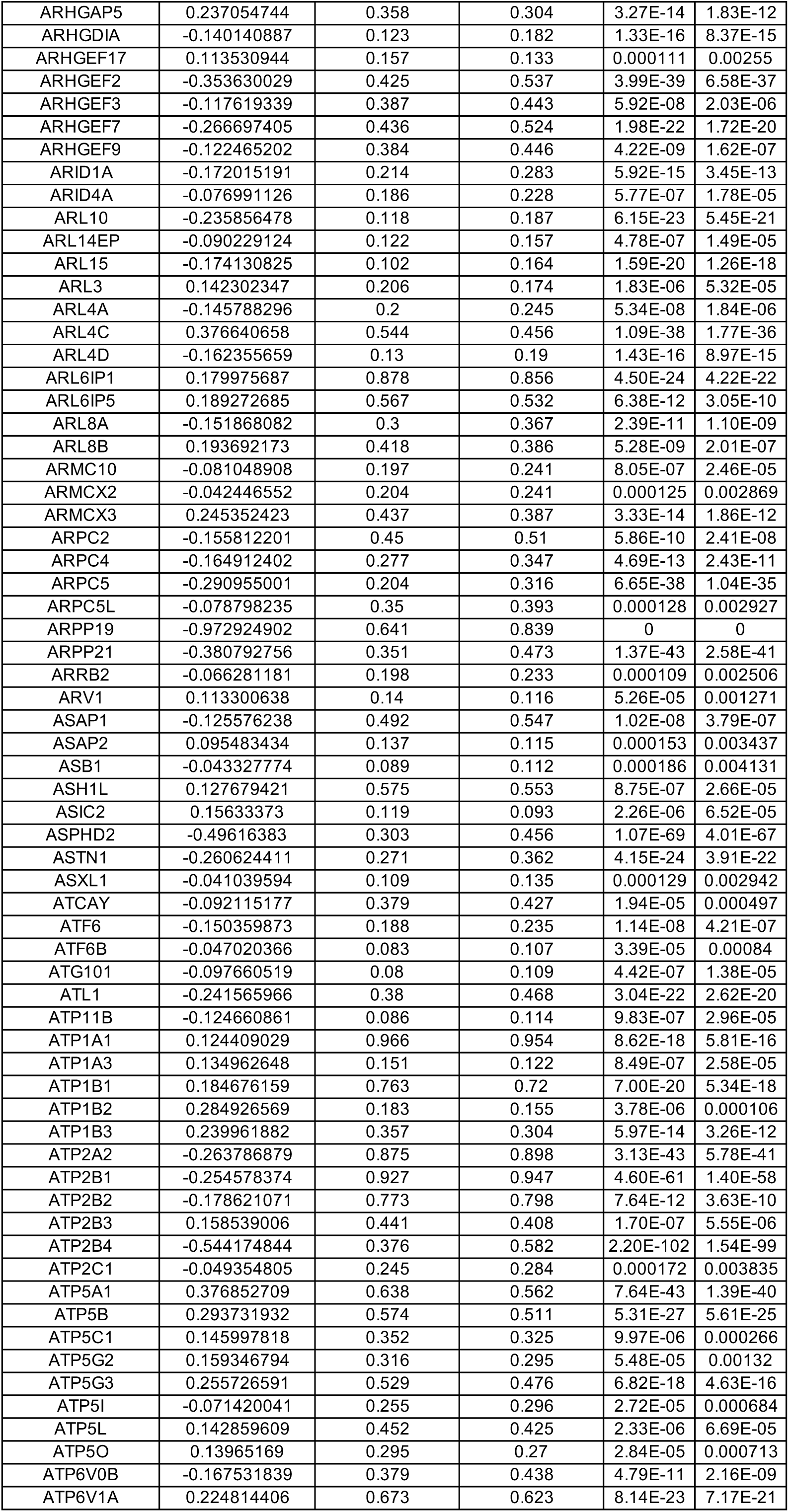
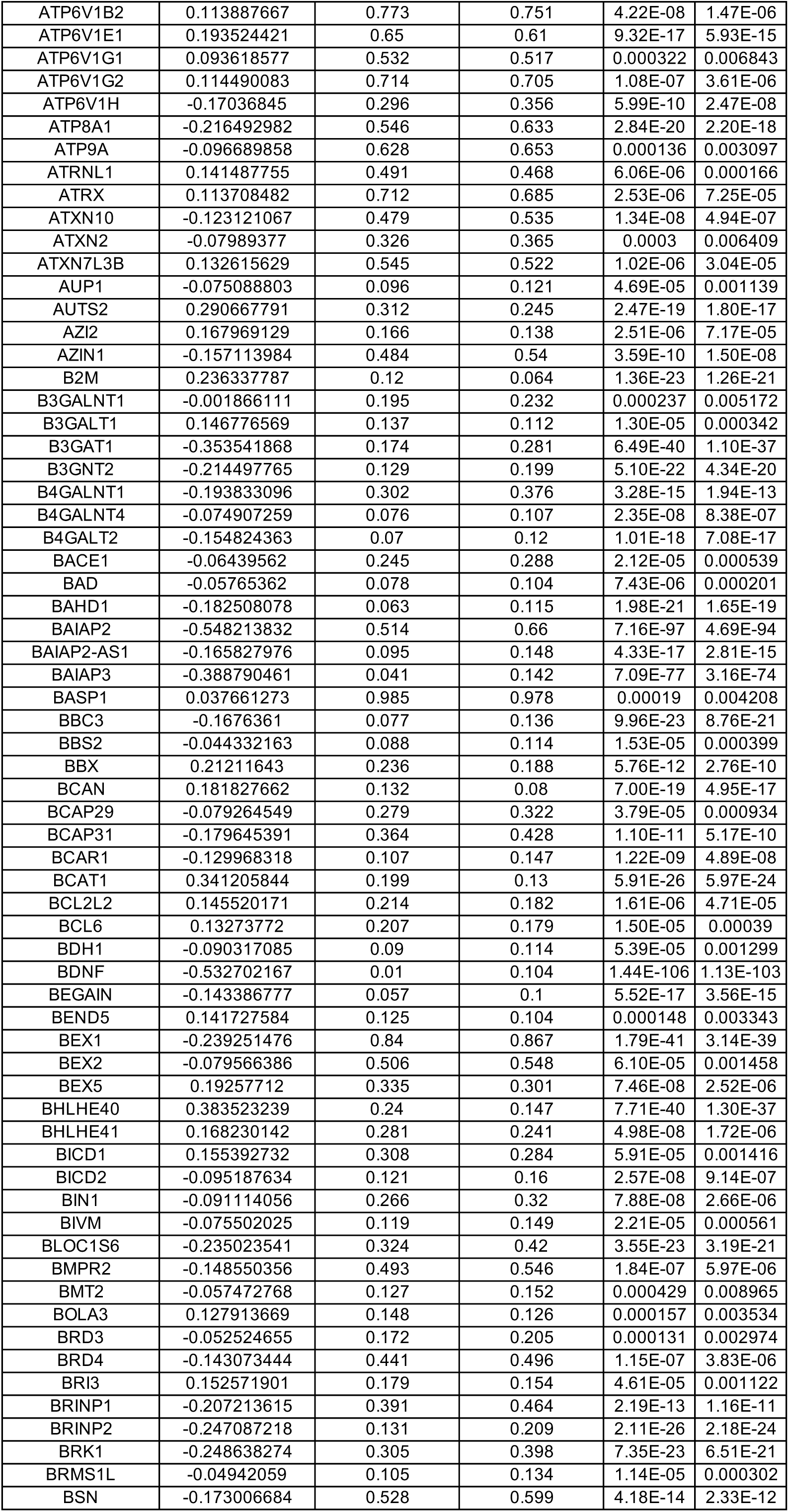
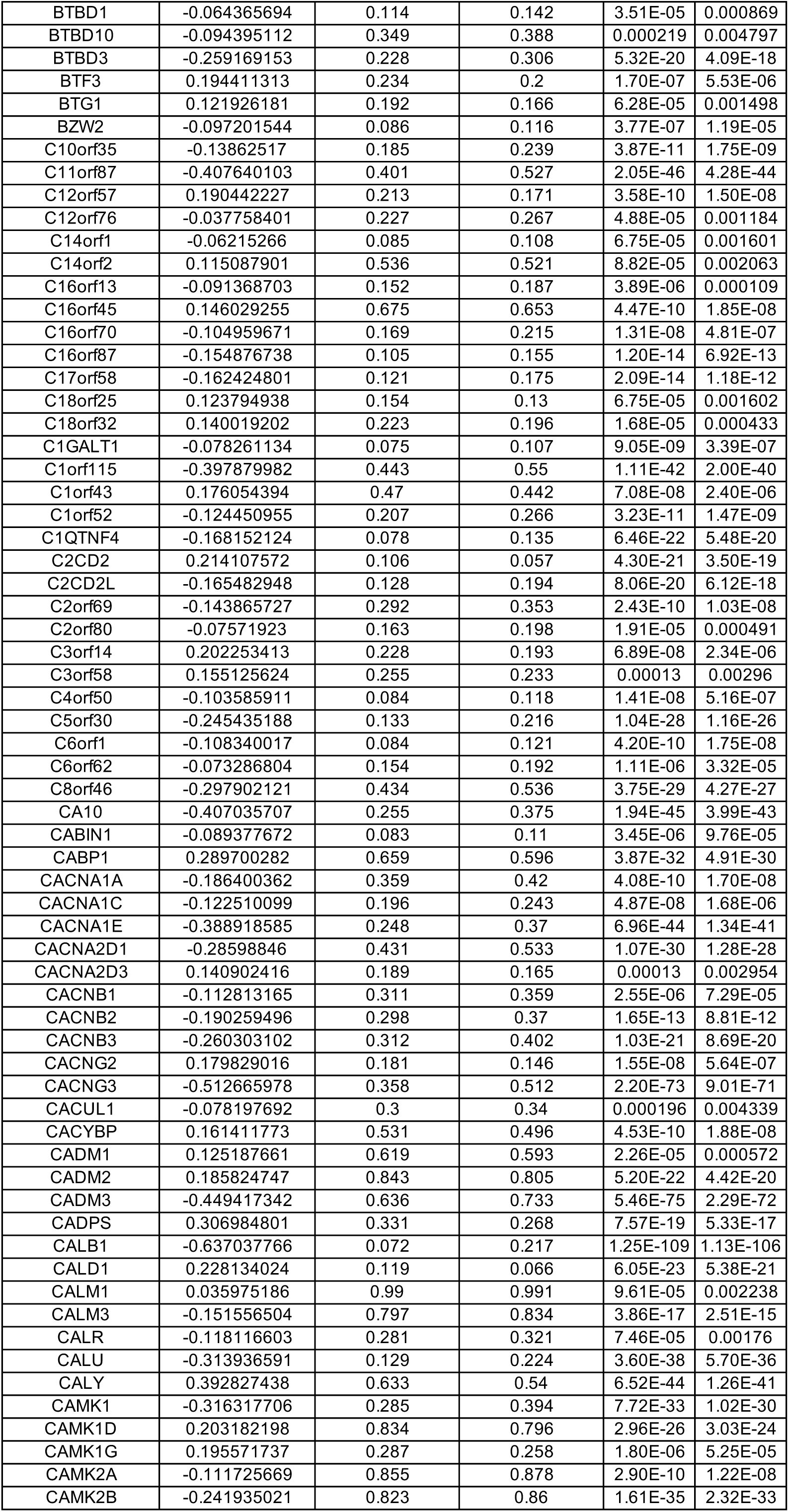
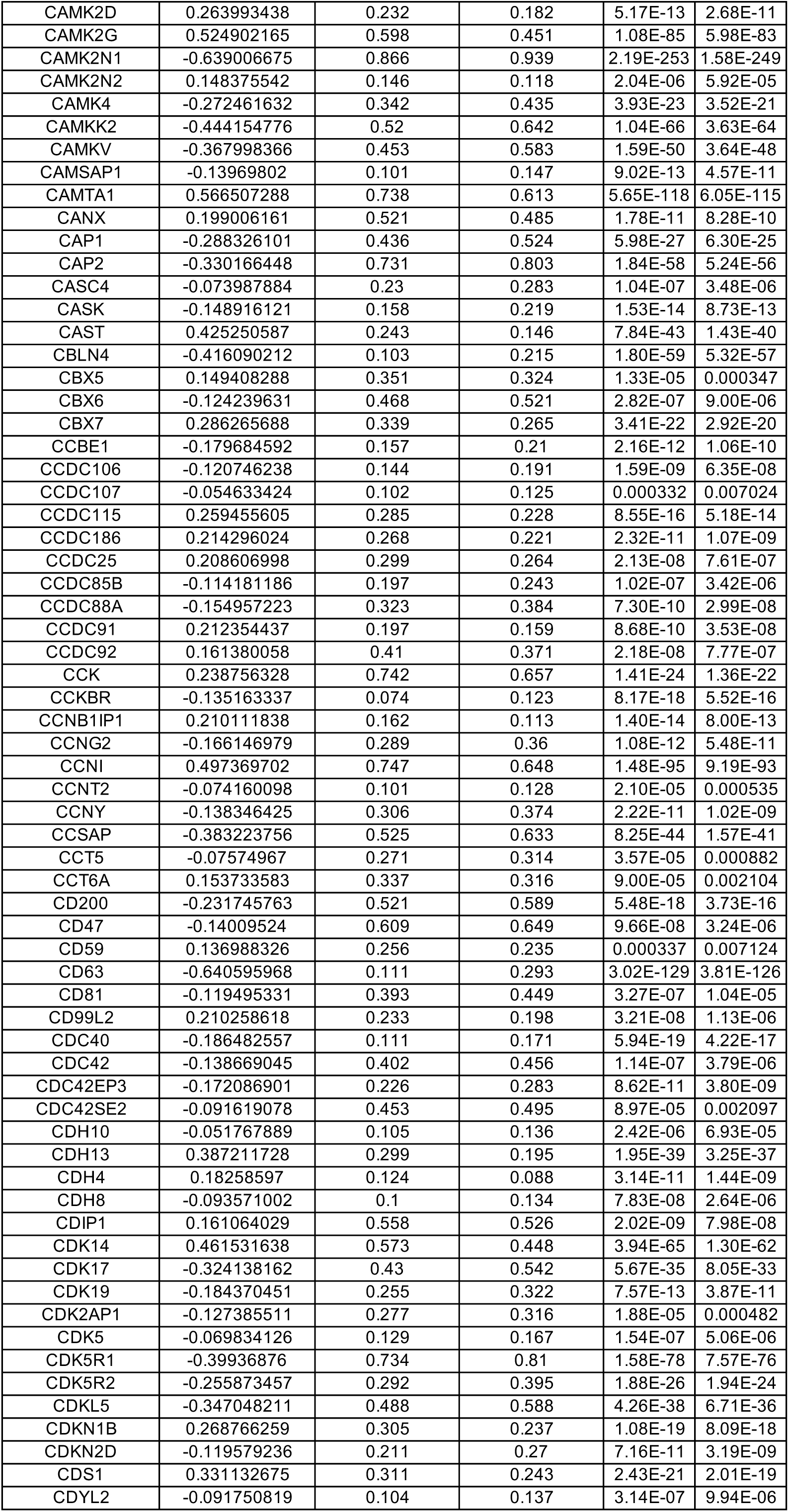
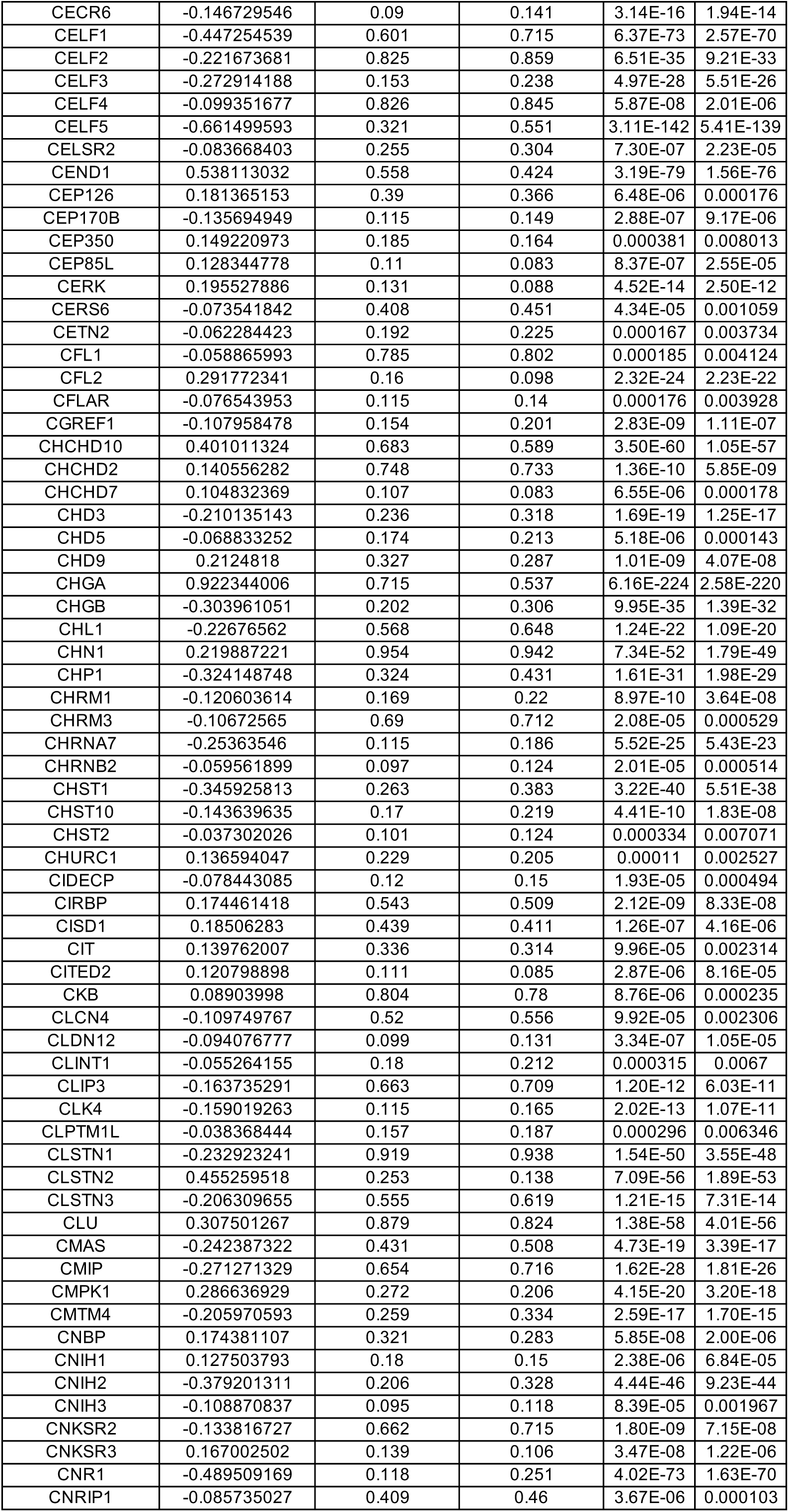
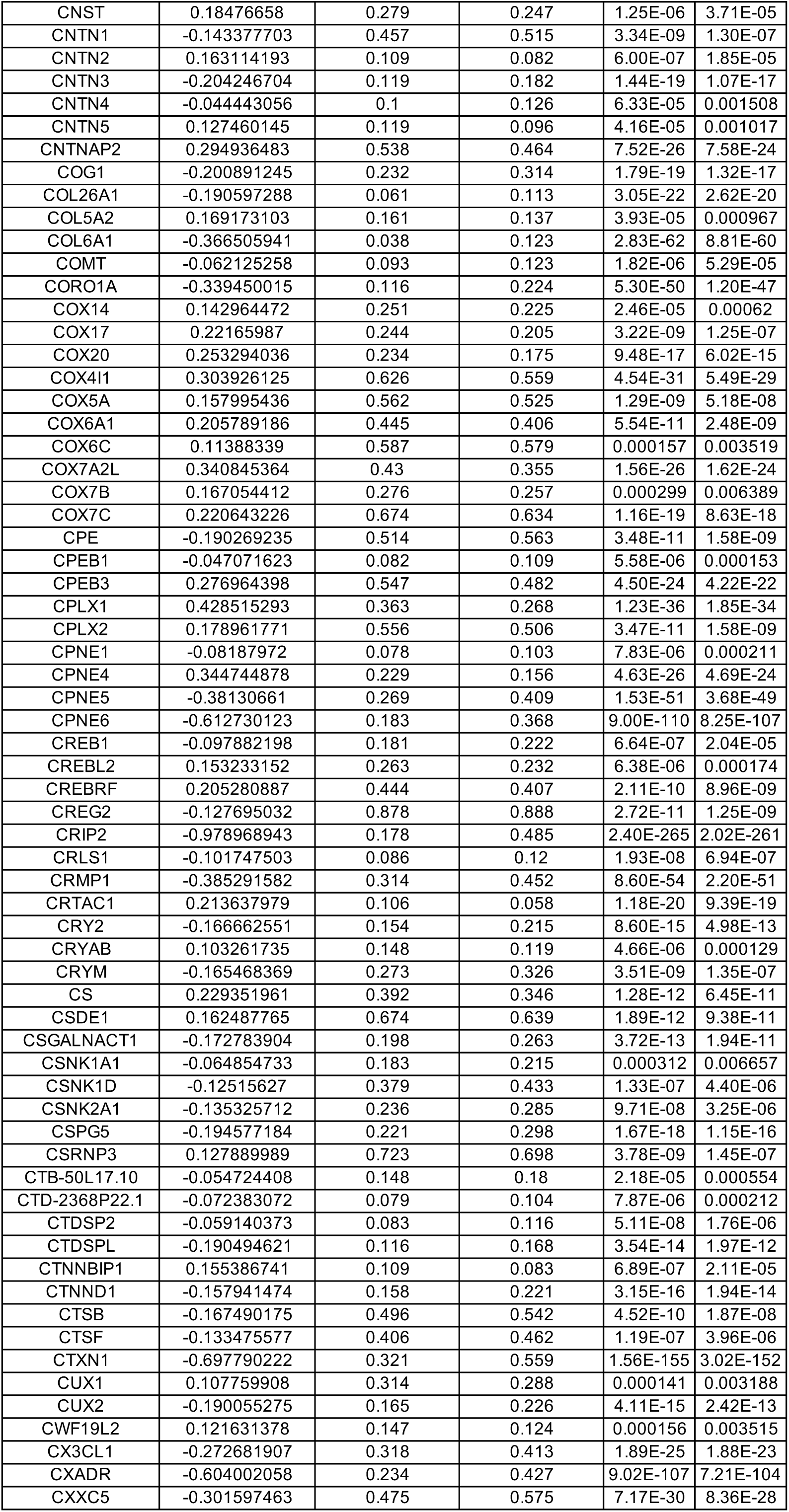
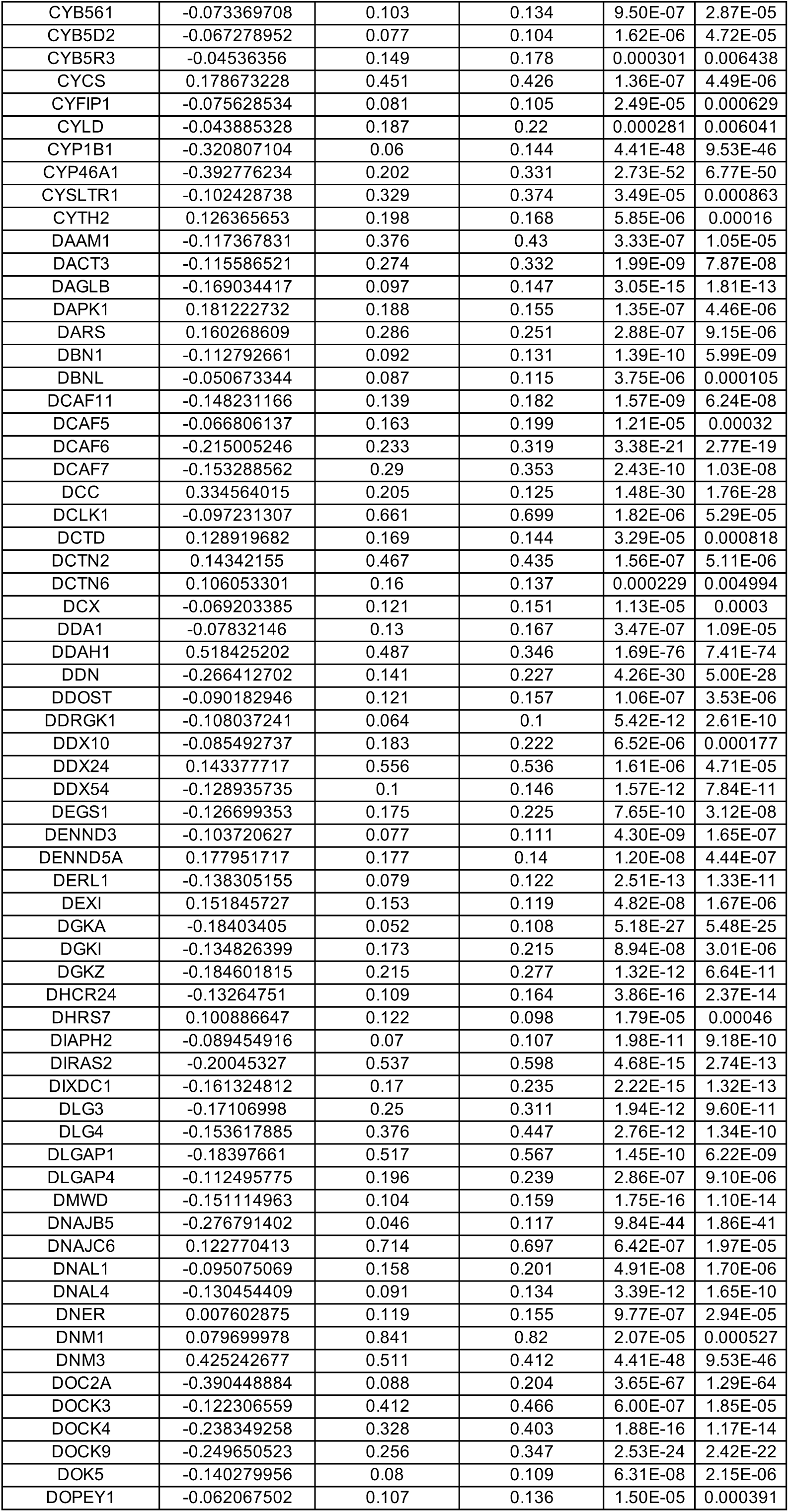
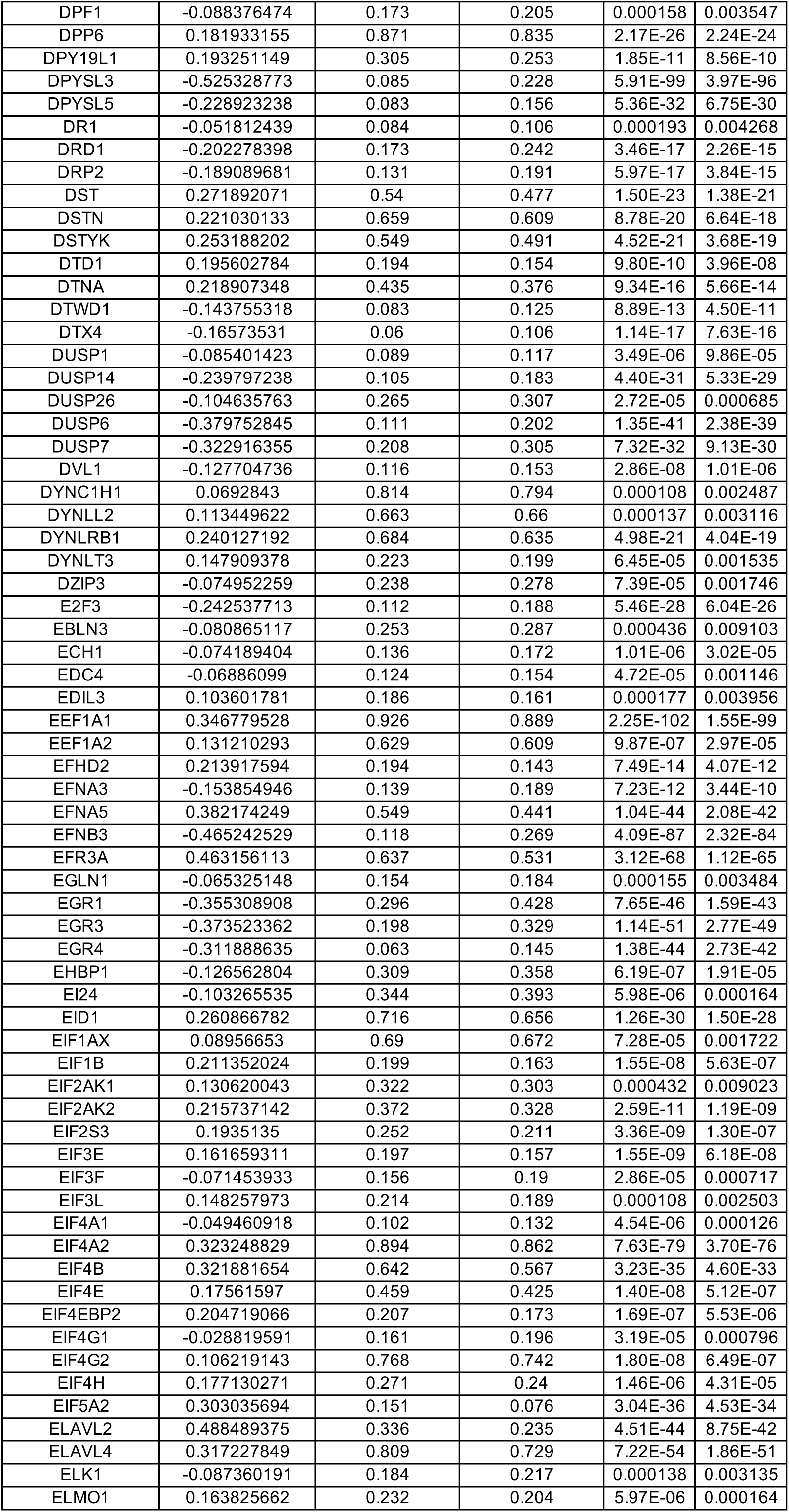
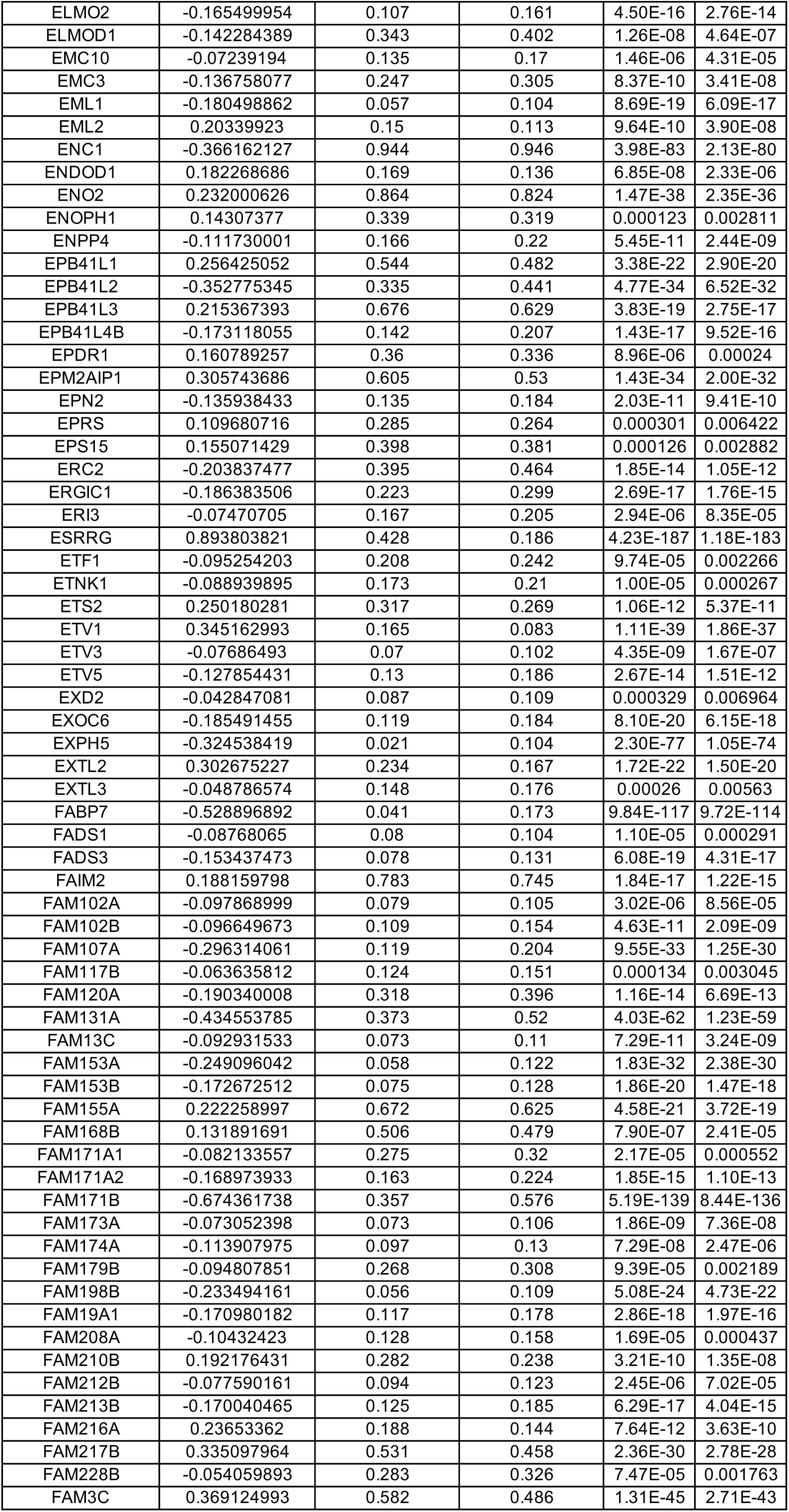
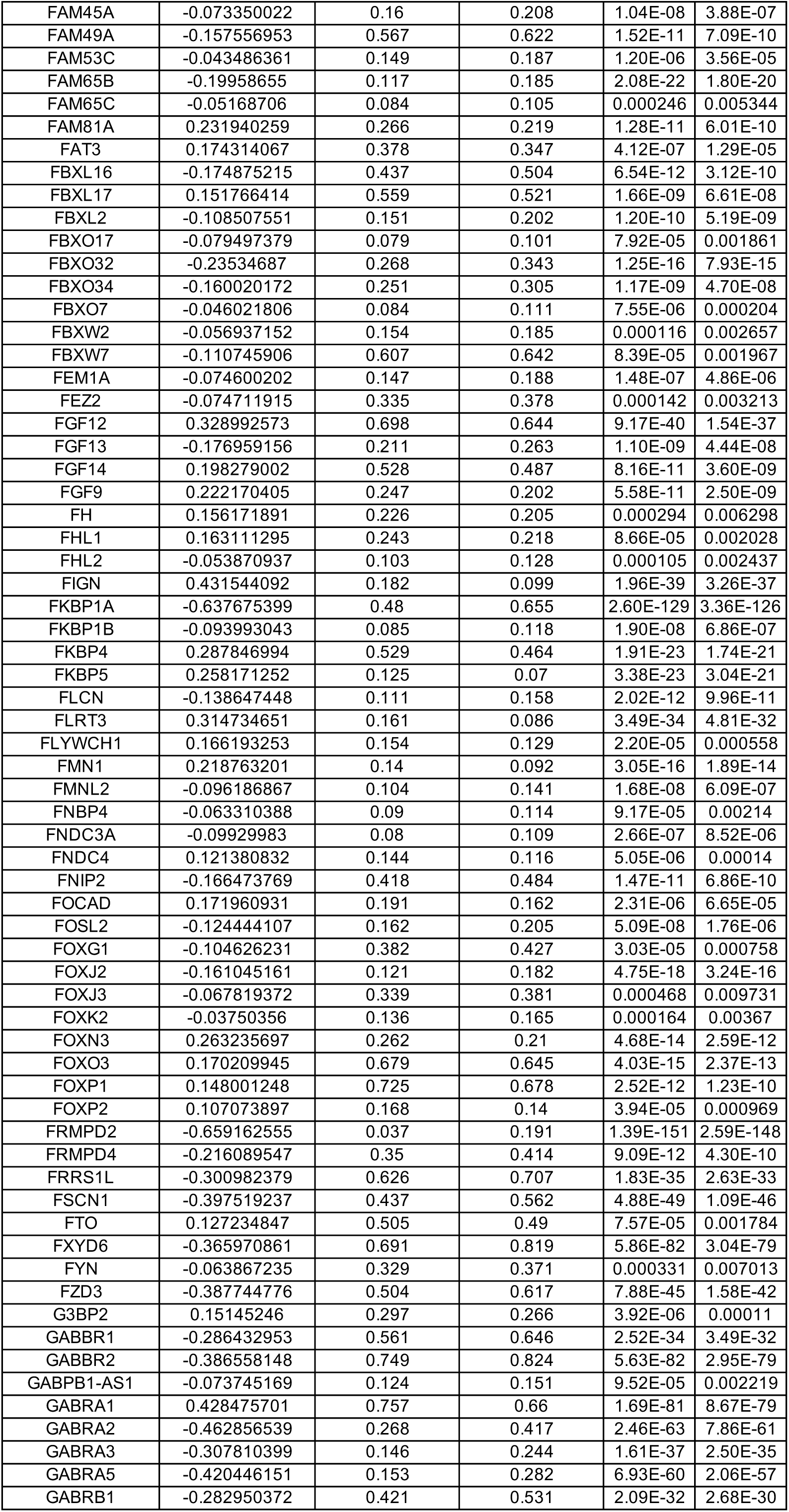
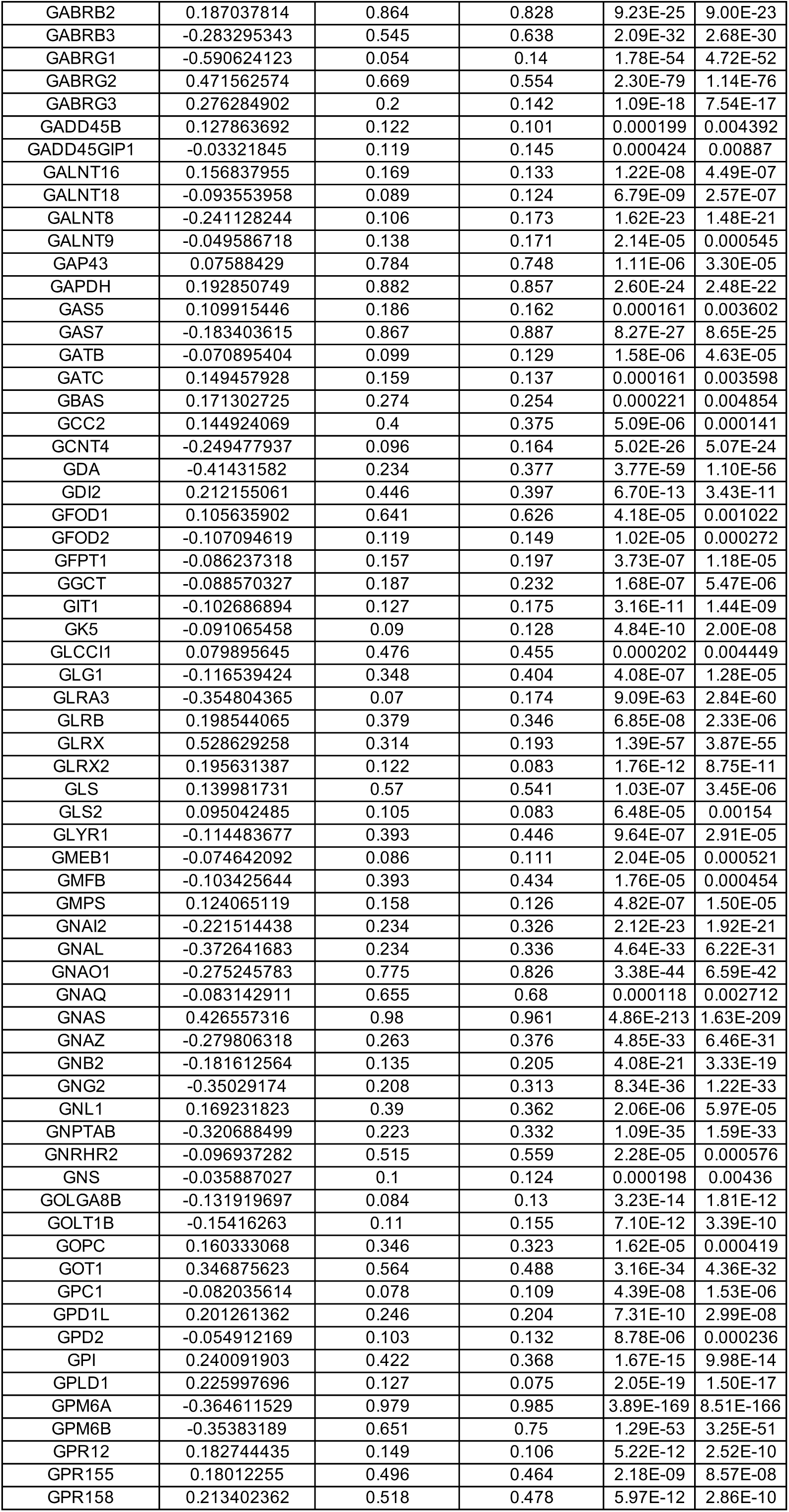
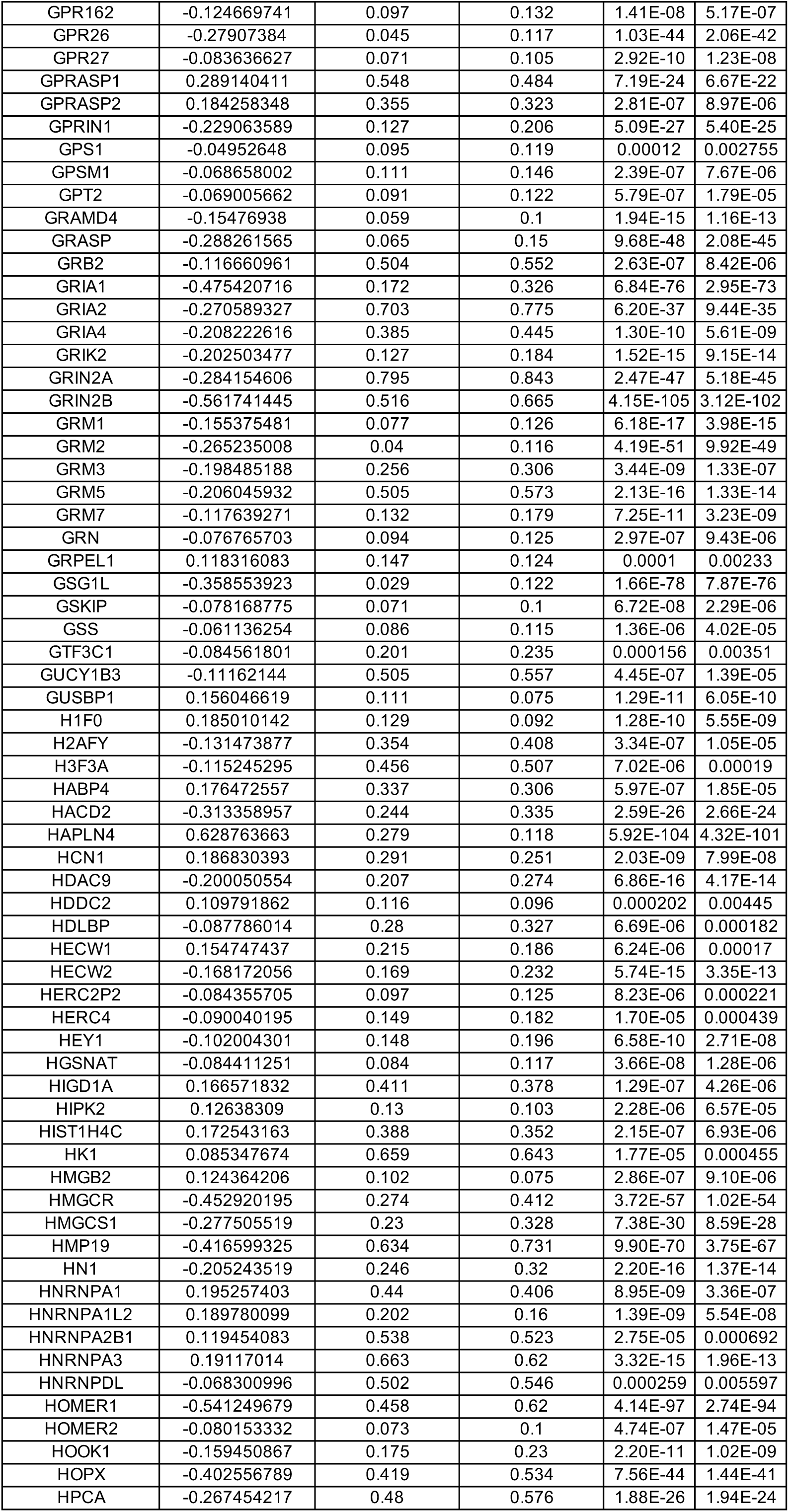
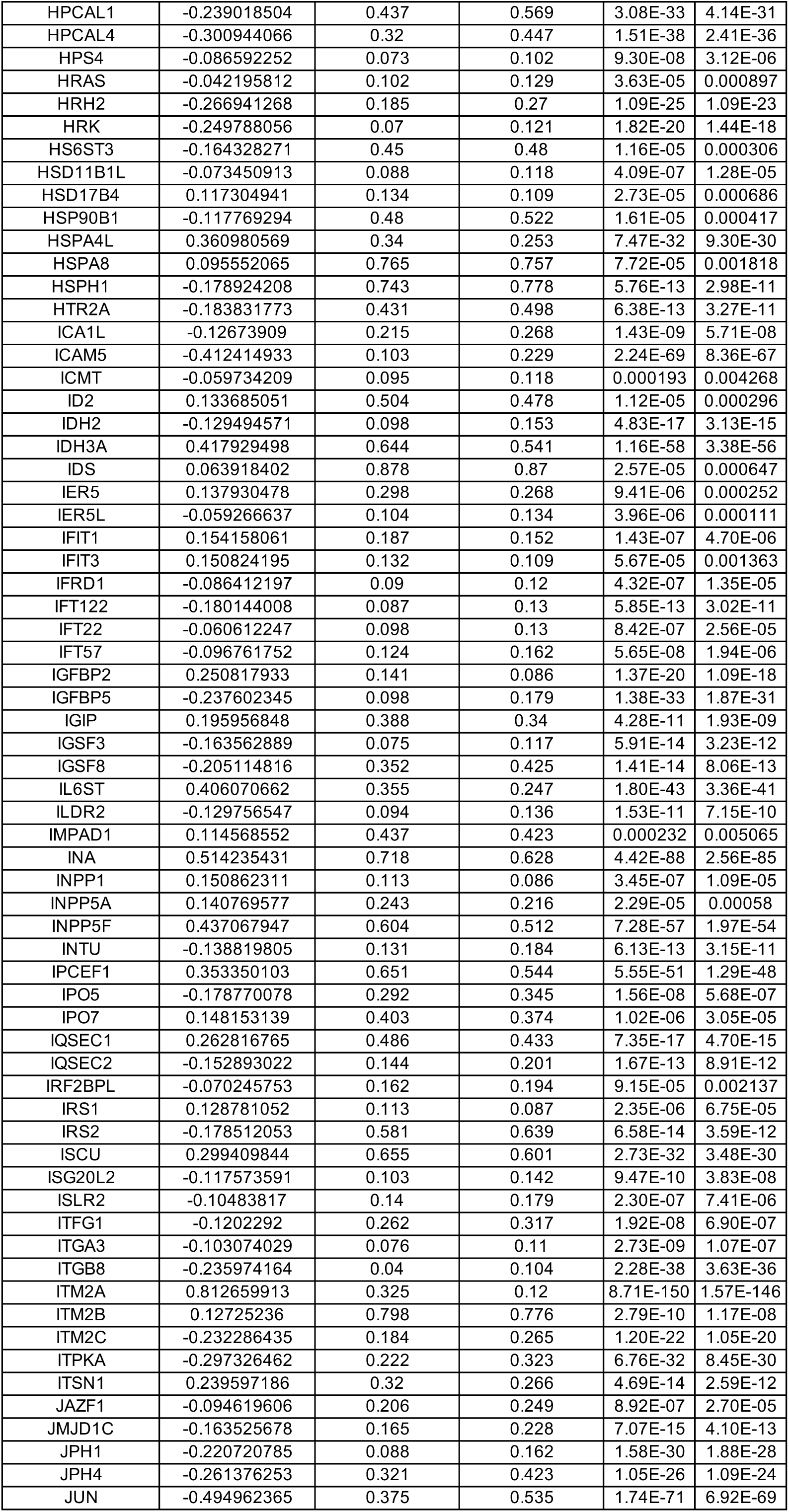
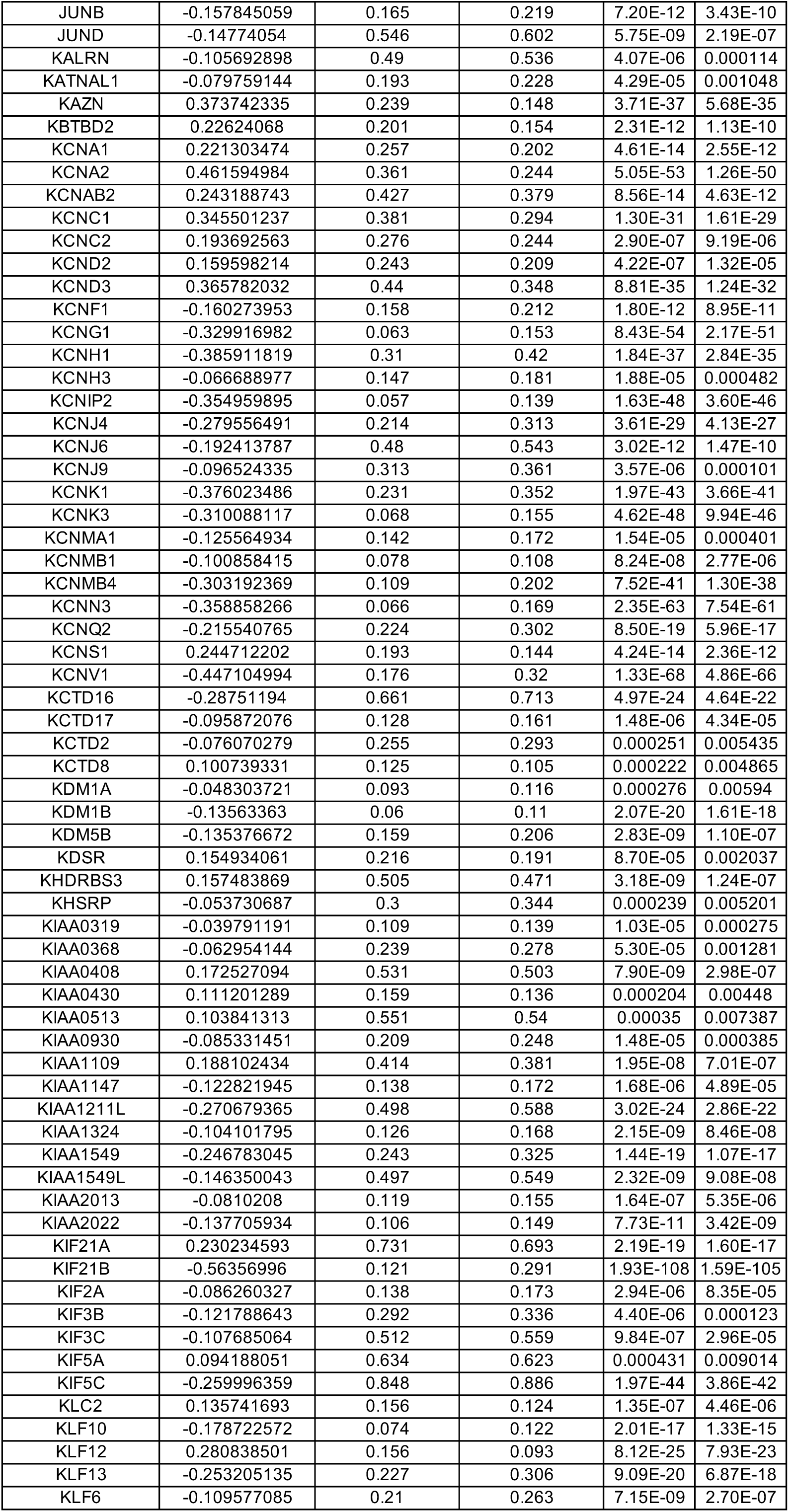
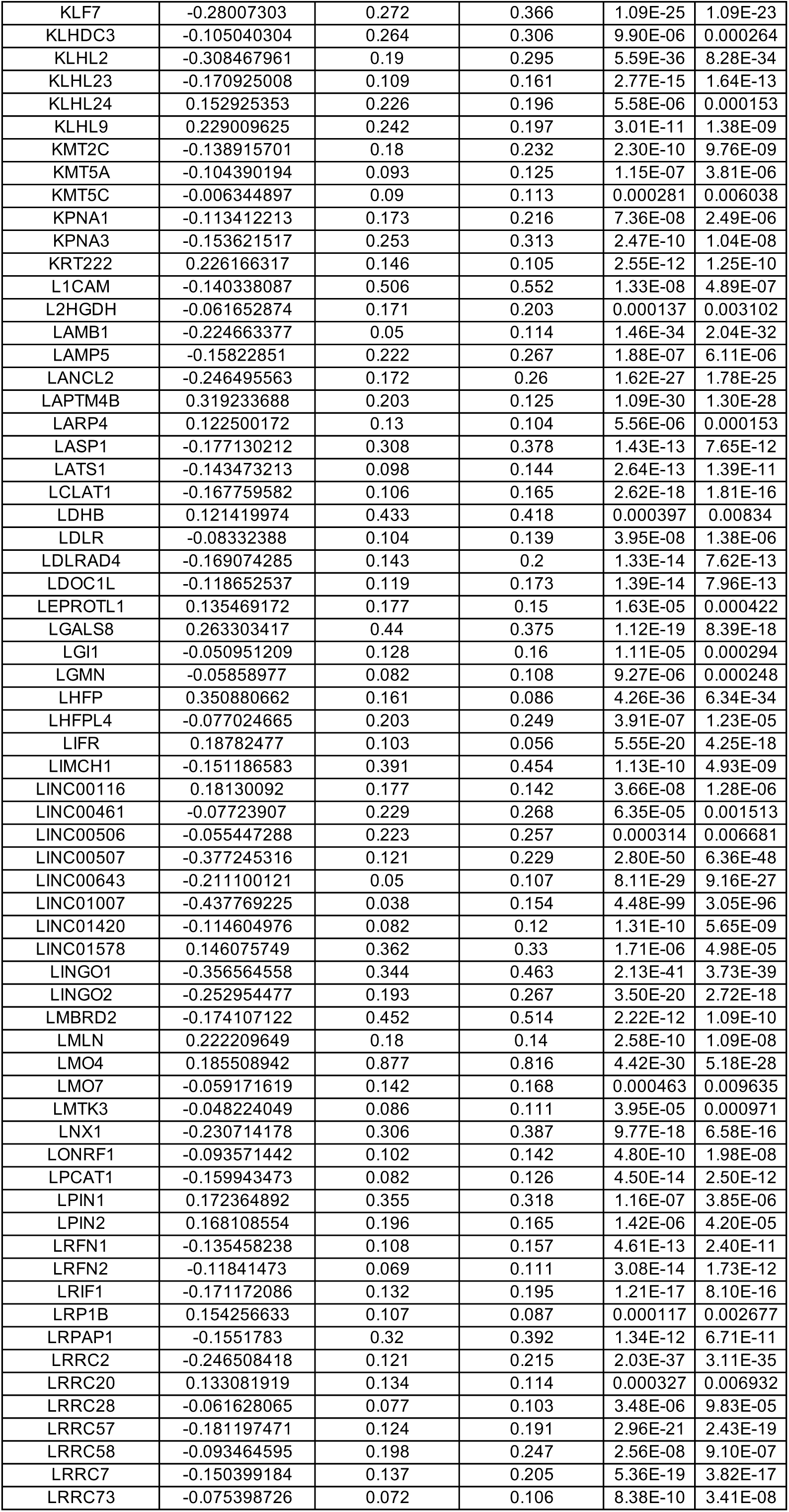
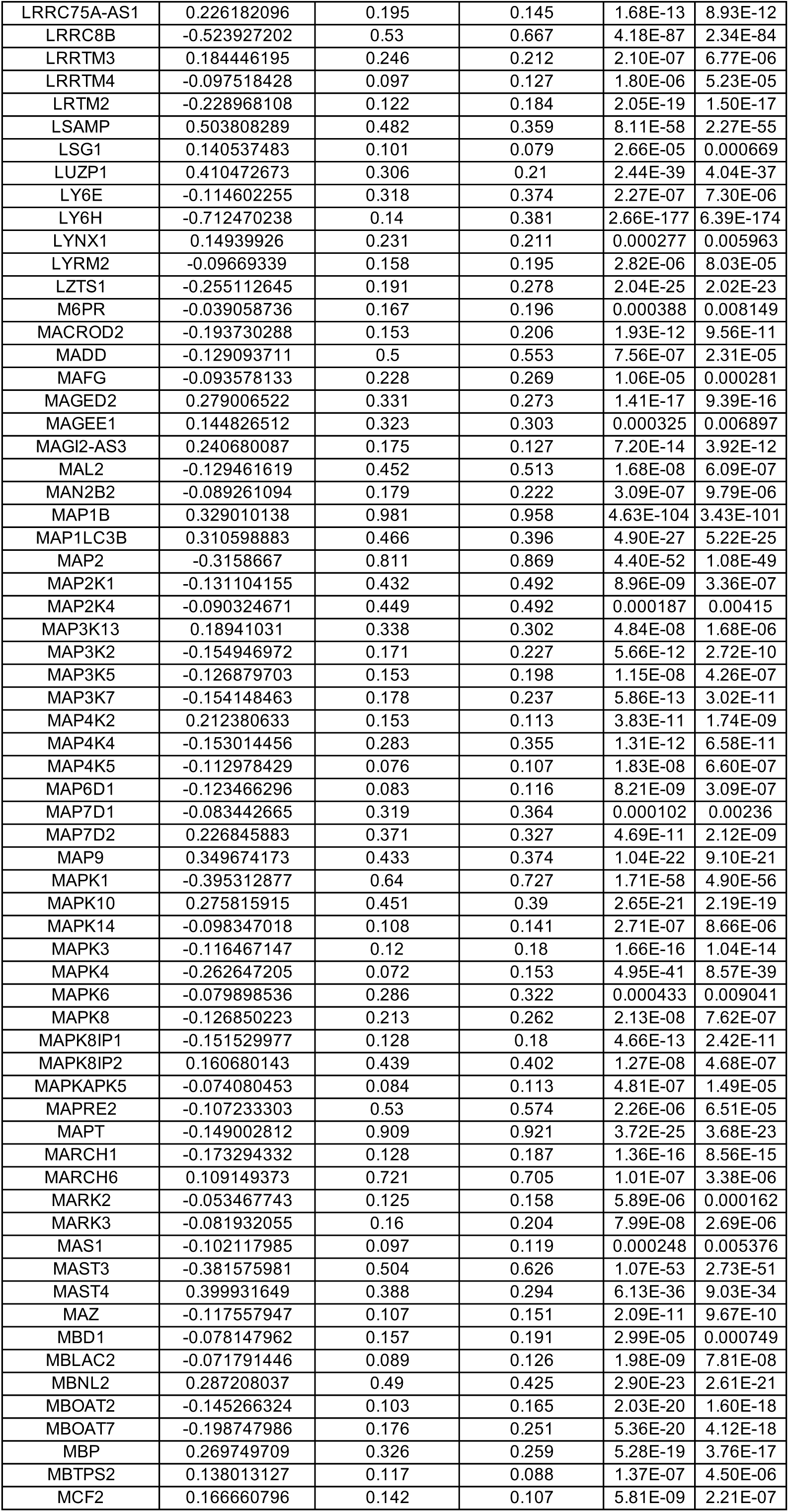
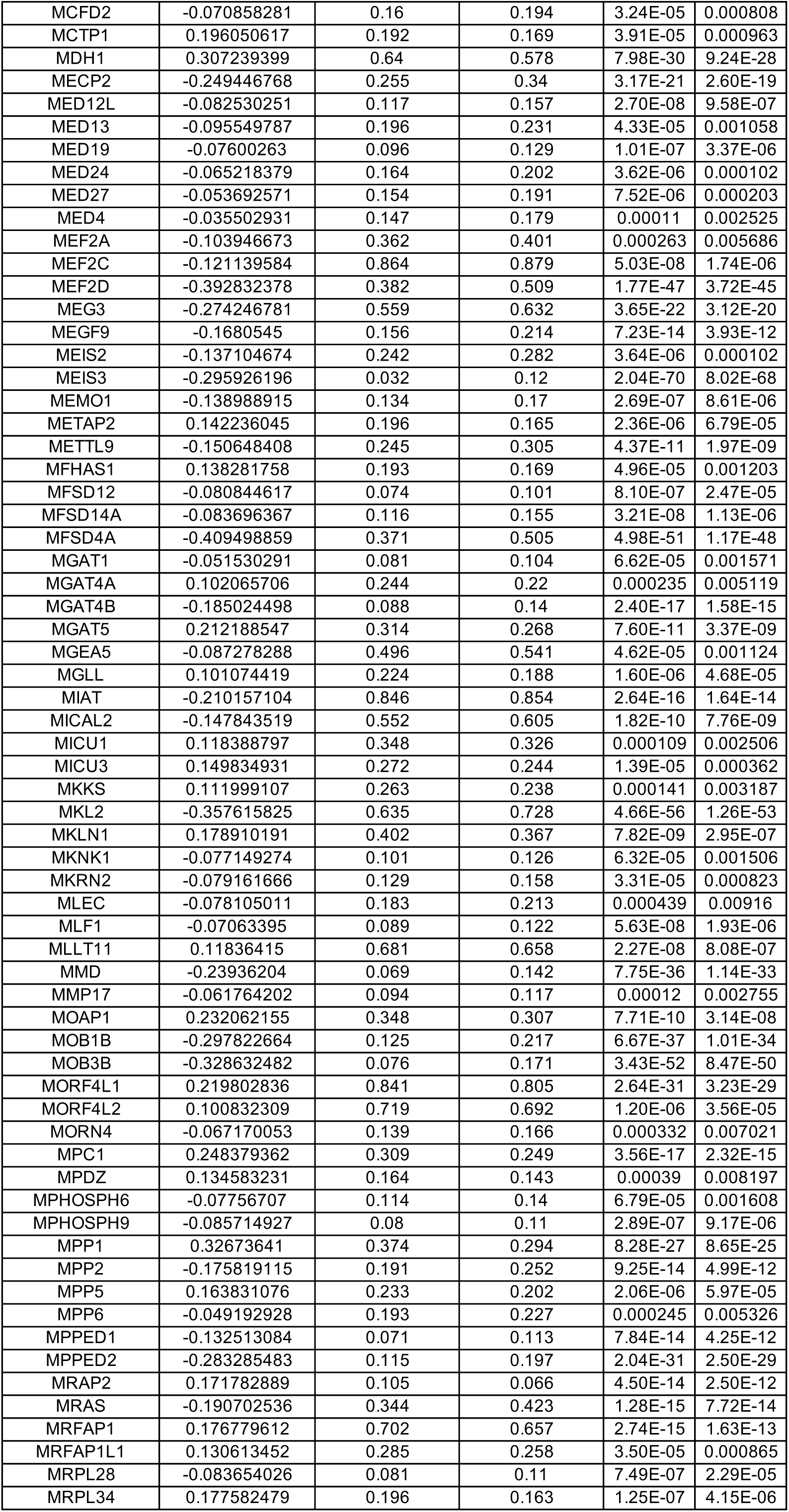
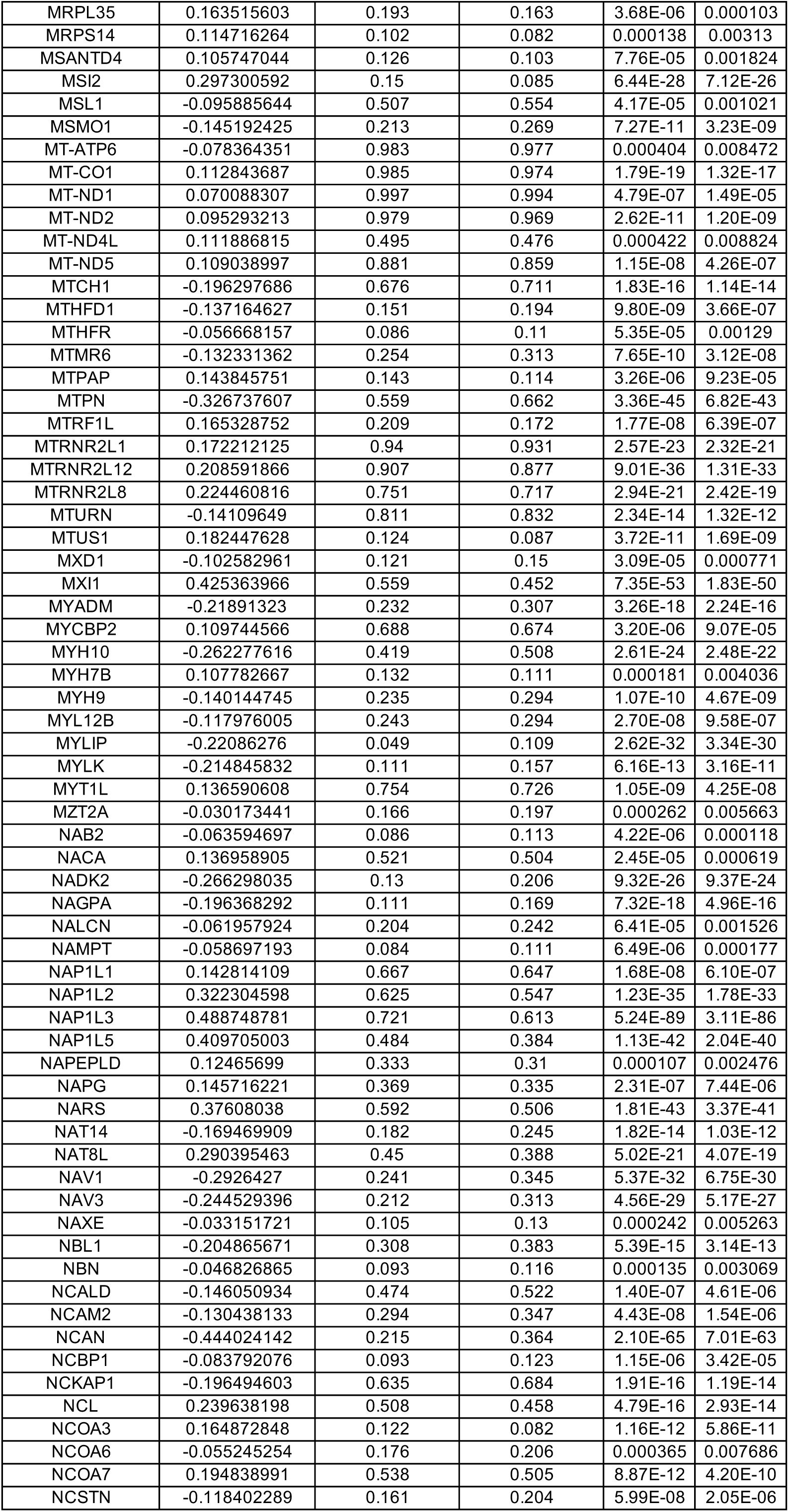
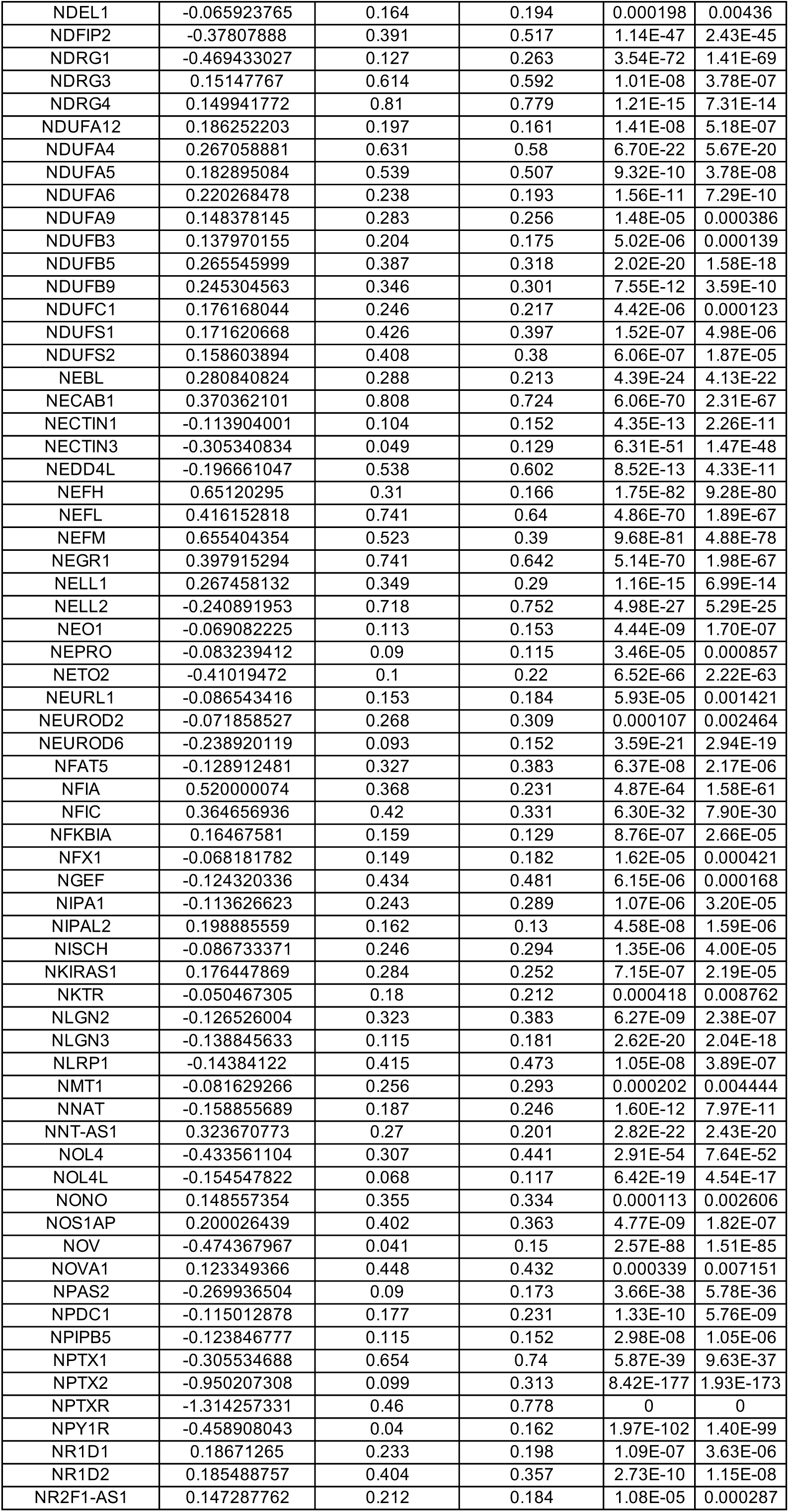
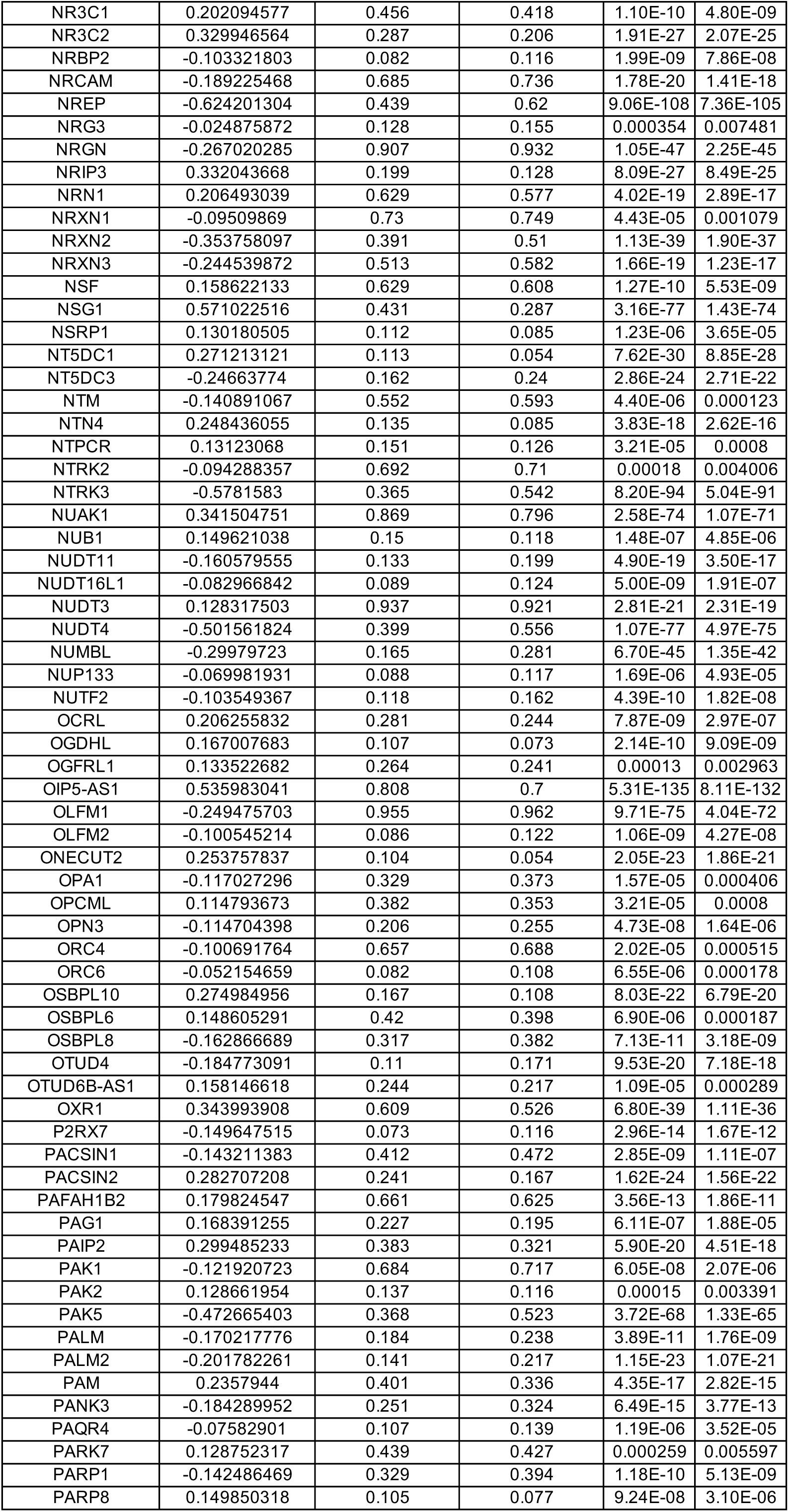
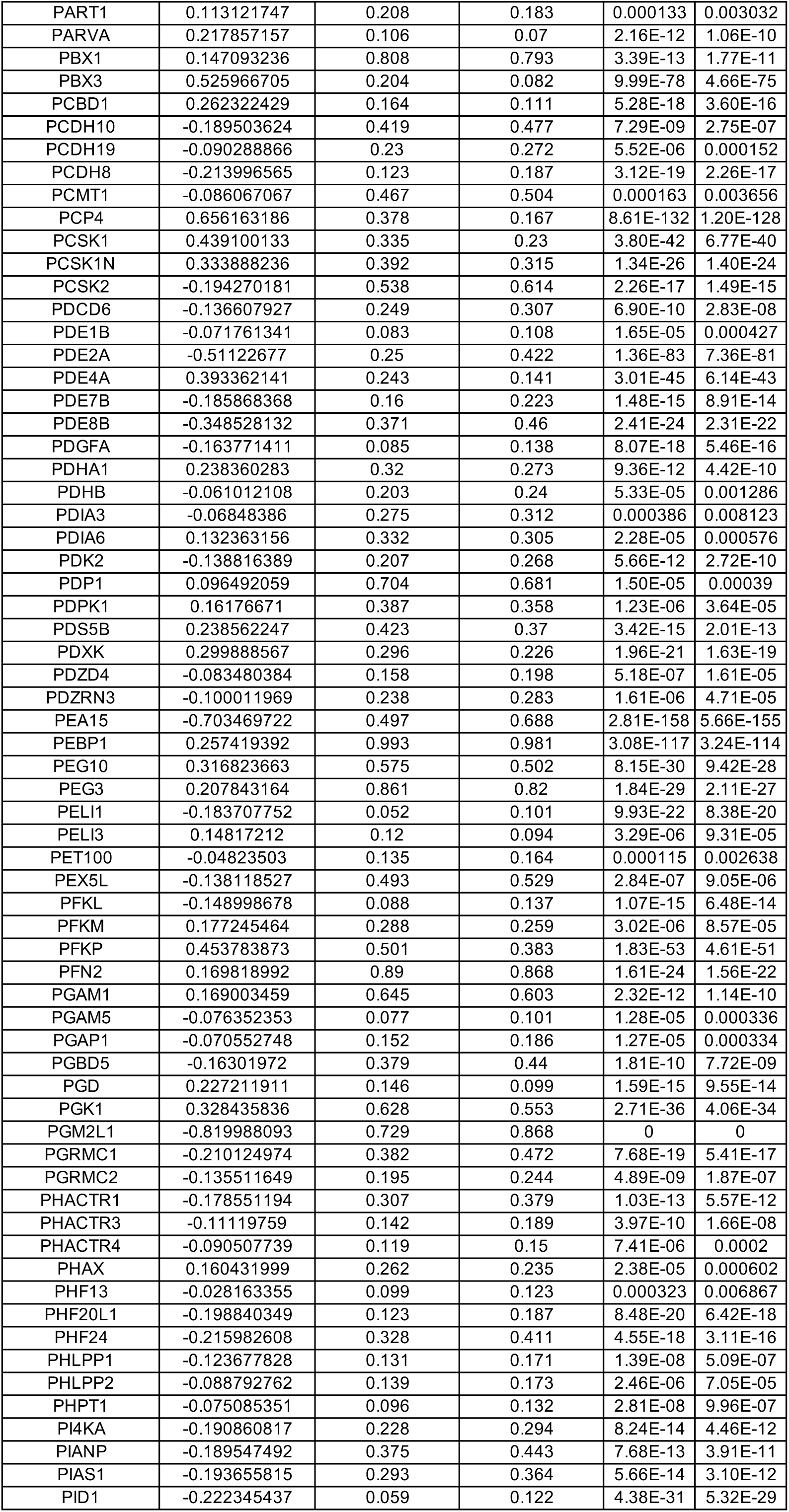
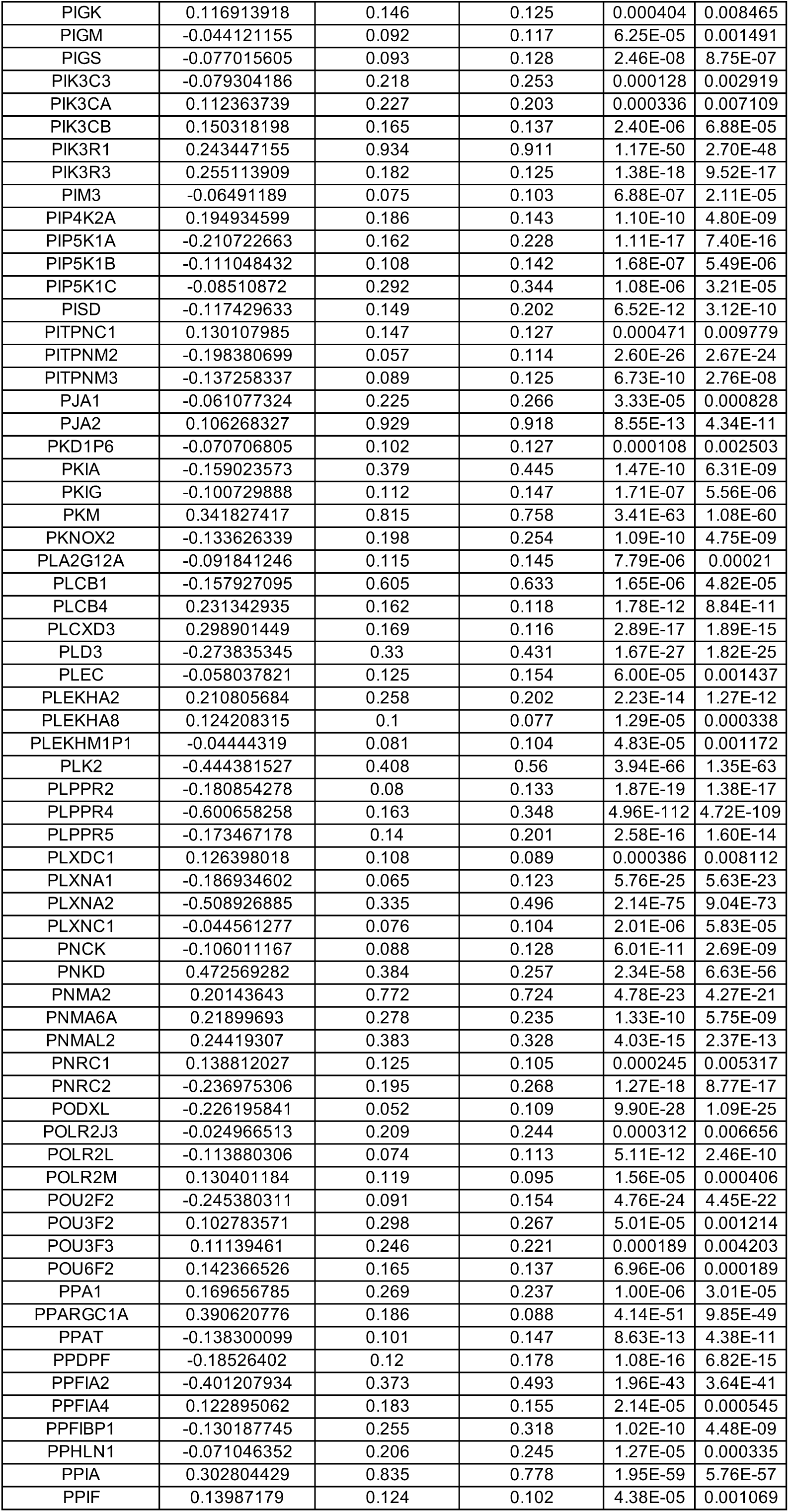
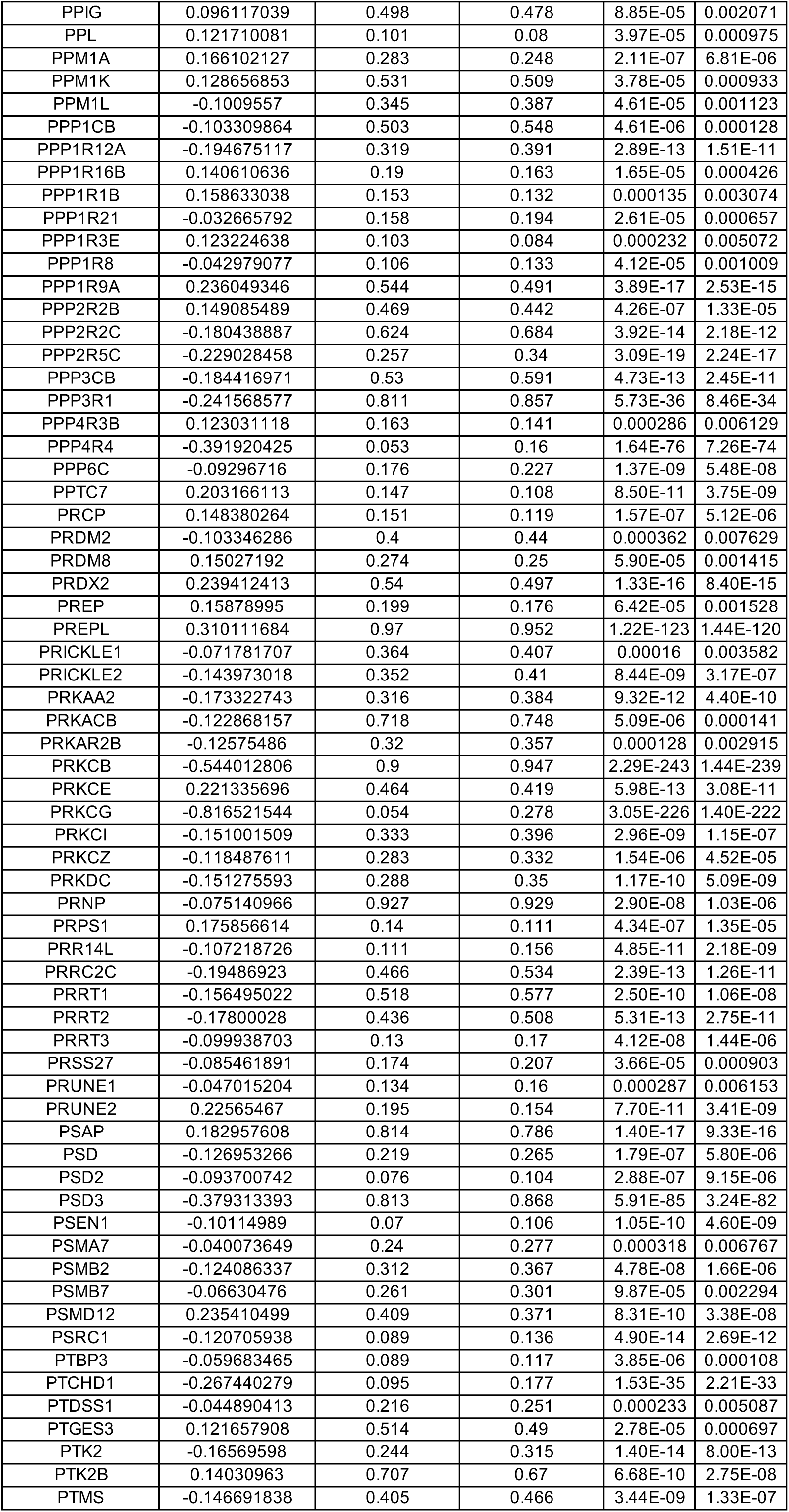
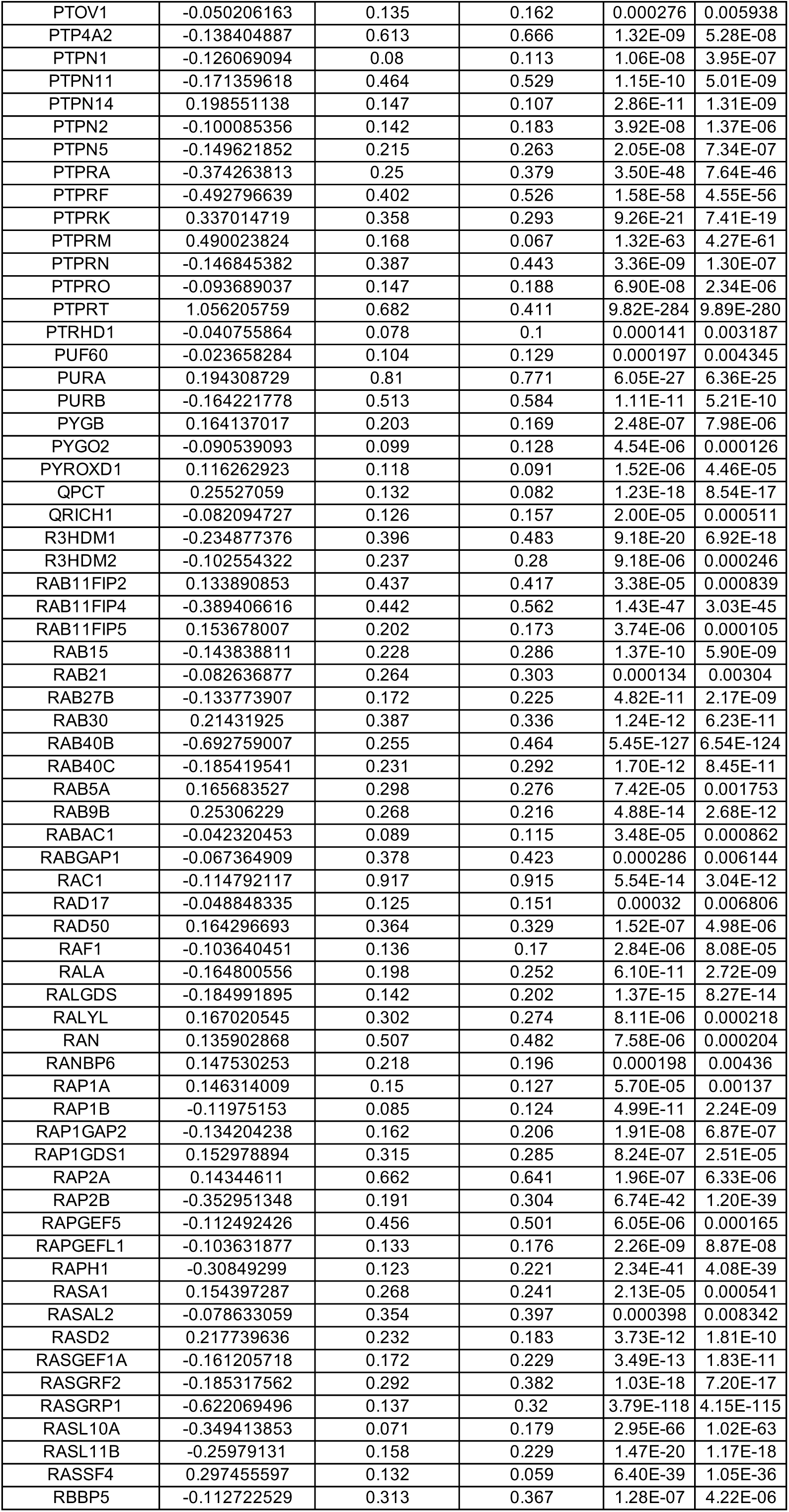
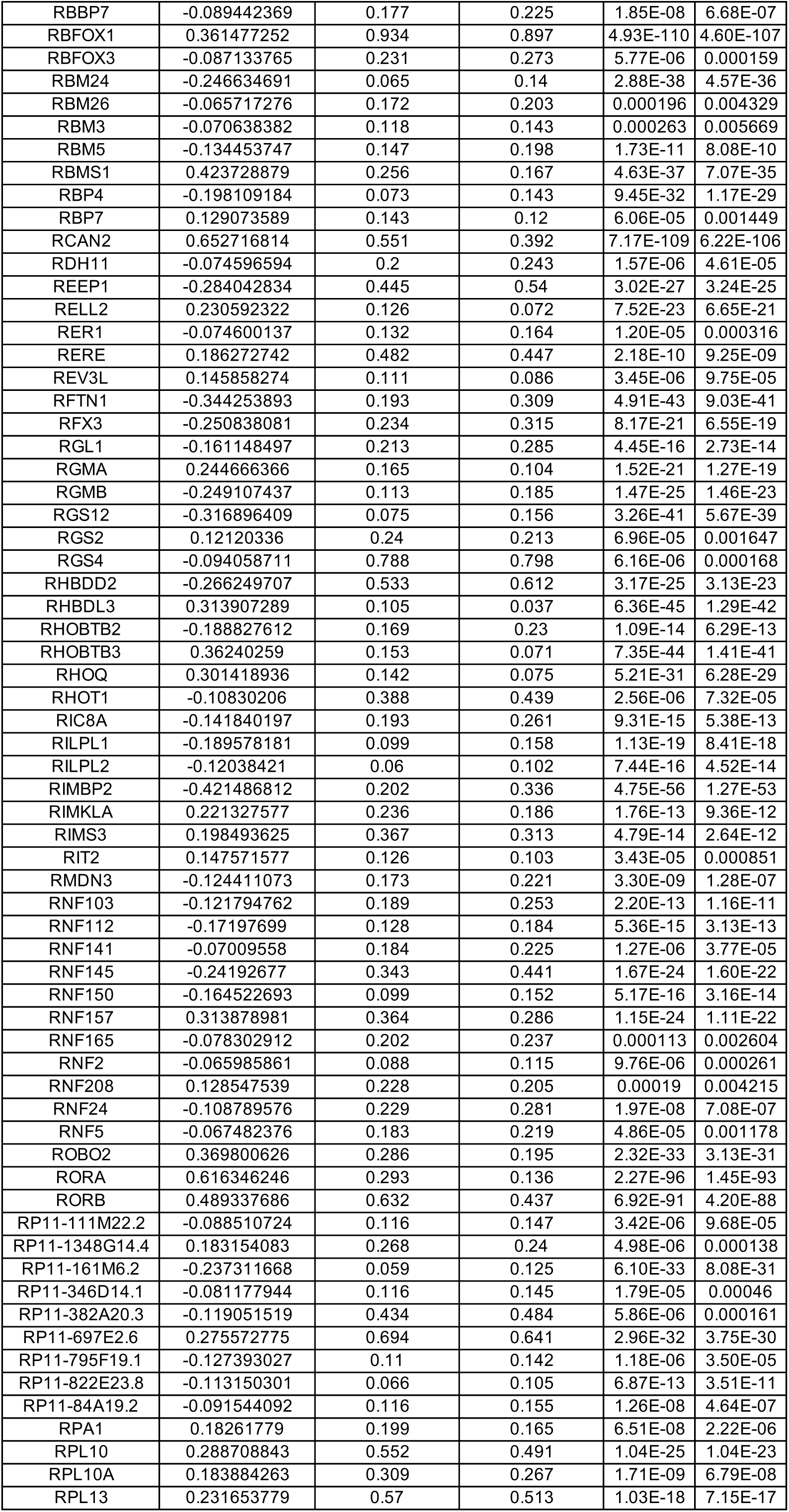
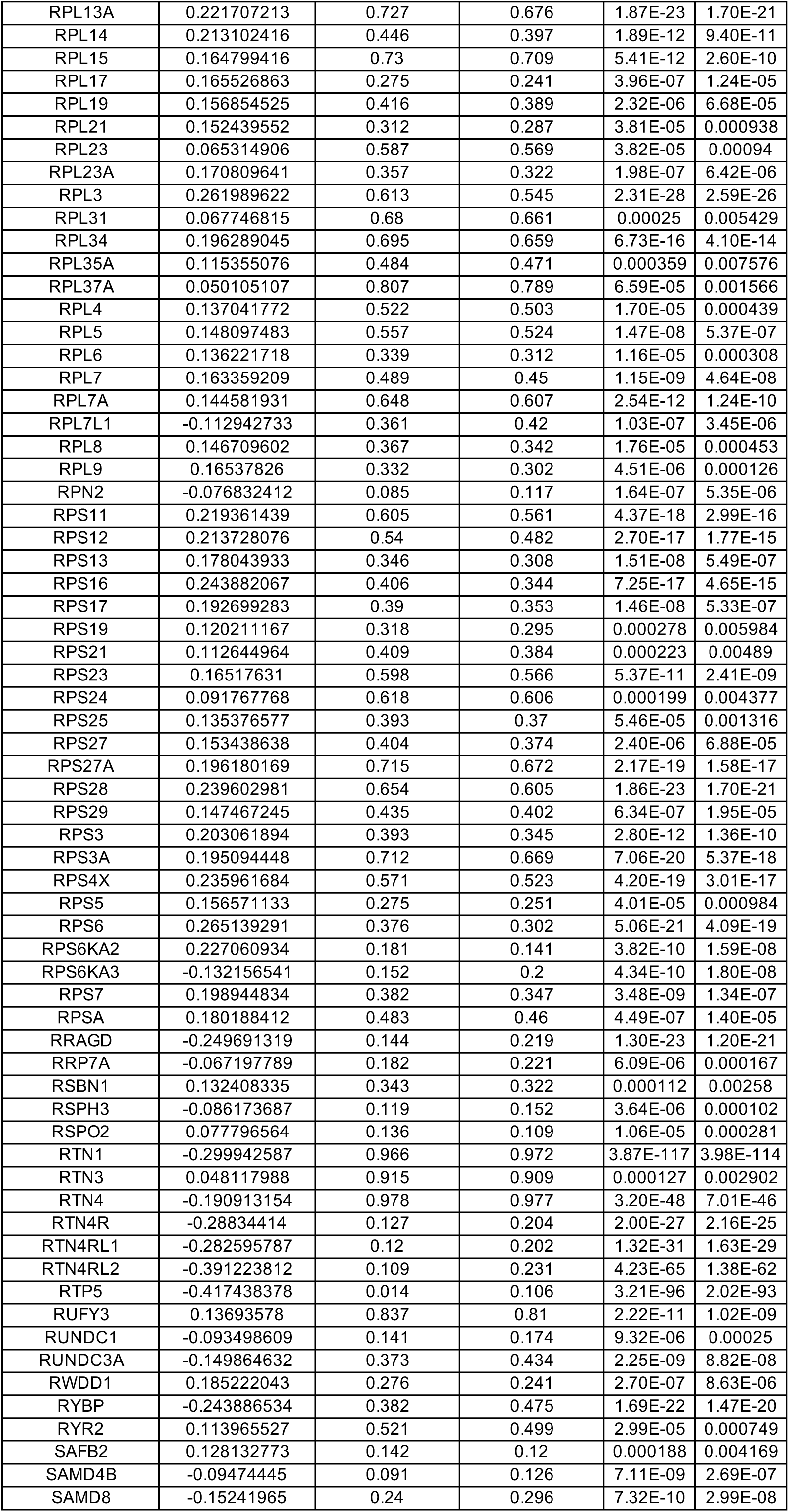
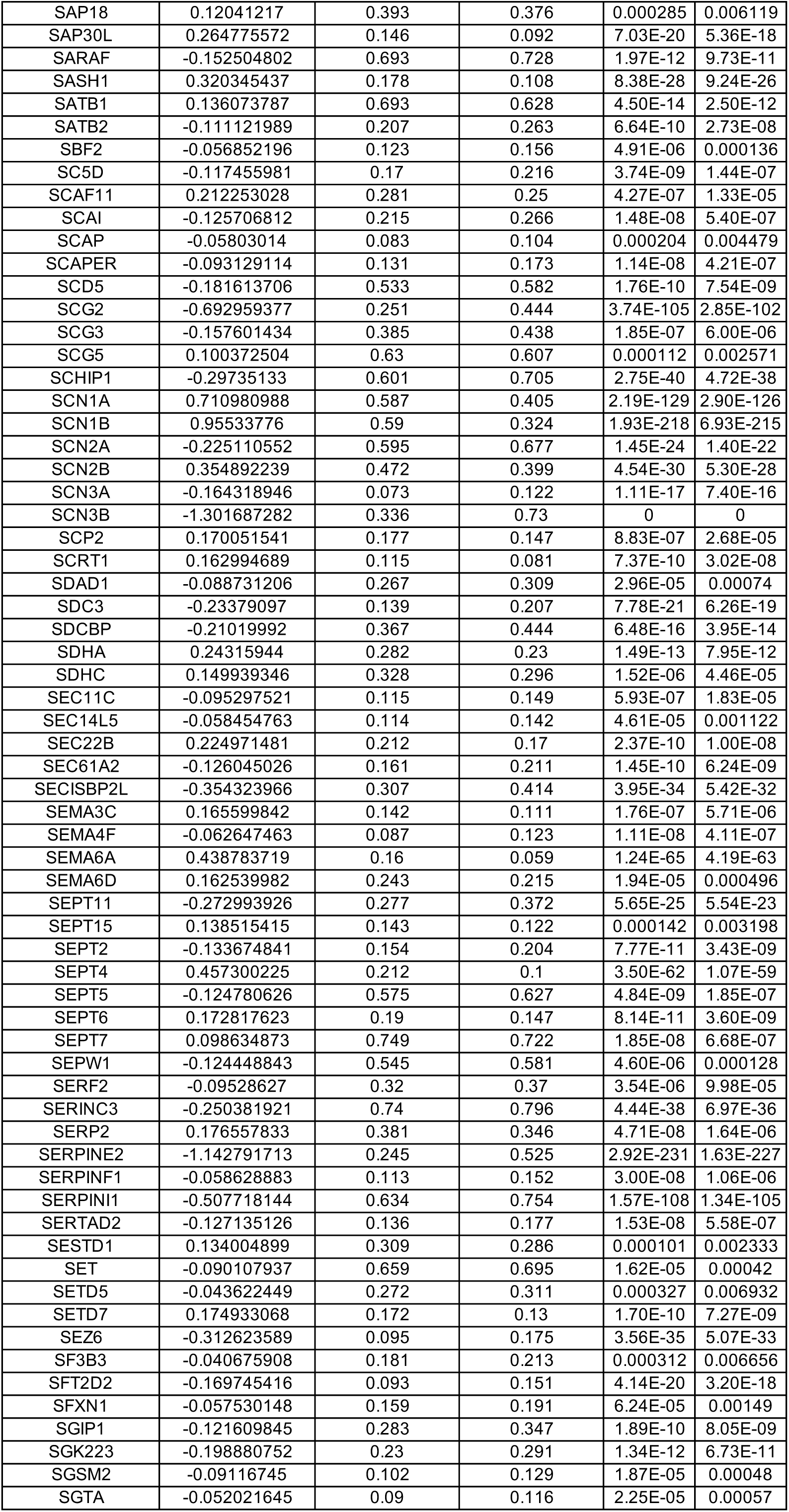
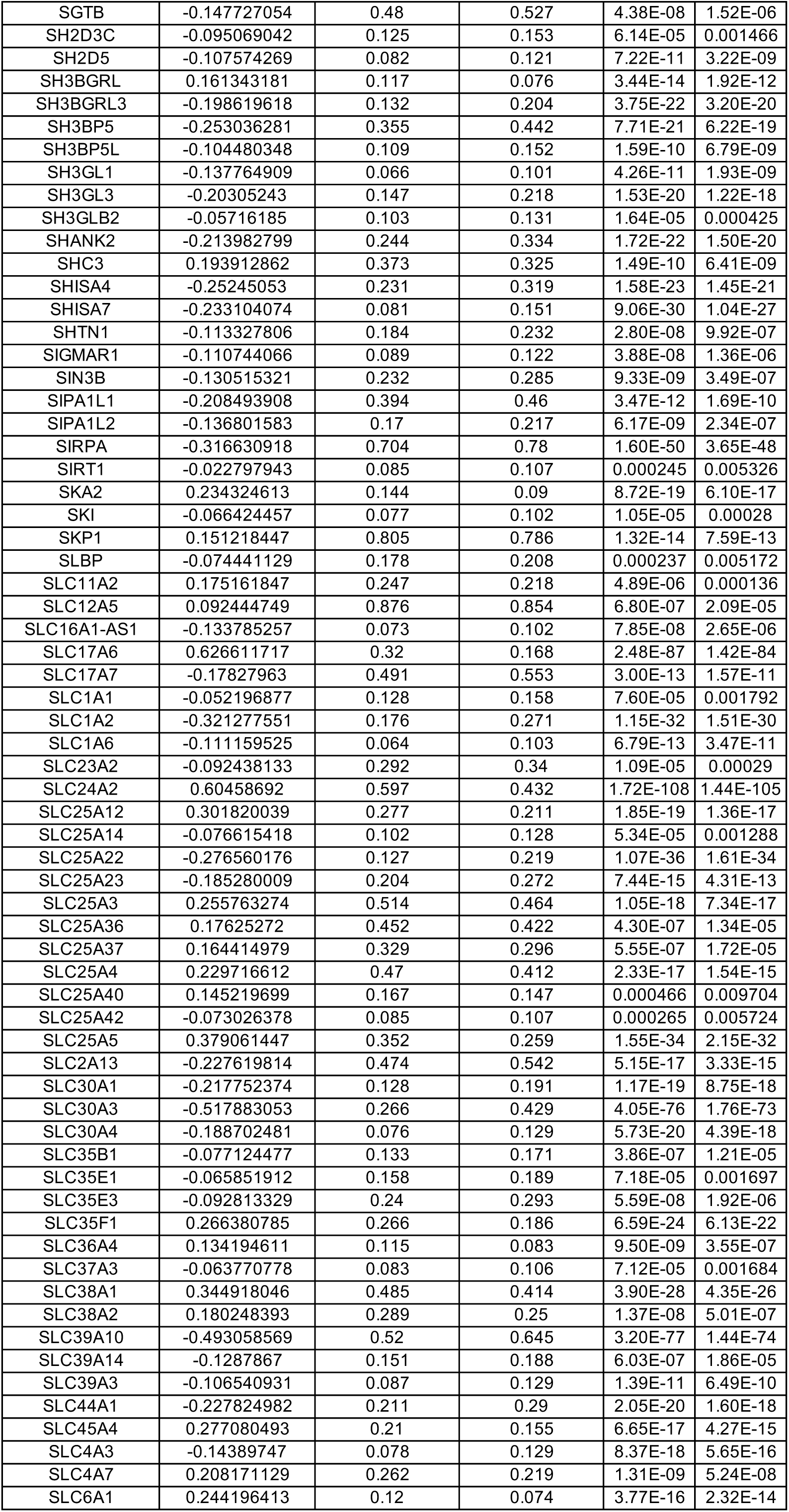
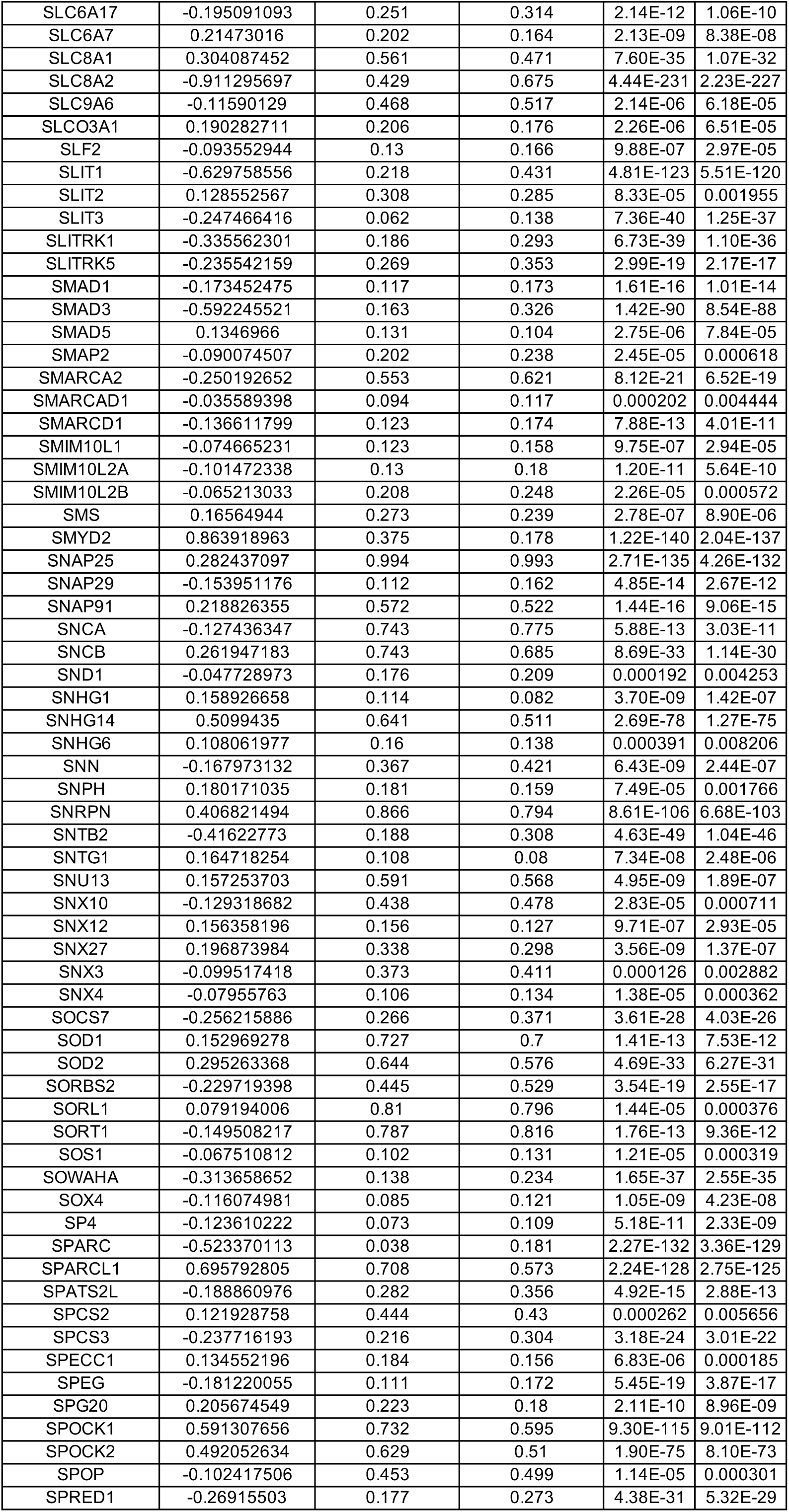
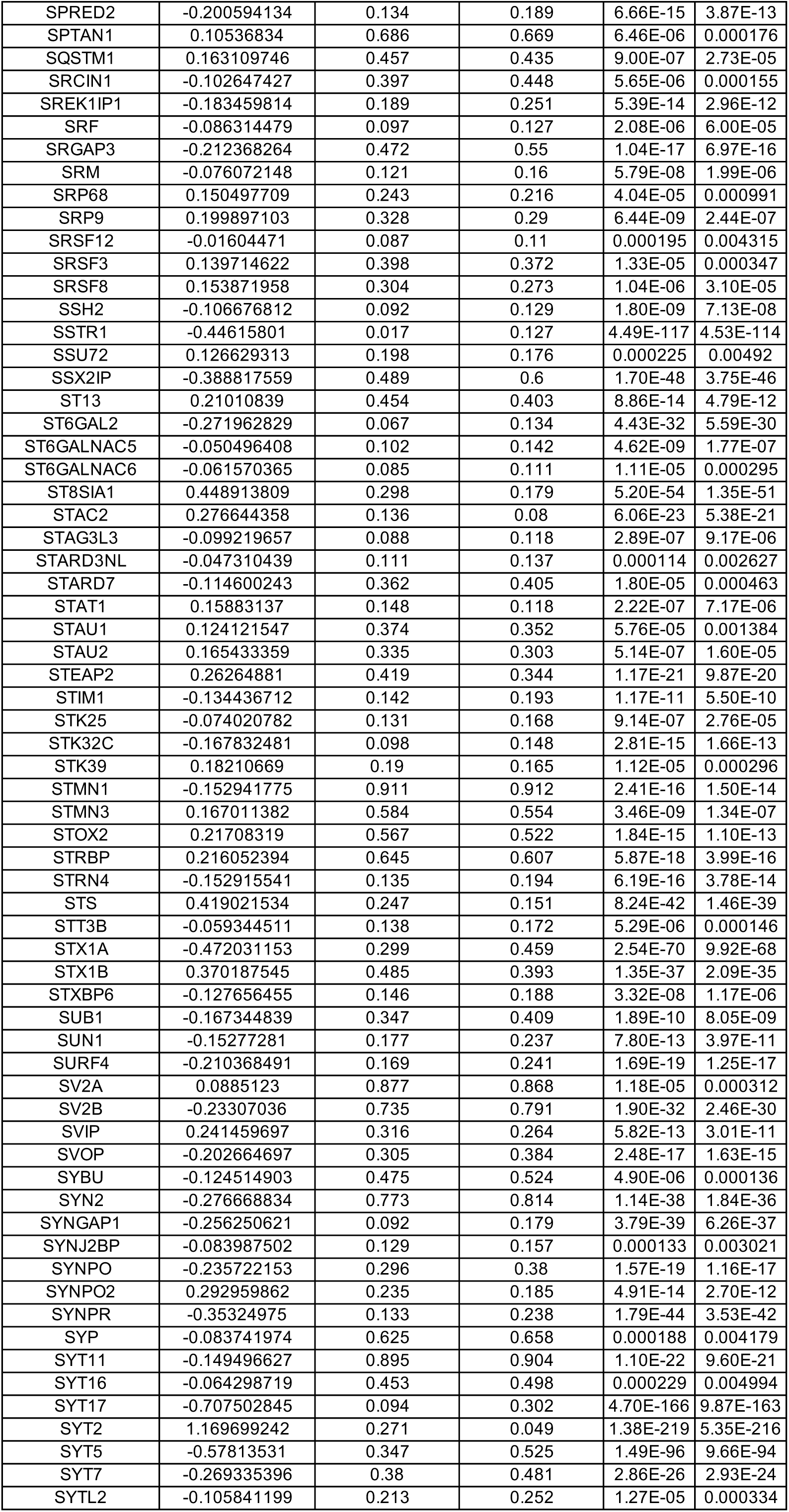
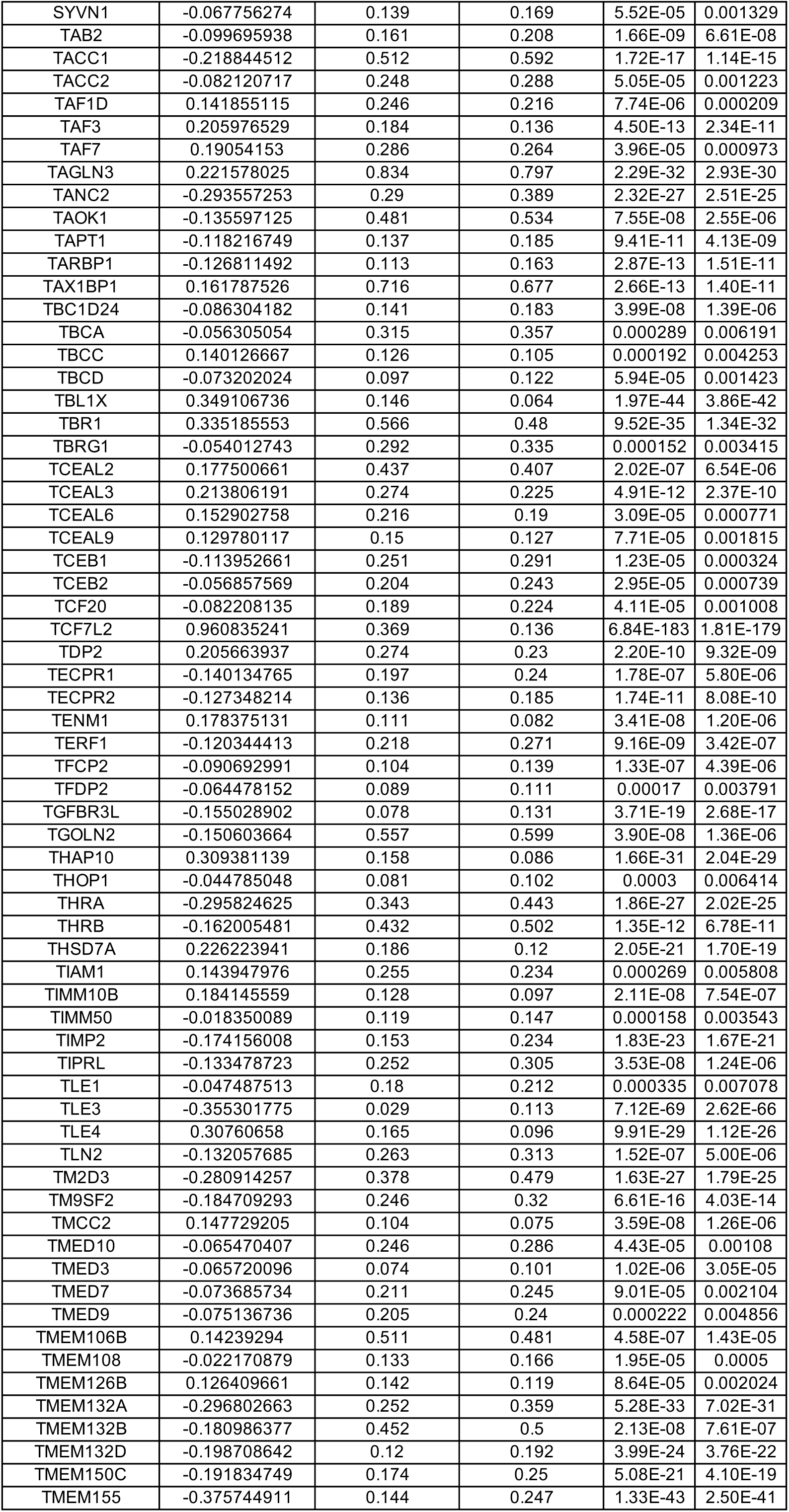
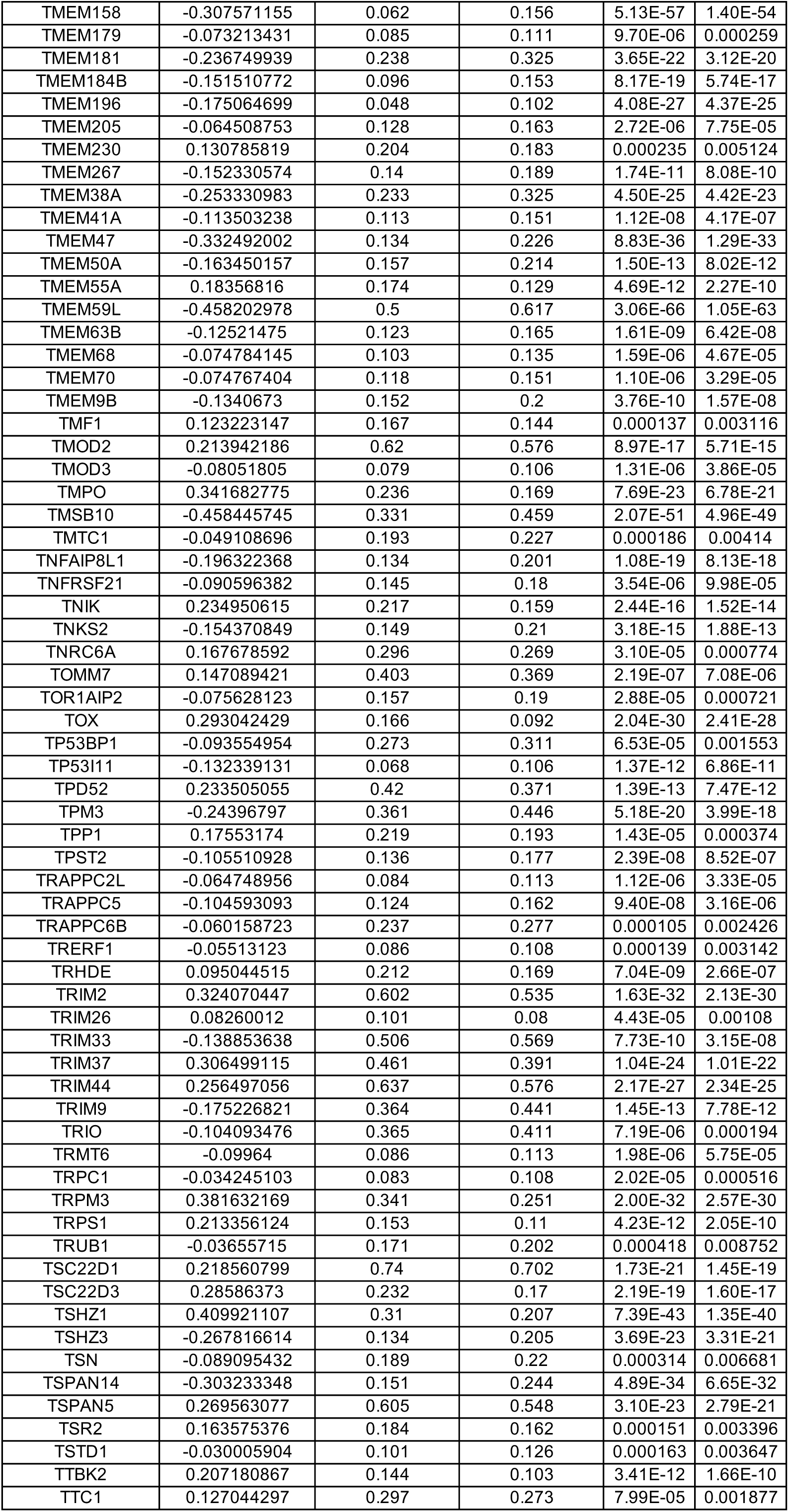
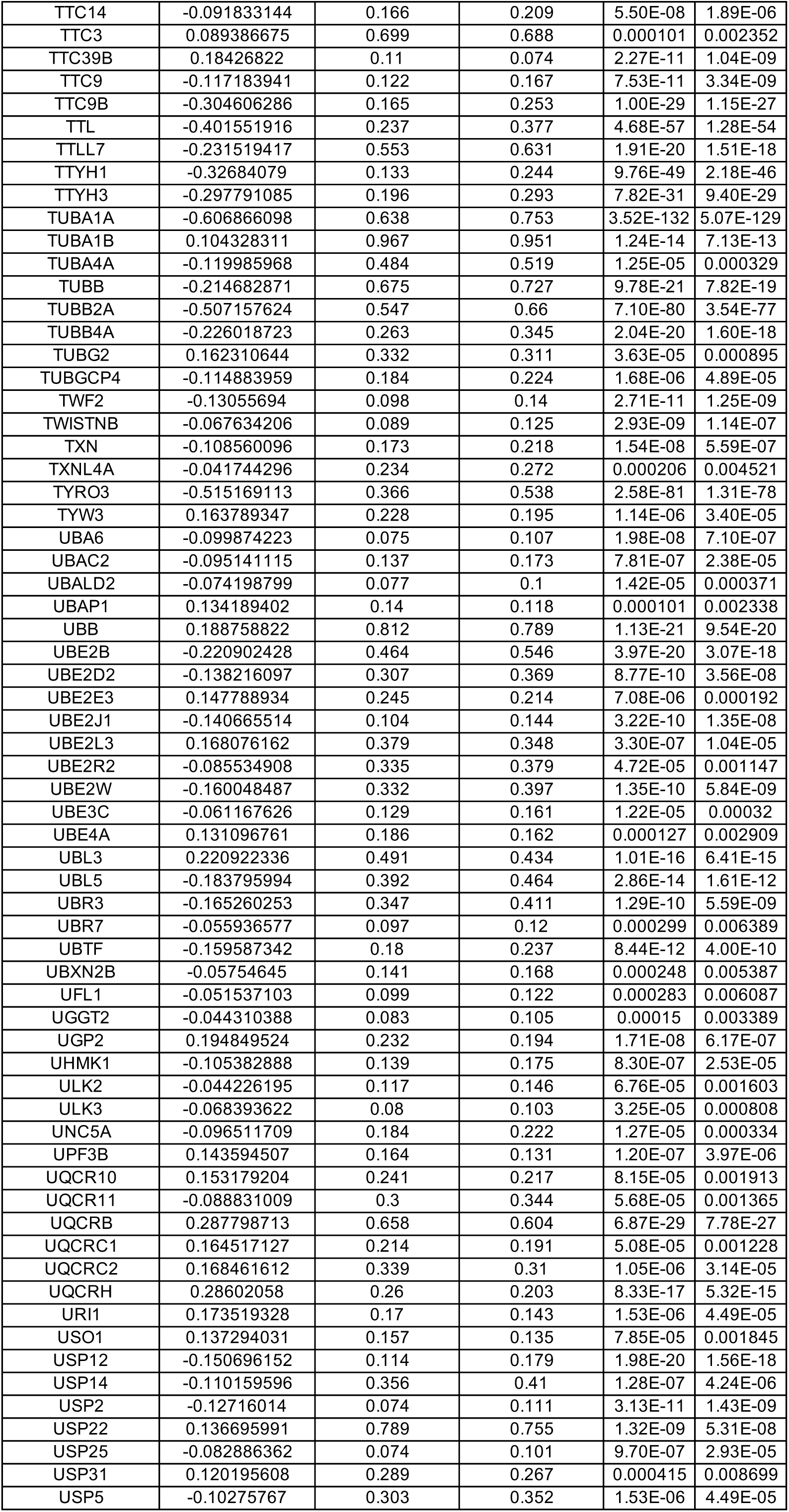
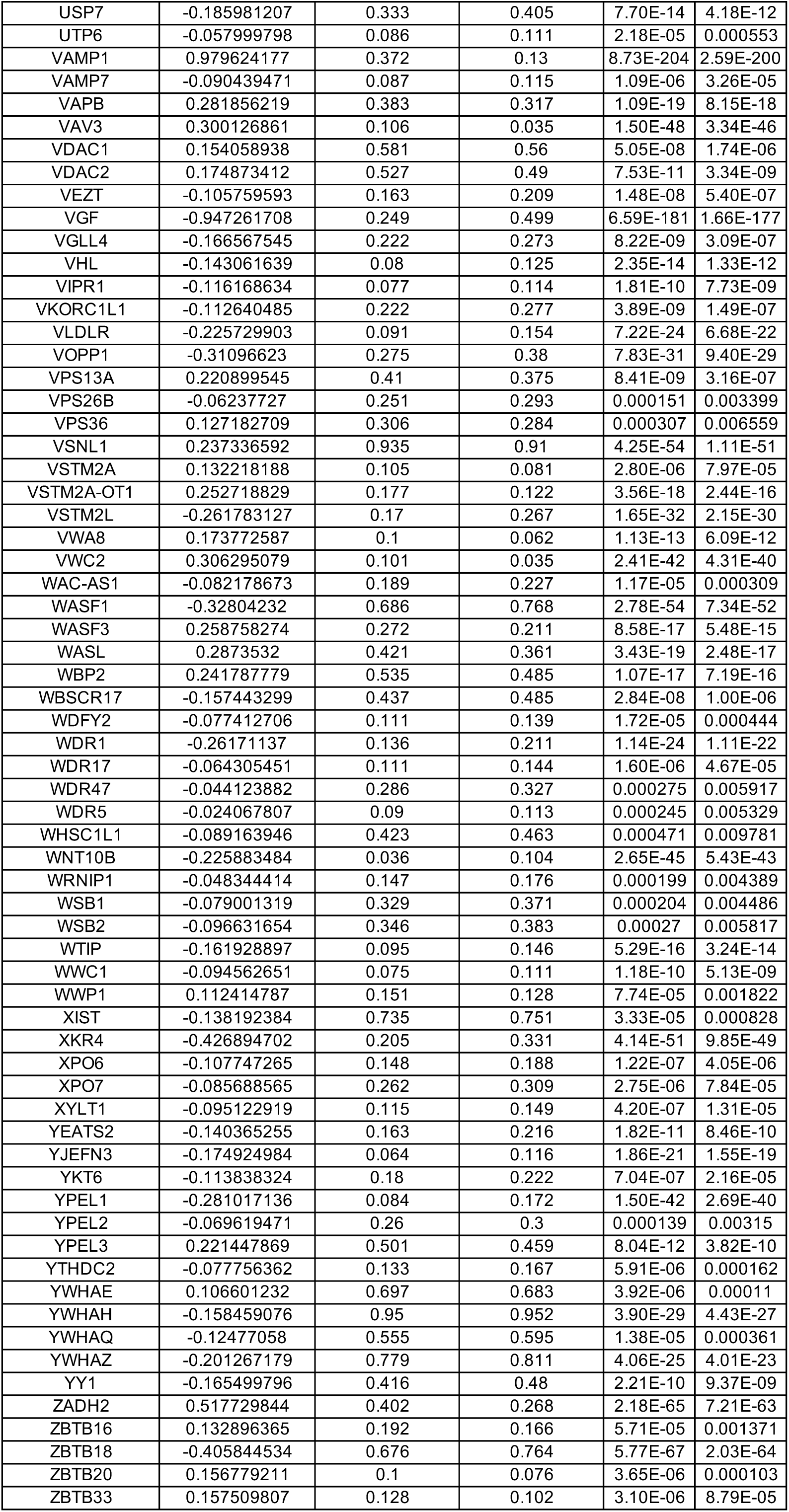
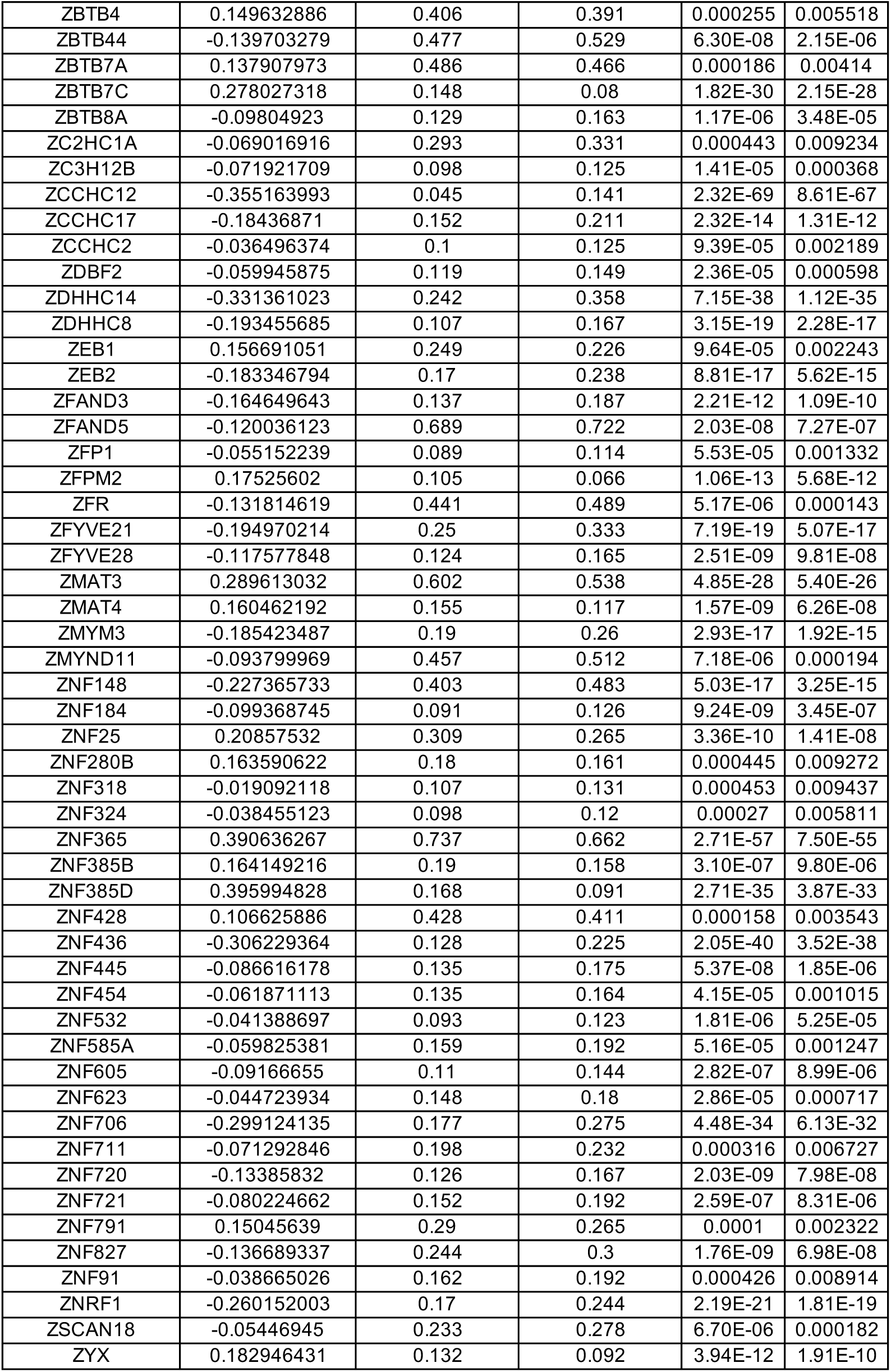
DEGs between wild-type (WT) and MECP2-null (KO) excitatory neurons of human OC (adjusted p-value < 0.01)

**Table S44.**
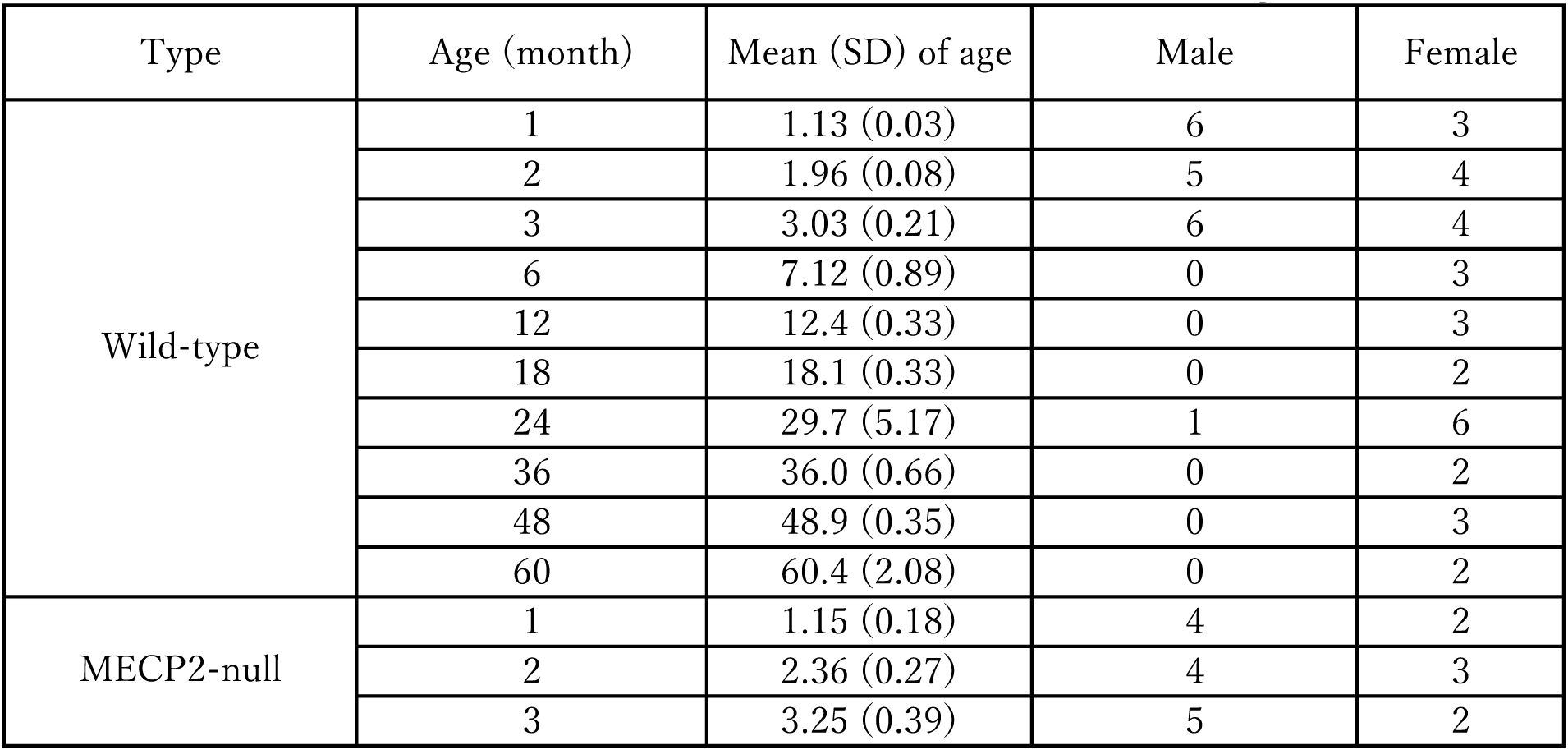
Common marmosets for evaluation of volume change over time.

**Table S45.**
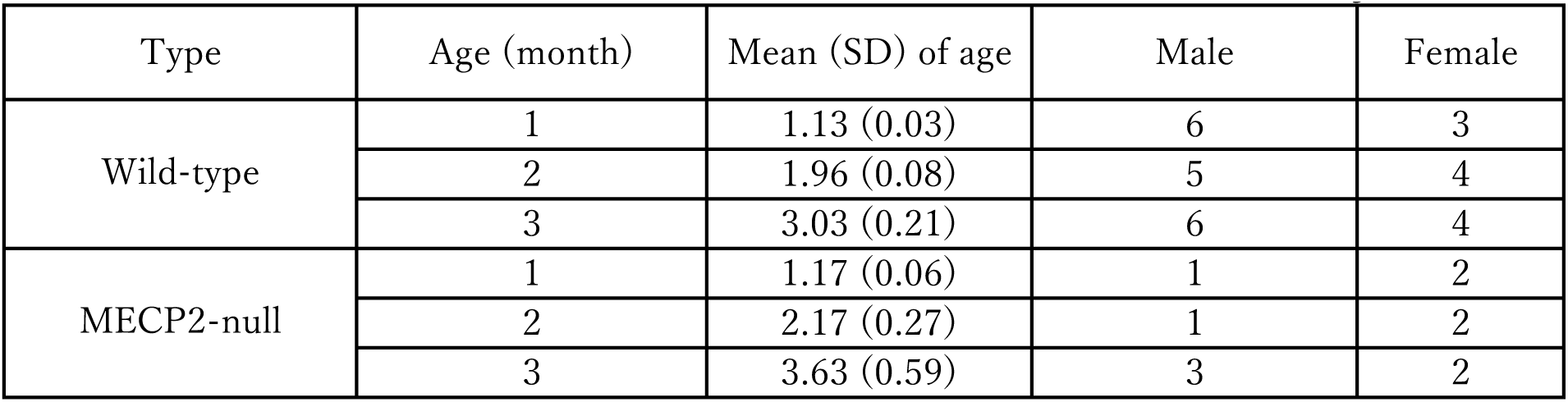
Common marmosets for evaluation of neural connectivity over time.

**Table S46.**
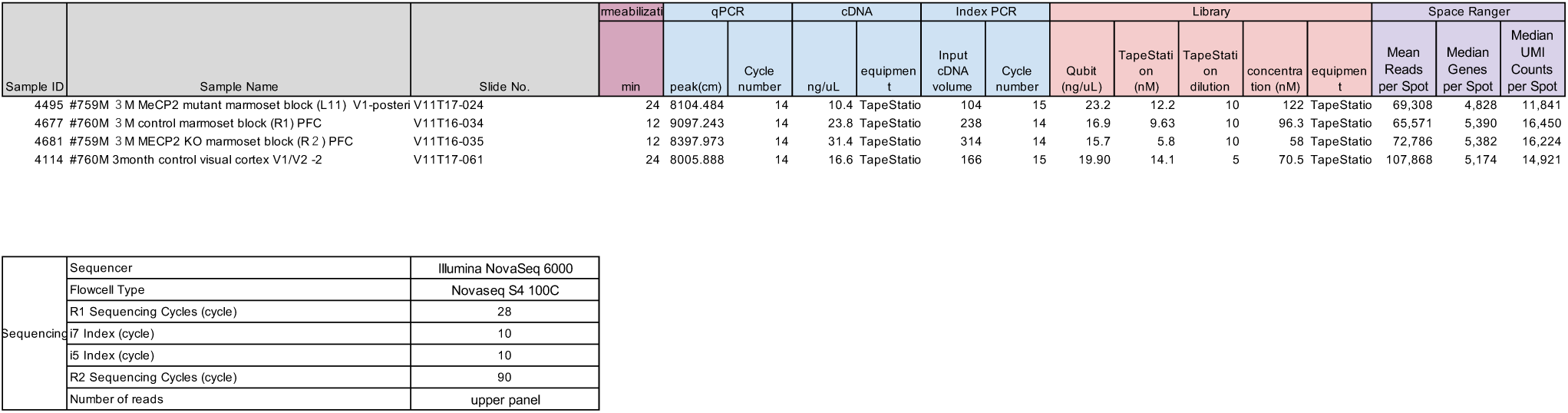
Visium detailed experimental conditions.

